# Intraspecific root and leaf trait variation with tropical forest successional status: consequences for community-weighted patterns

**DOI:** 10.1101/611640

**Authors:** J. Aaron Hogan, Oscar J. Valverde-Barrantes, Qiong Ding, Han Xu, Christopher Baraloto

## Abstract

1. Quantifying the dimensions and magnitude of intraspecific root trait variation is key to understanding the functional trade-offs in the belowground plant strategies of tropical forest trees. Additionally, accurately measuring how belowground functional trait variation relates to soil environment and forest age is crucial to tropical forest modeling efforts.
2. We sampled leaf and root morphologies from 423 juvenile trees of 72 species from 14 Angiosperm families along a 6.6 km transect that corresponded to an environmental gradient in decreasing soil fertility and texture with increasing forest age.
3. We observed within-lineage conservative functional trait-shifts in root and leaf morphological traits along the transect. From secondary to primary forest, average leaf area increased 7 cm^2^ and average root system diameter increased 0.4 mm. Mean specific leaf area decreased by 0.8 m^2^ kg^−1^, specific root length decreased by 3.5 m kg^−1^, and root branching intensity decreased by 0.3 tips cm^−1^. Leaf thickness and root tissue density showed no change.
4. We coupled trait measurements to a network of 164 1/16^th^-ha plots across a Chinese tropical forest reserve, to scale individual trait measurements up to the community-level, accounting for forest age.
5. For most traits, intraspecific trait variation negatively covaried with species compositional turnover between plots in younger versus older forest to compound and create greater community-weighted differences in trait values than would be observed if intraspecific variation in traits with forest age was not accounted for.
6. *Summary* Root morphologies are variable with local scale variation in soil fertility and texture. Accurately understanding broader (i.e. forest)-scale patterns in root functional traits, requires attention to underlying environmental variation in soil resources, which interacts with environmental filtering of plant communities.

## Introduction

Plant functional traits have been the principal approach applied to understating trade-offs in the physiology and biological functioning of plants, which scale with variation in their ecological life-histories (Keddy, 1992, Weiher et al., 1999, Westoby and Wright, 2006, Garnier et al., 2016, Shipley et al., 2016). Functional traits, such as specific-leaf area (Garnier et al., 2001) (SLA), or wood density (Swenson and Enquist, 2007, Chave et al., 2009) inform about resource (e.g. Carbon (C), Nitrogen (N)) allocation within the plant in relation to a fast or slow growth strategy (Reich, 2014, Wright et al., 2010), total photosynthetic potential (Shipley, 1995), and other measures of the relative plant performance across species (Weiher et al., 1999, Ackerly et al., 2002, Díaz et al., 2016). Interspecific tradeoffs in resource allocation in relation to the most-commonly measured function traits have been understood to fall out on two orthogonal axes of variation (Díaz et al., 2016): one encompassing the stem economics spectrum (Chave et al., 2009) related to whole plant size (King, 1996) and another related to the leaf economics spectrum (Wright et al., 2004, Reich et al., 1992).

The widespread measurement of functional traits of wild plant roots is a relatively recent development in plant functional ecology (Bardgett et al., 2014, Laliberté, 2017, Iversen et al. 2017). Until recently, we have known little of how root traits relate to variation in other (e.g., leaf, stem) traits and scale with plant strategies, because of the multidimensionality of the physiological tradeoffs in roots (Mommer and Weemstra, 2012, Reich, 2014, but see Freschet et al., 2010, Freschet et al., 2018) and their varying morphologies (Kramer-Walter et al., 2016). Much of the variation in root functional morphologies, and therefore root strategy among all land plants can be explained by continental-scale climate variation (Freschet et al., 2017, C. Wang et al., 2018, Jackson et al., 1996), yet a large amount remains unaccounted for (Freschet and Roumet, 2017, Valverde-Barrantes et al., 2017). At the individual plant scale, variation in root traits can be large; for example, interspecific variation in root diameter, specific root length (SRL) and link length within a community can be twenty-fold (Comas and Eissenstat, 2009, Guo et al., 2008). Such variation has potentially large ramifications for the ecologies of plant species (Schenk and Jackson, 2002, Craine et al., 2001, Comas and Eissenstat, 2009, Van Kleunen et al., 2010, Bardgett et al., 2014, C. Wang et al., 2018), including nutrient acquisition and use strategies and how species may respond differentially to modern Anthropogenic selections pressures.

The emerging paradigm with respect to root morphological traits it at is that low SRL, thick root diameter, high C:N ratios in tissues, and high root tissue density relate to low nutrient uptake capacities, low rates of root respiration, and long root lifespans, and are thus considered conservative trait should correspond to a K-selected, slow-growth plant strategy (Grime, 1977, Reich, 2014, Weemstra et al., 2016). In contrast, thin root diameters, high SRL, high root tissue N content, and low root tissue densities should relate to fast rates of root turnover and should be indicative fast-growth, pioneer-type strategies (Fig. S1). Recent studies have shown that root tissue density and diameter are positively-related to root lifespan and drought-resistance, but negatively related to nutrient uptake potential (Pérez-Harguindeguy et al., 2013, Kramer-Walter et al., 2016, Valverde-Barrantes and Blackwood, 2016). Most fine root functional trait values (e.g. diameter, SRL) should decrease in values from an acquisitive, R-selected, fast-growth plant strategy to a more conservative, K-selected, slower growth strategies (Weemstra et al., 2016, McCormack et al., 2012). This is because most morphological and architectural root traits are single-dimensional measurements, or mass-corrected single dimensional measurements, which correspond to the ability to explore a given soil volume per unit of plant investment in root tissue production (Lynch, 2005, Fitter et al., 1991).

Across taxa, the general pattern is that gymnosperms and basal angiosperms (e.g., Magnoliids) have thicker, denser, shorter roots, with fewer fine root tips than high-order angiosperms (Kong et al., 2014, Valverde-Barrantes et al., 2016, C. Wang et al., 2018). This may signify a greater reliance on mycorrhizal associations for nutrient acquisition (Eissenstat et al., 2015, Chen et al., 2016, Valverde-Barrantes et al., 2016, Kong et al., 2017), or may be a result of divergent evolutionary processes that have created a high degree of phylogenetic conservatism in Angiosperm root diameter (Ma et al., 2018, Lu and Hedin, 2019). Thus, studies investigating variation in root morphologies must account for plant lineage. Maherali (2017) showed that the standard deviation of root diameter increased with the average root diameter (which is largely phylogenetically-conserved), using family-level data from 581 species from 22 plant orders (Valverde-Barrantes et al 2017). Yet, there was substantial variation about this trend among families, and it is unclear how individual variation in root topology due to environment scales with root diameter, plant lineage, or across plant communities.

Nevertheless, irrespective of the relatively-consistent interspecific differences in root diameter, plants have been shown modulate root architecture in response to cues in the soil environment (i.e. soil moisture and fertility) (Fitter and Stickland, 1991, López-Bucio et al., 2003, Hodge et al., 2009). Nutrient deficiencies in the soil usually lead to the lengthening and architectural development of root systems (Fitter and Stickland, 1991, López-Bucio et al., 2003, Giehl et al., 2013). Forest communities typically move from acquisitive to conservative functional composition as species composition changes with increasing forest succession (Garnier et al. 2004, Swenson et al. 2012, Lohbeck et al. 2013, Letcher et al., 2015, Muscarella et al., 2016). To date, there have been few studies that look at above- and below-ground functional traits of tropical trees in relation to small-scale variation soils or in forest age, and it is unclear how intraspecific belowground trait variation may interact with species turnover among communities. One might hypothesize for roots of the same species to become more acquisitive (i.e. an intraspecific shift toward acquisitive root functional traits), as soil fertility decreases with increasing forest successional status. The logic is, that to meet the physiological requirements for growth, plants must adapt their root morphology to become more-acquisitive as root biomass and competition increases, and soil fertility decreases with forest biomass accretion and succession. This is the case with leaf functional traits, in that forests success, they increase in SLA. The increase in SLA, mainly driven by an increase in leaf area, with increasing forest biomass, canopy coverage and competition for light has been well studied (Givnish, 1984, Rijkers et al., 2000, Keenan and Niinemets, 2017). Yet, root systems are plant organs that must simultaneously forage for 14 mineral elements from the soil in addition to absorbing water (Lynch, 2005). Thus, it is not unreasonable to hypothesize that morphological plasticity of root systems could be multidimensional (Fitter, 1991, Weemstra et al., 2016, Maherali, 2017, Laliberté, 2017), and potentially have distinct abiotic and biotic controls that modulate functional trait expression.

Understanding how root morphology varies in the context of environmental variation and forest age is key to further understanding the ecology and functioning of tropical forests into the future (Bardgett et al., 2014, Warren et al., 2015, McCormack et al., 2017).

In that context, our two research questions and accompanying hypotheses were:

1. How does functional trait variation in roots and leaves relate to local-scale environmental variation in soil fertility and texture, forest successional status of montane tropical forest? *We hypothesized that intraspecific variation in roots should become more acquisitive as soil fertility decreases and competition for soil nutrients increases with forest succession. We hypothesized that intraspecific variation in understory leaves should become more acquisitive as light availability decreases with forest succession. Incorporated in these two independent hypotheses, is the idea that individuals coordinate root and leaf traits to become more acquisitive with increasing forest age*.
2. To what extent is community-weighted mean functional turnover across local-scale environmental gradients due to changes in community composition (i.e., forest age and/or environmental filtering effects) or intraspecific functional trait variation? *We expected changes in plant community composition and species relative abundances with forest age would influence estimates of community-weighted traits to a greater degree than would intraspecific trait variation (ITV)*.

We employed a paired sampling design that measured root and leaf morphological traits and soil chemistry along a 6.6-km transect in a subtropical, island forest ecosystem in southern China. Additionally, we incorporate data from a network of small vegetation plots established across the reserve (Xu et al., 2015a,b) to scale up trait measurements from plants across the transect to the forest community.

## Materials and methods

### Study site: Jianfengling, Hainan Island, China

The Jianfengling forest reserve (JFL), of south-west Hainan island, China (18°23’– 18°15’N & 108°36’–109°05’E, Fig. 1) is a 47,200 ha mountainous forest with a history of logging and forest resource extraction that dates back to 1957, where about two-thirds of the area was either clear-cut or selectively-logged, meaning 30-40% of large timber-valuable trees were extracted (Zhou, 1995, Xu et al., 2015b). All logging ceased in 1994 under a state-wide (Hainan island only) logging ban, followed by a Chinese national logging ban in 1998 (Zhou, 1995, Wenhua, 2004). The reserve encompasses several ecological life zones of vegetation, from tropical semi-deciduous monsoon forest at the lower elevations to mossy high elevation forest, with evergreen-monsoon forest dominated by Podocarpaceae intermixed throughout at elevations < 1000 m (Huang et al., 1995). The most common vegetation life zone is tropical montane rain forest, which occurs at elevations between 600 and 1100 m. It is characterized by a mix of palms (*Livistona saribus* (Lour.) Merr. ex A. Chev. is most common), broadleaf evergreen trees that reach an average canopy of height of 18 m (Jin et al., 2013).

**Figure 1:**
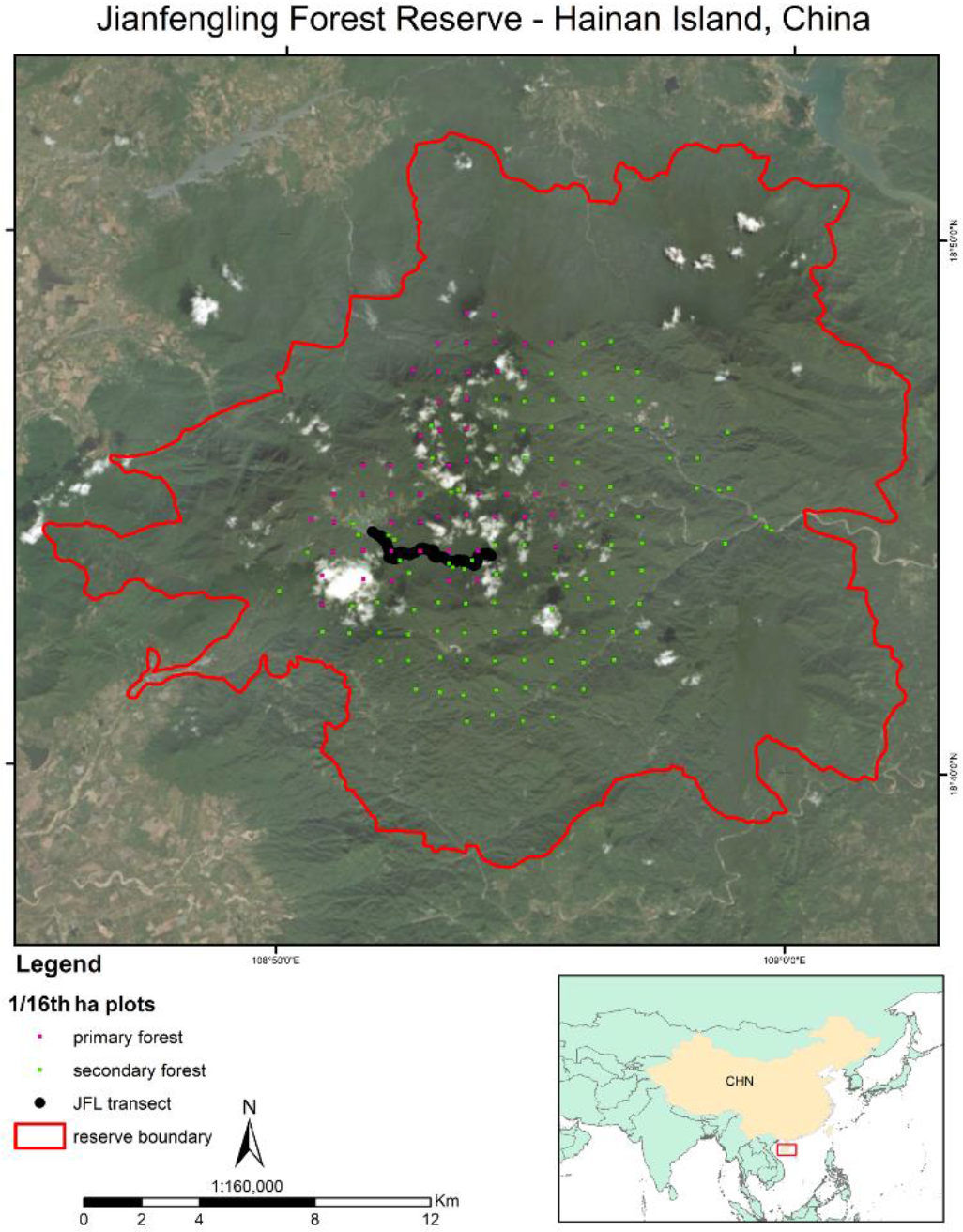
The study site: the Jianfengling Forest Reserve (JFL) of Hainan Island, China. Hainan Island is a small continental island off the southeastern coast of China, shown in the red box of the inset map. The 47,200 ha JFL boundary is shown in red. The 6.6 km transect of where functional traits of saplings were sampled is shown in black (see Appendix for an elevational profile of the transect). The first half of the transect is in secondary forest with a history of logging, and the second half of the transect is in unlogged, primary forest. Small plots (Han et al. 2015a, b) in primary and secondary forest are shown as pink and green points, respectively.

The climate in the area is a tropical monsoon climate with a seasonal rainfall regime where most of the rainfall occurs between May and October (Zeng, 1995). Cumulative 30-yr annual rainfall in the montane rain forest of JFL averages about 2700 mm (Wu, 1995). In the tropical montane rainforest lifezone, the soils are classified as lateritic and humic yellow soils, being derived from porphyritic granite (Wu, 1995). Such soils are characterized by surface accumulation of organic matter, slower rates of mineral and organic matter cycling than other tropical soils (e.g., latisols), intermediate rates of mineral leaching and some accumulation of Aluminum, and exchangeable base content of about 30 mL per kg of soil (Wu, 1995)

### Field Methods

During the summer (May 4 to June 30) of 2017, roots and leaves were of juvenile trees (individuals < 10 cm diameter at breast height, hereafter saplings) were sampled along a 6.6 km transect within the JFL reserve (Fig. 1, Fig. S2). A transect was positioned to capture a human disturbance gradient, roughly moving from secondary to primary forest and terminating in the 60-ha JFL permanent forest dynamics plot. A total of 423 individuals of 73 species from 14 tree families: Magnoliaceae, Annonaceae, Moraceae, Fagaceae, Juglandaceae, Sapindaceae, Rutaceae, Anacardiaceae, Burseraceae, Ebenaceae, Styracaceae, Sapotaceae, and Theaceae (see Fig. 2 for species Latin binomials), were collected in a paired approach, that sought to collect three individuals of each species in each half of the transect.

**Figure 2:**
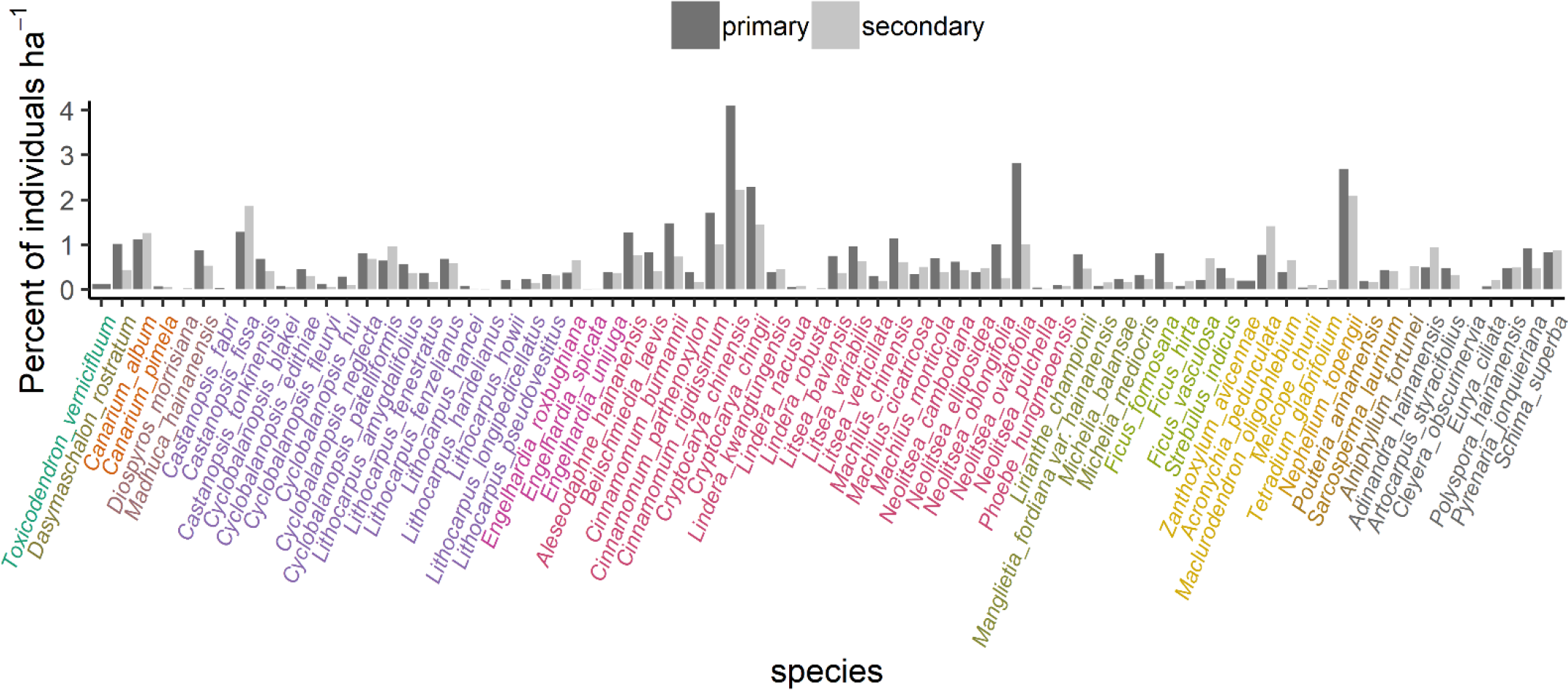
Relative abundances (per hectare) for the 72 species for which trait data were sampled. Relative abundances were calculated using abundance data from 164 1/16^th^-ha small plots established throughout the JFL reserve (see Figure 1 and Han et al. 2015 *Journal of Applied Ecology*). 112 of those plots are in secondary (i.e. previously logged or clear-cut) forest, while 52 are in primary forest. Colors of species names correspond to plant family in order from right to left: Anacardiaceae, Annonaceae. Burseraceae, Ebenaceae, Fagaceae, Juglandaceae, Lauraceae, Magnoliaceae, Moraceae, Rutaceae, Sapindaceae, Sapotaceae, Styracaceae, and Theaceae (see legend in Figure 3). Demographic data from those plots are used throughout for scaling intraspecific trait variably from individuals to the community.

For each individual, surface lateral fine roots in the top 10 cm of soil were gently excavated, tracing them from an identified individual. Excavation of living root systems from saplings was done carefully to preserve root morphology of the finest first order roots, using a hand shovel and spade, when necessary. Healthy leaves were manually cleaved off the plant at the base of petiole and collected. Leaves and roots were transported back to the lab in plastic bags for processing. Measurements of leaf morphologies were done immediately, and roots were placed in a refrigerator for storage until they could be processed. Following root collection, *ca*. 1 kg of surface soil from the immediate vicinity of the root sample was taken was collected.

### Functional trait measurements

For three leaves of each individual juvenile tree, leaf thickness was measured using a Vernier micrometer (Mitutoyo USA) precise to a thousandth of a millimeter. Leaves were scanned (single-sided scans) for leaf morphological measurements. Prior to scanning, roots were washed thoroughly by hand to remove soil. For each individual, three to five entire root systems, containing 3-4 root orders (ERSs *sensu* McCormack et al. 2015) were selected. Roots were placed in an acrylic root scanning tray with a cover glass, submerged in water and scanned at high resolution in black and white using a double-sided optical scanner (Epson Perfection V800, Epson America, Inc.). Scanned images of leaves and roots were analyzed using WinFolia (2007b version, Regent Instruments, Quebec, Canada) and WinRhizo (2016 version, Regent Instruments, Quebec, Canada) software. Following scanning, leaves and roots were dried in an oven at 70°C for at least 48 hours, before recording their dry mass. WinFolia measures leaf area, length width, perimeter, and aspect ratio. Specific leaf area (SLA) was calculated as the ratio of the surface area to the dry mass. WinRhizo measures root length, area, average diameter, volume and architecture (i.e., the number of root tips and forks) for each ERSs. Specific root length (SRL), specific root area (SRA), and specific root tip abundance were calculated by dividing root length, root area, and the number of root tips for each ERS, respectively, by its dry mass. Root branching intensity (RBI) and specific root tip abundance (SRTA) were calculated by dividing the number of root tips by the ERS root length and mass, respectively. Finally, root tissue density (RTD) was estimated by dividing ERS volume by its dry mass.

### Soil analytical methods

For 300 of the individuals of 50 species (6 individuals of each species paired across the gradient with 3 in each forest type), the collected soil samples were sieved (using a 1 mm soil sieve) and sent to the Guangzhou Xinhua Agricultural Technical Development Limited Company for analysis. Soil texture was measured for sand, silt and clay percentages using the international mechanical soil classification standard. Soil pH was measured using a glass electrode in 2.5:1 water to soil dilution. Total soil organic matter was measured using the high-temperature external heat potassium dichromate oxidation volumetric method. Total soil Nitrogen (N) was measured using Kelvin-distillation titration. Total Phosphorus (P), Available Potassium (K), and exchangeable sodium, calcium and magnesium were measured using an ammonium-acetate extraction, followed by flame atomic absorption spectrophotometry. Total K was measured by sodium-hydroxide melting- flame atomic absorption spectrophotometry. Alkaline-hydrolysable (i.e., available) N was measured via the alkali solution diffusion method. Available P was measured by doing a hydrochloric acid–ammonium fluoride extraction and the molybdenum antimony anti-coloring method. Lastly, soil cation exchange capacity (CEC) was measured using the ammonium acetate method.

### The network of small plots

We leverage data from a network of small plots previously established in the JFL forest reserve (Xu et al., 2015a,b) Between August 2007 and June 2009, a network of 164 1/16^th^-ha plots were established in the JFL reserve (Xu et al., 2015a) (Fig. 1). The plots span the local variability in vegetative life-zones and range in altitude from 259 to 1135 m. Roughly one third (52 of 164) of the small plots are in old growth forest, with no record or visible evidence of logging in the last 200 years (Xu et al., 2015b). The remainder of the plots (112) are in forest that was either selectively-logged or clear cut, with human land-use ceasing between 15 and 51 years ago (see Table 1 in Xu et al., 2015b). Selective-logging practices in the area typically result in the removal of 30-40% of the mature stems ≥ 40 cm in diameter of commercially-valuable timber species (Zhou, 1995, Xu et al., 2015b). In each of the 164 small plots, all free-standing stems ≥ 2.5 cm in diameter were mapped, measured, and identified to species.

**Table 1:** Mean and standard error for 14 soil variables from rhizosphere soil in the soil A horizon (0-10 cm depth) for 300 soil samples collected during root excavation in the Jianfengling Forest Reserve, Hainan Island, China during summer 2017. Abbreviations are TEB: total exchangeable bases, ECEC: effective cation exchange capacity, and BS: base saturation. (see Supplement 1)

| Transect area | Soil quality |  | Total soil nutrients |  |  | Available soil nutrients |  |  | Soil cations |  |  | Soil texture |  |  |
| --- | --- | --- | --- | --- | --- | --- | --- | --- | --- | --- | --- | --- | --- | --- |
|  | pH (in water) | Organic matter (g kg<sup>-1</sup>) | N (g kg<sup>-1</sup>) | P (g kg<sup>-1</sup>) | K (g kg<sup>-1</sup>) | N (mg kg<sup>-1</sup>) | P (mg kg<sup>-1</sup>) | K (cmol kg<sup>-1</sup>) | TEB (cmol kg<sup>-1</sup>) | ECEC (cmol kg<sup>-1</sup>) | BS (%) | % sand (0.5-2 mm) | % silt (0.002-0.5mm) | % clay (<0.002 mm) |
| Primary forest (n=150) | 4.27<br>±0.02 | 36.80<br>±1.45 | 1.39<br>±0.04 | 0.11<br>±<0.01 | 15.40<br>±0.70 | 186.56<br>±5.71 | 1.32<br>±0.08 | 0.19<br>±0.01 | 0.64<br>±0.03 | 8.60<br>±0.27 | 8.31<br>±0.41 | 73.01<br>±0.50 | 20.65<br>±0.38 | 6.34<br>±0.19 |
| Secondary forest (n=150) | 4.39<br>±0.02 | 36.56<br>±1.28 | 1.42<br>±0.04 | 0.13<br>±<0.01 | 19.89<br>±0.98 | 176.38<br>±4.12 | 1.58<br>±0.19 | 0.26<br>±0.01 | 1.28<br>±0.10 | 9.78<br>±0.21 | 13.94<br>±1.12 | 67.06<br>±0.55 | 24.63<br>±0.38 | 8.31<br>±0.25 |

### Data analyses: traits along the gradient & intraspecific trait variation (ITV) with forest age

Principal components analyses (PCA) were carried out on root and leaf functional trait matrices to help reduce dataset dimensionality and identify relationships between leaf and root functional traits (see Figs. S3 & S4). This was done separately for root and leaves because they have separate environmental controls (i.e. light vs. soil nutrients), and in our case, we were interested in them separately. The traits used in the PCA for leaves were leaf area, leaf perimeter, leaf width, leaf height, leaf mass, SLA and leaf thickness (see Table S1). The traits used in the root PCA were SRL, SRA, SRTA, RTD, BI, average diameter and root length per volume (see Table S2). PCAs were successful in reducing the dimensionality of morphological trait data, with the first two axes of each PCA explaining 62.1% of the variability in the leaf trait data (Fig. S3), and 69.8% of the variability in the root trait data (Fig. S4). Leaf morphologies were summarized by two main axes of variation: mass-based variation (SLA, dimension 2 in Fig. S3) and area-based variation (leaf area, dimension 1 in Fig. S3). Root morphological traits were not as orthogonally-organized as leaf traits (Fig. S4). Therefore, based on trait correlation with PCA-axes (Tables S1 and S2), three leaf traits: leaf area, SLA and leaf thickness, and four root traits: ERS average diameter (hereafter root diameter), RTD, SRL, and RBI were chosen for subsequent analyses. Subsequent analyses were not carried out using PCA axis loadings, because they can be hard to interpret.

Two-way, univariate analyses of variance were done separately with each of the seven selected functional traits as the response variable. Functional trait values were log_10_ transformed as necessary, as was the case for leaf area, SRL, root area, and RBI. Linear regression step-wise model selection was performed using all possible combinations of species lineage and forest type as factors. Lineage was represented as species nested within a plant family as a random factor variable and forest type was a two-level factor of primary or secondary. In all cases, the best-fitting models included the interaction between lineage and forest type. Those models were then used to predict the least-squared (i.e. marginal) mean trait values with respect to lineage and forest type. Analyses were carried out in R v.3.5.0 (R Core Team, 2018) and made use of the ‘emmeans’ package (Lenth, 2018) for predicting the least square mean values. Effect sizes for predictors were calculated using the ω^2^ estimator via the ‘sjstats’ package (Lüdecke, 2019).

### Data analyses: the network of small plots & Lepš’ Intraspecific variability effects

To address research question two and scale up our trait measurements from individual trees along the gradient to the scope of the JFL forest reserve using demographic data from the network of small plots, we used an assemblage (i.e. partial community) – weighted mean approach, using basal area species weights. We call this an assemblage-weighted mean in lieu of the classical community-weighted mean because we were unable to measure traits for all species found in the plant communities. We did, however, employ a sampling design that controlled for plant lineage, targeting species from 14 families across the Angiosperm phylogeny. Species for which we sampled functional traits accounting for 72 of 580 species and about 37 % of stems (20,673 of 65,144) in the array of small plots. Basal area (measured in August 2007 to June 2009) in the array of small plots ranged from 16.7 to 77.8, averaging 44.2 m^2^ ha^−1^. The basal area of species in those plots for which functional traits were sampled along the newly-established gradient was between <1 and 42.3 and averaged 18.4 m^2^ ha^−1^. Thus, the proportion of plot basal area for which there was trait coverage ranged from <1 to 82.3%, averaging 43.6%.

For each of the seven traits of interest, using mean trait values from all individuals for each species sampled along the transect(*x*_*i*_), we calculated assemblage-weighted means for each plot. We term these the fixed averages (after Lepš et al., 2011), thus 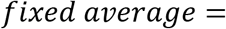 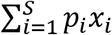, where *p*_*i*_ is the basal area fraction of that species within the plot (i.e. the community weight). Species weights were calculated after removing all species for which trait data was not sampled; this was necessary as weights must sum to one. We, then calculated assemblage-weighted mean trait values for each plot using habitat specific trait values for each species (*x*_*i_habitat*_). For example, for plots in secondary forests, trait values from saplings sampled in secondary forest. We term these the specific averages (again after Lepš et al., 2011), as such 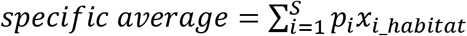. The intraspecific variability effects for the assemblages was calculated as the difference between the fixed and specific averages. Two-way analysis of variances (ANOVAs) assuming homoscedasticity were done with respect to each of the seven functional traits chosen. Then using the sum of squares of these tests, the variance in trait values was decomposed into intraspecific, and community (i.e. species turnover) portions and their covariation (i.e. interaction). This was done using the ‘trait flex ANOVA’ function (see supplemental material of Lepš et al., 2011) in R v.3.5.0 (R Core Team, 2018).

### Data analyses: gap-filling traits & validation of assemblage-weighted means

To validate the results regarding intraspecific variability effects using assemblage-weighted means, we employed an approach to gap-fill trait data for unsampled taxa. Many methods to do this are emerging (e.g. Schrodt et al., 2015, reviewed by Swenson, 2014). Due to the high degree of phylogenetic-conservatism in certain root traits (e.g. root diameter), we chose a use phylogenetic generalized linear model (pGLM) that uses phylogenetic covariance of taxa and a Brownian motion model of trait evolution to gap-fill missing trait values (Bruggeman et al., 2009). Parameter estimates from the phylogenetic and functional trait variance-covariance matrix are used to estimates missing data values. A rudimentary phylogeny for all 582 taxa in the plot dataset was obtained from Phylomatic v3 (http://www.phylodiversity.net/phylomatic, Webb and Donoghue, 2005), using the ‘slik2015’ base tree, which contains the most comprehensive phylogenetic skeleton for tropical trees to date (Slik et al., 2018). Wood specific gravity (WSG) was used as an additional trait in the pGLM, in that we wanted to have one trait for all taxa, and WSG has been shown to be a highly phylogenetically-conserved trait that can be readily used in pGLM to help constrain modeled trait values (Swenson and Enquist, 2007, Kraft et al., 2010). WSG values were obtained from the global database on wood density (Chave et al., 2009, Zanne et al., 2009), where available, and the CTFS Forest-GEO wood density database (http://ctfs.si.edu/Public/Datasets/CTFSWoodDensity/) via the ‘BIOMASS’ package (getWoodDensity() function, Réjou-Méchain et al., 2017). Where species WSG values were unavailable, higher taxonomic classification (e.g. genus or family) values were used (see Supplement 4).

WSG data was combined with three separate incomplete trait matrices, one using fixed trait values (*x*_*i*_), i.e., species mean values for all individuals measured along the transect, and two habitat-specific matrices (*x*_*i_habitat*_), with the habitats being secondary and primary forest, as delineated along the sampling transect. Then using the generated phylogeny, pGLMs were fit separately to each incomplete trait matrix and predicted to complete them for all 582 taxa using PhyloPars (http://zeus.few.vu.nl/programs/phylopars, Bruggeman et al., 2009). Regarding the model setting in PhyloPars, we allowed for correlated evolution of the different traits but did not allow for intraspecific variation in the models. We then used the gap-filled trait matrices along with basal area species weights (*p*_*i*_) from the whole community and recalculated fixed and specific averages. Complete statistical outputs from PhyloPars can be viewed in Supplement 4.

## Results

### Linking the transect to the network of small plots

Of the 72 species that were sampled for functional traits, plot community composition showed that 65% (47 species) preferred primary forest, while the remainder (25 species) favored plots in secondary forest (Fig. 2). Basal area in primary and secondary forest is roughly equal, averaging 40.5 (±0.2, standard error) m^2^ ha^−1^ for small plots in primary forest and 42.7 (±0.2) m^2^ ha^−1^ for those in secondary forest. Soils in primary forest area of the transect were significantly more acidic (ANOVA, F_(1,298)_ = 14.1, *p* < 0.001) and of coarser texture than soils from the secondary portion of the transect (% sand: F_(1,298)_ = 63.9, *p* ≪ 0.001, % silt: F_(1,298)_ = 53.8, *p* ≪ 0.001, % clay: F_(1,298)_ = 38.5, *p* ≪ 0.001) (Table 1). Soils in the secondary forest portion of the transect were significantly more fertile than those of the primary forest area, in that they measured higher in total exchangeable bases (TEB: F_(1,298)_ = 35.7, *p* ≪ 0.001), effective cation exchange capacity (ECEC: F_(1,298)_ = 11.62, *p* ≪ 0.001), and base saturation (BS: F_(1,298)_ = 22.5, *p* ≪ 0.001). The range of variability in soil conditions between the transect and the network of small vegetation plots was similar (Table 1, see supplement 1).

### Functional traits along the gradient, inter- & intraspecific variation

Analysis of 1315 leaves and 1949 entire root systems from 423 individual saplings from 72 species of 14 Angiosperm families showed significant interspecific trait variation. For example, predicted marginal-mean SLA values varied from 8.06 (± 0.55, standard error) to 20.47 (± 0.94) m^2^ kg^−1^ for Sapindaceae and Sytracaceae, respectively (Fig. 3). In the secondary forest portion of the transect, predicted marginal-mean SLA values for those two families increased to 9.25 (± 0.56) and 21.66 (± 0.94) m^2^ kg^−1^. In the primary forest, SLA values were 10.37 (± 0.37) and 10.55 (± 0.19) m^2^ kg^−1^ for Fagaceae and Lauraceae, respectively, the two most sampled families of plants along the transect. In secondary forest, those values increased 11.56 (± 0.26) m^2^ kg^−1^ for Fagaceae and 11.73 (± 0.18) m^2^ kg^−1^ for Lauraceae. Across all families, SLA was on average 1.18 m^2^ kg^−1^ greater in secondary than in primary forest (Fig. 3b). Results were similar for but inverse in directionality for leaf area (Fig. 3a), with predicted marginal-mean leaf area being 68.03 (±1.03) cm^2^ in secondary and 74.53 (±1.03) cm^2^ in primary forest, a difference of about 6.5 cm^2^ that was largely consistent across plant families (Fig. 3b). No differences in leaf thickness were measured with respect to forest age (Fig. 3c)

**Figure 3:**
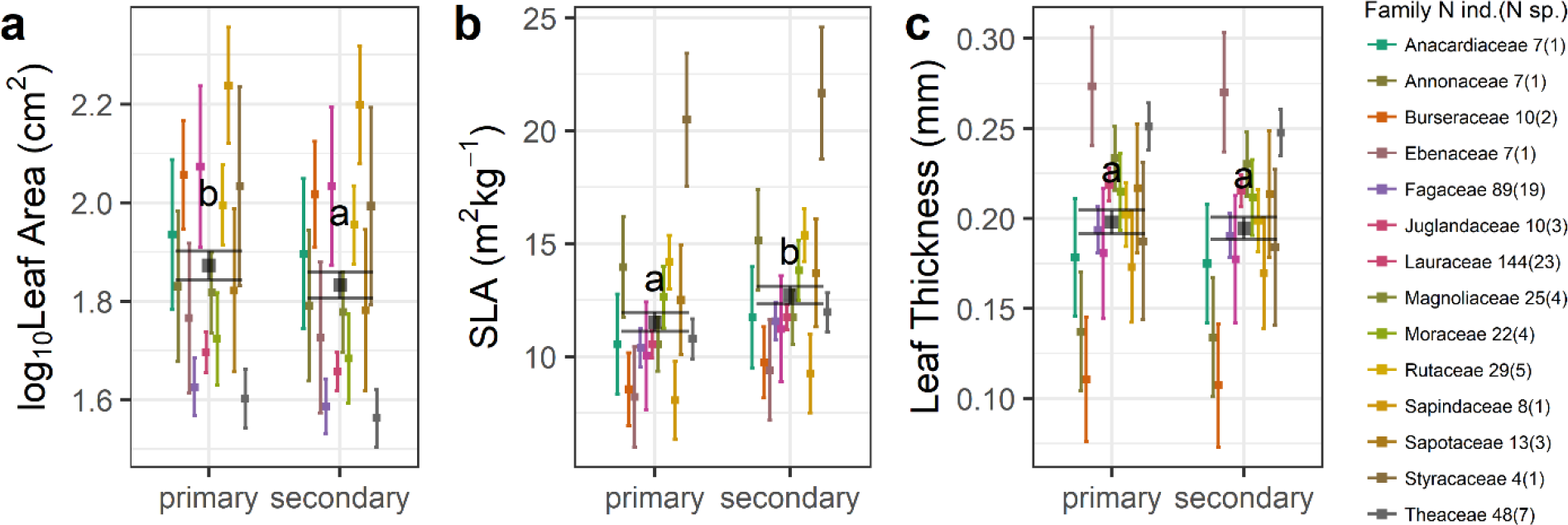
Least-square mean (points) with 95% confidence intervals (vertical bars) for three morphological leaf traits. Leaf area (a), specific leaf area (SLA) (b), and leaf thickness (c) by forest type and plant family using data from 423 individuals from 71 species sampled along a 6.6 km transect from secondary to primary forest. Values were predicted from linear models in the form of trait ~ forest type * species|family. Colored bars are predicted values by family and the grey bars are predicted values with respect to forest type. Count data for the number of individuals and species (in parenthesis) per family along the transect are shown in the legend. Letters denote the statistical groupings of post-hoc Tukey HSD tests. (See Supplement 2)

Regarding root morphology, the predicted-marginal mean difference in SRL, the belowground analog to SLA, was approximately 3.5 m kg^−1^ greater in secondary than in primary forest. In the secondary forest, values ranged from 25.96 (± 1.94) m kg^−1^ for species in the Annonaceae to 78.27 (± 1.14) for the Juglandaceae and was 32.97 (± 1.05) for Fagaceae species and 31.91 (± 1.03) for the Lauraceae family. In the primary forest, values ranged from 23.84 (± 1.14) m kg^−1^ for species in the Annonaceae to 71.87 (± 1.14) for the Juglandaceae and was 30.27 (± 1.05) for Fagaceae species and 29.30 (± 1.03) for Lauraceae saplings (Fig. 5b). Like the SLA-leaf area trends, root-system diameter had an opposite relationship to SRL, being generally 0.04 (±1.01) mm narrow in secondary than in primary forest. That trend was statistically significant (Table 2: F = 15.58, *p* < .001, ω^2^ = 0.004) and consistent across species (Fig. 4a). No difference in root tissue density with respect to forest type was detected, although there was substantial interfamilial variation (Fig. 4c). For example, of the 14 families samples, root tissue density was lowest for Sapotaceae, measuring 0.27 (±1.07) g cm^−3^ in both secondary and primary forest, and was greatest for Sapindaceae, measuring 0.64 (±1.05) g cm^−3^ in both secondary and primary forest. Root branching intensity averaged 1.72 (±1.02) tips cm^−1^ in secondary and 2.01 (±1.02) tips cm^−1^ in primary forest (Fig 5d). For measured Lauraceae saplings, least-squared mean root branching intensity was 1.72 (±1.02) tips cm^−1^ in the secondary forest and 1.48 (±1.02) tips cm^−1^ in the primary forest. For species in the Fagaceae, values were 2.09 (±1.03) tip cm^−1^ in the secondary forest and 1.80 (±1.03) tip cm^−1^. Those differences may seem small, but when ERS length is accounted for, being roughly 98 cm for all root systems measured, 94 cm for species in the Fagaceae family, and 95 cm for those in the Lauraceae, the 0.24 tip per cm difference in root topology scales up to between 23 and 24 root tips per 3-4 order root system.

**Table 2:**
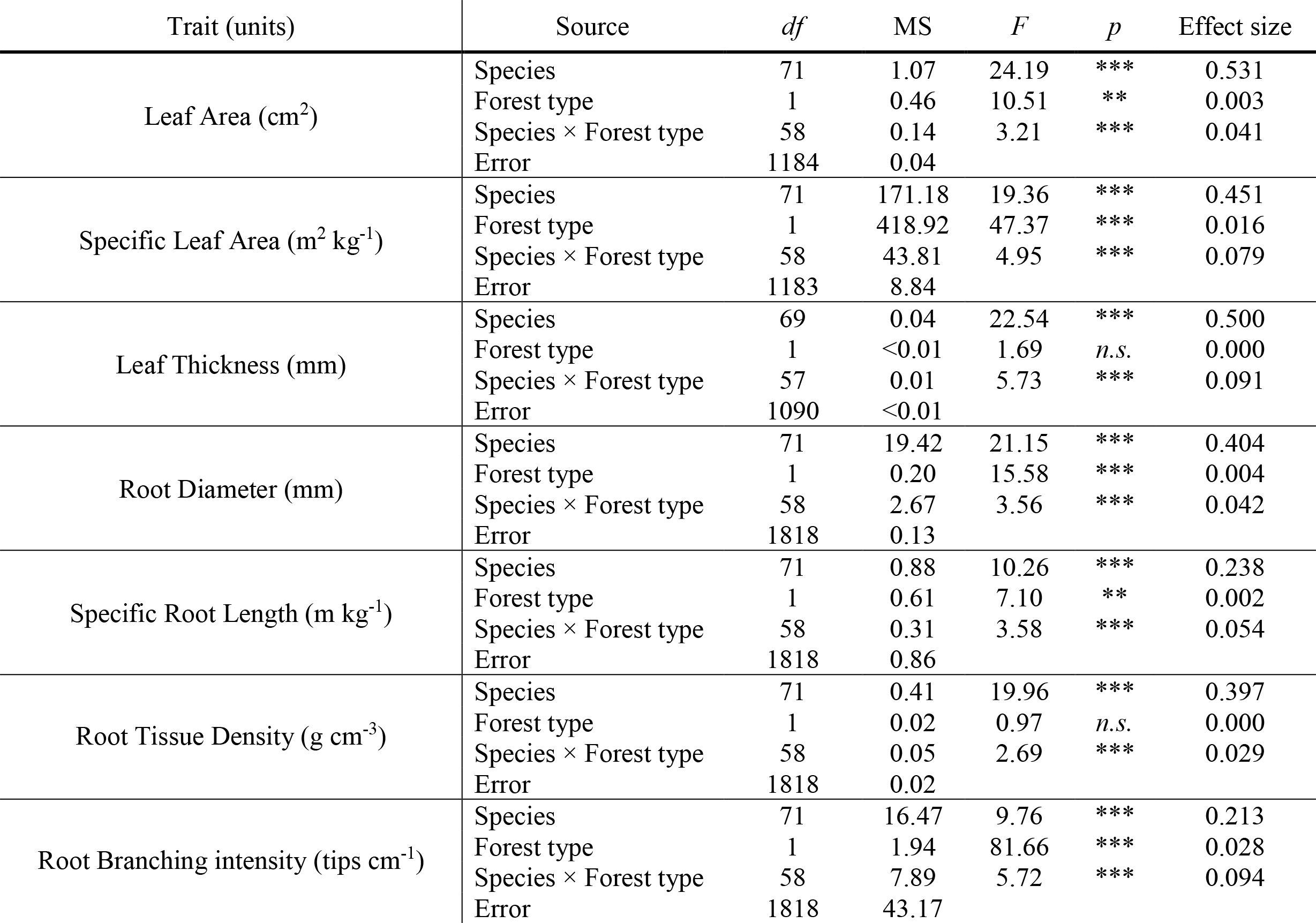
Analysis of variance table for linear models in the form: Trait ~ Species|Family × Forest type. Prior to model fitting, traits were log10 transformed in the case of leaf area, root diameter, SRL, root tissue density, and root branching intensity to improve data normality. *df* = degrees of freedom, MS = Mean squares, Effect size = ω^2^. \**p* <.05. \*\**p* < .01. \*\*\**p* < .001. *n.s*. = non-significant. Effect size (ω^2^) values near 0.01 are considered small, near 0.06 are considered medium and near 0.14 are considered large. (See Supplement 2)

**Figure 4:**
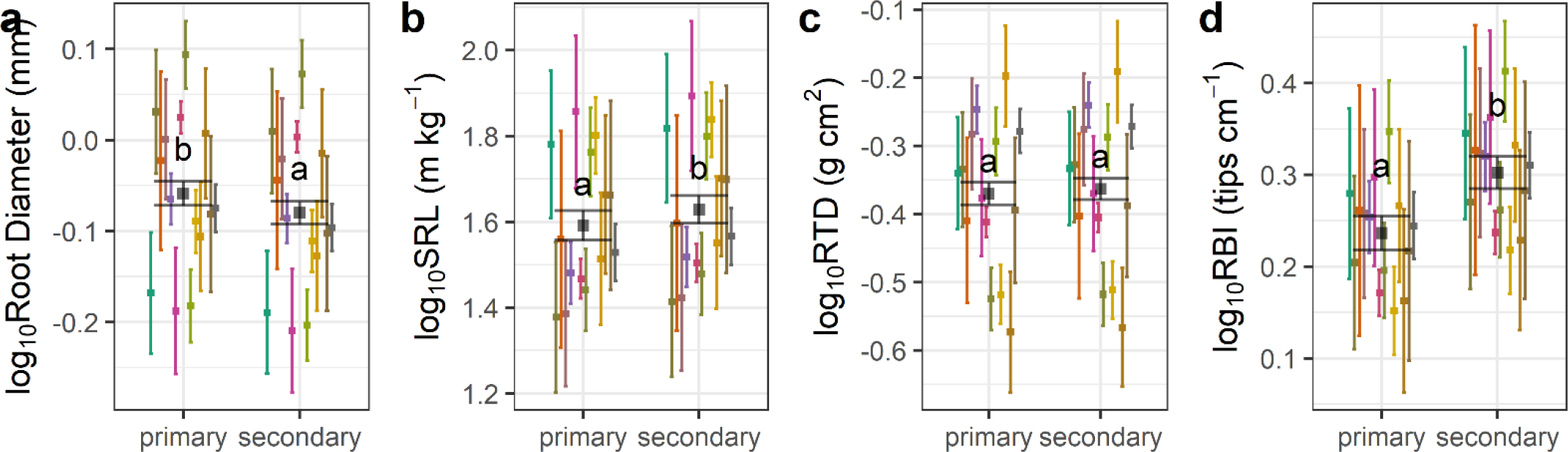
Least-squared means (points) and 95% confidence intervals (vertical bars) for four morphological root traits. Average root-system diameter (a), specific root length (SRL) (b), root tissue density (RTD) (c), and root branching intensity (RBI) (d) by forest type and plant family using data from 423 individuals from 71 species sampled along a 6.6 km gradient from secondary to primary forest. Colors correspond to the legend in Figure 3. Letters denote the statistical groupings of post-hoc Tukey HSD tests. (See Supplement 2)

**Figure 5:**
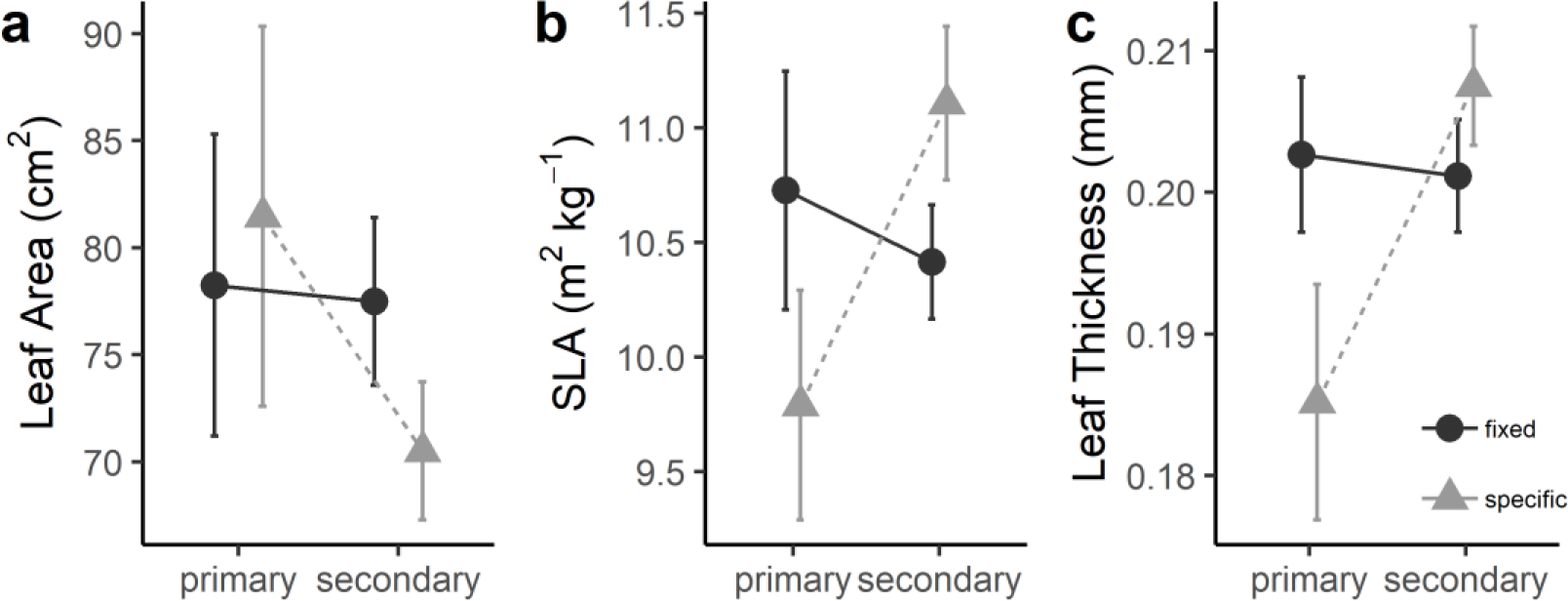
Assemblage-weighted mean trait values for leaf area, SLA and leaf thickness for the network of small plots (164 1/16th-ha plots; Fig. 1). Fifty-two of those plots are in primary forest, while 112 are in secondary forest with a history of logging. For each plot using data from 72 species, we weighted fixed and habitat-specific functional trait values (see methods) by basal area proportions; plot means with 95-percent confidence intervals are shown. (See Supplement 3).

Analysis of variance statistics for the three leaf and four root morphological traits measured (Table 2) confirmed the differences (Figs 3, 4). Forest type, as a two-level factor was found to be significant in two of the three leaf traits, excluding leaf thickness, and for three of the four root traits, excepting root tissue density. Effects sizes for forest type on leaf traits were small, being 0.003 for leaf area and 0.016 for SLA. For root traits, forest type effect sizes were 0.004 for root system diameter, 0.002 for SRL, and 0.028 for branching intensity. Interspecific variation in traits was always significant with very large effect sizes, which spanned from 0.213 for root branching intensity to 0.531 for leaf area. Interactive effects between species and forest type were small to medium, and statistically significant in all cases, even when forest type was non-significant, as was the case for leaf thickness and root tissue density.

### Scaling-up to the plots across the forest: Lepš’ Interspecific variability effects

We were interested in exploring how ITV by forest type translates to variation in traits at the community level. We used assemblage-weighted means (i.e. incomplete community-weighted means) and Lepš’ interspecific variability effects to quantify the amount of trait total interspecific variation due to species turnover and ITV by forest type. Both leaf and root morphologies were plastic with environment. Interspecific differences in organ morphologies outweighed differences due to forest type, however, the more-subtle differences due to forest-type were consistent. Effect sizes for forest type and forest type plant-lineage interactions as variables in the simple linear models of trait-species and environment relationships were nearly always statistically significant (Table 2). Their magnitude varied from small to medium in most cases, whereas the effect sizes for plant lineage (i.e. species) were always very large.

Differences in assemblage-weighted means among forest-types were larger between specific estimates (i.e. when forest type was accounted for in measuring functional traits) than for fixed ones (Figs. 6 & 7).

**Figure 6:**
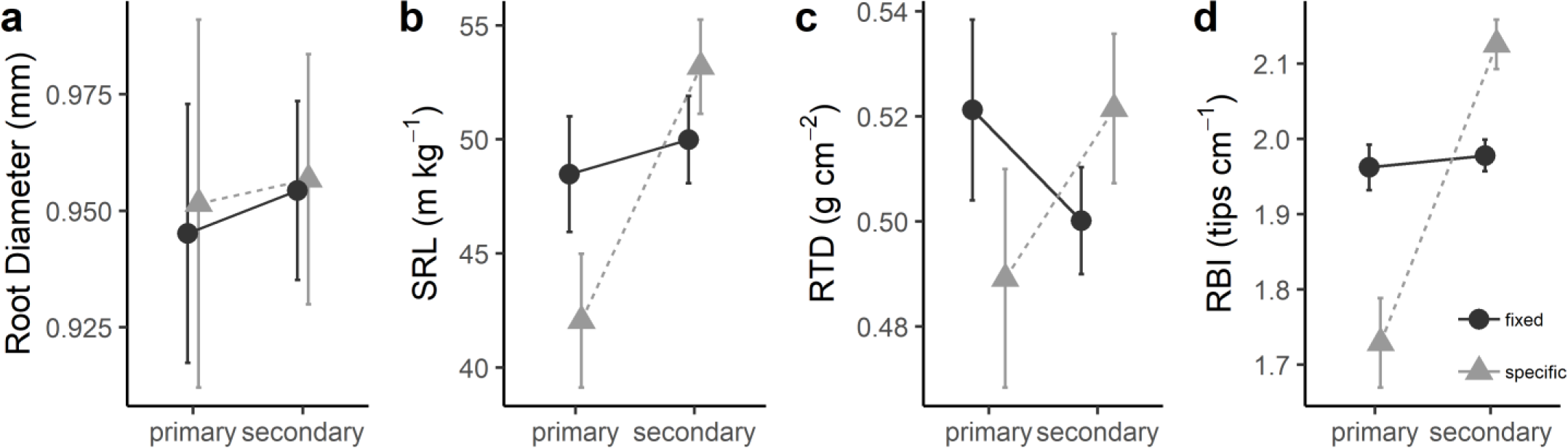
Assemblage-weighted mean trait values for root diameter, specific root length (SRL), root tissue density (RTD), and root branching intensity (RBI) by forest type for the network of small plots. Plot-level means, and 95-percent confidence intervals shown. (See Supplement 3)

For example, plot average assemblage weighted-mean estimates of leaf area were and 77.5 cm^2^ and 78.2 cm^2^(a 0.7 cm^2^ difference) for secondary and primary forest, respectively, when we used species trait means from the entire transect. Those estimates shifted to 70.5 cm ^2^ and 81.5 cm^2^(a difference of 11 cm^2^) when using forest-type-specific traits. This trend was consistent across all weighted-estimates of leaf traits (Fig. 5). For root traits, there was no difference in assemblage-weighted mean values by forest type for root diameter (Fig.6a), but differences were detected for SRL (Fig. 6b), RTD (Fig. 6c) and RBI (Fig 6d). The difference of specific SRL averages between secondary and primary forest plots was 11.1 m kg^−1^ as opposed to a difference of 1.5 m kg^−1^ for fixed weighted means. Those same respective differences (average difference between fixed and specific values between secondary and primary forest plots) were - 0.03 and 0.01 for root diameter, 0.03 and 0.02 g cm^−3^ for RTD and 0.40 and 0.02 for RBI.

The trait flex ANOVA statistics (Table 3) showed that forest type was statistically-significant in explaining trait variation in root branching intensity. In the case of branching intensity, forest type explained almost half of the trait variation (Table 4). For the other functional traits, the relative contribution of forest type (i.e. environmental variation in soil fertility and texture, Table 1) to trait variation ranged from 2 and 19 percent. At the plot level, nearly all this variation was due to ITV, as opposed to species turnover (i.e. compositional differences). Five of seven traits showed that ITV and species turnover covaried in a negative manner; Leaf area and SRL were the two excepting cases (Table 4). Residual variation in functional traits was large and was better explained by compositional turnover among plots within the two distinct forest types than ITV in assemblage-weighted mean trait values within them. There was negative co-variation between forest-type and ITV for leaf area, leaf thickness, root diameter, and SRL.

**Table 3:** Trait flex Analysis of Variance (ANOVA) table for forest type as a factor in assemblage-weighted functional trait variance. Statistics for residual variance not shown (see supplement 3). Demographic and functional trait data for 72 species (i.e. only those for which trait data was directly measured in the field). Fixed assemblage-weighted mean values use all functional trait data from along the transect, whereas specific assemblage-weighted means match functional trait data for plants to plots with corresponding forest-type classification. Intraspecific variability = fixed – specific. Abbreviations are as follows: *df* = degrees of freedom, SS = sum of squares, *F* = F-statistic, *p* = probability of obtaining the F statistic. \**p* <.05. \*\**p* < .01. \*\*\**p* < .001. *n.s*. = non-significant. Note that in this case, the mean squared error is equal to the sum of squares because there is 1 degree of freedom, therefore mean squared errors not shown. (See Supplement 3)

| Trait (units) | Fixed |  |  |  | Specific |  |  |  | Intraspecific variability |  |  |  |
| --- | --- | --- | --- | --- | --- | --- | --- | --- | --- | --- | --- | --- |
| | $df$ | SS | $F$ | $p$ | $df$ | SS | $F$ | $p$ | $df$ | SS | $F$ | $p$ |
| Leaf Area (cm<sup>2</sup>) | 1 | 20.00 | 0.04 | 0.84 | 1 | 4260.00 | 8.15 | ** | 1 | 3696.90 | 47.42 | *** |
| Specific Leaf Area (m<sup>2</sup> kg<sup>-1</sup>) | 1 | 3.46 | 1.50 | 0.22 | 1 | 61.50 | 19.10 | *** | 1 | 94.12 | 112.14 | *** |
| Leaf Thickness (mm) | 1 | 8.00E-05 | 0.19 | 0.67 | 1 | 0.02 | 28.28 | *** | 1 | 0.02 | 89.12 | *** |
| Root Diameter (mm) | 1 | 3.01E-03 | 0.29 | 0.59 | 1 | 0.04 | 3.15 | 0.08 | 1 | 0.07 | 18.42 | *** |
| Specific Root Length (m kg<sup>-1</sup>) | 1 | 81.30 | 0.83 | 0.36 | 1 | 4400.80 | 37.20 | *** | 1 | 3285.60 | 88.93 | *** |
| Root Tissue Density (g cm<sup>-3</sup>) | 1 | 8.99E-03 | 0.73 | 0.39 | 1 | 0.04 | 6.49 | * | 1 | 0.10 | 86.33 | *** |
| Branching Intensity (tips cm<sup>-1</sup>) | 1 | 1.59E-02 | 4.92 | * | 1 | 5.59 | 156.76 | *** | 1 | 5.15 | 275.00 | *** |

**Table 4:** Trait flex ANOVA results. The relative contribution of forest type vs unexplained variation in assemblage-weighted mean trait values. The relative contribution is calculated by diving the sum of squares attributable to that component by the total sum of squares for total variation (i.e., for specific traits, see Table S3). The statistical procedure follows Lepš et al. 2011 (see Supplement 3)

| Trait (units) | Relative Contribution of Forest Type |  |  |  | Relative Residual Contribution |  |  |  |
| --- | --- | --- | --- | --- | --- | --- | --- | --- |
|  | Turnover | Intraspecific variability | Covariation | Total = Specific average | Turnover | Intraspecific variability | Covariation | Total = Specific average |
| Leaf Area (cm<sup>2</sup>) | 0.02 | 4.16 | 0.61 | **4.79** | 91.28 | 14.21 | -10.28 | **95.21** |
| Specific Leaf Area (m<sup>2</sup> kg<sup>-1</sup>) | 0.59 | 16.14 | -6.18 | **10.54** | 63.94 | 23.31 | 2.21 | **89.46** |
| Leaf Thickness (mm) | 0.07 | 16.93 | -2.14 | **14.86** | 58.66 | 30.78 | -4.31 | **85.14** |
| Root Diameter (mm) | 0.14 | 3.09 | -1.32 | **1.91** | 78.42 | 27.16 | -7.49 | **98.09** |
| Specific Root Length (m kg<sup>-1</sup>) | 0.35 | 13.94 | 4.39 | **18.67** | 66.99 | 25.40 | -11.06 | **81.33** |
| Root Tissue Density (g cm<sup>-3</sup>) | 1.65 | 10.54 | -8.33 | **3.85** | 54.27 | 19.78 | 22.10 | **96.15** |
| Branching Intensity (tips cm<sup>-1</sup>) | 0.08 | 45.31 | 3.79 | **49.18** | 17.49 | 26.69 | 6.64 | **50.82** |

### Validation

A common issue in the linkage of plant functional traits to the community arises from sparse trait matrices (i.e. incomplete sampling of traits for taxa present in vegetation-monitoring plots). In this study, we directly measured functional traits for roughly one-eighth of the species in the tropical forest community (72 of 582 tree species). Accounting for ITV with environment (i.e. forest-type), we did so in a manner that targeted 14 families from across the Angiosperm phylogeny, and to validate the results from the Intraspecific Variability Effects analysis using our sub-sample of the community, we used a pGLM to impute missing trait values, then re-ran analyses with complete trait matrices. Missing trait values were imputed separately for fixed and specific trait matrices (i.e. by forest type).

Trends from the assemblage-weighted analyses were largely confirmed using imputed, complete matrices for functional traits. Variability among weighted trait estimates for plots decreased (i.e. 95% confidence interval breadth shrunk). For leaf traits (leaf area, SLA, and leaf thickness), no qualitative difference emerged from using imputed traits (Fig S5) versus only measured trait data (Fig. 5), however, the relative contribution of forest-type in explaining the total amount of trait variation increased markedly. Using only measured data, the total relative contribution of forest type for leaf area, SLA and leaf thickness was 5, 11, and 15 percent, respectively (Table 4). The relative contribution of forest-type for those three leaf traits increased to 26, 24, and 19 percent, respectively when pGLM-imputed trait matrices were used in the analysis (Table S4). Nearly all this variation was due to ITV, and not to species turnover (i.e. beta diversity, relative contributions < 1%) among the small vegetation plots (Table S4).

When using imputed trait matrices in the analyses, results for root traits tracked the same trend as leaf functional traits. Qualitatively, assemblage-weighted means (Fig. 6) were very similar to the community-weighted means using pGLM-imputed trait matrices (Fig. S6), with the only difference being in relation to root diameter. Using assemblage-weighted means, no difference was detected between fixed and specific estimates for root diameter (Fig 6a), however when the imputed data for the complete community were used, the specific estimates for root diameter increased slightly for primary forest and decreased for secondary forest, creating difference the estimates and between forest-types (Fig. S6a). With respect to the three other root traits (SRL, RTD, and RBI), differences were accentuated slightly, as compete functional trait matrices were used (e.g. Fig. S6 b, c & d). The relative contribution of forest-type to functional trait variation for root diameter, SRL, RTD and RBI was 2, 19, 4, and 49 percent (Table 4), respectively, using only directly-measured functional traits, whereas the relative contribution to forest-type increased to 12, 24, 9, and 77 percent, respectively, when using pGLM-imputed data. Again, this discrepancy in forest-type weighted-trait estimates was a result of ITV with environment, as opposed to species turnover in species composition among plots (Table S4). In many cases, for example for SLA, root diameter, SRL, and RTD, there was ITV and species turnover had negative covariance values (Table 4, Table S4), meaning that beta diversity in species composition drives weighted-trait values one way, while ITV in relation to environment causes shifts in the opposite direction.

## Discussion

### The utility of functional perspectives to species-rich tropical forests

The application of functional perspectives (Keddy, 1992, Westoby and Wright, 2006) to understand the ecology of plants in relation to environmental variation is a promising way to reduce complex and subtle changes in species composition and phenotypic plasticity in functional trait composition of communities into ecologically-meaningful information. This has been demonstrated regarding patterns of community assembly (Kraft et al., 2008, Spasojevic and Suding, 2012, Laughlin, 2014) and their underlying variability in plant traits (Letcher et al., 2015, Umaña et al., 2015, Muscarella et al., 2016). Our data confirm that interspecific trait variation in tropical tree communities is substantial (Figs. 4 and 5); such variation has been shown to account for key ecological differences among species (i.e. rates of photosynthesis and growth (Shipley et al. 2006, Wright et al. 2010), intrinsic rates of population growth (Adler et al., 2014), or variation in ecosystem productivity (Lohbeck et al., 2013, Poorter et al., 2017).

Yet, there has been a burgeoning interest in applying the utility of the functional perspective toward intraspecific, as opposed to interspecific trait variation (i.e. ecologically accounting for species trait variance rather than the mean trait values) (Albert et al., 2010, Violle et al., 2012, Siefert et al., 2015). Undoubtedly, some amount of ITV is related to environmental variation, but difficulty exists in quantifying dimensions of niche space, or specific ecosystem properties that systematically relate to this variation (Lavorel and Garnier, 2002, Shipley et al. 2016). Furthermore, there is the possibility of genetic variation, and genetic × environment interactive effects (Knustler et al., 2012, Soliveres et al., 2014, Yang et al. 2018). Regardless, the two approaches, a focus on interspecific or ITV, employ drastically different sampling approaches. On one hand, diversity-differences are emphasized, on the other, sampling effort is maximized at the expense of diversity-driven differences. In the tropics, the costs of this trade-off are particularly acute, as species diversity and environmental variation are both extensive. A second practical difficulty arises in that species compositional differences are often confounded with environmental variation (i.e. environmental filtering), making it difficult to assess ITV for all taxa, especially less-common ones. Despite this, Lepš et al (2011) propose a middle of the road solution that appeals, group trait sampling by *a priori* ecological classification (i.e. measure habitat or experimental-treatment specific trait values). This proved useful in the assessment of fertilization and mowing effects on the traits of European grasslands (Lepš, 1999, Lepš et al., 2011).

We employed a similar approach to assess how environmental variation in resource availability (i.e., soil nutrient content and light availability), roughly summarized by the successional progression of a forest (Odum, 1969, Christensen and Peet, 1984, Guariguata and Ostertag, 2001) leads to functional and compositional community change. In our case, the main measure of variation, irrespective of compositional differences among forest-types, was a gradient in soil fertility and texture, with soil texture and fertility decreasing along the transect from more-secondary to primary forest (Table 1), although observationally the understory light availability decreased as well. We did so with the principal objective of investing how variation root functional traits are with such environmental variation; an objective that has been identified as current research frontier in belowground functional ecology (Bardgett et al., 2014, Weemstra et al., 2016, Laliberté, 2017).

### Intraspecific leaf and root variation with tropical forest successional status

The intraspecific leaf trait variation measured along the JFL transect is consistent with previous research. For example, Fajardo & Siefert (2018) reported that ITV was greater than species turnover, accounting for 49% of the variation in leaf mass per area (1/SLA) in four temperate rainforests plots of southern Chile. Our estimates were 4% and 21% for assemblage-weighted, and community-weighted intraspecific SLA variation (Tables 4 and S4, respectively). Considerable differences in species richness exist between the temperate rainforests of Sothern Chile and our site, and there are notable differences sampling effort of traits and forest demography, yet, we still confirm their result that substantial intraspecific leaf-trait variation exists with environment. Subtle differences in leaf morphology with environment interact with species relative abundances across the landscape positively to the area-based leaf axis, and inverse to the mass-based axis of leaf morphological variation. This is shown by the positive covariation for leaf area and negative covariation of SLA, between species turnover and ITV (Table 4, Table S4), which is a similar result to Fajardo & Seifert (2018).

Generally, root morphologies varied more than leaf morphologies did with environmental variation in forest type. ITV in root diameter and root tissue density was less than that of SRL and root topology (i.e. branching intensity). These results suggest that, at least with respect to the four traits chosen in this study, there exist two or more axes of root morphological variation (Weemstra et al. 2016, Kramer-Walter et al. 2016). One that encompasses root diameter and root tissue density, which we found to be less-plastic and constrained by plant lineage (i.e. the evolved root strategy of the species), whether that be one of thicker or thinner roots (Valverde-Barrantes et al., 2017, Maherali, 2017). The second, which was more variable with soil-fertility and texture, seeks to optimize root structural investment into a topology that is most efficient for nutrient foraging, with some potential trade-off between purely abiotic and abiotically-assisted ways in doing so, (e.g. associations with mycorrhizae, chemical alterations of the rhizosphere, etc.) (Fitter 1991, Lynch et al 2005). SRL, or the distance a root can travel for a given amount of structural investment, competes with the production of absorptive fine-root tips (i.e. branching intensity).

We show that tropical forest juvenile trees modulate this axis of root architectural trade-offs in as they explore the soil for mineralized nutrients. In soil where nutrients are more abundant, root systems prioritize acquisitive architectures (i.e. higher SRLs and RBIs, Fig 5), yet where competition for soil nutrients is high (i.e. in primary forest), root systems engage more-conservative traits, such as increased root diameter, and likely try to maximize root lifespan to hold on to soil space. Other studies have found highly variable root system morphologies with the soil environment. Generally, N-depleted soils lead to increased lateral root elongation, while P-depleted soils cause more branched root systems (López-Bucio et al., 2003, Geihl et al. 2013). Notably, along very-strong Phosphorus (P) gradients, plant species have been shown to modulate root morphologies and nutrient acquisition strategies in relation to how they partition soil P, with more enzymatic activity and more-architecturally advanced roots in P-improvised soils (Lambers et al. 2006, Niu et al. 2013, Zemunik et al., 2015, Lambers et al., 2017). P-poor, highly-weathered, leached tropical soils, such as the yellow soils of JFL are also thought to have the ability to support a high diversity of plant species because of the way they allow plants to diversify and partition nutrient-use strategies in many ways within the soil environment (Laliberté et al., 2015, Turner, 2008, Turner et al., 2018). Moreover, in *Syzygium castaneum* roots in a tropical forest in Borneo, root diameter decreased, and specific root length and area and root phosphatase enzymatic activity increase with decreasing soil available P (Ushio et al., 2015). Soil Nitrogen, Phosphorus and base saturation all decrease from secondary to primary forest in JFL, both along the gradient (Table 1) and in the network of small plots (Supplement 1, Xu et al. 2015a).

### Consequences of ITV with tropical forest successional status

One convenient way in which plant traits are linked to communities is through community-weighted means (CWMs). On one-hand interpreting, CWM patterns of trait variation can help interpret ecological processes shaping community assembly from regional species pools (i.e. environmental filtering of species or even phenotypes) (Kraft et al. 2008, Muscarella and Uriarte, 2016). On the other, they are gross generalizations of the aggregate function of species assemblages, that may or may not reflect broader forest-level or ecosystem functioning (e.g. total photosynthetic, nutrient uptake, or biomass production capacity of a stand). Their accuracy depends on proper estimates of species traits in relation to the environment, which must account for both among and within species variation (Violle et al., 2012, Shipley et al. 2016, Yang et al. 2018). Measuring the morphology of all the plants within a community may one day be feasible but is currently not possible. Yet, the ability to accurately estimate trait values (e.g. leaf area, or more so even, belowground traits like root-tip abundance) is critical to accurately modeling how vegetation will respond to future changes in the biosphere (Warren et al. 2015, McCormack et al. 2017).

When we scaled-up trait measurements from along the JFL gradient to the forest using data from a network of small plots, we found that ITV compounded with compositional differences (i.e. species basal area weights) to create differences in community-weighted trait values with respect to forest successional status, where they would not have been detected has we not sampled functional traits along an environmental gradient consistent with the environmental variation found in the network of small plots (Figs. 6, 7, S4, S5). For example, along the JFL gradient, we measured a plant lineage-consistent decrease of about 3.5 m kg^−1^ in root-system SRL from secondary to primary forest. CWM difference in SRL between primary and secondary plots was >9 m kg^−1^ for the assemblage-weighted analysis and about 6.5 m kg^−1^ when pGLM-imputed data were used. Covariation between ITV and species turnover was positive for the assemblage-weighted mean estimate but was negative for more-complete analysis (Fig. 6, Table 4). Regardless, in the case of all traits as illustrated by the example of SRL, the difference in CWM trait estimates exceeded the magnitude of ITV measured in taxa across the JFL gradient. This is because successional shifts in species composition (Lohbeck et al., 2013, Letcher et al., 2015, Muscarella et al., 2016) compound with small change in ITV to move CWM trait values toward the conservative end of the plant economics spectrum (see Fig. S1, Table 4, Table S4).

In congruence with the second part of our first hypothesis, CWM trait shifts with forest successional status seemed to somewhat coordinate for roots and leaves, at least with respect to mass-area-based plant organ variation. SLA and SRL both showed consistently-conservative shifts in CWM values with increasing forest age. Both root diameter and leaf area showed acquisitive shits in within-species ITV, with roots becoming slightly thicker and leaves becoming slightly larger in primary forest. This is likely due to increased competition for light and nutrients as forest biomass increases, as well as differences in nutrient cycling dynamics that occur with increasing forest age (i.e. a shift from the exploitation of mineral nutrients to the recycling of organic nutrients), which could be coupled a shift in root strategy. Although no ITV in leaf thickness or RTD was measured along the gradient, which is consistent with previous research (Kramer-Walter et al., 2016b), including the study of root traits from many other Chinese forest sites (R. Wang et al., 2018), shifts in the relative abundance or species within plots (i.e. successional filtering effects), resulted in specific-CWM differences in trait values between forest types (Figs. 5c & S3c for leaf thickness & Figs. 6c & S4c). Our results speak to the need to create ecologically-relevant designs for the sampling of plant functional traits that account for environmental variation, especially as they relate to soil fertility, texture and forest age.

### Concluding remarks on the belowground functional ecology of tropical trees

Contrary to our first hypothesis, ITV in root traits became more conservative, in that entire root systems increased root-system diameter and were less topologically-developed (i.e. had fewer root tips per unit length). These results exemplify how, in an abiotic context (i.e. ignoring mycorrhizal and microorganisms in the rhizosphere), the soil environment feeds back in an abiotic to control on root morphology. Morphological shifts were observed along functional axes toward more-conservative root strategies, however one major limitation of this study lies in its inability to investigate if such shifts are accompanied with any compensation via biotic methods of acquiring nutrients (i.e. more association with mycorrhizal symbionts, increases in root enzymatic activity or shits in chemical partitioning of soil nutrients) among primary and secondary forest areas. Some research has shown root diameter to be correlated to mycorrhizal colonization and it would make sense that thicker roots, in later-successional forest, might be more-biotically reliant for in their nutrient-acquisition strategy (Phillips et al., 2013, Rosling et al., 2016, Kong et al., 2017).

Moreover, we are unable to assess if investment in root tissues (i.e. allocation to root biomass) is greater in primary than secondary forest, although some research has shown that root biomass increases with forest age (Odum 1969, Jackson et al. 1996). In such a case, it would be advantageous to have more a conservative strategy that maximizes root lifespan, as competition and soil space are come at a premium in late-successional forest, in comparison with more-secondary forest. Although we did find that differences in root diameter were highly-conserved and consistent across 14 Angiosperm families (Comas and Eissenstat, 2009, Kong et al., 2014, Valverde-Barrantes et al., 2017), we show some of the first evidence for tropical trees, that small-scale fluctuations in diameter are present and trade-off with investment of structural resources in root length (SRL) or tips (BI). However, we are not the first to demonstrate the utility of SRL as responsive root functional trait to soil environmental variation (Ostonen et al., 2007, Kong et al. 2014). In our case, we illustrate how minor differences in root morphology interact with environmental filters, in this case primarily a forest successional filter, to compound the effects of ITV, creating notable differences in forest-wide root functional morphologies that most-likely represent inherit differences in forest functioning (e.g. methods of nutrient acquisition and cycling dynamics).

## Supporting information

Supplement 1: Literate statistical document that compares the soils from 300 individuals from along the gradient to the soils of the 164 1/16th-hectar

Supplement 2: Literate statistical document with simple univariate Analyses of Variance and least-squared mean predicted values.

Supplement 3: Literate statistical document implementing the Trait Flex ANOVAS.

Supplement 4: Literate statistical document describing the gap-filling of functional traits and re-implementing the Trait Flex ANOVAS with gap-filled

## Acknowledgments

We thank the anonymous reviewers and Jan Lepš for comments that improved this work. JAH received support via a short-term fellowship from CTFS-ForestGEO at the Smithsonian. Additionally, we are grateful for many small personal donations that helped fund the soil analyses (http://www.experiment.com/chinaroots). We thank Sheyla Santana from the FIU GIS lab for help producing Figure 1. We acknowledge the assistance of the following people, who helped in the field and with the processing of plant samples: Shaojun Ling, Yaxin Xie, Jaming Wang, Siqi Yang, Wenguang Tang, Shitaing Ma, Qiqi Zhang, and Jiazhu Shi. Taxonomic field identification assistance was provided by Professor Yu from the Jianfengling Forest Bureau.

## Author Contributions Statement

JAH, OVB & CB designed the study, JAH, with help from QD and HX, collected the data, JAH analyzed the data and wrote the manuscript. All authors contributed intellectually and aided in the editing of manuscript drafts.

## Data Availability

Data have been archived on figshare: https://figshare.com/s/38440124ac7284ee3032 (DOI: 10.6084/m9.figshare.7996328)

## Supplementary Documents

Supplement 1: Literate statistical document that compares the soils from 300 individuals from along the gradient to the soils of the 164 1/16^th^-hectare small plots (data from XuDetto et al., 2015).

Supplement 3: Literate statistical document implementing the Trait Flex ANOVAS.

Supplement 4: Literate statistical document describing the gap-filling of functional traits and re-implementing the Trait Flex ANOVAS with gap-filled trait matrices.

## Appendix

**Figure S1:**
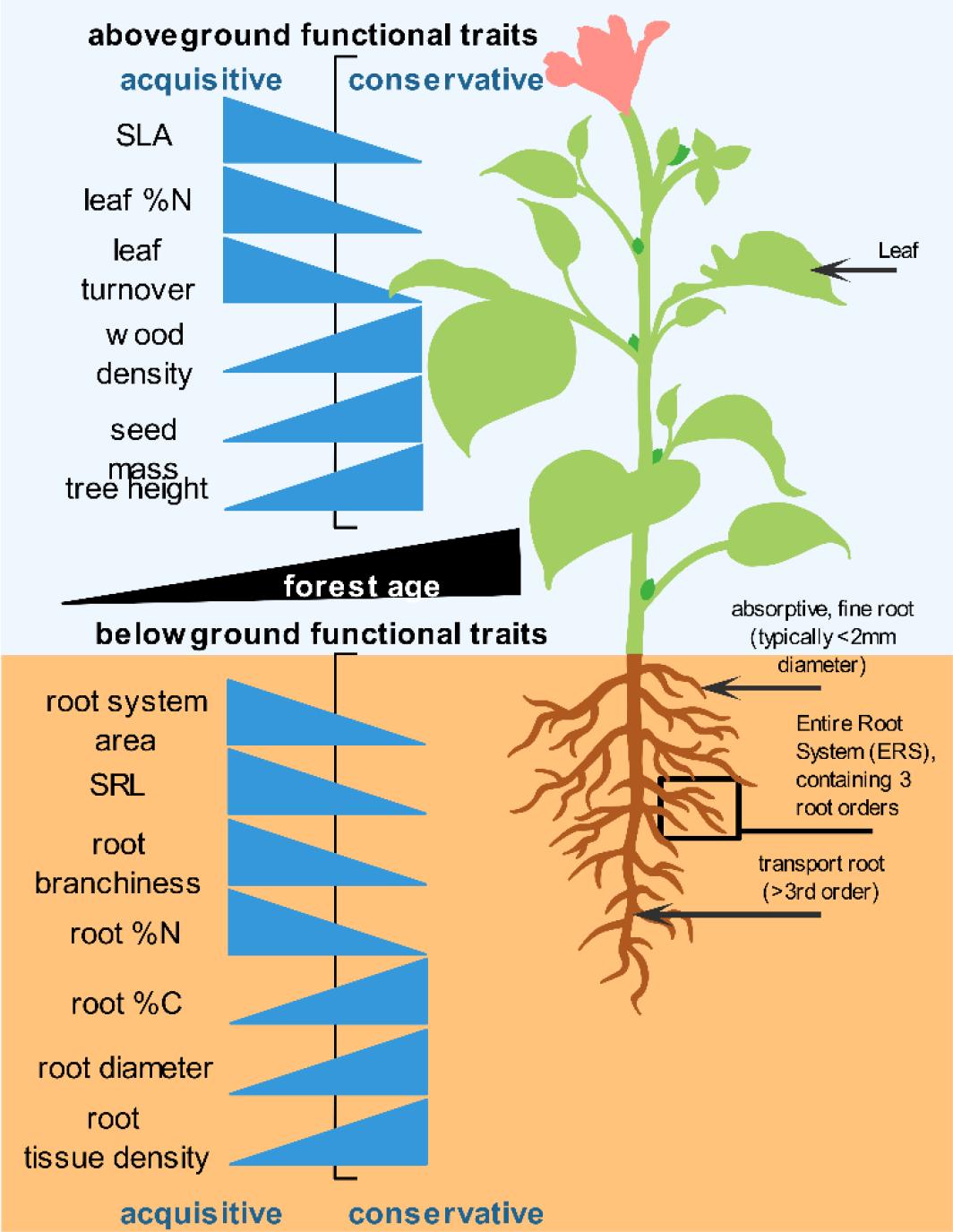
Conceptual diagram of some commonly-measured aboveground and belowground functional traits along an acquisitive-conservative life history continuum (i.e. fast-slow plant spectrum, *sensu* Reich, 2014). Increasing forest age (i.e. forest succession), shown in the black bar roughly corresponds to a shift in dominant plant strategies within a community from those with more-acquisitive traits to those with more conservative ones.

**Figure S2:**
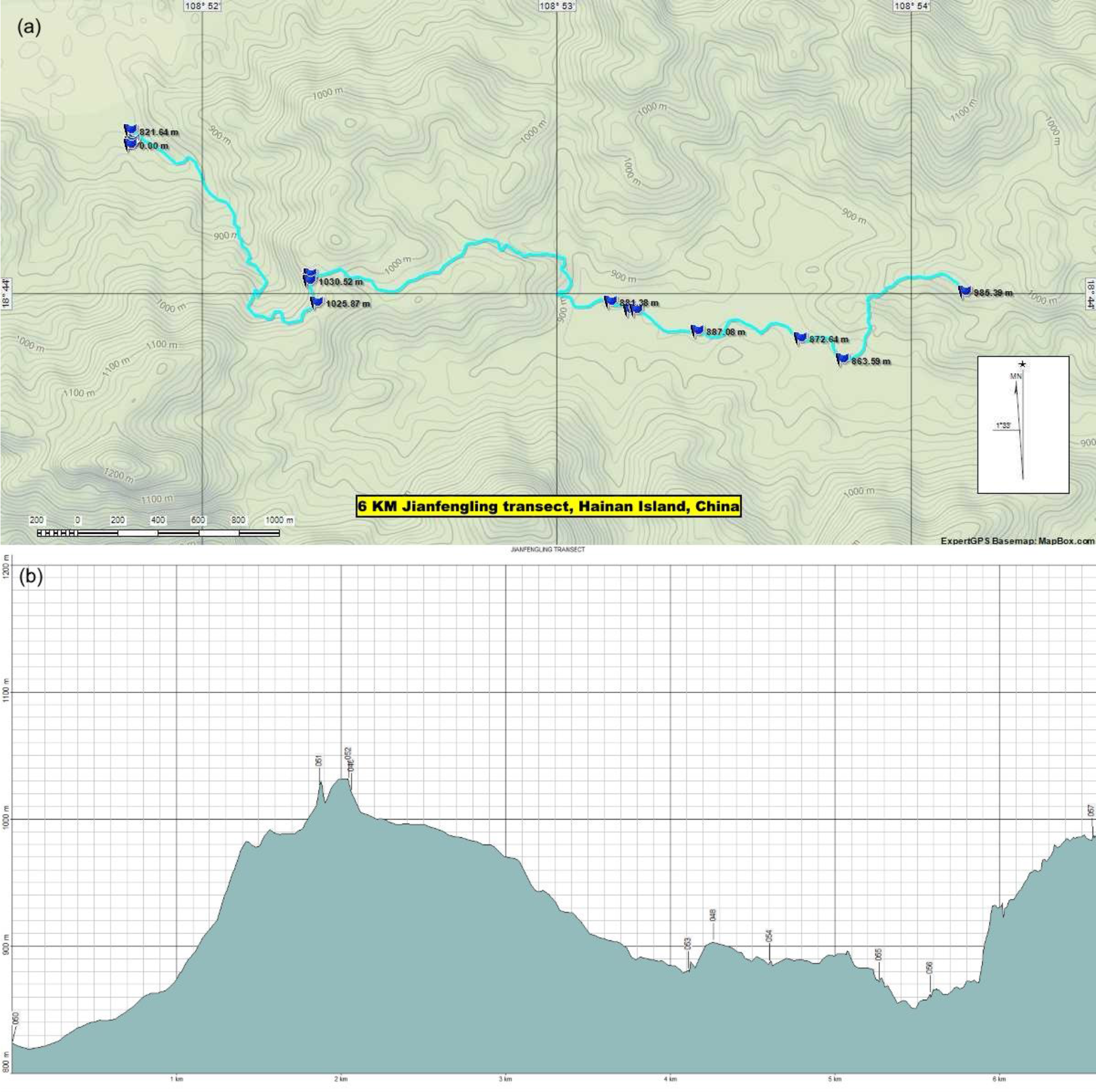
Detailed topographic map (a) and topographic profile (b) of the 6.6 km transect where functional traits of saplings were sampled. The transect started near the Jianfengling field house, at the entrance of the forest reserve. It progressed over one mountain, and over a stream (at km 4 of the transect) which delineates the secondary and primary parts of the forest. At roughly km 5.5 of the transect, the transect entered the 60-Ha CTFS-ForestGEO permanent forest dynamics plot.

**Figure S3:**
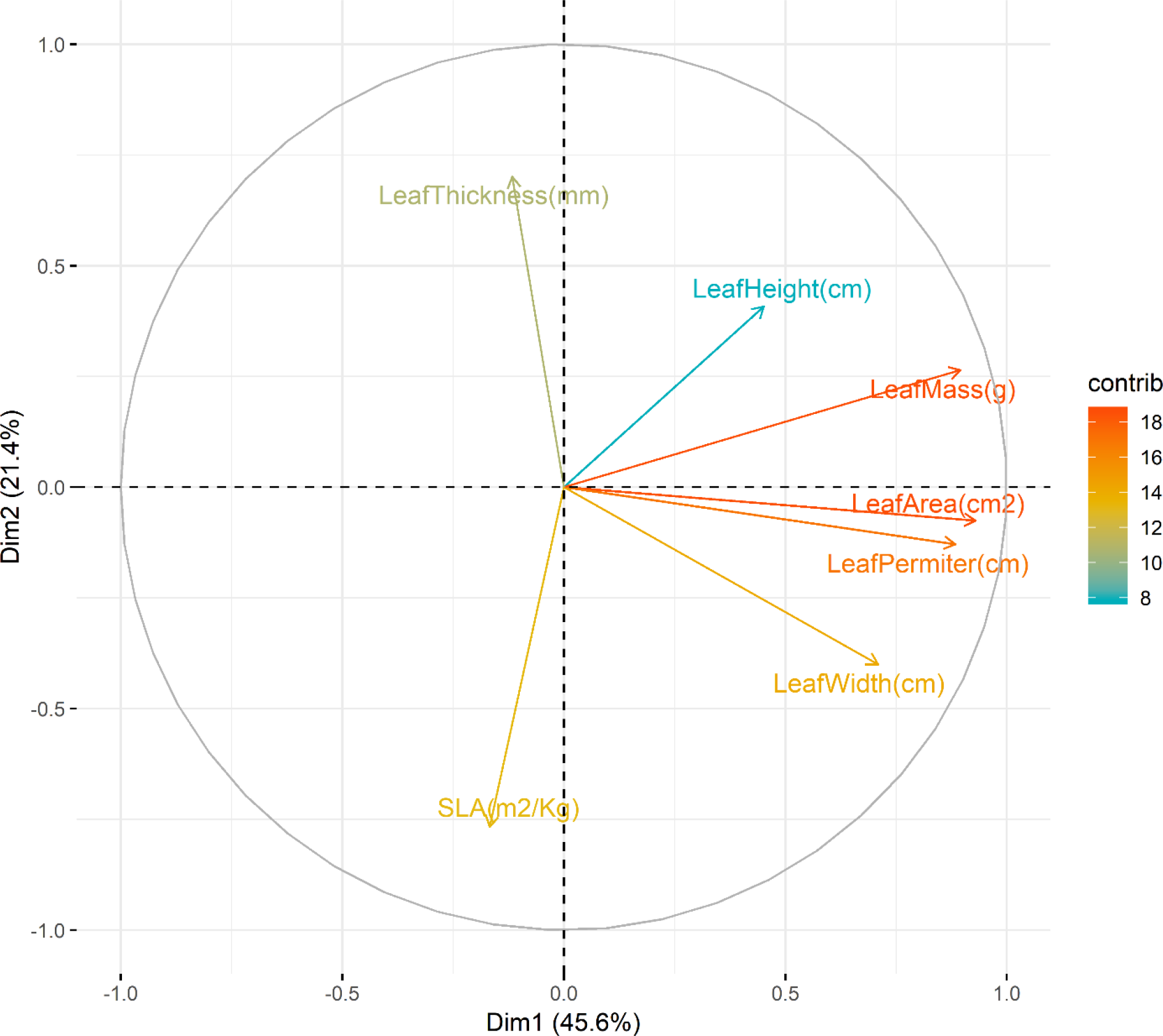
Principle components analysis (variable scores plotted) of 7 leaf traits measured on 423 saplings of 72 species. Raw trait measurements were first scaled and centered.

**Table S1:** PCA loading for 7 leaf traits. Traits used in downstream analyses bolded.

| Variable | PC1 | PC2 | PC3 |
| --- | --- | --- | --- |
| **LeafArea(cm<sup>2</sup>)** | 0.520 | -0.062 | 0.058 |
| LeafPerimeter(cm) | 0.494 | -0.105 | 0.057 |
| LeafWidth(cm) | 0.379 | -0.327 | -0.491 |
| LeafHeight(cm) | 0.253 | 0.333 | 0.738 |
| LeafMass(g) | 0.501 | 0.216 | -0.076 |
| **SLA(m<sup>2</sup>/kg)** | -0.094 | 0.336 | 0.336 |
| **LeafThickness(mm)** | -0.065 | 0.573 | -0.297 |

**Figure S4:**
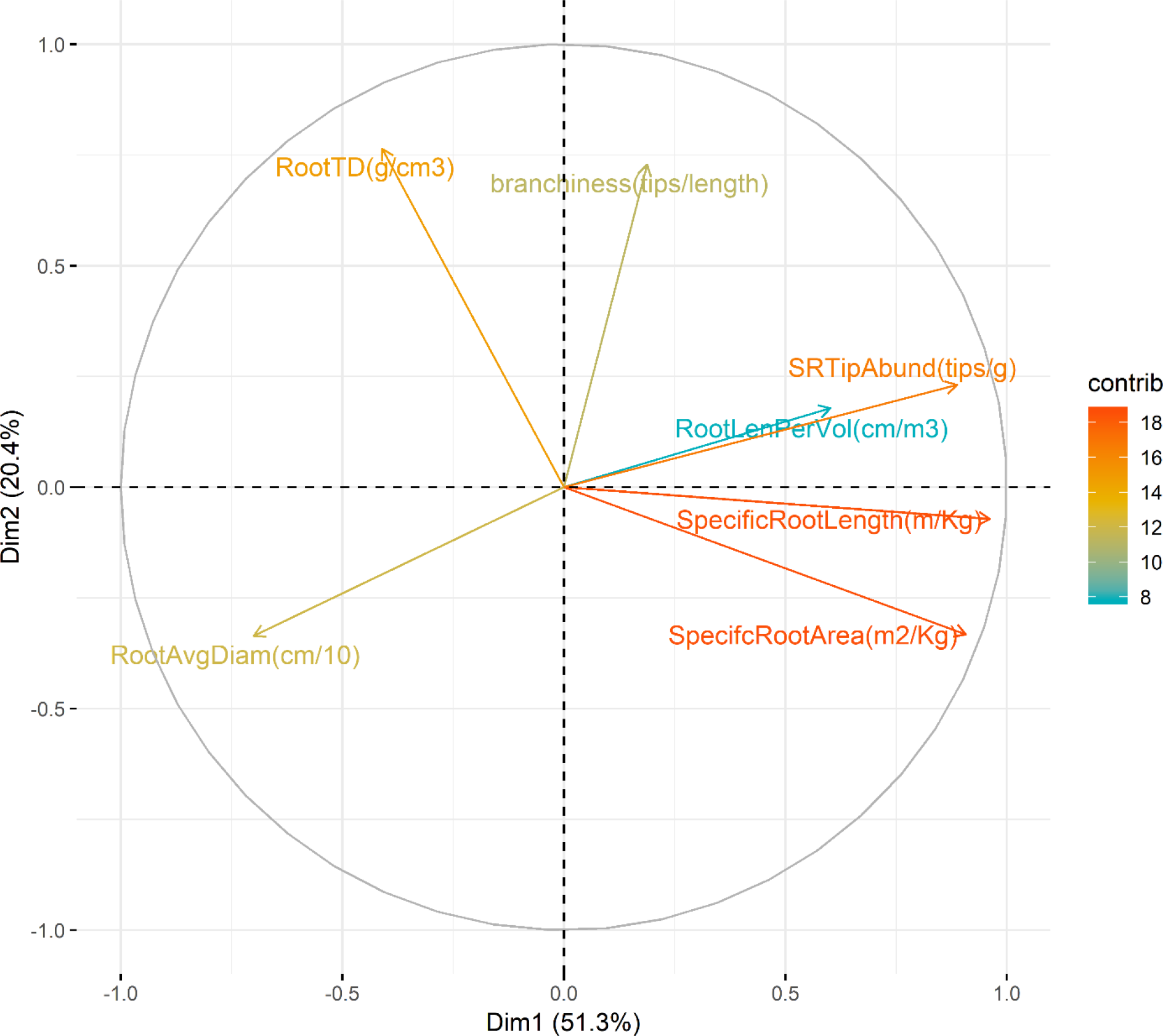
Principle components analysis (variable scores plotted) of 14 root traits measured on 423 saplings of 72 species. Raw trait measurements were first scaled and centered.

**Table S2:** PCA loadings for 7 root traits. Traits used in downstream analyses bolded.

| Variable | PC1 | PC2 | PC3 |
| --- | --- | --- | --- |
| **SpecificRootLength (m/kg)** | 0.508 | -0.060 | 0.012 |
| SpecifcRootArea (m<sup>2</sup>/kg) | 0.479 | -0.278 | 0.086 |
| **RootAvgDiam (mm)** | -0.369 | -0.281 | 0.372 |
| RootLenPerVol (cm/m<sup>3</sup>) | 0.317 | 0.149 | -0.532 |
| **RootTD (g/cm<sup>3</sup>)** | -0.216 | 0.639 | -0.345 |
| **Branchiness (tips/cm)** | 0.099 | 0.610 | 0.620 |
| SRTipAbund (tips/g) | 0.468 | 0.193 | 0.263 |

**Table S3:** Trait flex Analysis of Variance (ANOVA) table for forest type in community-weighted functional trait variance. Fore statistics for residual variance see supplement 4. Demographic and functional trait data for 582 species (i.e. using trait data was directly measured in the field). Fixed values use imputed functional trait data for a matrix containing species means along the entire transect, whereas specific weighted-means use imputed functional traits arising from matrices that limit species means for functional traits to either secondary or primary portions of the transect. Intraspecific variability = fixed – specific. Abbreviations are as follows: *df* = degrees of freedom, SS = sum of squares, *F* = F-statistic, *p* = probability of obtaining the F statistic. \**p* <.05. \*\**p* < .01. \*\*\**p* < .001. Note that in this case, the mean squared error is equal to the sum of squares because there is 1 degree of freedom, therefore mean squared errors not shown. (See Supplement 4)

| Trait (units) | Fixed |  |  |  | Specific |  |  |  | Intraspecific variability |  |  |  |
| --- | --- | --- | --- | --- | --- | --- | --- | --- | --- | --- | --- | --- |
|  | *df* | SS | *F* | *p* | *df* | SS | *F* | *p* | *df* | SS | *F* | *p* |
| Leaf Area (cm<sup>2</sup>) | 1 | 48.10 | 0.44 | 0.51 | 1 | 5548.40 | 56.97 | *** | 1 | 4563.30 | 271.24 | *** |
| Specific Leaf Area (m<sup>2</sup> kg<sup>-1</sup>) | 1 | 3.45E-4 | 5E-4 | 0.98 | 1 | 47.68 | 50.36 | *** | 1 | 47.91 | 317.85 | *** |
| Leaf Thickness (mm) | 1 | 6.76E-5 | 0.53 | 0.47 | 1 | 6.24E-3 | 38.69 | *** | 1 | 5.02E-3 | 150.38 | *** |
| Root Diameter (mm) | 1 | 9.01E-3 | 2.24 | 0.14 | 1 | 0.09 | 21.26 | *** | 1 | 0.16 | 192.09 | *** |
| Specific Root Length (m kg<sup>-1</sup>) | 1 | 50.10 | 1.63 | 0.20 | 1 | 1515.60 | 51.59 | *** | 1 | 2116.90 | 361.98 | *** |
| Root Tissue Density (g cm<sup>-3</sup>) | 1 | 1.91E-3 | 1.80 | 0.18 | 1 | 0.03 | 15.42 | *** | 1 | 0.04 | 132.44 | *** |
| Branching Intensity (tips cm<sup>-1</sup>) | 1 | 5.68E-3 | 1.50 | 0.22 | 1 | 4.58 | 528.60 | *** | 1 | 4.26 | 1480.70 | *** |

**Table S4:** Trait flex ANOVA results. The relative contribution of forest type vs unexplained variation in community-weighted mean trait values. The analysis uses complete functional trait matrices (i.e. those imputed using pGLMS). The relative contribution is calculated by diving the sum of squares attributable to that component by the total sum of squares for total variation (i.e., for specific traits, see Table S3). The statistical procedure follows Lepš et al. 2011 (see Supplement 4).

| Trait (units) | Relative Contribution of Forest Type |  |  |  | Relative Residual Contribution |  |  |  |
| --- | --- | --- | --- | --- | --- | --- | --- | --- |
|  | Turnover | Intraspecific variability | Covariation | Total = Specific average | Turnover | Intraspecific variability | Covariation | Total = Specific average |
| Leaf Area (cm<sup>2</sup>) | 0.23 | 21.40 | 4.39 | **26.02** | 82.38 | 12.78 | -21.19 | **73.98** |
| Specific Leaf Area (m<sup>2</sup> kg<sup>-1</sup>) | 1.72E-04 | 23.84 | -0.13 | **23.72** | 5.66E+01 | 12.15 | 7.49 | **76.28** |
| Leaf Thickness (mm) | 0.21 | 15.48 | 3.59 | **19.28** | 63.61 | 16.67 | 0.44 | **80.72** |
| Root Diameter (mm) | 1.16 | 20.10 | -9.65 | **11.60** | 83.80 | 16.95 | -12.35 | **88.40** |
| Specific Root Length (m kg<sup>-1</sup>) | 0.80 | 33.74 | -10.38 | **24.15** | 79.59 | 15.10 | -18.84 | **75.85** |
| Root Tissue Density (g cm<sup>-3</sup>) | 0.62 | 13.96 | -5.89 | **8.69** | 56.01 | 17.08 | 18.23 | **91.31** |
| Branching Intensity (tips cm<sup>-1</sup>) | 0.09 | 71.25 | 5.20 | **76.54** | 10.27 | 7.80 | 5.40 | **23.46** |

**Figure S5:**
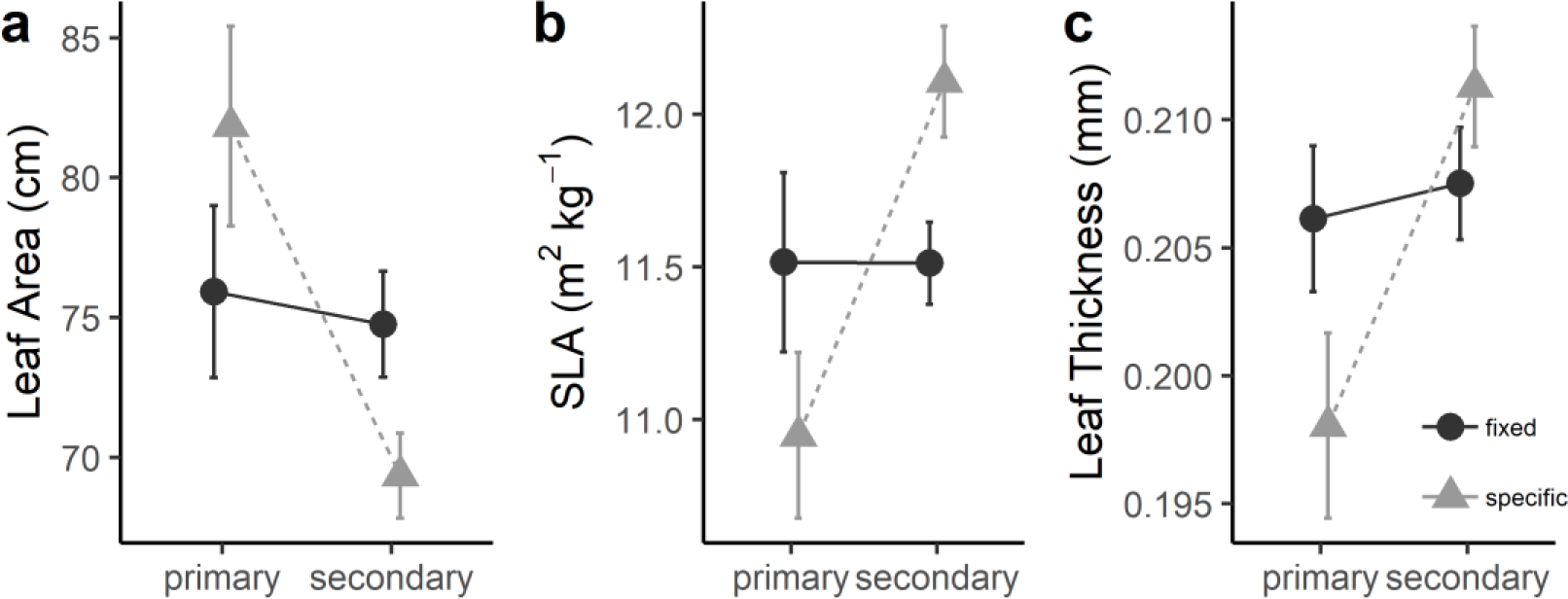
Community-weighted mean trait values for leaf area, SLA and leaf thickness for the network of small plots (164 1/16th-ha plots; see Fig. 1). Fifty-two of those plots are in primary forest, while 112 are in secondary forest with a history of logging. Compete functional trait matrices used in these analyses (i.e. those using trait imputation via pGLM). We weighted fixed and habitat-specific functional trait values (see methods) by basal area proportions; plot means with 95-percent confidence intervals are shown. (See Supplement 4)

**Figure S6:**
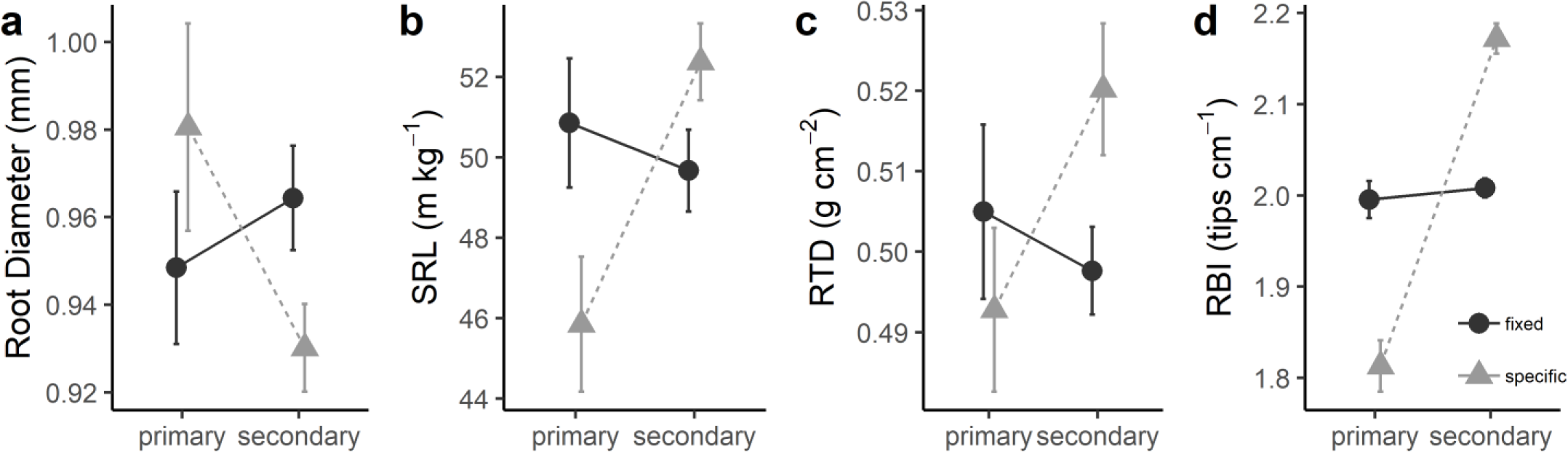
Community-weighted mean trait values for root diameter, specific root length (SRL), root tissue density (RTD), and root branching intensity (RBI) by forest type for the network of small plots. Complete functional trait matrices for all 582 species present in the plant community data for the network of small plots used in these analyses. Imputation of traits done using PhyloPars (pGLM) (Bruggeman et al., 2009). Means of CWM trait values and 95-percent confidence intervals shown. (See Supplement 3).

