## Supplement 1: Literate statistical document that compares the soils from 300 individuals from along the gradient to the soils of the 164 1/16th-hectar for "Intraspecific root and leaf trait variation with tropical forest successional status: consequences for community-weighted patterns"

### JFL Gradient vs. Small Plots - Soil Analyses

*JAH*

*October 9, 2017*

#### Background

In this literate statistical document, we look at the soil data from the Jianfengling (JFL) transect vs. the data from Dr. Han Xu's network of 164 1/16th of a Ha (25 x 25m) plots that are distributed throughout Jianfengling Forest Reserve.

His data are available as supplementary material for two key publications:

1. *Journal of Ecology* **103** 1325-1333. "Habitat hotspots of common and rare species along climatic and edaphic gradients"
2. *Journal of Applied Ecology* **52** 1014-1052. "Partial recovery of a tropical rain forest a half-century after clear-cut and selective logging"

We compare standaridized values of measured soil variables between 300 sites along a 6.6 km graident along the road from near the field station to well into the 60-Ha plot in Jianfengling to those of previously published.

- We look at these soil variables (units in parentheses):
  - pH (unitless)
  - soil [organic] Carbon (g/kg)
  - total Nitrogen (g/kg)
  - total Phosphorus (g/kg)
  - total Potassium (g/kg)
  - exchangeable Calcium (cmol(1/2Ca2+)/kg)
  - exchangeable Magnesium (cmol(1/2Mg2+)/kg)
  - available Potassium (cmolK+/kg)
  - alkali-hydrolyzable (available) Nitrogen (mg/kg)
  - available Phosphorus (mg/kg)

#### Summary Stats

##### Xu Han's 1/16th-Ha Plots

```
##           vars   n   mean    sd median trimmed   mad   min    max  range
## pH           1 163   4.81  0.29   4.77    4.80  0.22  3.81   6.09   2.28
## org.mat      2 163  15.55  5.82  14.80   15.20  5.71  4.10  39.80  35.70
## total.N      3 163   1.18  0.33   1.15    1.16  0.28  0.30   2.19   1.89
## total.P      4 163   0.12  0.05   0.11    0.11  0.03  0.06   0.58   0.52
## total.K      5 163  22.15 13.16  21.92   21.40 16.54  2.00  52.43  50.44
## exch.Ca      6 163   0.30  0.78   0.13    0.17  0.09  0.00   9.28   9.28
## exch.Mg      7 163   0.25  0.34   0.16    0.19  0.09  0.04   3.90   3.86
## avail.K      8 163   0.37  0.18   0.35    0.36  0.19  0.08   0.91   0.83
## alk-hyd.N    9 163 158.04 44.42 151.55  155.18 45.86 61.86 324.75 262.89
## avail.P     10 163   1.85  1.02   1.67    1.72  0.79  0.65   7.82   7.17
##           skew kurtosis   se
## pH        0.88     4.43 0.02
## org.mat   0.83     1.40 0.46
```

```
## total.N   0.60     0.79 0.03
## total.P   5.28    44.31 0.00
## total.K   0.37    -0.86 1.03
## exch.Ca   9.54   103.26 0.06
## exch.Mg   7.99    80.98 0.03
## avail.K   0.64    -0.21 0.01
## alk-hyd.N 0.65     0.39 3.48
## avail.P   2.33     8.88 0.08
```

##### soil300 - JFL Transect

```
##           vars   n   mean    sd median trimmed   mad   min    max  range
## pH           1 300   4.33  0.29   4.36    4.33  0.29  3.66   5.26   1.60
## org.mat      2 300  36.68 16.74  33.20   34.44 10.48 12.72 163.59 150.87
## total.N      3 300   1.41  0.49   1.31    1.36  0.39  0.54   3.55   3.02
## total.P      4 300   0.12  0.03   0.12    0.12  0.03  0.05   0.22   0.17
## total.K      5 300  17.64 10.64  18.23   17.52 10.73  0.33  42.32  41.99
## exch.Ca      6 300   0.51  0.78   0.26    0.33  0.21  0.01   7.00   6.99
## exch.Mg      7 300   0.20  0.17   0.16    0.17  0.10  0.01   1.49   1.48
## avail.K      8 300   0.22  0.11   0.19    0.21  0.08  0.06   0.67   0.60
## alk-hyd.N    9 300 181.47 61.13 168.15  174.06 48.35 71.17 463.08 391.91
## avail.P     10 300   1.45  1.79   1.10    1.15  1.30  0.10  15.95  15.85
##            skew kurtosis   se
## pH        -0.10    -0.51 0.02
## org.mat    2.47    11.73 0.97
## total.N    1.22     2.18 0.03
## total.P    0.64     0.28 0.00
## total.K   -0.01    -0.84 0.61
## exch.Ca    4.23    23.54 0.04
## exch.Mg    3.08    14.69 0.01
## avail.K    1.35     1.75 0.01
## alk-hyd.N  1.26     2.16 3.53
## avail.P    3.99    24.05 0.10
```

###### JFL-transect secondary

```
##           vars   n   mean    sd median trimmed   mad   min    max  range
## pH           1 150   4.39  0.28   4.40    4.40  0.28  3.80   5.26   1.46
## org.mat      2 150  36.56 15.67  33.66   34.90 12.70 12.72  94.09  81.37
## total.N      3 150   1.42  0.49   1.36    1.39  0.46  0.54   2.76   2.22
## total.P      4 150   0.13  0.03   0.13    0.13  0.03  0.05   0.22   0.17
## total.K      5 150  19.89 11.98  20.99   20.06 14.49  0.68  42.32  41.64
## exch.Ca      6 150   0.75  1.03   0.34    0.52  0.32  0.01   7.00   6.99
## exch.Mg      7 150   0.24  0.19   0.18    0.21  0.13  0.02   1.14   1.12
## avail.K      8 150   0.26  0.13   0.24    0.25  0.12  0.06   0.67   0.60
## alk-hyd.N    9 150 176.38 50.50 175.30  174.29 50.04 71.17 318.33 247.16
## avail.P     10 150   1.58  2.33   0.88    1.09  1.07  0.10  15.95  15.85
##            skew kurtosis   se
## pH        -0.01    -0.28 0.02
## org.mat    1.11     1.36 1.28
## total.N    0.56    -0.15 0.04
## total.P    0.34    -0.13 0.00
```

```
## total.K   -0.22    -1.10 0.98
## exch.Ca    2.97    11.15 0.08
## exch.Mg    1.78     4.47 0.02
## avail.K    0.79     0.07 0.01
## alk-hyd.N  0.45     0.11 4.12
## avail.P    3.46    15.08 0.19
```

**JFL-transect primary**

```
##           vars   n   mean    sd median trimmed   mad   min    max  range
## pH           1 150   4.27  0.29   4.31    4.27  0.31  3.66   4.86   1.20
## org.mat      2 150  36.80 17.80  32.48   33.97  9.22 15.31 163.59 148.28
## total.N      3 150   1.39  0.49   1.28    1.32  0.33  0.70   3.55   2.86
## total.P      4 150   0.11  0.02   0.11    0.11  0.02  0.07   0.17   0.11
## total.K      5 150  15.40  8.58  16.57   15.45  7.75  0.33  34.52  34.19
## exch.Ca      6 150   0.26  0.19   0.22    0.24  0.15  0.01   0.89   0.88
## exch.Mg      7 150   0.16  0.15   0.13    0.15  0.07  0.01   1.49   1.48
## avail.K      8 150   0.19  0.06   0.17    0.18  0.05  0.08   0.49   0.40
## alk-hyd.N    9 150 186.56 69.98 162.43  176.42 46.65 84.33 463.08 378.75
## avail.P     10 150   1.32  0.99   1.30    1.23  1.26  0.10   3.85   3.75
##            skew kurtosis   se
## pH        -0.15    -0.96 0.02
## org.mat    3.35    17.37 1.45
## total.N    1.86     4.53 0.04
## total.P    0.52     0.12 0.00
## total.K   -0.19    -0.74 0.70
## exch.Ca    1.22     1.03 0.02
## exch.Mg    5.72    44.84 0.01
## avail.K    1.45     3.29 0.01
## alk-hyd.N  1.41     1.76 5.71
## avail.P    0.55    -0.57 0.08
```

### Histograms

For the following histograms, the transect soil data are shown in red, and Xu Han's 1/16th-Ha plot data are shown in blue.

**pH**

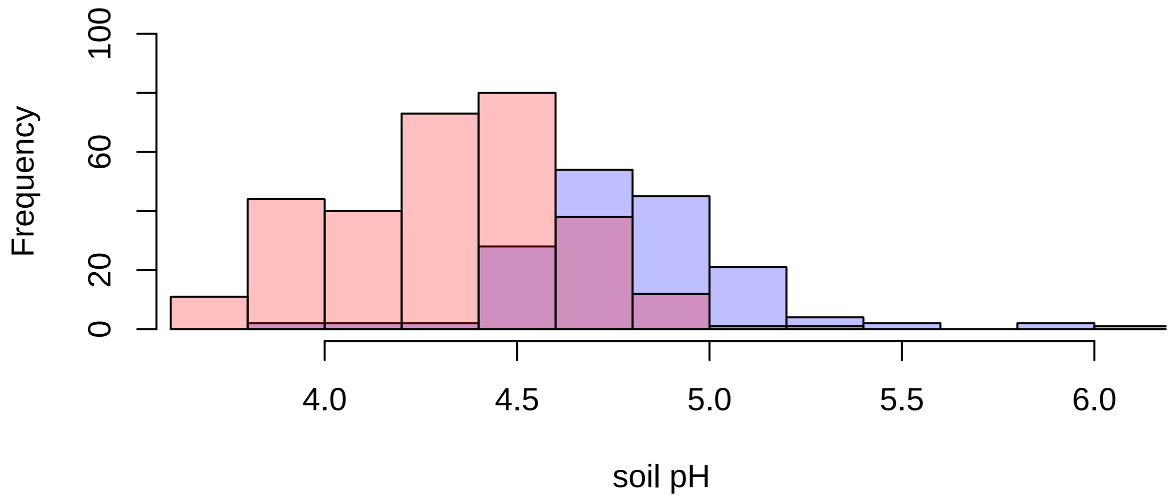


**Organic Carbon**

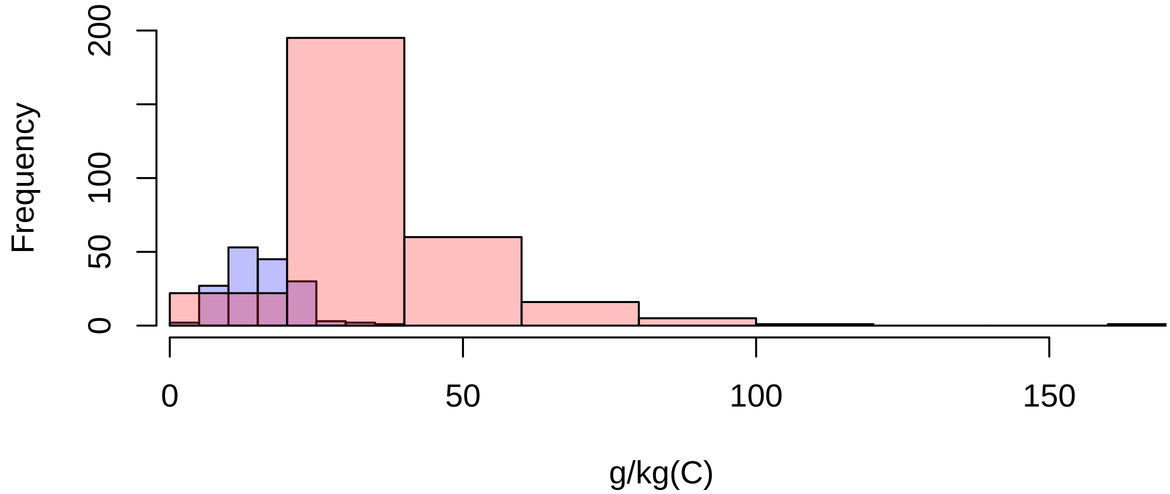

**Total Nitrogen**

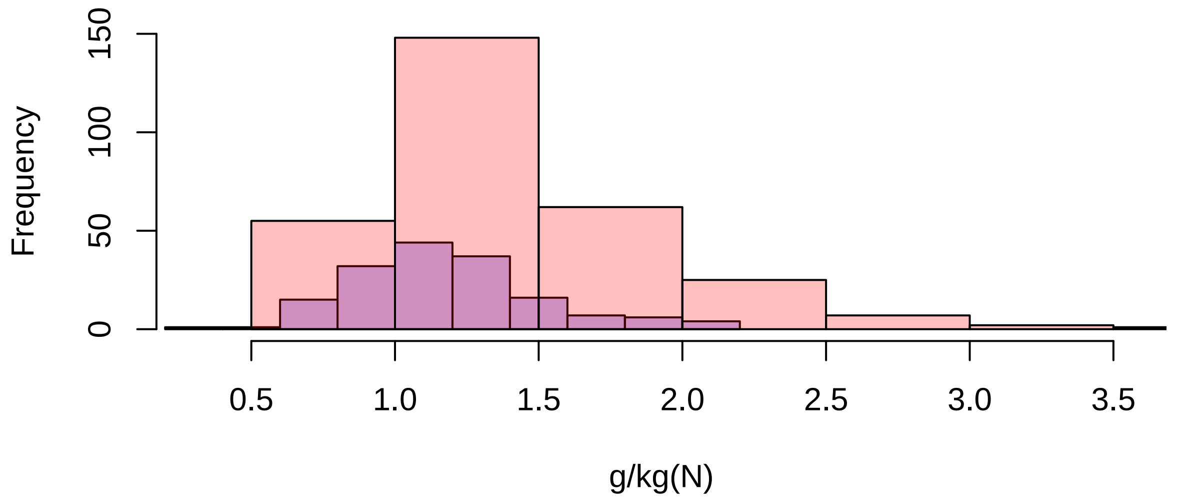


**Total Phosphorus**

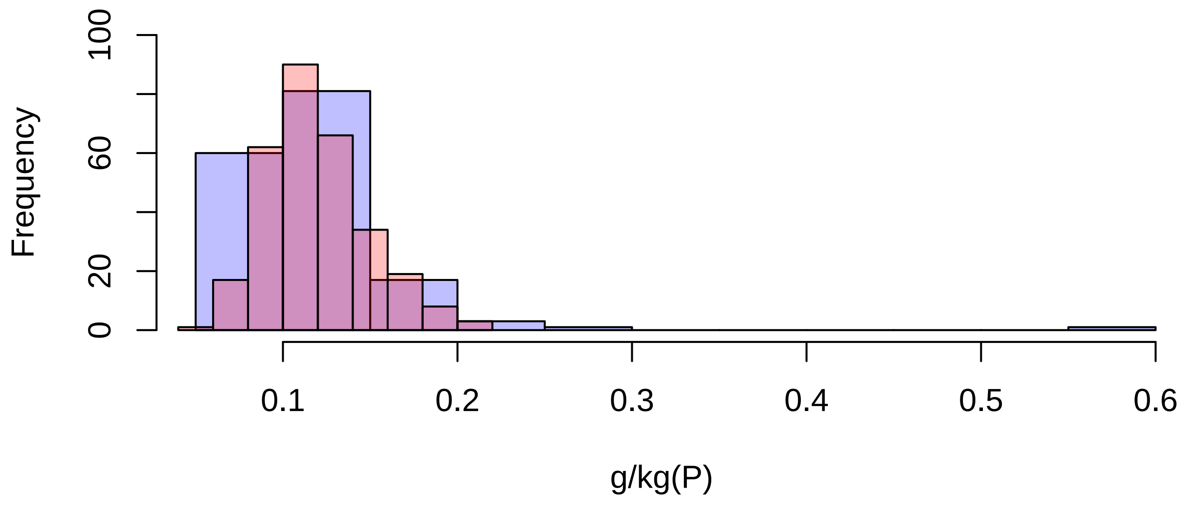

**Total Potassium**

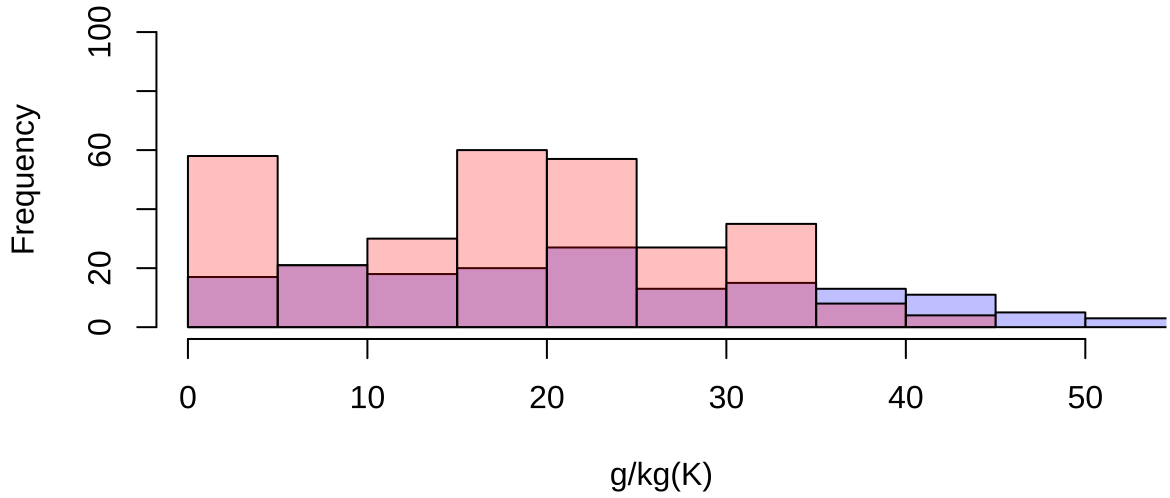


**Exchangeable Calcium**

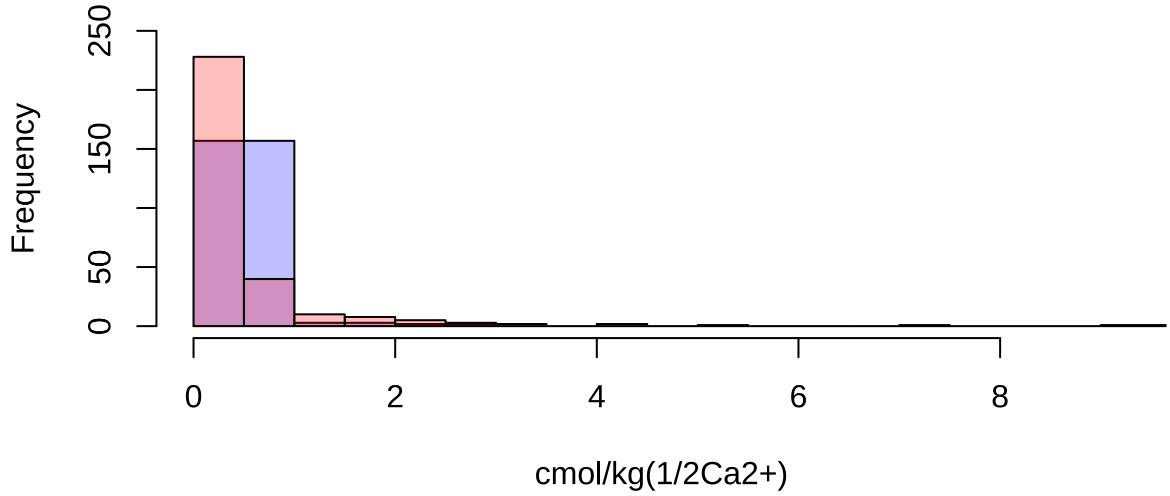

**Exchangeable Magnesium**

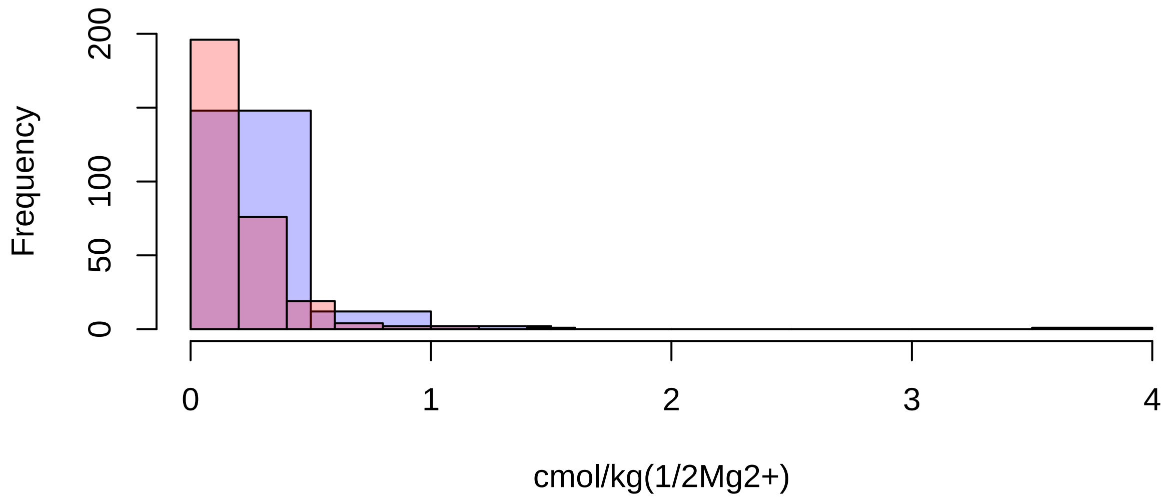


**Available Potassium**

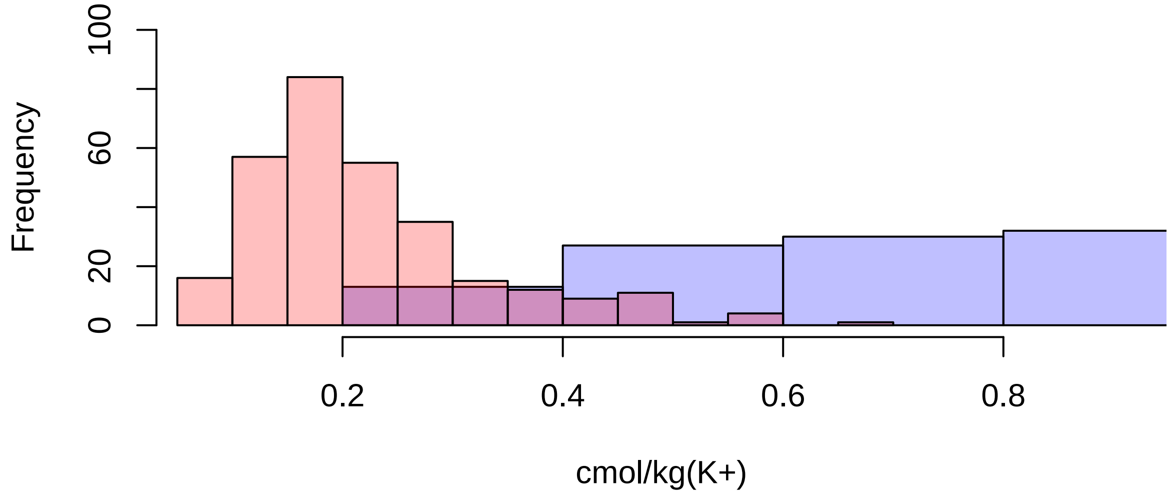

**Alkali-hydrolyzable Nitrogen**

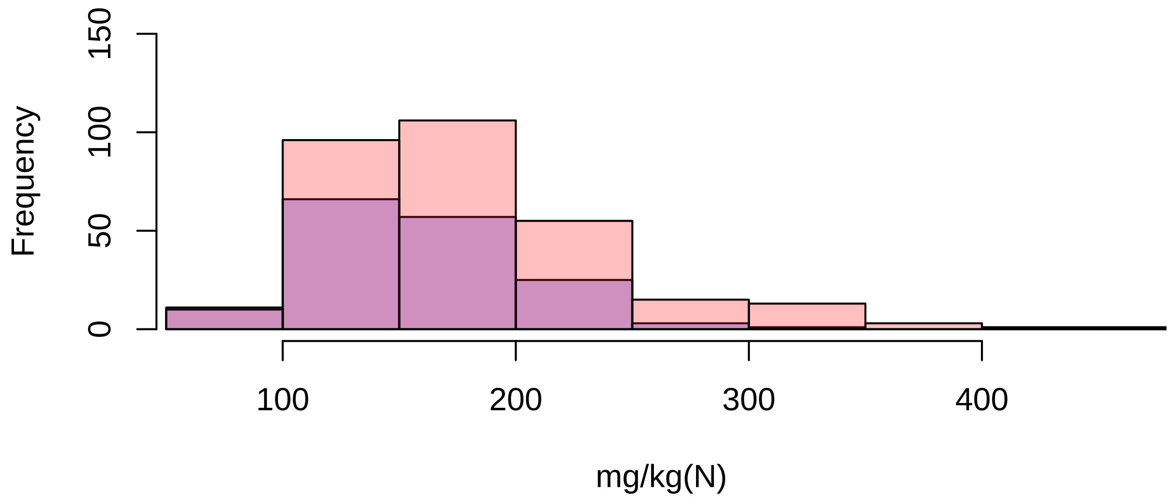


**Available Phosphorus**

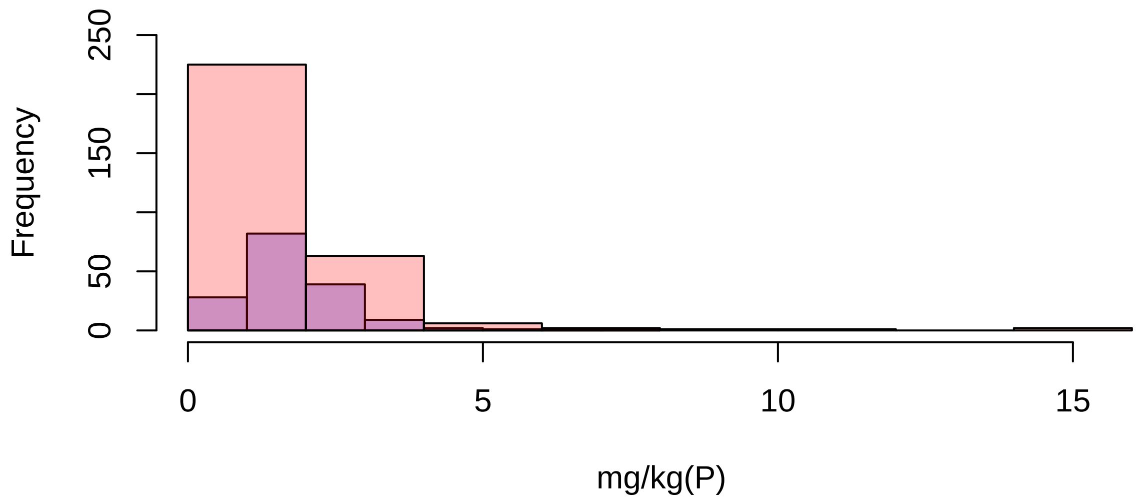
