## Supplement 2: Literate statistical document with simple univariate Analyses of Variance and least-squared mean predicted values. for "Intraspecific root and leaf trait variation with tropical forest successional status: consequences for community-weighted patterns"

### JFL_Univariate ANOVAS

*JAH*

*July 26, 2018*

#### Background

In this analysis, we use some simple univariate ANOVAS (linear models) to test for among species and foest types using the Jianfengling trait data (collected by JAH during the suimmer of 2017). The driving quesion is how plastic are the traits? within species and between the two forest types - primary vs. seconday.

We use two-way analysis of variance (assuming homoscedasticity) and test two models with respect to several traits. Principle components analyses showed that the main orthogonal leaf traits wer SLA and Leaf Area, and the orthogonal were SRL and Root Area, so we test those first.

##### Testing 2 models:

1. trait ~ species + forest.type this model tests for intraspecific diffenrences due to forest type (i.e. first vs. second half of the gradient)
2. trait ~ species|family + forest.type
   this modesl does the same as model1, but has species nested within family to kind of control for phylogenetic relatedness (becuase certain Angiosperm clades were targeted in the sampling design)

#### SLA

##### plots

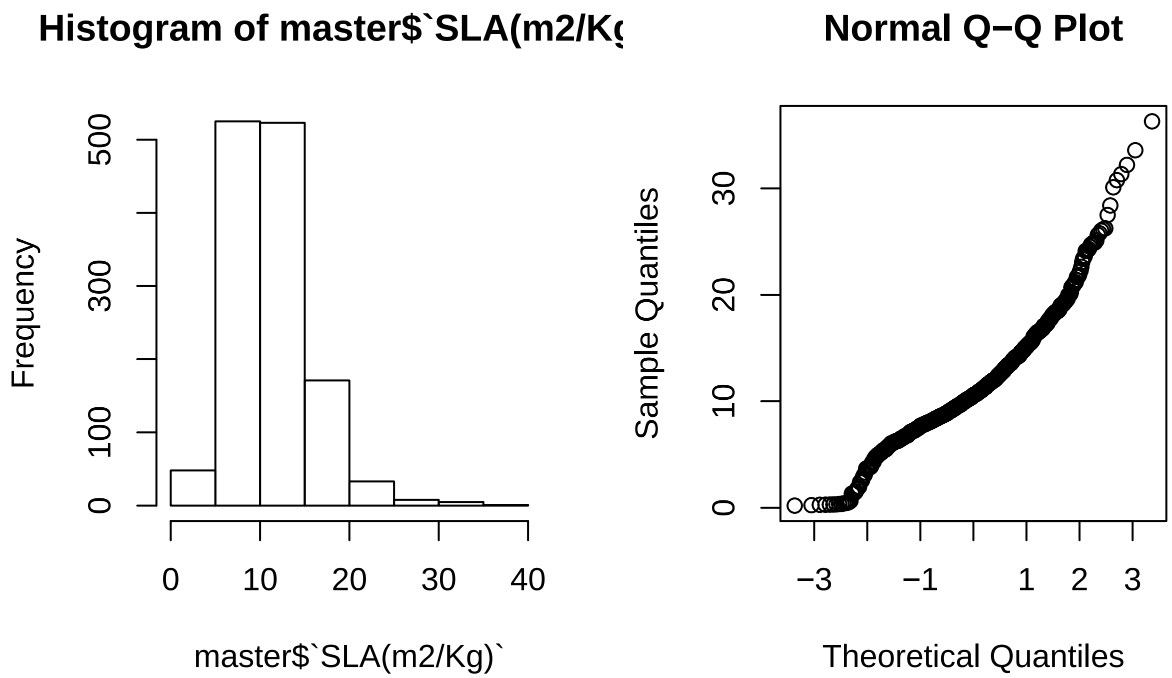
Histogram of master$`SLA(m2/Kg
Frequency
0
100
300
500
0
10
20
30
40
master$`SLA(m2/Kg)`
Normal Q−Q Plot
Sample Quantiles
0
10
20
30
−3
−1
1
2
3
Theoretical Quantiles

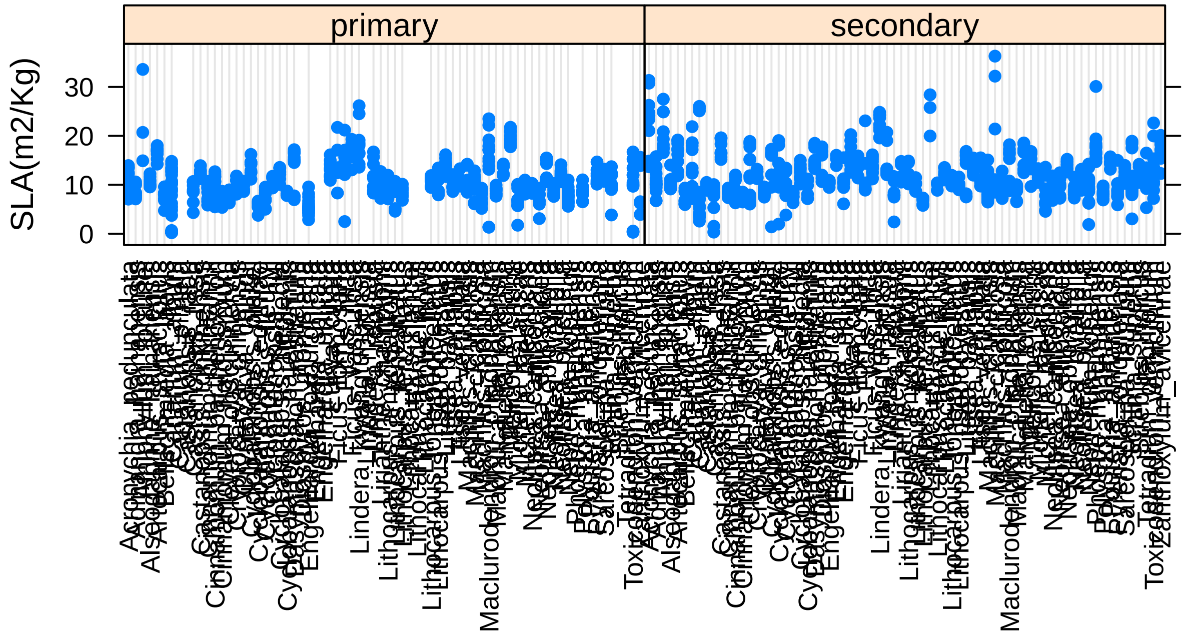
primary
secondary
SLA(m2/Kg)
0
10
20
30

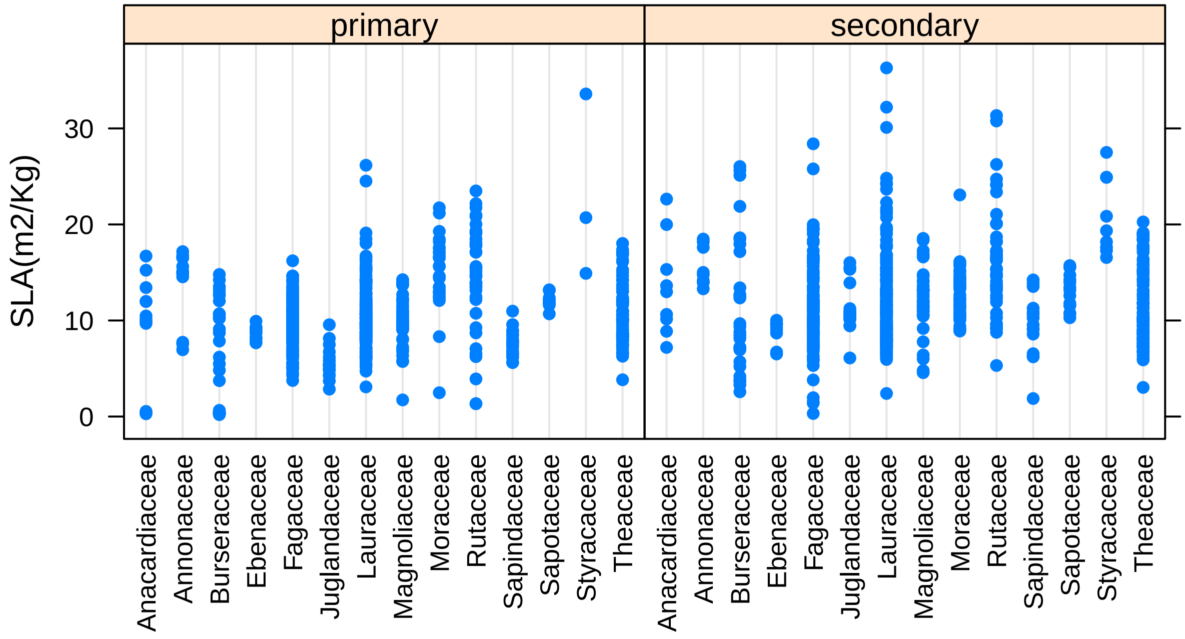


##### models

```
## Analysis of Variance Table
##
## Model 1: `SLA(m2/Kg)` ~ Gen.sp + forest.type
## Model 2: `SLA(m2/Kg)` ~ Gen.sp/Family + forest.type
## Model 3: `SLA(m2/Kg)` ~ Gen.sp/Family * forest.type
##   Res.Df   RSS Df Sum of Sq      F    Pr(>F)
## 1   1241 13002
## 2   1241 13002  0       0.0
## 3   1183 10461 58    2541.1 4.9545 < 2.2e-16 ***
## ---
## Signif. codes:  0 '***' 0.001 '**' 0.01 '*' 0.05 '.' 0.1 ' ' 1
```

```
## Analysis of Variance Table
##
## Response: SLA(m2/Kg)
##                        Df  Sum Sq Mean Sq F value    Pr(>F)
## Gen.sp                 71 12153.7  171.18 19.3577 < 2.2e-16 ***
## forest.type             1   418.9  418.92 47.3730 9.497e-12 ***
## Gen.sp:forest.type     58  2541.1   43.81  4.9545 < 2.2e-16 ***
## Residuals            1183 10461.2    8.84
## ---
## Signif. codes:  0 '***' 0.001 '**' 0.01 '*' 0.05 '.' 0.1 ' ' 1
```

**predicted marginal means**

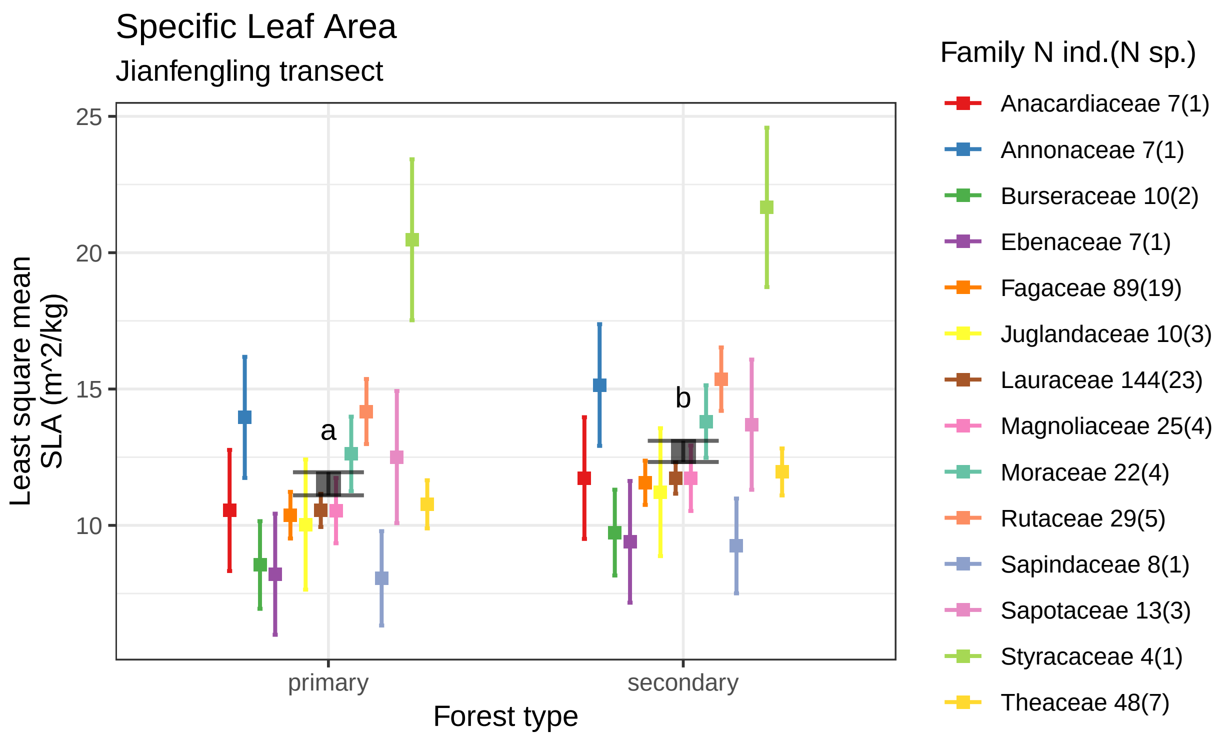


Specific leaf area estimates by forest type using data 423 tropical saplings from 72 species of 14 families.
Boxes indicate the LS mean. Error bars indicate the 95% confidence interval of the LS mean.
Means sharing a letter are not significantly different Tukey−adjusted comparisons

#### Leaf Area

##### plots

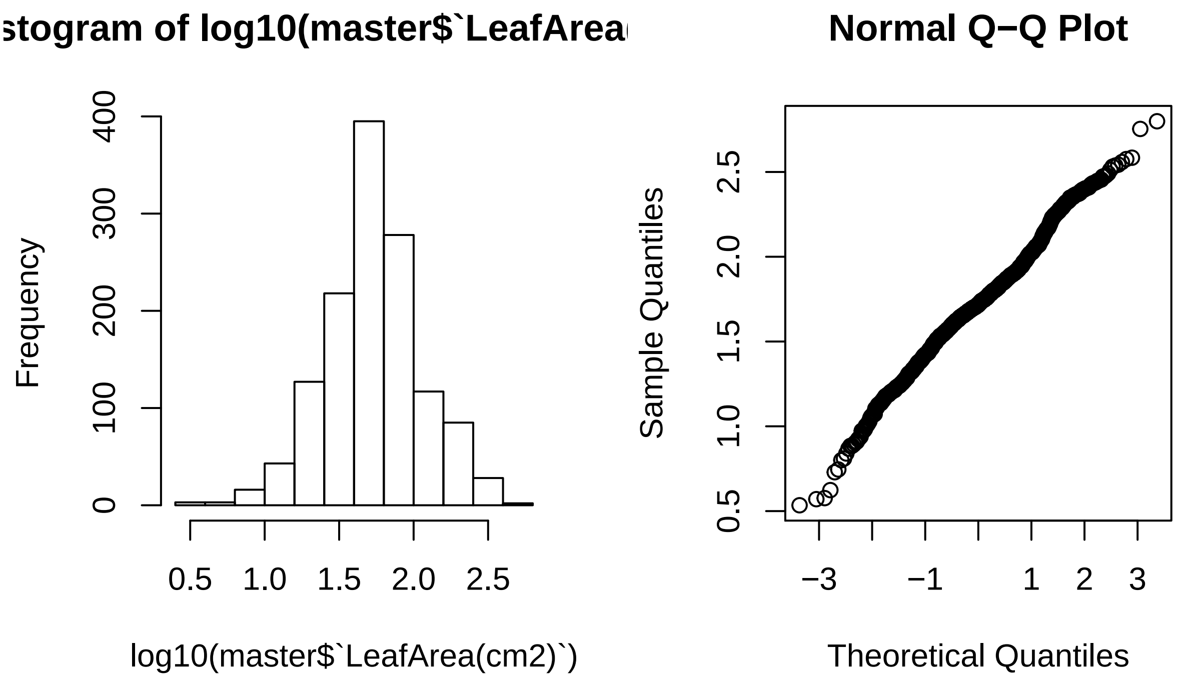


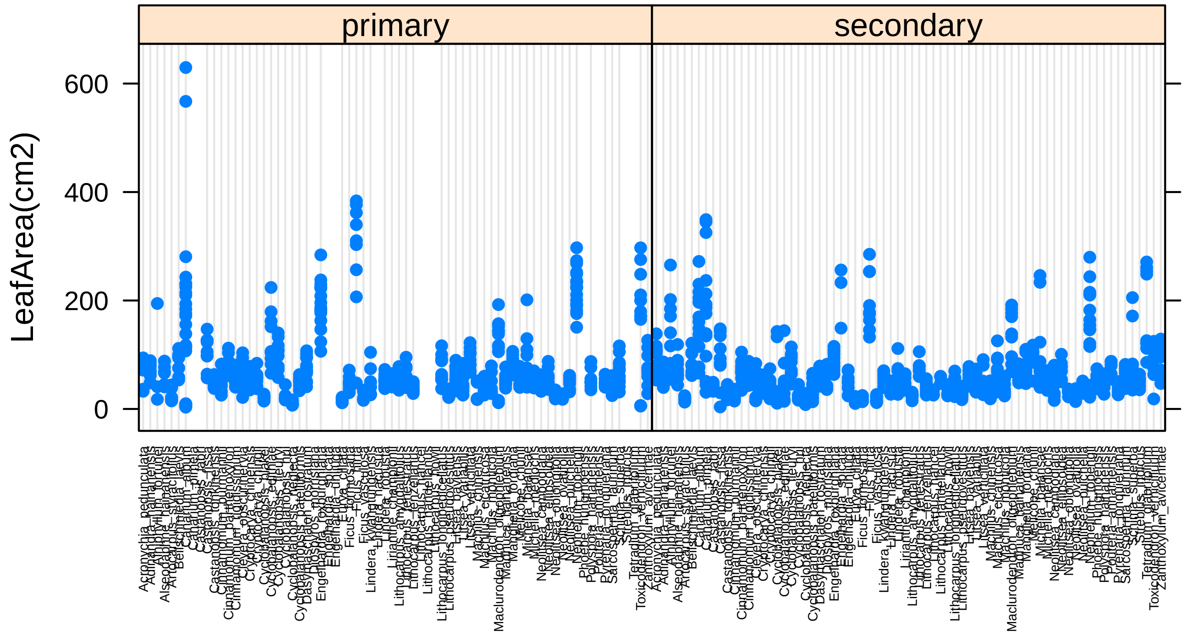

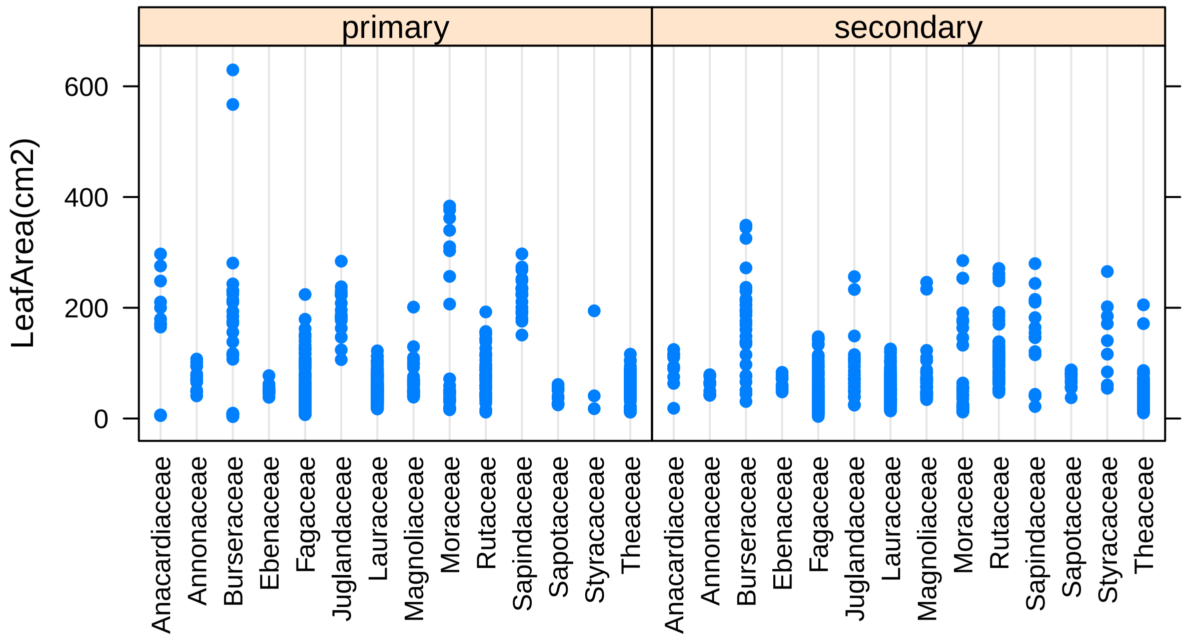


#### models

```
## Analysis of Variance Table
##
## Model 1: log10(`LeafArea(cm2)`) ~ Gen.sp + forest.type
## Model 2: log10(`LeafArea(cm2)`) ~ Gen.sp/Family + forest.type
## Model 3: log10(`LeafArea(cm2)`) ~ Gen.sp/Family * forest.type
##   Res.Df    RSS Df Sum of Sq      F    Pr(>F)
## 1   1242 60.799
## 2   1242 60.799  0    0.0000
## 3   1184 52.532 58    8.2662 3.2122 5.616e-14 ***
## ---
## Signif. codes:  0 '***' 0.001 '**' 0.01 '*' 0.05 '.' 0.1 ' ' 1
```

```
## Analysis of Variance Table
##
## Response: log10(`LeafArea(cm2)`)
##                     Df Sum Sq Mean Sq F value    Pr(>F)
## Gen.sp              71 76.199 1.07323 24.1890 < 2.2e-16 ***
## forest.type          1  0.466 0.46626 10.5089  0.001221 **
## Gen.sp:forest.type  58  8.266 0.14252  3.2122 5.616e-14 ***
## Residuals         1184 52.532 0.04437
## ---
## Signif. codes:  0 '***' 0.001 '**' 0.01 '*' 0.05 '.' 0.1 ' ' 1
```

**predicted marginal means**

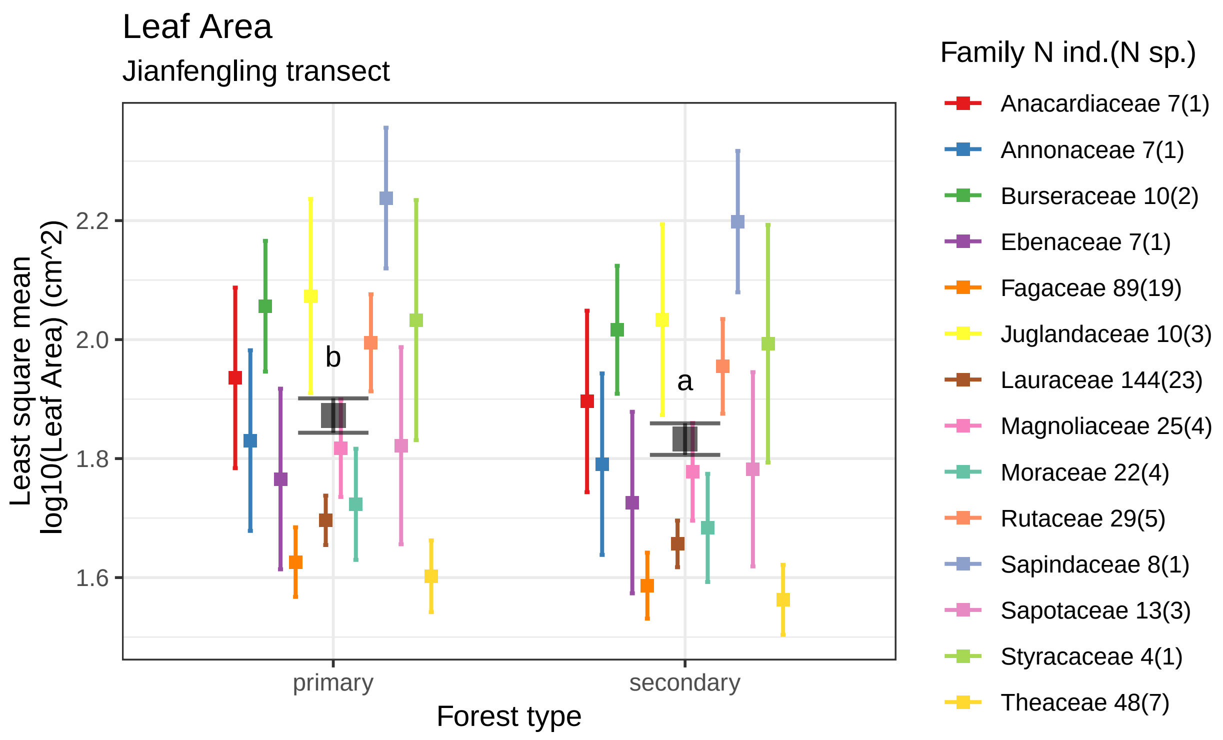


Leaf area estimates by forest type using data 423 tropical saplings from 72 species of 14 families.
Boxes indicate the LS mean. Error bars indicate the 95% confidence interval of the LS mean.
Means sharing a letter are not significantly different Tukey−adjusted comparisons

#### Leaf Thickness

##### plots

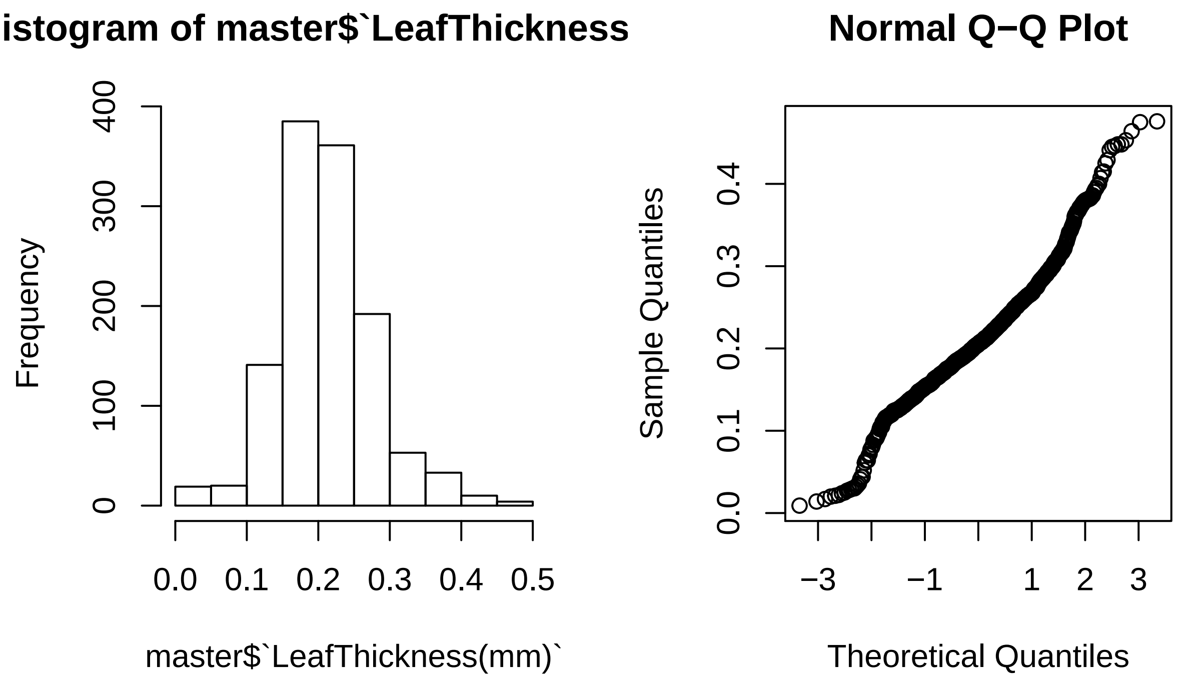


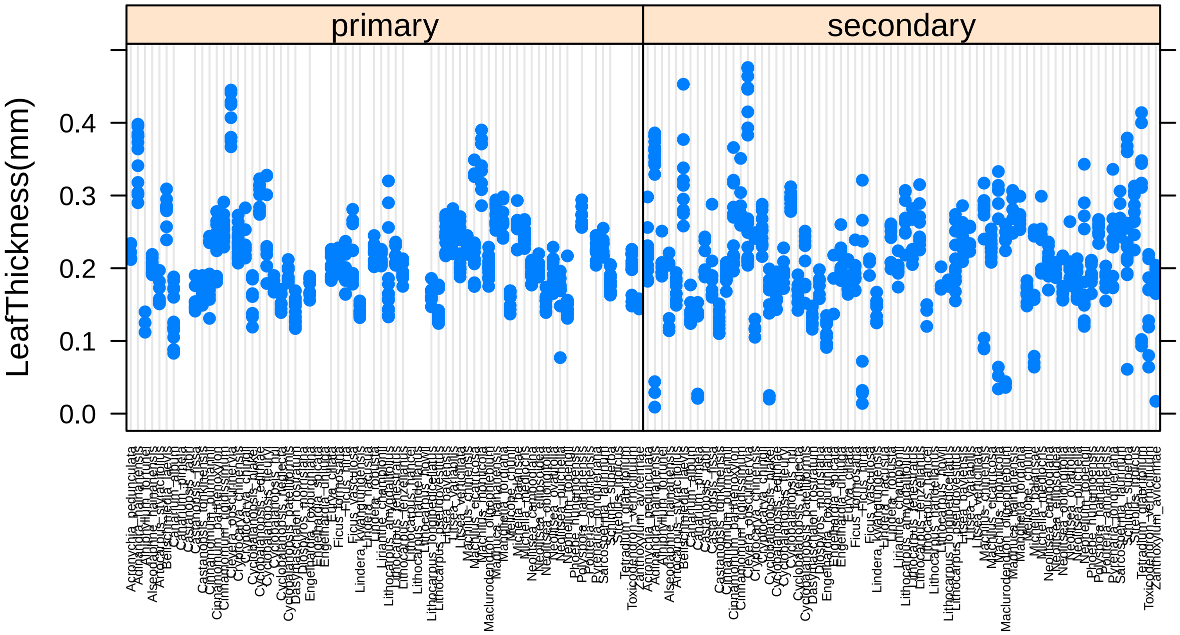

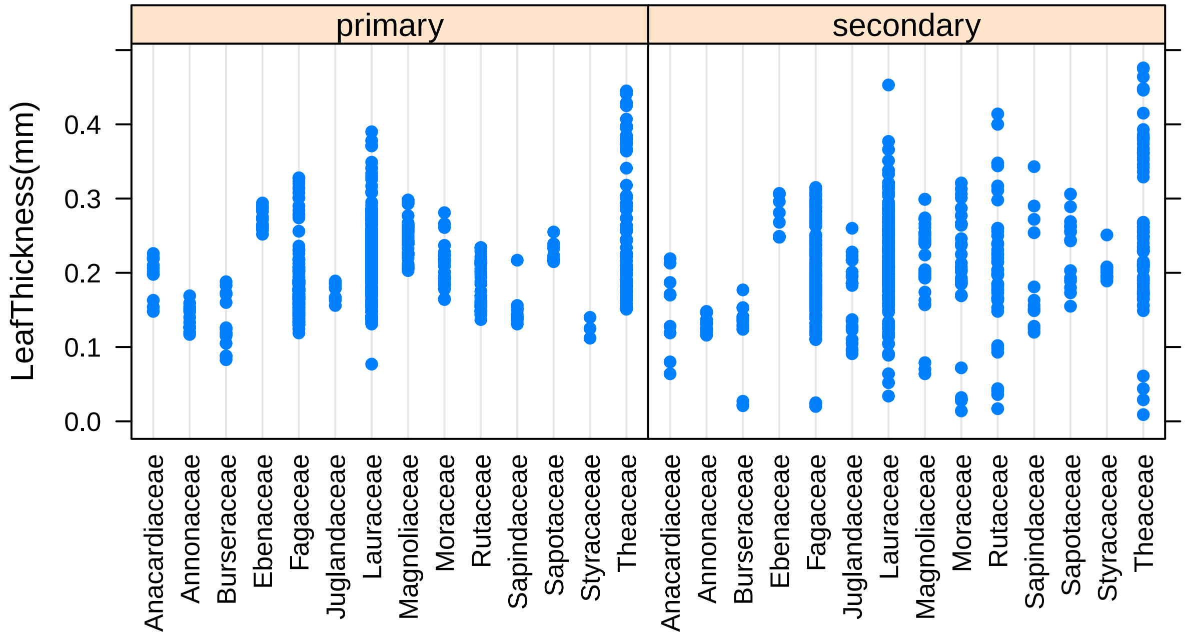


##### models

```
## Analysis of Variance Table
##
## Model 1: `LeafThickness(mm)` ~ Gen.sp + forest.type
## Model 2: `LeafThickness(mm)` ~ Gen.sp/Family + forest.type
## Model 3: `LeafThickness(mm)` ~ Gen.sp/Family * forest.type
##   Res.Df    RSS Df Sum of Sq      F    Pr(>F)
## 1   1147 2.6251
## 2   1147 2.6251  0   0.00000
## 3   1090 2.0196 57   0.60546 5.7328 < 2.2e-16 ***
## ---
## Signif. codes:  0 '***' 0.001 '**' 0.01 '*' 0.05 '.' 0.1 ' ' 1
```

```
## Analysis of Variance Table
##
## Response: LeafThickness(mm)
##                       Df  Sum Sq  Mean Sq F value Pr(>F)
## Gen.sp                69 2.88103 0.041754 22.5351 <2e-16 ***
## forest.type            1 0.00312 0.003123  1.6855 0.1945
## Gen.sp:forest.type    57 0.60546 0.010622  5.7328 <2e-16 ***
## Residuals           1090 2.01961 0.001853
## ---
## Signif. codes:  0 '***' 0.001 '**' 0.01 '*' 0.05 '.' 0.1 ' ' 1
```

**predicted marginal means**

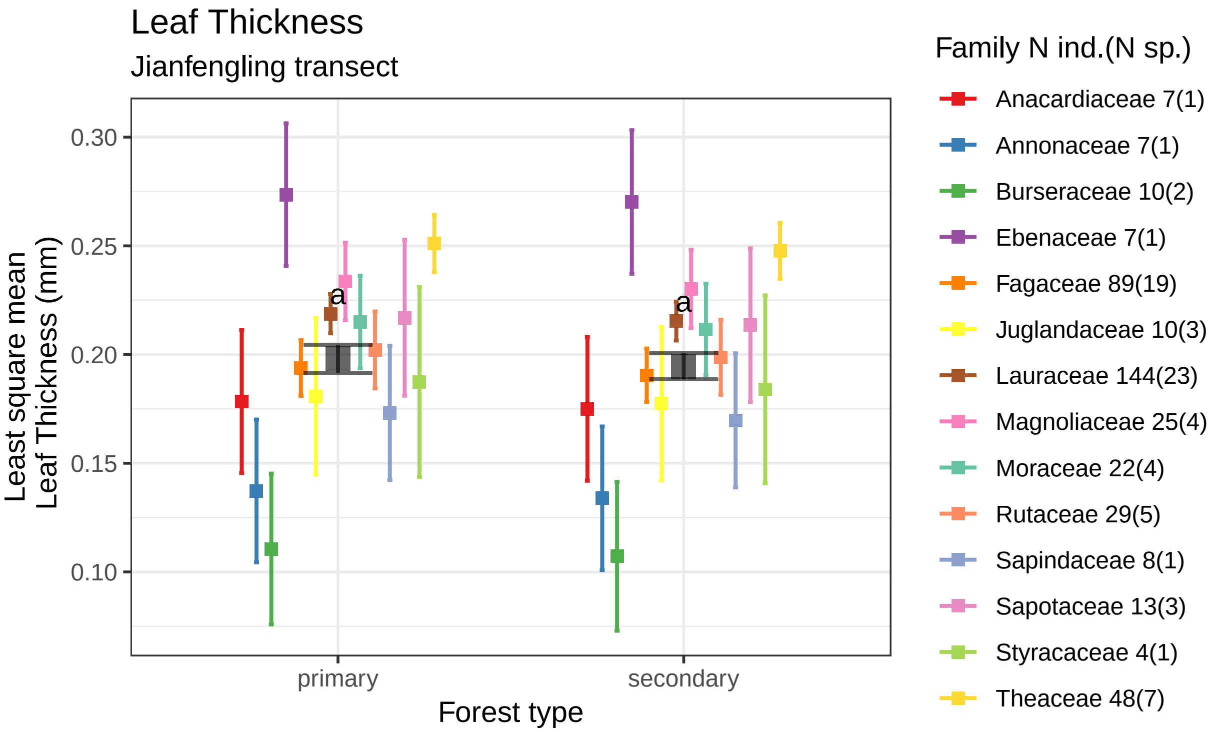


Leaf Thickness estimates by forest type using data 423 tropical saplings from 72 species of 14 families.
Boxes indicate the LS mean. Error bars indicate the 95% confidence interval of the LS mean.
Means sharing a letter are not significantly different Tukey−adjusted comparisons

#### Root Diameter

##### plots

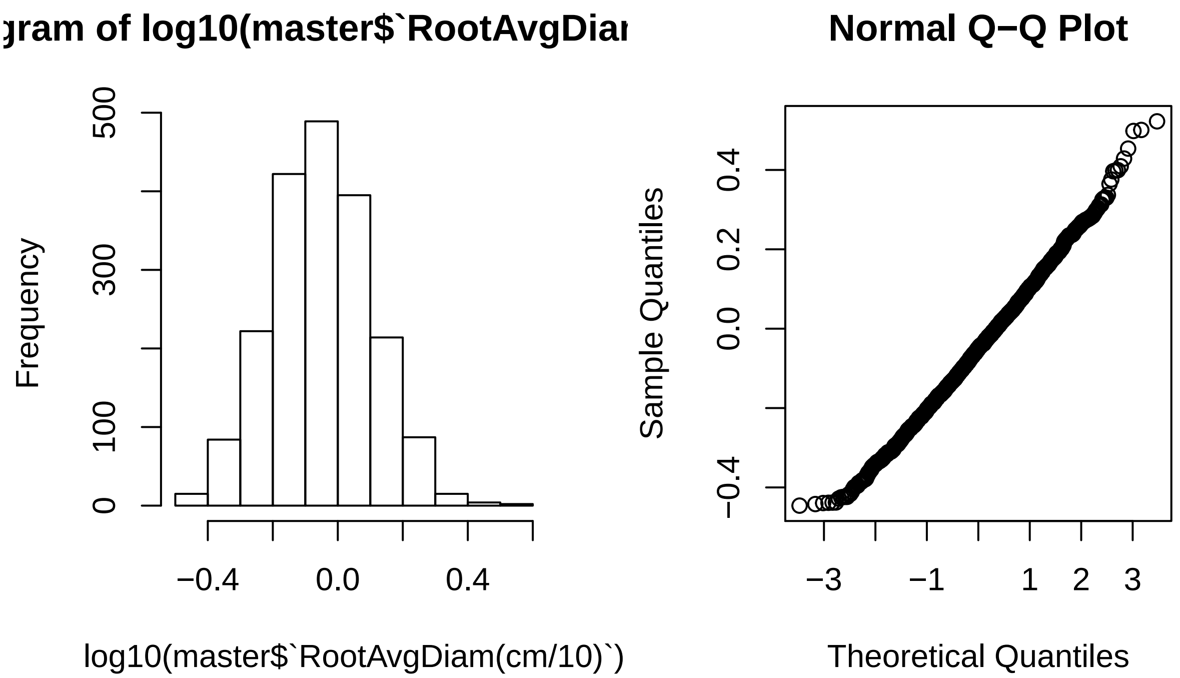


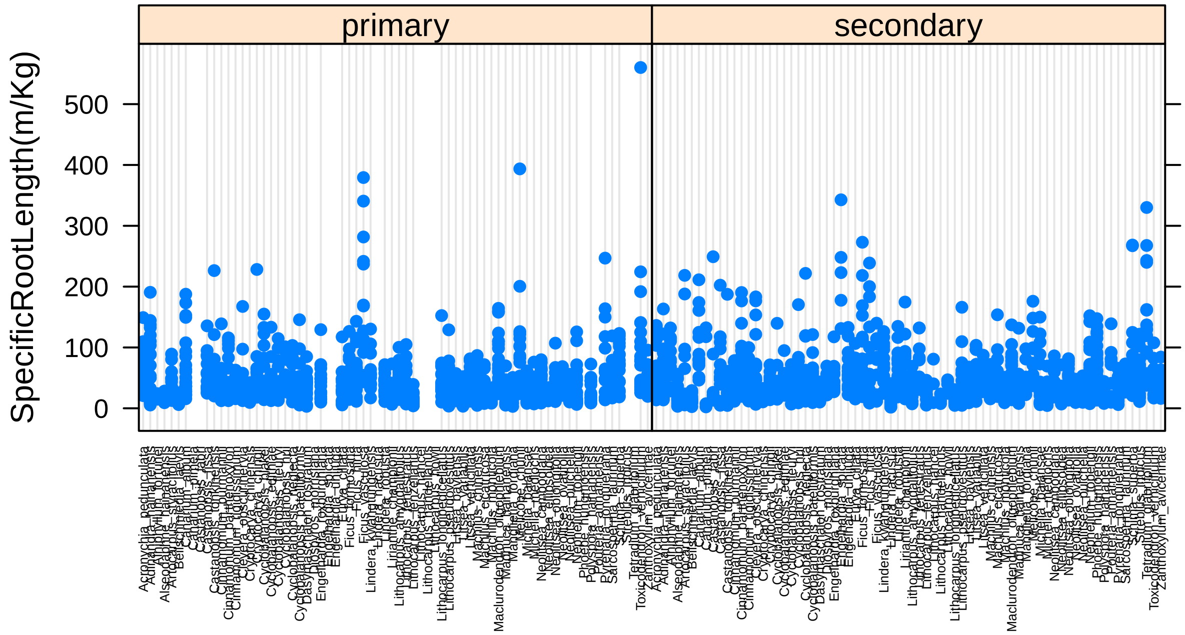

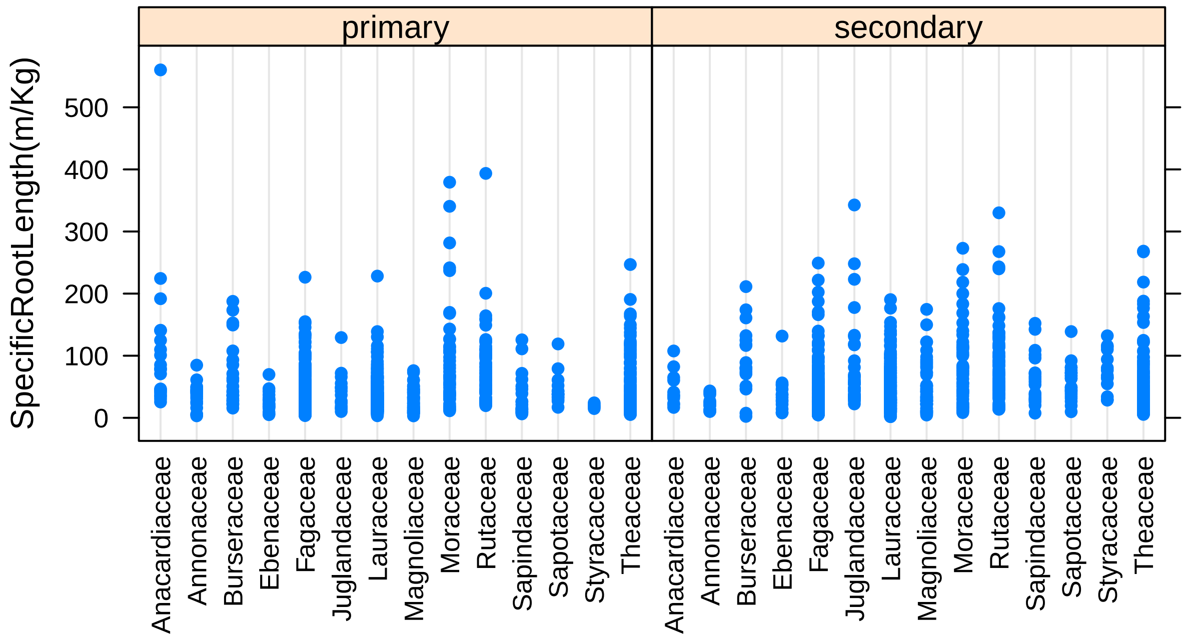


##### models

```
## Analysis of Variance Table
##
## Model 1: log10(`RootAvgDiam(cm/10)`) ~ Gen.sp + forest.type
## Model 2: log10(`RootAvgDiam(cm/10)`) ~ Gen.sp/Family + forest.type
## Model 3: log10(`RootAvgDiam(cm/10)`) ~ Gen.sp/Family * forest.type
##   Res.Df    RSS Df Sum of Sq      F    Pr(>F)
## 1   1876 26.190
## 2   1876 26.190  0    0.0000
## 3   1818 23.518 58    2.6719 3.5611 < 2.2e-16 ***
## ---
## Signif. codes:  0 '***' 0.001 '**' 0.01 '*' 0.05 '.' 0.1 ' ' 1
```

```
## Analysis of Variance Table
##
## Response: log10(`RootAvgDiam(cm/10)`)
##                       Df  Sum Sq  Mean Sq F value    Pr(>F)
## Gen.sp                71 19.4227 0.273559 21.1470 < 2.2e-16 ***
## forest.type            1  0.2015 0.201483 15.5753  8.23e-05 ***
## Gen.sp:forest.type    58  2.6719 0.046067  3.5611 < 2.2e-16 ***
## Residuals           1818 23.5177 0.012936
## ---
## Signif. codes:  0 '***' 0.001 '**' 0.01 '*' 0.05 '.' 0.1 ' ' 1
```

**predicted marginal means**

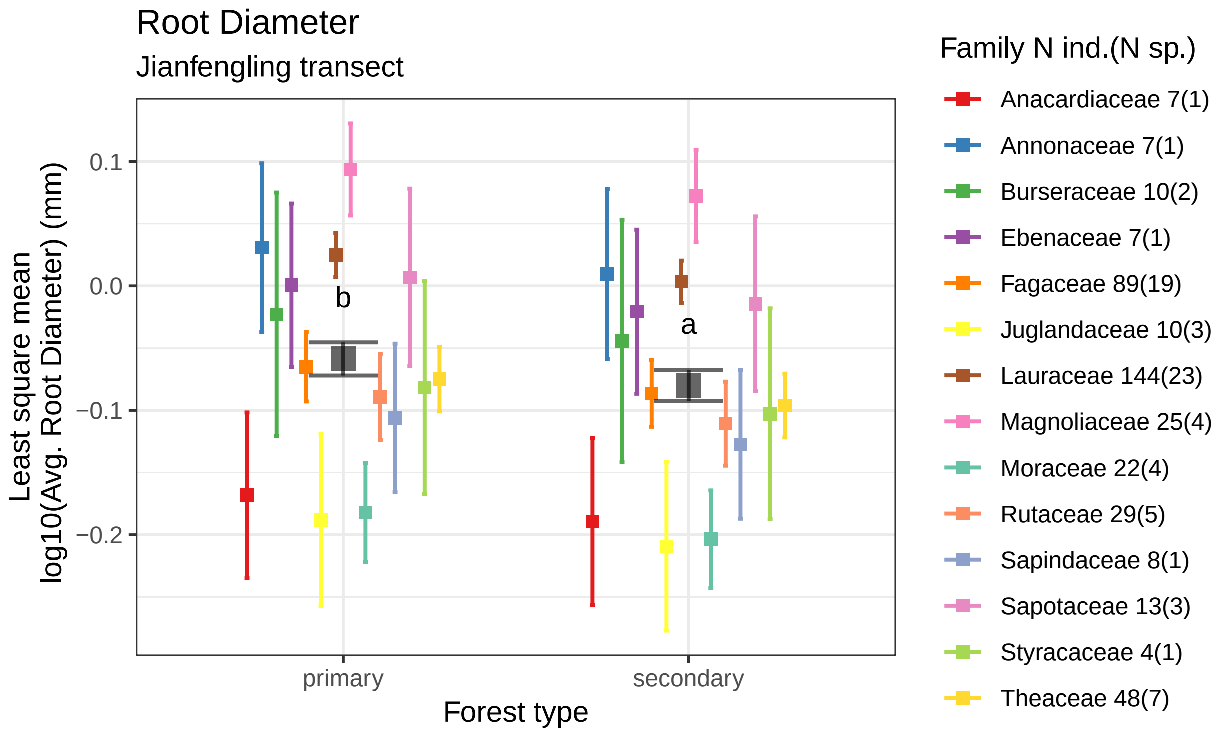


Mean ERS root diameter estimates by forest type using data 423 tropical saplings from 72 species of 14
Boxes indicate the LS mean. Error bars indicate the 95% confidence interval of the LS mean.
Means sharing a letter are not significantly different Tukey−adjusted comparisons

```
## pdf
##   2
```

### SRL

#### plots

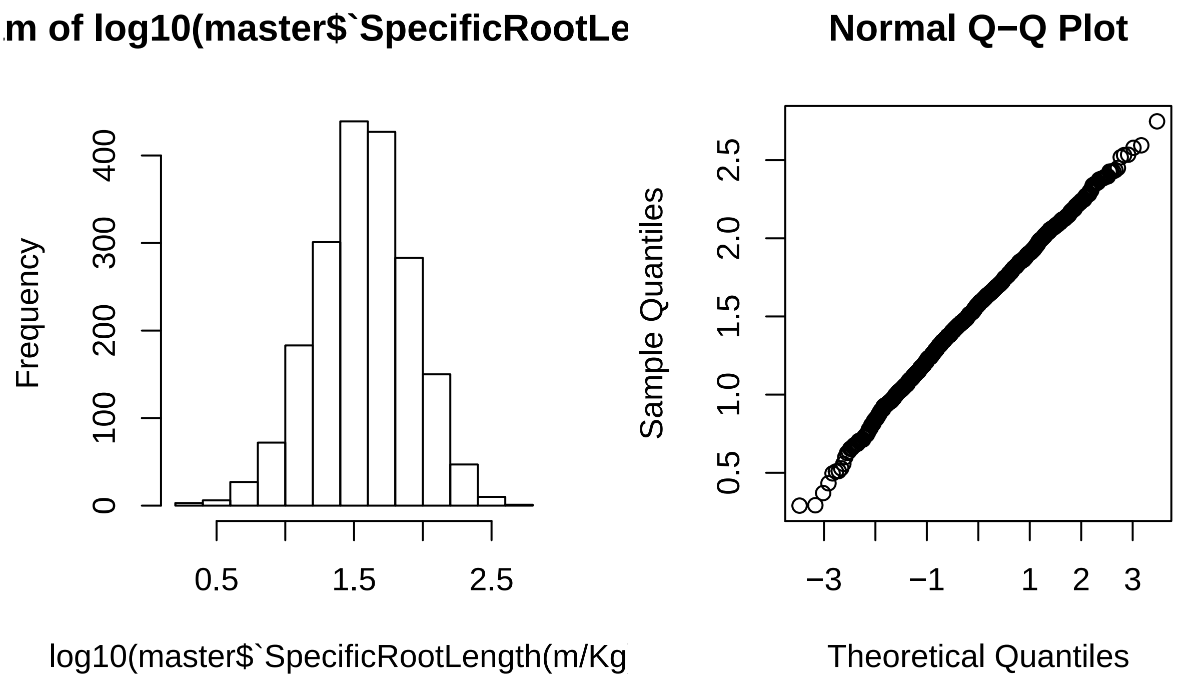

m of log10(master$`SpecificRootLe
Frequency
log10(master$`SpecificRootLength(m/Kg
Normal Q−Q Plot
Sample Quantiles
Theoretical Quantiles


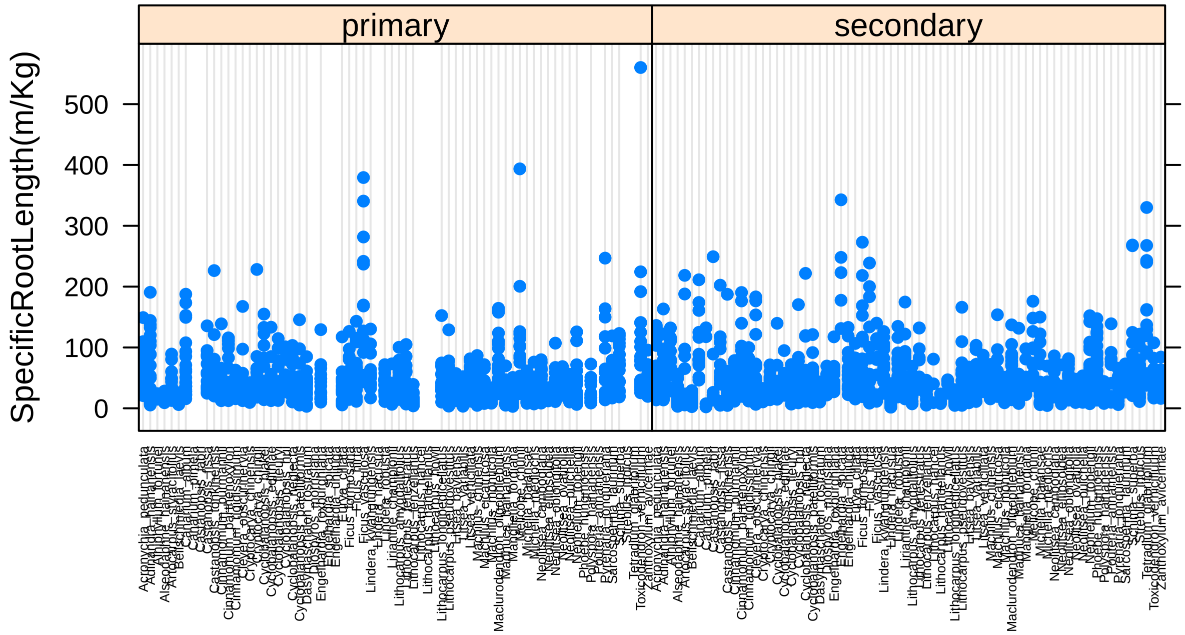

primary
secondary
SpecificRootLength(m/Kg)

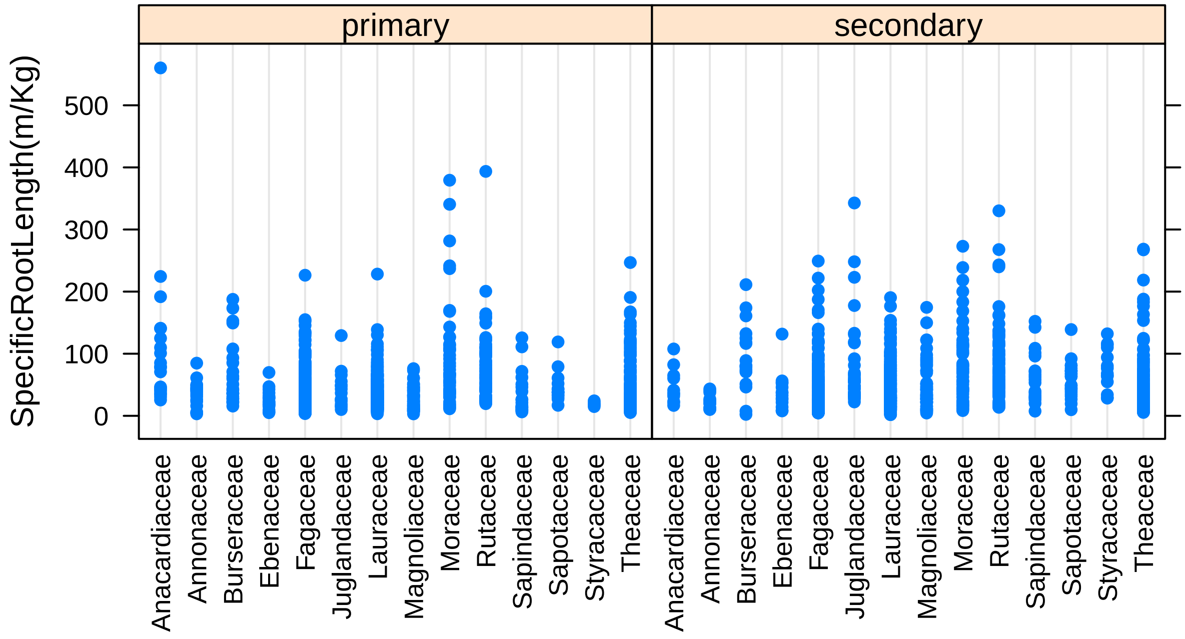


##### models

```
## Analysis of Variance Table
##
## Model 1: log10(`SpecificRootLength(m/Kg)`) ~ Gen.sp + forest.type
## Model 2: log10(`SpecificRootLength(m/Kg)`) ~ Gen.sp/Family + forest.type
## Model 3: log10(`SpecificRootLength(m/Kg)`) ~ Gen.sp/Family * forest.type
##   Res.Df    RSS Df Sum of Sq      F    Pr(>F)
## 1   1876 174.35
## 2   1876 174.35  0     0.000
## 3   1818 156.46 58    17.891 3.5844 < 2.2e-16 ***
## ---
## Signif. codes:  0 '***' 0.001 '**' 0.01 '*' 0.05 '.' 0.1 ' ' 1
```

```
## Analysis of Variance Table
##
## Response: log10(`SpecificRootLength(m/Kg)`)
##                       Df  Sum Sq Mean Sq F value    Pr(>F)
## Gen.sp                71  62.660 0.88254 10.2551 < 2.2e-16 ***
## forest.type            1   0.611 0.61089  7.0985  0.007783 **
## Gen.sp:forest.type    58  17.891 0.30847  3.5844 < 2.2e-16 ***
## Residuals           1818 156.455 0.08606
## ---
## Signif. codes:  0 '***' 0.001 '**' 0.01 '*' 0.05 '.' 0.1 ' ' 1
```

**predicted marginal means**

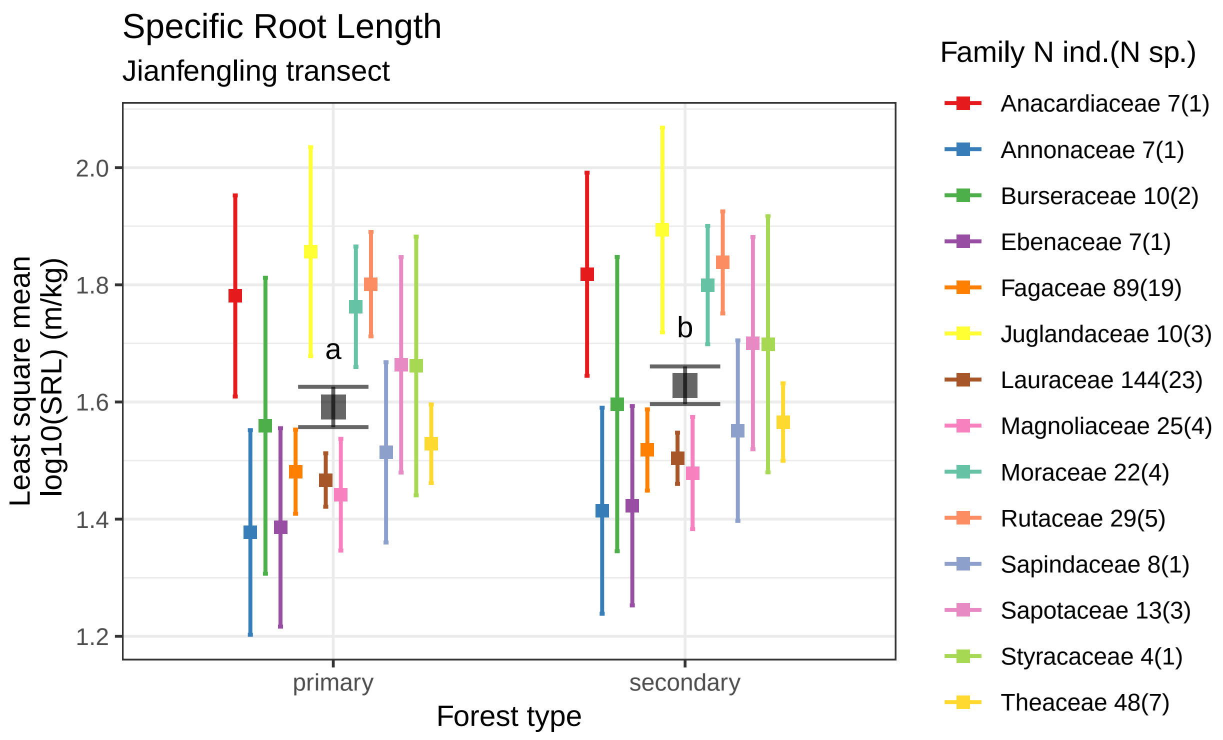


Specific root length estimates by forest type using data 423 tropical saplings from 72 species of 14 families
Boxes indicate the LS mean. Error bars indicate the 95% confidence interval of the LS mean.
Means sharing a letter are not significantly different Tukey−adjusted comparisons

#### Root Area

##### plots

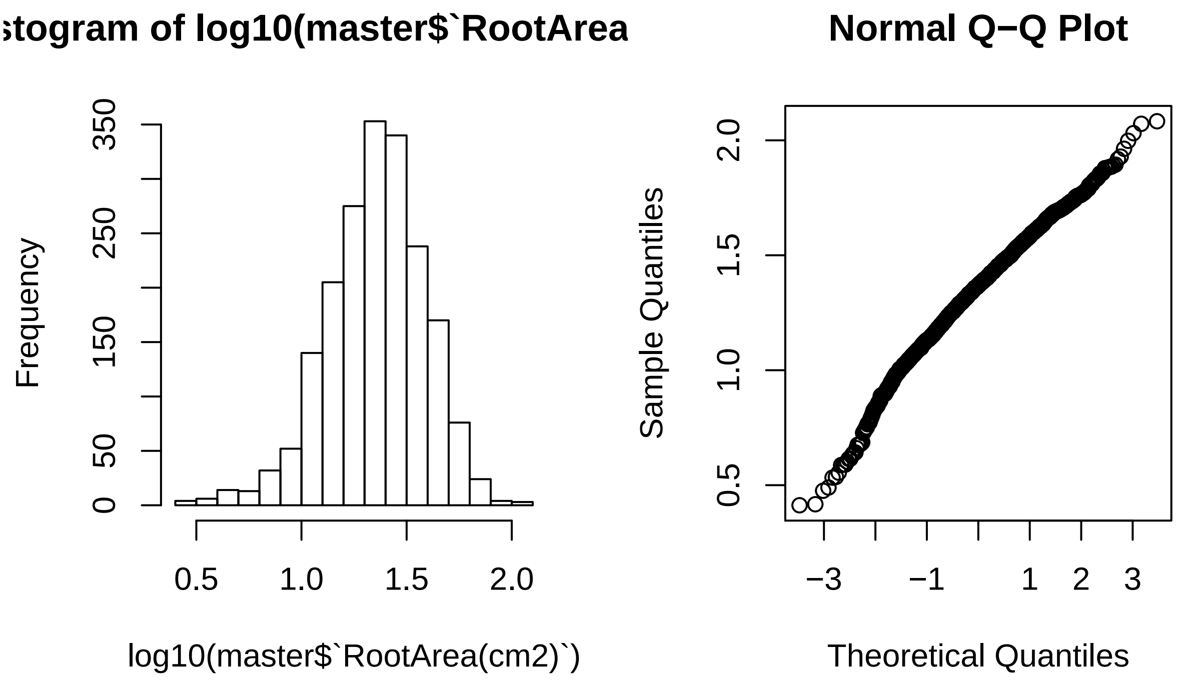


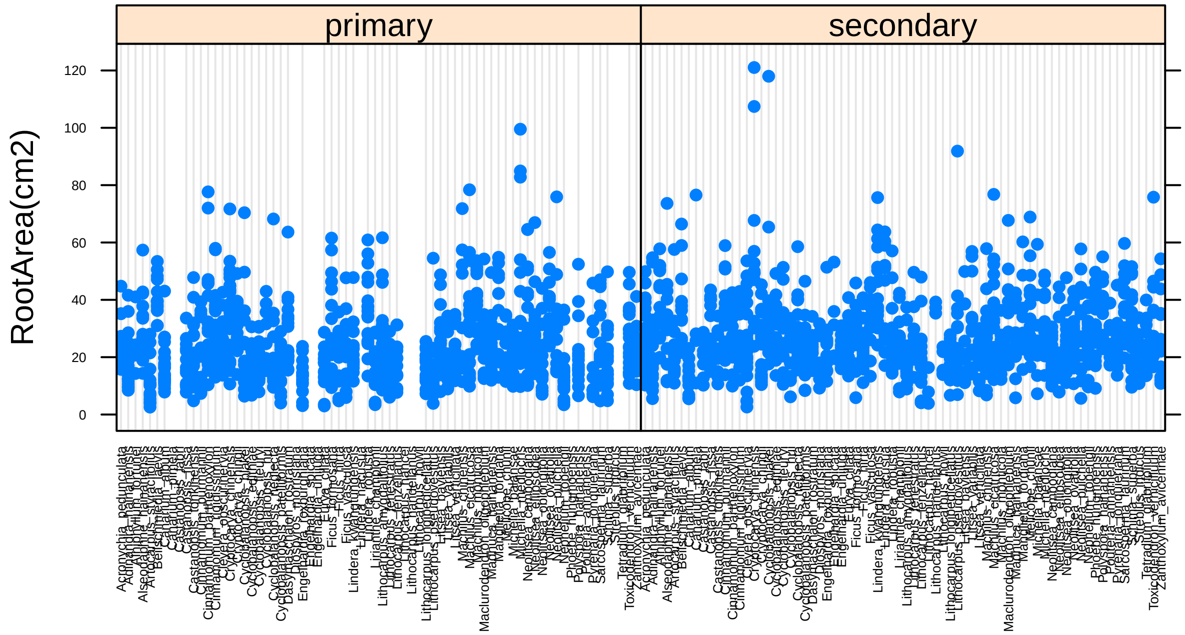

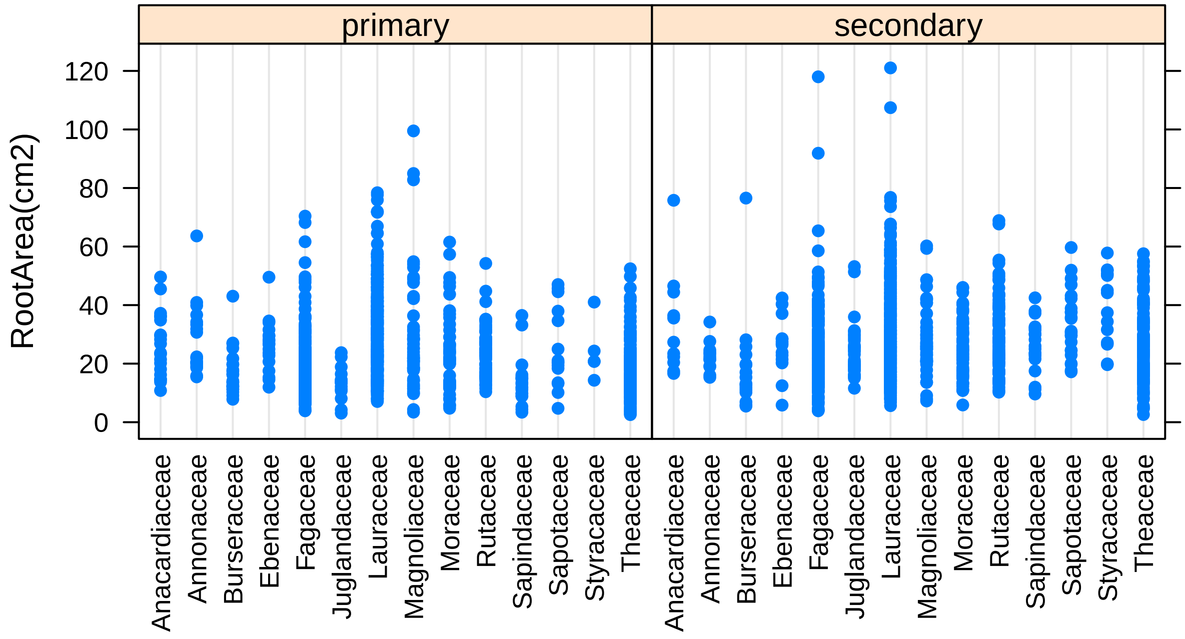


##### models

```
## Analysis of Variance Table
##
## Model 1: log10(`RootArea(cm2)`) ~ Gen.sp + forest.type
## Model 2: log10(`RootArea(cm2)`) ~ Gen.sp/Family + forest.type
## Model 3: log10(`RootArea(cm2)`) ~ Gen.sp/Family * forest.type
##   Res.Df    RSS Df Sum of Sq      F    Pr(>F)
## 1   1876 84.773
## 2   1876 84.773  0     0.000
## 3   1818 74.461 58    10.312 4.3407 < 2.2e-16 ***
## ---
## Signif. codes:  0 '***' 0.001 '**' 0.01 '*' 0.05 '.' 0.1 ' ' 1
```

```
## Analysis of Variance Table
##
## Response: log10(`RootArea(cm2)`)
##                     Df Sum Sq Mean Sq F value    Pr(>F)
## Gen.sp              71 19.409 0.27336  6.6742 < 2.2e-16 ***
## forest.type          1  2.215 2.21517 54.0844 2.891e-13 ***
## Gen.sp:forest.type  58 10.311 0.17778  4.3407 < 2.2e-16 ***
## Residuals         1818 74.461 0.04096
## ---
## Signif. codes:  0 '***' 0.001 '**' 0.01 '*' 0.05 '.' 0.1 ' ' 1
```

**predicted marginal means**

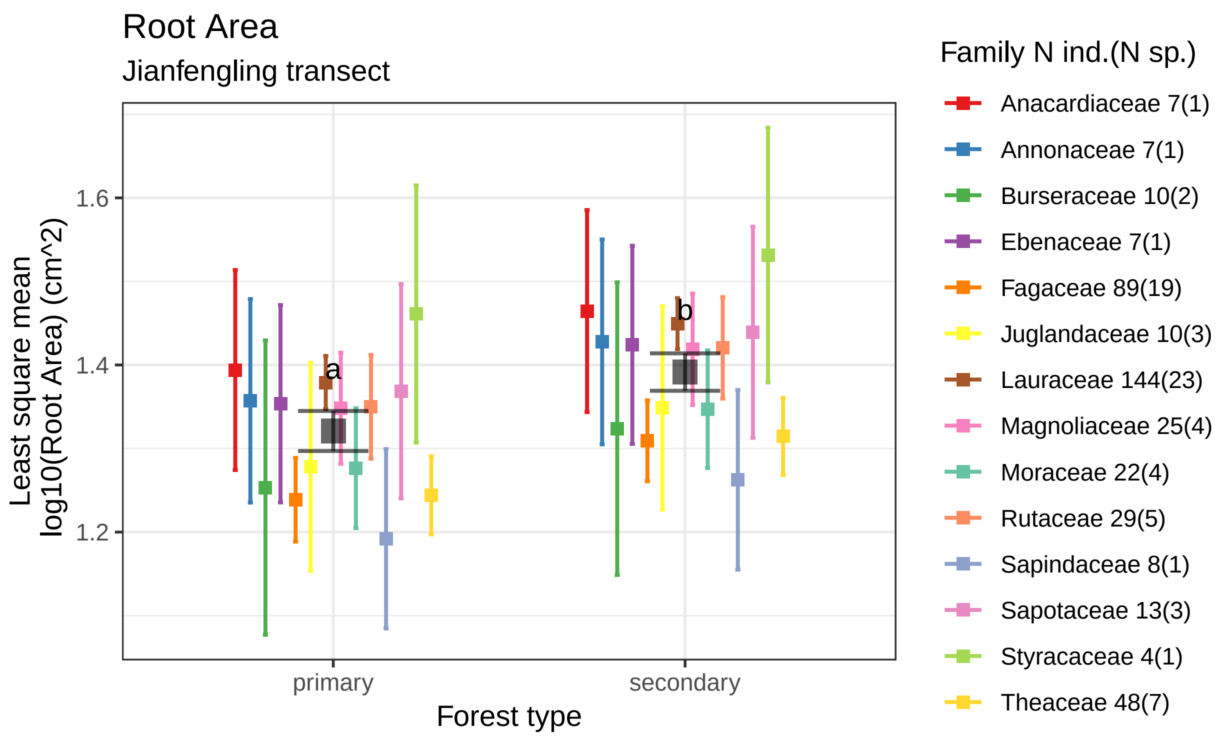


Root area estimates by forest type using data 423 tropical saplings from 72 species of 14 families.
Boxes indicate the LS mean. Error bars indicate the 95% confidence interval of the LS mean.
Means sharing a letter are not significantly different Tukey−adjusted comparisons

#### Root Branchiness

##### plots

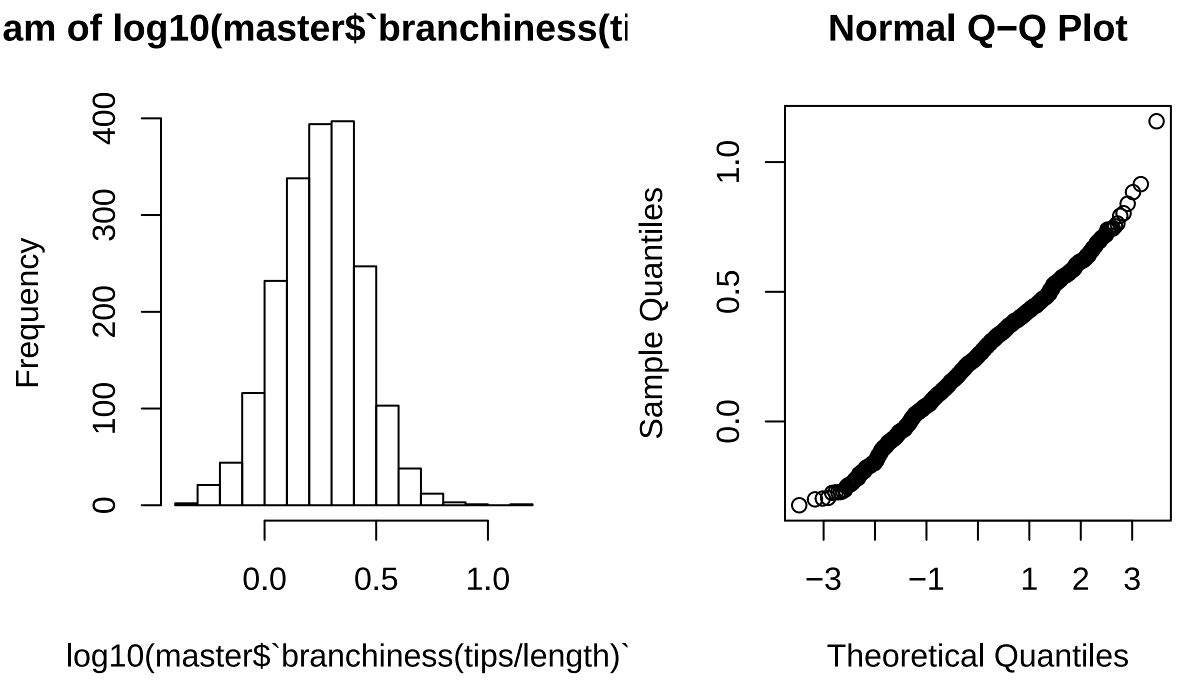


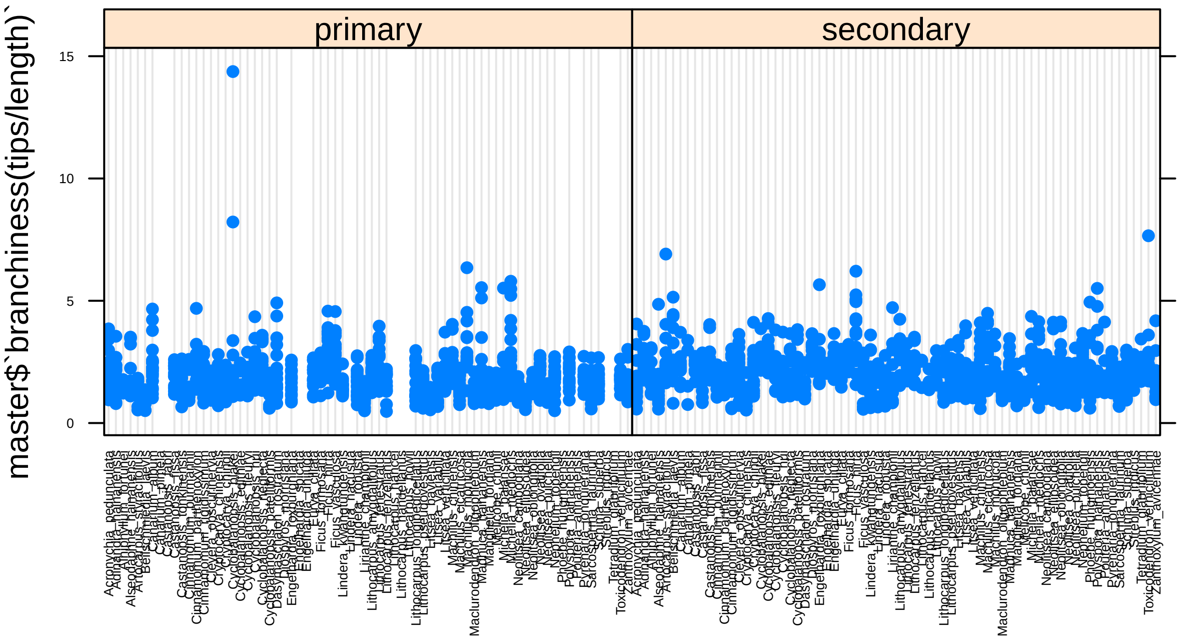

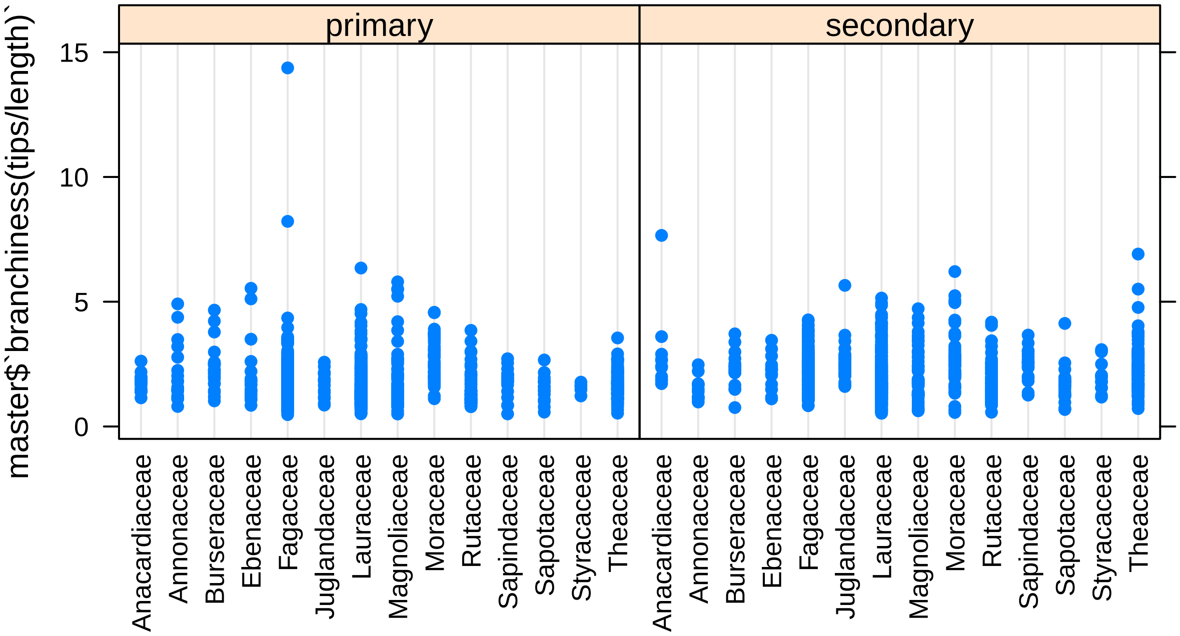


##### models

```
## Analysis of Variance Table
##
## Model 1: log10(`branchiness(tips/length)`) ~ Gen.sp + forest.type
## Model 2: log10(`branchiness(tips/length)`) ~ Gen.sp/Family + forest.type
## Model 3: log10(`branchiness(tips/length)`) ~ Gen.sp/Family * forest.type
##   Res.Df    RSS Df Sum of Sq    F    Pr(>F)
## 1   1876 51.079
## 2   1876 51.079  0    0.0000
## 3   1818 43.196 58    7.8827 5.72 < 2.2e-16 ***
## ---
## Signif. codes:  0 '***' 0.001 '**' 0.01 '*' 0.05 '.' 0.1 ' ' 1
```

```
## Analysis of Variance Table
##
## Response: log10(`branchiness(tips/length)`)
##                     Df Sum Sq Mean Sq F value    Pr(>F)
## Gen.sp              71 16.468 0.23195  9.7619 < 2.2e-16 ***
## forest.type          1  1.940 1.94032 81.6619 < 2.2e-16 ***
## Gen.sp:forest.type  58  7.883 0.13591  5.7200 < 2.2e-16 ***
## Residuals         1818 43.196 0.02376
## ---
## Signif. codes:  0 '***' 0.001 '**' 0.01 '*' 0.05 '.' 0.1 ' ' 1
```

**predicted marginal means**

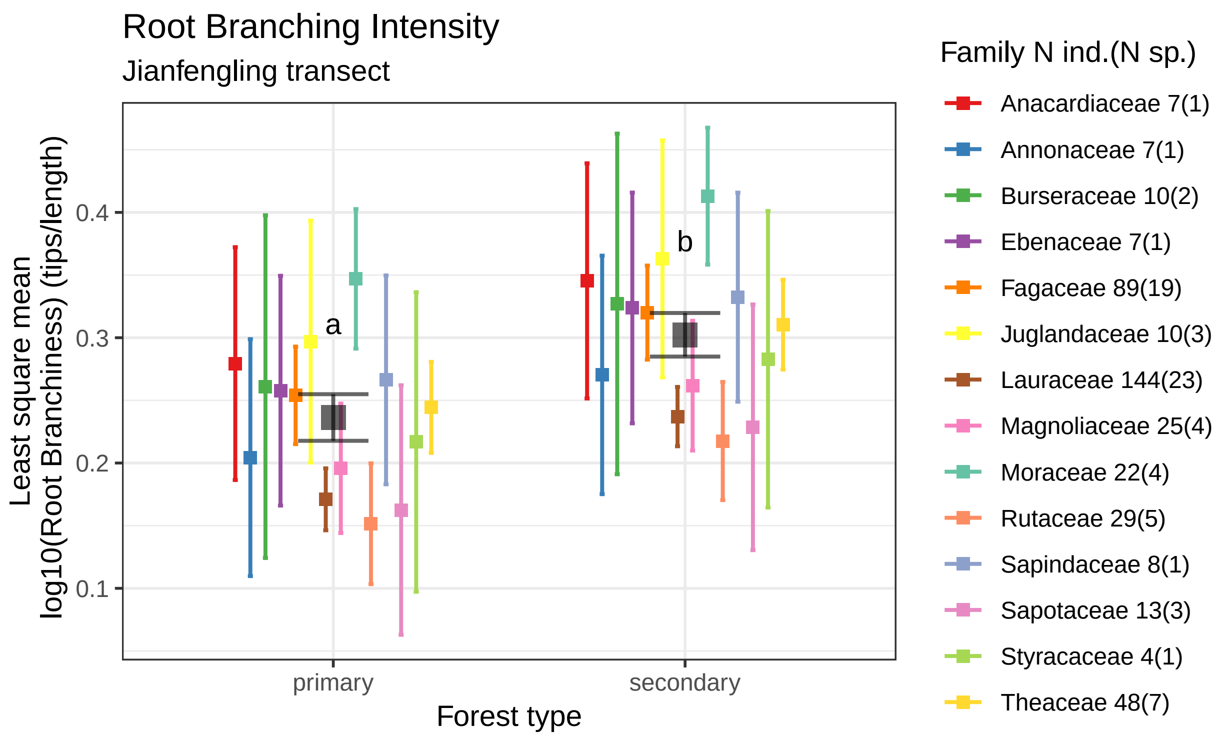


Number of root tips per length by forest type using data 423 tropical saplings from 72 species of 14 familie
Boxes indicate the LS mean. Error bars indicate the 95% confidence interval of the LS mean.
Means sharing a letter are not significantly different Tukey−adjusted comparisons

#### Nroot Tips

##### plots

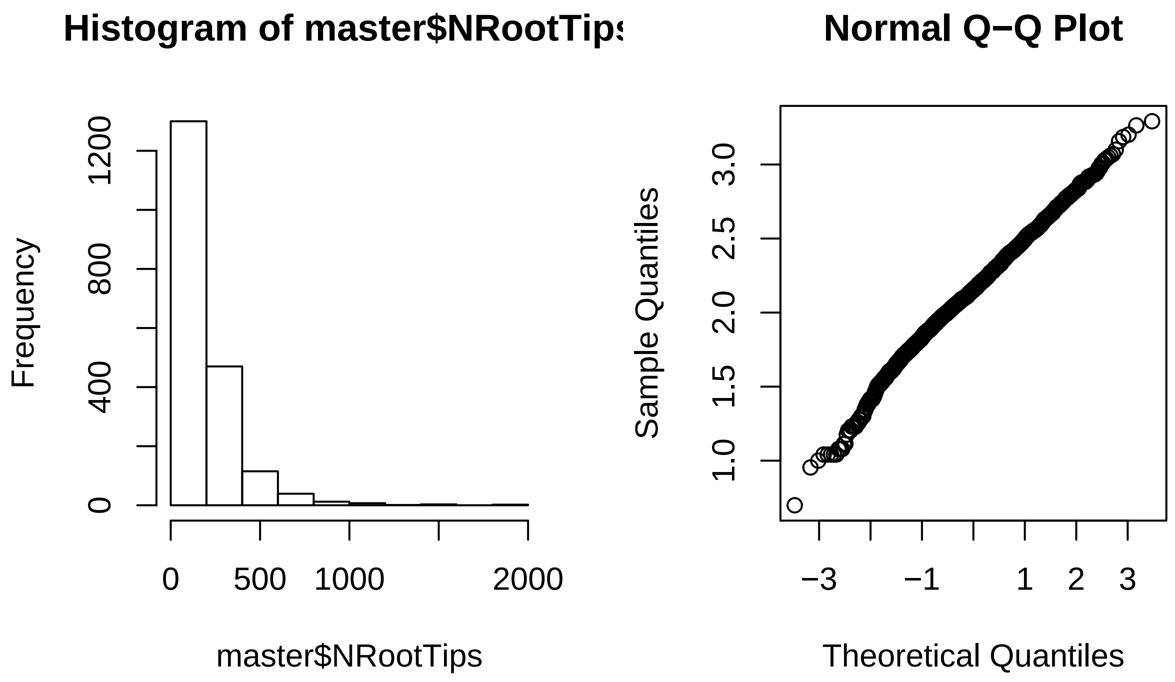


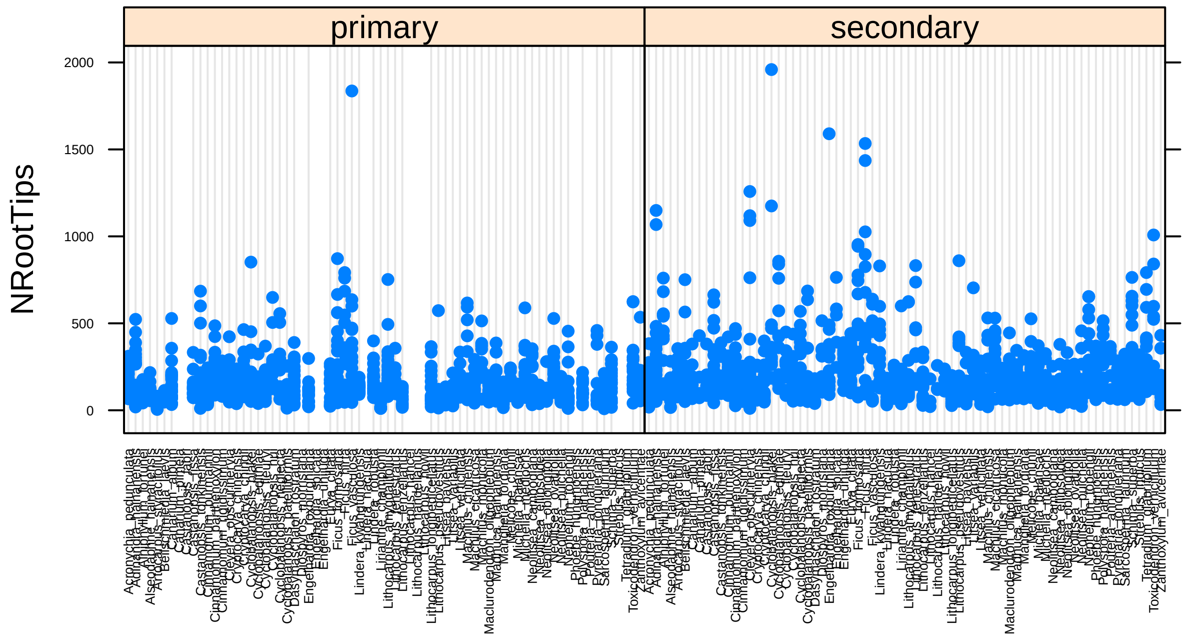

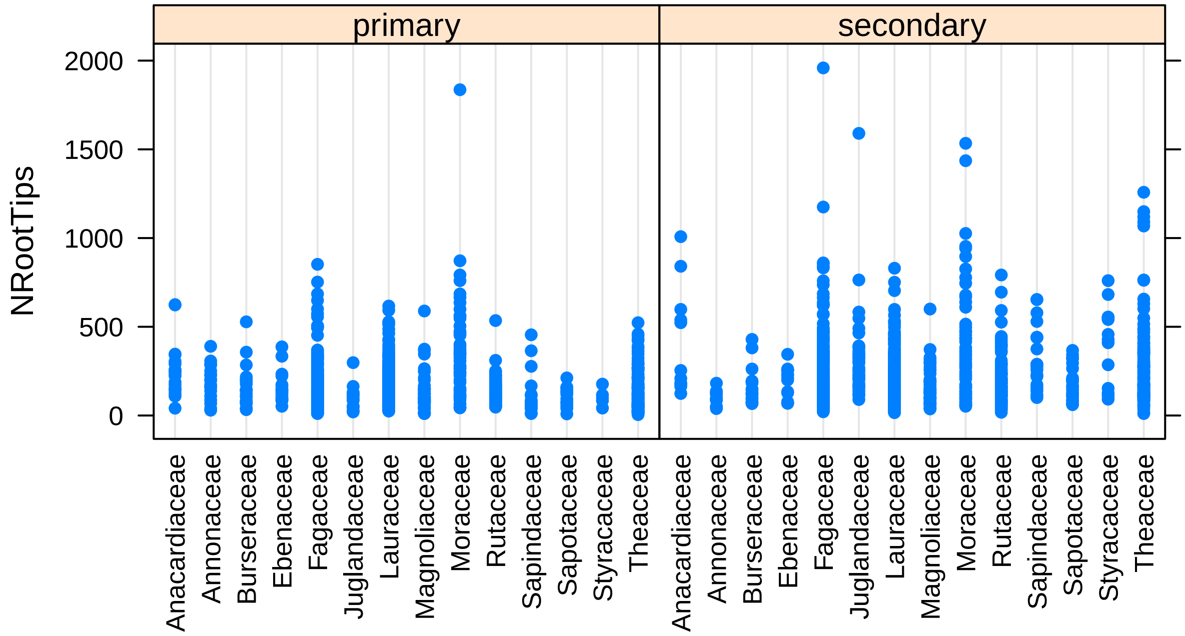


#### models

```
## Analysis of Variance Table
##
## Model 1: NRootTips ~ Gen.sp + forest.type
## Model 2: NRootTips ~ Gen.sp/Family + forest.type
## Model 3: NRootTips ~ Gen.sp/Family * forest.type
##   Res.Df      RSS Df Sum of Sq      F    Pr(>F)
## 1   1876 46575191
## 2   1876 46575191  0         0
## 3   1818 41362303 58   5212889 3.9504 < 2.2e-16 ***
## ---
## Signif. codes:  0 '***' 0.001 '**' 0.01 '*' 0.05 '.' 0.1 ' ' 1
```

```
## Analysis of Variance Table
##
## Response: NRootTips
##                       Df   Sum Sq Mean Sq F value    Pr(>F)
## Gen.sp                71 11772629  165812  7.2879 < 2.2e-16 ***
## forest.type            1  2205802 2205802 96.9518 < 2.2e-16 ***
## Gen.sp:forest.type    58  5212889   89877  3.9504 < 2.2e-16 ***
## Residuals           1818 41362303   22752
## ---
## Signif. codes:  0 '***' 0.001 '**' 0.01 '*' 0.05 '.' 0.1 ' ' 1
```

**predicted marginal means**

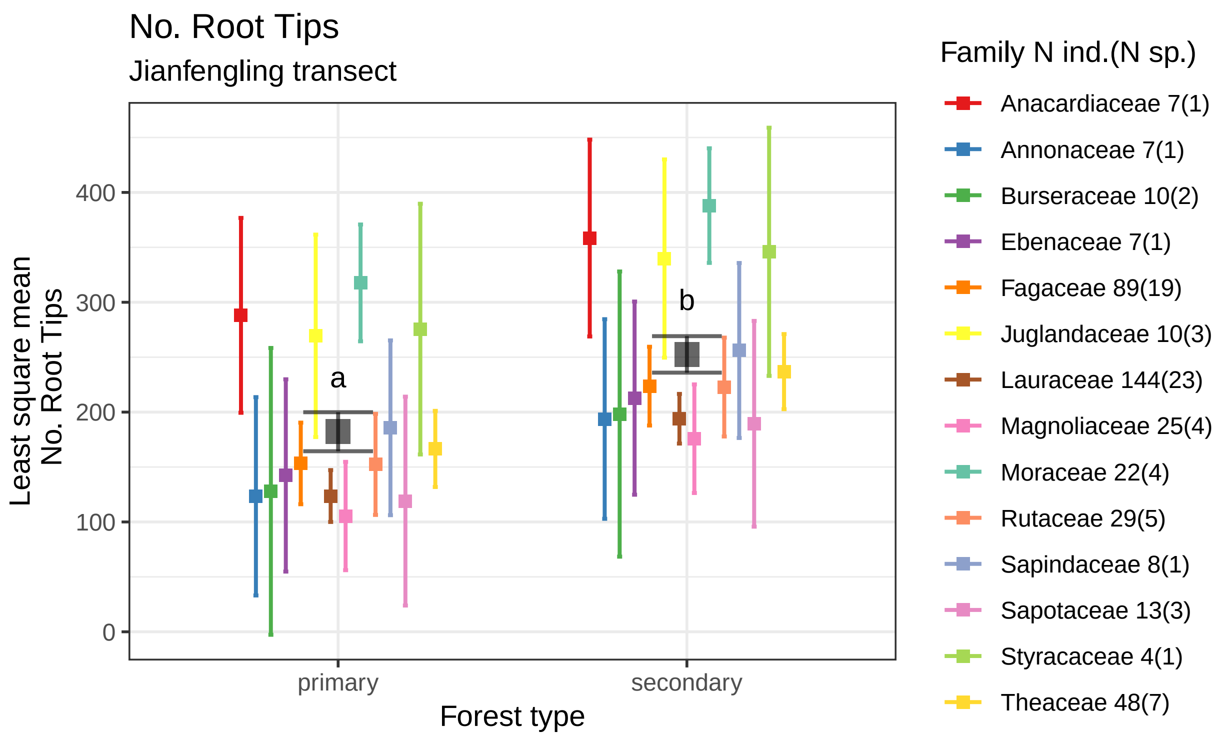


No. root tips estimates by forest type using data 423 tropical saplings from 72 species of 14 families.
Boxes indicate the LS mean. Error bars indicate the 95% confidence interval of the LS mean.
Means sharing a letter are not significantly different Tukey−adjusted comparisons

#### Root Tissue Density

##### plots

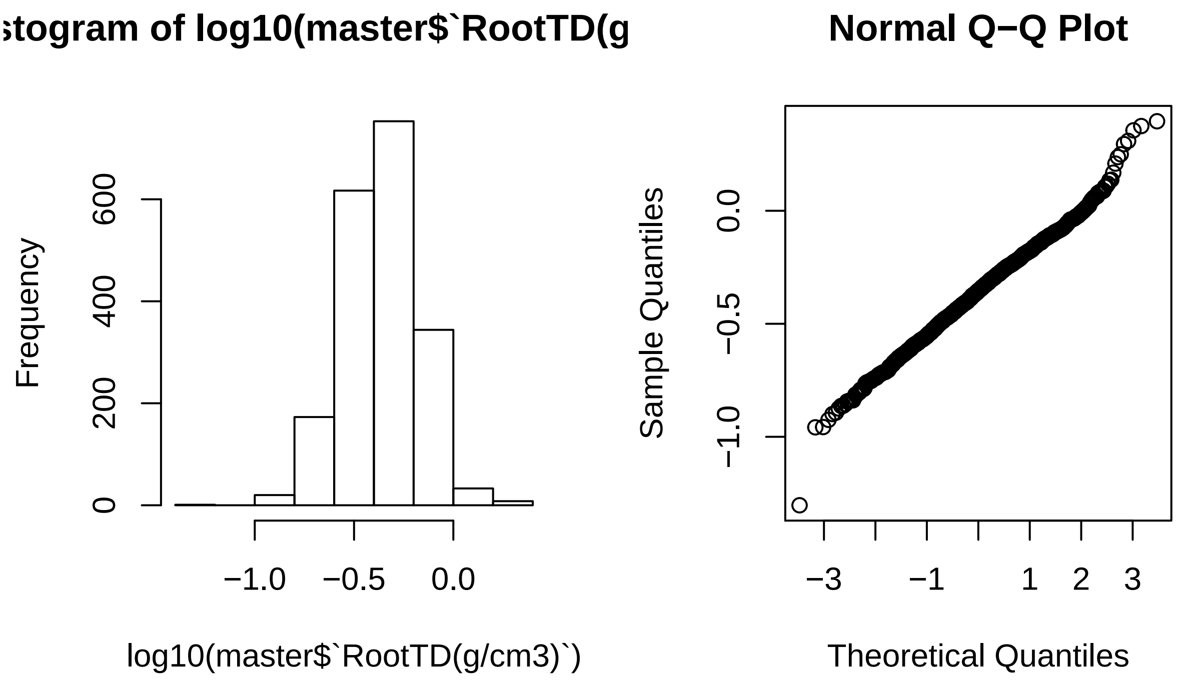


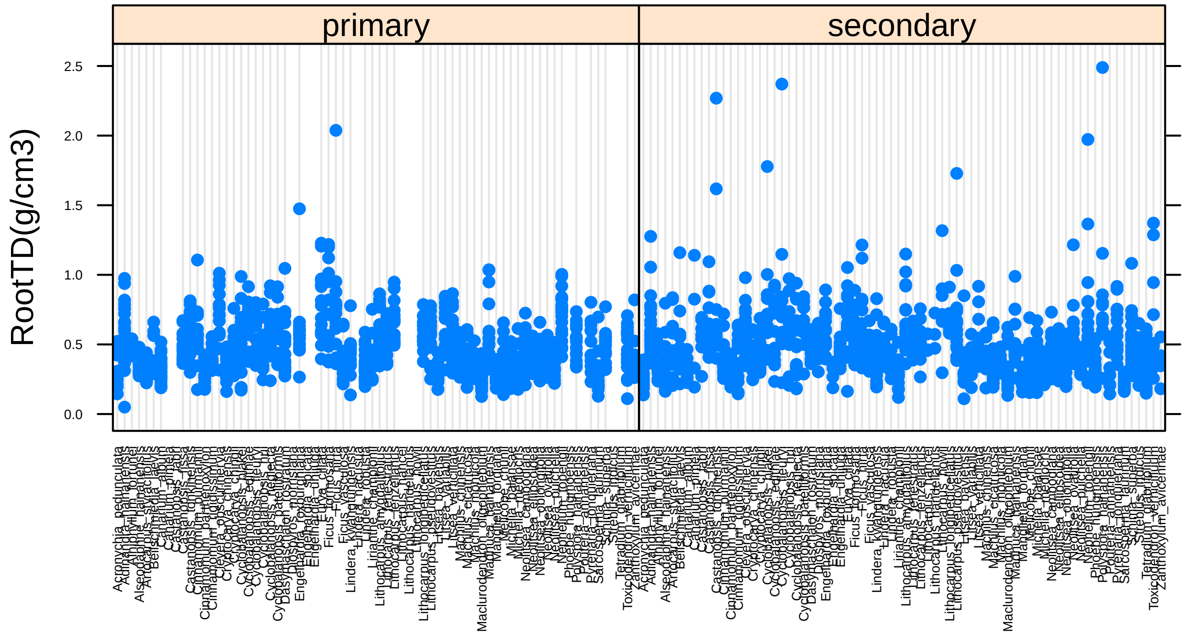

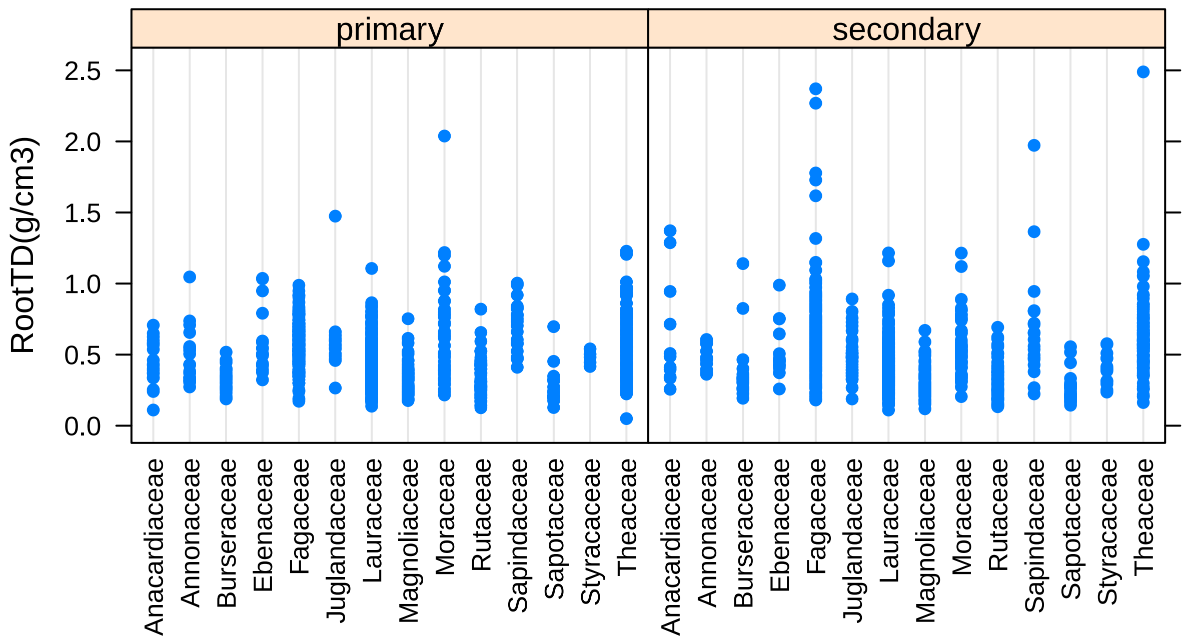


##### models

```
## Analysis of Variance Table
##
## Model 1: log10(`RootTD(g/cm3)`) ~ Gen.sp + forest.type
## Model 2: log10(`RootTD(g/cm3)`) ~ Gen.sp/Family + forest.type
## Model 3: log10(`RootTD(g/cm3)`) ~ Gen.sp/Family * forest.type
##   Res.Df    RSS Df Sum of Sq      F    Pr(>F)
## 1   1876 40.322
## 2   1876 40.322  0    0.0000
## 3   1818 37.145 58    3.1777 2.6815 2.724e-10 ***
## ---
## Signif. codes:  0 '***' 0.001 '**' 0.01 '*' 0.05 '.' 0.1 ' ' 1
```

```
## Analysis of Variance Table
##
## Response: log10(`RootTD(g/cm3)`)
##                       Df Sum Sq Mean Sq F value    Pr(>F)
## Gen.sp                71 28.946 0.40769 19.9537 < 2.2e-16 ***
## forest.type            1  0.020 0.01971  0.9649    0.3261
## Gen.sp:forest.type    58  3.178 0.05479  2.6815 2.724e-10 ***
## Residuals           1818 37.145 0.02043
## ---
## Signif. codes:  0 '***' 0.001 '**' 0.01 '*' 0.05 '.' 0.1 ' ' 1
```

**predicted marginal means**

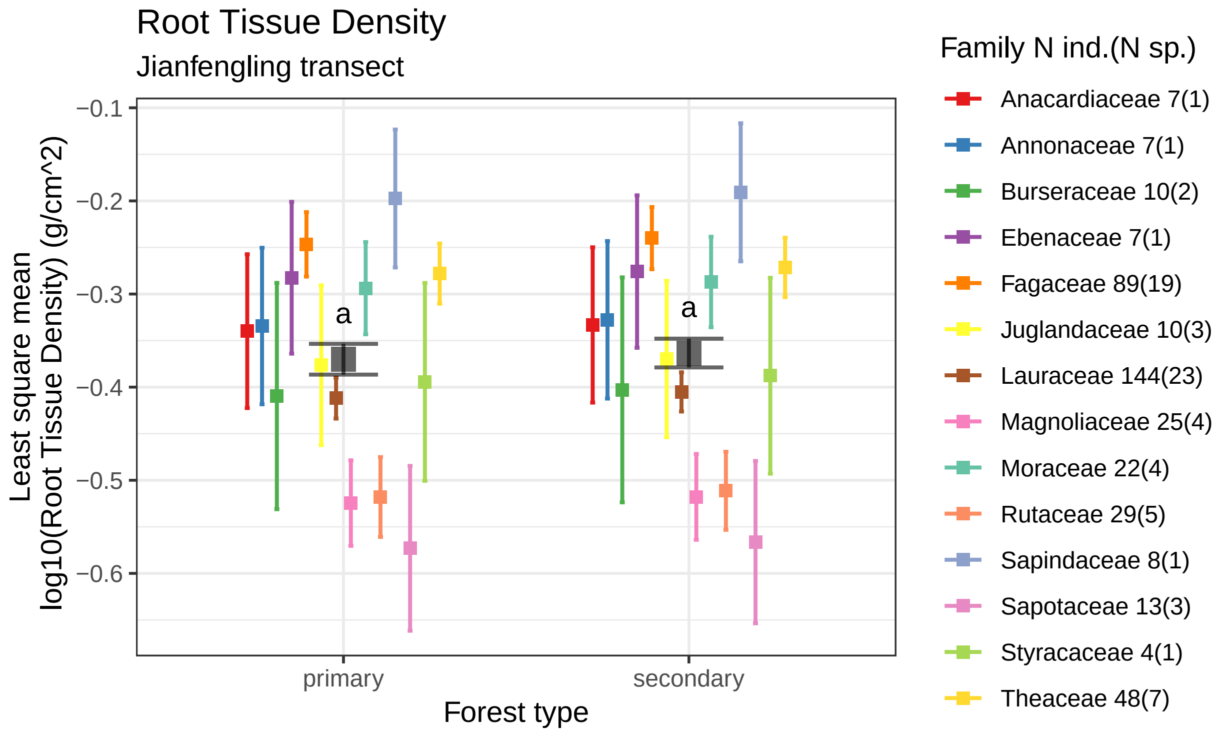


Number of root tips per length by forest type using data 423 tropical saplings from 72 species of 14 famili
Boxes indicate the LS mean. Error bars indicate the 95% confidence interval of the LS mean.
Means sharing a letter are not significantly different Tukey−adjusted comparisons
