## Supplement 3: Literate statistical document implementing the Trait Flex ANOVAS. for "Intraspecific root and leaf trait variation with tropical forest successional status: consequences for community-weighted patterns"

### JFL_Intraspecific_Variability_Effects

*JAH*

*July 20, 2018*

#### Background

In this literate stastitical document, we calculate Jan Leps *et al.*'s Intraspecific varability effects for some trait data collected in Jianfengling, Hainan Island China during the summer 2017. Above and belowground traits were collected including: leaf thickness, leaf dry mass, leaf area, leaf width, leaf height, leaf perimeter, SLA, root mass, average root diameter, number of root tips, number of root forks, root area, root length, root volume, specific root length, specific root area, specific root tip abundance, root length per volume, root tissue density, and root branchiness.

We have abundance and basal area data from a network of small plots established by Han Xu (Chinese Academy of Forestry, Guangzhou, China) - see:

- Xu et al. (2015) Journal of Applied Ecology 52:1044-1052.
- Xu et al. (2015) Journal of Ecology 103:1325-1333.

Briefly, abundances and sizes (for basal area calcuialtion) of the species on which traits were measured, were determined from plot inventory data for 164 0.0625 Ha plots established across the Jianfengling forest. 52 were in pirmary forest, while the remaining 112 were in secondary forest.

We use those basal area proporions to calcuate the "intraspecific variability effect" (see Jan Leps et al. 2011, Ecography 34: 856-863), which is defined as:

$Intraspecific\ variability\ effect = Specific\ average - Fixed\ average$, where the Fixed average is a site-independent weghted trait mean (in this case independent of forest type), thus we can note mathematically:

$$Fixed\ average = \sum_{i=1}^{s} p_i x_i$$

, where $p_i$ is the portion of the $ith$ species in the community and $x_i$ is the fixed point mean trait value. Similarly, the Specific average is the site-(or habitat, in this case Forest type)-specific weighted trait mean, given as:

$$Specific\ average = \sum_{i=1}^{s} p_i x_{i\_habitat}$$

The method goes futher to statisticall test the degree to which (intraspecific) trait variability interacts with species tunover, and decomposes that variation to allow for ecologially meaninful interpretaion of trait-variability, beta diversity interactions along environmental gradients.
The function used to do that (which is implemented here is trait.flex.anova() available in the supplemnetary material for Leps et al 2011. link here: www.oikosoffice.lu.se/appendix - Appendix E6904 in *Ecography*).

### Trait Flex ANOVAS

#### Leaf Mass

```
##
##  Decomposing trait sum of squares into composition turnover
##  effect, intraspecific trait variability, and their covariation:
```

```
##               Turnover Intraspec. Covariation   Total
## forest.type 2.1517e-05     2.1389   -0.013568  2.1254
## Residuals   1.5399e+01     4.9238   -3.512308 16.8103
## Total       1.5399e+01     7.0627   -3.525877 18.9357
##
##  Relative contributions:
##              Turnover Intraspec. Covariation  Total
## forest.type 1.136e-06      0.113  -0.0007165 0.1122
## Residuals   8.132e-01      0.260  -0.1854860 0.8878
## Total       8.132e-01      0.373  -0.1862026 1.0000
##
##  Significance of testable effects:
##             Turnover Intraspec.     Total
## forest.type  0.98801 2.2881e-14 1.1598e-05
```

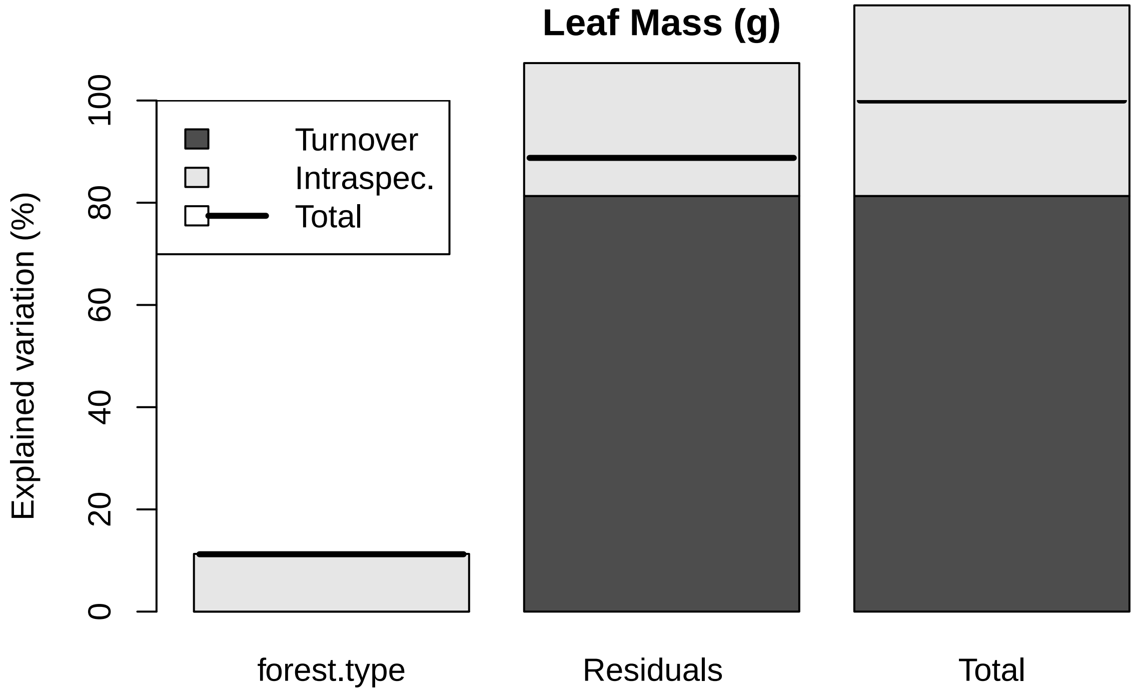


##### Leaf Area

```
##
##  Decomposing trait sum of squares into composition turnover
##  effect, intraspecific trait variability, and their covariation:
##              Turnover Intraspec. Covariation   Total
## forest.type    19.937     3696.9      542.98  4259.8
## Residuals   81149.233    12628.9    -9135.77 84642.4
## Total       81169.171    16325.8    -8592.79 88902.2
##
```

```
##  Relative contributions:
##              Turnover Intraspec. Covariation   Total
## forest.type 0.0002243    0.04158    0.006108 0.04792
## Residuals   0.9127925    0.14205   -0.102762 0.95208
## Total       0.9130167    0.18364   -0.096654 1.00000
##
##  Significance of testable effects:
##             Turnover Intraspec.    Total
## forest.type  0.84212 1.1972e-10 0.004862
```

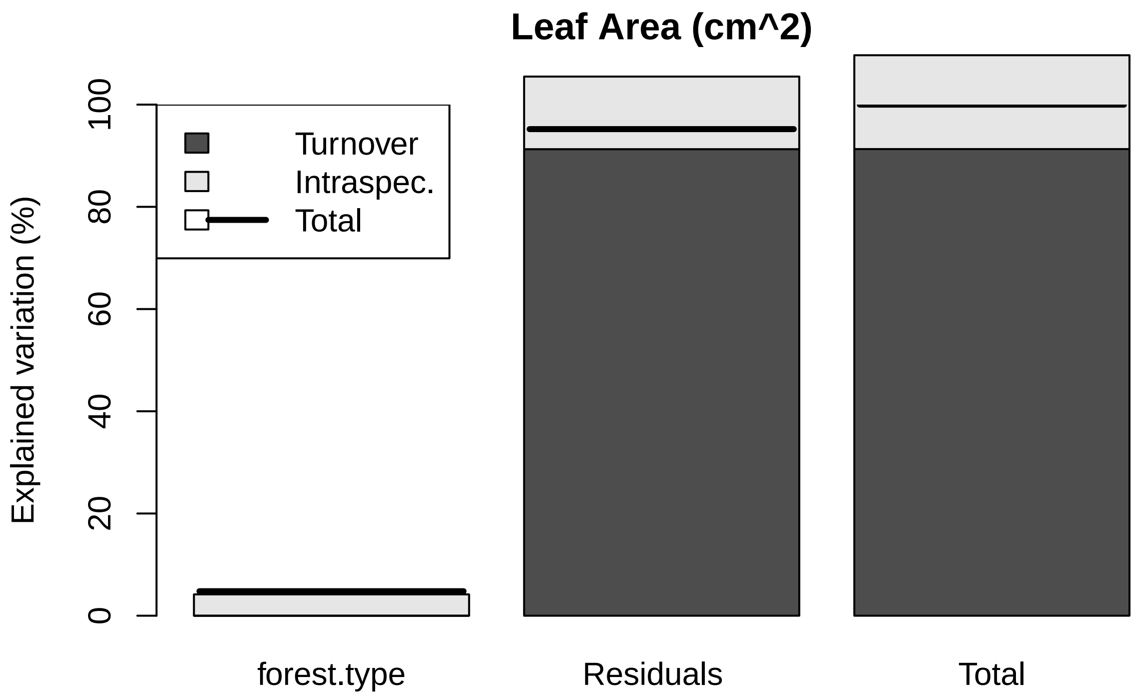


##### SLA

```
##
##  Decomposing trait sum of squares into composition turnover
##  effect, intraspecific trait variability, and their covariation:
##             Turnover Intraspec. Covariation   Total
## forest.type   3.4563     94.118     -36.072  61.502
## Residuals   372.9233    135.971      12.875 521.769
## Total       376.3796    230.088     -23.198 583.270
##
##  Relative contributions:
##             Turnover Intraspec. Covariation  Total
## forest.type 0.005926     0.1614    -0.06184 0.1054
## Residuals   0.639366     0.2331     0.02207 0.8946
## Total       0.645292     0.3945    -0.03977 1.0000
```

```
##
##  Significance of testable effects:
##             Turnover Intraspec.      Total
## forest.type  0.22223 3.0372e-20 2.2137e-05
```

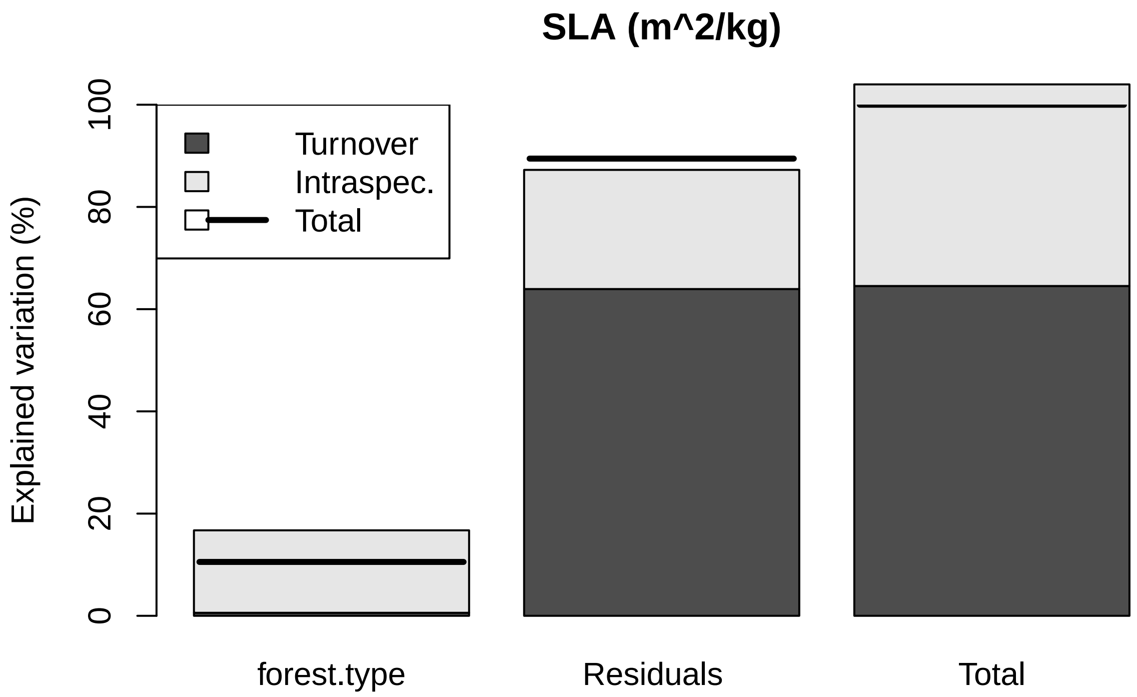


##### Leaf Perimiter

```
##
##  Decomposing trait sum of squares into composition turnover
##  effect, intraspecific trait variability, and their covariation:
##               Turnover Intraspec. Covariation    Total
## forest.type 1.6166e+00     6358.3     -202.77   6157.1
## Residuals   2.0246e+05    33842.2   -31340.68 204960.2
## Total       2.0246e+05    40200.5   -31543.45 211117.3
##
##  Relative contributions:
##              Turnover Intraspec. Covariation   Total
## forest.type 7.658e-06    0.03012  -0.0009605 0.02916
## Residuals   9.590e-01    0.16030  -0.1484515 0.97084
## Total       9.590e-01    0.19042  -0.1494119 1.00000
##
##  Significance of testable effects:
##             Turnover Intraspec.    Total
## forest.type  0.97135 1.3405e-07 0.028789
```

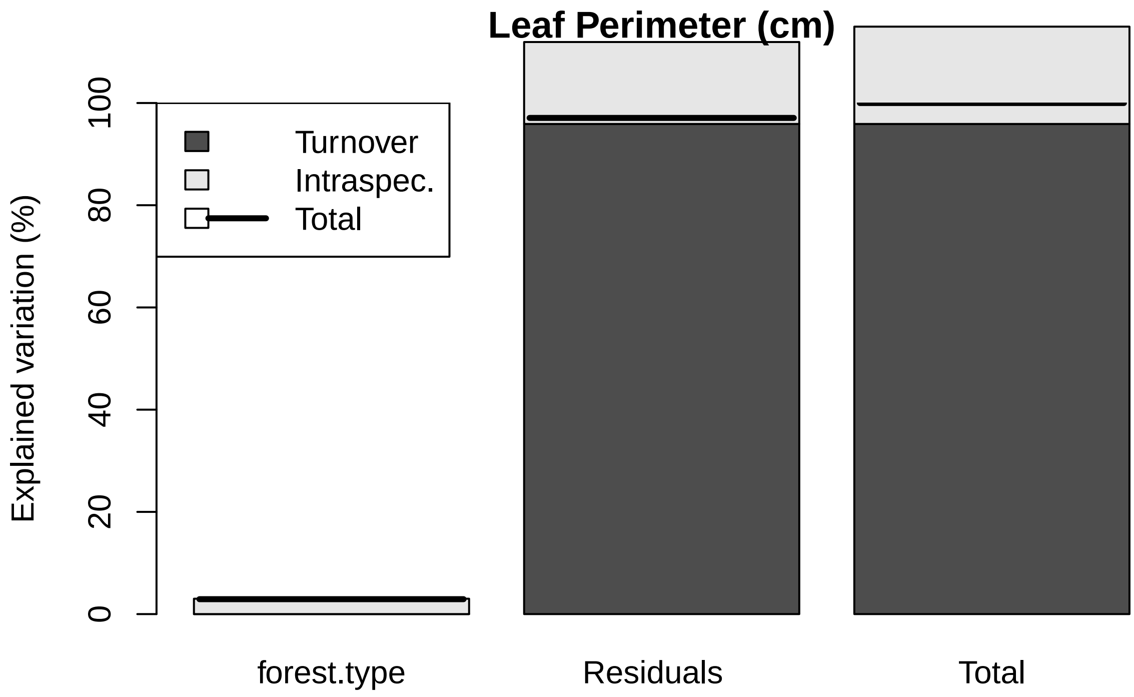


##### Leaf Width

```
##
##  Decomposing trait sum of squares into composition turnover
##  effect, intraspecific trait variability, and their covariation:
##               Turnover Intraspec. Covariation    Total
## forest.type    0.19403      62.12     -6.9435   55.371
## Residuals   1481.23115     326.08   -121.4866 1685.821
## Total       1481.42518     388.20   -128.4302 1741.191
##
##  Relative contributions:
##              Turnover Intraspec. Covariation  Total
## forest.type 0.0001114    0.03568   -0.003988 0.0318
## Residuals   0.8506998    0.18727   -0.069772 0.9682
## Total       0.8508113    0.22295   -0.073760 1.0000
##
##  Significance of testable effects:
##             Turnover Intraspec.    Total
## forest.type  0.88436 1.1147e-07 0.022338
```

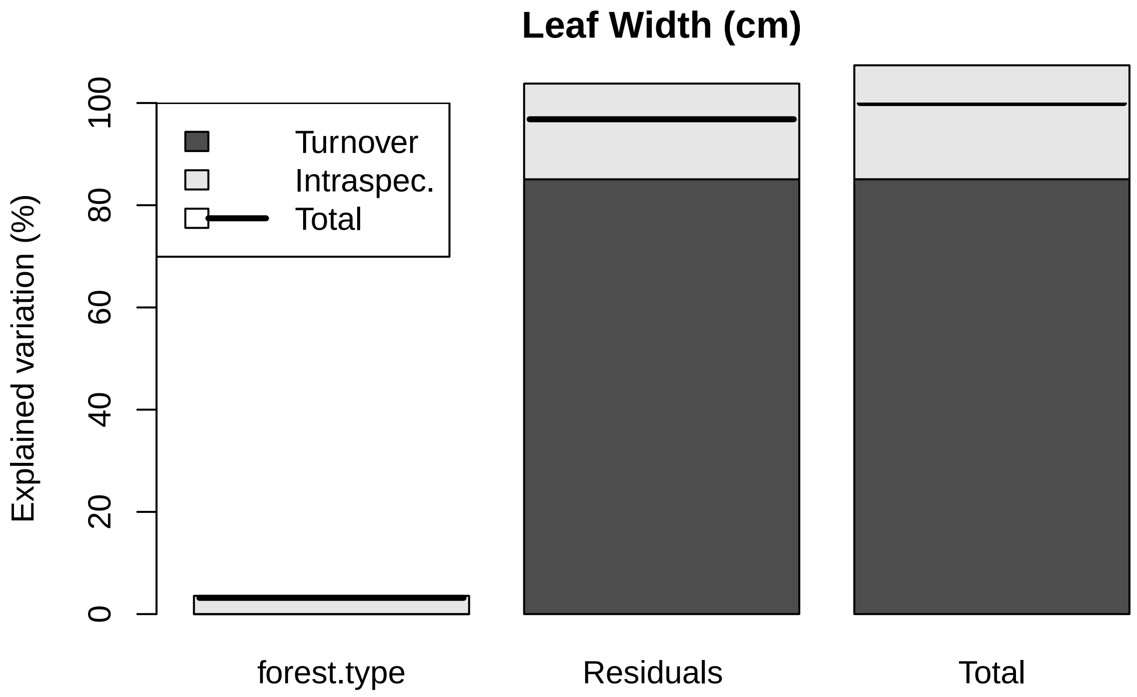


##### Leaf Height

```
##
##  Decomposing trait sum of squares into composition turnover
##  effect, intraspecific trait variability, and their covariation:
##              Turnover Intraspec. Covariation   Total
## forest.type   0.57013     87.025     -14.088  73.508
## Residuals   254.40949    180.629     -68.093 366.945
## Total       254.97961    267.654     -82.181 440.452
##
##  Relative contributions:
##             Turnover Intraspec. Covariation  Total
## forest.type 0.001294     0.1976    -0.03198 0.1669
## Residuals   0.577609     0.4101    -0.15460 0.8331
## Total       0.578904     0.6077    -0.18658 1.0000
##
##  Significance of testable effects:
##             Turnover Intraspec.      Total
## forest.type  0.54767 1.5899e-15 5.6203e-08
```

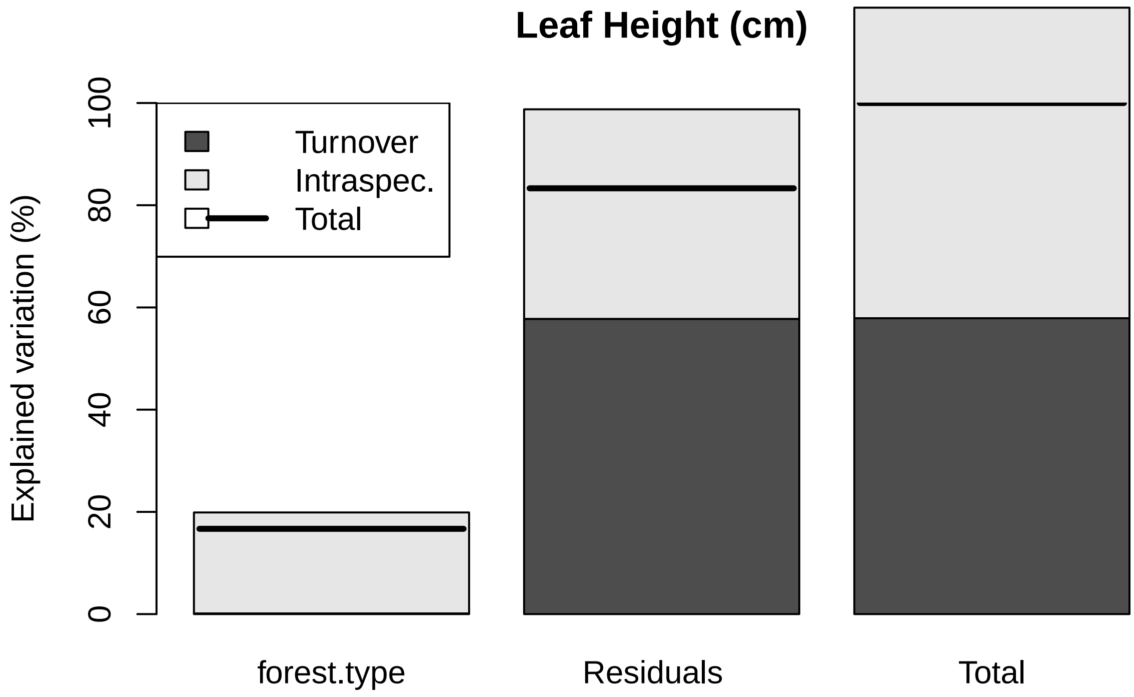


##### Leaf Thickness

```
##
##  Decomposing trait sum of squares into composition turnover
##  effect, intraspecific trait variability, and their covariation:
##               Turnover Intraspec. Covariation    Total
## forest.type 8.0338e-05   0.020167  -0.0025457 0.017702
## Residuals   6.9865e-02   0.036661  -0.0051287 0.101398
## Total       6.9946e-02   0.056828  -0.0076744 0.119099
##
##  Relative contributions:
##              Turnover Intraspec. Covariation  Total
## forest.type 0.0006745     0.1693    -0.02137 0.1486
## Residuals   0.5866146     0.3078    -0.04306 0.8514
## Total       0.5872891     0.4771    -0.06444 1.0000
##
##  Significance of testable effects:
##             Turnover Intraspec.      Total
## forest.type   0.6666 3.9616e-17 3.4362e-07
```

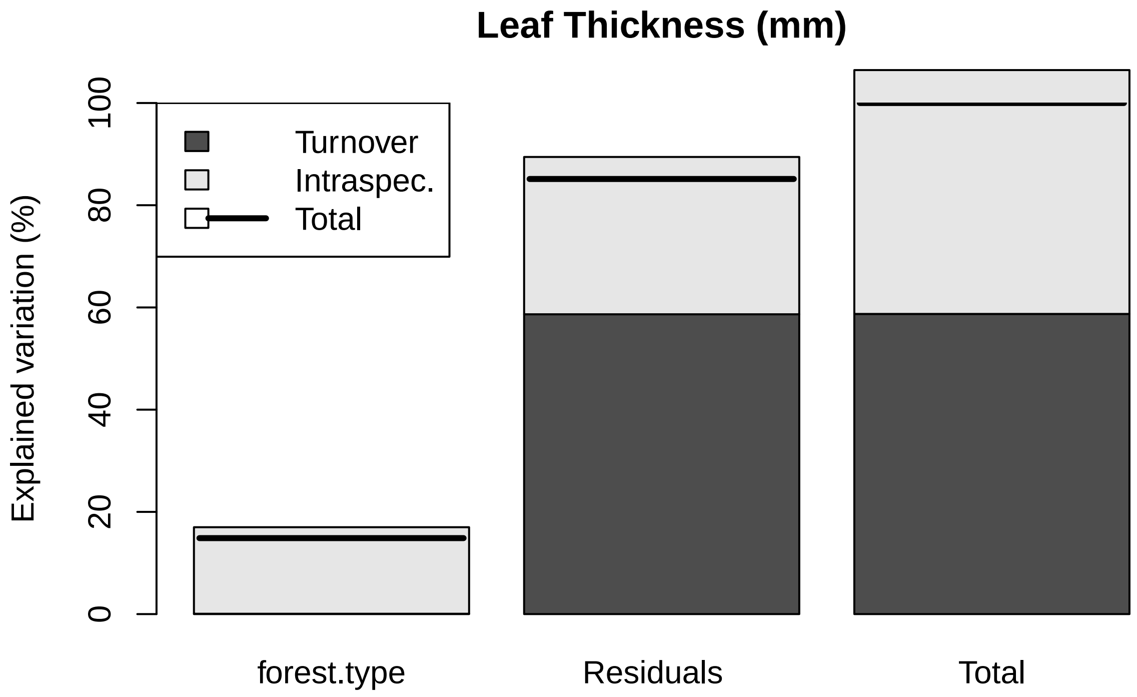


##### Root Mass

```
##
##  Decomposing trait sum of squares into composition turnover
##  effect, intraspecific trait variability, and their covariation:
##               Turnover Intraspec. Covariation   Total
## forest.type 0.00096802    0.36055   -0.037364 0.32415
## Residuals   0.41840428    0.34641    0.083230 0.84805
## Total       0.41937230    0.70696    0.045866 1.17220
##
##  Relative contributions:
##              Turnover Intraspec. Covariation  Total
## forest.type 0.0008258     0.3076    -0.03188 0.2765
## Residuals   0.3569403     0.2955     0.07100 0.7235
## Total       0.3577661     0.6031     0.03913 1.0000
##
##  Significance of testable effects:
##             Turnover Intraspec.      Total
## forest.type  0.54126 7.0217e-27 4.8079e-13
```

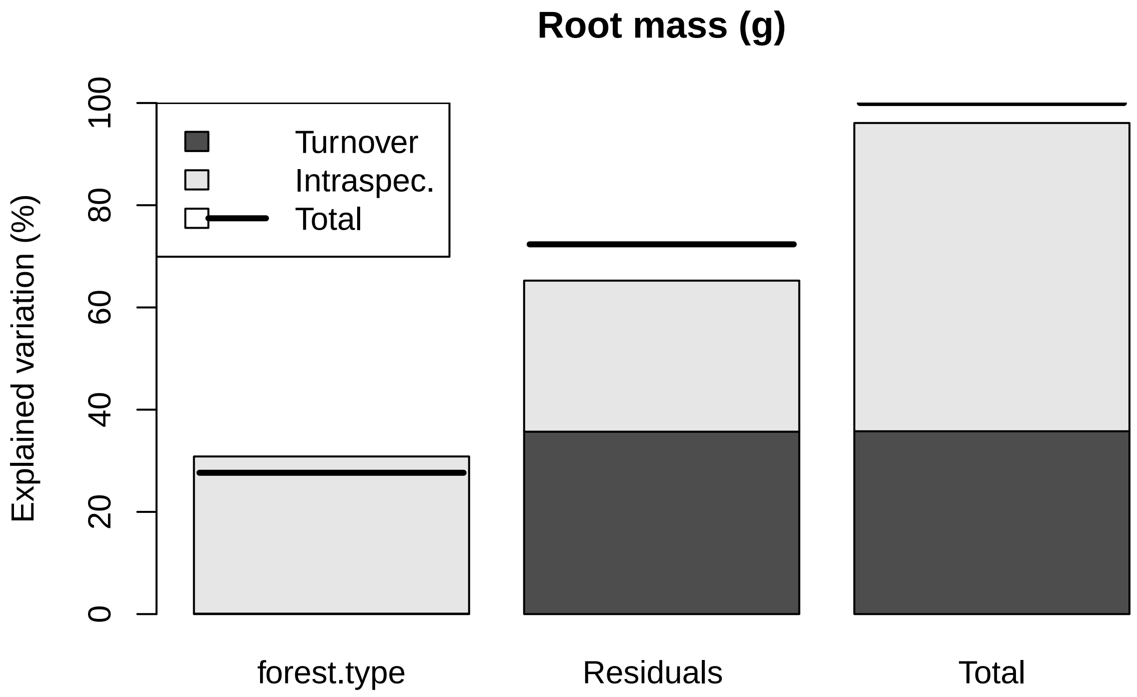


##### Root Avg Diameter

```
##
##  Decomposing trait sum of squares into composition turnover
##  effect, intraspecific trait variability, and their covariation:
##              Turnover Intraspec. Covariation    Total
## forest.type 0.0030078   0.065729   -0.028121 0.040615
## Residuals   1.6687548   0.577961   -0.159367 2.087349
## Total       1.6717627   0.643690   -0.187489 2.127964
##
##  Relative contributions:
##             Turnover Intraspec. Covariation   Total
## forest.type 0.001413    0.03089    -0.01322 0.01909
## Residuals   0.784203    0.27160    -0.07489 0.98091
## Total       0.785616    0.30249    -0.08811 1.00000
##
##  Significance of testable effects:
##             Turnover Intraspec.    Total
## forest.type  0.58969  3.035e-05 0.077704
```

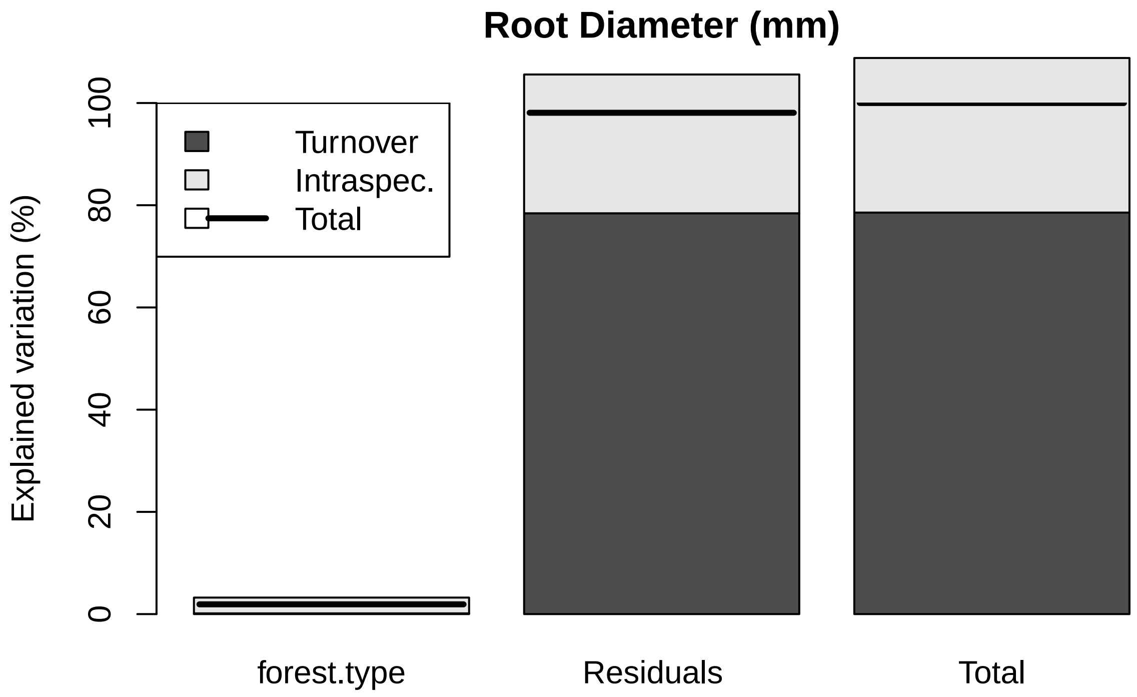


##### Number of Root Tips

```
##
##  Decomposing trait sum of squares into composition turnover
##  effect, intraspecific trait variability, and their covariation:
##              Turnover Intraspec. Covariation  Total
## forest.type    948.09     339022      -35857 304114
## Residuals   114996.21     103893      -30342 188547
## Total       115944.30     442915      -66199 492660
##
##  Relative contributions:
##             Turnover Intraspec. Covariation  Total
## forest.type 0.001924     0.6881    -0.07278 0.6173
## Residuals   0.233419     0.2109    -0.06159 0.3827
## Total       0.235343     0.8990    -0.13437 1.0000
##
##  Significance of testable effects:
##             Turnover Intraspec.      Total
## forest.type  0.24951 7.0026e-53 1.2948e-35
```

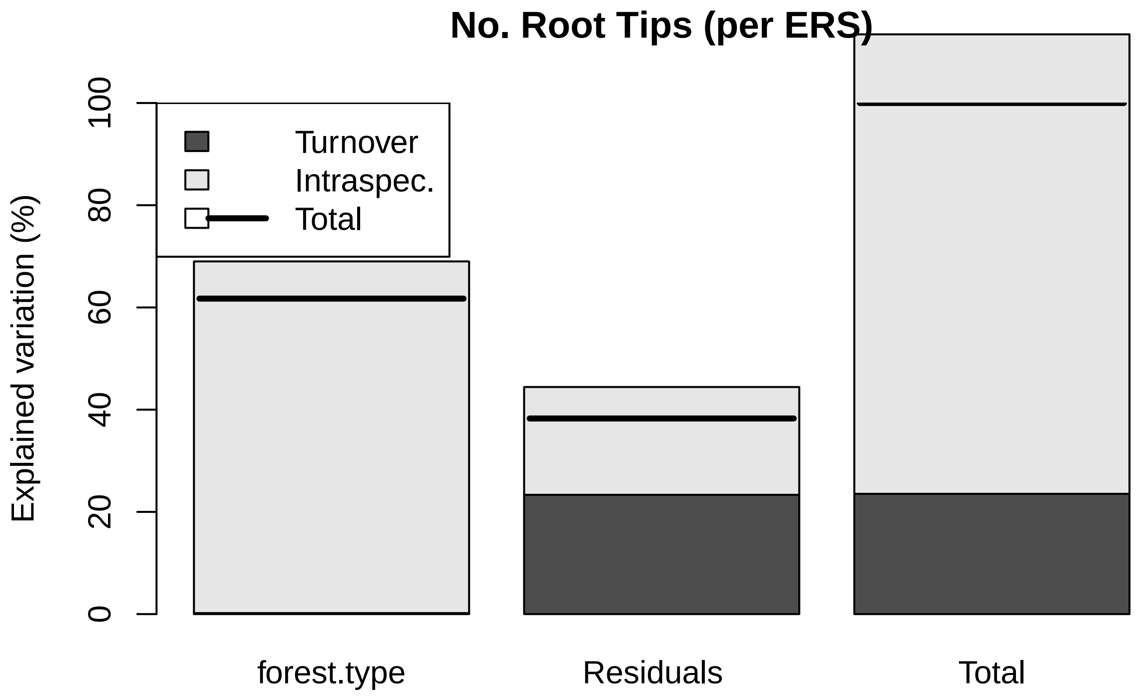


##### Number of Root Forks

```
##
##  Decomposing trait sum of squares into composition turnover
##  effect, intraspecific trait variability, and their covariation:
##             Turnover Intraspec. Covariation  Total
## forest.type   8659.2     194912      -82166 121406
## Residuals   428109.7     253159     -154146 527123
## Total       436768.9     448072     -236311 648529
##
##  Relative contributions:
##             Turnover Intraspec. Covariation  Total
## forest.type  0.01335     0.3005     -0.1267 0.1872
## Residuals    0.66012     0.3904     -0.2377 0.8128
## Total        0.67348     0.6909     -0.3644 1.0000
##
##  Significance of testable effects:
##             Turnover Intraspec.      Total
## forest.type 0.072122 7.7566e-22 7.2131e-09
```

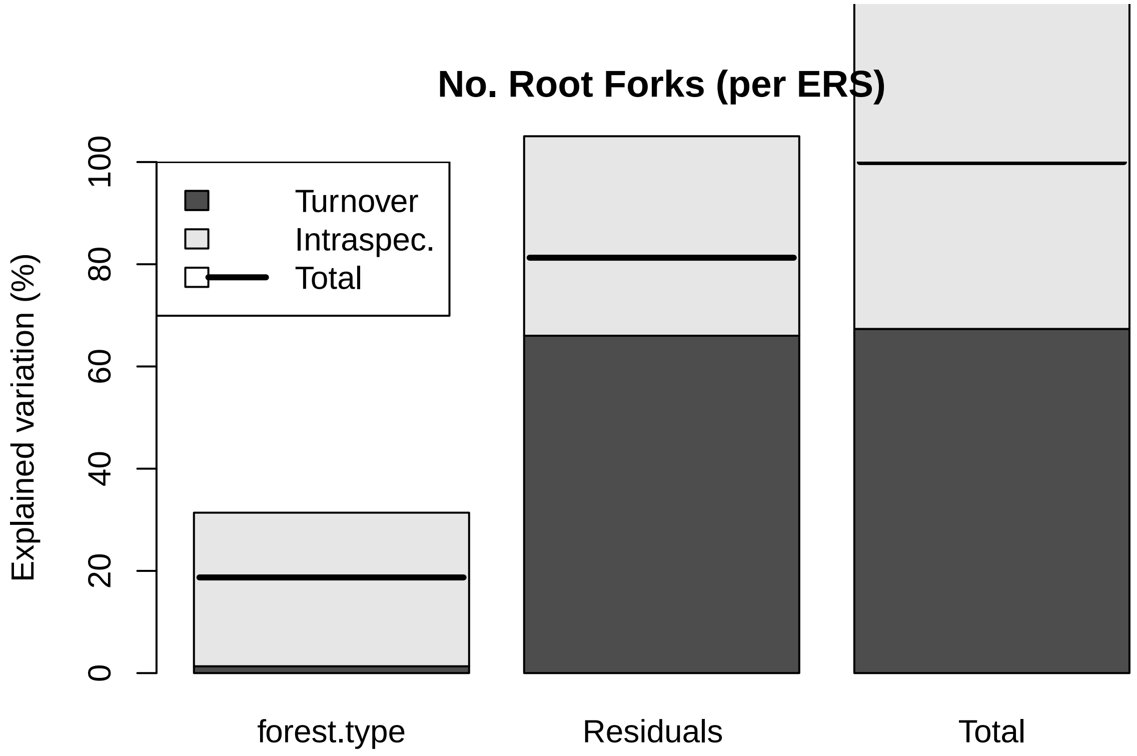


##### Root Area

```
##
##  Decomposing trait sum of squares into composition turnover
##  effect, intraspecific trait variability, and their covariation:
##             Turnover Intraspec. Covariation  Total
## forest.type   1.2226    1746.81     -92.427 1655.6
## Residuals   983.5863     631.11     -22.827 1591.9
## Total       984.8089    2377.92    -115.254 3247.5
##
##  Relative contributions:
##              Turnover Intraspec. Covariation  Total
## forest.type 0.0003765     0.5379   -0.028461 0.5098
## Residuals   0.3028771     0.1943   -0.007029 0.4902
## Total       0.3032536     0.7322   -0.035490 1.0000
##
##  Significance of testable effects:
##             Turnover Intraspec.     Total
## forest.type  0.65422 1.5813e-48 7.237e-27
```

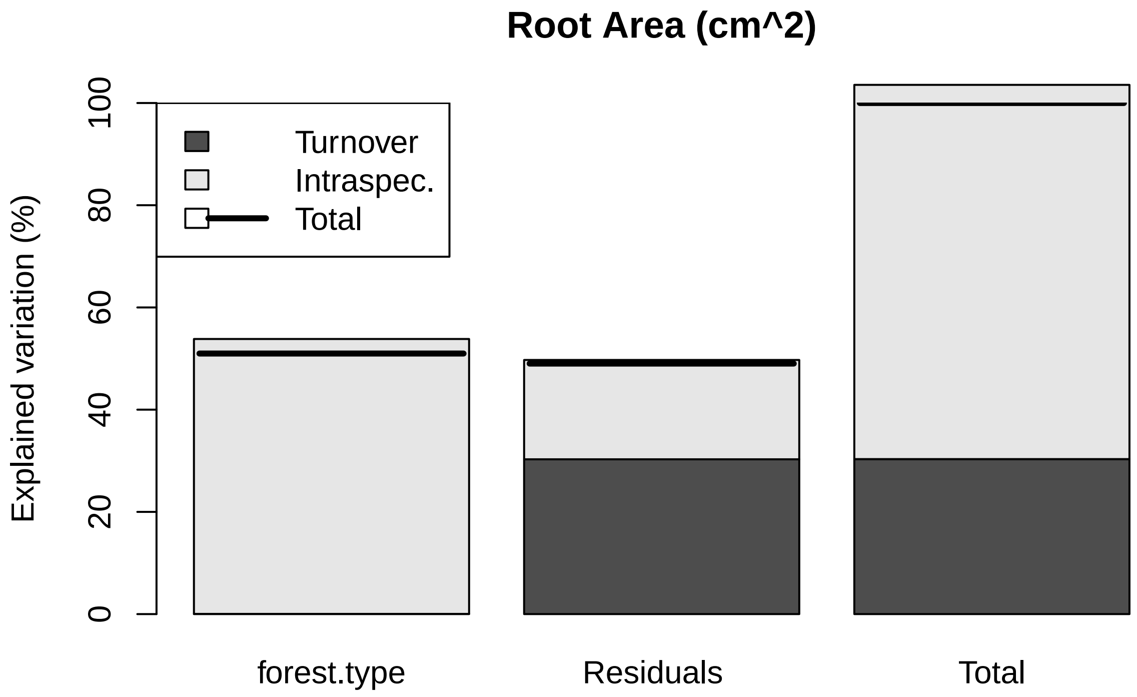


##### Root Length

```
##
##  Decomposing trait sum of squares into composition turnover
##  effect, intraspecific trait variability, and their covariation:
##             Turnover Intraspec. Covariation Total
## forest.type    198.6      40316     -5659.3 34856
## Residuals    17729.3      16747     -8706.6 25769
## Total        17927.9      57063    -14365.9 60625
##
##  Relative contributions:
##             Turnover Intraspec. Covariation  Total
## forest.type 0.003276     0.6650    -0.09335 0.5749
## Residuals   0.292443     0.2762    -0.14361 0.4251
## Total       0.295719     0.9412    -0.23696 1.0000
##
##  Significance of testable effects:
##             Turnover Intraspec.      Total
## forest.type  0.17983 5.5479e-45 6.5956e-32
```

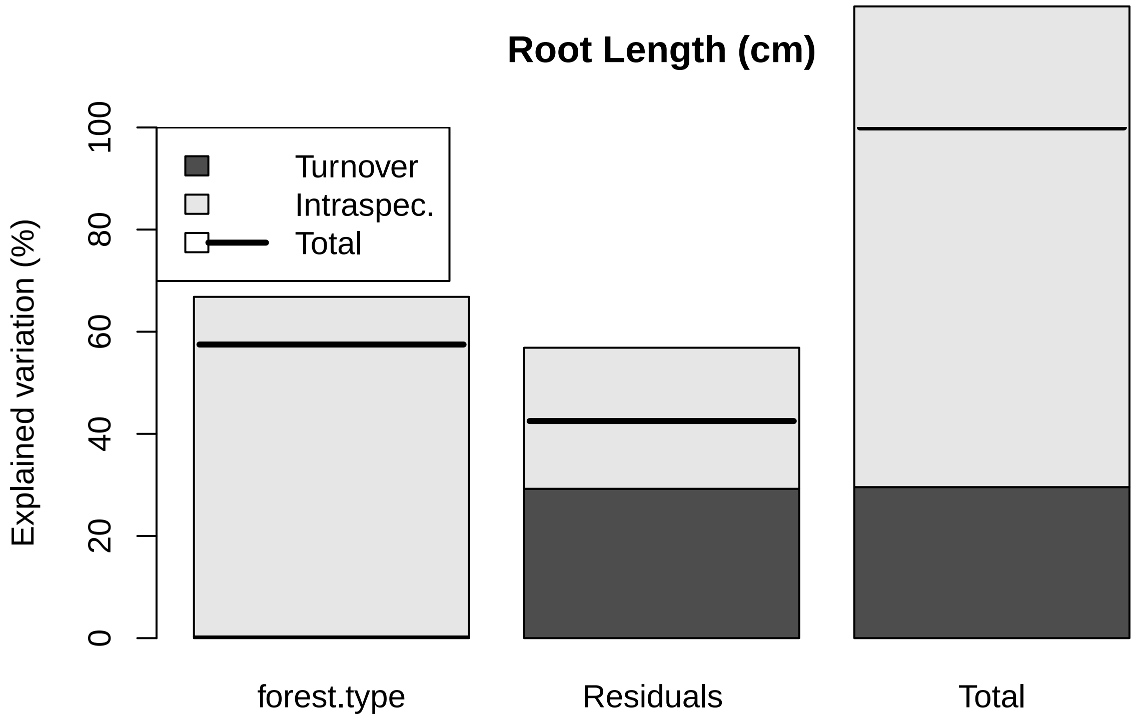


##### Root Volume

```
##
##  Decomposing trait sum of squares into composition turnover
##  effect, intraspecific trait variability, and their covariation:
##             Turnover Intraspec. Covariation   Total
## forest.type 0.003203    0.62339    0.089369 0.71596
## Residuals   2.303241    0.51094    0.246439 3.06062
## Total       2.306444    1.13433    0.335808 3.77658
##
##  Relative contributions:
##              Turnover Intraspec. Covariation  Total
## forest.type 0.0008481     0.1651     0.02366 0.1896
## Residuals   0.6098750     0.1353     0.06525 0.8104
## Total       0.6107231     0.3004     0.08892 1.0000
##
##  Significance of testable effects:
##             Turnover Intraspec.      Total
## forest.type  0.63568 7.3841e-30 5.6557e-09
```

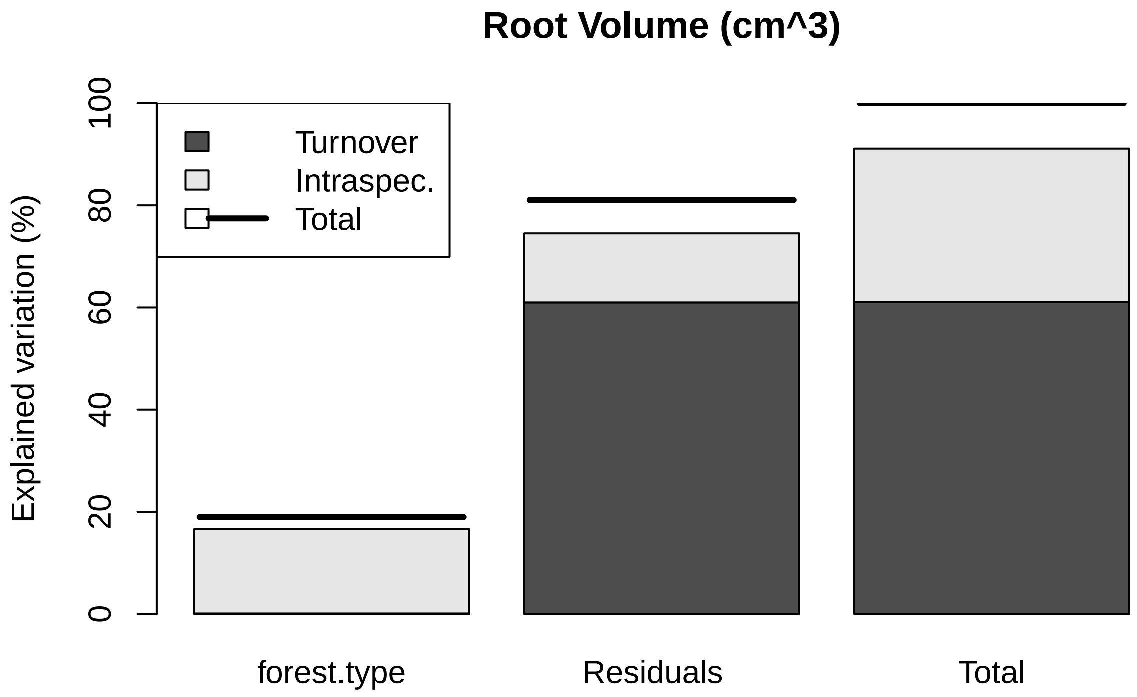


##### SRL

```
##
##  Decomposing trait sum of squares into composition turnover
##  effect, intraspecific trait variability, and their covariation:
##              Turnover Intraspec. Covariation   Total
## forest.type    81.328     3285.6      1033.9  4400.8
## Residuals   15787.988     5985.7     -2606.2 19167.5
## Total       15869.316     9271.3     -1572.4 23568.3
##
##  Relative contributions:
##             Turnover Intraspec. Covariation  Total
## forest.type 0.003451     0.1394     0.04387 0.1867
## Residuals   0.669883     0.2540    -0.11058 0.8133
## Total       0.673334     0.3934    -0.06671 1.0000
##
##  Significance of testable effects:
##             Turnover Intraspec.     Total
## forest.type  0.36233 4.2147e-17 7.572e-09
```

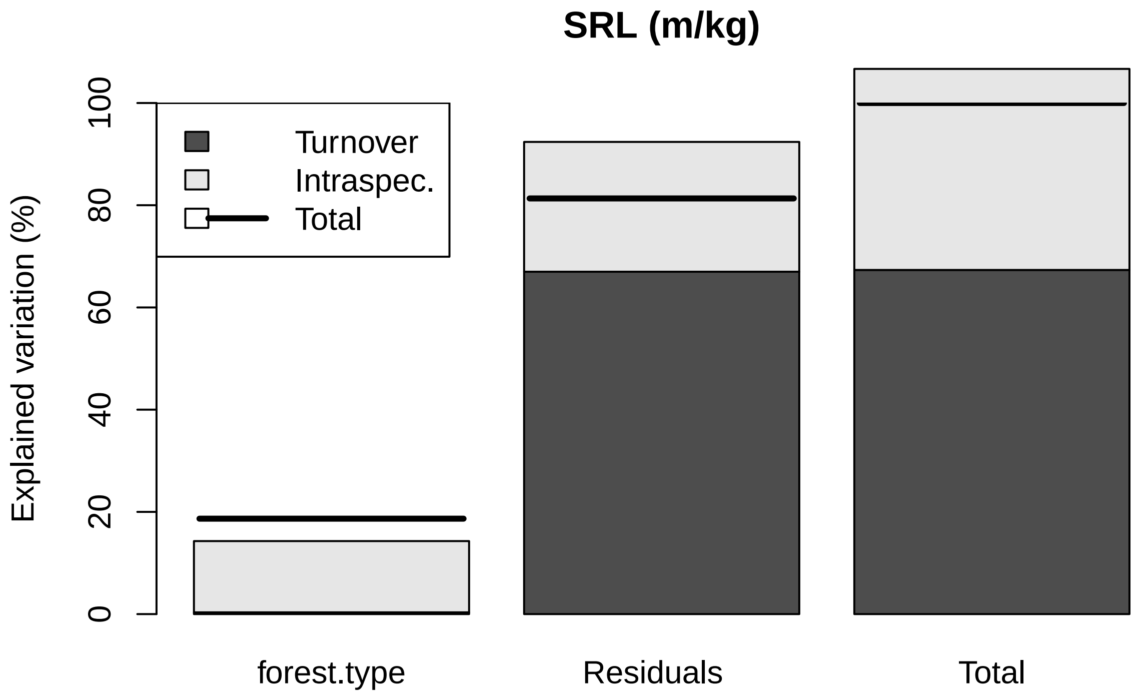


##### SRA

```
##
##  Decomposing trait sum of squares into composition turnover
##  effect, intraspecific trait variability, and their covariation:
##             Turnover Intraspec. Covariation  Total
## forest.type   4.2499     73.129      35.258 112.64
## Residuals   367.3441    197.914     -69.147 496.11
## Total       371.5940    271.043     -33.889 608.75
##
##  Relative contributions:
##             Turnover Intraspec. Covariation Total
## forest.type 0.006981     0.1201     0.05792 0.185
## Residuals   0.603442     0.3251    -0.11359 0.815
## Total       0.610423     0.4452    -0.05567 1.000
##
##  Significance of testable effects:
##             Turnover Intraspec.      Total
## forest.type  0.17289 1.0293e-12 9.0016e-09
```

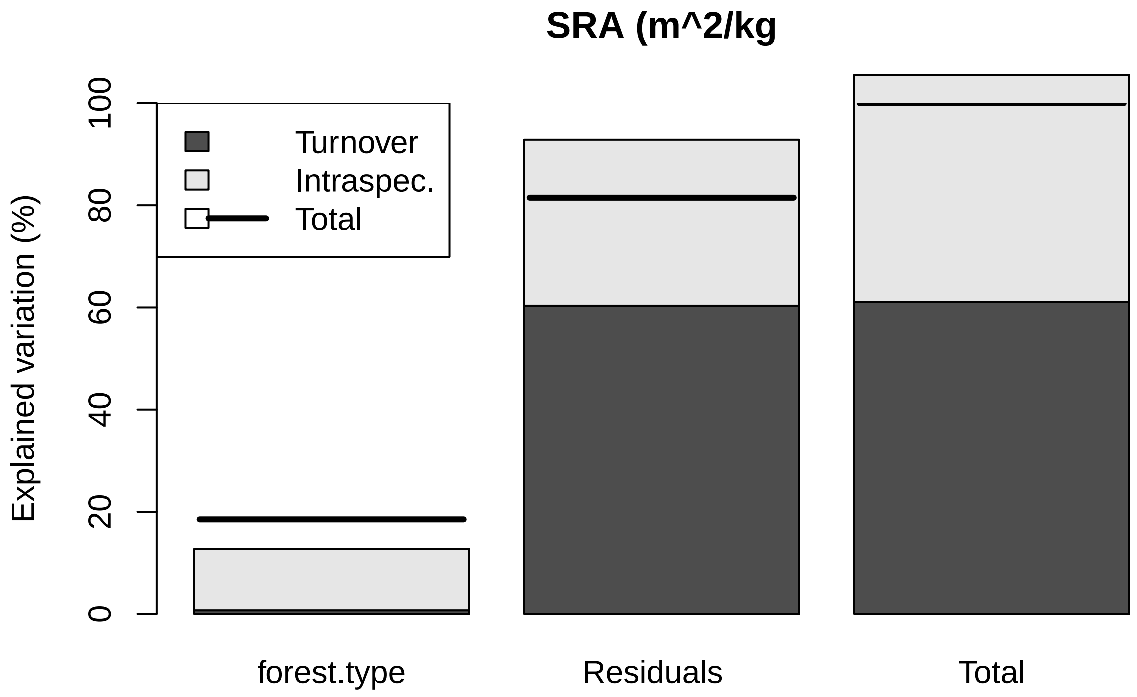


##### SR Tip Abundance

```
##
##  Decomposing trait sum of squares into composition turnover
##  effect, intraspecific trait variability, and their covariation:
##             Turnover Intraspec. Covariation    Total
## forest.type    63932    3763684      981064  4808680
## Residuals    9454703    2975285     -387383 12042604
## Total        9518635    6738968      593681 16851285
##
##  Relative contributions:
##             Turnover Intraspec. Covariation  Total
## forest.type 0.003794     0.2233     0.05822 0.2854
## Residuals   0.561067     0.1766    -0.02299 0.7146
## Total       0.564861     0.3999     0.03523 1.0000
##
##  Significance of testable effects:
##             Turnover Intraspec.      Total
## forest.type  0.29683 1.4477e-30 1.7519e-13
```

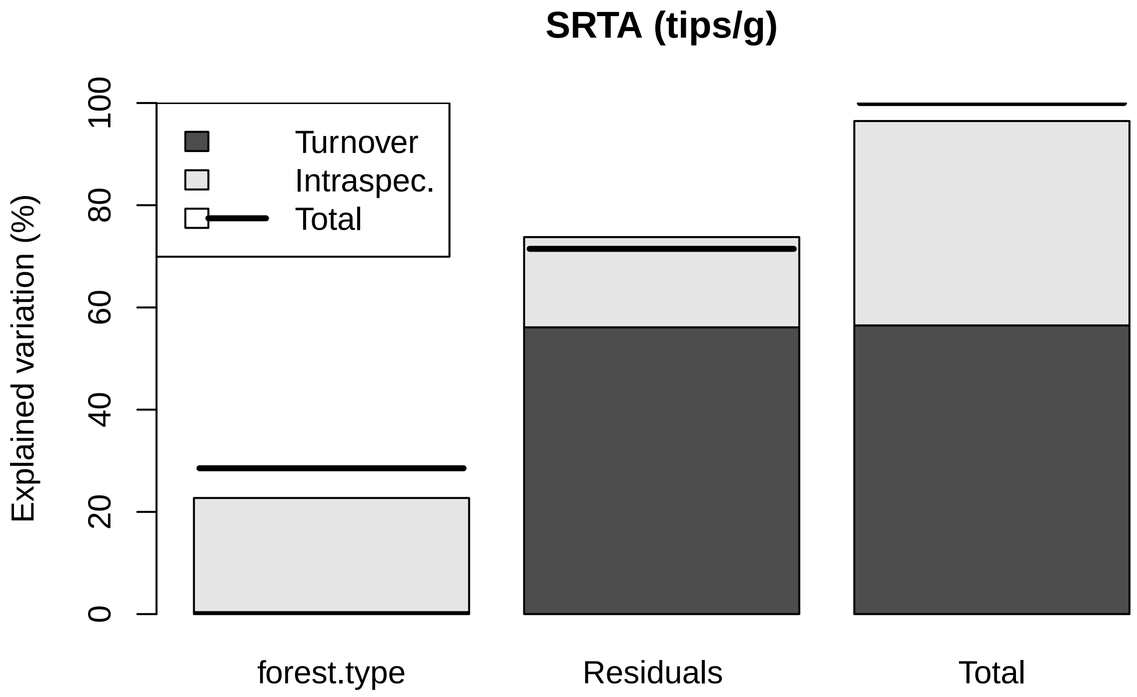


##### Root Length Per Volume

```
##
##  Decomposing trait sum of squares into composition turnover
##  effect, intraspecific trait variability, and their covariation:
##              Turnover Intraspec. Covariation Total
## forest.type    198.6      40316     -5659.3 34856
## Residuals    17729.3      16747     -8706.6 25769
## Total        17927.9      57063    -14365.9 60625
##
##  Relative contributions:
##              Turnover Intraspec. Covariation  Total
## forest.type 0.003276     0.6650    -0.09335 0.5749
## Residuals   0.292443     0.2762    -0.14361 0.4251
## Total       0.295719     0.9412    -0.23696 1.0000
##
##  Significance of testable effects:
##              Turnover Intraspec.      Total
## forest.type  0.17983 5.5479e-45 6.5956e-32
```

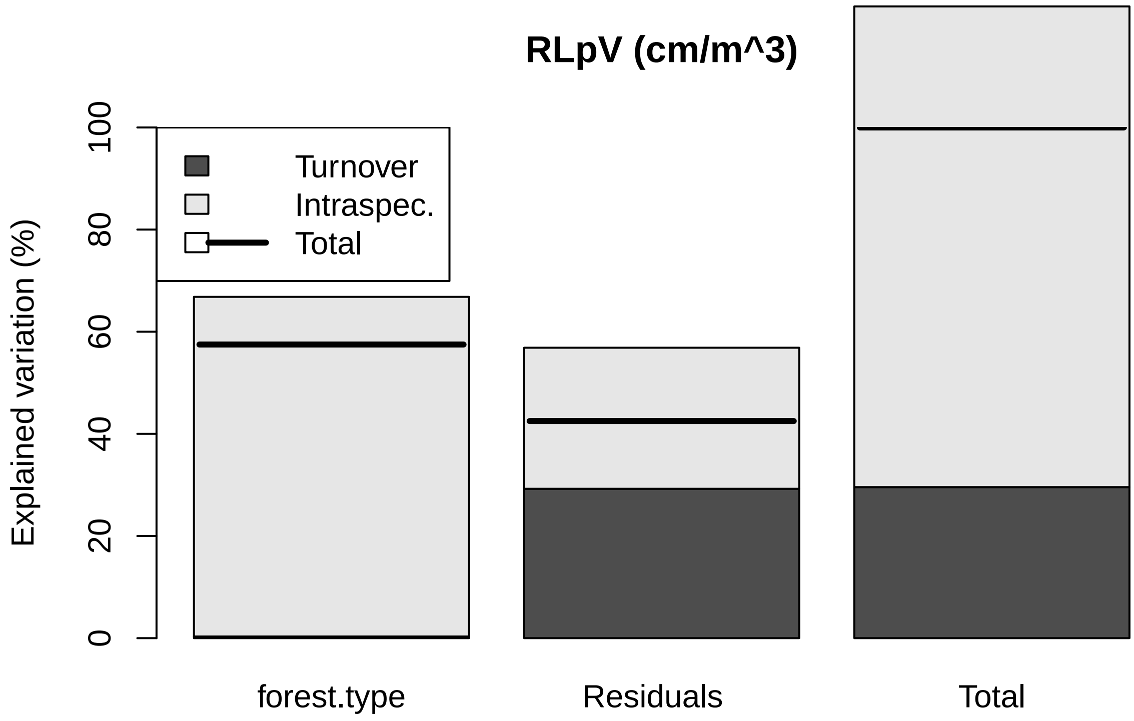


##### Root Tissue Density

```
##
##  Decomposing trait sum of squares into composition turnover
##  effect, intraspecific trait variability, and their covariation:
##             Turnover Intraspec. Covariation    Total
## forest.type  0.01585    0.10143    -0.08019 0.037088
## Residuals    0.52233    0.19034     0.21271 0.925385
## Total        0.53818    0.29177     0.13252 0.962473
##
##  Relative contributions:
##             Turnover Intraspec. Covariation   Total
## forest.type  0.01647     0.1054    -0.08332 0.03853
## Residuals    0.54270     0.1978     0.22101 0.96147
## Total        0.55917     0.3031     0.13769 1.00000
##
##  Significance of testable effects:
##             Turnover Intraspec.    Total
## forest.type 0.028005 9.8843e-17 0.011762
```

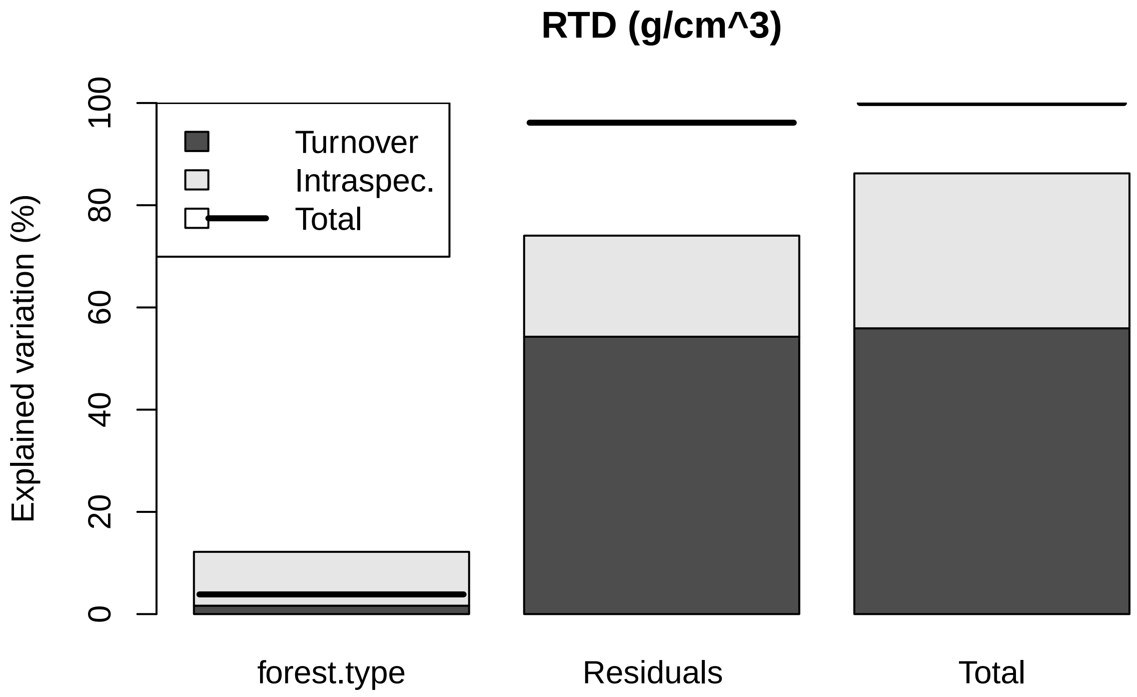


##### Root Branchiness

```
##
##  Decomposing trait sum of squares into composition turnover
##  effect, intraspecific trait variability, and their covariation:
##              Turnover Intraspec. Covariation   Total
## forest.type 0.0089921     5.1527     0.43050  5.5922
## Residuals   1.9887151     3.0355     0.75499  5.7792
## Total       1.9977072     8.1882     1.18549 11.3714
##
##  Relative contributions:
##              Turnover Intraspec. Covariation  Total
## forest.type 0.0007908     0.4531     0.03786 0.4918
## Residuals   0.1748878     0.2669     0.06639 0.5082
## Total       0.1756785     0.7201     0.10425 1.0000
##
##  Significance of testable effects:
##             Turnover Intraspec.      Total
## forest.type  0.39334   9.72e-37 1.3746e-25
```

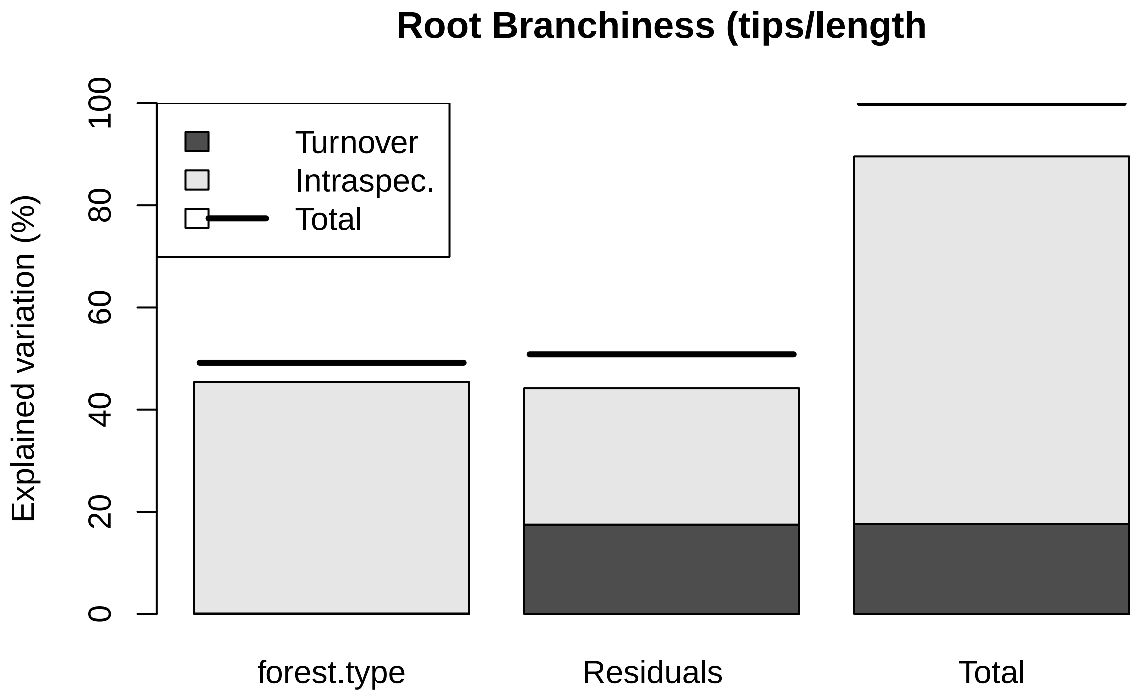
Root Branchiness (tips/length
Explained variation (%)
0
20
40
60
80
100
Turnover
Intraspec.
Total
forest.type
Residuals
Total

### PLOTTING weighted averages

#### Leaf Mass

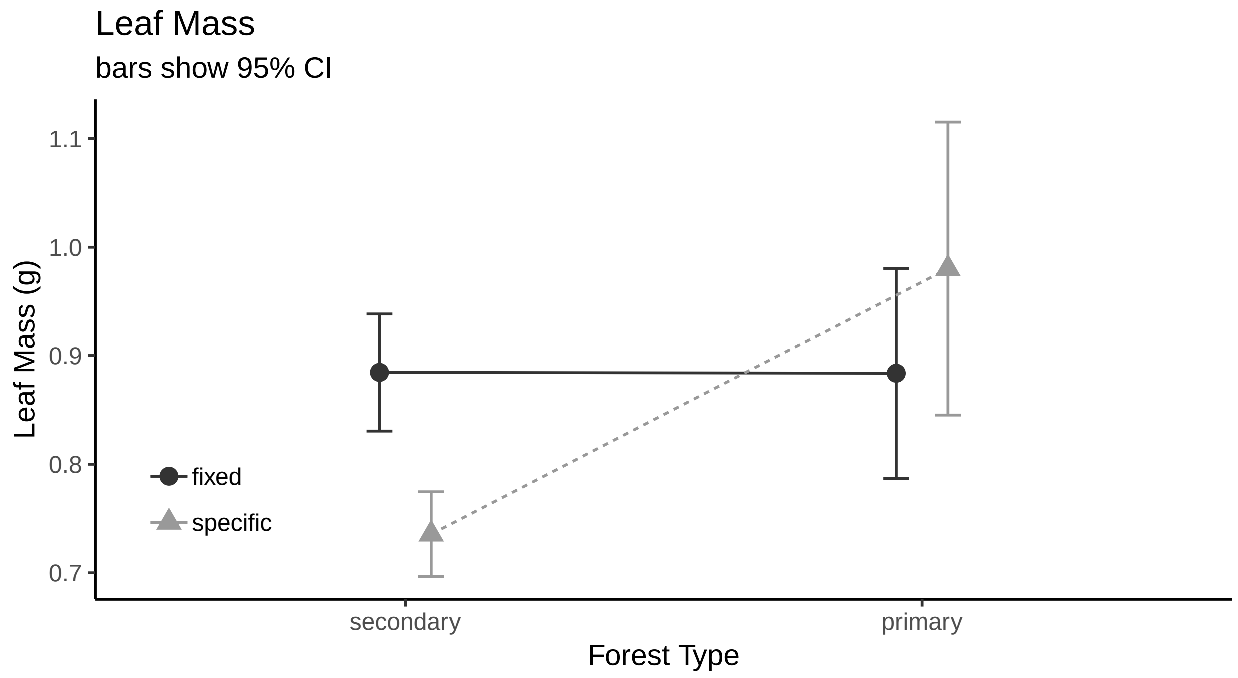


##### histogram

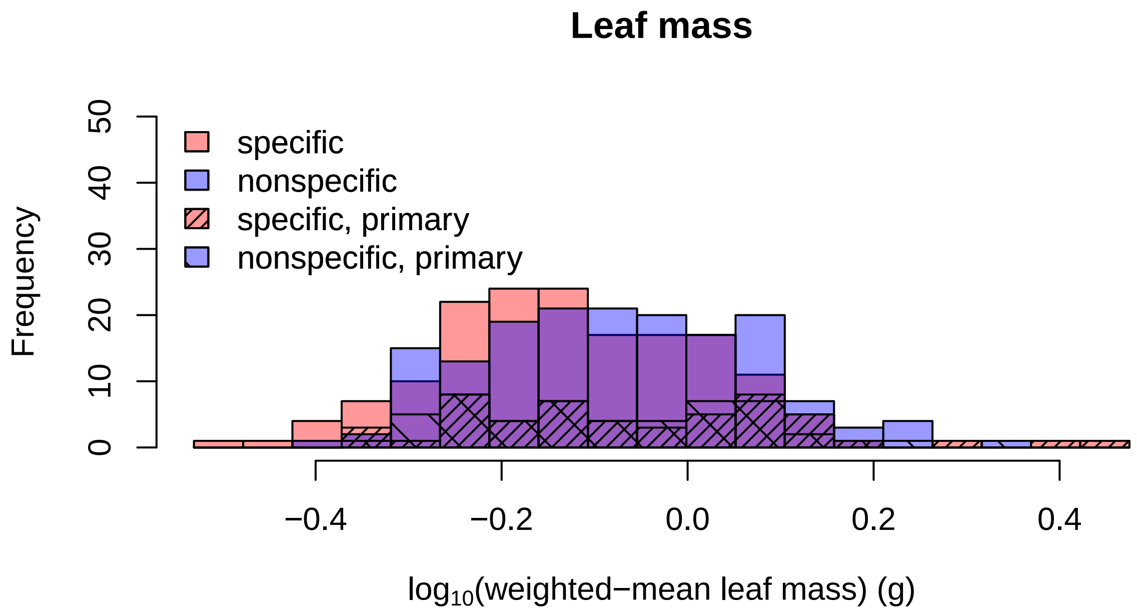

##### Leaf Area

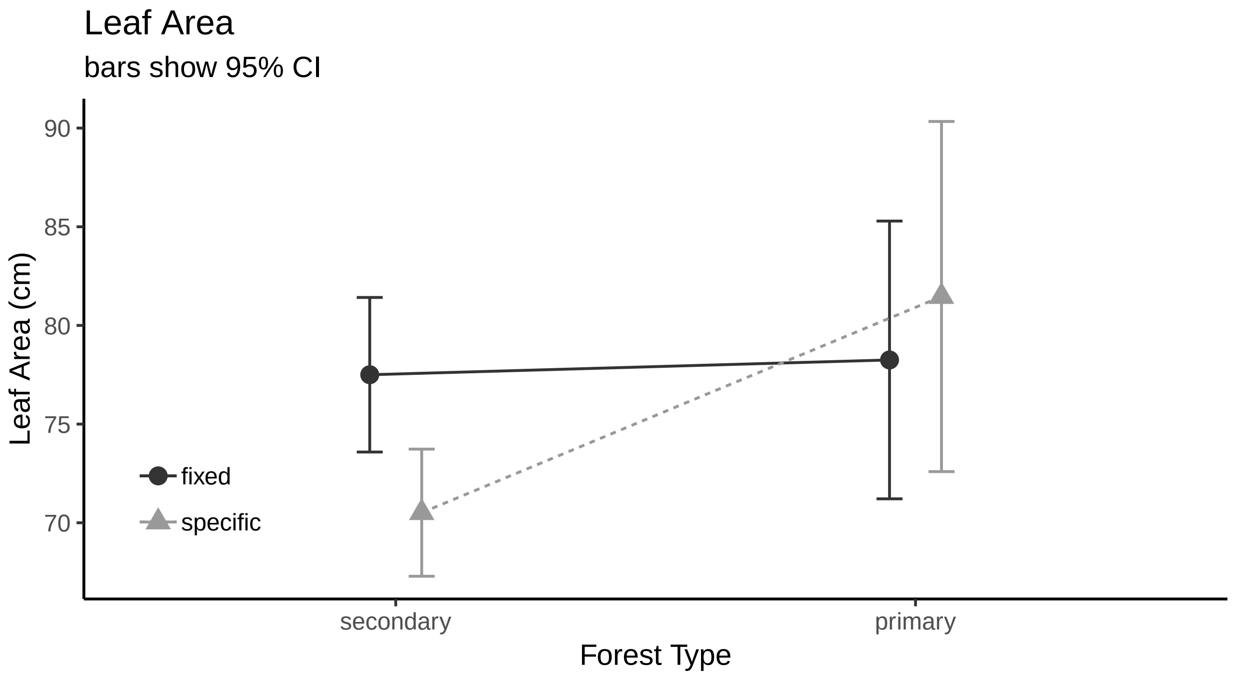


###### histogram

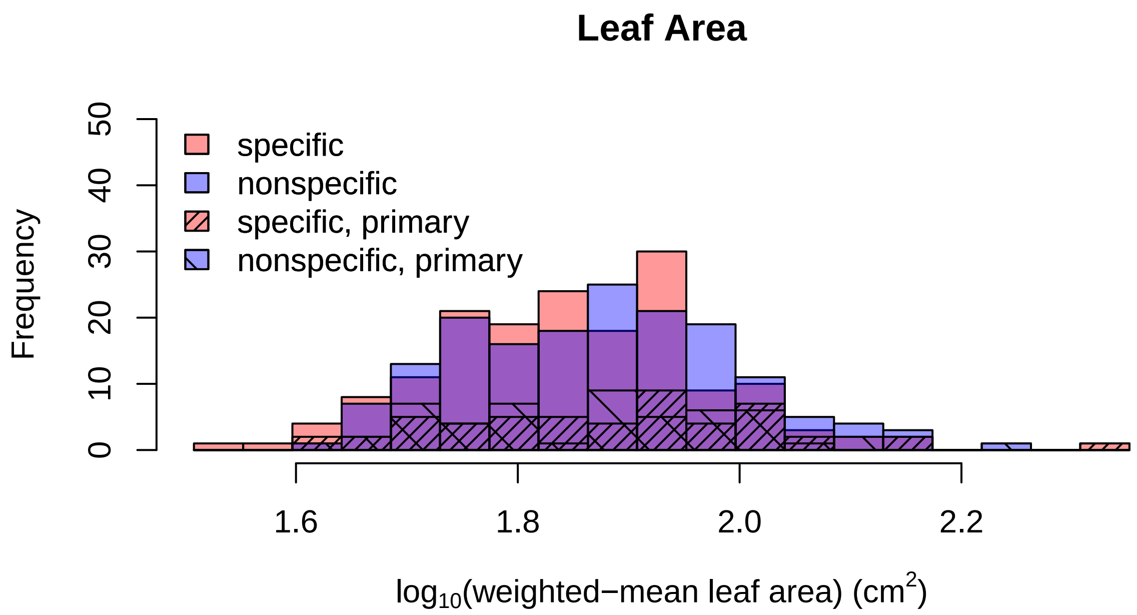

##### SLA

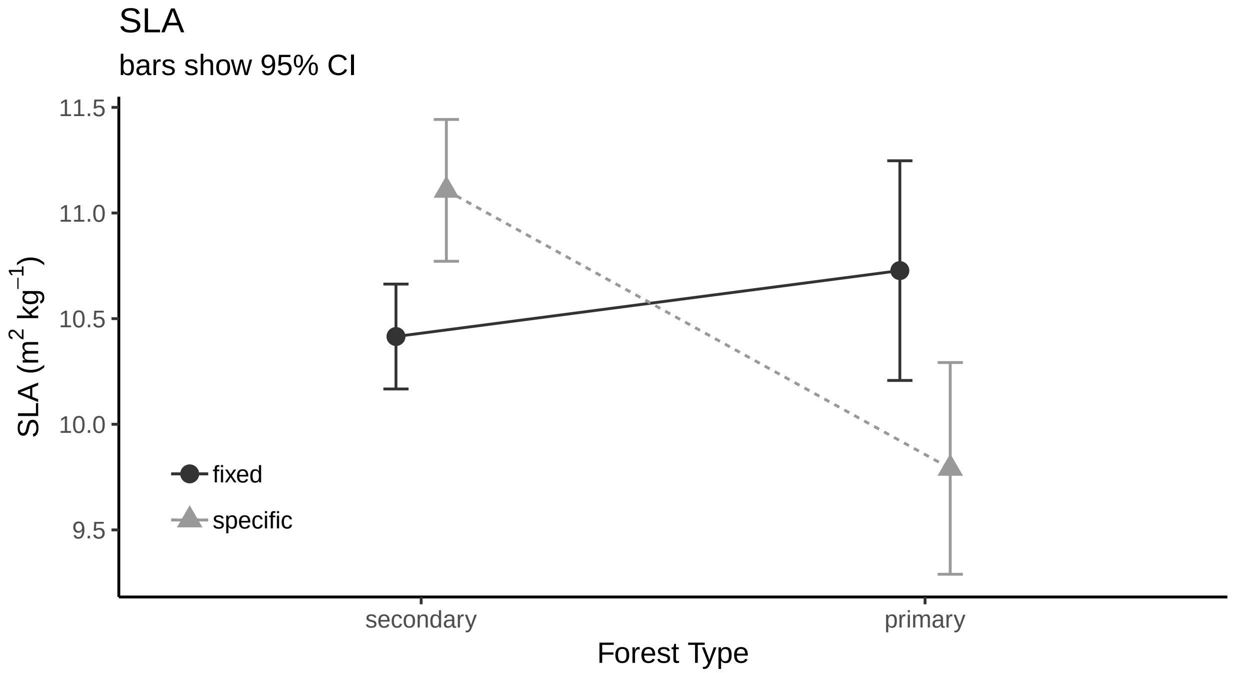


###### histogram

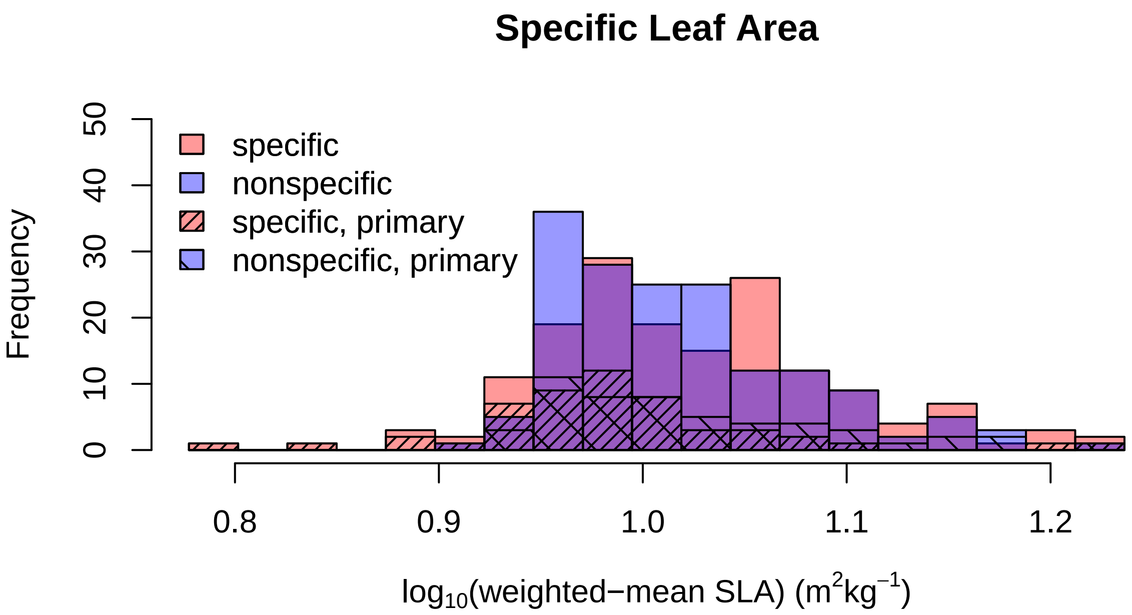

##### Leaf Perimeter

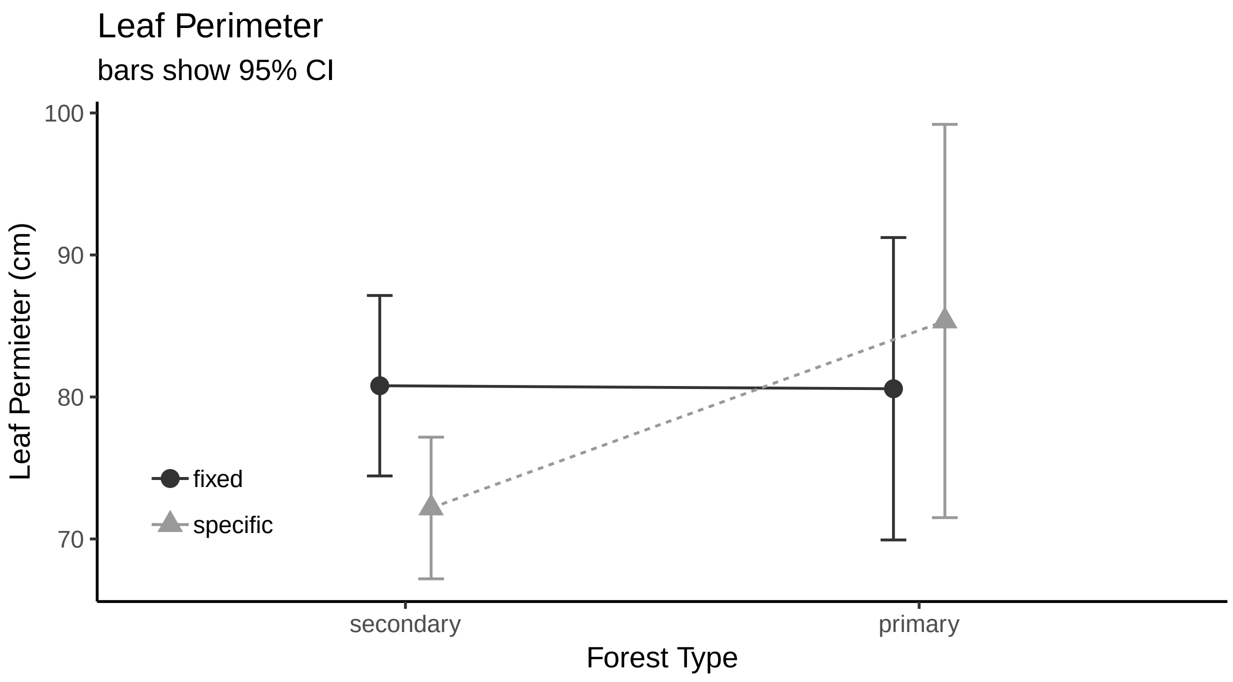


###### histogram

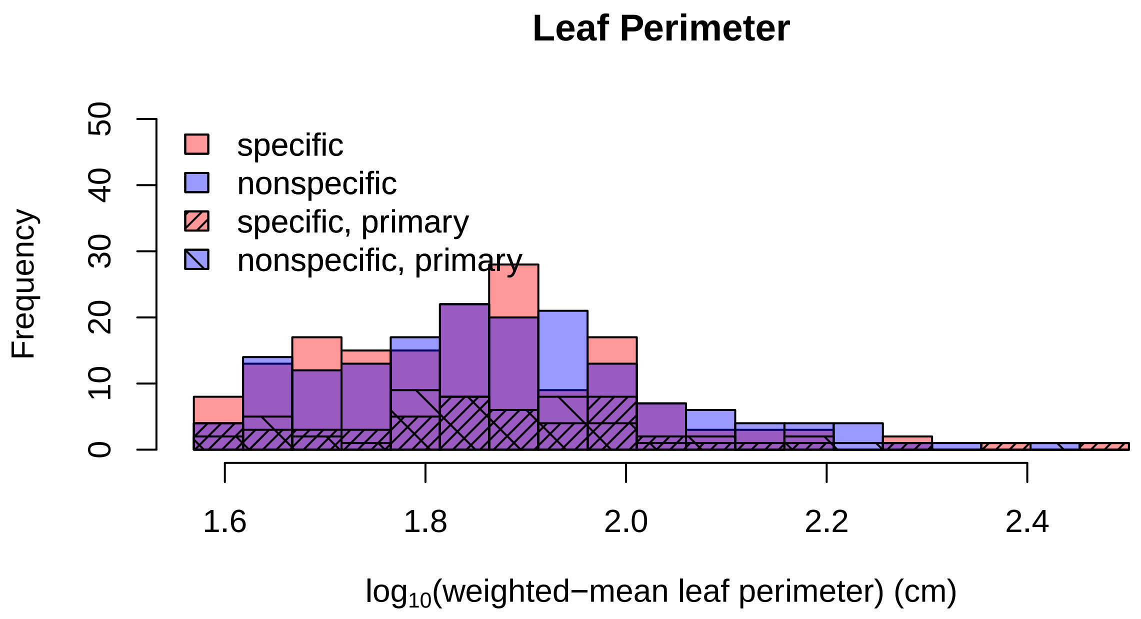

##### Leaf Width

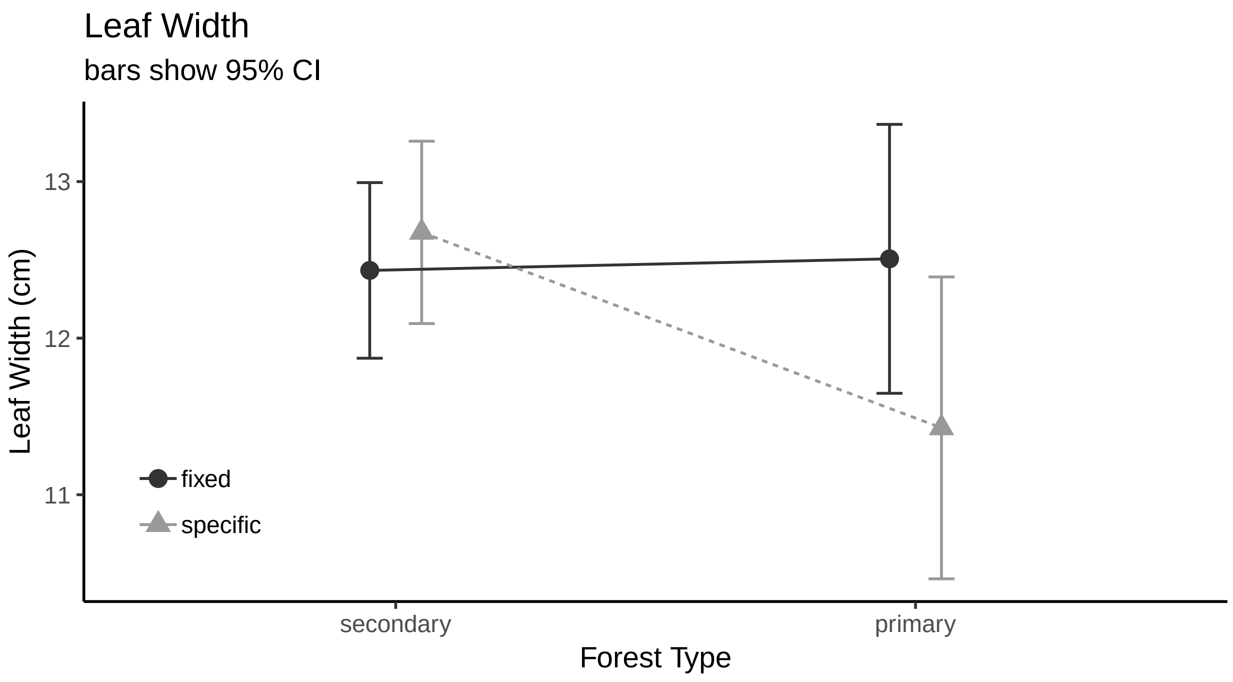


###### histogram

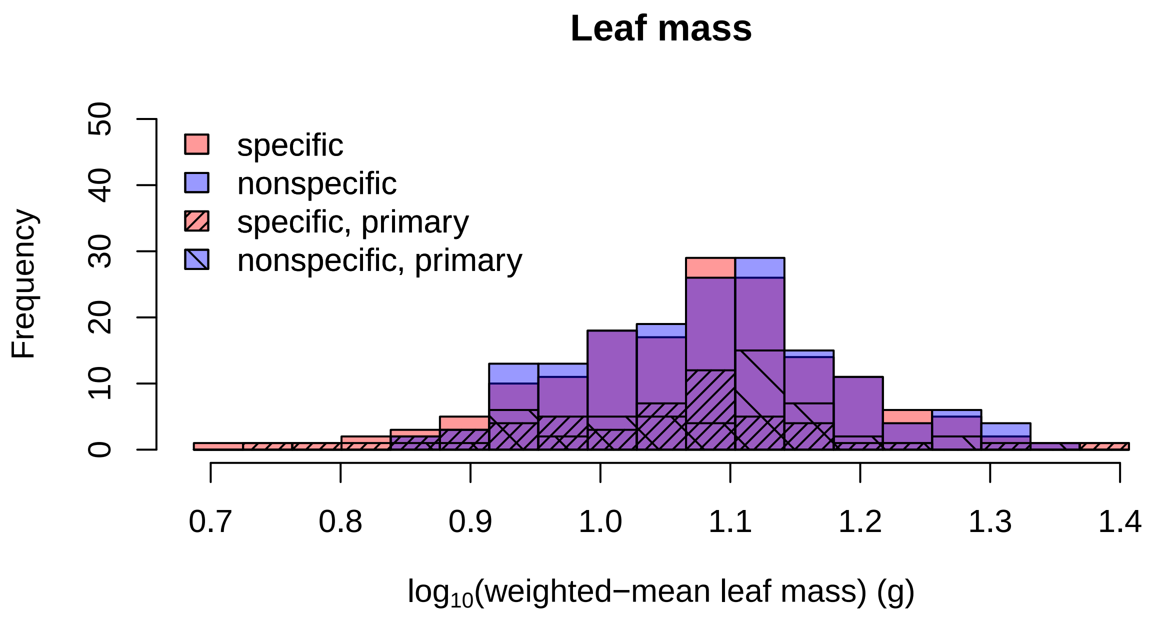

##### Leaf Height

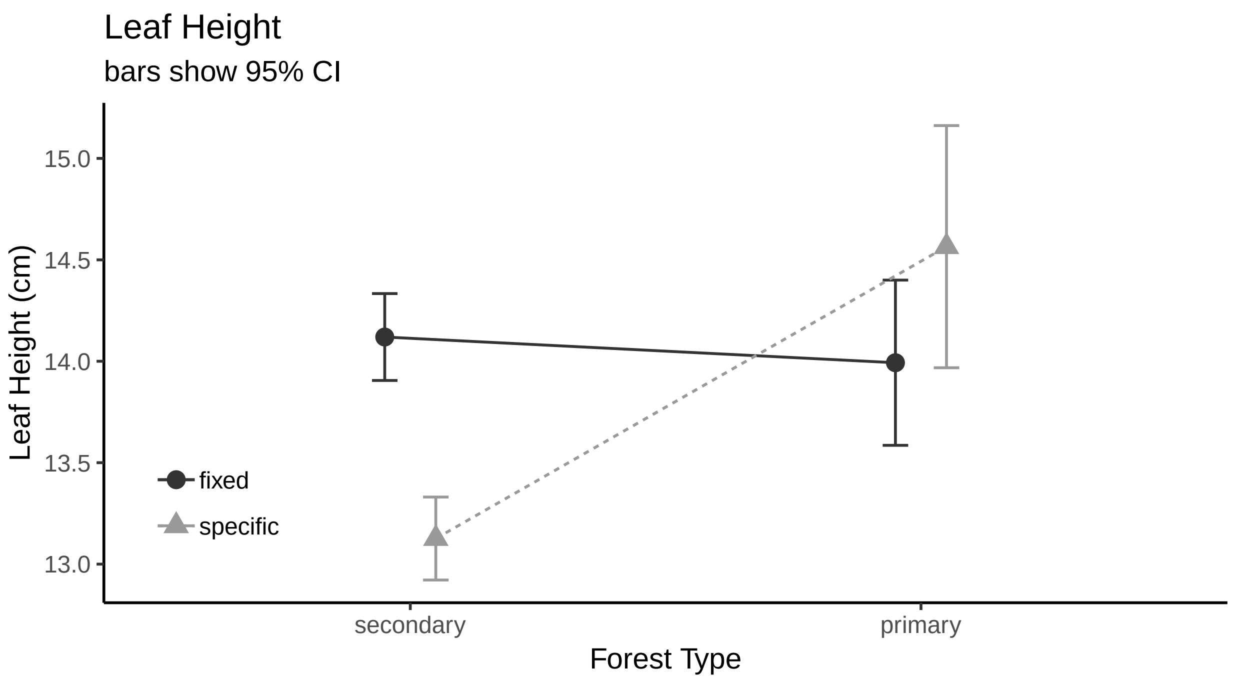


##### histogram

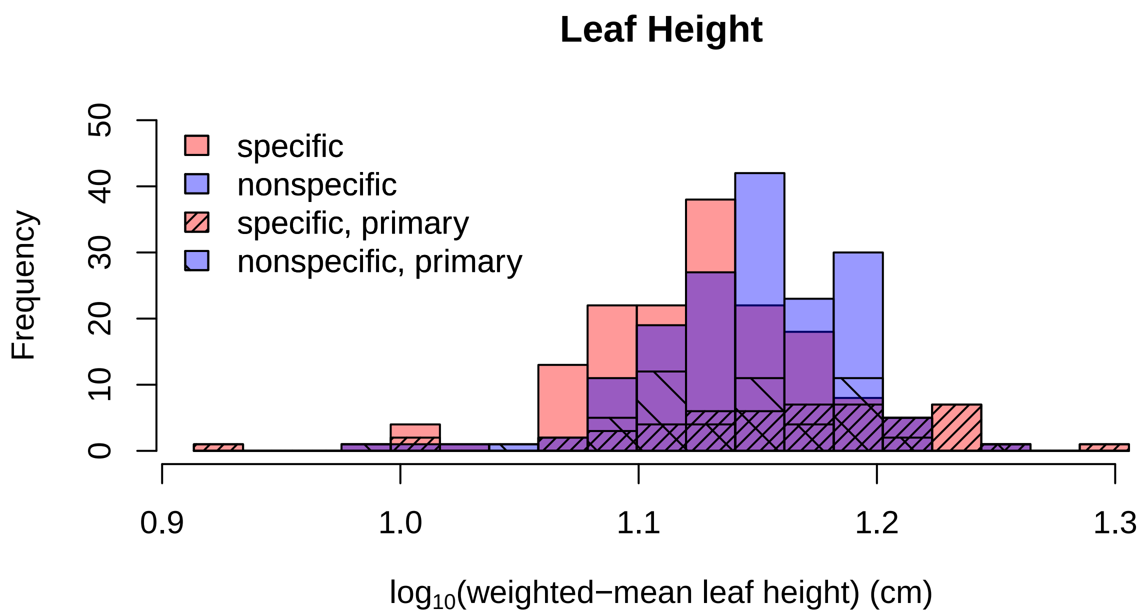

##### Leaf Thickness

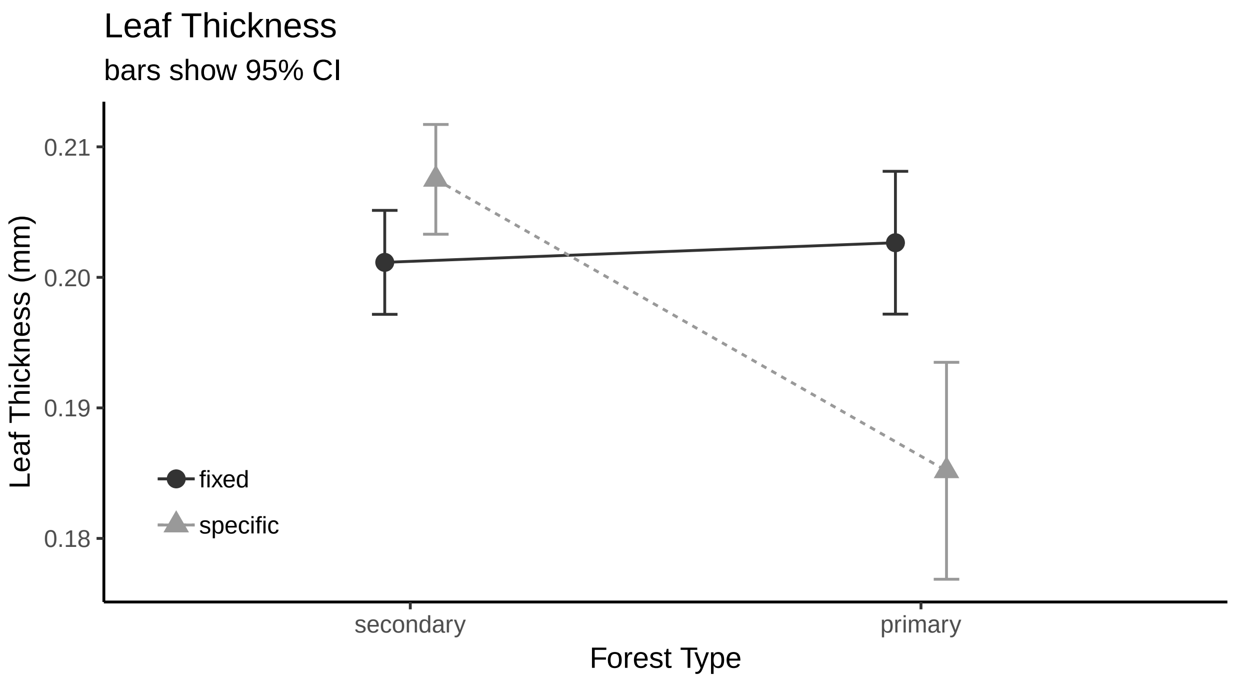


###### histogram

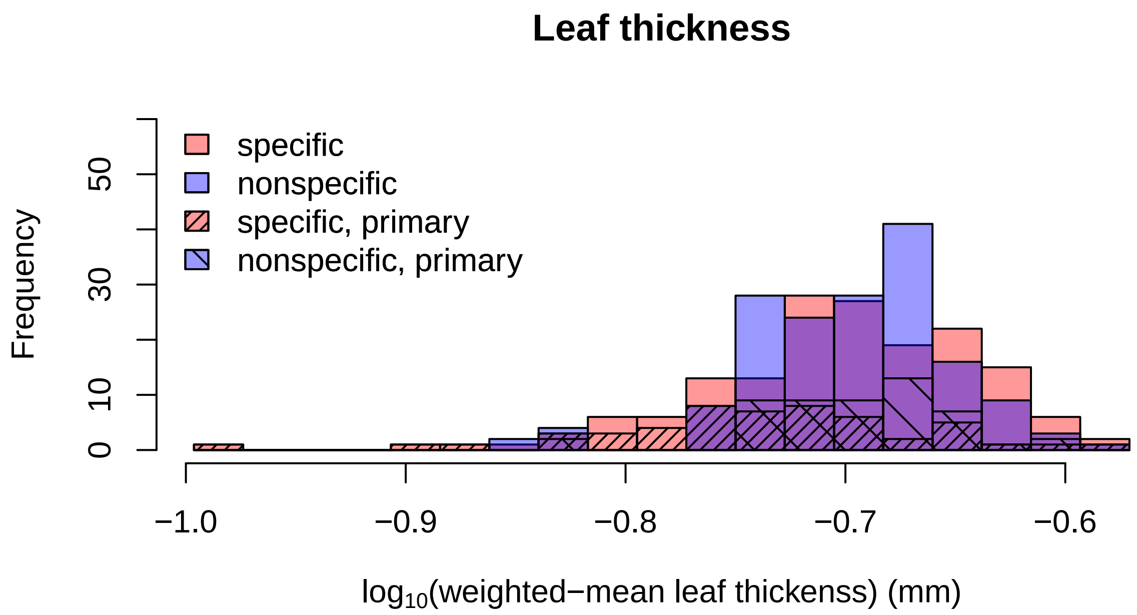

##### Root Mass

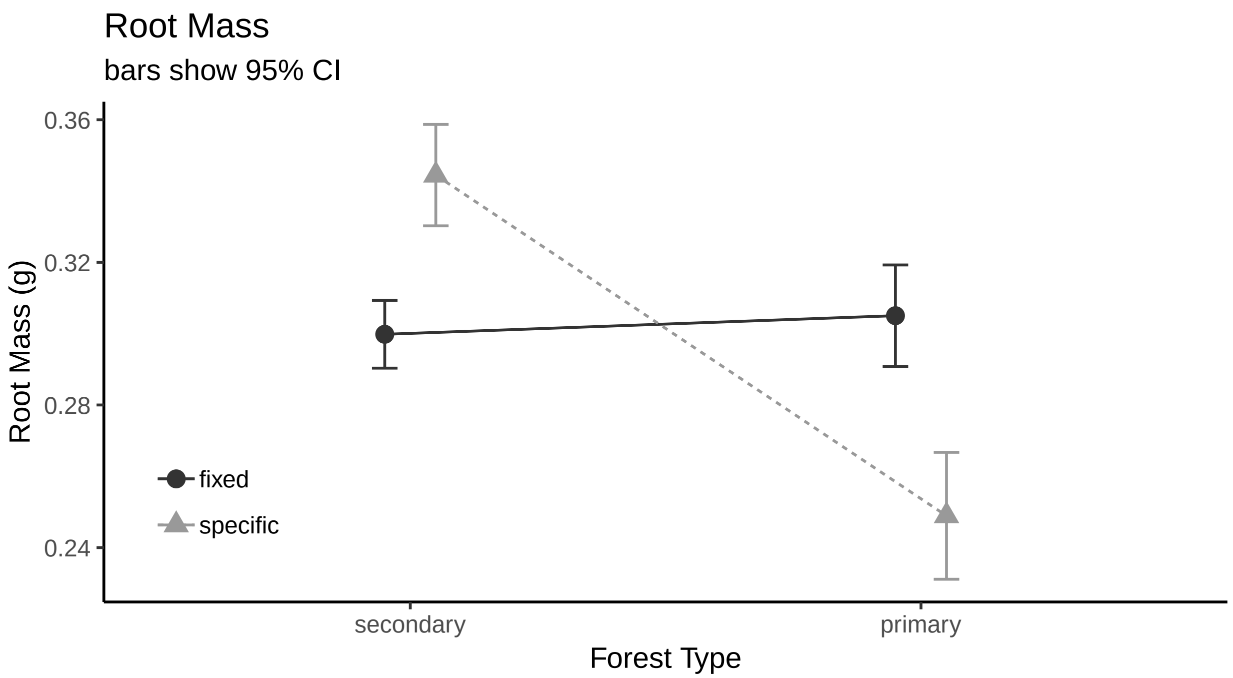


###### histogram

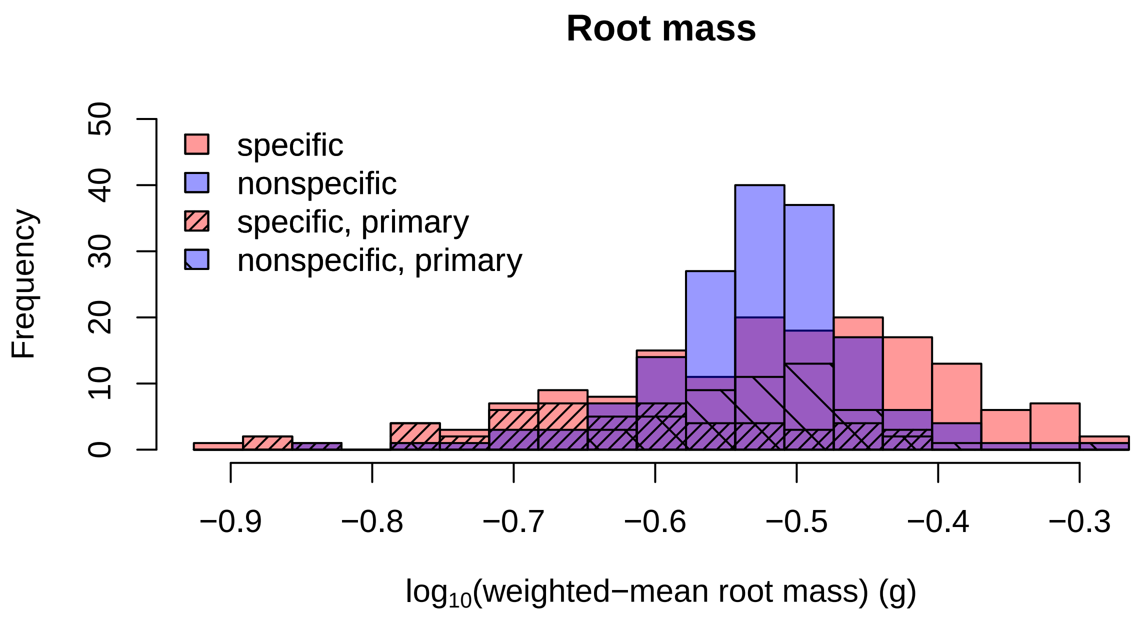

**Avg. Root Diameter**

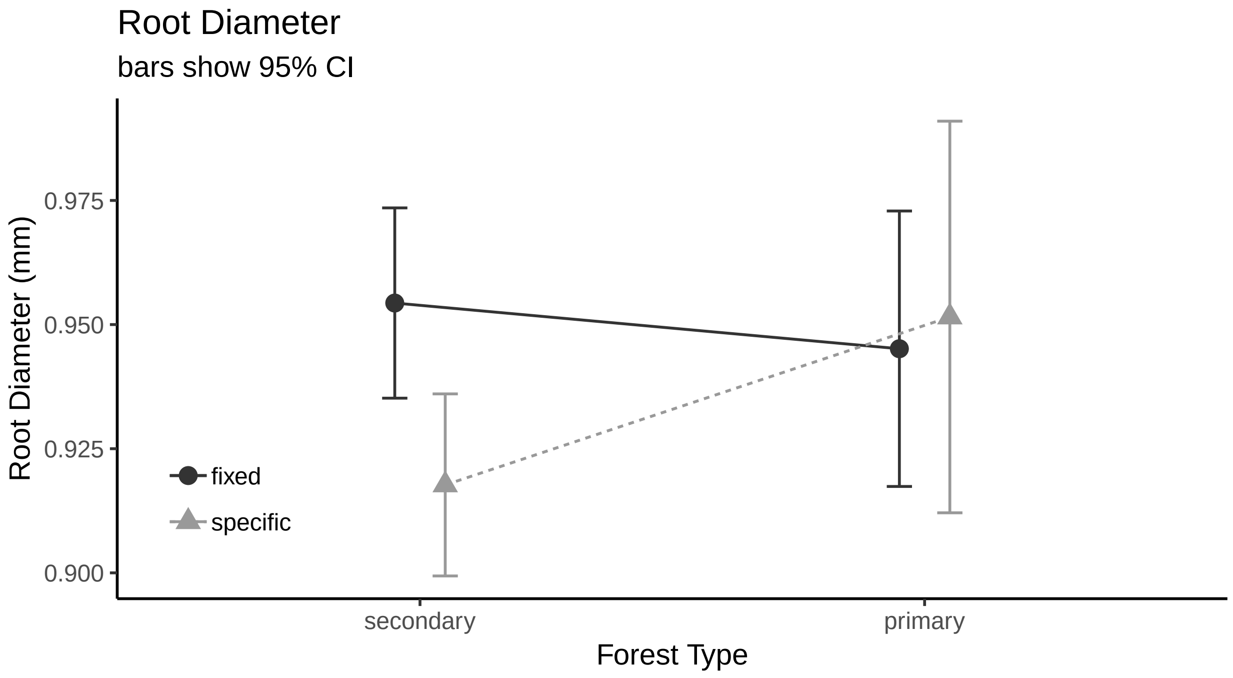


**histogram**

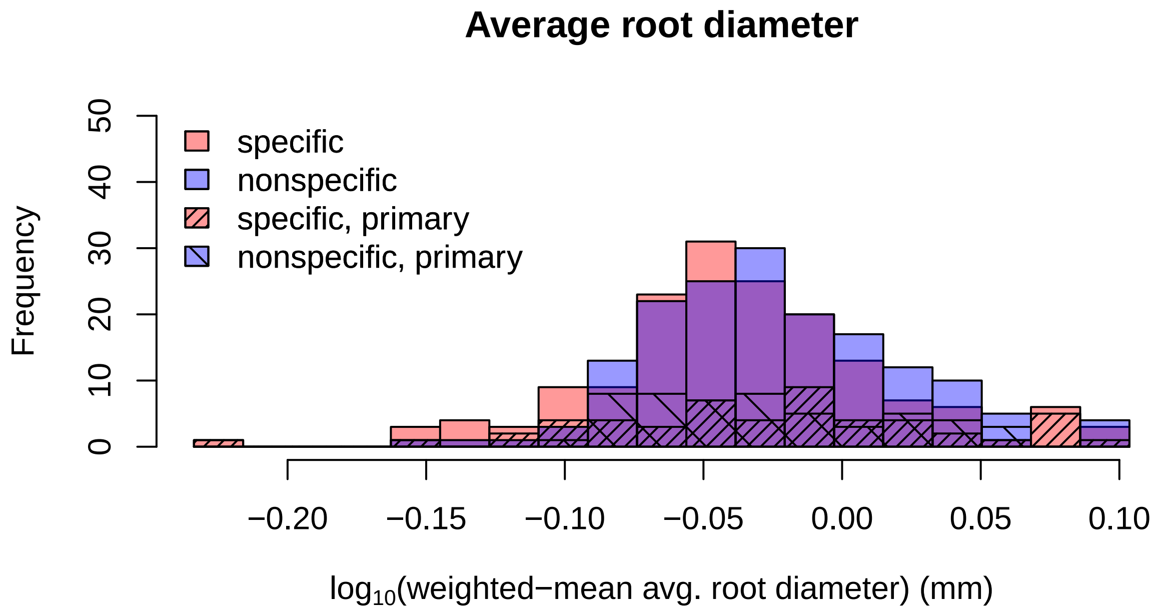

**Number of Root Tips**

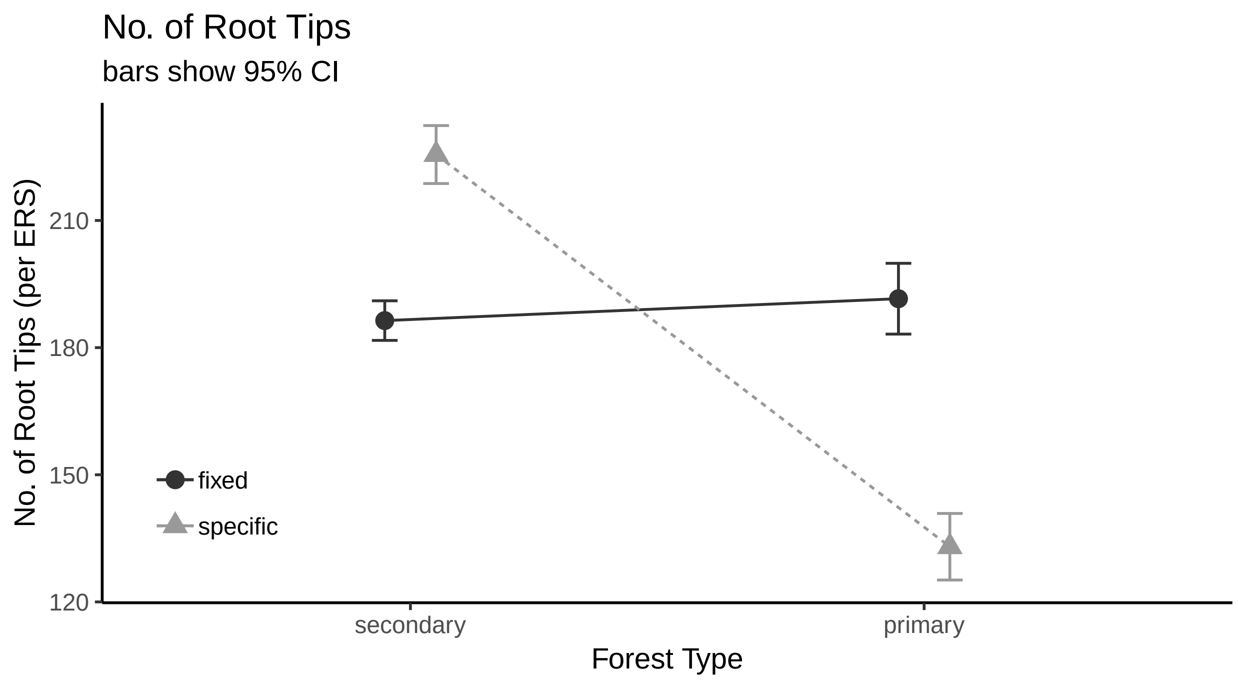


**histogram**

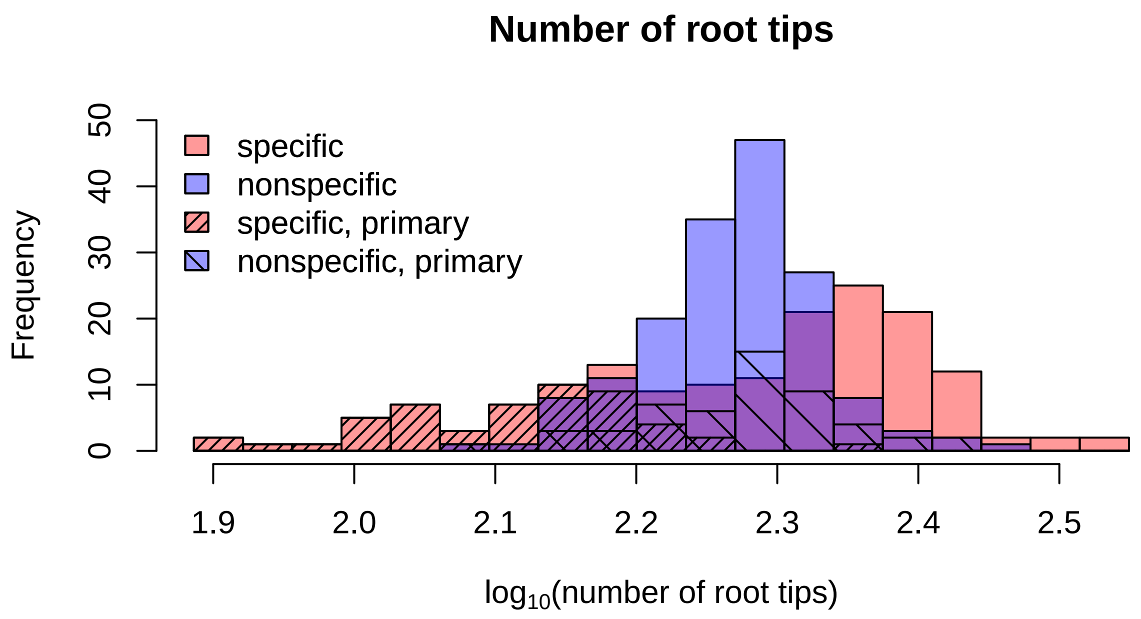

##### Number of Root Forks

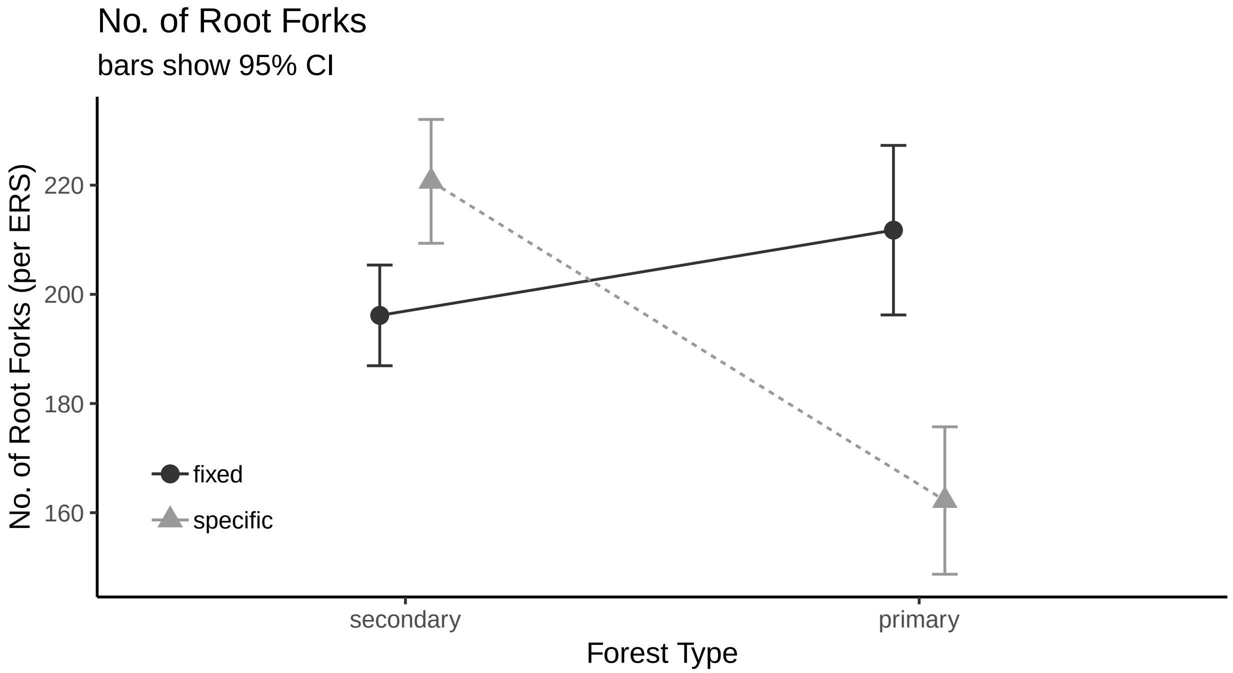


###### histogram

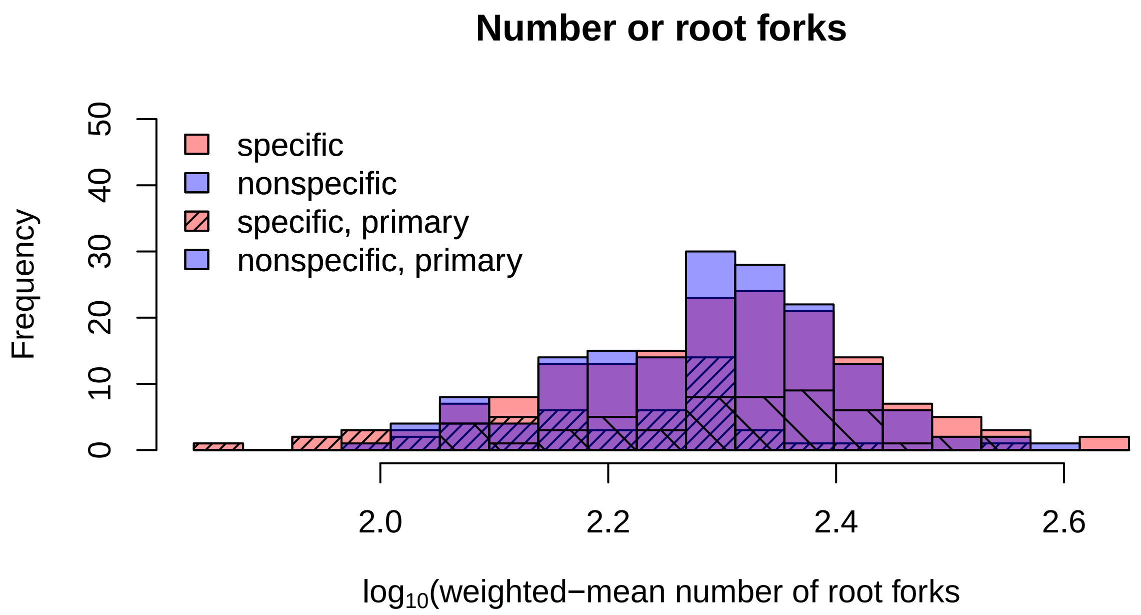

##### Root Area

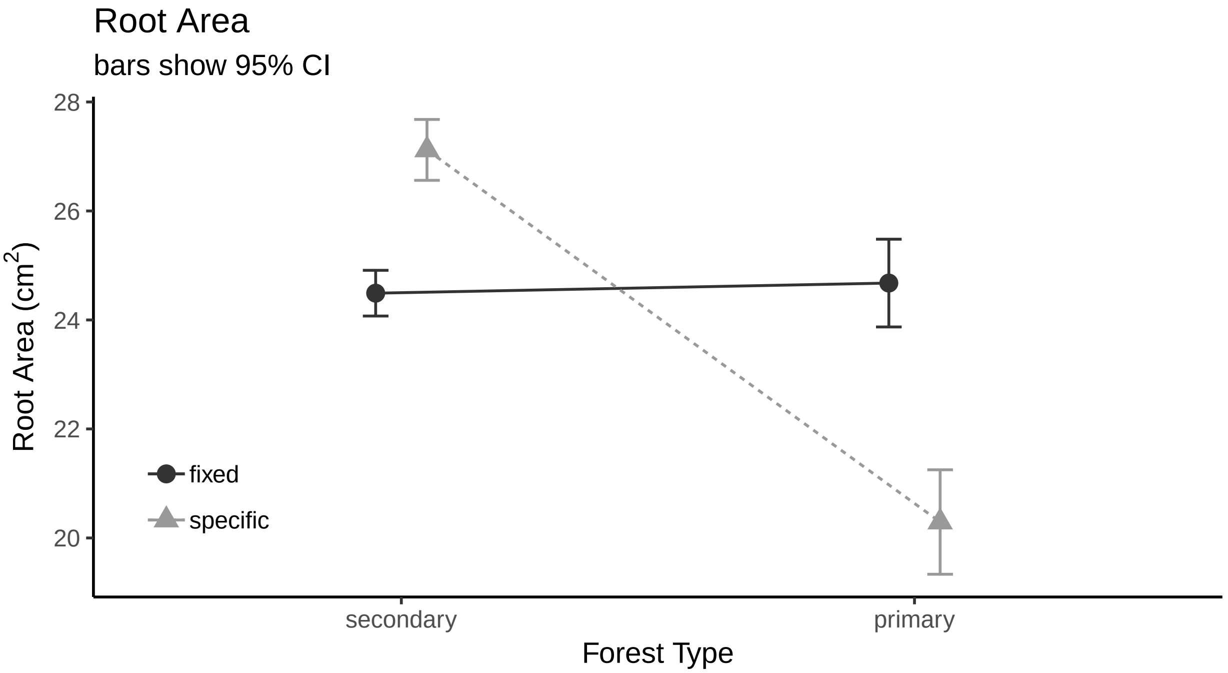


###### histogram

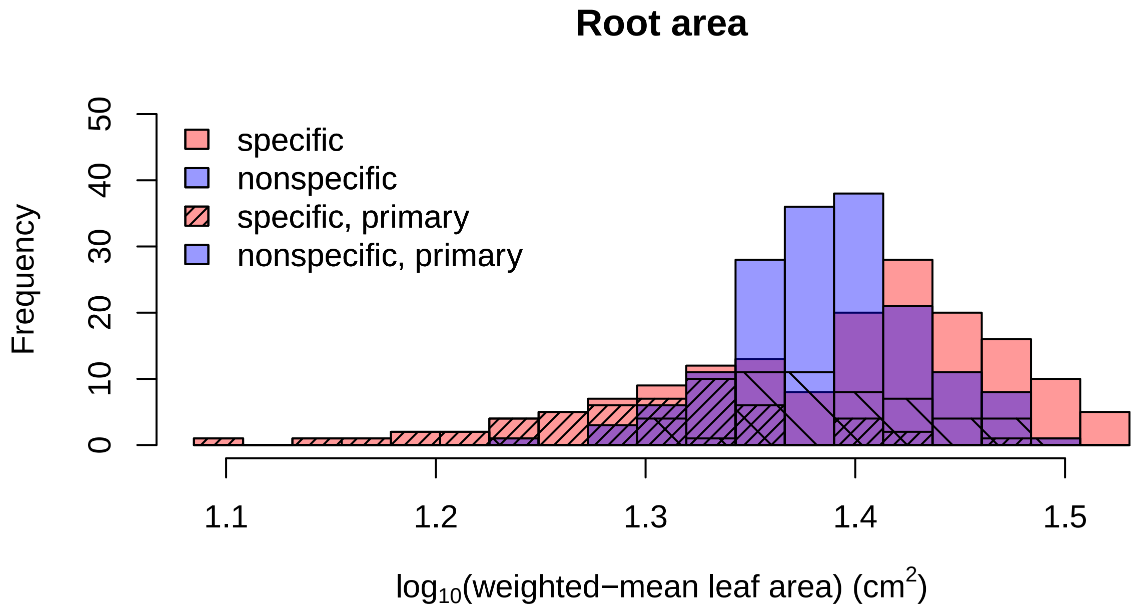

##### Root Length

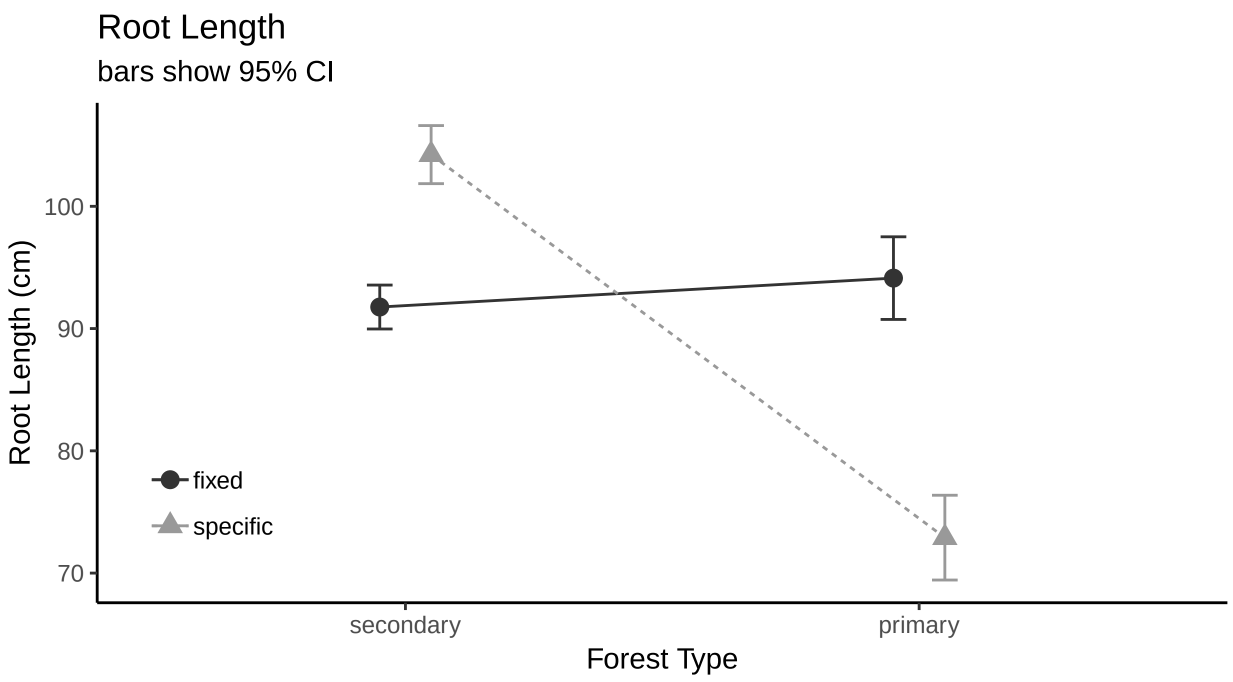


##### histogram

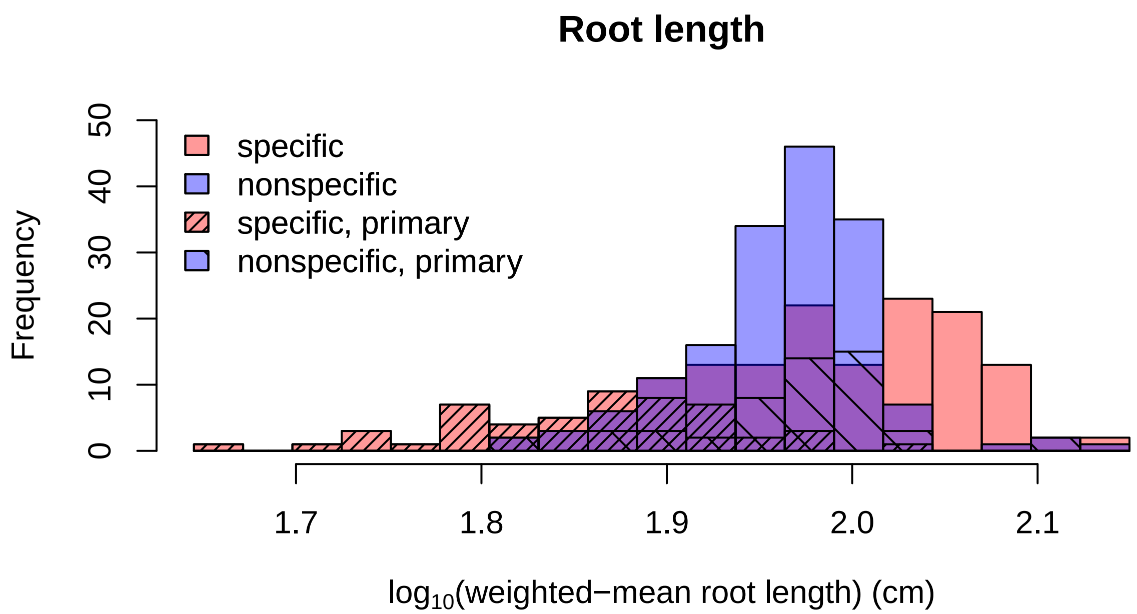

#### Root Volumne

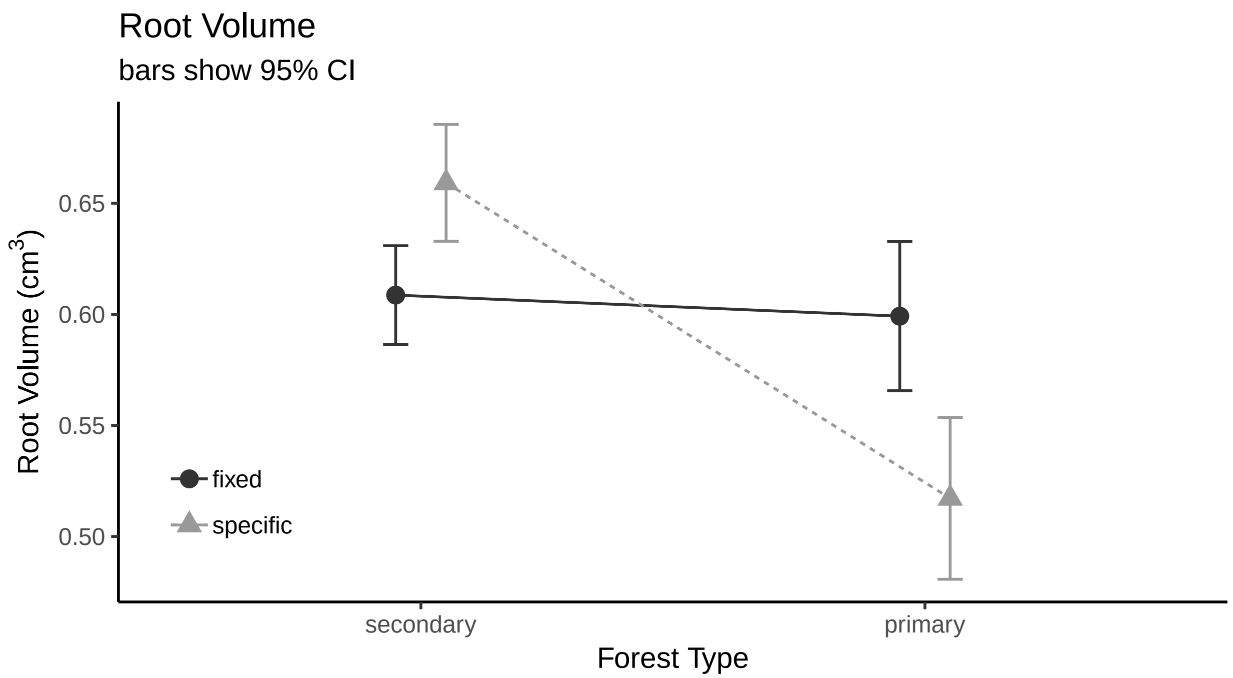


#### histogram

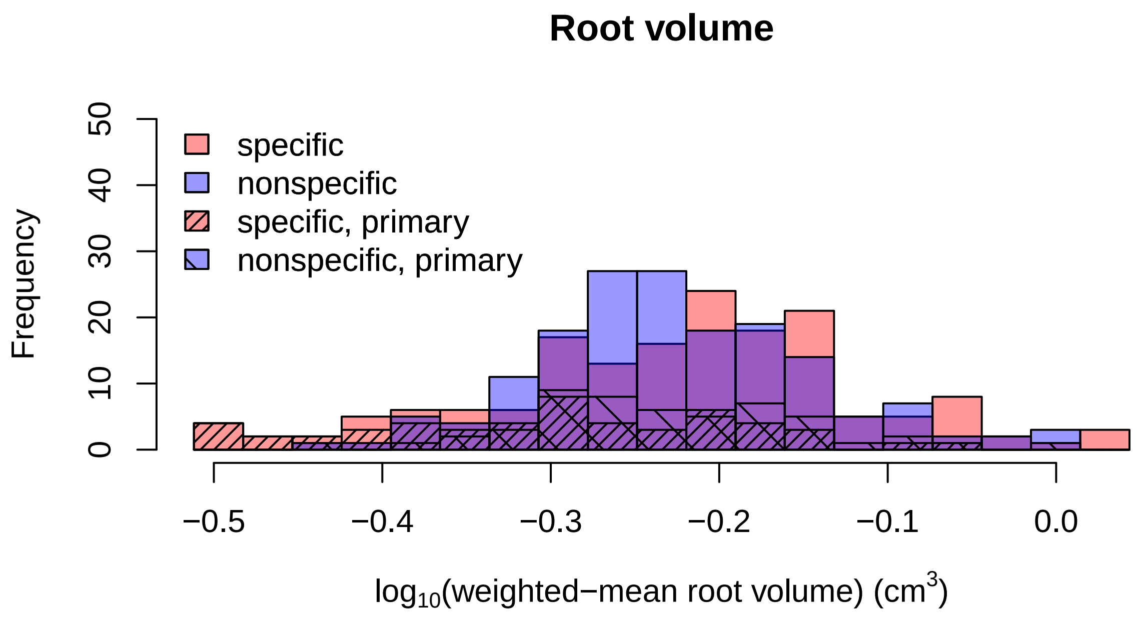

#### SRL

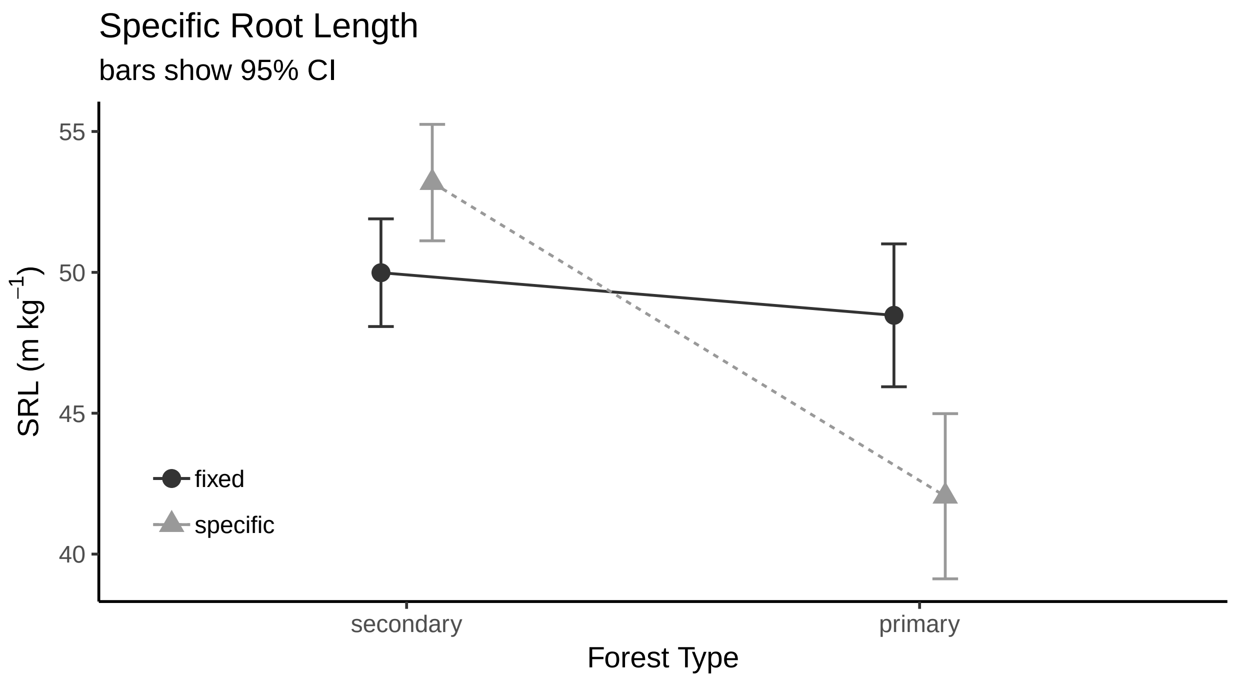


##### histogram

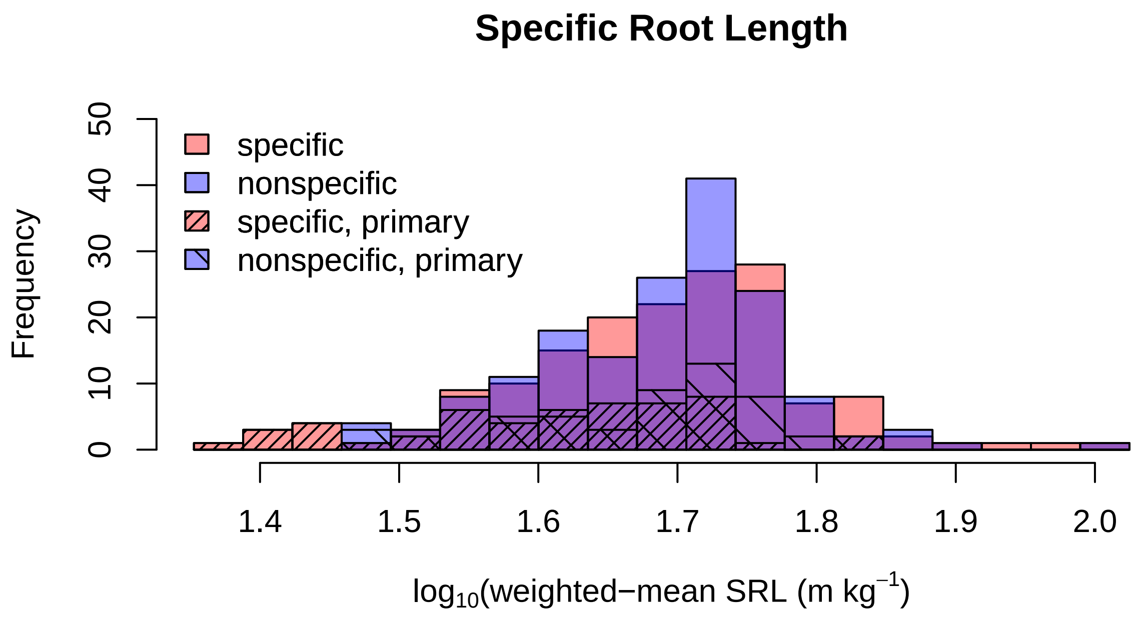

**SRA**

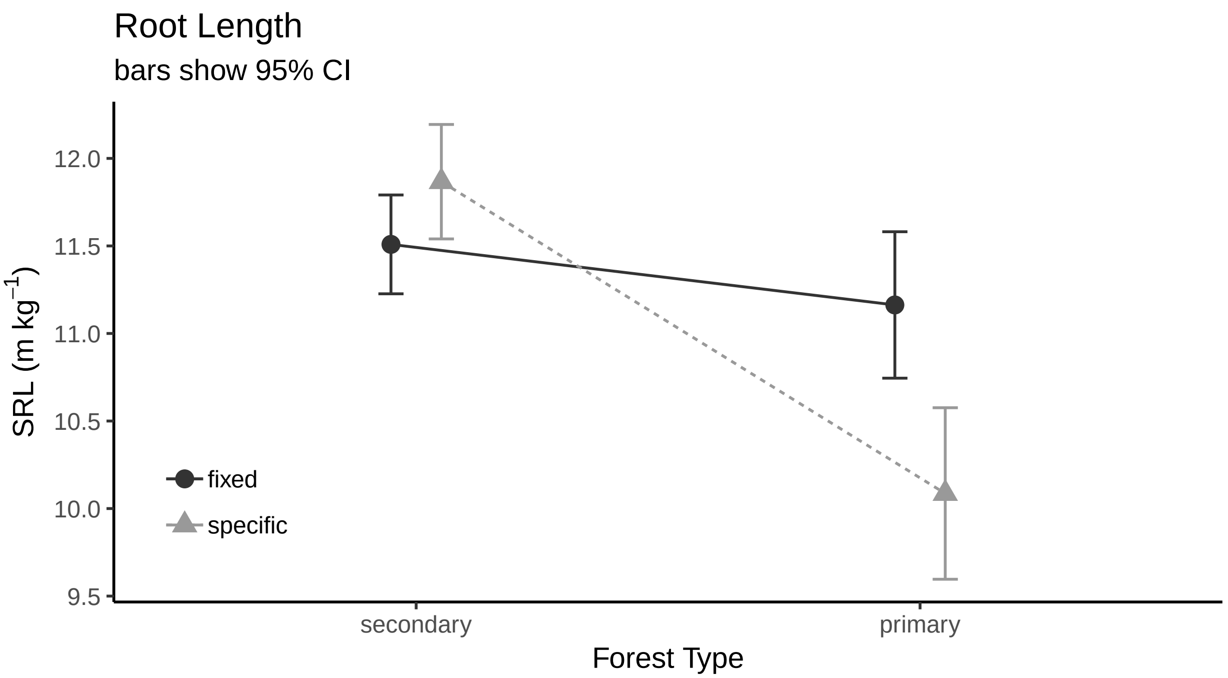


**histogram**

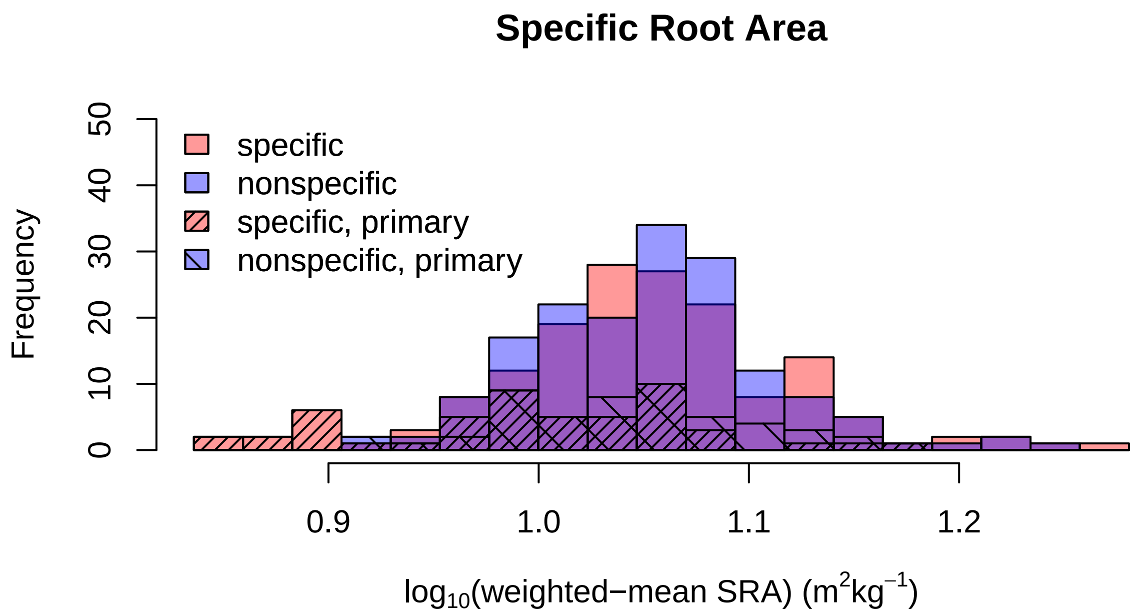

**Specific Root Tip Abundance**

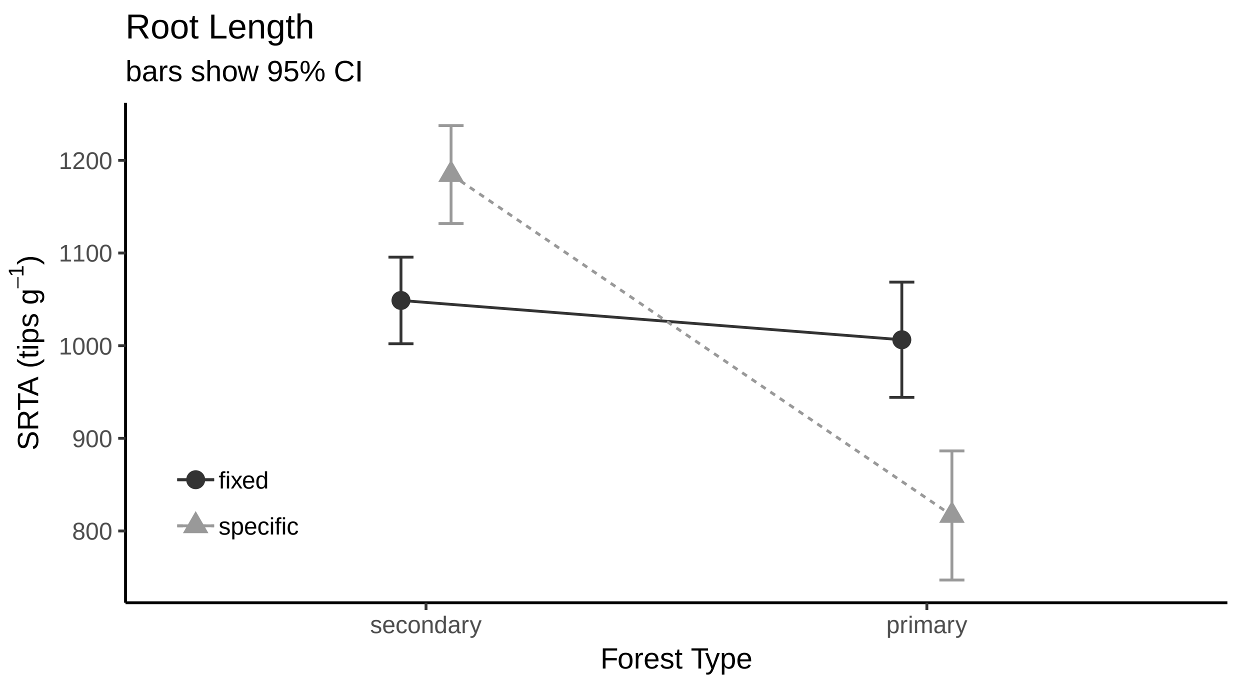


**histogram**

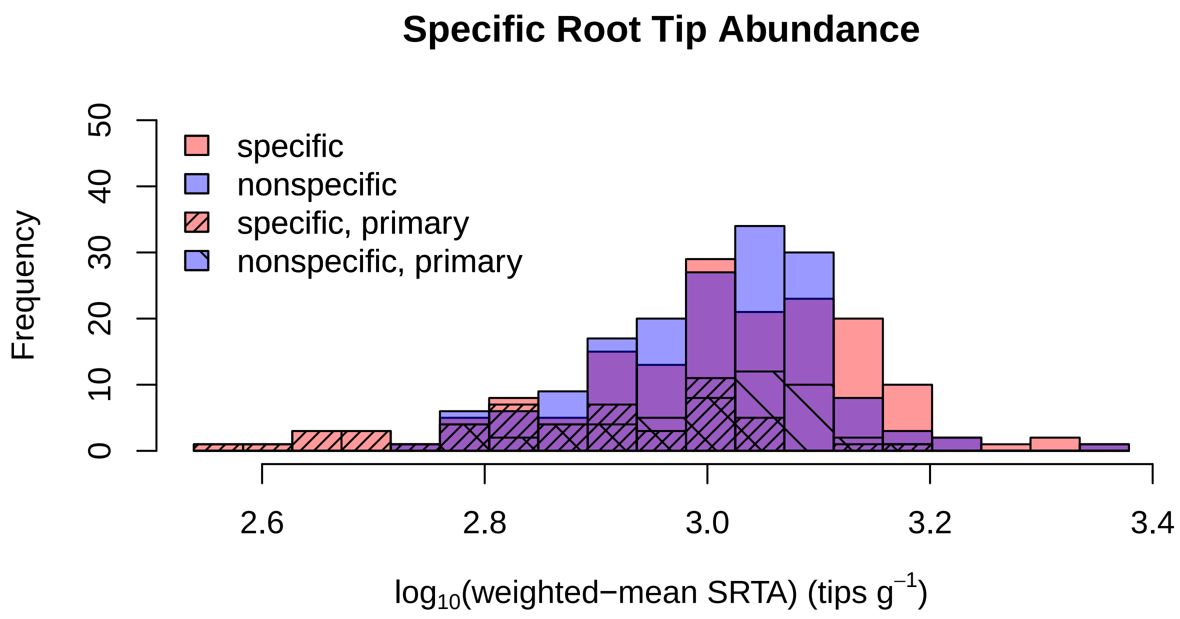

##### Root Length Per Volume

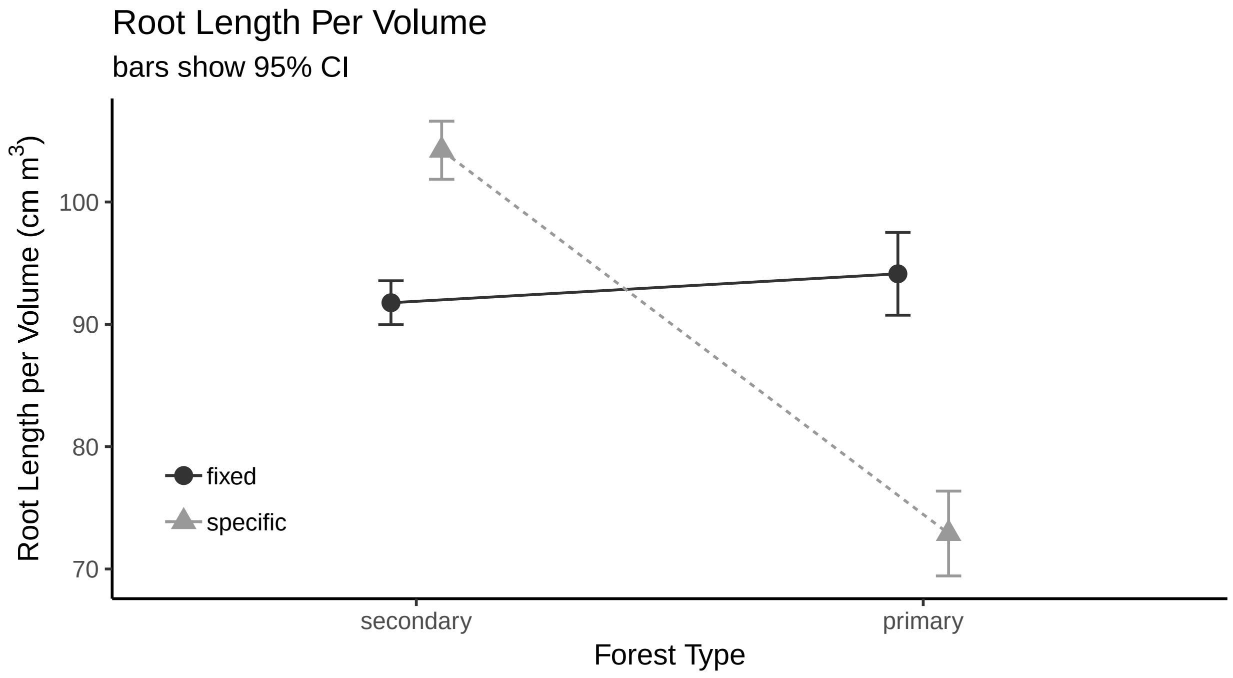


##### histogram

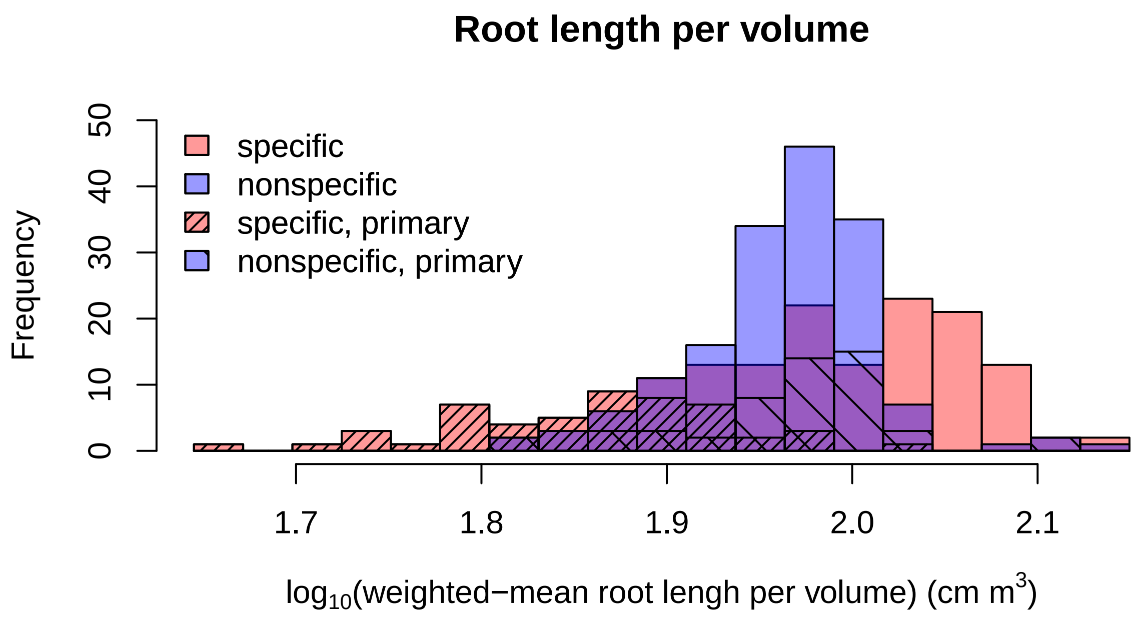

##### Root Tissue Density

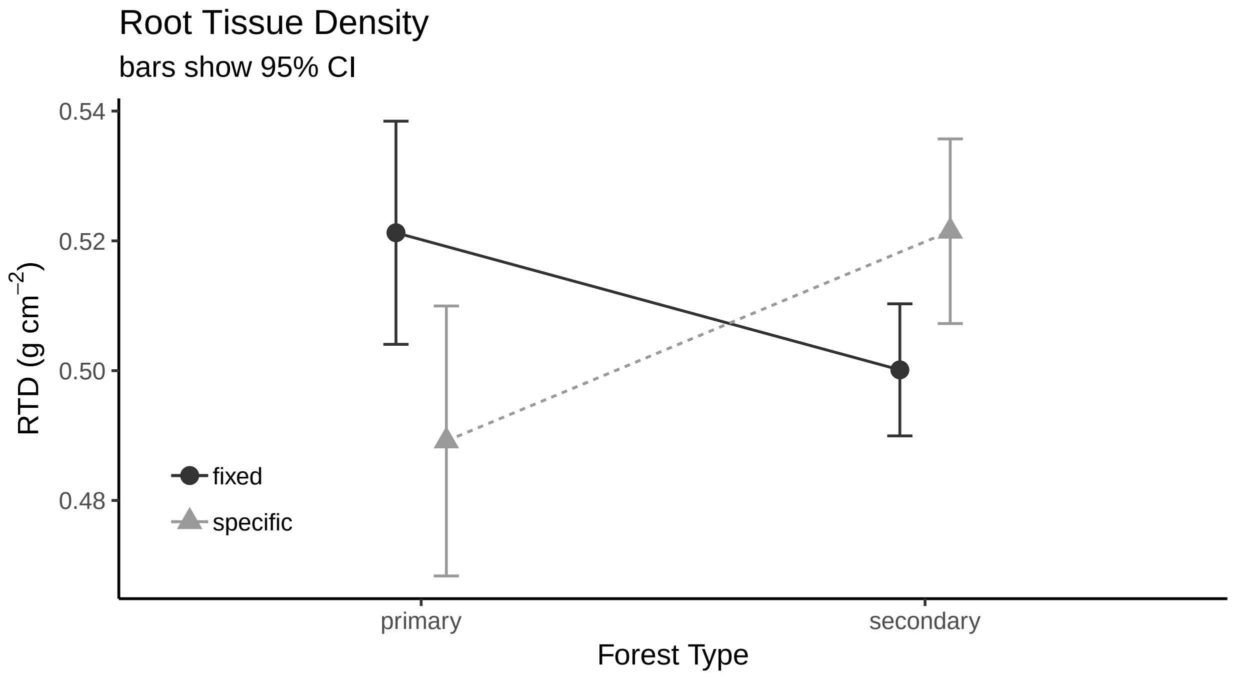


###### histogram

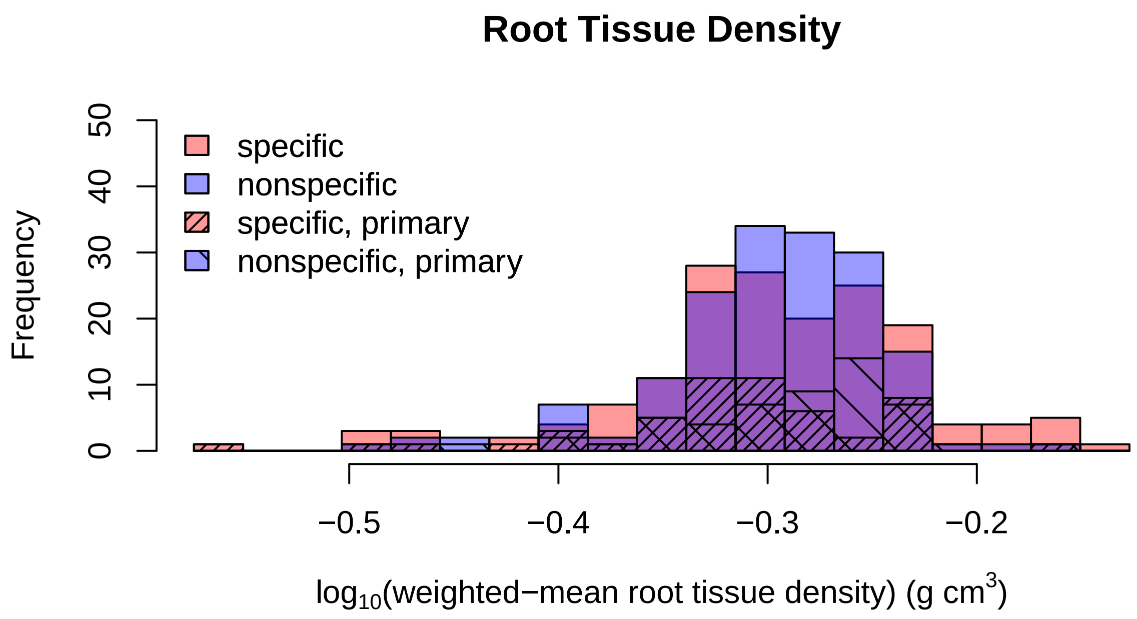

##### Root Branchiness

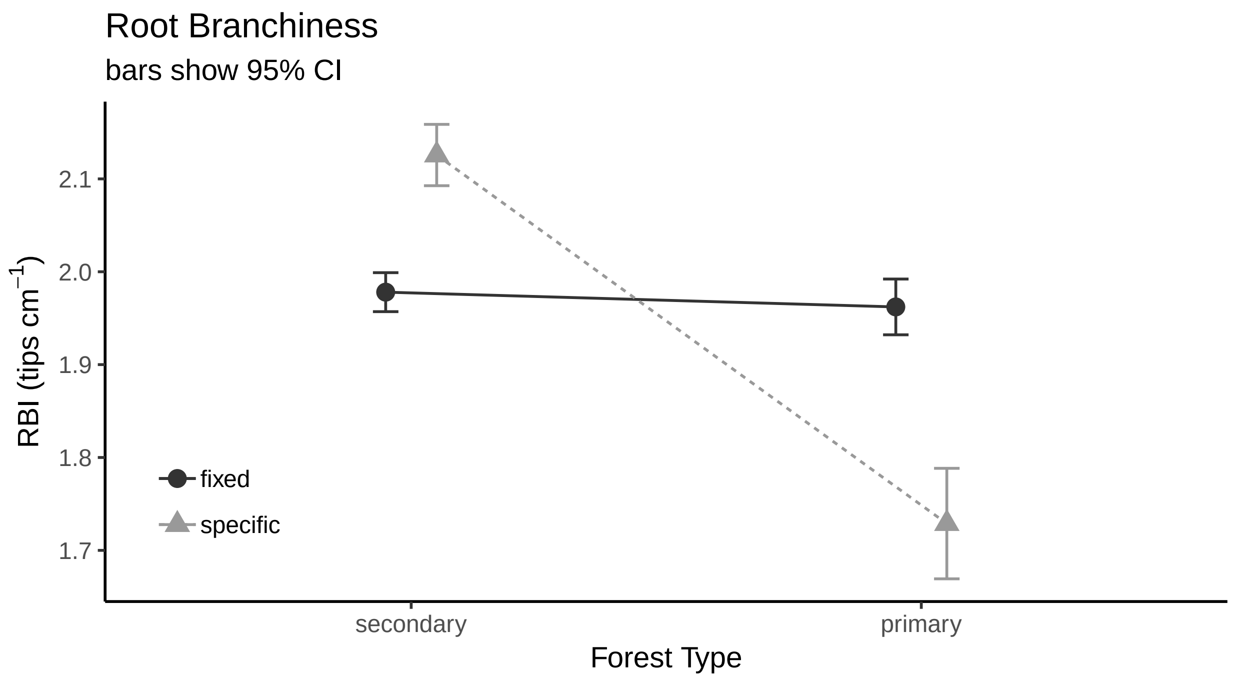


###### histogram

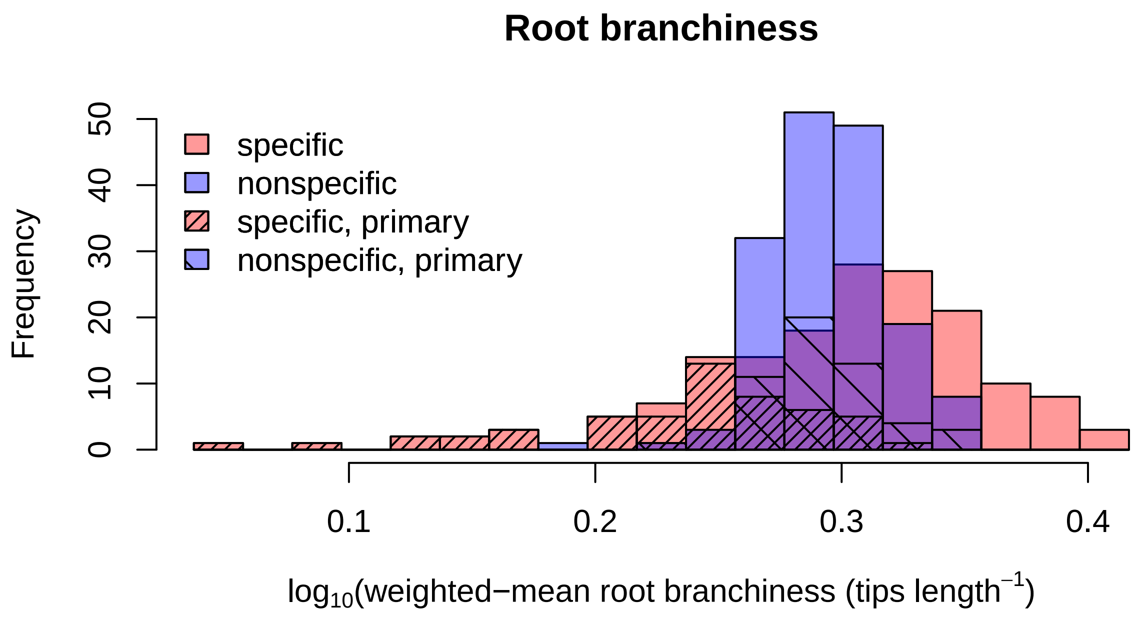
