## Supplement 4: Literate statistical document describing the gap-filling of functional traits and re-implementing the Trait Flex ANOVAS with gap-filled for "Intraspecific root and leaf trait variation with tropical forest successional status: consequences for community-weighted patterns"

### JFL_Intraspecific_Variability_Effects_2_WHOLEJFLCOMMU

*JAH*

*January 22, 2019*

#### Background

This document compliments the "JFL_Intraspecific_Variability_Effects.pdf", were I looked used Jan Leps' analysis for paritioning the varaiance in functional trait and community compositional turnover using data for the Jianfengling (JFL) plant community. The forest composition data come from Dr Han Xu's network of small plots. The trait data were collected during a summer 2017 trait campaign to JFL.

Here we re-work that analysis using data from the entire community. The reason behind re-working the analysis is that it was unlear if using a subset of the community baised the results in any particular way. One may contend that community-weighted analyses should indluce all species (or at least majority of the basal area in a sampled community). This poses a major increase in sampling effort, especially in species-rich tropical forests (>580 species in the JFL plot dataset).

Therefore, we used a phylogeny to predict traits for species that we missing data from. The phylogeny for all species in the JFL community was generated from Phylomatic (http://phylodiversity.net/phylomatic/) using the Slik2015 base tree (Slik et al. 2018) using method = phylomatic.

We fed the phylogeny and incomplete trait matrices into PhyloPars (www.ibi.vu.nl/programs/phylopars/) - a program for estimating missing parameter values using phylogeny (Bruggeman et al. 2009). This was done separately for the three trait matricies, 1) (the fixed) traits irrespective of forest type (species average trait values from all individuals sampled), and (2,3) for primary and secondary forest matrices (where trait averages were constrained to individuals collected in that forest type along the transect). We allowed correalated evolution of the different traits, but did not allow for intraspecific variability of features (in the model). Trait matrices had the 7 traits of interest (Leaf Area, SLA, Leaf Thickness, SRL, Root Diameter, Root Tissue Density, and Root Branching Intensity), plus Wood Specific Gravity. In order to have one trait with complete information for all species, we used Wood Specific Gravity. Data we obtained from either the Global Wood Density Database (Chave et al. 2009/ Zanne et al. 2009) or from the CTFS Wood Density dataset (http://ctfs.si.edu/Public/Datasets/CTFSWoodDensity/) using the getWoodDensity function in the BIOMASS package (Rjou-Mechain et al. 2017). In cases where species values were unattainable, genus, family or dataset values were used. Genus values were used for 278 species; Family values were used for 137 speices, and dataset values were used for 15 species.

**Important Sources:**


- Jorn Bruggeman, Haap Heringa & Bernd W. Brandt. 2009. PhyloPars: estimation of missing parameter values using phylogeny. Nucleic Acids Research, 37(2):W179-W184. https://doi.org/10.1093/nar/gkp370
- Jerome Chave et al. 2009. Towards a worldwide wood economics spectrum. Ecology Letters 12(4):351-366
- Maxime Rjou-Mechain et al. 2017. BIOMASS: an R package for estimating above-ground biomass and its uncertainty in tropical forests. Methods in Ecology and Evolution, 9(8):1163-1167. http://doi.wiley.com/10.1111/2041-210X.12753
- J.W. Ferry Slik et al. 2018. Phylogenetic classification of the world's tropical forests. PNAS 115(8):1837-1842. https://doi.org/10.1073/pnas.1714977115
- Amy E. Zanne et al. Data from: Towards a worldwide wood economics spectrum. Dryad Digital Repository. https://doi.org/10.5061/dryad.234

The rest of the analytical details follow what is oulined in "JFL_Intraspecific_Variability_Effects.pdf". The only differnece is that the results in this document use the trait and community datasets for all (>581) taxa.

At the end of this document, we have appened statisistical output from the PhyloPars phylogenetic-GLMs.

### Trait Flex ANOVAS

#### Leaf Area

```
##
##  Decomposing trait sum of squares into composition turnover
##  effect, intraspecific trait variability, and their covariation:
##              Turnover Intraspec. Covariation   Total
## forest.type    48.095     4563.3      936.96  5548.4
## Residuals   17565.513     2725.5    -4518.07 15773.0
## Total       17613.608     7288.9    -3581.11 21321.4
##
##  Relative contributions:
##             Turnover Intraspec. Covariation  Total
## forest.type 0.002256     0.2140     0.04394 0.2602
## Residuals   0.823845     0.1278    -0.21190 0.7398
## Total       0.826101     0.3419    -0.16796 1.0000
##
##  Significance of testable effects:
##             Turnover Intraspec.      Total
## forest.type  0.50636 1.9621e-36 3.0101e-12
```

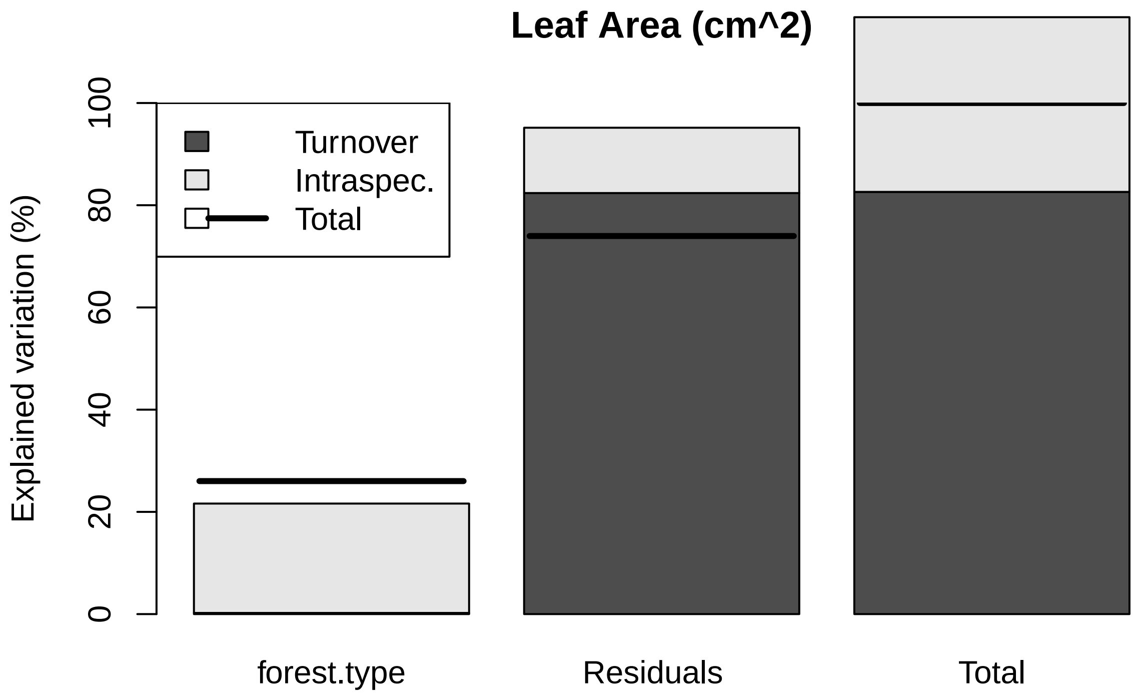


##### SLA

```
##
##  Decomposing trait sum of squares into composition turnover
##  effect, intraspecific trait variability, and their covariation:
##               Turnover Intraspec. Covariation   Total
## forest.type 3.4506e-04     47.941    -0.25724  47.684
## Residuals   1.1389e+02     24.434    15.05579 153.382
## Total       1.1389e+02     72.375    14.79856 201.066
##
##  Relative contributions:
##              Turnover Intraspec. Covariation  Total
## forest.type 1.716e-06     0.2384   -0.001279 0.2372
## Residuals   5.664e-01     0.1215    0.074880 0.7628
## Total       5.664e-01     0.3600    0.073600 1.0000
##
##  Significance of testable effects:
##             Turnover Intraspec.      Total
## forest.type  0.98235 4.8543e-40 3.7864e-11
```

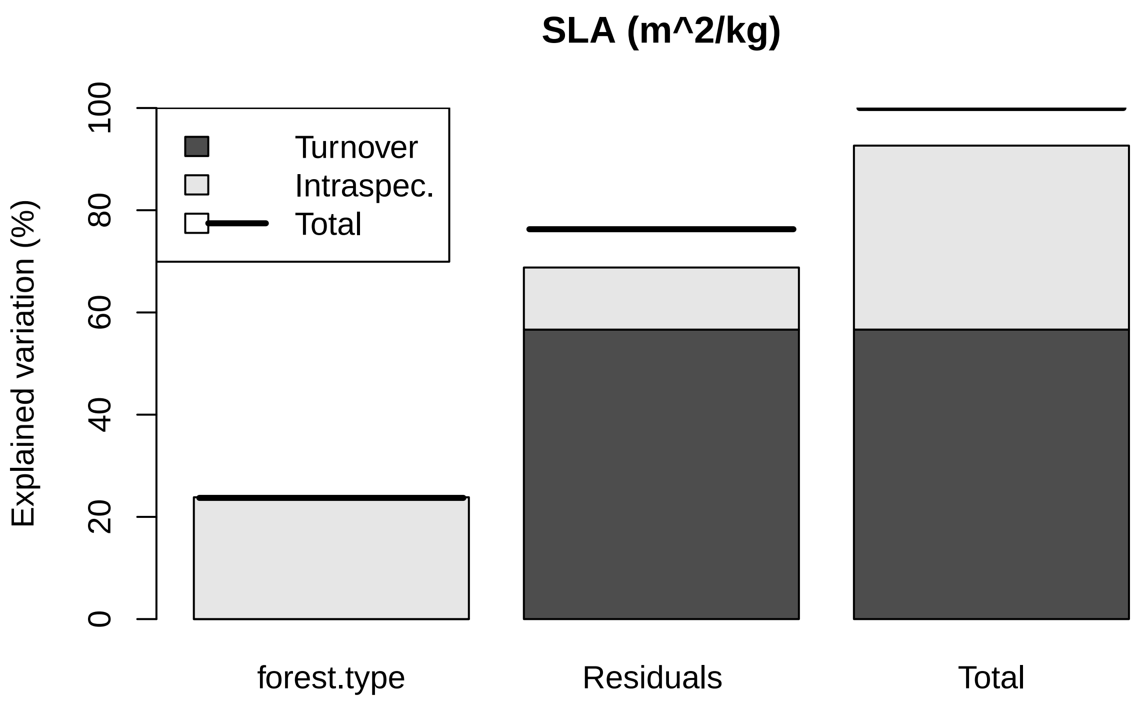


##### Leaf Thickness

```
##
##  Decomposing trait sum of squares into composition turnover
##  effect, intraspecific trait variability, and their covariation:
##               Turnover Intraspec. Covariation     Total
## forest.type 6.7635e-05  0.0050150  0.00116481 0.0062475
## Residuals   2.0613e-02  0.0054026  0.00014102 0.0261567
## Total       2.0681e-02  0.0104176  0.00130583 0.0324042
##
##  Relative contributions:
##             Turnover Intraspec. Covariation  Total
## forest.type 0.002087     0.1548    0.035946 0.1928
## Residuals   0.636124     0.1667    0.004352 0.8072
## Total       0.638212     0.3215    0.040298 1.0000
##
##  Significance of testable effects:
##             Turnover Intraspec.      Total
## forest.type  0.46701 7.1422e-25 4.0645e-09
```

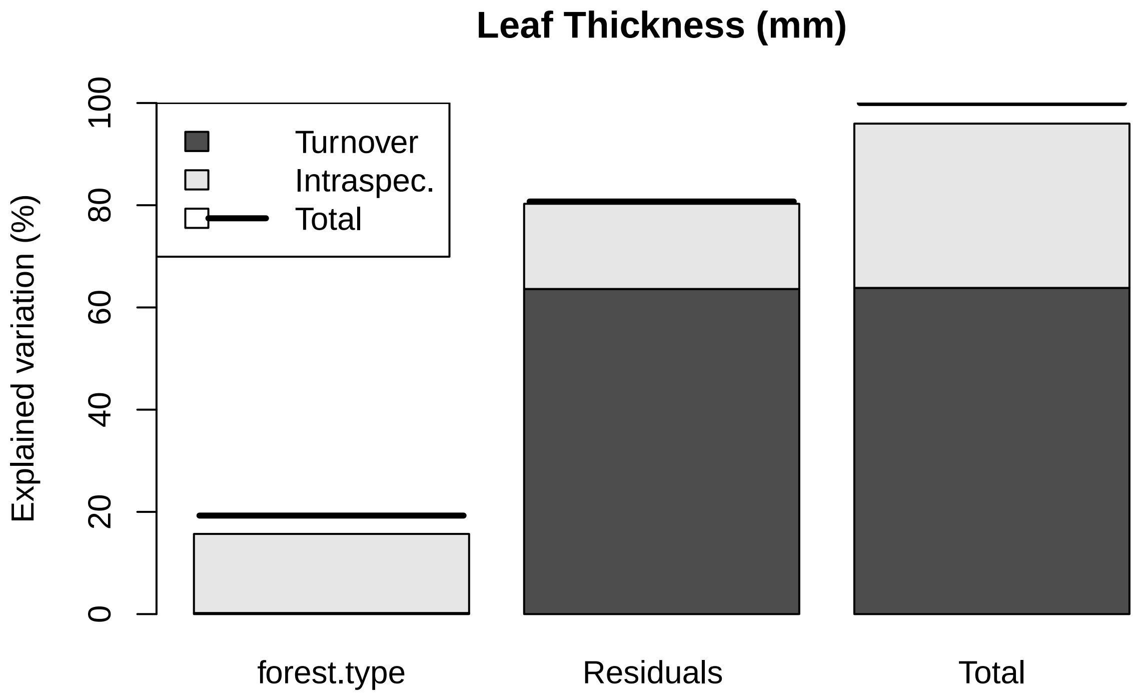


##### Root Avg Diameter

```
##
##  Decomposing trait sum of squares into composition turnover
##  effect, intraspecific trait variability, and their covariation:
##              Turnover Intraspec. Covariation    Total
## forest.type 0.0090128    0.15630   -0.075065 0.090244
## Residuals   0.6517348    0.13181   -0.096028 0.687521
## Total       0.6607477    0.28811   -0.171093 0.777766
##
##  Relative contributions:
##             Turnover Intraspec. Covariation Total
## forest.type  0.01159     0.2010    -0.09651 0.116
## Residuals    0.83796     0.1695    -0.12347 0.884
## Total        0.84955     0.3704    -0.21998 1.000
##
##  Significance of testable effects:
##             Turnover Intraspec.      Total
## forest.type   0.1364 2.6296e-29 8.0796e-06
```

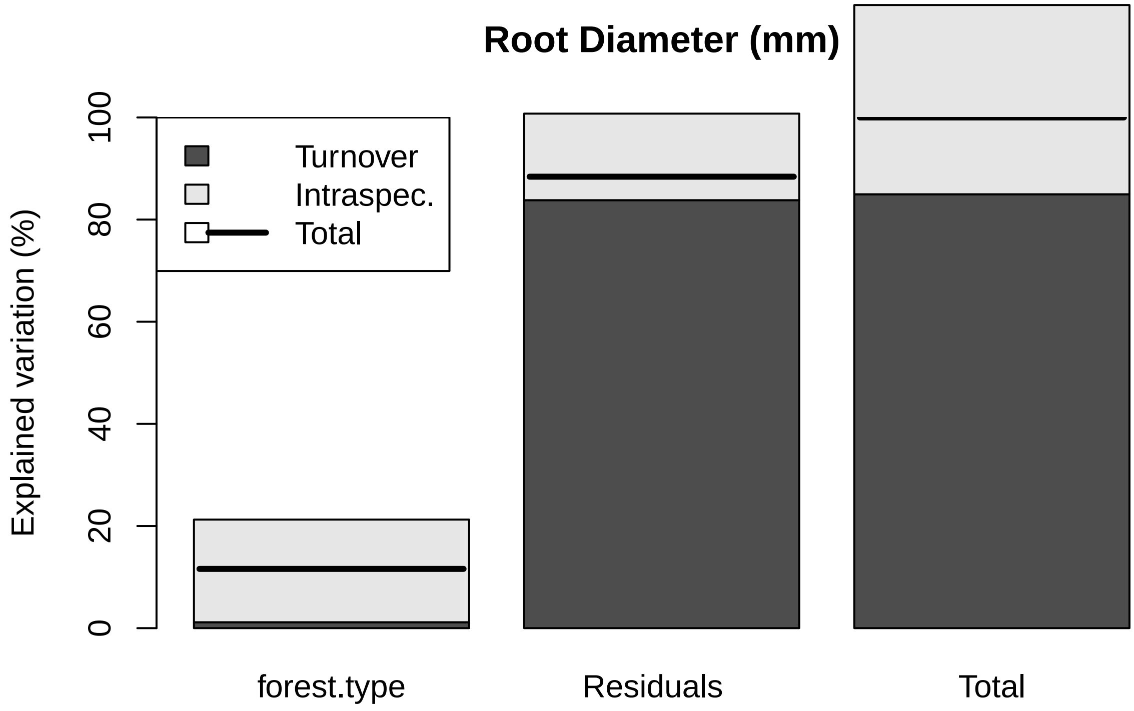


#### SRL

```
##
##  Decomposing trait sum of squares into composition turnover
##  effect, intraspecific trait variability, and their covariation:
##             Turnover Intraspec. Covariation  Total
## forest.type   50.122    2116.90     -651.47 1515.6
## Residuals   4994.058     947.38    -1182.22 4759.2
## Total       5044.180    3064.28    -1833.69 6274.8
##
##  Relative contributions:
##             Turnover Intraspec. Covariation  Total
## forest.type 0.007988     0.3374     -0.1038 0.2415
## Residuals   0.795894     0.1510     -0.1884 0.7585
## Total       0.803882     0.4883     -0.2922 1.0000
##
##  Significance of testable effects:
##             Turnover Intraspec.      Total
## forest.type   0.2041 3.8161e-43 2.3552e-11
```

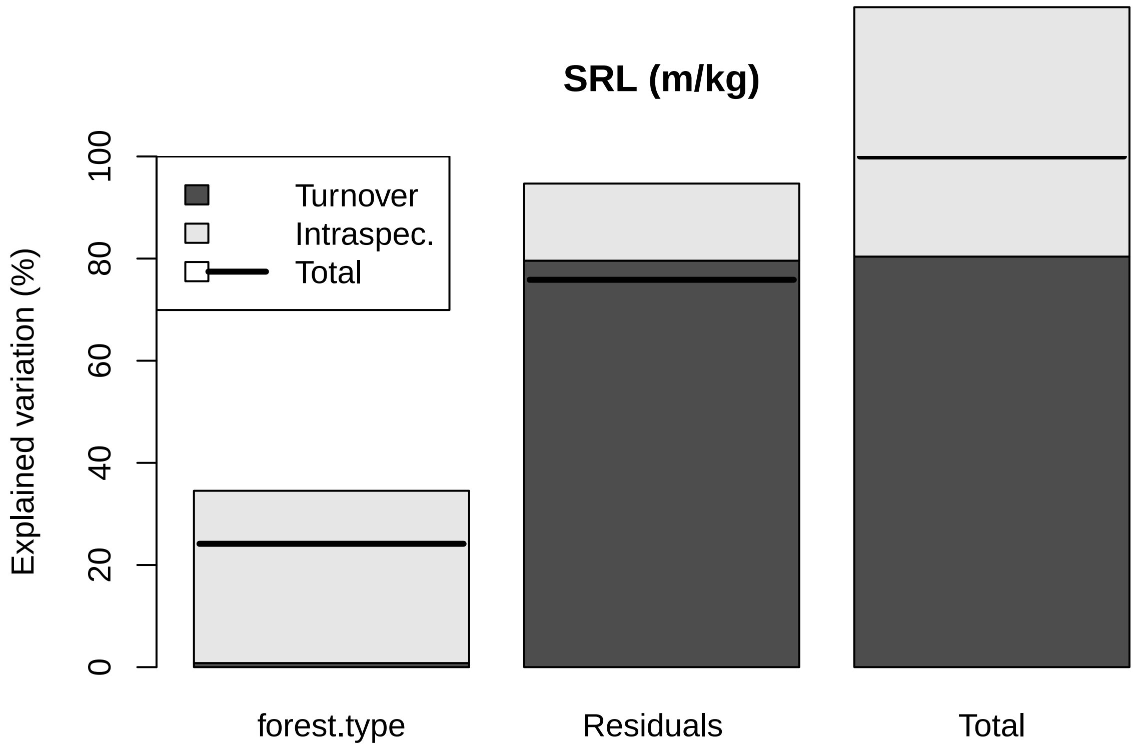


##### Root Tissue Density

```
##
##  Decomposing trait sum of squares into composition turnover
##  effect, intraspecific trait variability, and their covariation:
##              Turnover Intraspec. Covariation    Total
## forest.type 0.0019099   0.042871   -0.018098 0.026683
## Residuals   0.1720033   0.052441    0.055986 0.280430
## Total       0.1739132   0.095312    0.037889 0.307114
##
##  Relative contributions:
##             Turnover Intraspec. Covariation   Total
## forest.type 0.006219     0.1396    -0.05893 0.08688
## Residuals   0.560064     0.1708     0.18230 0.91312
## Total       0.566283     0.3103     0.12337 1.00000
##
##  Significance of testable effects:
##             Turnover Intraspec.      Total
## forest.type  0.18173 8.8935e-23 0.00012742
```

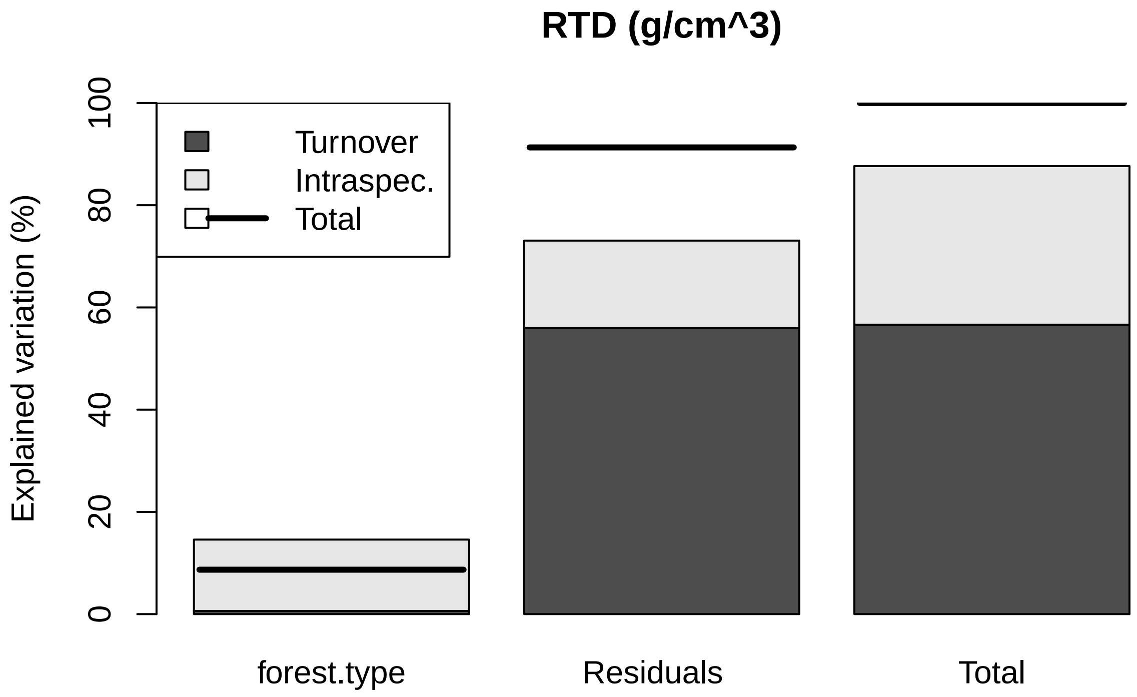


##### Root Branchiness

```
##
##  Decomposing trait sum of squares into composition turnover
##  effect, intraspecific trait variability, and their covariation:
##              Turnover Intraspec. Covariation  Total
## forest.type 0.0056787    4.26406     0.31122 4.5810
## Residuals   0.6144781    0.46652     0.32291 1.4039
## Total       0.6201567    4.73058     0.63413 5.9849
##
##  Relative contributions:
##              Turnover Intraspec. Covariation  Total
## forest.type 0.0009488    0.71247     0.05200 0.7654
## Residuals   0.1026720    0.07795     0.05396 0.2346
## Total       0.1036208    0.79042     0.10596 1.0000
##
##  Significance of testable effects:
##             Turnover Intraspec.      Total
## forest.type  0.22289 2.1345e-83 7.0294e-53
```

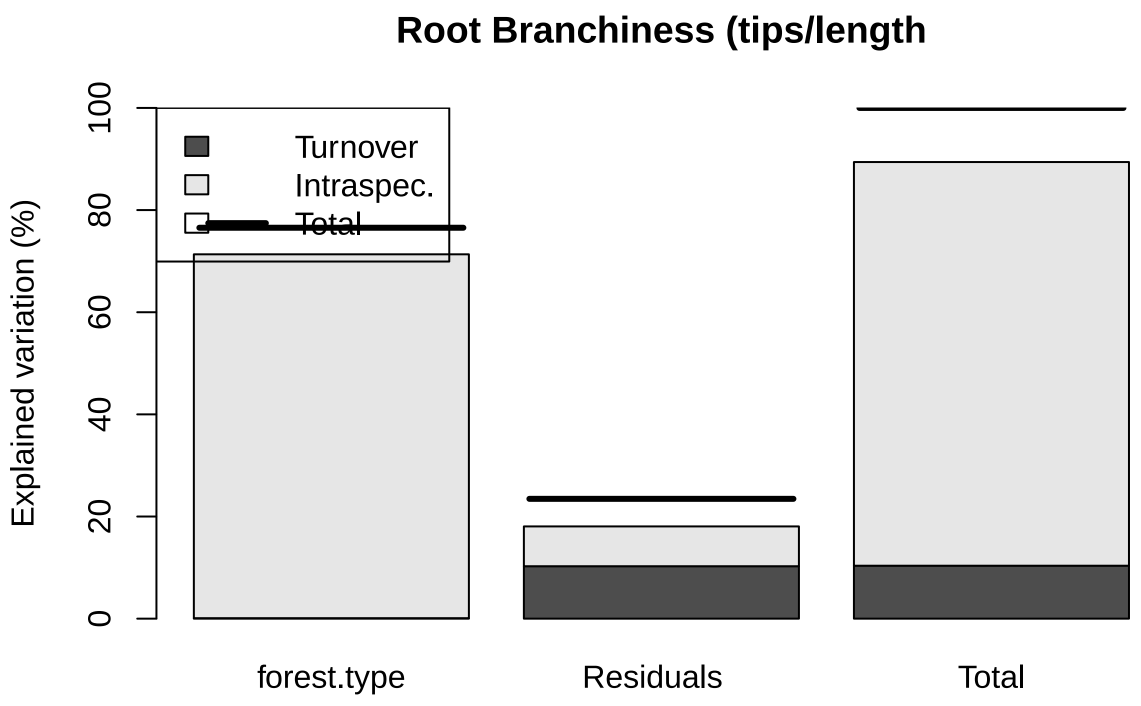
Root Branchiness (tips/length
Explained variation (%)
0
20
40
60
80
100
Turnover
Intraspec.
Total
forest.type
Residuals
Total

#### PLOTTING weighted averages

##### Leaf Area

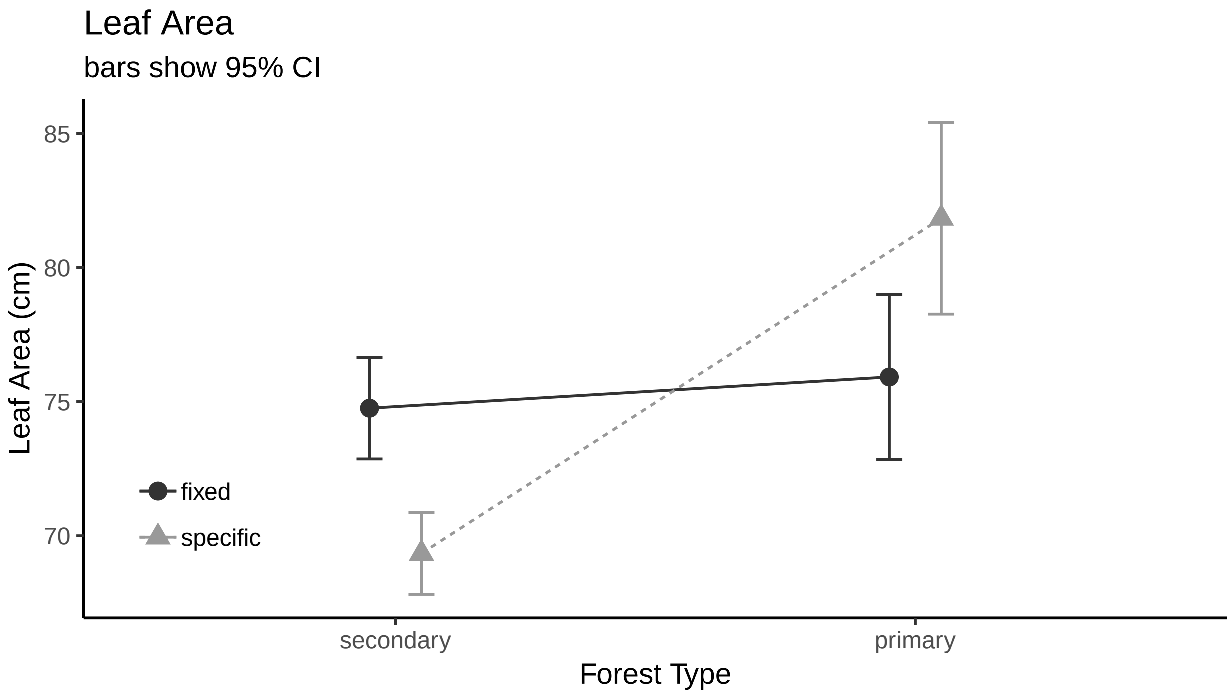


##### SLA

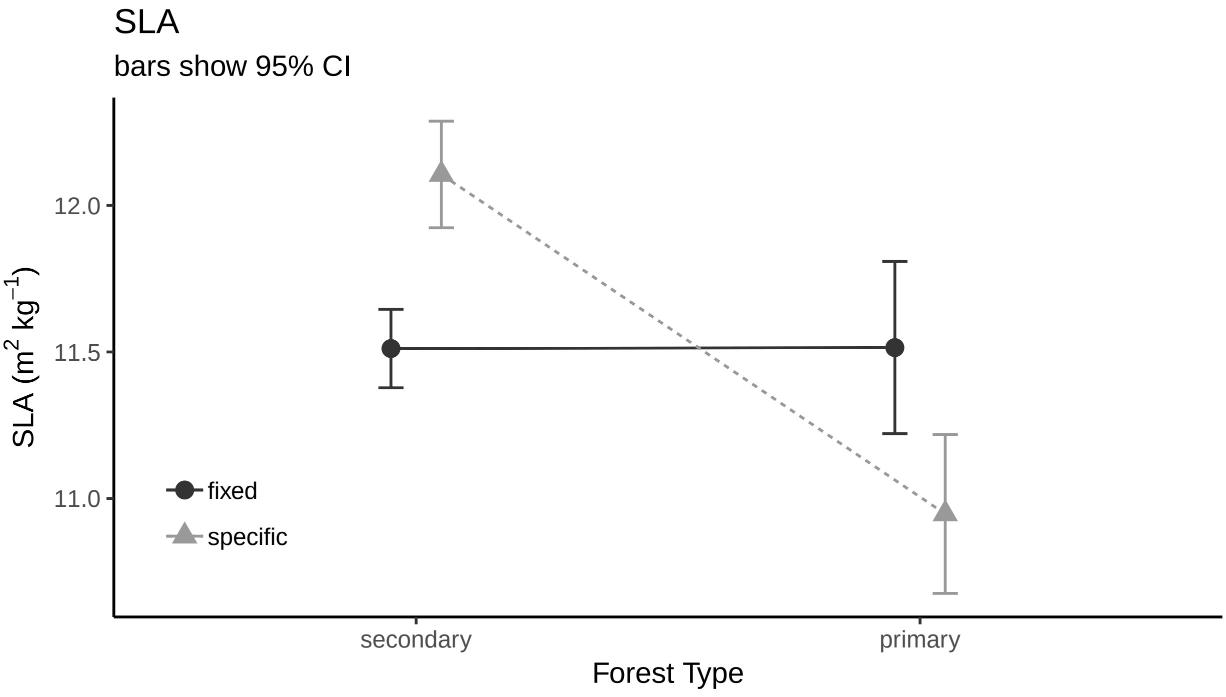

##### Leaf Thickness

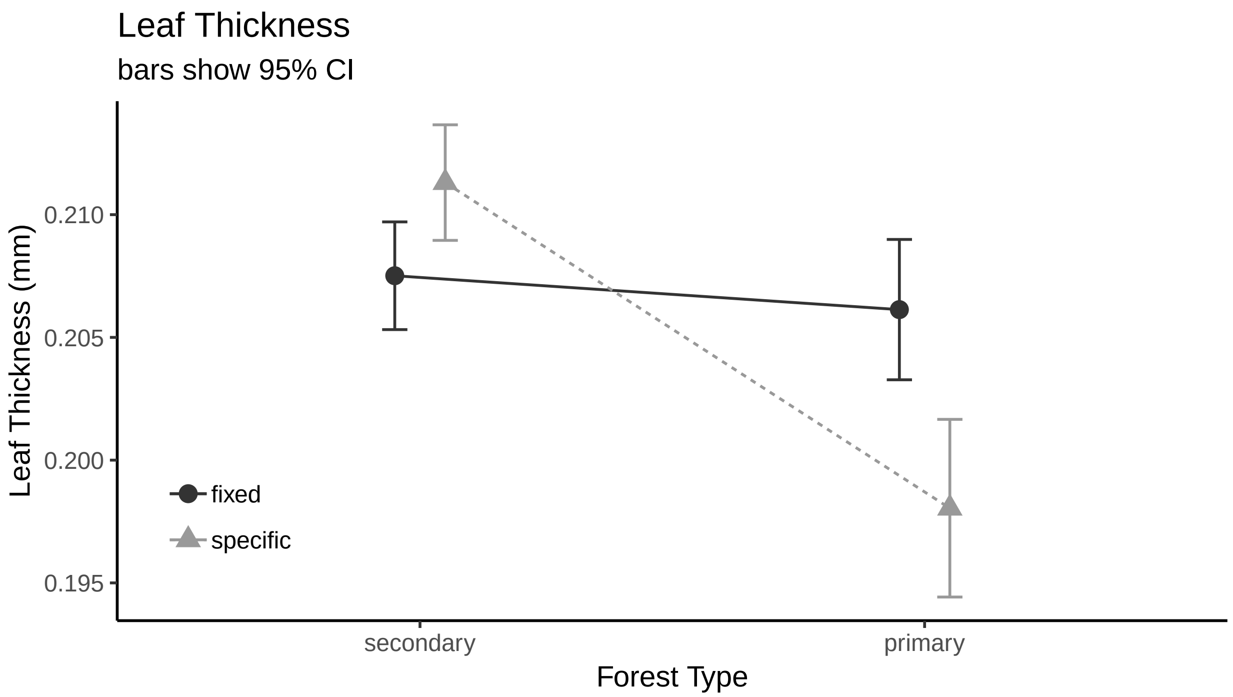


##### Avg. Root Diameter

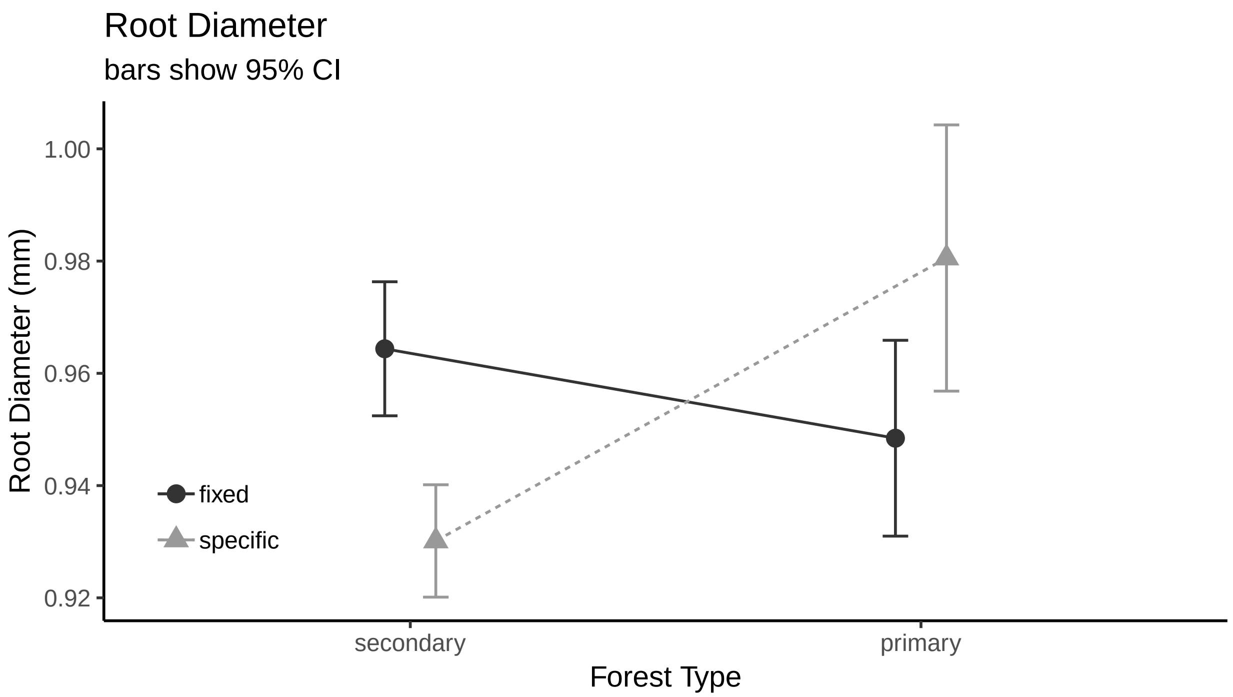

##### SRL

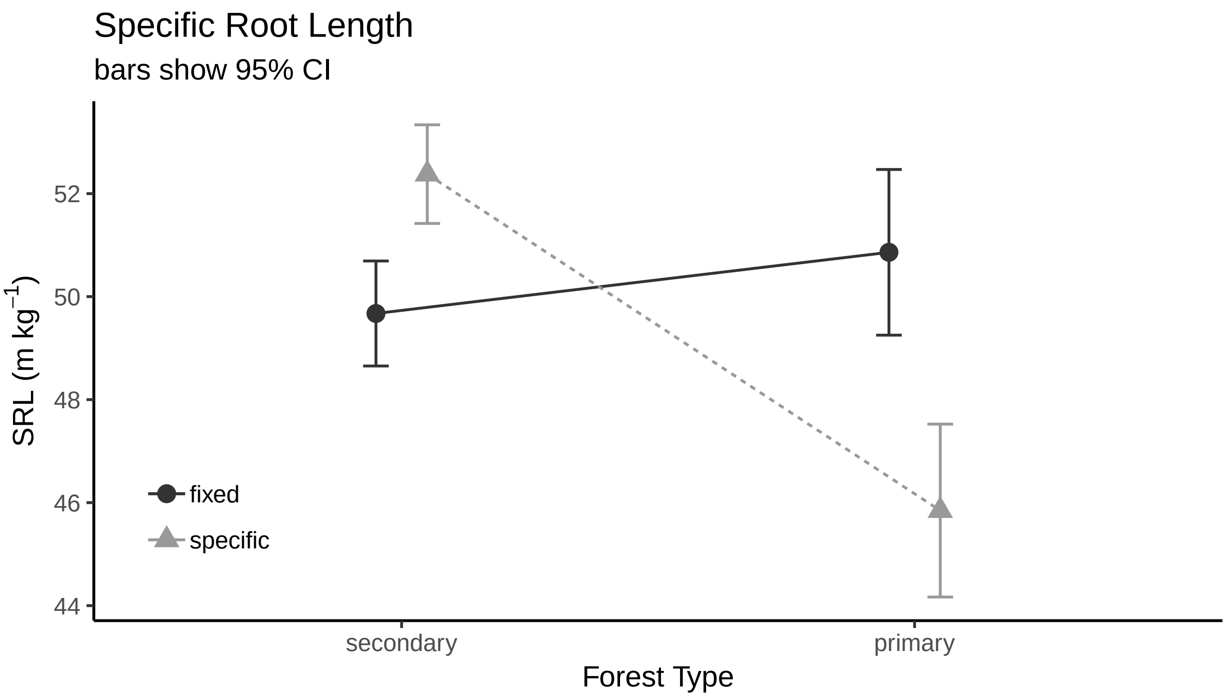


##### Root Tissue Density

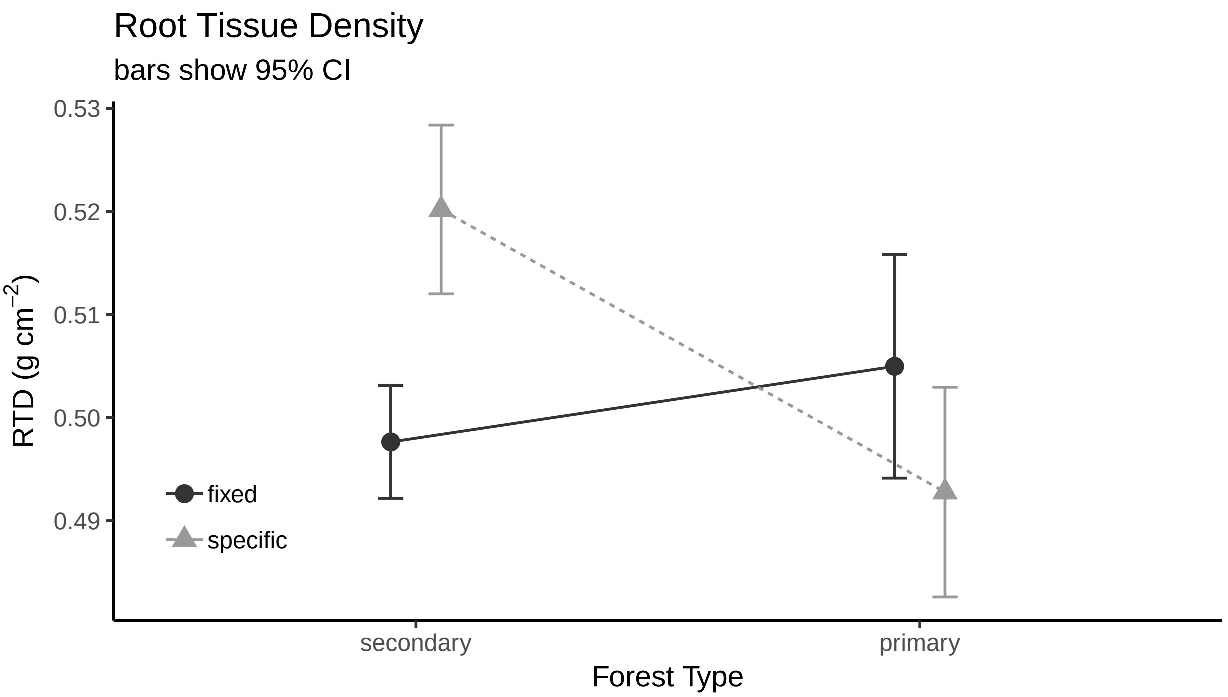

**Root Branchiness**

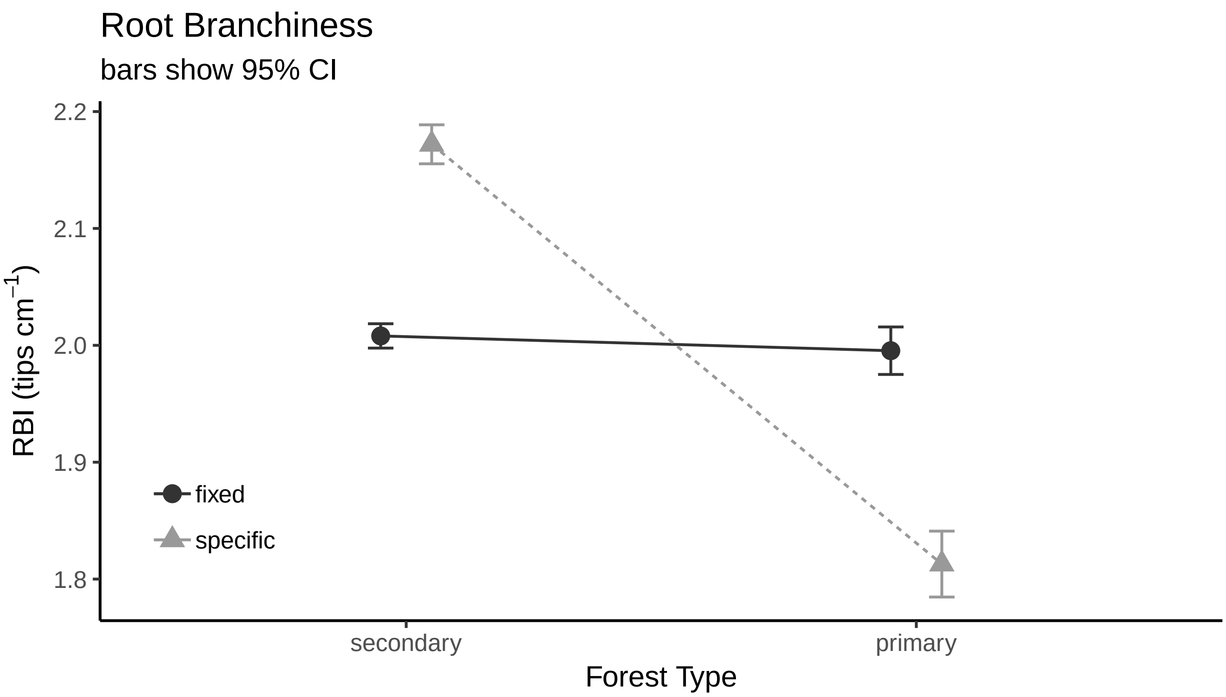

Centre for Integrative Bioinformatics VU
vrije Universiteit amsterdam
IBIVU - www.ibi.vu.nl



### PhyloPars results Model uses traits for all individuals (both primary and secondary forest)

#### Contents

Phylogenetic variability
Estimates for missing parameters
Cross-validation details

**Please cite:** Bruggeman J, Heringa J and Brandt BW. (2009) PhyloPars: estimation of missing parameter values using phylogeny.
Nucleic Acids Research 37: W179-W184.

#### Phylogenetic variability

PhyloPars first estimates the parameters of the evolutionary model, i.e., the phylogenetic covariances. These are subsequently used to estimate missing feature values and to perform cross-valida

| feature | phylogenetic s.d. | | cross-validation results | | | |
|---|---|---|---|---|---|---|
| | | | error | | bias | |
| meanWD | 0.049 | g/cm3 | 0.0497 | g/cm3 | -0.000606 | g/cm3 |
| meanLeafArea | 30.3 | cm2 | 29.3 | cm2 | -1.43 | cm2 |
| meanSLA | 2.547 | m2/Kg | 2.4 | m2/Kg | -0.0925 | m2/Kg |
| meanLeafThickness | 0.08613 | mm | 0.064 | mm | 5.55e-05 | mm |
| meanRootAvgDiam | 0.1652 | cm/10 | 0.124 | cm/10 | 0.00097 | cm/10 |
| meanSpecificRootLength | 20.49 | m/Kg | 13.3 | m/Kg | 0.0833 | m/Kg |
| meanRootTD | 0.07609 | g/cm3 | 0.076 | g/cm3 | -0.00175 | g/cm3 |
| meanBranchiness | 0.2629 | tips/length | 0.287 | tips/length | -0.00118 | tips/length |

##### Phylogenetic correlations

| | meanWD | meanLeafArea | meanSLA | meanLeafThickness | meanRootAvgDiam | meanSpecificRootLength | meanRootTD | meanBr |
|---|---|---|---|---|---|---|---|---|
| **meanWD** | 1 | | | | | | | |
| **meanLeafArea** | -0.055 | 1 | | | | | | |
| **meanSLA** | 0.053 | -0.136 | 1 | | | | | |
| **meanLeafThickness** | 0.042 | -0.031 | -0.224 | 1 | | | | |
| **meanRootAvgDiam** | 0.062 | -0.176 | -0.056 | 0.512 | 1 | | | |
| **meanSpecificRootLength** | -0.170 | 0.310 | 0.295 | -0.209 | -0.556 | 1 | | |
| **meanRootTD** | 0.220 | 0.058 | -0.195 | 0.044 | 0.078 | -0.465 | 1 | |
| **meanBranchiness** | 0.122 | 0.237 | -0.029 | -0.198 | -0.333 | 0.058 | 0.115 | 1 |

##### Phylogenetic regression coefficients

Rows represent independent variables and columns dependent variables. A regression of feature *i* on *j* thus can be found at row *j*, column *i*.

| | meanWD | meanLeafArea | meanSLA | meanLeafThickness | meanRootAvgDiam | meanSpecificRootLength | meanRootTD | meanBr |
|---|---|---|---|---|---|---|---|---|
| **meanWD** | 1 | -33.778 | 2.749 | 0.073 | 0.209 | -71.166 | 0.342 | 0.656 |
| **meanLeafArea** | -0.000 | 1 | -0.011 | -0.000 | -0.001 | 0.209 | 0.000 | 0.002 |
| **meanSLA** | 0.001 | -1.619 | 1 | -0.008 | -0.004 | 2.374 | -0.006 | -0.003 |
| **meanLeafThickness** | 0.024 | -10.796 | -6.628 | 1 | 0.982 | -49.702 | 0.039 | -0.605 |
| **meanRootAvgDiam** | 0.018 | -32.221 | -0.858 | 0.267 | 1 | -69.045 | 0.036 | -0.530 |
| **meanSpecificRootLength** | -0.000 | 0.458 | 0.037 | -0.001 | -0.004 | 1 | -0.002 | 0.001 |
| **meanRootTD** | 0.142 | 23.139 | -6.543 | 0.050 | 0.169 | -125.196 | 1 | 0.399 |
| **meanBranchiness** | 0.023 | 27.358 | -0.283 | -0.065 | -0.209 | 4.543 | 0.033 | 1 |

#### Estimates for missing parameters

Using the optimal phylogenetic covariances listed above, PhyloPars estimates the values that were originally missing in the feature matrix. The table below lists all estimated values. If one or more have been provided for a value, this is denoted by a trailing asterisk symbol (*). You can click on an entry to view the contribution of individual observations to the estimate, and to retrieve details su standard deviation of the estimate. You can also download the complete table as a single text file.

| | meanWD (g/cm3) | meanLeafArea (cm2) | meanSLA (m2/Kg) | meanLeafThickness (mm) | meanRootAvgDiam (cm/10) | meanSpecificRootLength (m/Kg) | meanRootTD (g/cm3) | mean (tip |
|---|---|---|---|---|---|---|---|---|

| | | | | | | | |
|---|---|---|---|---|---|---|---|
| root | 0.5269 | 68.37 | 11.80 | 0.2246 | 1.137 | 41.33 | 0.4136 |
| fagales | 0.6168 | 69.68 | 11.53 | 0.2043 | 0.8501 | 57.71 | 0.5530 |
| fagaceae | 0.6190 | 59.51 | 10.85 | 0.2040 | 0.8328 | 51.14 | 0.5960 |
| lithocarpus | 0.6897 | 45.09 | 11.92 | 0.2499 | 0.9760 | 35.77 | 0.6153 |
| Lithocarpusamygdalifolius | 0.7580* | 41.10* | 10.80* | 0.2134* | 0.7643* | 44.24* | 0.6286* |
| Lithocarpusbacgiangensis | 0.6773* | 45.51 | 11.88 | 0.2490 | 0.9734 | 36.66 | 0.6111 |
| Lithocarpusbrachystachyus | 0.6773* | 45.51 | 11.88 | 0.2490 | 0.9734 | 36.66 | 0.6111 |
| Lithocarpuschiungchungensis | 0.6773* | 45.51 | 11.88 | 0.2490 | 0.9734 | 36.66 | 0.6111 |
| Lithocarpuscorneus | 0.8180* | 40.75 | 12.27 | 0.2592 | 1.003 | 26.65 | 0.6592 |
| Lithocarpuselaeagnifolius | 0.6773* | 45.51 | 11.88 | 0.2490 | 0.9734 | 36.66 | 0.6111 |
| Lithocarpusfenestratus | 0.6773* | 68.18* | 8.638* | 0.2241* | 0.7912* | 54.19* | 0.5391* |
| Lithocarpusfenzelianus | 0.6773* | 38.27* | 7.581* | 0.2253* | 1.189* | 21.23* | 0.5987* |
| Lithocarpushancei | 0.6773* | 29.48* | 24.72* | 0.1373* | 1.017* | 43.43* | 0.4978* |
| Lithocarpushandelianus | 0.7610* | 55.54* | 9.797* | 0.6953* | 1.310* | 14.49* | 0.5898* |
| Lithocarpushowii | 0.6773* | 34.51* | 12.46* | 0.1830* | 0.7426* | 37.12* | 0.7950* |
| Lithocarpuslitseifolius | 0.6773* | 45.51 | 11.88 | 0.2490 | 0.9734 | 36.66 | 0.6111 |
| Lithocarpuslongipedicellatus | 0.5995* | 58.98* | 10.97* | 0.1678* | 0.9867* | 32.67* | 0.6309* |
| Lithocarpuspseudovestitus | 0.6773* | 28.93* | 10.55* | 0.1756* | 1.059* | 36.63* | 0.6186* |
| Lithocarpussp0502 | 0.6773* | 45.51 | 11.88 | 0.2490 | 0.9734 | 36.66 | 0.6111 |
| castanopsis | 0.5502 | 65.27 | 11.19 | 0.1766 | 0.8179 | 51.46 | 0.6351 |
| Castanopsiscarlesii | 0.4407* | 68.97 | 10.88 | 0.1686 | 0.7951 | 59.25 | 0.5976 |
| Castanopsisfissa | 0.4446* | 98.66* | 7.733* | 0.1736* | 0.8078* | 58.74* | 0.5658* |
| Castanopsishainanensis | 0.6473* | 61.99 | 11.45 | 0.1837 | 0.8383 | 44.55 | 0.6683 |
| Castanopsishystrix | 0.5650* | 64.77 | 11.23 | 0.1777 | 0.8210 | 50.41 | 0.6402 |
| Castanopsisjianfenglingensis | 0.5680* | 64.67 | 11.23 | 0.1779 | 0.8217 | 50.19 | 0.6412 |
| Castanopsisjucunda | 0.5680* | 64.67 | 11.23 | 0.1779 | 0.8217 | 50.19 | 0.6412 |
| Castanopsistonkinensis | 0.5680* | 39.51* | 14.34* | 0.1536* | 0.7728* | 51.74* | 0.6980* |
| quercus | 0.7003 | 53.82 | 11.40 | 0.2184 | 0.8864 | 41.75 | 0.6314 |
| Quercusacutissima | 0.7334* | 52.70 | 11.49 | 0.2208 | 0.8934 | 39.39 | 0.6427 |
| Cyclobalanopsis | 0.6076 | 51.57 | 10.28 | 0.2008 | 0.8152 | 47.11 | 0.6146 |
| Cyclobalanopsisbella | 0.6062* | 51.62 | 10.28 | 0.2007 | 0.8149 | 47.21 | 0.6141 |
| Cyclobalanopsisblakei | 0.6062* | 34.31* | 13.60* | 0.1417* | 0.7440* | 54.32* | 0.6249* |
| Cyclobalanopsisedithiae | 0.6062* | 91.65* | 9.141* | 0.2309* | 0.8085* | 39.83* | 0.6595* |
| Cyclobalanopsisfleuryi | 0.6062* | 91.91* | 8.118* | 0.2207* | 0.8503* | 36.52* | 0.7028* |
| Cyclobalanopsishui | 0.6062* | 26.97* | 8.857* | 0.2483* | 0.7869* | 52.08* | 0.5501* |
| Cyclobalanopsisneglecta | 0.6062* | 12.37* | 12.44* | 0.1606* | 0.8209* | 53.16* | 0.5258* |
| Cyclobalanopsispatelliformis | 0.6062* | 44.18* | 8.974* | 0.1994* | 0.8638* | 42.49* | 0.6443* |
| Cyclobalanopsisphanera | 0.6062* | 51.62 | 10.28 | 0.2007 | 0.8149 | 47.21 | 0.6141 |
| betulaceae | 0.5671 | 80.55 | 11.14 | 0.1945 | 0.7952 | 68.36 | 0.5416 |
| betula | 0.5567 | 80.91 | 11.11 | 0.1937 | 0.7930 | 69.10 | 0.5381 |
| Betulaalnoides | 0.5515* | 81.08 | 11.09 | 0.1934 | 0.7919 | 69.47 | 0.5363 |
| juglandaceae | 0.5280 | 110.4 | 11.21 | 0.1797 | 0.7158 | 91.78 | 0.4973 |
| engelhardia | 0.4916 | 130.6 | 11.23 | 0.1691 | 0.6606 | 108.1 | 0.4642 |
| Engelhardiahainanensis | 0.4879* | 130.7 | 11.22 | 0.1688 | 0.6599 | 108.4 | 0.4630 |
| Engelhardiaroxburghiana | 0.4879* | 141.5* | 8.116* | 0.1122* | 0.7735* | 46.39* | 0.6088* |
| Engelhardiaspicata | 0.4879* | 212.8* | 15.09* | 0.1803* | 0.4544* | 224.5* | 0.3024* |
| EngelhardiaspicatavarAceriflora | 0.4879* | 130.7 | 11.22 | 0.1688 | 0.6599 | 108.4 | 0.4630 |
| Engelhardiaunijuga | 0.4879* | 47.36* | 10.51* | 0.2100* | 0.7280* | 61.17* | 0.4674* |
| rosales | 0.6484 | 66.41 | 12.48 | 0.2135 | 0.9044 | 55.43 | 0.5219 |
| moraceae | 0.5627 | 55.33 | 13.43 | 0.2198 | 0.9180 | 63.74 | 0.4429 |
| broussonetia | 0.3365 | 79.99 | 13.27 | 0.1970 | 0.7719 | 84.65 | 0.4744 |
| Broussonetiapapyrifera | 0.2900* | 81.56 | 13.15 | 0.1936 | 0.7622 | 87.96 | 0.4585 |
| ficus | 0.3949 | 95.04 | 13.90 | 0.1949 | 0.6853 | 85.32 | 0.6032 |
| Ficusaltissima | 0.4730* | 92.40 | 14.11 | 0.2006 | 0.7016 | 79.76 | 0.6299 |
| Ficusauriculata | 0.4680* | 92.57 | 14.10 | 0.2002 | 0.7006 | 80.12 | 0.6282 |
| Ficusfistulosa | 0.3800* | 95.55 | 13.86 | 0.1938 | 0.6822 | 86.38 | 0.5981 |
| Ficusformosana | 0.4055* | 33.56* | 13.81* | 0.2024* | 0.5994* | 79.08* | 0.6734* |
| Ficusglaberrima | 0.4055* | 94.68 | 13.93 | 0.1957 | 0.6875 | 84.56 | 0.6069 |
| Ficusheteropleura | 0.4055* | 94.68 | 13.93 | 0.1957 | 0.6875 | 84.56 | 0.6069 |
| Ficushirta | 0.4055* | 242.5* | 12.83* | 0.1561* | 0.6230* | 75.48* | 0.7416* |
| Ficuslangkokensis | 0.4055* | 94.68 | 13.93 | 0.1957 | 0.6875 | 84.56 | 0.6069 |
| Ficusnervosa | 0.2800* | 98.92 | 13.58 | 0.1865 | 0.6613 | 93.50 | 0.5639 |
| Ficuspandurata | 0.4055* | 94.68 | 13.93 | 0.1957 | 0.6875 | 84.56 | 0.6069 |

| | | | | | | | |
|---|---|---|---|---|---|---|---|
| Ficuspubigera | 0.4055* | 94.68 | 13.93 | 0.1957 | 0.6875 | 84.56 | 0.6069 |
| Ficussimplicissima | 0.4055* | 94.68 | 13.93 | 0.1957 | 0.6875 | 84.56 | 0.6069 |
| Ficussubpisocarpa | 0.4055* | 94.68 | 13.93 | 0.1957 | 0.6875 | 84.56 | 0.6069 |
| Ficustinctoria | 0.4055* | 94.68 | 13.93 | 0.1957 | 0.6875 | 84.56 | 0.6069 |
| Ficustuphapensis | 0.4055* | 94.68 | 13.93 | 0.1957 | 0.6875 | 84.56 | 0.6069 |
| Ficusvariegata | 0.3270* | 97.34 | 13.71 | 0.1899 | 0.6711 | 90.15 | 0.5800 |
| Ficusvariolosa | 0.4055* | 94.68 | 13.93 | 0.1957 | 0.6875 | 84.56 | 0.6069 |
| Ficusvasculosa | 0.3000* | 20.09* | 15.09* | 0.2177* | 0.7687* | 109.1* | 0.4239* |
| antiaris | 0.3884 | 86.75 | 13.65 | 0.1976 | 0.7333 | 83.37 | 0.5466 |
| Antiaristoxicaria | 0.3831* | 86.93 | 13.63 | 0.1972 | 0.7322 | 83.75 | 0.5447 |
| artocarpus | 0.4959 | 34.22 | 15.35 | 0.1824 | 1.134 | 53.36 | 0.3831 |
| Artocarpusnitidus | 0.4800* | 34.76 | 15.30 | 0.1812 | 1.131 | 54.50 | 0.3776 |
| Artocarpusstyracifolius | 0.5077* | 22.15* | 16.43* | 0.1669* | 1.252* | 44.97* | 0.3686* |
| Artocarpustonkinensis | 0.4667* | 35.21 | 15.27 | 0.1802 | 1.128 | 55.45 | 0.3731 |
| streblus | 0.6898 | 51.41 | 12.74 | 0.2511 | 0.8852 | 60.28 | 0.4406 |
| Streblusilicifolius | 0.7321* | 49.98 | 12.86 | 0.2542 | 0.8941 | 57.27 | 0.4551 |
| Streblusindicus | 0.7321* | 50.36* | 11.81* | 0.2761* | 0.8347* | 62.86* | 0.4092* |
| Streblustaxoides | 0.7321* | 49.98 | 12.86 | 0.2542 | 0.8941 | 57.27 | 0.4551 |
| urticaceae | 0.4506 | 61.91 | 12.89 | 0.2091 | 0.8883 | 71.27 | 0.4145 |
| Oreocnide | 0.3900 | 63.96 | 12.72 | 0.2047 | 0.8756 | 75.58 | 0.3938 |
| Oreocnidetonkinensis | 0.3294* | 66.01 | 12.55 | 0.2003 | 0.8629 | 79.90 | 0.3730 |
| debregeasia | 0.3536 | 65.19 | 12.62 | 0.2020 | 0.8680 | 78.17 | 0.3813 |
| Debregeasiasquamata | 0.3294* | 66.01 | 12.55 | 0.2003 | 0.8629 | 79.90 | 0.3730 |
| cannabaceae | 0.6187 | 59.03 | 13.11 | 0.2189 | 0.9171 | 58.87 | 0.4820 |
| aphananthe | 0.6543 | 57.83 | 13.21 | 0.2215 | 0.9246 | 56.33 | 0.4942 |
| Aphananthecuspidata | 0.6900* | 56.62 | 13.31 | 0.2241 | 0.9320 | 53.79 | 0.5064 |
| celtis | 0.6608 | 57.61 | 13.23 | 0.2220 | 0.9259 | 55.87 | 0.4964 |
| Celtisphilippensis | 0.7030* | 56.18 | 13.34 | 0.2250 | 0.9347 | 52.87 | 0.5108 |
| gironniera | 0.5141 | 62.56 | 12.82 | 0.2112 | 0.8952 | 66.31 | 0.4462 |
| Gironnierasubaequalis | 0.4617* | 64.33 | 12.68 | 0.2074 | 0.8843 | 70.04 | 0.4283 |
| rhamnaceae | 0.7808 | 59.14 | 13.08 | 0.2257 | 0.9384 | 46.44 | 0.5573 |
| rhamnella | 0.7594 | 59.87 | 13.02 | 0.2241 | 0.9339 | 47.97 | 0.5500 |
| Rhamnellarubrinervis | 0.7421* | 60.45 | 12.98 | 0.2229 | 0.9303 | 49.20 | 0.5440 |
| ventilago | 0.9216 | 54.39 | 13.47 | 0.2360 | 0.9678 | 36.43 | 0.6055 |
| Ventilagoleiocarpa | 0.9800* | 52.42 | 13.63 | 0.2403 | 0.9801 | 32.27 | 0.6255 |
| rosaceae | 0.6498 | 66.37 | 12.48 | 0.2136 | 0.9047 | 55.33 | 0.5224 |
| photinia | 0.7217 | 63.94 | 12.68 | 0.2189 | 0.9197 | 50.21 | 0.5470 |
| Photiniabenthamiana | 0.7235* | 63.88 | 12.69 | 0.2190 | 0.9201 | 50.08 | 0.5476 |
| Photiniaprunifolia | 0.7235* | 63.88 | 12.69 | 0.2190 | 0.9201 | 50.08 | 0.5476 |
| eriobotrya | 0.7683 | 62.36 | 12.81 | 0.2223 | 0.9295 | 46.89 | 0.5630 |
| Eriobotryadeflexa | 0.7755* | 62.12 | 12.83 | 0.2228 | 0.9310 | 46.38 | 0.5654 |
| laurocerasus | 0.6522 | 66.28 | 12.49 | 0.2138 | 0.9052 | 55.16 | 0.5232 |
| Laurocerasusphaeosticta | 0.6481* | 66.42 | 12.48 | 0.2135 | 0.9043 | 55.45 | 0.5218 |
| pygeum | 0.6522 | 66.28 | 12.49 | 0.2138 | 0.9052 | 55.16 | 0.5232 |
| Pygeumtopengii | 0.6481* | 66.42 | 12.48 | 0.2135 | 0.9043 | 55.45 | 0.5218 |
| Rhaphiolepis | 0.6487 | 66.41 | 12.48 | 0.2135 | 0.9045 | 55.41 | 0.5220 |
| Rhaphiolepisferruginea | 0.6481* | 66.42 | 12.48 | 0.2135 | 0.9043 | 55.45 | 0.5218 |
| Rhaphiolepisindica | 0.6481* | 66.42 | 12.48 | 0.2135 | 0.9043 | 55.45 | 0.5218 |
| fabales | 0.6463 | 72.37 | 12.31 | 0.2106 | 0.9121 | 54.46 | 0.5251 |
| fabaceae | 0.6541 | 72.11 | 12.33 | 0.2111 | 0.9137 | 53.90 | 0.5278 |
| archidendron | 0.4954 | 77.47 | 11.90 | 0.1995 | 0.8805 | 65.20 | 0.4734 |
| Archidendronclypearia | 0.3233* | 83.28 | 11.42 | 0.1870 | 0.8445 | 77.44 | 0.4146 |
| Archidendronlucidum | 0.5814* | 74.57 | 12.13 | 0.2058 | 0.8985 | 59.08 | 0.5029 |
| Archidendronutile | 0.5814* | 74.57 | 12.13 | 0.2058 | 0.8985 | 59.08 | 0.5029 |
| albizia | 0.4780 | 78.06 | 11.85 | 0.1983 | 0.8769 | 66.43 | 0.4675 |
| Albiziachinensis | 0.3000* | 84.07 | 11.36 | 0.1853 | 0.8397 | 79.10 | 0.4066 |
| Albiziaodoratissima | 0.6385* | 72.64 | 12.29 | 0.2100 | 0.9105 | 55.01 | 0.5224 |
| peltophorum | 0.8257 | 66.32 | 12.81 | 0.2237 | 0.9496 | 41.69 | 0.5865 |
| Peltophorumtonkinense | 0.8486* | 65.54 | 12.87 | 0.2253 | 0.9544 | 40.06 | 0.5943 |
| sindora | 0.8410 | 65.80 | 12.85 | 0.2248 | 0.9528 | 40.60 | 0.5917 |
| Sindoraglabra | 0.8486* | 65.54 | 12.87 | 0.2253 | 0.9544 | 40.06 | 0.5943 |
| ormosia | 0.6027 | 73.85 | 12.19 | 0.2074 | 0.9030 | 57.56 | 0.5101 |
| Ormosiabalansae | 0.4505* | 78.99 | 11.77 | 0.1962 | 0.8711 | 68.39 | 0.4581 |
| Ormosiafordiana | 0.8486* | 65.54 | 12.87 | 0.2253 | 0.9544 | 40.06 | 0.5943 |

| | | | | | | | |
|---|---|---|---|---|---|---|---|
| Ormosiapinnata | **0.5683**** | **75.01** | **12.10** | **0.2049** | **0.8958** | **60.01** | **0.4984** |
| Ormosiasemicastrata | **0.6213*** | **73.22** | **12.24** | **0.2087** | **0.9069** | **56.23** | **0.5165** |
| Ormosiaxylocarpa | **0.5055*** | **77.13** | **11.93** | **0.2003** | **0.8826** | **64.48** | **0.4769** |
| dalbergia | **0.6828** | **71.14** | **12.41** | **0.2132** | **0.9197** | **51.86** | **0.5376** |
| Dalbergiahainanensis | **0.6888*** | **70.94** | **12.43** | **0.2137** | **0.9210** | **51.44** | **0.5396** |
| polygalaceae | **0.6705** | **71.56** | **12.38** | **0.2123** | **0.9171** | **52.74** | **0.5334** |
| xanthophyllum | **0.6786** | **71.28** | **12.40** | **0.2129** | **0.9188** | **52.16** | **0.5361** |
| Xanthophyllumhainanense | **0.6867*** | **71.01** | **12.42** | **0.2135** | **0.9205** | **51.58** | **0.5389** |
| malpighiales | **0.6420** | **75.62** | **12.38** | **0.2099** | **0.9256** | **54.09** | **0.5176** |
| euphorbiaceae | **0.5468** | **78.83** | **12.11** | **0.2030** | **0.9057** | **60.86** | **0.4850** |
| mallotus | **0.4875** | **80.83** | **11.95** | **0.1986** | **0.8933** | **65.08** | **0.4647** |
| Mallotusanomalus | **0.5038*** | **80.28** | **12.00** | **0.1998** | **0.8967** | **63.92** | **0.4703** |
| Mallotuspaniculatus | **0.3450*** | **85.65** | **11.56** | **0.1882** | **0.8635** | **75.22** | **0.4159** |
| Mallotusphilippensis | **0.6033*** | **76.92** | **12.27** | **0.2071** | **0.9175** | **56.84** | **0.5043** |
| Mallotusyunnanensis | **0.5038*** | **80.28** | **12.00** | **0.1998** | **0.8967** | **63.92** | **0.4703** |
| macaranga | **0.4576** | **81.84** | **11.87** | **0.1965** | **0.8870** | **67.21** | **0.4544** |
| Macarangadenticulata | **0.4335*** | **82.66** | **11.80** | **0.1947** | **0.8820** | **68.92** | **0.4462** |
| hancea | **0.5215** | **79.69** | **12.04** | **0.2011** | **0.9004** | **62.66** | **0.4763** |
| Hanceahookeriana | **0.5431*** | **78.96** | **12.10** | **0.2027** | **0.9049** | **61.12** | **0.4837** |
| cleidion | **0.5048** | **80.25** | **12.00** | **0.1999** | **0.8969** | **63.85** | **0.4706** |
| Cleidionbrevipetiolatum | **0.5167*** | **79.85** | **12.03** | **0.2008** | **0.8994** | **63.01** | **0.4747** |
| koilodepas | **0.5339** | **79.27** | **12.08** | **0.2020** | **0.9030** | **61.78** | **0.4806** |
| Koilodepashainanense | **0.5431*** | **78.96** | **12.10** | **0.2027** | **0.9049** | **61.12** | **0.4837** |
| claoxylon | **0.3813** | **84.42** | **11.66** | **0.1909** | **0.8711** | **72.64** | **0.4284** |
| Claoxylonindicum | **0.3550*** | **85.31** | **11.59** | **0.1890** | **0.8656** | **74.51** | **0.4193** |
| alchornea | **0.4163** | **83.24** | **11.76** | **0.1934** | **0.8784** | **70.15** | **0.4403** |
| Alchornearugosa | **0.4084*** | **83.50** | **11.73** | **0.1929** | **0.8768** | **70.71** | **0.4376** |
| croton | **0.5191** | **79.77** | **12.04** | **0.2009** | **0.8999** | **62.83** | **0.4755** |
| Crotoncascarilloides | **0.5104*** | **80.06** | **12.01** | **0.2003** | **0.8981** | **63.45** | **0.4725** |
| Crotonlaevigatus | **0.5300*** | **79.40** | **12.07** | **0.2017** | **0.9022** | **62.06** | **0.4792** |
| ostodes | **0.3628** | **85.04** | **11.61** | **0.1895** | **0.8672** | **73.95** | **0.4220** |
| Ostodespaniculata | **0.3390*** | **85.85** | **11.54** | **0.1878** | **0.8622** | **75.65** | **0.4139** |
| suregada | **0.5980** | **77.10** | **12.26** | **0.2067** | **0.9164** | **57.22** | **0.5025** |
| Suregadamultiflora | **0.6470*** | **75.45** | **12.39** | **0.2103** | **0.9266** | **53.73** | **0.5193** |
| triadica | **0.5409** | **79.03** | **12.10** | **0.2025** | **0.9045** | **61.28** | **0.4830** |
| Triadicacochinchinensis | **0.5431*** | **78.96** | **12.10** | **0.2027** | **0.9049** | **61.12** | **0.4837** |
| endospermum | **0.3836** | **84.34** | **11.67** | **0.1910** | **0.8716** | **72.47** | **0.4291** |
| Endospermumchinense | **0.3475*** | **85.56** | **11.57** | **0.1884** | **0.8640** | **75.04** | **0.4168** |
| Trevia | **0.5450** | **78.89** | **12.11** | **0.2028** | **0.9053** | **60.99** | **0.4844** |
| Trevianudiflora | **0.5431*** | **78.96** | **12.10** | **0.2027** | **0.9049** | **61.12** | **0.4837** |
| Euphorbiaceaesp9 | **0.5431*** | **78.96** | **12.10** | **0.2027** | **0.9049** | **61.12** | **0.4837** |
| Lasiococca | **0.5450** | **78.89** | **12.11** | **0.2028** | **0.9053** | **60.99** | **0.4844** |
| Lasiococcacomberi | **0.5431*** | **78.96** | **12.10** | **0.2027** | **0.9049** | **61.12** | **0.4837** |
| Epiprinus | **0.5450** | **78.89** | **12.11** | **0.2028** | **0.9053** | **60.99** | **0.4844** |
| Epiprinussiletianus | **0.5431*** | **78.96** | **12.10** | **0.2027** | **0.9049** | **61.12** | **0.4837** |
| Euphorbiaceaesp11 | **0.5431*** | **78.96** | **12.10** | **0.2027** | **0.9049** | **61.12** | **0.4837** |
| Euphorbiaceaesp3 | **0.5431*** | **78.96** | **12.10** | **0.2027** | **0.9049** | **61.12** | **0.4837** |
| Euphorbiaceaesp4 | **0.5431*** | **78.96** | **12.10** | **0.2027** | **0.9049** | **61.12** | **0.4837** |
| Euphorbiaceaesp5 | **0.5431*** | **78.96** | **12.10** | **0.2027** | **0.9049** | **61.12** | **0.4837** |
| phyllanthaceae | **0.6010** | **77.00** | **12.26** | **0.2069** | **0.9170** | **57.00** | **0.5035** |
| antidesma | **0.6284** | **76.07** | **12.34** | **0.2089** | **0.9228** | **55.05** | **0.5129** |
| Antidesmamaclurei | **0.5900*** | **77.37** | **12.23** | **0.2061** | **0.9147** | **57.79** | **0.4998** |
| Antidesmamontanum | **0.5900*** | **77.37** | **12.23** | **0.2061** | **0.9147** | **57.79** | **0.4998** |
| Antidesmasp1 | **0.6575*** | **75.09** | **12.42** | **0.2111** | **0.9288** | **52.98** | **0.5229** |
| Antidesmasp2 | **0.6575*** | **75.09** | **12.42** | **0.2111** | **0.9288** | **52.98** | **0.5229** |
| Antidesmahainanense | **0.6575*** | **75.09** | **12.42** | **0.2111** | **0.9288** | **52.98** | **0.5229** |
| aporosa | **0.4232** | **83.01** | **11.77** | **0.1939** | **0.8798** | **69.66** | **0.4427** |
| Aporosadioica | **0.3700*** | **84.80** | **11.63** | **0.1901** | **0.8687** | **73.44** | **0.4245** |
| baccaurea | **0.5097** | **80.08** | **12.01** | **0.2003** | **0.8979** | **63.50** | **0.4723** |
| Baccaurearamiflora | **0.5431*** | **78.96** | **12.10** | **0.2027** | **0.9049** | **61.12** | **0.4837** |
| bischofia | **0.5772** | **77.80** | **12.20** | **0.2052** | **0.9120** | **58.70** | **0.4954** |
| Bischofiapolycarpa | **0.5699*** | **78.05** | **12.18** | **0.2047** | **0.9105** | **59.22** | **0.4929** |
| breynia | **0.6215** | **76.31** | **12.32** | **0.2084** | **0.9213** | **55.54** | **0.5105** |
| Breyniafruticosa | **0.6315*** | **75.97** | **12.35** | **0.2092** | **0.9234** | **54.83** | **0.5140** |

| | | | | | | | |
|---|---|---|---|---|---|---|---|
| Breyniarostrata | 0.6315* | 75.97 | 12.35 | 0.2092 | 0.9234 | 54.83 | 0.5140 |
| glochidion | 0.5486 | 78.77 | 12.12 | 0.2031 | 0.9061 | 60.73 | 0.4856 |
| Glochidioncoccineum | 0.5576* | 78.47 | 12.14 | 0.2038 | 0.9079 | 60.10 | 0.4887 |
| Glochidionhirsutum | 0.5576* | 78.47 | 12.14 | 0.2038 | 0.9079 | 60.10 | 0.4887 |
| Glochidionsphaerogynum | 0.5576* | 78.47 | 12.14 | 0.2038 | 0.9079 | 60.10 | 0.4887 |
| Glochidiontriandrum | 0.5576* | 78.47 | 12.14 | 0.2038 | 0.9079 | 60.10 | 0.4887 |
| Glochidionzeylanicum | 0.4800* | 81.09 | 11.93 | 0.1981 | 0.8917 | 65.61 | 0.4621 |
| phyllanthus | 0.6033 | 76.92 | 12.27 | 0.2071 | 0.9175 | 56.84 | 0.5043 |
| Phyllanthuspachyphyllus | 0.6126* | 76.61 | 12.30 | 0.2078 | 0.9194 | 56.18 | 0.5075 |
| bridelia | 0.6132 | 76.59 | 12.30 | 0.2078 | 0.9196 | 56.14 | 0.5077 |
| Brideliabalansae | 0.6053* | 76.86 | 12.28 | 0.2072 | 0.9179 | 56.70 | 0.5050 |
| cleistanthus | 0.6404 | 75.67 | 12.37 | 0.2098 | 0.9253 | 54.20 | 0.5170 |
| Cleistanthusconcinnus | 0.6518* | 75.28 | 12.40 | 0.2106 | 0.9276 | 53.39 | 0.5209 |
| leptopus | 0.6282 | 76.08 | 12.34 | 0.2089 | 0.9227 | 55.07 | 0.5128 |
| Leptopushainanensis | 0.6315* | 75.97 | 12.35 | 0.2092 | 0.9234 | 54.83 | 0.5140 |
| actephila | 0.6163 | 76.48 | 12.31 | 0.2080 | 0.9202 | 55.92 | 0.5088 |
| Actephilamerrilliana | 0.6315* | 75.97 | 12.35 | 0.2092 | 0.9234 | 54.83 | 0.5140 |
| ixonanthaceae | 0.6116 | 76.64 | 12.29 | 0.2077 | 0.9192 | 56.25 | 0.5072 |
| ixonanthes | 0.6191 | 76.39 | 12.31 | 0.2083 | 0.9208 | 55.72 | 0.5097 |
| Ixonanthesreticulata | 0.6265* | 76.14 | 12.33 | 0.2088 | 0.9224 | 55.19 | 0.5123 |
| salicaceae | 0.6564 | 75.13 | 12.42 | 0.2110 | 0.9286 | 53.06 | 0.5225 |
| casearia | 0.6394 | 75.70 | 12.37 | 0.2097 | 0.9251 | 54.27 | 0.5167 |
| Caseariamembranacea | 0.6500* | 75.34 | 12.40 | 0.2105 | 0.9273 | 53.52 | 0.5203 |
| Caseariavelutina | 0.6246* | 76.20 | 12.33 | 0.2087 | 0.9220 | 55.32 | 0.5116 |
| homalium | 0.6849 | 74.17 | 12.49 | 0.2131 | 0.9346 | 51.03 | 0.5322 |
| Homaliummollissimum | 0.6957* | 73.80 | 12.52 | 0.2139 | 0.9368 | 50.26 | 0.5360 |
| Homaliumpaniculiflorum | 0.6957* | 73.80 | 12.52 | 0.2139 | 0.9368 | 50.26 | 0.5360 |
| Homaliumphanerophlebium | 0.6400* | 75.68 | 12.37 | 0.2098 | 0.9252 | 54.23 | 0.5169 |
| Homaliumstenophyllum | 0.6957* | 73.80 | 12.52 | 0.2139 | 0.9368 | 50.26 | 0.5360 |
| scolopia | 0.8052 | 70.10 | 12.82 | 0.2218 | 0.9597 | 42.47 | 0.5734 |
| Scolopiasaeva | 0.8300* | 69.26 | 12.89 | 0.2237 | 0.9649 | 40.71 | 0.5819 |
| flacourtia | 0.7518 | 71.90 | 12.68 | 0.2180 | 0.9486 | 46.27 | 0.5552 |
| Flacourtiarukam | 0.7500* | 71.97 | 12.67 | 0.2178 | 0.9482 | 46.40 | 0.5545 |
| achariaceae | 0.6310 | 75.99 | 12.35 | 0.2091 | 0.9233 | 54.87 | 0.5138 |
| hydnocarpus | 0.6311 | 75.98 | 12.35 | 0.2091 | 0.9233 | 54.86 | 0.5138 |
| Hydnocarpushainanensis | 0.6313* | 75.98 | 12.35 | 0.2091 | 0.9234 | 54.85 | 0.5139 |
| hypericaceae | 0.6854 | 74.15 | 12.50 | 0.2131 | 0.9347 | 51.00 | 0.5324 |
| cratoxylum | 0.6915 | 73.94 | 12.51 | 0.2135 | 0.9359 | 50.56 | 0.5345 |
| Cratoxylumcochinchinense | 0.6700* | 74.67 | 12.45 | 0.2120 | 0.9315 | 52.09 | 0.5271 |
| Cratoxylumformosum | 0.7150* | 73.15 | 12.58 | 0.2153 | 0.9409 | 48.89 | 0.5425 |
| calophyllaceae | 0.6797 | 74.34 | 12.48 | 0.2127 | 0.9335 | 51.41 | 0.5305 |
| calophyllum | 0.6700 | 74.67 | 12.45 | 0.2120 | 0.9315 | 52.09 | 0.5271 |
| Calophyllummembranaceum | 0.6692* | 74.70 | 12.45 | 0.2119 | 0.9313 | 52.15 | 0.5269 |
| Calophyllumsp1 | 0.6692* | 74.70 | 12.45 | 0.2119 | 0.9313 | 52.15 | 0.5269 |
| clusiaceae | 0.6745 | 74.52 | 12.47 | 0.2123 | 0.9324 | 51.77 | 0.5287 |
| garcinia | 0.6831 | 74.23 | 12.49 | 0.2129 | 0.9342 | 51.16 | 0.5316 |
| Garciniamultiflora | 0.7416* | 72.25 | 12.65 | 0.2172 | 0.9464 | 47.00 | 0.5517 |
| Garciniaoblongifolia | 0.6268* | 76.13 | 12.33 | 0.2088 | 0.9224 | 55.17 | 0.5124 |
| Clusiaceaesp1 | 0.6692* | 74.70 | 12.45 | 0.2119 | 0.9313 | 52.15 | 0.5269 |
| ochnaceae | 0.7051 | 73.48 | 12.55 | 0.2145 | 0.9388 | 49.60 | 0.5391 |
| ochna | 0.7375 | 72.39 | 12.64 | 0.2169 | 0.9456 | 47.29 | 0.5502 |
| Ochnaintegerrima | 0.7440* | 72.17 | 12.66 | 0.2174 | 0.9469 | 46.83 | 0.5525 |
| Campylospermum | 0.7201 | 72.98 | 12.59 | 0.2156 | 0.9419 | 48.53 | 0.5443 |
| Campylospermumserratum | 0.7351* | 72.47 | 12.63 | 0.2167 | 0.9451 | 47.46 | 0.5494 |
| rhizophoraceae | 0.6974 | 73.74 | 12.53 | 0.2140 | 0.9372 | 50.14 | 0.5365 |
| carallia | 0.6737 | 74.54 | 12.46 | 0.2122 | 0.9322 | 51.83 | 0.5284 |
| Caralliabrachiata | 0.6659* | 74.81 | 12.44 | 0.2117 | 0.9306 | 52.39 | 0.5257 |
| erythroxylaceae | 0.7334 | 72.53 | 12.63 | 0.2166 | 0.9447 | 47.58 | 0.5488 |
| erythroxylum | 0.7615 | 71.58 | 12.70 | 0.2187 | 0.9506 | 45.58 | 0.5585 |
| Erythroxylumsinense | 0.7896* | 70.63 | 12.78 | 0.2207 | 0.9565 | 43.58 | 0.5681 |
| pandaceae | 0.6324 | 75.94 | 12.35 | 0.2092 | 0.9236 | 54.77 | 0.5143 |
| microdesmis | 0.6162 | 76.49 | 12.31 | 0.2080 | 0.9202 | 55.92 | 0.5087 |
| Microdesmiscaseariifolia | 0.6000* | 77.03 | 12.26 | 0.2069 | 0.9168 | 57.07 | 0.5032 |
| putranjivaceae | 0.6839 | 74.20 | 12.49 | 0.2130 | 0.9343 | 51.11 | 0.5319 |

| | | | | | | | |
|---|---|---|---|---|---|---|---|
| drypetes | 0.6978 | 73.73 | 12.53 | 0.2140 | 0.9373 | 50.11 | 0.5367 |
| Drypetescumingii | 0.6945* | 73.84 | 12.52 | 0.2138 | 0.9366 | 50.35 | 0.5355 |
| Drypeteshainanensis | 0.7183* | 73.04 | 12.59 | 0.2155 | 0.9416 | 48.65 | 0.5437 |
| Drypetesperreticulata | 0.6945* | 73.84 | 12.52 | 0.2138 | 0.9366 | 50.35 | 0.5355 |
| oxalidales | 0.6122 | 76.62 | 12.29 | 0.2077 | 0.9194 | 56.21 | 0.5073 |
| elaeocarpaceae | 0.5533 | 78.61 | 12.13 | 0.2034 | 0.9070 | 60.40 | 0.4872 |
| elaeocarpus | 0.4924 | 80.67 | 11.96 | 0.1990 | 0.8943 | 64.73 | 0.4664 |
| Elaeocarpusangustifolius | 0.4029* | 83.69 | 11.72 | 0.1925 | 0.8756 | 71.10 | 0.4357 |
| Elaeocarpusdubius | 0.5240* | 79.60 | 12.05 | 0.2013 | 0.9009 | 62.48 | 0.4772 |
| Elaeocarpusglabripetalus | 0.5032* | 80.30 | 11.99 | 0.1998 | 0.8966 | 63.96 | 0.4701 |
| Elaeocarpushowii | 0.5032* | 80.30 | 11.99 | 0.1998 | 0.8966 | 63.96 | 0.4701 |
| Elaeocarpusjaponicus | 0.5032* | 80.30 | 11.99 | 0.1998 | 0.8966 | 63.96 | 0.4701 |
| Elaeocarpuslimitaneus | 0.5032* | 80.30 | 11.99 | 0.1998 | 0.8966 | 63.96 | 0.4701 |
| Elaeocarpusnitentifolius | 0.5032* | 80.30 | 11.99 | 0.1998 | 0.8966 | 63.96 | 0.4701 |
| Elaeocarpuspoilanei | 0.5032* | 80.30 | 11.99 | 0.1998 | 0.8966 | 63.96 | 0.4701 |
| Elaeocarpussylvestris | 0.4750* | 81.26 | 11.92 | 0.1977 | 0.8907 | 65.97 | 0.4604 |
| sloanea | 0.5458 | 78.86 | 12.11 | 0.2029 | 0.9055 | 60.93 | 0.4847 |
| Sloaneaintegrifolia | 0.6095* | 76.71 | 12.29 | 0.2075 | 0.9188 | 56.40 | 0.5064 |
| Sloaneasinensis | 0.4785* | 81.14 | 11.93 | 0.1980 | 0.8914 | 65.72 | 0.4616 |
| connaraceae | 0.5881 | 77.43 | 12.23 | 0.2060 | 0.9143 | 57.92 | 0.4991 |
| ellipanthus | 0.5847 | 77.55 | 12.22 | 0.2057 | 0.9136 | 58.16 | 0.4980 |
| Ellipanthusglabrifolius | 0.5814* | 77.66 | 12.21 | 0.2055 | 0.9129 | 58.40 | 0.4968 |
| celastrales | 0.6422 | 75.61 | 12.38 | 0.2099 | 0.9256 | 54.07 | 0.5176 |
| celastraceae | 0.6503 | 75.34 | 12.40 | 0.2105 | 0.9273 | 53.50 | 0.5204 |
| Celastraceaesp2 | 0.6797* | 74.34 | 12.48 | 0.2127 | 0.9335 | 51.41 | 0.5304 |
| Celastraceaesp3 | 0.6797* | 74.34 | 12.48 | 0.2127 | 0.9335 | 51.41 | 0.5304 |
| salacia | 0.7478 | 72.04 | 12.67 | 0.2177 | 0.9477 | 46.56 | 0.5538 |
| Salaciachinensis | 0.7600* | 71.63 | 12.70 | 0.2185 | 0.9503 | 45.69 | 0.5579 |
| euonymus | 0.5874 | 77.46 | 12.23 | 0.2059 | 0.9142 | 57.97 | 0.4989 |
| Euonymusgibber | 0.5665* | 78.17 | 12.17 | 0.2044 | 0.9098 | 59.46 | 0.4917 |
| Euonymuslaxiflorus | 0.5665* | 78.17 | 12.17 | 0.2044 | 0.9098 | 59.46 | 0.4917 |
| Euonymusnitidus | 0.5665* | 78.17 | 12.17 | 0.2044 | 0.9098 | 59.46 | 0.4917 |
| myrtales | 0.6326 | 87.57 | 12.57 | 0.2053 | 0.9560 | 54.43 | 0.4987 |
| melastomataceae | 0.6854 | 85.78 | 12.72 | 0.2092 | 0.9671 | 50.67 | 0.5168 |
| blastus | 0.6081 | 88.39 | 12.50 | 0.2035 | 0.9509 | 56.17 | 0.4904 |
| Blastuscochinchinensis | 0.6094* | 88.35 | 12.51 | 0.2036 | 0.9512 | 56.08 | 0.4908 |
| melastoma | 0.4452 | 93.90 | 12.06 | 0.1916 | 0.9168 | 67.77 | 0.4346 |
| Melastomamalabathricum | 0.4400* | 94.07 | 12.04 | 0.1912 | 0.9157 | 68.13 | 0.4328 |
| Melastomapenicillatum | 0.4400* | 94.07 | 12.04 | 0.1912 | 0.9157 | 68.13 | 0.4328 |
| Melastomasanguineum | 0.4400* | 94.07 | 12.04 | 0.1912 | 0.9157 | 68.13 | 0.4328 |
| medinilla | 0.6142 | 88.19 | 12.52 | 0.2040 | 0.9522 | 55.74 | 0.4925 |
| Medinillaassamica | 0.6094* | 88.35 | 12.51 | 0.2036 | 0.9512 | 56.08 | 0.4908 |
| Allomorphia | 0.6474 | 87.07 | 12.61 | 0.2064 | 0.9591 | 53.37 | 0.5038 |
| Allomorphiabalansae | 0.6094* | 88.35 | 12.51 | 0.2036 | 0.9512 | 56.08 | 0.4908 |
| memecylon | 0.7510 | 83.57 | 12.90 | 0.2140 | 0.9808 | 46.00 | 0.5393 |
| Memecylonhainanense | 0.7728* | 82.83 | 12.96 | 0.2155 | 0.9853 | 44.45 | 0.5467 |
| Memecylonligustrifolium | 0.7728* | 82.83 | 12.96 | 0.2155 | 0.9853 | 44.45 | 0.5467 |
| Memecylonnigrescens | 0.7728* | 82.83 | 12.96 | 0.2155 | 0.9853 | 44.45 | 0.5467 |
| myrtaceae | 0.6869 | 85.73 | 12.72 | 0.2093 | 0.9674 | 50.56 | 0.5173 |
| rhodamnia | 0.8244 | 81.09 | 13.10 | 0.2193 | 0.9961 | 40.78 | 0.5644 |
| Rhodamniadumetorum | 0.8787* | 79.25 | 13.25 | 0.2233 | 1.007 | 36.91 | 0.5830 |
| rhodomyrtus | 0.7069 | 85.06 | 12.78 | 0.2107 | 0.9716 | 49.14 | 0.5242 |
| Rhodomyrtustomentosa | 0.6797* | 85.98 | 12.70 | 0.2087 | 0.9659 | 51.08 | 0.5149 |
| decaspermum | 0.7256 | 84.42 | 12.83 | 0.2121 | 0.9755 | 47.81 | 0.5306 |
| Decaspermummontanum | 0.7170* | 84.72 | 12.80 | 0.2115 | 0.9737 | 48.42 | 0.5276 |
| syzygium | 0.6693 | 86.33 | 12.67 | 0.2080 | 0.9637 | 51.82 | 0.5113 |
| Syzygiumacuminatissimum | 0.6644* | 86.49 | 12.66 | 0.2076 | 0.9627 | 52.17 | 0.5096 |
| Syzygiumaraiocladum | 0.7360* | 84.07 | 12.86 | 0.2129 | 0.9776 | 47.07 | 0.5341 |
| Syzygiumbullockii | 0.6644* | 86.49 | 12.66 | 0.2076 | 0.9627 | 52.17 | 0.5096 |
| Syzygiumbuxifolium | 0.6644* | 86.49 | 12.66 | 0.2076 | 0.9627 | 52.17 | 0.5096 |
| Syzygiumchampionii | 0.6644* | 86.49 | 12.66 | 0.2076 | 0.9627 | 52.17 | 0.5096 |
| Syzygiumchunianum | 0.6644* | 86.49 | 12.66 | 0.2076 | 0.9627 | 52.17 | 0.5096 |
| Syzygiumclaviflorum | 0.6235* | 87.87 | 12.55 | 0.2046 | 0.9541 | 55.07 | 0.4956 |
| Syzygiumcumini | 0.6727* | 86.21 | 12.68 | 0.2082 | 0.9644 | 51.58 | 0.5125 |

| | | | | | | | |
|---|---|---|---|---|---|---|---|
| Syzygiumglobiflorum | 0.6644* | 86.49 | 12.66 | 0.2076 | 0.9627 | 52.17 | 0.5096 |
| Syzygiumhancei | 0.6644* | 86.49 | 12.66 | 0.2076 | 0.9627 | 52.17 | 0.5096 |
| Syzygiumjambos | 0.7000* | 85.29 | 12.76 | 0.2102 | 0.9701 | 49.63 | 0.5218 |
| Syzygiumjienfunicum | 0.6644* | 86.49 | 12.66 | 0.2076 | 0.9627 | 52.17 | 0.5096 |
| Syzygiumlevinei | 0.6644* | 86.49 | 12.66 | 0.2076 | 0.9627 | 52.17 | 0.5096 |
| Syzygiumodoratum | 0.6644* | 86.49 | 12.66 | 0.2076 | 0.9627 | 52.17 | 0.5096 |
| Syzygiumrehderianum | 0.6644* | 86.49 | 12.66 | 0.2076 | 0.9627 | 52.17 | 0.5096 |
| Syzygiumrysopodum | 0.6644* | 86.49 | 12.66 | 0.2076 | 0.9627 | 52.17 | 0.5096 |
| Syzygiumsp | 0.6792* | 85.99 | 12.70 | 0.2087 | 0.9658 | 51.11 | 0.5147 |
| Syzygiumsterrophyllum | 0.6644* | 86.49 | 12.66 | 0.2076 | 0.9627 | 52.17 | 0.5096 |
| Syzygiumtephrodes | 0.6644* | 86.49 | 12.66 | 0.2076 | 0.9627 | 52.17 | 0.5096 |
| Syzygiumtsoongii | 0.6644* | 86.49 | 12.66 | 0.2076 | 0.9627 | 52.17 | 0.5096 |
| sapindales | 0.5606 | 111.8 | 12.65 | 0.1868 | 0.9552 | 63.66 | 0.4601 |
| meliaceae | 0.5769 | 122.8 | 13.36 | 0.1871 | 0.9236 | 67.73 | 0.4062 |
| aphanamixis | 0.5849 | 122.5 | 13.38 | 0.1877 | 0.9253 | 67.16 | 0.4089 |
| Aphanamixispolystachya | 0.5765* | 122.8 | 13.36 | 0.1871 | 0.9235 | 67.76 | 0.4060 |
| aglaia | 0.5994 | 122.0 | 13.42 | 0.1888 | 0.9283 | 66.12 | 0.4139 |
| Aglaiaelaeagnoidea | 0.6300* | 121.0 | 13.51 | 0.1910 | 0.9347 | 63.95 | 0.4243 |
| Aglaiaspectabilis | 0.5750* | 122.9 | 13.35 | 0.1870 | 0.9232 | 67.86 | 0.4055 |
| dysoxylum | 0.5722 | 123.0 | 13.35 | 0.1868 | 0.9226 | 68.07 | 0.4045 |
| Dysoxylumgotadhora | 0.5974* | 122.1 | 13.42 | 0.1886 | 0.9279 | 66.27 | 0.4132 |
| Dysoxylummollissimum | 0.5189* | 124.8 | 13.20 | 0.1829 | 0.9115 | 71.86 | 0.3863 |
| walsura | 0.8326 | 114.2 | 14.06 | 0.2058 | 0.9771 | 49.53 | 0.4937 |
| Walsurapinnata | 0.8680* | 113.0 | 14.16 | 0.2084 | 0.9845 | 47.01 | 0.5058 |
| Walsurarobusta | 0.8680* | 113.0 | 14.16 | 0.2084 | 0.9845 | 47.01 | 0.5058 |
| melia | 0.4818 | 126.0 | 13.10 | 0.1802 | 0.9037 | 74.50 | 0.3736 |
| Meliaazedarach | 0.4378* | 127.5 | 12.98 | 0.1769 | 0.8945 | 77.63 | 0.3586 |
| Heynea | 0.5856 | 122.5 | 13.38 | 0.1877 | 0.9254 | 67.11 | 0.4091 |
| Heyneatrijuga | 0.5944* | 122.2 | 13.41 | 0.1884 | 0.9273 | 66.49 | 0.4121 |
| Meliaceaesp1 | 0.5944* | 122.2 | 13.41 | 0.1884 | 0.9273 | 66.49 | 0.4121 |
| simaroubaceae | 0.5340 | 124.2 | 13.24 | 0.1840 | 0.9146 | 70.78 | 0.3915 |
| brucea | 0.4532 | 127.0 | 13.02 | 0.1781 | 0.8978 | 76.53 | 0.3638 |
| Bruceamollis | 0.4330* | 127.7 | 12.96 | 0.1766 | 0.8935 | 77.97 | 0.3569 |
| rutaceae | 0.5497 | 122.2 | 13.69 | 0.1878 | 0.9021 | 71.36 | 0.3645 |
| murraya | 0.6589 | 109.8 | 14.29 | 0.1976 | 0.9130 | 64.05 | 0.4083 |
| Murrayaalata | 0.7537* | 106.6 | 14.55 | 0.2046 | 0.9328 | 57.30 | 0.4407 |
| glycosmis | 0.4703 | 116.1 | 13.77 | 0.1838 | 0.8735 | 77.47 | 0.3437 |
| Glycosmissp1 | 0.4390* | 117.2 | 13.69 | 0.1816 | 0.8670 | 79.70 | 0.3330 |
| Glycosmiscochinchinensis | 0.4390* | 117.2 | 13.69 | 0.1816 | 0.8670 | 79.70 | 0.3330 |
| Glycosmiscraibii | 0.4390* | 117.2 | 13.69 | 0.1816 | 0.8670 | 79.70 | 0.3330 |
| clausena | 0.5231 | 114.3 | 13.92 | 0.1877 | 0.8846 | 73.72 | 0.3618 |
| Clausenaexcavata | 0.4820* | 115.7 | 13.81 | 0.1847 | 0.8760 | 76.64 | 0.3477 |
| micromelum | 0.6121 | 111.3 | 14.16 | 0.1942 | 0.9032 | 67.38 | 0.3922 |
| Micromelumfalcatum | 0.6600* | 109.7 | 14.30 | 0.1977 | 0.9132 | 63.97 | 0.4087 |
| tetradium | 0.2887 | 163.5 | 11.91 | 0.2535 | 0.6539 | 137.0 | 0.3369 |
| Tetradiumglabrifolium | 0.2320* | 177.9* | 11.34* | 0.2697* | 0.5995* | 152.6* | 0.3298* |
| acronychia | 0.4687 | 84.00 | 17.01 | 0.2085 | 0.9436 | 70.10 | 0.2989 |
| Acronychiapedunculata | 0.4587* | 83.94* | 17.38* | 0.2197* | 0.9998* | 64.12* | 0.2804* |
| melicope | 0.4921 | 78.75 | 16.44 | 0.1741 | 0.7653 | 89.87 | 0.3472 |
| Melicopechunii | 0.4953* | 73.38* | 16.59* | 0.1621* | 0.6993* | 97.69* | 0.3584* |
| zanthoxylum | 0.5667 | 92.83 | 14.36 | 0.1737 | 0.9267 | 58.40 | 0.3792 |
| Zanthoxylumavicennae | 0.6177* | 79.86* | 14.53* | 0.1584* | 0.9823* | 42.09* | 0.3927* |
| Maclurodendron | 0.5499 | 125.2 | 13.95 | 0.1895 | 0.8924 | 72.81 | 0.3290 |
| Maclurodendronoligophlebium | 0.5500* | 128.1* | 14.21* | 0.1912* | 0.8826* | 74.26* | 0.2936* |
| sapindaceae | 0.5646 | 128.3 | 12.62 | 0.1840 | 0.9263 | 66.69 | 0.4465 |
| mischocarpus | 0.7465 | 154.7 | 10.46 | 0.2016 | 0.8693 | 51.77 | 0.6184 |
| Mischocarpushainanensis | 0.7307* | 155.2 | 10.42 | 0.2005 | 0.8660 | 52.90 | 0.6130 |
| Mischocarpuspentapetalus | 0.7307* | 155.2 | 10.42 | 0.2005 | 0.8660 | 52.90 | 0.6130 |
| Mischocarpussundaicus | 0.7800* | 153.6 | 10.56 | 0.2041 | 0.8763 | 49.39 | 0.6298 |
| nephelium | 0.7724 | 171.9 | 9.057 | 0.2059 | 0.8219 | 48.83 | 0.6881 |
| Nepheliumtopengii | 0.7782* | 175.3* | 8.778* | 0.2068* | 0.8126* | 48.20* | 0.7023* |
| dimocarpus | 0.7344 | 166.0 | 9.544 | 0.2022 | 0.8351 | 51.97 | 0.6508 |
| Dimocarpuslongan | 0.7000* | 167.1 | 9.449 | 0.1997 | 0.8279 | 54.42 | 0.6390 |

| | | | | | | | |
|---|---|---|---|---|---|---|---|
| litchi | 0.8115 | 163.4 | 9.755 | 0.2078 | 0.8512 | 46.48 | 0.6772 |
| Litchichinensis | 0.8541* | 161.9 | 9.873 | 0.2109 | 0.8601 | 43.45 | 0.6918 |
| lepisanthes | 0.6471 | 161.7 | 9.894 | 0.1949 | 0.8380 | 58.62 | 0.5965 |
| Lepisanthesrubiginosa | 0.6300* | 162.3 | 9.847 | 0.1936 | 0.8344 | 59.84 | 0.5907 |
| amesiodendron | 0.8166 | 141.5 | 11.54 | 0.2053 | 0.9156 | 47.44 | 0.6058 |
| Amesiodendronchinense | 0.8345* | 140.9 | 11.59 | 0.2066 | 0.9194 | 46.17 | 0.6119 |
| paranephelium | 0.8037 | 141.9 | 11.51 | 0.2044 | 0.9129 | 48.36 | 0.6014 |
| Paranepheliumhainanense | 0.8267* | 141.1 | 11.57 | 0.2061 | 0.9177 | 46.72 | 0.6093 |
| acer | 0.4879 | 138.1 | 11.82 | 0.1794 | 0.8891 | 71.71 | 0.4446 |
| Acerfabri | 0.5145* | 137.2 | 11.89 | 0.1813 | 0.8947 | 69.81 | 0.4537 |
| Acerlaurinum | 0.4300* | 140.1 | 11.66 | 0.1752 | 0.8770 | 75.83 | 0.4248 |
| anacardiaceae | 0.5240 | 130.6 | 12.45 | 0.1654 | 0.9849 | 68.94 | 0.4800 |
| toxicodendron | 0.5620 | 130.6 | 11.20 | 0.1758 | 0.7264 | 79.83 | 0.5484 |
| Toxicodendronvernicifluum | 0.5662* | 130.6* | 11.06* | 0.1769* | 0.6976* | 81.04* | 0.5560* |
| choerospondias | 0.4932 | 131.7 | 12.37 | 0.1632 | 0.9785 | 71.13 | 0.4695 |
| Choerospondiasaxillaris | 0.4870* | 131.9 | 12.35 | 0.1627 | 0.9772 | 71.57 | 0.4673 |
| burseraceae | 0.5159 | 133.9 | 12.69 | 0.1567 | 1.072 | 66.05 | 0.4816 |
| canarium | 0.4860 | 144.4 | 12.93 | 0.1327 | 1.245 | 62.31 | 0.5029 |
| Canariumalbum | 0.4170* | 162.5* | 10.45* | 0.1150* | 0.8478* | 79.27* | 0.3229* |
| Canariumpimela | 0.5450* | 129.9* | 15.49* | 0.1423* | 1.699* | 44.09* | 0.6899* |
| malvales | 0.5699 | 106.0 | 12.61 | 0.1908 | 0.9536 | 61.97 | 0.4668 |
| thymelaeaceae | 0.5296 | 107.4 | 12.50 | 0.1878 | 0.9452 | 64.84 | 0.4530 |
| aquilaria | 0.4134 | 111.3 | 12.18 | 0.1793 | 0.9209 | 73.11 | 0.4132 |
| Aquilariasinensis | 0.3660* | 112.9 | 12.05 | 0.1759 | 0.9110 | 76.48 | 0.3970 |
| wikstroemia | 0.5262 | 107.5 | 12.49 | 0.1876 | 0.9445 | 65.09 | 0.4518 |
| Wikstroemiahainanensis | 0.5294* | 107.4 | 12.50 | 0.1878 | 0.9451 | 64.85 | 0.4529 |
| Wikstroemiaindica | 0.5294* | 107.4 | 12.50 | 0.1878 | 0.9451 | 64.85 | 0.4529 |
| Wikstroemianutans | 0.5294* | 107.4 | 12.50 | 0.1878 | 0.9451 | 64.85 | 0.4529 |
| Wikstroemiapachyrachis | 0.5294* | 107.4 | 12.50 | 0.1878 | 0.9451 | 64.85 | 0.4529 |
| malvaceae | 0.5740 | 105.9 | 12.62 | 0.1911 | 0.9545 | 61.68 | 0.4682 |
| reevesia | 0.4716 | 109.3 | 12.34 | 0.1836 | 0.9330 | 68.97 | 0.4331 |
| Reevesialancifolia | 0.5075* | 108.1 | 12.44 | 0.1862 | 0.9406 | 66.41 | 0.4454 |
| Reevesiathyrsoidea | 0.4350* | 110.6 | 12.24 | 0.1809 | 0.9254 | 71.57 | 0.4206 |
| sterculia | 0.4567 | 109.8 | 12.30 | 0.1825 | 0.9299 | 70.03 | 0.4281 |
| Sterculiahainanensis | 0.4048* | 111.6 | 12.15 | 0.1787 | 0.9191 | 73.72 | 0.4103 |
| Sterculialanceolata | 0.5000* | 108.4 | 12.42 | 0.1857 | 0.9390 | 66.95 | 0.4429 |
| pterospermum | 0.4925 | 108.6 | 12.40 | 0.1851 | 0.9374 | 67.48 | 0.4403 |
| Pterospermumheterophyllum | 0.4460* | 110.2 | 12.27 | 0.1817 | 0.9277 | 70.79 | 0.4244 |
| Pterospermumlanceifolium | 0.5146* | 107.9 | 12.46 | 0.1867 | 0.9420 | 65.91 | 0.4478 |
| Pterospermumxiaoye | 0.5146* | 107.9 | 12.46 | 0.1867 | 0.9420 | 65.91 | 0.4478 |
| microcos | 0.4888 | 108.8 | 12.39 | 0.1848 | 0.9366 | 67.75 | 0.4390 |
| Micrococschungii | 0.4819* | 109.0 | 12.37 | 0.1843 | 0.9352 | 68.24 | 0.4367 |
| dipterocarpaceae | 0.7336 | 100.5 | 13.06 | 0.2027 | 0.9878 | 50.32 | 0.5228 |
| vatica | 0.7577 | 99.68 | 13.12 | 0.2045 | 0.9929 | 48.61 | 0.5310 |
| Vaticamangachapoi | 0.7500* | 99.94 | 13.10 | 0.2039 | 0.9913 | 49.16 | 0.5284 |
| hopea | 0.8353 | 97.05 | 13.34 | 0.2102 | 1.009 | 43.08 | 0.5576 |
| Hopeahainanensis | 0.8900* | 95.21 | 13.49 | 0.2142 | 1.021 | 39.19 | 0.5763 |
| brassicales | 0.5866 | 105.5 | 12.65 | 0.1920 | 0.9571 | 60.79 | 0.4725 |
| capparaceae | 0.6725 | 102.6 | 12.89 | 0.1983 | 0.9751 | 54.67 | 0.5019 |
| Capparaceaesp1 | 0.6832* | 102.2 | 12.92 | 0.1991 | 0.9773 | 53.91 | 0.5056 |
| crossosomatales | 0.5384 | 96.19 | 12.38 | 0.1951 | 0.9399 | 62.16 | 0.4630 |
| staphyleaceae | 0.4807 | 98.14 | 12.22 | 0.1909 | 0.9278 | 66.27 | 0.4432 |
| turpinia | 0.4229 | 100.1 | 12.07 | 0.1867 | 0.9157 | 70.38 | 0.4235 |
| Turpiniamontana | 0.3940* | 101.1 | 11.99 | 0.1846 | 0.9097 | 72.44 | 0.4136 |
| saxifragales | 0.6081 | 78.25 | 12.45 | 0.2128 | 0.9690 | 51.73 | 0.4888 |
| hamamelidaceae | 0.5996 | 78.54 | 12.42 | 0.2122 | 0.9673 | 52.33 | 0.4859 |
| eustigma | 0.6330 | 77.41 | 12.52 | 0.2147 | 0.9742 | 49.96 | 0.4973 |
| Eustigmaoblongifolium | 0.6372* | 77.27 | 12.53 | 0.2150 | 0.9751 | 49.66 | 0.4987 |
| exbucklandia | 0.5715 | 79.49 | 12.35 | 0.2102 | 0.9614 | 54.34 | 0.4763 |
| Exbucklandiatonkinensis | 0.5480* | 80.28 | 12.28 | 0.2085 | 0.9565 | 56.01 | 0.4682 |
| chunia | 0.6161 | 77.98 | 12.47 | 0.2134 | 0.9707 | 51.16 | 0.4915 |
| Chuniabucklandioides | 0.6372* | 77.27 | 12.53 | 0.2150 | 0.9751 | 49.66 | 0.4987 |
| daphniphyllaceae | 0.5520 | 80.14 | 12.29 | 0.2087 | 0.9573 | 55.72 | 0.4696 |
| daphniphyllum | 0.5292 | 80.92 | 12.23 | 0.2071 | 0.9525 | 57.35 | 0.4618 |

| | | | | | | | |
|---|---|---|---|---|---|---|---|
| Daphniphyllumcalycinum | 0.5063* | 81.69 | 12.17 | 0.2054 | 0.9477 | 58.98 | 0.4539 |
| altingiaceae | 0.6460 | 76.97 | 12.55 | 0.2156 | 0.9770 | 49.04 | 0.5017 |
| altingia | 0.6731 | 76.05 | 12.63 | 0.2176 | 0.9826 | 47.11 | 0.5110 |
| Altingiaobovata | 0.7002* | 75.14 | 12.70 | 0.2196 | 0.9883 | 45.17 | 0.5203 |
| iteaceae | 0.5875 | 78.94 | 12.39 | 0.2113 | 0.9647 | 53.20 | 0.4817 |
| itea | 0.5834 | 79.08 | 12.38 | 0.2110 | 0.9639 | 53.49 | 0.4803 |
| Iteamacrophylla | 0.5814* | 79.15 | 12.37 | 0.2109 | 0.9634 | 53.63 | 0.4796 |
| Iteaxiaoye | 0.5814* | 79.15 | 12.37 | 0.2109 | 0.9634 | 53.63 | 0.4796 |
| saxifragaceae | 0.5848 | 79.04 | 12.38 | 0.2111 | 0.9642 | 53.39 | 0.4808 |
| Saxifragaceaesp1 | 0.5814* | 79.15 | 12.37 | 0.2109 | 0.9634 | 53.63 | 0.4796 |
| dilleniales | 0.6104 | 75.82 | 12.46 | 0.2160 | 0.9804 | 49.86 | 0.4870 |
| dilleniaceae | 0.6094 | 75.86 | 12.46 | 0.2159 | 0.9801 | 49.94 | 0.4867 |
| dillenia | 0.6063 | 75.96 | 12.45 | 0.2157 | 0.9795 | 50.15 | 0.4856 |
| Dilleniapentagyna | 0.5953* | 76.33 | 12.42 | 0.2149 | 0.9772 | 50.94 | 0.4819 |
| Dilleniaturbinata | 0.6162* | 75.62 | 12.48 | 0.2164 | 0.9816 | 49.45 | 0.4890 |
| gentianales | 0.6145 | 66.54 | 12.78 | 0.2270 | 0.9441 | 47.32 | 0.4996 |
| rubiaceae | 0.6438 | 65.55 | 12.86 | 0.2292 | 0.9502 | 45.24 | 0.5096 |
| Benkara | 0.6406 | 65.66 | 12.85 | 0.2289 | 0.9496 | 45.47 | 0.5085 |
| Benkarahainanensis | 0.6374* | 65.77 | 12.84 | 0.2287 | 0.9489 | 45.69 | 0.5075 |
| antirhea | 0.6404 | 65.66 | 12.85 | 0.2289 | 0.9495 | 45.48 | 0.5085 |
| Antirheachinensis | 0.6374* | 65.77 | 12.84 | 0.2287 | 0.9489 | 45.69 | 0.5075 |
| pertusadina | 0.6847 | 64.17 | 12.97 | 0.2322 | 0.9588 | 42.32 | 0.5237 |
| Pertusadinametcalfii | 0.6800* | 64.33 | 12.96 | 0.2318 | 0.9578 | 42.66 | 0.5220 |
| adina | 0.7215 | 62.92 | 13.08 | 0.2349 | 0.9665 | 39.71 | 0.5362 |
| Adinarubella | 0.7351* | 62.46 | 13.11 | 0.2359 | 0.9693 | 38.74 | 0.5409 |
| nauclea | 0.6423 | 65.60 | 12.86 | 0.2291 | 0.9499 | 45.35 | 0.5091 |
| naucleaofficinalis | 0.6374* | 65.77 | 12.84 | 0.2287 | 0.9489 | 45.69 | 0.5075 |
| diplospora | 0.7015 | 63.60 | 13.02 | 0.2334 | 0.9623 | 41.13 | 0.5294 |
| Diplosporadubia | 0.7000* | 63.65 | 13.02 | 0.2333 | 0.9620 | 41.24 | 0.5289 |
| catunaregam | 0.6913 | 63.94 | 12.99 | 0.2327 | 0.9602 | 41.86 | 0.5259 |
| Catunaregamspinosa | 0.6880* | 64.05 | 12.98 | 0.2324 | 0.9595 | 42.09 | 0.5248 |
| gardenia | 0.6722 | 64.59 | 12.94 | 0.2313 | 0.9562 | 43.22 | 0.5194 |
| Gardeniahainanensis | 0.6681* | 64.73 | 12.93 | 0.2310 | 0.9553 | 43.51 | 0.5180 |
| Gardeniasootepensis | 0.6681* | 64.73 | 12.93 | 0.2310 | 0.9553 | 43.51 | 0.5180 |
| aidia | 0.7404 | 62.29 | 13.13 | 0.2362 | 0.9704 | 38.36 | 0.5427 |
| Aidiacanthioides | 0.7527* | 61.87 | 13.16 | 0.2371 | 0.9730 | 37.49 | 0.5469 |
| Aidiapycnantha | 0.7527* | 61.87 | 13.16 | 0.2371 | 0.9730 | 37.49 | 0.5469 |
| pavetta | 0.6447 | 65.52 | 12.86 | 0.2292 | 0.9504 | 45.17 | 0.5100 |
| Pavettaarenosa | 0.6374* | 65.77 | 12.84 | 0.2287 | 0.9489 | 45.69 | 0.5075 |
| Pavettahongkongensis | 0.6374* | 65.77 | 12.84 | 0.2287 | 0.9489 | 45.69 | 0.5075 |
| tarenna | 0.6783 | 64.38 | 12.96 | 0.2317 | 0.9575 | 42.78 | 0.5215 |
| Tarennaattenuata | 0.6798* | 64.33 | 12.96 | 0.2318 | 0.9578 | 42.68 | 0.5220 |
| Tarennalancilimba | 0.6798* | 64.33 | 12.96 | 0.2318 | 0.9578 | 42.68 | 0.5220 |
| Tarennatsangii | 0.6798* | 64.33 | 12.96 | 0.2318 | 0.9578 | 42.68 | 0.5220 |
| ixora | 0.7755 | 61.10 | 13.22 | 0.2388 | 0.9778 | 35.86 | 0.5547 |
| Ixoranienkui | 0.7929* | 60.51 | 13.27 | 0.2401 | 0.9814 | 34.63 | 0.5607 |
| canthium | 0.6477 | 65.42 | 12.87 | 0.2295 | 0.9511 | 44.96 | 0.5110 |
| Canthiumhorridum | 0.6367* | 65.79 | 12.84 | 0.2287 | 0.9488 | 45.74 | 0.5072 |
| Canthiumsimile | 0.6367* | 65.79 | 12.84 | 0.2287 | 0.9488 | 45.74 | 0.5072 |
| psydrax | 0.7505 | 61.94 | 13.15 | 0.2370 | 0.9726 | 37.64 | 0.5462 |
| Psydraxdicocca | 0.7628* | 61.53 | 13.19 | 0.2379 | 0.9751 | 36.77 | 0.5504 |
| Tarennoidea | 0.6406 | 65.66 | 12.85 | 0.2289 | 0.9496 | 45.47 | 0.5085 |
| Tarennoideawallichii | 0.6374* | 65.77 | 12.84 | 0.2287 | 0.9489 | 45.69 | 0.5075 |
| hedyotis | 0.6367 | 65.79 | 12.84 | 0.2287 | 0.9488 | 45.74 | 0.5072 |
| Hedyotiscathayana | 0.6374* | 65.77 | 12.84 | 0.2287 | 0.9489 | 45.69 | 0.5075 |
| saprosma | 0.6372 | 65.77 | 12.84 | 0.2287 | 0.9489 | 45.71 | 0.5074 |
| Saprosmacrassipes | 0.6374* | 65.77 | 12.84 | 0.2287 | 0.9489 | 45.69 | 0.5075 |
| Saprosmahainanensis | 0.6374* | 65.77 | 12.84 | 0.2287 | 0.9489 | 45.69 | 0.5075 |
| Saprosmamerrillii | 0.6374* | 65.77 | 12.84 | 0.2287 | 0.9489 | 45.69 | 0.5075 |
| Saprosmayueyan | 0.6374* | 65.77 | 12.84 | 0.2287 | 0.9489 | 45.69 | 0.5075 |
| chassalia | 0.6309 | 65.98 | 12.83 | 0.2282 | 0.9475 | 46.16 | 0.5052 |
| Chassaliacurviflora | 0.6374* | 65.77 | 12.84 | 0.2287 | 0.9489 | 45.69 | 0.5075 |
| psychotria | 0.5730 | 67.94 | 12.67 | 0.2240 | 0.9354 | 50.27 | 0.4854 |
| Psychotriaasiatica | 0.5636* | 68.26 | 12.64 | 0.2233 | 0.9335 | 50.94 | 0.4822 |

| | | | | | | | |
|---|---|---|---|---|---|---|---|
| Psychotriastraminea | 0.5636* | 68.26 | 12.64 | 0.2233 | 0.9335 | 50.94 | 0.4822 |
| prismatomeris | 0.6304 | 66.00 | 12.82 | 0.2282 | 0.9474 | 46.19 | 0.5051 |
| Prismatomeristetrandra | 0.6374* | 65.77 | 12.84 | 0.2287 | 0.9489 | 45.69 | 0.5075 |
| lasianthus | 0.6373 | 65.77 | 12.84 | 0.2287 | 0.9489 | 45.70 | 0.5074 |
| Lasianthuschevalieri | 0.6374* | 65.77 | 12.84 | 0.2287 | 0.9489 | 45.69 | 0.5075 |
| Lasianthuscurtisii | 0.6374* | 65.77 | 12.84 | 0.2287 | 0.9489 | 45.69 | 0.5075 |
| Lasianthushirsutus | 0.6374* | 65.77 | 12.84 | 0.2287 | 0.9489 | 45.69 | 0.5075 |
| Lasianthusjaponicus | 0.6374* | 65.77 | 12.84 | 0.2287 | 0.9489 | 45.69 | 0.5075 |
| Lasianthuslancifolius | 0.6374* | 65.77 | 12.84 | 0.2287 | 0.9489 | 45.69 | 0.5075 |
| Lasianthusrhinocerotis | 0.6374* | 65.77 | 12.84 | 0.2287 | 0.9489 | 45.69 | 0.5075 |
| Lasianthustrichophlebus | 0.6374* | 65.77 | 12.84 | 0.2287 | 0.9489 | 45.69 | 0.5075 |
| Caelospermum | 0.6406 | 65.66 | 12.85 | 0.2289 | 0.9496 | 45.47 | 0.5085 |
| Caelospermumtruncatum | 0.6374* | 65.77 | 12.84 | 0.2287 | 0.9489 | 45.69 | 0.5075 |
| Wendlandia | 0.6764 | 64.45 | 12.95 | 0.2316 | 0.9571 | 42.91 | 0.5208 |
| Wendlandiamerrilliana | 0.7337* | 62.51 | 13.11 | 0.2357 | 0.9690 | 38.84 | 0.5404 |
| Wendlandiauvariifolia | 0.6518* | 65.28 | 12.88 | 0.2298 | 0.9519 | 44.66 | 0.5124 |
| apocynaceae | 0.6109 | 66.66 | 12.77 | 0.2268 | 0.9434 | 47.58 | 0.4984 |
| rauvolfia | 0.5001 | 70.40 | 12.47 | 0.2187 | 0.9202 | 55.46 | 0.4605 |
| Rauvolfiaverticillata | 0.4863* | 70.87 | 12.43 | 0.2177 | 0.9173 | 56.44 | 0.4558 |
| kopsia | 0.5740 | 67.91 | 12.67 | 0.2241 | 0.9356 | 50.20 | 0.4858 |
| Kopsiaarborea | 0.5929* | 67.27 | 12.72 | 0.2255 | 0.9396 | 48.86 | 0.4922 |
| wrightia | 0.3411 | 75.77 | 12.03 | 0.2071 | 0.8870 | 66.78 | 0.4061 |
| Wrightialaevis | 0.3120* | 76.76 | 11.95 | 0.2049 | 0.8809 | 68.85 | 0.3961 |
| tabernaemontana | 0.5587 | 68.42 | 12.63 | 0.2230 | 0.9324 | 51.29 | 0.4805 |
| Tabernaemontanabovina | 0.5656* | 68.19 | 12.65 | 0.2235 | 0.9339 | 50.80 | 0.4829 |
| Tabernaemontanabufalina | 0.5656* | 68.19 | 12.65 | 0.2235 | 0.9339 | 50.80 | 0.4829 |
| alstonia | 0.5095 | 70.08 | 12.49 | 0.2194 | 0.9222 | 54.79 | 0.4637 |
| Alstoniarostrata | 0.4640* | 71.62 | 12.37 | 0.2160 | 0.9126 | 58.03 | 0.4481 |
| Hunteria | 0.6654 | 64.82 | 12.92 | 0.2308 | 0.9548 | 43.70 | 0.5171 |
| Hunteriazeylanica | 0.7200* | 62.97 | 13.07 | 0.2348 | 0.9662 | 39.81 | 0.5357 |
| lamiales | 0.6055 | 66.84 | 12.76 | 0.2264 | 0.9422 | 47.96 | 0.4965 |
| bignoniaceae | 0.5817 | 67.64 | 12.69 | 0.2246 | 0.9373 | 49.65 | 0.4884 |
| radermachera | 0.5246 | 69.57 | 12.53 | 0.2205 | 0.9253 | 53.72 | 0.4689 |
| Radermacherasinica | 0.6255* | 66.17 | 12.81 | 0.2278 | 0.9464 | 46.54 | 0.5034 |
| Radermacherafrondosa | 0.4592* | 71.78 | 12.35 | 0.2157 | 0.9117 | 58.37 | 0.4465 |
| Radermacherahainanensis | 0.4592* | 71.78 | 12.35 | 0.2157 | 0.9117 | 58.37 | 0.4465 |
| markhamia | 0.6451 | 65.50 | 12.87 | 0.2293 | 0.9505 | 45.14 | 0.5101 |
| Markhamiastipulata | 0.6755* | 64.48 | 12.95 | 0.2315 | 0.9569 | 42.98 | 0.5205 |
| lamiaceae | 0.5300 | 69.39 | 12.55 | 0.2209 | 0.9265 | 53.33 | 0.4707 |
| gmelina | 0.5624 | 68.30 | 12.64 | 0.2232 | 0.9332 | 51.03 | 0.4818 |
| Gmelinahainanensis | 0.5947* | 67.21 | 12.73 | 0.2256 | 0.9400 | 48.73 | 0.4928 |
| Tsoongia | 0.5533 | 68.60 | 12.61 | 0.2226 | 0.9313 | 51.68 | 0.4787 |
| Tsoongiaaxillariflora | 0.5766* | 67.82 | 12.68 | 0.2243 | 0.9362 | 50.02 | 0.4866 |
| callicarpa | 0.4100 | 73.44 | 12.22 | 0.2121 | 0.9014 | 61.87 | 0.4296 |
| Callicarpabrevipes | 0.3500* | 75.47 | 12.05 | 0.2077 | 0.8888 | 66.14 | 0.4091 |
| Callicarpasp2 | 0.3500* | 75.47 | 12.05 | 0.2077 | 0.8888 | 66.14 | 0.4091 |
| vitex | 0.6354 | 65.83 | 12.84 | 0.2286 | 0.9485 | 45.84 | 0.5068 |
| Vitexpierreana | 0.8545* | 58.43 | 13.44 | 0.2446 | 0.9943 | 30.24 | 0.5818 |
| Vitexquinata | 0.4513* | 72.05 | 12.33 | 0.2151 | 0.9100 | 58.93 | 0.4438 |
| clerodendrum | 0.5702 | 68.04 | 12.66 | 0.2238 | 0.9348 | 50.48 | 0.4844 |
| Clerodendrumhainanense | 0.5735* | 67.92 | 12.67 | 0.2240 | 0.9355 | 50.24 | 0.4856 |
| Clerodendrumkwangtungense | 0.5735* | 67.92 | 12.67 | 0.2240 | 0.9355 | 50.24 | 0.4856 |
| verbenaceae | 0.6258 | 66.16 | 12.81 | 0.2279 | 0.9465 | 46.52 | 0.5035 |
| Verbenaceaesp | 0.6426* | 65.59 | 12.86 | 0.2291 | 0.9500 | 45.32 | 0.5092 |
| oleaceae | 0.6711 | 64.63 | 12.94 | 0.2312 | 0.9560 | 43.30 | 0.5190 |
| olea | 0.6935 | 63.87 | 13.00 | 0.2328 | 0.9606 | 41.70 | 0.5267 |
| Oleabrachiata | 0.6300* | 66.01 | 12.82 | 0.2282 | 0.9474 | 46.22 | 0.5049 |
| Oleaneriifolia | 0.7431* | 62.19 | 13.13 | 0.2364 | 0.9710 | 38.17 | 0.5436 |
| Oleaparvilimba | 0.7431* | 62.19 | 13.13 | 0.2364 | 0.9710 | 38.17 | 0.5436 |
| Oleatsoongii | 0.6300* | 66.01 | 12.82 | 0.2282 | 0.9474 | 46.22 | 0.5049 |
| osmanthus | 0.8323 | 59.18 | 13.38 | 0.2430 | 0.9897 | 31.82 | 0.5741 |
| Osmanthusdidymopetalus | 0.8415* | 58.87 | 13.41 | 0.2436 | 0.9916 | 31.17 | 0.5773 |
| Osmanthushainanensis | 0.8415* | 58.87 | 13.41 | 0.2436 | 0.9916 | 31.17 | 0.5773 |
| Osmanthusmarginatus | 0.8415* | 58.87 | 13.41 | 0.2436 | 0.9916 | 31.17 | 0.5773 |

| | | | | | | | |
|---|---|---|---|---|---|---|---|
| Osmanthusmatsumuranus | 0.8415* | 58.87 | 13.41 | 0.2436 | 0.9916 | 31.17 | 0.5773 |
| chionanthus | 0.7094 | 63.33 | 13.04 | 0.2340 | 0.9640 | 40.57 | 0.5321 |
| Chionanthusbrachythyrsus | 0.6805* | 64.31 | 12.96 | 0.2319 | 0.9579 | 42.63 | 0.5222 |
| Chionanthusramiflorus | 0.7530* | 61.86 | 13.16 | 0.2372 | 0.9731 | 37.46 | 0.5470 |
| boraginales | 0.5693 | 68.06 | 12.66 | 0.2237 | 0.9347 | 50.54 | 0.4842 |
| boraginaceae | 0.5336 | 69.27 | 12.56 | 0.2211 | 0.9272 | 53.08 | 0.4719 |
| ehretia | 0.5140 | 69.93 | 12.50 | 0.2197 | 0.9231 | 54.47 | 0.4652 |
| Ehretialongiflora | 0.5195* | 69.75 | 12.52 | 0.2201 | 0.9242 | 54.08 | 0.4671 |
| cordia | 0.4745 | 71.27 | 12.40 | 0.2168 | 0.9148 | 57.28 | 0.4517 |
| Cordiadichotoma | 0.4512* | 72.05 | 12.33 | 0.2151 | 0.9100 | 58.94 | 0.4437 |
| garryales | 0.5291 | 69.42 | 12.55 | 0.2208 | 0.9263 | 53.40 | 0.4704 |
| icacinaceae | 0.5004 | 70.39 | 12.47 | 0.2187 | 0.9203 | 55.44 | 0.4606 |
| apodytes | 0.5735 | 67.92 | 12.67 | 0.2240 | 0.9355 | 50.24 | 0.4856 |
| Apodytesdimidiata | 0.6100* | 66.69 | 12.77 | 0.2267 | 0.9432 | 47.64 | 0.4981 |
| Platea | 0.4351 | 72.60 | 12.29 | 0.2139 | 0.9066 | 60.09 | 0.4382 |
| Platealatifolia | 0.3400* | 75.81 | 12.03 | 0.2070 | 0.8867 | 66.86 | 0.4057 |
| Plateaparvifolia | 0.4650* | 71.59 | 12.37 | 0.2161 | 0.9129 | 57.96 | 0.4485 |
| apiales | 0.5389 | 69.09 | 12.57 | 0.2215 | 0.9283 | 52.70 | 0.4738 |
| pittosporaceae | 0.5381 | 69.12 | 12.57 | 0.2215 | 0.9281 | 52.76 | 0.4735 |
| pittosporum | 0.6077 | 66.77 | 12.76 | 0.2265 | 0.9427 | 47.80 | 0.4973 |
| Pittosporumbalansae | 0.6135* | 66.57 | 12.78 | 0.2270 | 0.9439 | 47.39 | 0.4993 |
| Pittosporumcrispulum | 0.6135* | 66.57 | 12.78 | 0.2270 | 0.9439 | 47.39 | 0.4993 |
| Pittosporumperryanum | 0.6135* | 66.57 | 12.78 | 0.2270 | 0.9439 | 47.39 | 0.4993 |
| araliaceae | 0.4737 | 71.29 | 12.39 | 0.2168 | 0.9147 | 57.34 | 0.4514 |
| heteropanax | 0.3638 | 75.01 | 12.09 | 0.2087 | 0.8917 | 65.17 | 0.4138 |
| Heteropanaxfragrans | 0.3440* | 75.67 | 12.04 | 0.2073 | 0.8876 | 66.57 | 0.4071 |
| schefflera | 0.3975 | 73.87 | 12.18 | 0.2112 | 0.8987 | 62.77 | 0.4253 |
| Scheffleraheinanensis | 0.3946* | 73.97 | 12.18 | 0.2110 | 0.8981 | 62.97 | 0.4244 |
| Scheffleraheptaphylla | 0.3946* | 73.97 | 12.18 | 0.2110 | 0.8981 | 62.97 | 0.4244 |
| dendropanax | 0.4178 | 73.18 | 12.24 | 0.2127 | 0.9030 | 61.32 | 0.4323 |
| Dendropanaxhainanensis | 0.4198* | 73.11 | 12.25 | 0.2128 | 0.9034 | 61.17 | 0.4330 |
| dipsacales | 0.5508 | 68.69 | 12.61 | 0.2224 | 0.9308 | 51.86 | 0.4778 |
| adoxaceae | 0.5537 | 68.59 | 12.61 | 0.2226 | 0.9314 | 51.65 | 0.4788 |
| sambucus | 0.4821 | 71.01 | 12.42 | 0.2174 | 0.9164 | 56.74 | 0.4543 |
| Sambucusjavanica | 0.4463* | 72.22 | 12.32 | 0.2148 | 0.9090 | 59.29 | 0.4421 |
| viburnum | 0.5923 | 67.29 | 12.72 | 0.2254 | 0.9395 | 48.90 | 0.4920 |
| Viburnumpunctatum | 0.6310* | 65.98 | 12.83 | 0.2282 | 0.9476 | 46.15 | 0.5053 |
| escalloniales | 0.5514 | 68.67 | 12.61 | 0.2224 | 0.9309 | 51.81 | 0.4780 |
| escalloniaceae | 0.5515 | 68.66 | 12.61 | 0.2224 | 0.9310 | 51.80 | 0.4781 |
| polyosma | 0.5519 | 68.65 | 12.61 | 0.2225 | 0.9310 | 51.78 | 0.4782 |
| Polyosmacambodiana | 0.5520* | 68.65 | 12.61 | 0.2225 | 0.9311 | 51.77 | 0.4782 |
| aquifoliales | 0.5599 | 68.38 | 12.63 | 0.2231 | 0.9327 | 51.20 | 0.4810 |
| stemonuraceae | 0.5250 | 69.56 | 12.53 | 0.2205 | 0.9254 | 53.69 | 0.4690 |
| gomphandra | 0.4905 | 70.73 | 12.44 | 0.2180 | 0.9182 | 56.15 | 0.4572 |
| Gomphandratetrandra | 0.4560* | 71.89 | 12.35 | 0.2155 | 0.9110 | 58.60 | 0.4454 |
| cardiopteridaceae | 0.5935 | 67.25 | 12.72 | 0.2255 | 0.9397 | 48.82 | 0.4924 |
| gonocaryum | 0.6275 | 66.10 | 12.82 | 0.2280 | 0.9468 | 46.40 | 0.5041 |
| Gonocaryumlobbianum | 0.6615* | 64.95 | 12.91 | 0.2305 | 0.9539 | 43.98 | 0.5157 |
| aquifoliaceae | 0.5659 | 68.18 | 12.65 | 0.2235 | 0.9340 | 50.78 | 0.4830 |
| ilex | 0.5689 | 68.08 | 12.66 | 0.2237 | 0.9346 | 50.57 | 0.4840 |
| Ilexsp12 | 0.5627* | 68.29 | 12.64 | 0.2233 | 0.9333 | 51.01 | 0.4819 |
| Ilexsp2 | 0.5627* | 68.29 | 12.64 | 0.2233 | 0.9333 | 51.01 | 0.4819 |
| Ilexsp5 | 0.5627* | 68.29 | 12.64 | 0.2233 | 0.9333 | 51.01 | 0.4819 |
| Ilexsterrophylla | 0.6485* | 65.39 | 12.87 | 0.2295 | 0.9512 | 44.90 | 0.5113 |
| Ilextriflora | 0.5627* | 68.29 | 12.64 | 0.2233 | 0.9333 | 51.01 | 0.4819 |
| Ilexangulata | 0.5627* | 68.29 | 12.64 | 0.2233 | 0.9333 | 51.01 | 0.4819 |
| Ilexcochinchinensis | 0.6030* | 66.93 | 12.75 | 0.2262 | 0.9417 | 48.14 | 0.4957 |
| Ilexelmerrilliana | 0.5627* | 68.29 | 12.64 | 0.2233 | 0.9333 | 51.01 | 0.4819 |
| Ilexficoidea | 0.5627* | 68.29 | 12.64 | 0.2233 | 0.9333 | 51.01 | 0.4819 |
| Ilexgodajam | 0.5627* | 68.29 | 12.64 | 0.2233 | 0.9333 | 51.01 | 0.4819 |
| Ilexgoshiensis | 0.5627* | 68.29 | 12.64 | 0.2233 | 0.9333 | 51.01 | 0.4819 |
| Ilexhainanensis | 0.5627* | 68.29 | 12.64 | 0.2233 | 0.9333 | 51.01 | 0.4819 |
| Ilexkobuskiana | 0.5627* | 68.29 | 12.64 | 0.2233 | 0.9333 | 51.01 | 0.4819 |
| Ilexlancilimba | 0.5627* | 68.29 | 12.64 | 0.2233 | 0.9333 | 51.01 | 0.4819 |

| | | | | | | | |
|---|---|---|---|---|---|---|---|
| Ilexnuculicava | **0.5627*** | **68.29** | **12.64** | **0.2233** | **0.9333** | **51.01** | **0.4819** |
| Ilexpubescens | **0.5627*** | **68.29** | **12.64** | **0.2233** | **0.9333** | **51.01** | **0.4819** |
| Ilexrotunda | **0.5627*** | **68.29** | **12.64** | **0.2233** | **0.9333** | **51.01** | **0.4819** |
| Ilexsp | **0.5627*** | **68.29** | **12.64** | **0.2233** | **0.9333** | **51.01** | **0.4819** |
| Ilexsp0502 | **0.5627*** | **68.29** | **12.64** | **0.2233** | **0.9333** | **51.01** | **0.4819** |
| Ilexsp1 | **0.5627*** | **68.29** | **12.64** | **0.2233** | **0.9333** | **51.01** | **0.4819** |
| ericales | **0.5577** | **67.10** | **12.69** | **0.2244** | **0.9227** | **51.25** | **0.4829** |
| pentaphyllacaceae | **0.5621** | **52.02** | **12.42** | **0.2507** | **0.8650** | **48.90** | **0.5600** |
| pentaphylax | **0.5481** | **52.49** | **12.38** | **0.2497** | **0.8621** | **49.90** | **0.5552** |
| Pentaphylaxeuryoides | **0.5340*** | **52.97** | **12.34** | **0.2487** | **0.8591** | **50.90** | **0.5503** |
| ternstroemia | **0.5878** | **47.03** | **12.34** | **0.2627** | **0.8486** | **46.36** | **0.6082** |
| Ternstroemiagymnanthera | **0.5814*** | **47.25** | **12.32** | **0.2622** | **0.8473** | **46.82** | **0.6060** |
| Ternstroemiahainanensis | **0.5814*** | **47.25** | **12.32** | **0.2622** | **0.8473** | **46.82** | **0.6060** |
| anneslea | **0.6425** | **45.19** | **12.49** | **0.2667** | **0.8601** | **42.47** | **0.6269** |
| Annesleafragrans | **0.6843*** | **43.77** | **12.60** | **0.2698** | **0.8688** | **39.50** | **0.6412** |
| eurya | **0.5241** | **26.15** | **13.73** | **0.2172** | **0.8021** | **42.90** | **0.6947** |
| Euryaciliata | **0.5200*** | **16.83*** | **14.58*** | **0.1914*** | **0.7956*** | **39.54*** | **0.7278*** |
| Euryacuneata | **0.5200*** | **26.29** | **13.72** | **0.2169** | **0.8013** | **43.19** | **0.6934** |
| Euryagroffii | **0.5200*** | **26.29** | **13.72** | **0.2169** | **0.8013** | **43.19** | **0.6934** |
| Euryahainanensis | **0.5200*** | **26.29** | **13.72** | **0.2169** | **0.8013** | **43.19** | **0.6934** |
| Euryaloquaiana | **0.5200*** | **26.29** | **13.72** | **0.2169** | **0.8013** | **43.19** | **0.6934** |
| Euryanitida | **0.5300*** | **25.95** | **13.74** | **0.2176** | **0.8034** | **42.48** | **0.6968** |
| adinandra | **0.5331** | **55.97** | **10.49** | **0.3147** | **0.7535** | **56.68** | **0.6438** |
| Adinandraangustifolia | **0.5814*** | **54.34** | **10.62** | **0.3182** | **0.7636** | **53.24** | **0.6604** |
| Adinandrahainanensis | **0.4700*** | **63.97*** | **9.792*** | **0.3204*** | **0.6944*** | **65.35*** | **0.6323*** |
| cleyera | **0.5578** | **48.76** | **10.60** | **0.3315** | **0.8343** | **49.49** | **0.6372** |
| Cleyeraobscurinervia | **0.5677*** | **47.91*** | **10.14*** | **0.3576*** | **0.8662*** | **47.54*** | **0.6355*** |
| ebenaceae | **0.6046** | **56.66** | **13.43** | **0.2078** | **0.9083** | **43.58** | **0.4722** |
| diospyros | **0.5900** | **54.86** | **13.91** | **0.1884** | **0.8702** | **40.61** | **0.4919** |
| Diospyrossusarticulata | **0.5643*** | **55.73** | **13.84** | **0.1865** | **0.8649** | **42.43** | **0.4831** |
| Diospyroschunii | **0.5643*** | **55.73** | **13.84** | **0.1865** | **0.8649** | **42.43** | **0.4831** |
| Diospyroseriantha | **0.7026*** | **51.06** | **14.22** | **0.1966** | **0.8938** | **32.59** | **0.5304** |
| Diospyroshainanensis | **0.5643*** | **55.73** | **13.84** | **0.1865** | **0.8649** | **42.43** | **0.4831** |
| Diospyrosinflata | **0.5643*** | **55.73** | **13.84** | **0.1865** | **0.8649** | **42.43** | **0.4831** |
| Diospyroslongibracteata | **0.5100*** | **57.56** | **13.69** | **0.1825** | **0.8535** | **46.30** | **0.4645** |
| Diospyrosmaclurei | **0.7040*** | **51.01** | **14.22** | **0.1967** | **0.8941** | **32.49** | **0.5309** |
| Diospyrosmorrisiana | **0.5643*** | **54.58*** | **14.10*** | **0.1773*** | **0.8474*** | **40.42*** | **0.4955*** |
| Diospyrosstrigosa | **0.5643*** | **55.73** | **13.84** | **0.1865** | **0.8649** | **42.43** | **0.4831** |
| primulaceae | **0.6407** | **56.59** | **13.27** | **0.2196** | **0.9333** | **43.02** | **0.4721** |
| myrsine | **0.7145** | **54.10** | **13.47** | **0.2250** | **0.9487** | **37.77** | **0.4974** |
| Myrsineseguinii | **0.7410*** | **53.20** | **13.54** | **0.2269** | **0.9543** | **35.88** | **0.5065** |
| Myrsinestolonifera | **0.7410*** | **53.20** | **13.54** | **0.2269** | **0.9543** | **35.88** | **0.5065** |
| ardisia | **0.6111** | **57.58** | **13.19** | **0.2174** | **0.9271** | **45.12** | **0.4620** |
| Ardisiacrassinervosa | **0.6265*** | **57.07** | **13.23** | **0.2186** | **0.9303** | **44.03** | **0.4673** |
| Ardisiadensilepidotula | **0.6265*** | **57.07** | **13.23** | **0.2186** | **0.9303** | **44.03** | **0.4673** |
| Ardisiaobtusa | **0.5909*** | **58.27** | **13.13** | **0.2160** | **0.9229** | **46.56** | **0.4551** |
| Ardisiaquinquegona | **0.5909*** | **58.27** | **13.13** | **0.2160** | **0.9229** | **46.56** | **0.4551** |
| Ardisiavillosa | **0.5909*** | **58.27** | **13.13** | **0.2160** | **0.9229** | **46.56** | **0.4551** |
| Ardisiavirens | **0.5909*** | **58.27** | **13.13** | **0.2160** | **0.9229** | **46.56** | **0.4551** |
| Embelia | **0.6400** | **56.61** | **13.27** | **0.2195** | **0.9331** | **43.07** | **0.4719** |
| Embeliavestita | **0.6393*** | **56.64** | **13.26** | **0.2195** | **0.9330** | **43.12** | **0.4717** |
| maesa | **0.6672** | **55.69** | **13.34** | **0.2215** | **0.9388** | **41.13** | **0.4812** |
| Maesaacuminatissima | **0.6760*** | **55.39** | **13.36** | **0.2222** | **0.9407** | **40.50** | **0.4842** |
| Maesaconsanguinea | **0.6760*** | **55.39** | **13.36** | **0.2222** | **0.9407** | **40.50** | **0.4842** |
| Maesaperlarius | **0.6760*** | **55.39** | **13.36** | **0.2222** | **0.9407** | **40.50** | **0.4842** |
| sapotaceae | **0.5887** | **60.77** | **12.66** | **0.2267** | **0.9832** | **51.03** | **0.3814** |
| planchonella | **0.6952** | **64.22** | **11.00** | **0.2351** | **0.9713** | **50.19** | **0.4392** |
| Planchonellaclemensii | **0.7148*** | **63.56** | **11.05** | **0.2365** | **0.9754** | **48.80** | **0.4459** |
| chrysophyllum | **0.5699** | **68.45** | **10.65** | **0.2260** | **0.9451** | **59.11** | **0.3963** |
| Chrysophyllumlanceolatum | **0.5230*** | **70.04** | **10.52** | **0.2225** | **0.9353** | **62.45** | **0.3802** |
| madhuca | **0.8592** | **59.24** | **9.505** | **0.2605** | **0.9979** | **37.44** | **0.5355** |
| Madhucahainanensis | **0.9257*** | **57.27*** | **8.717*** | **0.2721*** | **1.008*** | **32.16*** | **0.5783*** |
| pouteria | **0.6678** | **70.80** | **13.82** | **0.2002** | **0.9506** | **61.60** | **0.3507** |

| | | | | | | | |
|---|---|---|---|---|---|---|---|
| Pouteriaannamensis | 0.6878* | 76.34* | 14.82* | 0.1822* | 0.9321* | 68.54* | 0.3186* |
| sarcosperma | 0.5277 | 57.63 | 12.32 | 0.2367 | 1.040 | 49.85 | 0.3188 |
| Sarcospermalaurinum | 0.4667* | 54.48* | 11.97* | 0.2467* | 1.097* | 48.67* | 0.2562* |
| ericaceae | 0.5276 | 68.01 | 12.88 | 0.2242 | 0.8835 | 53.48 | 0.4897 |
| rhododendron | 0.4984 | 69.00 | 12.80 | 0.2221 | 0.8774 | 55.55 | 0.4797 |
| Rhododendronmoulmainense | 0.4955* | 69.10 | 12.79 | 0.2219 | 0.8768 | 55.76 | 0.4787 |
| symplocaceae | 0.5244 | 77.26 | 14.11 | 0.2194 | 0.8685 | 53.58 | 0.4964 |
| symplocos | 0.5317 | 77.02 | 14.13 | 0.2200 | 0.8700 | 53.06 | 0.4989 |
| Symplocoscongesta | 0.5335* | 76.96 | 14.14 | 0.2201 | 0.8704 | 52.93 | 0.4995 |
| Symplocoscrassilimba | 0.5335* | 76.96 | 14.14 | 0.2201 | 0.8704 | 52.93 | 0.4995 |
| Symplocoseuryoides | 0.5335* | 76.96 | 14.14 | 0.2201 | 0.8704 | 52.93 | 0.4995 |
| Symplocosglauca | 0.5335* | 76.96 | 14.14 | 0.2201 | 0.8704 | 52.93 | 0.4995 |
| Symplocosheishanensis | 0.5335* | 76.96 | 14.14 | 0.2201 | 0.8704 | 52.93 | 0.4995 |
| Symplocoslancifolia | 0.4635* | 79.32 | 13.95 | 0.2150 | 0.8558 | 57.92 | 0.4755 |
| Symplocospendula | 0.5335* | 76.96 | 14.14 | 0.2201 | 0.8704 | 52.93 | 0.4995 |
| Symplocospoilanei | 0.5335* | 76.96 | 14.14 | 0.2201 | 0.8704 | 52.93 | 0.4995 |
| Symplocospseudobarberina | 0.5335* | 76.96 | 14.14 | 0.2201 | 0.8704 | 52.93 | 0.4995 |
| Symplocosracemosa | 0.5335* | 76.96 | 14.14 | 0.2201 | 0.8704 | 52.93 | 0.4995 |
| Symplocossp1 | 0.5335* | 76.96 | 14.14 | 0.2201 | 0.8704 | 52.93 | 0.4995 |
| Symplocossumuntia | 0.5335* | 76.96 | 14.14 | 0.2201 | 0.8704 | 52.93 | 0.4995 |
| Symplocosviridissima | 0.5335* | 76.96 | 14.14 | 0.2201 | 0.8704 | 52.93 | 0.4995 |
| Symplocoswikstroemiifolia | 0.5335* | 76.96 | 14.14 | 0.2201 | 0.8704 | 52.93 | 0.4995 |
| Symplocosadenophylla | 0.6500* | 73.02 | 14.46 | 0.2286 | 0.8948 | 44.64 | 0.5394 |
| Symplocosanomala | 0.4815* | 78.71 | 14.00 | 0.2163 | 0.8595 | 56.64 | 0.4817 |
| Symplocoscochinchinensis | 0.5150* | 77.58 | 14.09 | 0.2187 | 0.8665 | 54.25 | 0.4932 |
| Symplocaceaesp7 | 0.5335* | 76.96 | 14.14 | 0.2201 | 0.8704 | 52.93 | 0.4995 |
| styracaceae | 0.4394 | 93.18 | 16.06 | 0.2063 | 0.8495 | 58.79 | 0.4598 |
| alniphyllum | 0.3940 | 120.8 | 20.30 | 0.1891 | 0.8374 | 60.34 | 0.4292 |
| Alniphyllumfortunei | 0.3827* | 127.7* | 21.36* | 0.1848* | 0.8344* | 60.73* | 0.4215* |
| styrax | 0.4165 | 93.95 | 16.00 | 0.2046 | 0.8447 | 60.42 | 0.4519 |
| Styraxagrestis | 0.4050* | 94.34 | 15.97 | 0.2038 | 0.8423 | 61.24 | 0.4480 |
| Styraxsuberifolius | 0.4050* | 94.34 | 15.97 | 0.2038 | 0.8423 | 61.24 | 0.4480 |
| theaceae | 0.5304 | 66.64 | 12.10 | 0.2257 | 0.8573 | 54.30 | 0.5175 |
| camellia | 0.5499 | 55.25 | 9.747 | 0.2440 | 0.9546 | 40.20 | 0.5945 |
| Camelliacaudata | 0.5500* | 55.25 | 9.747 | 0.2440 | 0.9546 | 40.20 | 0.5945 |
| Camelliaoleifera | 0.5500* | 55.25 | 9.747 | 0.2440 | 0.9546 | 40.20 | 0.5945 |
| Camelliapaucipunctata | 0.5500* | 55.25 | 9.747 | 0.2440 | 0.9546 | 40.20 | 0.5945 |
| Camelliasinensis | 0.5500* | 55.25 | 9.747 | 0.2440 | 0.9546 | 40.20 | 0.5945 |
| Camelliasp1 | 0.5500* | 55.25 | 9.747 | 0.2440 | 0.9546 | 40.20 | 0.5945 |
| Camelliasp3 | 0.5500* | 55.25 | 9.747 | 0.2440 | 0.9546 | 40.20 | 0.5945 |
| Camelliaxanthochroma | 0.5500* | 55.25 | 9.747 | 0.2440 | 0.9546 | 40.20 | 0.5945 |
| polyspora | 0.5613 | 54.56 | 9.052 | 0.2529 | 1.023 | 31.39 | 0.6266 |
| Polysporaaxillaris | 0.5677* | 54.35 | 9.070 | 0.2534 | 1.024 | 30.93 | 0.6287 |
| Polysporahainanensis | 0.5677* | 54.05* | 8.345* | 0.2614* | 1.091* | 22.92* | 0.6570* |
| pyrenaria | 0.5225 | 50.25 | 10.38 | 0.2324 | 0.8568 | 52.75 | 0.5554 |
| Pyrenariajonquieriana | 0.5182* | 44.16* | 10.35* | 0.2304* | 0.8299* | 55.64* | 0.5524* |
| Pyrenariamicrocarpa | 0.5182* | 50.39 | 10.37 | 0.2320 | 0.8559 | 53.05 | 0.5539 |
| Pyrenariaspectabilis | 0.5182* | 50.39 | 10.37 | 0.2320 | 0.8559 | 53.05 | 0.5539 |
| schima | 0.5430 | 67.56 | 10.80 | 0.2208 | 0.7406 | 66.38 | 0.5144 |
| Schimaremotiserrata | 0.5526* | 67.24 | 10.83 | 0.2215 | 0.7426 | 65.70 | 0.5176 |
| Schimasuperba | 0.5375* | 70.38* | 10.59* | 0.2163* | 0.6863* | 72.91* | 0.5011* |
| cornales | 0.5439 | 70.28 | 12.53 | 0.2203 | 0.9389 | 52.45 | 0.4727 |
| nyssaceae | 0.5066 | 71.54 | 12.42 | 0.2176 | 0.9311 | 55.11 | 0.4599 |
| mastixia | 0.4989 | 71.80 | 12.40 | 0.2171 | 0.9295 | 55.66 | 0.4573 |
| Mastixiapentandra | 0.4950* | 71.93 | 12.39 | 0.2168 | 0.9287 | 55.93 | 0.4560 |
| cornaceae | 0.4976 | 71.85 | 12.40 | 0.2170 | 0.9293 | 55.75 | 0.4569 |
| alangium | 0.4397 | 73.80 | 12.24 | 0.2127 | 0.9172 | 59.87 | 0.4371 |
| Alangiumchinense | 0.4017* | 75.09 | 12.13 | 0.2099 | 0.9092 | 62.58 | 0.4240 |
| Alangiumkurzii | 0.4200* | 74.47 | 12.19 | 0.2113 | 0.9130 | 61.27 | 0.4303 |
| cornus | 0.5425 | 70.33 | 12.52 | 0.2202 | 0.9387 | 52.55 | 0.4723 |
| Cornussp1 | 0.5875* | 68.81 | 12.65 | 0.2235 | 0.9481 | 49.35 | 0.4876 |
| santalales | 0.6368 | 71.22 | 12.60 | 0.2225 | 0.9871 | 46.17 | 0.4963 |
| olacaceae | 0.7150 | 68.58 | 12.81 | 0.2282 | 1.003 | 40.60 | 0.5230 |
| olax | 0.7523 | 67.32 | 12.92 | 0.2309 | 1.011 | 37.95 | 0.5358 |

| | | | | | | | |
|---|---|---|---|---|---|---|---|
| Olaximbricata | **0.7710*** | **66.69** | **12.97** | **0.2323** | **1.015** | **36.62** | **0.5422** |
| santalaceae | **0.6849** | **69.60** | **12.73** | **0.2260** | **0.9972** | **42.75** | **0.5127** |
| scleropyrum | **0.6975** | **69.17** | **12.77** | **0.2269** | **0.9998** | **41.85** | **0.5170** |
| Scleropyrumwallichianum | **0.6993*** | **69.11** | **12.77** | **0.2271** | **1.000** | **41.72** | **0.5176** |
| buxales | **0.6429** | **70.39** | **12.44** | **0.2243** | **1.039** | **42.65** | **0.4850** |
| buxaceae | **0.6693** | **69.50** | **12.52** | **0.2263** | **1.044** | **40.78** | **0.4941** |
| buxus | **0.6956** | **68.61** | **12.59** | **0.2282** | **1.050** | **38.90** | **0.5031** |
| Buxusmyrica | **0.7220*** | **67.72** | **12.66** | **0.2301** | **1.055** | **37.02** | **0.5121** |
| proteales | **0.5483** | **71.60** | **12.08** | **0.2203** | **1.060** | **46.19** | **0.4421** |
| proteaceae | **0.5517** | **71.49** | **12.08** | **0.2206** | **1.061** | **45.95** | **0.4433** |
| helicia | **0.5887** | **70.24** | **12.19** | **0.2233** | **1.068** | **43.32** | **0.4559** |
| Heliciacochinchinensis | **0.5510*** | **71.51** | **12.08** | **0.2205** | **1.061** | **46.00** | **0.4430** |
| Heliciaformosana | **0.6051*** | **69.69** | **12.23** | **0.2245** | **1.072** | **42.15** | **0.4615** |
| Heliciahainanensis | **0.6051*** | **69.69** | **12.23** | **0.2245** | **1.072** | **42.15** | **0.4615** |
| Helicialongipetiolata | **0.6051*** | **69.69** | **12.23** | **0.2245** | **1.072** | **42.15** | **0.4615** |
| Heliciaobovatifolia | **0.5777*** | **70.61** | **12.16** | **0.2225** | **1.066** | **44.10** | **0.4521** |
| Heliciareticulata | **0.6051*** | **69.69** | **12.23** | **0.2245** | **1.072** | **42.15** | **0.4615** |
| heliciopsis | **0.4612** | **74.55** | **11.84** | **0.2140** | **1.042** | **52.39** | **0.4123** |
| Heliciopsislobata | **0.4300*** | **75.60** | **11.75** | **0.2117** | **1.035** | **54.61** | **0.4016** |
| Heliciopsisterminalis | **0.4767*** | **74.03** | **11.88** | **0.2151** | **1.045** | **51.29** | **0.4176** |
| sabiaceae | **0.5297** | **72.24** | **12.02** | **0.2190** | **1.056** | **47.52** | **0.4357** |
| meliosma | **0.5110** | **72.87** | **11.97** | **0.2176** | **1.052** | **48.84** | **0.4293** |
| Meliosmaangustifolia | **0.5483*** | **71.61** | **12.08** | **0.2203** | **1.060** | **46.19** | **0.4421** |
| Meliosmadumicola | **0.4789*** | **73.95** | **11.88** | **0.2153** | **1.045** | **51.13** | **0.4183** |
| Meliosmafordii | **0.4789*** | **73.95** | **11.88** | **0.2153** | **1.045** | **51.13** | **0.4183** |
| Meliosmalaui | **0.4789*** | **73.95** | **11.88** | **0.2153** | **1.045** | **51.13** | **0.4183** |
| Meliosmarigida | **0.5960*** | **69.99** | **12.21** | **0.2238** | **1.070** | **42.80** | **0.4584** |
| Meliosmasquamulata | **0.4985*** | **73.29** | **11.94** | **0.2167** | **1.050** | **49.73** | **0.4250** |
| Meliosmathorelii | **0.4789*** | **73.95** | **11.88** | **0.2153** | **1.045** | **51.13** | **0.4183** |
| monocots | **0.5388** | **68.96** | **11.88** | **0.2240** | **1.119** | **42.08** | **0.4230** |
| arecales | **0.5312** | **69.21** | **11.86** | **0.2235** | **1.118** | **42.62** | **0.4203** |
| arecaceae | **0.5368** | **69.02** | **11.88** | **0.2239** | **1.119** | **42.22** | **0.4223** |
| pinanga | **0.5518** | **68.51** | **11.92** | **0.2250** | **1.122** | **41.15** | **0.4274** |
| Pinangabaviensis | **0.5518*** | **68.52** | **11.92** | **0.2250** | **1.122** | **41.15** | **0.4274** |
| livistona | **0.5737** | **67.78** | **11.98** | **0.2266** | **1.127** | **39.59** | **0.4349** |
| Livistonasaribus | **0.5814*** | **67.52** | **12.00** | **0.2271** | **1.128** | **39.05** | **0.4375** |
| licuala | **0.5533** | **68.46** | **11.92** | **0.2251** | **1.122** | **41.04** | **0.4279** |
| Licualafordiana | **0.5518*** | **68.52** | **11.92** | **0.2250** | **1.122** | **41.15** | **0.4274** |
| Licualahainanensis | **0.5518*** | **68.52** | **11.92** | **0.2250** | **1.122** | **41.15** | **0.4274** |
| caryota | **0.5512** | **68.54** | **11.92** | **0.2249** | **1.122** | **41.20** | **0.4272** |
| Caryotamaxima | **0.5518*** | **68.52** | **11.92** | **0.2250** | **1.122** | **41.15** | **0.4274** |
| arenga | **0.5512** | **68.54** | **11.92** | **0.2249** | **1.122** | **41.20** | **0.4272** |
| Arengapinnata | **0.5518*** | **68.52** | **11.92** | **0.2250** | **1.122** | **41.15** | **0.4274** |
| asparagales | **0.5104** | **69.91** | **11.81** | **0.2220** | **1.113** | **44.10** | **0.4132** |
| asparagaceae | **0.4349** | **72.46** | **11.60** | **0.2164** | **1.098** | **49.47** | **0.3874** |
| dracaena | **0.3594** | **75.01** | **11.39** | **0.2109** | **1.082** | **54.84** | **0.3616** |
| Dracaenaangustifolia | **0.3500*** | **75.33** | **11.37** | **0.2102** | **1.080** | **55.51** | **0.3584** |
| magnoliids | **0.5339** | **67.14** | **11.76** | **0.2266** | **1.159** | **39.23** | **0.4107** |
| magnoniales | **0.5151** | **66.11** | **11.70** | **0.2270** | **1.226** | **36.49** | **0.3812** |
| annonaceae | **0.5547** | **65.96** | **12.59** | **0.2031** | **1.233** | **31.92** | **0.4134** |
| miliusa | **0.6161** | **63.88** | **12.76** | **0.2076** | **1.246** | **27.55** | **0.4344** |
| Miliusahorsfieldii | **0.6100*** | **64.09** | **12.74** | **0.2072** | **1.245** | **27.98** | **0.4323** |
| popowia | **0.5643** | **65.63** | **12.62** | **0.2038** | **1.235** | **31.24** | **0.4167** |
| Popowiapisocarpa | **0.5450*** | **66.29** | **12.57** | **0.2024** | **1.231** | **32.61** | **0.4101** |
| alphonsea | **0.7078** | **60.79** | **13.01** | **0.2143** | **1.265** | **21.02** | **0.4658** |
| Alphonseahainanensis | **0.7380*** | **59.77** | **13.10** | **0.2165** | **1.271** | **18.87** | **0.4761** |
| mitrephora | **0.6614** | **62.36** | **12.89** | **0.2109** | **1.255** | **24.33** | **0.4499** |
| Mitrephoratomentosa | **0.6800*** | **61.73** | **12.94** | **0.2123** | **1.259** | **23.00** | **0.4563** |
| orophea | **0.5835** | **64.99** | **12.67** | **0.2052** | **1.239** | **29.87** | **0.4233** |
| Oropheahainanensis | **0.5656*** | **65.59** | **12.62** | **0.2039** | **1.235** | **31.14** | **0.4171** |
| polyalthia | **0.6103** | **64.08** | **12.74** | **0.2072** | **1.245** | **27.96** | **0.4324** |
| Polyalthiacerasoides | **0.7550*** | **59.19** | **13.14** | **0.2178** | **1.275** | **17.66** | **0.4820** |
| Polyalthialaui | **0.5541*** | **65.98** | **12.59** | **0.2031** | **1.233** | **31.96** | **0.4132** |
| Polyalthiaobliqua | **0.5541*** | **65.98** | **12.59** | **0.2031** | **1.233** | **31.96** | **0.4132** |

| | | | | | | | |
|---|---|---|---|---|---|---|---|
| Polyalthiarumphii | 0.5800* | 65.10 | 12.66 | 0.2050 | 1.238 | 30.12 | 0.4221 |
| disepalum | 0.5637 | 65.95 | 12.85 | 0.1953 | 1.225 | 30.99 | 0.4269 |
| Disepalumplagioneurum | 0.5656* | 65.88 | 12.85 | 0.1954 | 1.225 | 30.86 | 0.4275 |
| uvaria | 0.5649 | 66.78 | 13.54 | 0.1698 | 1.194 | 30.03 | 0.4585 |
| Uvariaboniana | 0.5656* | 66.76 | 13.55 | 0.1699 | 1.194 | 29.99 | 0.4588 |
| dasymaschalon | 0.5648 | 67.66 | 14.24 | 0.1443 | 1.163 | 29.18 | 0.4897 |
| Dasymaschalonrostratum | 0.5656* | 67.93* | 14.47* | 0.1358* | 1.153* | 28.83* | 0.5004* |
| Chieniodendron | 0.5602 | 65.77 | 12.61 | 0.2035 | 1.234 | 31.53 | 0.4153 |
| Chieniodendronhainanense | 0.5656* | 65.59 | 12.62 | 0.2039 | 1.235 | 31.14 | 0.4171 |
| magnoliaceae | 0.4798 | 67.63 | 11.56 | 0.2306 | 1.289 | 37.52 | 0.3336 |
| Manglietia | 0.4517 | 72.08 | 13.03 | 0.2523 | 1.331 | 36.76 | 0.3023 |
| Manglietiafordiana | 0.4235* | 76.53* | 14.50* | 0.2741* | 1.374* | 36.00* | 0.2710* |
| Michelia | 0.5013 | 73.19 | 10.10 | 0.2267 | 1.321 | 35.63 | 0.3318 |
| Micheliabalansae | 0.5423* | 102.6* | 9.512* | 0.2078* | 1.260* | 40.85* | 0.3500* |
| Micheliagioi | 0.4880* | 73.64 | 10.07 | 0.2257 | 1.318 | 36.58 | 0.3273 |
| Micheliamediocris | 0.4950* | 48.91* | 9.273* | 0.2426* | 1.417* | 27.59* | 0.3164* |
| Lirianthe | 0.4607 | 58.52 | 11.34 | 0.2181 | 1.249 | 41.40 | 0.3350 |
| Lirianthechampionii | 0.4415* | 49.41* | 11.12* | 0.2057* | 1.209* | 45.28* | 0.3364* |
| laurales | 0.5292 | 64.01 | 11.49 | 0.2318 | 1.190 | 35.66 | 0.4058 |
| lauraceae | 0.5482 | 58.15 | 10.94 | 0.2438 | 1.156 | 31.47 | 0.4412 |
| phoebe | 0.5206 | 51.82 | 16.15 | 0.2074 | 0.9201 | 66.01 | 0.3346 |
| Phoebehungmoensis | 0.4900* | 54.74* | 18.02* | 0.1933* | 0.8283* | 78.57* | 0.3097* |
| Phoebetavoyana | 0.5382* | 51.23 | 16.19 | 0.2087 | 0.9238 | 64.76 | 0.3406 |
| machilus | 0.5626 | 42.81 | 10.74 | 0.2452 | 1.152 | 34.92 | 0.3826 |
| Machiluschinensis | 0.5643* | 31.43* | 10.27* | 0.2350* | 1.046* | 37.02* | 0.4035* |
| Machiluscicatricosa | 0.5643* | 51.24* | 12.20* | 0.2306* | 1.072* | 40.06* | 0.3732* |
| Machilusfoonchewii | 0.5643* | 42.75 | 10.75 | 0.2454 | 1.153 | 34.79 | 0.3832 |
| Machilusgamblei | 0.5643* | 42.75 | 10.75 | 0.2454 | 1.153 | 34.79 | 0.3832 |
| Machilusmonticola | 0.5643* | 41.76* | 8.163* | 0.2814* | 1.392* | 19.96* | 0.3777* |
| Machiluspomifera | 0.5643* | 42.75 | 10.75 | 0.2454 | 1.153 | 34.79 | 0.3832 |
| Machilusrobusta | 0.5643* | 42.75 | 10.75 | 0.2454 | 1.153 | 34.79 | 0.3832 |
| Machilussp | 0.5643* | 42.75 | 10.75 | 0.2454 | 1.153 | 34.79 | 0.3832 |
| Machilusvelutina | 0.5643* | 42.75 | 10.75 | 0.2454 | 1.153 | 34.79 | 0.3832 |
| neolitsea | 0.5343 | 41.99 | 10.39 | 0.1905 | 1.030 | 35.10 | 0.4224 |
| Neolitseacambodiana | 0.5415* | 49.33* | 9.229* | 0.1990* | 0.9128* | 43.39* | 0.4360* |
| Neolitseachui | 0.5415* | 41.75 | 10.41 | 0.1910 | 1.031 | 34.59 | 0.4249 |
| Neolitseaellipsoidea | 0.4527* | 65.40* | 8.420* | 0.1981* | 1.138* | 27.08* | 0.4204* |
| Neolitseaoblongifolia | 0.5300* | 29.38* | 13.64* | 0.1651* | 0.9812* | 42.86* | 0.3957* |
| Neolitseaovatifolia | 0.5415* | 21.77* | 8.694* | 0.2006* | 0.9572* | 34.23* | 0.4754* |
| Neolitseaphanerophlebia | 0.5900* | 40.11 | 10.54 | 0.1946 | 1.042 | 31.13 | 0.4415 |
| Neolitseapulchella | 0.5415* | 43.90* | 11.50* | 0.1740* | 1.115* | 31.73* | 0.3713* |
| alseodaphne | 0.5678 | 61.62 | 11.00 | 0.2174 | 1.384 | 27.71 | 0.3761 |
| Alseodaphnehainanensis | 0.5970* | 67.65* | 10.44* | 0.1881* | 1.542* | 19.80* | 0.3629* |
| lindera | 0.4822 | 49.97 | 14.13 | 0.3554 | 1.115 | 44.30 | 0.4154 |
| Linderacommunis | 0.4519* | 50.99 | 14.04 | 0.3532 | 1.109 | 46.45 | 0.4051 |
| Linderakwangtungensis | 0.5778* | 52.23* | 20.47* | 0.1408* | 0.7655* | 65.38* | 0.4106* |
| Linderametcalfiana | 0.4519* | 50.99 | 14.04 | 0.3532 | 1.109 | 46.45 | 0.4051 |
| Linderanacusua | 0.4519* | 37.51* | 14.26* | 0.7712* | 1.649* | 15.61* | 0.5077* |
| Linderarobusta | 0.4519* | 54.19* | 9.894* | 0.2263* | 0.9277* | 50.51* | 0.3614* |
| cinnamomum | 0.4926 | 57.99 | 8.895 | 0.2393 | 0.9560 | 49.96 | 0.3951 |
| Cinnamomumbejolghota | 0.4671* | 58.85 | 8.825 | 0.2374 | 0.9507 | 51.77 | 0.3864 |
| Cinnamomumburmannii | 0.4900* | 36.26* | 8.503* | 0.1902* | 0.8803* | 44.75* | 0.4905* |
| Cinnamomumliangii | 0.4671* | 58.85 | 8.825 | 0.2374 | 0.9507 | 51.77 | 0.3864 |
| Cinnamomumparthenoxylon | 0.5800* | 73.82* | 9.368* | 0.2447* | 0.9115* | 65.62* | 0.3101* |
| Cinnamomumrigidissimum | 0.4671* | 63.39* | 7.393* | 0.2658* | 0.9922* | 40.30* | 0.4002* |
| Cinnamomumsubavenium | 0.5000* | 57.74 | 8.915 | 0.2398 | 0.9576 | 49.43 | 0.3977 |
| Cinnamomumtsoi | 0.4671* | 58.85 | 8.825 | 0.2374 | 0.9507 | 51.77 | 0.3864 |
| cryptocarya | 0.5675 | 49.43 | 10.08 | 0.2354 | 1.162 | 28.87 | 0.4376 |
| Cryptocaryachinensis | 0.5000* | 48.49* | 10.17* | 0.2062* | 1.413* | 20.35* | 0.3618* |
| Cryptocaryachingii | 0.5400* | 46.83* | 8.966* | 0.2450* | 0.9028* | 43.63* | 0.4632* |
| Cryptocaryadensiflora | 0.5357* | 50.51 | 9.994 | 0.2330 | 1.155 | 31.13 | 0.4267 |
| Cryptocaryaimpressinervia | 0.5613* | 49.64 | 10.06 | 0.2349 | 1.161 | 29.31 | 0.4354 |
| Cryptocaryamaclurei | 0.5613* | 49.64 | 10.06 | 0.2349 | 1.161 | 29.31 | 0.4354 |
| Cryptocaryametcalfiana | 0.7600* | 42.93 | 10.61 | 0.2494 | 1.202 | 15.17 | 0.5034 |

| | | | | | | | |
|---|---|---|---|---|---|---|---|
| Cryptocaryasp9 | 0.5613* | 49.64 | 10.06 | 0.2349 | 1.161 | 29.31 | 0.4354 |
| litsea | 0.4348 | 61.01 | 11.81 | 0.2379 | 1.047 | 38.99 | 0.4436 |
| Litseabaviensis | 0.4352* | 61.42* | 13.56* | 0.2447* | 1.212* | 35.46* | 0.3002* |
| Litseacubeba | 0.3100* | 65.23 | 11.46 | 0.2288 | 1.021 | 47.87 | 0.4009 |
| Litseaelongata | 0.4255* | 61.33 | 11.78 | 0.2372 | 1.045 | 39.65 | 0.4404 |
| Litsealancilimba | 0.5317* | 57.74 | 12.07 | 0.2450 | 1.067 | 32.10 | 0.4768 |
| Litseamonopetala | 0.4228* | 61.42 | 11.77 | 0.2370 | 1.044 | 39.84 | 0.4395 |
| Litseapseudoelongata | 0.4255* | 61.33 | 11.78 | 0.2372 | 1.045 | 39.65 | 0.4404 |
| Litseavariabilischinensis | 0.4255* | 61.33 | 11.78 | 0.2372 | 1.045 | 39.65 | 0.4404 |
| Litseavariabilis | 0.4255* | 49.43* | 11.83* | 0.2440* | 1.053* | 39.19* | 0.4981* |
| Litseaverticillata | 0.4255* | 73.14* | 11.31* | 0.2235* | 0.7952* | 43.78* | 0.5604* |
| beilschmiedia | 0.5632 | 79.07 | 8.116 | 0.2907 | 1.450 | 18.06 | 0.4461 |
| Beilschmiediaappendiculata | 0.5631* | 79.07 | 8.116 | 0.2907 | 1.450 | 18.07 | 0.4461 |
| Beilschmiediaglauca | 0.5631* | 79.07 | 8.116 | 0.2907 | 1.450 | 18.07 | 0.4461 |
| Beilschmiediaintermedia | 0.5148* | 80.71 | 7.983 | 0.2871 | 1.440 | 21.51 | 0.4295 |
| Beilschmiedialaevis | 0.5631* | 84.75* | 7.437* | 0.3014* | 1.525* | 15.16* | 0.4442* |
| Beilschmiediaobconica | 0.5631* | 79.07 | 8.116 | 0.2907 | 1.450 | 18.07 | 0.4461 |
| Beilschmiediapercoriacea | 0.5631* | 79.07 | 8.116 | 0.2907 | 1.450 | 18.07 | 0.4461 |
| Beilschmiediapergamentacea | 0.5631* | 79.07 | 8.116 | 0.2907 | 1.450 | 18.07 | 0.4461 |
| Beilschmiediaroxburghiana | 0.5631* | 79.07 | 8.116 | 0.2907 | 1.450 | 18.07 | 0.4461 |
| Beilschmiediasp | 0.6063* | 77.61 | 8.234 | 0.2938 | 1.459 | 14.99 | 0.4608 |
| Beilschmiediasp0502 | 0.5631* | 79.07 | 8.116 | 0.2907 | 1.450 | 18.07 | 0.4461 |
| Beilschmiediasp4 | 0.5631* | 79.07 | 8.116 | 0.2907 | 1.450 | 18.07 | 0.4461 |
| Beilschmiediasp6 | 0.5631* | 79.07 | 8.116 | 0.2907 | 1.450 | 18.07 | 0.4461 |
| Beilschmiediatsangii | 0.5631* | 79.07 | 8.116 | 0.2907 | 1.450 | 18.07 | 0.4461 |
| Beilschmiediatungfangensis | 0.5631* | 79.07 | 8.116 | 0.2907 | 1.450 | 18.07 | 0.4461 |
| Beilschmiediawangii | 0.5631* | 79.07 | 8.116 | 0.2907 | 1.450 | 18.07 | 0.4461 |
| endiandra | 0.5923 | 72.40 | 8.875 | 0.2820 | 1.381 | 18.90 | 0.4579 |
| Endiandrahainanensis | 0.6150* | 71.64 | 8.937 | 0.2837 | 1.386 | 17.28 | 0.4657 |
| Lauraceaesp5 | 0.5643* | 57.61 | 10.98 | 0.2450 | 1.159 | 30.32 | 0.4467 |
| Lauraceaesp8 | 0.5643* | 57.61 | 10.98 | 0.2450 | 1.159 | 30.32 | 0.4467 |
| austrobaileyales | 0.5515 | 67.54 | 11.86 | 0.2264 | 1.142 | 39.58 | 0.4220 |
| schisandraceae | 0.5698 | 66.92 | 11.92 | 0.2278 | 1.146 | 38.27 | 0.4283 |
| illicium | 0.5759 | 66.71 | 11.93 | 0.2282 | 1.147 | 37.84 | 0.4304 |
| Illiciumsp1 | 0.5790* | 66.61 | 11.94 | 0.2284 | 1.148 | 37.62 | 0.4314 |
| Illiciumternstroemioides | 0.5790* | 66.61 | 11.94 | 0.2284 | 1.148 | 37.62 | 0.4314 |
| gymnosperms | 0.5044 | 69.13 | 11.74 | 0.2230 | 1.133 | 42.93 | 0.4059 |
| pinales | 0.4858 | 69.75 | 11.68 | 0.2216 | 1.129 | 44.25 | 0.3995 |
| pinaceae | 0.5028 | 69.18 | 11.73 | 0.2229 | 1.132 | 43.05 | 0.4053 |
| pinus | 0.5537 | 67.46 | 11.87 | 0.2266 | 1.143 | 39.42 | 0.4228 |
| Pinuscaribaea | 0.5707* | 66.89 | 11.92 | 0.2278 | 1.146 | 38.21 | 0.4286 |
| podocarpaceae | 0.4644 | 70.48 | 11.63 | 0.2201 | 1.124 | 45.77 | 0.3922 |
| podocarpus | 0.4769 | 70.06 | 11.66 | 0.2210 | 1.127 | 44.89 | 0.3965 |
| Podocarpusneriifolius | 0.4769* | 70.06 | 11.66 | 0.2210 | 1.127 | 44.89 | 0.3965 |
| dacrycarpus | 0.4463 | 71.09 | 11.58 | 0.2187 | 1.120 | 47.06 | 0.3860 |
| Dacrycarpusimbricatus | 0.4133* | 72.20 | 11.48 | 0.2163 | 1.113 | 49.41 | 0.3747 |
| dacrydium | 0.5502 | 67.58 | 11.86 | 0.2263 | 1.142 | 39.67 | 0.4216 |
| Dacrydiumpectinatum | 0.5856* | 66.38 | 11.96 | 0.2289 | 1.150 | 37.15 | 0.4337 |
| cupressaceae | 0.3732 | 73.56 | 11.37 | 0.2134 | 1.105 | 52.26 | 0.3610 |
| cunninghamia | 0.3445 | 74.53 | 11.30 | 0.2113 | 1.099 | 54.31 | 0.3512 |
| Cunninghamialanceolata | 0.3157* | 75.50 | 11.22 | 0.2092 | 1.093 | 56.36 | 0.3413 |
| monilophyte | 0.5347 | 68.10 | 11.82 | 0.2252 | 1.139 | 40.77 | 0.4163 |
| cyatheales | 0.5619 | 67.18 | 11.89 | 0.2272 | 1.145 | 38.84 | 0.4256 |
| cyatheaceae | 0.5736 | 66.79 | 11.93 | 0.2280 | 1.147 | 38.00 | 0.4296 |
| Alsophila | 0.5775 | 66.66 | 11.94 | 0.2283 | 1.148 | 37.73 | 0.4309 |
| Alsophilapodophylla | 0.5814* | 66.53 | 11.95 | 0.2286 | 1.149 | 37.45 | 0.4322 |

#### Cross-validation details

In addition to estimating missing values, PhyloPars performs cross-validation: it temporarily excludes each observation from the available information, in order to calculate its best estimate given al observations and the optimal phylogenetic covariances. This provides a detailed estimate of the error and bias one may expect for the estimated missing feature values. The distributions of estima calculated through cross-validation are shown below; these are compared with two null models (red and green curves) in order to assess the improvement resulting from the PhyloPars evolutionar

##### Meanwd

**Cross-validation error**

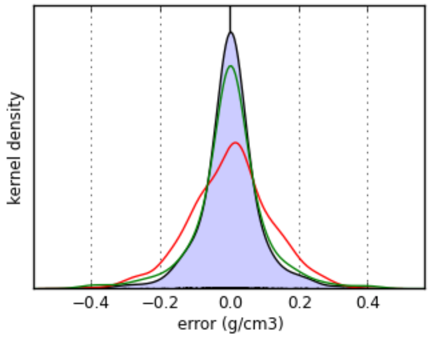


| | evolutionary model | mean model | nearest neighbor model |
|---|---|---|---|
| mean bias | **-0.000606 g/cm3** | 3.3e-16 g/cm3 | -0.000865 g/cm3 |
| mean error | **0.0497 g/cm3** | 0.0884 g/cm3 | 0.0591 g/cm3 |

**Meanleafarea**

**Cross-validation error**

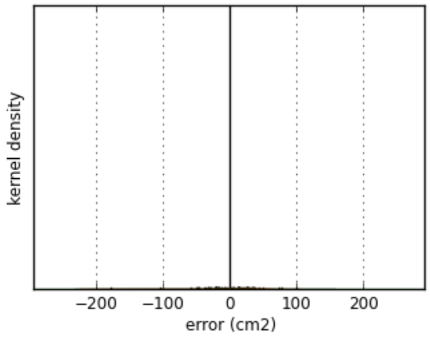


| | evolutionary model | mean model | nearest neighbor model |
|---|---|---|---|
| mean bias | **-1.43 cm2** | 1.02e-14 cm2 | -0.682 cm2 |
| mean error | **29.3 cm2** | 32.3 cm2 | 33.9 cm2 |

**Meansla**

**Cross-validation error**

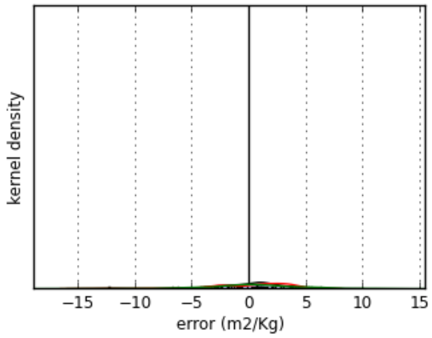


| | evolutionary model | mean model | nearest neighbor model |
|---|---|---|---|
| mean bias | **-0.0925 m2/Kg** | 4.57e-15 m2/Kg | 0.106 m2/Kg |
| mean error | **2.4 m2/Kg** | 2.74 m2/Kg | 3.22 m2/Kg |

**Meanleafthickness**

**Cross-validation error**

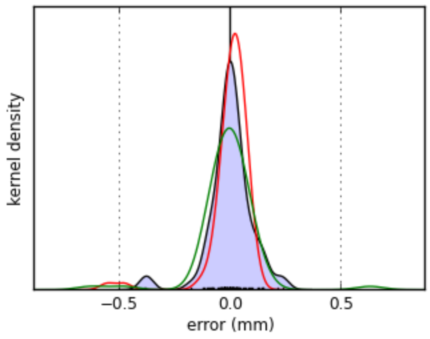


| | evolutionary model | mean model | nearest neighbor model |
|---|---|---|---|
| mean bias | **5.55e-05 mm** | -7.82e-17 mm | -0.01 mm |
| mean error | **0.064 mm** | 0.0538 mm | 0.0724 mm |

**Meanrootavgdiam**

**Cross-validation error**

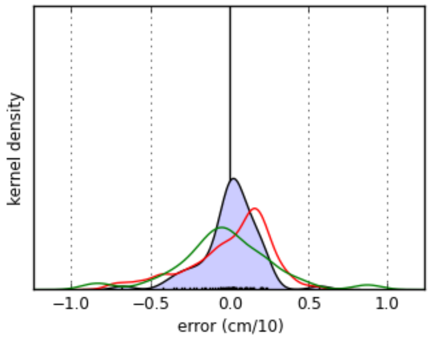


| | evolutionary model | mean model | nearest neighbor model |
|---|---|---|---|
| mean bias | **0.00097 cm/10** | 1.09e-17 cm/10 | -0.0363 cm/10 |
| mean error | **0.124 cm/10** | 0.206 cm/10 | 0.228 cm/10 |

**Meanspecificrootlength**

**Cross-validation error**

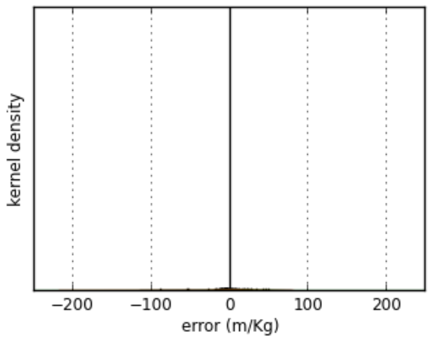


| | evolutionary model | mean model | nearest neighbor model |
|---|---|---|---|
| mean bias | **0.0833 m/Kg** | 0 m/Kg | 2.39 m/Kg |
| mean error | **13.3 m/Kg** | 19.5 m/Kg | 24.1 m/Kg |

**Meanroottd**

**Cross-validation error**

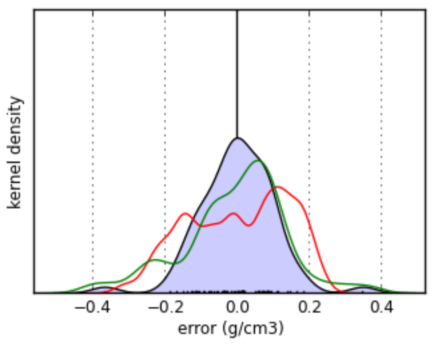


| | evolutionary model | mean model | nearest neighbor model |
|---|---|---|---|
| mean bias | **-0.00175 g/cm3** | 5.79e-17 g/cm3 | -0.00891 g/cm3 |
| mean error | **0.076 g/cm3** | 0.121 g/cm3 | 0.109 g/cm3 |

#### Meanbranchiness

**Cross-validation error**

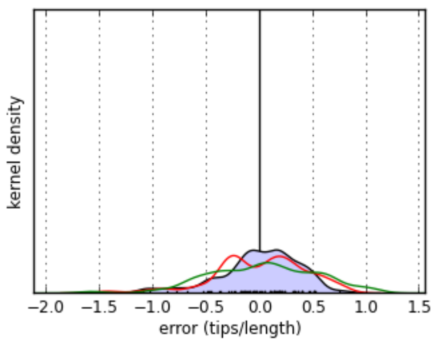


| | evolutionary model | mean model | nearest neighbor model |
|---|---|---|---|
| mean bias | **-0.00118 tips/length** | 7.07e-16 tips/length | 0.0244 tips/length |
| mean error | **0.287 tips/length** | 0.344 tips/length | 0.406 tips/length |

The PhyloPars tool is (c) Jorn Bruggeman 2019. If you are experiencing problems with this tool, please contact jbr [at] pml.ac.uk or bwbrandt [at] few.vu.nl.

### PhyloPars results

### Model uses traits for individuals only found in the secondary forest portion of the transect (i.e. beginning of the JFL transect)

#### Contents

Phylogenetic variability
Estimates for missing parameters
Cross-validation details

**Please cite:** Bruggeman J, Heringa J and Brandt BW. (2009) PhyloPars: estimation of missing parameter values using phylogeny.
Nucleic Acids Research 37: W179-W184.

#### Phylogenetic variability

PhyloPars first estimates the parameters of the evolutionary model, i.e., the phylogenetic covariances. These are subsequently used to estimate missing feature values and to perform cross-valida

| feature | phylogenetic s.d. | | cross-validation results | | | |
|---|---|---|---|---|---|---|
| | | | error | | bias | |
| meanWD | 0.049 | g/cm3 | 0.0496 | g/cm3 | -0.000581 | g/cm3 |
| meanLeafArea | 27.71 | cm2 | 23.1 | cm2 | -0.579 | cm2 |
| meanSLA | 2.795 | m2/Kg | 2.89 | m2/Kg | -0.0725 | m2/Kg |
| meanLeafThickness | 0.102 | mm | 0.0801 | mm | 0.000495 | mm |
| meanRootAvgDiam | 0.1838 | cm/10 | 0.143 | cm/10 | 0.000635 | cm/10 |
| meanSpecificRootLength | 22.13 | m/Kg | 13.7 | m/Kg | 0.0936 | m/Kg |
| meanRootTD | 0.08217 | g/cm3 | 0.0809 | g/cm3 | -0.00257 | g/cm3 |
| meanBranchiness | 0.312 | tips/length | 0.349 | tips/length | -0.00825 | tips/length |

##### Phylogenetic correlations

| | meanWD | meanLeafArea | meanSLA | meanLeafThickness | meanRootAvgDiam | meanSpecificRootLength | meanRootTD | meanBr |
|---|---|---|---|---|---|---|---|---|
| **meanWD** | 1 | | | | | | | |
| **meanLeafArea** | -0.063 | 1 | | | | | | |
| **meanSLA** | 0.087 | -0.059 | 1 | | | | | |
| **meanLeafThickness** | -0.003 | -0.054 | -0.241 | 1 | | | | |
| **meanRootAvgDiam** | 0.027 | -0.204 | -0.079 | 0.287 | 1 | | | |
| **meanSpecificRootLength** | -0.105 | 0.531 | 0.207 | -0.155 | -0.632 | 1 | | |
| **meanRootTD** | 0.201 | -0.082 | -0.129 | 0.084 | 0.117 | -0.443 | 1 | |
| **meanBranchiness** | 0.172 | 0.278 | 0.090 | -0.271 | -0.373 | 0.111 | 0.243 | 1 |

##### Phylogenetic regression coefficients

Rows represent independent variables and columns dependent variables. A regression of feature *i* on *j* thus can be found at row *j*, column *i*.

| | meanWD | meanLeafArea | meanSLA | meanLeafThickness | meanRootAvgDiam | meanSpecificRootLength | meanRootTD | meanBr |
|---|---|---|---|---|---|---|---|---|
| **meanWD** | 1 | -35.450 | 4.966 | -0.007 | 0.100 | -47.544 | 0.337 | 1.097 |
| **meanLeafArea** | -0.000 | 1 | -0.006 | -0.000 | -0.001 | 0.424 | -0.000 | 0.003 |
| **meanSLA** | 0.002 | -0.588 | 1 | -0.009 | -0.005 | 1.639 | -0.004 | 0.010 |
| **meanLeafThickness** | -0.002 | -14.608 | -6.591 | 1 | 0.516 | -33.543 | 0.067 | -0.828 |
| **meanRootAvgDiam** | 0.007 | -30.704 | -1.204 | 0.159 | 1 | -76.148 | 0.052 | -0.633 |
| **meanSpecificRootLength** | -0.000 | 0.665 | 0.026 | -0.001 | -0.005 | 1 | -0.002 | 0.002 |
| **meanRootTD** | 0.120 | -27.718 | -4.388 | 0.104 | 0.261 | -119.344 | 1 | 0.923 |
| **meanBranchiness** | 0.027 | 24.646 | 0.806 | -0.089 | -0.220 | 7.875 | 0.064 | 1 |

#### Estimates for missing parameters

Using the optimal phylogenetic covariances listed above, PhyloPars estimates the values that were originally missing in the feature matrix. The table below lists all estimated values. If one or more have been provided for a value, this is denoted by a trailing asterisk symbol (*). You can click on an entry to view the contribution of individual observations to the estimate, and to retrieve details su standard deviation of the estimate. You can also download the complete table as a single text file.

| | meanWD (g/cm3) | meanLeafArea (cm2) | meanSLA (m2/Kg) | meanLeafThickness (mm) | meanRootAvgDiam (cm/10) | meanSpecificRootLength (m/Kg) | meanRootTD (g/cm3) | mean (tip |
|---|---|---|---|---|---|---|---|---|

| | | | | | | | |
|---|---|---|---|---|---|---|---|
| root | 0.5269 | 65.56 | 12.54 | 0.2348 | 1.055 | 46.74 | 0.4094 |
| fagales | 0.6168 | 62.38 | 12.14 | 0.1981 | 0.8387 | 58.15 | 0.5807 |
| fagaceae | 0.6190 | 52.88 | 11.50 | 0.1979 | 0.8243 | 50.56 | 0.6324 |
| lithocarpus | 0.6897 | 41.99 | 12.11 | 0.2661 | 0.9440 | 36.57 | 0.6378 |
| Lithocarpusamygdalifolius | 0.7580* | 24.54* | 12.28* | 0.2686* | 0.8195* | 34.95* | 0.7305* |
| Lithocarpusbacgiangensis | 0.6773* | 42.43 | 12.05 | 0.2662 | 0.9428 | 37.16 | 0.6336 |
| Lithocarpusbrachystachyus | 0.6773* | 42.43 | 12.05 | 0.2662 | 0.9428 | 37.16 | 0.6336 |
| Lithocarpuschiungchungensis | 0.6773* | 42.43 | 12.05 | 0.2662 | 0.9428 | 37.16 | 0.6336 |
| Lithocarpuscorneus | 0.8180* | 37.45 | 12.75 | 0.2652 | 0.9569 | 30.47 | 0.6810 |
| Lithocarpuselaeagnifolius | 0.6773* | 42.43 | 12.05 | 0.2662 | 0.9428 | 37.16 | 0.6336 |
| Lithocarpusfenestratus | 0.6773* | 77.81* | 9.665* | 0.2360* | 0.6147* | 74.06* | 0.5505* |
| Lithocarpusfenzelianus | 0.6773* | 38.24* | 6.726* | 0.2573* | 1.012* | 28.23* | 0.5282* |
| Lithocarpushancei | 0.6773* | 29.48* | 24.72* | 0.1373* | 1.017* | 43.43* | 0.4978* |
| Lithocarpushandelianus | 0.7610* | 55.54* | 9.797* | 0.6953* | 1.310* | 14.49* | 0.5898* |
| Lithocarpushowii | 0.6773* | 34.51* | 12.46* | 0.1830* | 0.7426* | 37.12* | 0.7950* |
| Lithocarpuslitseifolius | 0.6773* | 42.43 | 12.05 | 0.2662 | 0.9428 | 37.16 | 0.6336 |
| Lithocarpuslongipedicellatus | 0.5995* | 43.94* | 11.13* | 0.1793* | 1.214* | 15.25* | 0.6772* |
| Lithocarpuspseudovestitus | 0.6773* | 28.00* | 9.888* | 0.2090* | 0.8552* | 42.49* | 0.6808* |
| Lithocarpussp0502 | 0.6773* | 42.43 | 12.05 | 0.2662 | 0.9428 | 37.16 | 0.6336 |
| castanopsis | 0.5502 | 57.93 | 12.18 | 0.1734 | 0.8678 | 48.92 | 0.7506 |
| Castanopsiscarlesii | 0.4408* | 61.81 | 11.64 | 0.1742 | 0.8568 | 54.12 | 0.7138 |
| Castanopsisfissa | 0.4446* | 92.08* | 6.749* | 0.1909* | 0.8780* | 58.79* | 0.6483* |
| Castanopsishainanensis | 0.6473* | 54.49 | 12.66 | 0.1727 | 0.8775 | 44.30 | 0.7833 |
| Castanopsishystrix | 0.5650* | 57.41 | 12.25 | 0.1733 | 0.8692 | 48.21 | 0.7556 |
| Castanopsisjianfenglingensis | 0.5680* | 57.30 | 12.27 | 0.1733 | 0.8695 | 48.07 | 0.7566 |
| Castanopsisjucunda | 0.5680* | 57.30 | 12.27 | 0.1733 | 0.8695 | 48.07 | 0.7566 |
| Castanopsistonkinensis | 0.5680* | 30.22* | 17.47* | 0.1280* | 0.8345* | 44.50* | 0.8792* |
| quercus | 0.7003 | 47.98 | 12.13 | 0.2134 | 0.8719 | 42.96 | 0.6747 |
| Quercusacutissima | 0.7334* | 46.81 | 12.29 | 0.2131 | 0.8752 | 41.38 | 0.6858 |
| Cyclobalanopsis | 0.6076 | 45.56 | 10.95 | 0.1921 | 0.8106 | 45.46 | 0.6539 |
| Cyclobalanopsisbella | 0.6062* | 45.61 | 10.95 | 0.1921 | 0.8104 | 45.53 | 0.6534 |
| Cyclobalanopsisblakei | 0.6062* | 47.71* | 13.55* | 0.1207* | 0.7251* | 42.63* | 0.7559* |
| Cyclobalanopsisedithiae | 0.6062* | 53.48* | 12.50* | 0.1648* | 0.7369* | 43.45* | 0.6638* |
| Cyclobalanopsisfleuryi | 0.6062* | 82.32* | 9.116* | 0.1951* | 0.8842* | 29.66* | 0.7938* |
| Cyclobalanopsishui | 0.6062* | 28.83* | 7.788* | 0.2947* | 0.8210* | 48.86* | 0.5960* |
| Cyclobalanopsisneglecta | 0.6062* | 14.42* | 12.53* | 0.1656* | 0.8387* | 61.18* | 0.4899* |
| Cyclobalanopsispatelliformis | 0.6062* | 39.14* | 9.706* | 0.2061* | 0.8440* | 41.75* | 0.6464* |
| Cyclobalanopsisphanera | 0.6062* | 45.61 | 10.95 | 0.1921 | 0.8104 | 45.53 | 0.6534 |
| betulaceae | 0.5671 | 71.52 | 11.75 | 0.1915 | 0.7834 | 68.07 | 0.5675 |
| betula | 0.5567 | 71.89 | 11.70 | 0.1915 | 0.7824 | 68.57 | 0.5640 |
| Betulaalnoides | 0.5515* | 72.08 | 11.67 | 0.1916 | 0.7819 | 68.82 | 0.5622 |
| juglandaceae | 0.5280 | 97.08 | 11.92 | 0.1780 | 0.6935 | 92.54 | 0.5103 |
| engelhardia | 0.4916 | 114.5 | 11.99 | 0.1690 | 0.6326 | 109.3 | 0.4688 |
| Engelhardiahainanensis | 0.4879* | 114.6 | 11.97 | 0.1691 | 0.6322 | 109.5 | 0.4675 |
| Engelhardiaroxburghiana | 0.4879* | 91.77* | 10.44* | 0.1122* | 0.6855* | 50.40* | 0.6182* |
| Engelhardiaspicata | 0.4879* | 212.8* | 15.09* | 0.1803* | 0.4544* | 224.5* | 0.3024* |
| EngelhardiaspicatavarAceriflora | 0.4879* | 114.6 | 11.97 | 0.1691 | 0.6322 | 109.5 | 0.4675 |
| Engelhardiaunijuga | 0.4879* | 47.36* | 10.51* | 0.2100* | 0.7280* | 61.17* | 0.4674* |
| rosales | 0.6484 | 59.96 | 12.96 | 0.2051 | 0.8967 | 57.13 | 0.5442 |
| moraceae | 0.5627 | 49.68 | 13.24 | 0.2178 | 0.9459 | 63.89 | 0.4610 |
| broussonetia | 0.3365 | 67.74 | 12.45 | 0.1969 | 0.8265 | 78.95 | 0.4758 |
| Broussonetiapapyrifera | 0.2900* | 69.39 | 12.22 | 0.1972 | 0.8218 | 81.16 | 0.4602 |
| ficus | 0.3949 | 75.71 | 13.07 | 0.1739 | 0.7357 | 80.48 | 0.5864 |
| Ficusaltissima | 0.4730* | 72.95 | 13.46 | 0.1734 | 0.7435 | 76.77 | 0.6127 |
| Ficusauriculata | 0.4680* | 73.12 | 13.44 | 0.1734 | 0.7430 | 77.01 | 0.6110 |
| Ficusfistulosa | 0.3800* | 76.24 | 13.00 | 0.1740 | 0.7342 | 81.19 | 0.5814 |
| Ficusformosana | 0.4055* | 19.37* | 13.75* | 0.1929* | 0.5279* | 105.8* | 0.6085* |
| Ficusglaberrima | 0.4055* | 75.34 | 13.13 | 0.1738 | 0.7367 | 79.98 | 0.5900 |
| Ficusheteropleura | 0.4055* | 75.34 | 13.13 | 0.1738 | 0.7367 | 79.98 | 0.5900 |
| Ficushirta | 0.4055* | 197.5* | 11.62* | 0.1144* | 0.6138* | 90.73* | 0.6988* |
| Ficuslangkokensis | 0.4055* | 75.34 | 13.13 | 0.1738 | 0.7367 | 79.98 | 0.5900 |
| Ficusnervosa | 0.2800* | 79.79 | 12.50 | 0.1747 | 0.7241 | 85.95 | 0.5477 |
| Ficuspandurata | 0.4055* | 75.34 | 13.13 | 0.1738 | 0.7367 | 79.98 | 0.5900 |

| | | | | | | | |
|---|---|---|---|---|---|---|---|
| Ficuspubigera | 0.4055* | 75.34 | 13.13 | 0.1738 | 0.7367 | 79.98 | 0.5900 |
| Ficussimplicissima | 0.4055* | 75.34 | 13.13 | 0.1738 | 0.7367 | 79.98 | 0.5900 |
| Ficussubpisocarpa | 0.4055* | 75.34 | 13.13 | 0.1738 | 0.7367 | 79.98 | 0.5900 |
| Ficustinctoria | 0.4055* | 75.34 | 13.13 | 0.1738 | 0.7367 | 79.98 | 0.5900 |
| Ficustuphapensis | 0.4055* | 75.34 | 13.13 | 0.1738 | 0.7367 | 79.98 | 0.5900 |
| Ficusvariegata | 0.3270* | 78.12 | 12.74 | 0.1744 | 0.7288 | 83.71 | 0.5635 |
| Ficusvariolosa | 0.4055* | 75.34 | 13.13 | 0.1738 | 0.7367 | 79.98 | 0.5900 |
| Ficusvasculosa | 0.3000* | 17.87* | 13.66* | 0.2037* | 1.010* | 50.53* | 0.4725* |
| antiaris | 0.3884 | 70.92 | 12.87 | 0.1852 | 0.7833 | 78.64 | 0.5387 |
| Antiaristoxicaria | 0.3831* | 71.11 | 12.85 | 0.1853 | 0.7828 | 78.89 | 0.5369 |
| artocarpus | 0.4959 | 29.99 | 15.12 | 0.1861 | 1.187 | 56.79 | 0.4346 |
| Artocarpusnitidus | 0.4800* | 30.56 | 15.04 | 0.1862 | 1.186 | 57.55 | 0.4293 |
| Artocarpusstyracifolius | 0.5077* | 18.55* | 16.29* | 0.1699* | 1.312* | 51.10* | 0.4366* |
| Artocarpustonkinensis | 0.4667* | 31.03 | 14.98 | 0.1863 | 1.184 | 58.18 | 0.4248 |
| streblus | 0.6898 | 48.52 | 12.74 | 0.2466 | 0.8945 | 61.36 | 0.4494 |
| Streblusilicifolius | 0.7321* | 47.02 | 12.95 | 0.2463 | 0.8988 | 59.35 | 0.4636 |
| Streblusindicus | 0.7321* | 50.36* | 11.81* | 0.2761* | 0.8347* | 62.86* | 0.4092* |
| Streblustaxoides | 0.7321* | 47.02 | 12.95 | 0.2463 | 0.8988 | 59.35 | 0.4636 |
| urticaceae | 0.4506 | 56.32 | 12.54 | 0.2162 | 0.9231 | 68.68 | 0.4341 |
| Oreocnide | 0.3900 | 58.47 | 12.24 | 0.2166 | 0.9170 | 71.56 | 0.4137 |
| Oreocnidetonkinensis | 0.3294* | 60.62 | 11.94 | 0.2170 | 0.9109 | 74.44 | 0.3933 |
| debregeasia | 0.3536 | 59.76 | 12.06 | 0.2169 | 0.9133 | 73.29 | 0.4015 |
| Debregeasiasquamata | 0.3294* | 60.62 | 11.94 | 0.2170 | 0.9109 | 74.44 | 0.3933 |
| cannabaceae | 0.6187 | 53.03 | 13.23 | 0.2126 | 0.9284 | 60.16 | 0.5016 |
| aphananthe | 0.6543 | 51.76 | 13.41 | 0.2123 | 0.9320 | 58.46 | 0.5136 |
| Aphananthecuspidata | 0.6900* | 50.50 | 13.59 | 0.2120 | 0.9355 | 56.76 | 0.5256 |
| celtis | 0.6608 | 51.53 | 13.44 | 0.2122 | 0.9326 | 58.15 | 0.5158 |
| Celtisphilippensis | 0.7030* | 50.04 | 13.65 | 0.2119 | 0.9369 | 56.15 | 0.5299 |
| gironniera | 0.5141 | 56.73 | 12.71 | 0.2133 | 0.9179 | 65.13 | 0.4664 |
| Gironnierasubaequalis | 0.4618* | 58.59 | 12.45 | 0.2137 | 0.9126 | 67.62 | 0.4488 |
| rhamnaceae | 0.7808 | 52.61 | 13.75 | 0.2066 | 0.9216 | 51.37 | 0.5779 |
| rhamnella | 0.7594 | 53.37 | 13.65 | 0.2067 | 0.9194 | 52.40 | 0.5706 |
| Rhamnellarubrinervis | 0.7421* | 53.98 | 13.56 | 0.2068 | 0.9177 | 53.22 | 0.5648 |
| ventilago | 0.9216 | 47.62 | 14.45 | 0.2055 | 0.9357 | 44.68 | 0.6252 |
| Ventilagoleiocarpa | 0.9800* | 45.55 | 14.74 | 0.2051 | 0.9415 | 41.91 | 0.6449 |
| rosaceae | 0.6498 | 59.92 | 12.96 | 0.2051 | 0.8968 | 57.07 | 0.5446 |
| photinia | 0.7217 | 57.37 | 13.32 | 0.2046 | 0.9041 | 53.65 | 0.5688 |
| Photiniabenthamiana | 0.7235* | 57.30 | 13.33 | 0.2046 | 0.9042 | 53.57 | 0.5694 |
| Photiniaprunifolia | 0.7235* | 57.30 | 13.33 | 0.2046 | 0.9042 | 53.57 | 0.5694 |
| eriobotrya | 0.7683 | 55.71 | 13.55 | 0.2042 | 0.9087 | 51.43 | 0.5845 |
| Eriobotryadeflexa | 0.7755* | 55.46 | 13.59 | 0.2042 | 0.9095 | 51.09 | 0.5869 |
| laurocerasus | 0.6522 | 59.83 | 12.97 | 0.2051 | 0.8971 | 56.95 | 0.5454 |
| Laurocerasusphaeosticta | 0.6481* | 59.98 | 12.95 | 0.2051 | 0.8967 | 57.15 | 0.5441 |
| pygeum | 0.6522 | 59.83 | 12.97 | 0.2051 | 0.8971 | 56.95 | 0.5454 |
| Pygeumtopengii | 0.6481* | 59.98 | 12.95 | 0.2051 | 0.8967 | 57.15 | 0.5441 |
| Rhaphiolepis | 0.6487 | 59.96 | 12.96 | 0.2051 | 0.8967 | 57.12 | 0.5442 |
| Rhaphiolepisferruginea | 0.6481* | 59.98 | 12.95 | 0.2051 | 0.8967 | 57.15 | 0.5441 |
| Rhaphiolepisindica | 0.6481* | 59.98 | 12.95 | 0.2051 | 0.8967 | 57.15 | 0.5441 |
| fabales | 0.6463 | 66.05 | 12.92 | 0.2027 | 0.8950 | 56.14 | 0.5470 |
| fabaceae | 0.6541 | 65.77 | 12.96 | 0.2026 | 0.8958 | 55.77 | 0.5497 |
| archidendron | 0.4954 | 71.40 | 12.17 | 0.2038 | 0.8799 | 63.32 | 0.4962 |
| Archidendronclypearia | 0.3233* | 77.50 | 11.32 | 0.2050 | 0.8626 | 71.50 | 0.4384 |
| Archidendronlucidum | 0.5814* | 68.35 | 12.60 | 0.2032 | 0.8885 | 59.23 | 0.5252 |
| Archidendronutile | 0.5814* | 68.35 | 12.60 | 0.2032 | 0.8885 | 59.23 | 0.5252 |
| albizia | 0.4780 | 72.02 | 12.09 | 0.2039 | 0.8781 | 64.14 | 0.4904 |
| Albiziachinensis | 0.3000* | 78.33 | 11.20 | 0.2052 | 0.8603 | 72.60 | 0.4305 |
| Albiziaodoratissima | 0.6385* | 66.33 | 12.88 | 0.2027 | 0.8942 | 56.51 | 0.5444 |
| peltophorum | 0.8257 | 59.69 | 13.81 | 0.2014 | 0.9130 | 47.61 | 0.6074 |
| Peltophorumtonkinense | 0.8486* | 58.88 | 13.93 | 0.2012 | 0.9153 | 46.52 | 0.6151 |
| sindora | 0.8410 | 59.15 | 13.89 | 0.2013 | 0.9146 | 46.88 | 0.6126 |
| Sindoraglabra | 0.8486* | 58.88 | 13.93 | 0.2012 | 0.9153 | 46.52 | 0.6151 |
| ormosia | 0.6027 | 67.60 | 12.71 | 0.2030 | 0.8906 | 58.21 | 0.5324 |
| Ormosiabalansae | 0.4505* | 72.99 | 11.95 | 0.2041 | 0.8754 | 65.45 | 0.4811 |
| Ormosiafordiana | 0.8486* | 58.88 | 13.93 | 0.2012 | 0.9153 | 46.52 | 0.6151 |

| | | | | | | | |
|---|---|---|---|---|---|---|---|
| Ormosiapinnata | 0.5683* | 68.81 | 12.53 | 0.2033 | 0.8872 | 59.85 | 0.5208 |
| Ormosiasemicastrata | 0.6213* | 66.94 | 12.80 | 0.2029 | 0.8925 | 57.33 | 0.5386 |
| Ormosiaxylocarpa | 0.5055* | 71.04 | 12.22 | 0.2037 | 0.8809 | 62.83 | 0.4997 |
| dalbergia | 0.6828 | 64.76 | 13.10 | 0.2024 | 0.8987 | 54.41 | 0.5593 |
| Dalbergiahainanensis | 0.6887* | 64.55 | 13.13 | 0.2024 | 0.8993 | 54.12 | 0.5613 |
| polygalaceae | 0.6705 | 65.19 | 13.04 | 0.2025 | 0.8974 | 54.99 | 0.5552 |
| xanthophyllum | 0.6786 | 64.91 | 13.08 | 0.2025 | 0.8983 | 54.61 | 0.5579 |
| Xanthophyllumhainanense | 0.6867* | 64.62 | 13.12 | 0.2024 | 0.8991 | 54.22 | 0.5606 |
| malpighiales | 0.6420 | 69.55 | 13.02 | 0.2027 | 0.9047 | 55.78 | 0.5383 |
| euphorbiaceae | 0.5468 | 72.92 | 12.55 | 0.2034 | 0.8951 | 60.31 | 0.5063 |
| mallotus | 0.4875 | 75.03 | 12.25 | 0.2038 | 0.8892 | 63.13 | 0.4863 |
| Mallotusanomalus | 0.5038* | 74.45 | 12.33 | 0.2037 | 0.8908 | 62.35 | 0.4918 |
| Mallotuspaniculatus | 0.3450* | 80.08 | 11.54 | 0.2048 | 0.8749 | 69.91 | 0.4384 |
| Mallotusphilippensis | 0.6033* | 70.92 | 12.83 | 0.2030 | 0.9008 | 57.62 | 0.5253 |
| Mallotusyunnanensis | 0.5038* | 74.45 | 12.33 | 0.2037 | 0.8908 | 62.35 | 0.4918 |
| macaranga | 0.4576 | 76.09 | 12.10 | 0.2040 | 0.8862 | 64.55 | 0.4762 |
| Macarangadenticulata | 0.4335* | 76.94 | 11.98 | 0.2042 | 0.8837 | 65.70 | 0.4681 |
| hancea | 0.5215 | 73.82 | 12.42 | 0.2036 | 0.8926 | 61.52 | 0.4977 |
| Hanceahookeriana | 0.5431* | 73.06 | 12.53 | 0.2034 | 0.8947 | 60.49 | 0.5050 |
| cleidion | 0.5048 | 74.41 | 12.34 | 0.2037 | 0.8909 | 62.31 | 0.4921 |
| Cleidionbrevipetiolatum | 0.5167* | 73.99 | 12.40 | 0.2036 | 0.8921 | 61.74 | 0.4961 |
| koilodepas | 0.5339 | 73.38 | 12.48 | 0.2035 | 0.8938 | 60.92 | 0.5019 |
| Koilodepashainanense | 0.5431* | 73.06 | 12.53 | 0.2034 | 0.8947 | 60.49 | 0.5050 |
| claoxylon | 0.3813 | 78.79 | 11.73 | 0.2046 | 0.8785 | 68.18 | 0.4506 |
| Claoxylonindicum | 0.3550* | 79.72 | 11.59 | 0.2048 | 0.8759 | 69.43 | 0.4417 |
| alchornea | 0.4163 | 77.55 | 11.90 | 0.2043 | 0.8820 | 66.52 | 0.4624 |
| Alchornearugosa | 0.4084* | 77.83 | 11.86 | 0.2044 | 0.8812 | 66.89 | 0.4597 |
| croton | 0.5191 | 73.91 | 12.41 | 0.2036 | 0.8923 | 61.63 | 0.4969 |
| Crotoncascarilloides | 0.5104* | 74.21 | 12.37 | 0.2037 | 0.8915 | 62.04 | 0.4940 |
| Crotonlaevigatus | 0.5300* | 73.52 | 12.46 | 0.2035 | 0.8934 | 61.11 | 0.5006 |
| ostodes | 0.3628 | 79.45 | 11.63 | 0.2047 | 0.8766 | 69.06 | 0.4444 |
| Ostodespaniculata | 0.3390* | 80.29 | 11.51 | 0.2049 | 0.8743 | 70.19 | 0.4363 |
| suregada | 0.5980 | 71.11 | 12.80 | 0.2030 | 0.9002 | 57.88 | 0.5235 |
| Suregadamultiflora | 0.6470* | 69.37 | 13.04 | 0.2027 | 0.9052 | 55.55 | 0.5400 |
| triadica | 0.5409 | 73.13 | 12.52 | 0.2034 | 0.8945 | 60.59 | 0.5043 |
| Triadicacochinchinensis | 0.5431* | 73.06 | 12.53 | 0.2034 | 0.8947 | 60.49 | 0.5050 |
| endospermum | 0.3836 | 78.71 | 11.74 | 0.2046 | 0.8787 | 68.07 | 0.4514 |
| Endospermumchinense | 0.3475* | 79.99 | 11.56 | 0.2048 | 0.8751 | 69.79 | 0.4392 |
| Trevia | 0.5450 | 72.99 | 12.54 | 0.2034 | 0.8949 | 60.40 | 0.5057 |
| Trevianudiflora | 0.5431* | 73.06 | 12.53 | 0.2034 | 0.8947 | 60.49 | 0.5050 |
| Euphorbiaceaesp9 | 0.5431* | 73.06 | 12.53 | 0.2034 | 0.8947 | 60.49 | 0.5050 |
| Lasiococca | 0.5450 | 72.99 | 12.54 | 0.2034 | 0.8949 | 60.40 | 0.5057 |
| Lasiococcacomberi | 0.5431* | 73.06 | 12.53 | 0.2034 | 0.8947 | 60.49 | 0.5050 |
| Epiprinus | 0.5450 | 72.99 | 12.54 | 0.2034 | 0.8949 | 60.40 | 0.5057 |
| Epiprinussiletianus | 0.5431* | 73.06 | 12.53 | 0.2034 | 0.8947 | 60.49 | 0.5050 |
| Euphorbiaceaesp11 | 0.5431* | 73.06 | 12.53 | 0.2034 | 0.8947 | 60.49 | 0.5050 |
| Euphorbiaceaesp3 | 0.5431* | 73.06 | 12.53 | 0.2034 | 0.8947 | 60.49 | 0.5050 |
| Euphorbiaceaesp4 | 0.5431* | 73.06 | 12.53 | 0.2034 | 0.8947 | 60.49 | 0.5050 |
| Euphorbiaceaesp5 | 0.5431* | 73.06 | 12.53 | 0.2034 | 0.8947 | 60.49 | 0.5050 |
| phyllanthaceae | 0.6010 | 71.00 | 12.82 | 0.2030 | 0.9006 | 57.73 | 0.5245 |
| antidesma | 0.6284 | 70.03 | 12.95 | 0.2028 | 0.9033 | 56.43 | 0.5337 |
| Antidesmamaclurei | 0.5900* | 71.39 | 12.76 | 0.2031 | 0.8994 | 58.26 | 0.5208 |
| Antidesmamontanum | 0.5900* | 71.39 | 12.76 | 0.2031 | 0.8994 | 58.26 | 0.5208 |
| Antidesmasp1 | 0.6575* | 69.00 | 13.10 | 0.2026 | 0.9062 | 55.05 | 0.5435 |
| Antidesmasp2 | 0.6575* | 69.00 | 13.10 | 0.2026 | 0.9062 | 55.05 | 0.5435 |
| Antidesmahainanense | 0.6575* | 69.00 | 13.10 | 0.2026 | 0.9062 | 55.05 | 0.5435 |
| aporosa | 0.4232 | 77.31 | 11.93 | 0.2043 | 0.8827 | 66.19 | 0.4647 |
| Aporosadioica | 0.3700* | 79.19 | 11.67 | 0.2047 | 0.8774 | 68.72 | 0.4468 |
| baccaurea | 0.5097 | 74.24 | 12.36 | 0.2037 | 0.8914 | 62.07 | 0.4938 |
| Baccaurearamiflora | 0.5431* | 73.06 | 12.53 | 0.2034 | 0.8947 | 60.49 | 0.5050 |
| bischofia | 0.5772 | 71.85 | 12.70 | 0.2032 | 0.8982 | 58.87 | 0.5165 |
| Bischofiapolycarpa | 0.5699* | 72.11 | 12.66 | 0.2032 | 0.8974 | 59.21 | 0.5140 |
| breynia | 0.6215 | 70.28 | 12.92 | 0.2029 | 0.9026 | 56.76 | 0.5314 |
| Breyniafruticosa | 0.6315* | 69.92 | 12.97 | 0.2028 | 0.9036 | 56.28 | 0.5348 |

| | | | | | | | |
|---|---|---|---|---|---|---|---|
| Breyniarostrata | 0.6315* | 69.92 | 12.97 | 0.2028 | 0.9036 | 56.28 | 0.5348 |
| glochidion | 0.5486 | 72.86 | 12.56 | 0.2034 | 0.8953 | 60.23 | 0.5069 |
| Glochidioncoccineum | 0.5576* | 72.54 | 12.60 | 0.2033 | 0.8962 | 59.80 | 0.5099 |
| Glochidionhirsutum | 0.5576* | 72.54 | 12.60 | 0.2033 | 0.8962 | 59.80 | 0.5099 |
| Glochidionsphaerogynum | 0.5576* | 72.54 | 12.60 | 0.2033 | 0.8962 | 59.80 | 0.5099 |
| Glochidiontriandrum | 0.5576* | 72.54 | 12.60 | 0.2033 | 0.8962 | 59.80 | 0.5099 |
| Glochidionzeylanicum | 0.4800* | 75.29 | 12.22 | 0.2039 | 0.8884 | 63.49 | 0.4838 |
| phyllanthus | 0.6033 | 70.92 | 12.83 | 0.2030 | 0.9008 | 57.62 | 0.5253 |
| Phyllanthuspachyphyllus | 0.6126* | 70.59 | 12.87 | 0.2029 | 0.9017 | 57.18 | 0.5284 |
| bridelia | 0.6132 | 70.57 | 12.88 | 0.2029 | 0.9018 | 57.16 | 0.5286 |
| Brideliabalansae | 0.6053* | 70.85 | 12.84 | 0.2030 | 0.9010 | 57.53 | 0.5259 |
| cleistanthus | 0.6404 | 69.61 | 13.01 | 0.2027 | 0.9045 | 55.86 | 0.5378 |
| Cleistanthusconcinnus | 0.6518* | 69.20 | 13.07 | 0.2026 | 0.9057 | 55.32 | 0.5416 |
| leptopus | 0.6282 | 70.04 | 12.95 | 0.2028 | 0.9033 | 56.44 | 0.5337 |
| Leptopushainanensis | 0.6315* | 69.92 | 12.97 | 0.2028 | 0.9036 | 56.28 | 0.5348 |
| actephila | 0.6163 | 70.46 | 12.89 | 0.2029 | 0.9021 | 57.01 | 0.5297 |
| Actephilamerrilliana | 0.6315* | 69.92 | 12.97 | 0.2028 | 0.9036 | 56.28 | 0.5348 |
| ixonanthaceae | 0.6116 | 70.63 | 12.87 | 0.2029 | 0.9016 | 57.23 | 0.5281 |
| ixonanthes | 0.6191 | 70.36 | 12.91 | 0.2029 | 0.9024 | 56.88 | 0.5306 |
| Ixonanthesreticulata | 0.6265* | 70.10 | 12.94 | 0.2028 | 0.9031 | 56.52 | 0.5331 |
| salicaceae | 0.6564 | 69.04 | 13.09 | 0.2026 | 0.9061 | 55.10 | 0.5432 |
| casearia | 0.6394 | 69.64 | 13.01 | 0.2027 | 0.9044 | 55.91 | 0.5374 |
| Caseariamembranacea | 0.6500* | 69.27 | 13.06 | 0.2027 | 0.9055 | 55.40 | 0.5410 |
| Caseariavelutina | 0.6246* | 70.17 | 12.93 | 0.2028 | 0.9029 | 56.61 | 0.5325 |
| homalium | 0.6849 | 68.03 | 13.23 | 0.2024 | 0.9090 | 53.75 | 0.5527 |
| Homaliummollissimum | 0.6957* | 67.64 | 13.29 | 0.2023 | 0.9101 | 53.23 | 0.5564 |
| Homaliumpaniculiflorum | 0.6957* | 67.64 | 13.29 | 0.2023 | 0.9101 | 53.23 | 0.5564 |
| Homaliumphanerophlebium | 0.6400* | 69.62 | 13.01 | 0.2027 | 0.9045 | 55.88 | 0.5376 |
| Homaliumstenophyllum | 0.6957* | 67.64 | 13.29 | 0.2023 | 0.9101 | 53.23 | 0.5564 |
| scolopia | 0.8052 | 63.77 | 13.83 | 0.2015 | 0.9210 | 48.03 | 0.5932 |
| Scolopiasaeva | 0.8300* | 62.89 | 13.95 | 0.2014 | 0.9235 | 46.85 | 0.6016 |
| flacourtia | 0.7518 | 65.66 | 13.57 | 0.2019 | 0.9157 | 50.56 | 0.5753 |
| Flacourtiarukam | 0.7500* | 65.72 | 13.56 | 0.2019 | 0.9155 | 50.65 | 0.5747 |
| achariaceae | 0.6310 | 69.94 | 12.96 | 0.2028 | 0.9036 | 56.31 | 0.5346 |
| hydnocarpus | 0.6311 | 69.93 | 12.97 | 0.2028 | 0.9036 | 56.30 | 0.5347 |
| Hydnocarpushainanensis | 0.6313* | 69.93 | 12.97 | 0.2028 | 0.9036 | 56.29 | 0.5347 |
| hypericaceae | 0.6854 | 68.01 | 13.23 | 0.2024 | 0.9090 | 53.72 | 0.5529 |
| cratoxylum | 0.6915 | 67.80 | 13.27 | 0.2024 | 0.9096 | 53.43 | 0.5550 |
| Cratoxylumcochinchinense | 0.6700* | 68.56 | 13.16 | 0.2025 | 0.9075 | 54.45 | 0.5477 |
| Cratoxylumformosum | 0.7150* | 66.96 | 13.38 | 0.2022 | 0.9120 | 52.31 | 0.5629 |
| calophyllaceae | 0.6797 | 68.21 | 13.21 | 0.2024 | 0.9084 | 53.99 | 0.5510 |
| calophyllum | 0.6700 | 68.56 | 13.16 | 0.2025 | 0.9075 | 54.45 | 0.5477 |
| Calophyllummembranaceum | 0.6692* | 68.59 | 13.15 | 0.2025 | 0.9074 | 54.49 | 0.5475 |
| Calophyllumsp1 | 0.6692* | 68.59 | 13.15 | 0.2025 | 0.9074 | 54.49 | 0.5475 |
| clusiaceae | 0.6745 | 68.40 | 13.18 | 0.2025 | 0.9079 | 54.24 | 0.5493 |
| garcinia | 0.6831 | 68.09 | 13.22 | 0.2024 | 0.9088 | 53.83 | 0.5522 |
| Garciniamultiflora | 0.7416* | 66.02 | 13.51 | 0.2020 | 0.9147 | 51.05 | 0.5718 |
| Garciniaoblongifolia | 0.6268* | 70.09 | 12.94 | 0.2028 | 0.9031 | 56.51 | 0.5332 |
| Clusiaceaesp1 | 0.6692* | 68.59 | 13.15 | 0.2025 | 0.9074 | 54.49 | 0.5475 |
| ochnaceae | 0.7051 | 67.31 | 13.33 | 0.2023 | 0.9110 | 52.79 | 0.5595 |
| ochna | 0.7375 | 66.16 | 13.49 | 0.2020 | 0.9143 | 51.24 | 0.5705 |
| Ochnaintegerrima | 0.7440* | 65.93 | 13.53 | 0.2020 | 0.9149 | 50.93 | 0.5726 |
| Campylospermum | 0.7201 | 66.78 | 13.41 | 0.2022 | 0.9125 | 52.07 | 0.5646 |
| Campylospermumserratum | 0.7351* | 66.25 | 13.48 | 0.2020 | 0.9140 | 51.36 | 0.5696 |
| rhizophoraceae | 0.6974 | 67.59 | 13.29 | 0.2023 | 0.9102 | 53.15 | 0.5570 |
| carallia | 0.6737 | 68.42 | 13.18 | 0.2025 | 0.9079 | 54.28 | 0.5490 |
| Caralliabrachiata | 0.6659* | 68.70 | 13.14 | 0.2025 | 0.9071 | 54.65 | 0.5463 |
| erythroxylaceae | 0.7334 | 66.31 | 13.47 | 0.2021 | 0.9138 | 51.44 | 0.5691 |
| erythroxylum | 0.7615 | 65.31 | 13.61 | 0.2019 | 0.9167 | 50.10 | 0.5785 |
| Erythroxylumsinense | 0.7896* | 64.32 | 13.75 | 0.2017 | 0.9195 | 48.77 | 0.5880 |
| pandaceae | 0.6324 | 69.89 | 12.97 | 0.2028 | 0.9037 | 56.24 | 0.5351 |
| microdesmis | 0.6162 | 70.46 | 12.89 | 0.2029 | 0.9021 | 57.01 | 0.5296 |
| Microdesmiscaseariifolia | 0.6000* | 71.04 | 12.81 | 0.2030 | 0.9005 | 57.78 | 0.5242 |
| putranjivaceae | 0.6839 | 68.07 | 13.23 | 0.2024 | 0.9089 | 53.79 | 0.5524 |

| | | | | | | | |
|---|---|---|---|---|---|---|---|
| drypetes | 0.6978 | 67.57 | 13.30 | 0.2023 | 0.9103 | 53.13 | 0.5571 |
| Drypetescumingii | 0.6945* | 67.69 | 13.28 | 0.2023 | 0.9099 | 53.29 | 0.5560 |
| Drypeteshainanensis | 0.7183* | 66.84 | 13.40 | 0.2022 | 0.9123 | 52.16 | 0.5640 |
| Drypetesperreticulata | 0.6945* | 67.69 | 13.28 | 0.2023 | 0.9099 | 53.29 | 0.5560 |
| oxalidales | 0.6122 | 70.61 | 12.87 | 0.2029 | 0.9017 | 57.20 | 0.5283 |
| elaeocarpaceae | 0.5533 | 72.69 | 12.58 | 0.2034 | 0.8958 | 60.00 | 0.5085 |
| elaeocarpus | 0.4924 | 74.85 | 12.28 | 0.2038 | 0.8896 | 62.90 | 0.4880 |
| Elaeocarpusangustifolius | 0.4029* | 78.03 | 11.83 | 0.2044 | 0.8807 | 67.15 | 0.4578 |
| Elaeocarpusdubius | 0.5240* | 73.73 | 12.43 | 0.2036 | 0.8928 | 61.39 | 0.4986 |
| Elaeocarpusglabripetalus | 0.5032* | 74.47 | 12.33 | 0.2037 | 0.8907 | 62.38 | 0.4916 |
| Elaeocarpushowii | 0.5032* | 74.47 | 12.33 | 0.2037 | 0.8907 | 62.38 | 0.4916 |
| Elaeocarpusjaponicus | 0.5032* | 74.47 | 12.33 | 0.2037 | 0.8907 | 62.38 | 0.4916 |
| Elaeocarpuslimitaneus | 0.5032* | 74.47 | 12.33 | 0.2037 | 0.8907 | 62.38 | 0.4916 |
| Elaeocarpusnitentifolius | 0.5032* | 74.47 | 12.33 | 0.2037 | 0.8907 | 62.38 | 0.4916 |
| Elaeocarpuspoilanei | 0.5032* | 74.47 | 12.33 | 0.2037 | 0.8907 | 62.38 | 0.4916 |
| Elaeocarpussylvestris | 0.4750* | 75.47 | 12.19 | 0.2039 | 0.8879 | 63.72 | 0.4821 |
| sloanea | 0.5458 | 72.96 | 12.54 | 0.2034 | 0.8950 | 60.36 | 0.5060 |
| Sloaneaintegrifolia | 0.6095* | 70.70 | 12.86 | 0.2029 | 0.9014 | 57.33 | 0.5274 |
| Sloaneasinensis | 0.4785* | 75.35 | 12.21 | 0.2039 | 0.8883 | 63.56 | 0.4833 |
| connaraceae | 0.5881 | 71.46 | 12.75 | 0.2031 | 0.8993 | 58.35 | 0.5202 |
| ellipanthus | 0.5847 | 71.58 | 12.74 | 0.2031 | 0.8989 | 58.51 | 0.5190 |
| Ellipanthusglabrifolius | 0.5814* | 71.70 | 12.72 | 0.2031 | 0.8986 | 58.67 | 0.5179 |
| celastrales | 0.6422 | 69.54 | 13.02 | 0.2027 | 0.9047 | 55.78 | 0.5384 |
| celastraceae | 0.6503 | 69.26 | 13.06 | 0.2027 | 0.9055 | 55.39 | 0.5411 |
| Celastraceaesp2 | 0.6797* | 68.21 | 13.21 | 0.2024 | 0.9084 | 53.99 | 0.5510 |
| Celastraceaesp3 | 0.6797* | 68.21 | 13.21 | 0.2024 | 0.9084 | 53.99 | 0.5510 |
| salacia | 0.7478 | 65.80 | 13.54 | 0.2020 | 0.9153 | 50.75 | 0.5739 |
| Salaciachinensis | 0.7600* | 65.37 | 13.61 | 0.2019 | 0.9165 | 50.17 | 0.5780 |
| euonymus | 0.5874 | 71.48 | 12.75 | 0.2031 | 0.8992 | 58.38 | 0.5200 |
| Euonymusgibber | 0.5665* | 72.23 | 12.64 | 0.2033 | 0.8971 | 59.37 | 0.5129 |
| Euonymuslaxiflorus | 0.5665* | 72.23 | 12.64 | 0.2033 | 0.8971 | 59.37 | 0.5129 |
| Euonymusnitidus | 0.5665* | 72.23 | 12.64 | 0.2033 | 0.8971 | 59.37 | 0.5129 |
| myrtales | 0.6326 | 81.69 | 13.33 | 0.1991 | 0.9271 | 55.82 | 0.5182 |
| melastomataceae | 0.6854 | 79.81 | 13.59 | 0.1987 | 0.9325 | 53.30 | 0.5360 |
| blastus | 0.6081 | 82.55 | 13.21 | 0.1993 | 0.9247 | 56.98 | 0.5099 |
| Blastuscochinchinensis | 0.6094* | 82.51 | 13.22 | 0.1993 | 0.9248 | 56.92 | 0.5104 |
| melastoma | 0.4452 | 88.33 | 12.40 | 0.2005 | 0.9083 | 64.73 | 0.4551 |
| Melastomamalabathricum | 0.4400* | 88.51 | 12.37 | 0.2005 | 0.9078 | 64.97 | 0.4534 |
| Melastomapenicillatum | 0.4400* | 88.51 | 12.37 | 0.2005 | 0.9078 | 64.97 | 0.4534 |
| Melastomasanguineum | 0.4400* | 88.51 | 12.37 | 0.2005 | 0.9078 | 64.97 | 0.4534 |
| medinilla | 0.6142 | 82.34 | 13.24 | 0.1992 | 0.9253 | 56.69 | 0.5120 |
| Medinillaassamica | 0.6094* | 82.51 | 13.22 | 0.1993 | 0.9248 | 56.92 | 0.5104 |
| Allomorphia | 0.6474 | 81.16 | 13.40 | 0.1990 | 0.9286 | 55.11 | 0.5232 |
| Allomorphiabalansae | 0.6094* | 82.51 | 13.22 | 0.1993 | 0.9248 | 56.92 | 0.5104 |
| memecylon | 0.7510 | 77.49 | 13.92 | 0.1983 | 0.9390 | 50.19 | 0.5580 |
| Memecylonhainanense | 0.7728* | 76.72 | 14.03 | 0.1981 | 0.9412 | 49.15 | 0.5654 |
| Memecylonligustrifolium | 0.7728* | 76.72 | 14.03 | 0.1981 | 0.9412 | 49.15 | 0.5654 |
| Memecylonnigrescens | 0.7728* | 76.72 | 14.03 | 0.1981 | 0.9412 | 49.15 | 0.5654 |
| myrtaceae | 0.6869 | 79.76 | 13.60 | 0.1987 | 0.9326 | 53.23 | 0.5365 |
| rhodamnia | 0.8244 | 74.89 | 14.28 | 0.1977 | 0.9464 | 46.69 | 0.5828 |
| Rhodamniadumetorum | 0.8787* | 72.96 | 14.55 | 0.1973 | 0.9519 | 44.11 | 0.6010 |
| rhodomyrtus | 0.7069 | 79.05 | 13.70 | 0.1986 | 0.9346 | 52.28 | 0.5432 |
| Rhodomyrtustomentosa | 0.6797* | 80.02 | 13.56 | 0.1988 | 0.9319 | 53.58 | 0.5340 |
| decaspermum | 0.7256 | 78.39 | 13.79 | 0.1984 | 0.9365 | 51.39 | 0.5495 |
| Decaspermummontanum | 0.7170* | 78.69 | 13.75 | 0.1985 | 0.9356 | 51.80 | 0.5466 |
| syzygium | 0.6693 | 80.38 | 13.51 | 0.1988 | 0.9308 | 54.07 | 0.5305 |
| Syzygiumacuminatissimum | 0.6644* | 80.56 | 13.49 | 0.1989 | 0.9303 | 54.30 | 0.5289 |
| Syzygiumaraiocladum | 0.7360* | 78.02 | 13.84 | 0.1984 | 0.9375 | 50.90 | 0.5530 |
| Syzygiumbullockii | 0.6644* | 80.56 | 13.49 | 0.1989 | 0.9303 | 54.30 | 0.5289 |
| Syzygiumbuxifolium | 0.6644* | 80.56 | 13.49 | 0.1989 | 0.9303 | 54.30 | 0.5289 |
| Syzygiumchampionii | 0.6644* | 80.56 | 13.49 | 0.1989 | 0.9303 | 54.30 | 0.5289 |
| Syzygiumchunianum | 0.6644* | 80.56 | 13.49 | 0.1989 | 0.9303 | 54.30 | 0.5289 |
| Syzygiumclaviflorum | 0.6235* | 82.01 | 13.29 | 0.1992 | 0.9262 | 56.25 | 0.5151 |
| Syzygiumcumini | 0.6727* | 80.27 | 13.53 | 0.1988 | 0.9312 | 53.91 | 0.5317 |

| | | | | | | | |
|---|---|---|---|---|---|---|---|
| Syzygiumglobiflorum | 0.6644* | 80.56 | 13.49 | 0.1989 | 0.9303 | 54.30 | 0.5289 |
| Syzygiumhancei | 0.6644* | 80.56 | 13.49 | 0.1989 | 0.9303 | 54.30 | 0.5289 |
| Syzygiumjambos | 0.7000* | 79.30 | 13.67 | 0.1986 | 0.9339 | 52.61 | 0.5409 |
| Syzygiumjienfunicum | 0.6644* | 80.56 | 13.49 | 0.1989 | 0.9303 | 54.30 | 0.5289 |
| Syzygiumlevinei | 0.6644* | 80.56 | 13.49 | 0.1989 | 0.9303 | 54.30 | 0.5289 |
| Syzygiumodoratum | 0.6644* | 80.56 | 13.49 | 0.1989 | 0.9303 | 54.30 | 0.5289 |
| Syzygiumrehderianum | 0.6644* | 80.56 | 13.49 | 0.1989 | 0.9303 | 54.30 | 0.5289 |
| Syzygiumrysopodum | 0.6644* | 80.56 | 13.49 | 0.1989 | 0.9303 | 54.30 | 0.5289 |
| Syzygiumsp | 0.6792* | 80.04 | 13.56 | 0.1988 | 0.9318 | 53.60 | 0.5339 |
| Syzygiumsterrophyllum | 0.6644* | 80.56 | 13.49 | 0.1989 | 0.9303 | 54.30 | 0.5289 |
| Syzygiumtephrodes | 0.6644* | 80.56 | 13.49 | 0.1989 | 0.9303 | 54.30 | 0.5289 |
| Syzygiumtsoongii | 0.6644* | 80.56 | 13.49 | 0.1989 | 0.9303 | 54.30 | 0.5289 |
| sapindales | 0.5606 | 104.7 | 13.46 | 0.1850 | 0.9330 | 62.05 | 0.4844 |
| meliaceae | 0.5769 | 120.9 | 13.90 | 0.1806 | 0.9147 | 64.17 | 0.4304 |
| aphanamixis | 0.5849 | 120.7 | 13.94 | 0.1806 | 0.9155 | 63.79 | 0.4331 |
| Aphanamixispolystachya | 0.5765* | 121.0 | 13.90 | 0.1806 | 0.9147 | 64.19 | 0.4303 |
| aglaia | 0.5994 | 120.1 | 14.01 | 0.1805 | 0.9170 | 63.10 | 0.4380 |
| Aglaiaelaeagnoidea | 0.6300* | 119.1 | 14.16 | 0.1803 | 0.9201 | 61.64 | 0.4483 |
| Aglaiaspectabilis | 0.5750* | 121.0 | 13.89 | 0.1806 | 0.9145 | 64.26 | 0.4298 |
| dysoxylum | 0.5722 | 121.1 | 13.88 | 0.1807 | 0.9142 | 64.39 | 0.4288 |
| Dysoxylumgotadhora | 0.5974* | 120.2 | 14.00 | 0.1805 | 0.9168 | 63.19 | 0.4373 |
| Dysoxylummollissimum | 0.5189* | 123.0 | 13.61 | 0.1810 | 0.9089 | 66.93 | 0.4109 |
| walsura | 0.8326 | 111.9 | 15.17 | 0.1788 | 0.9404 | 52.01 | 0.5165 |
| Walsurapinnata | 0.8680* | 110.6 | 15.34 | 0.1785 | 0.9439 | 50.33 | 0.5284 |
| Walsurarobusta | 0.8680* | 110.6 | 15.34 | 0.1785 | 0.9439 | 50.33 | 0.5284 |
| melia | 0.4818 | 124.3 | 13.43 | 0.1813 | 0.9052 | 68.69 | 0.3984 |
| Meliaazedarach | 0.4378* | 125.9 | 13.21 | 0.1816 | 0.9008 | 70.78 | 0.3836 |
| Heynea | 0.5856 | 120.6 | 13.94 | 0.1806 | 0.9156 | 63.76 | 0.4334 |
| Heyneatrijuga | 0.5944* | 120.3 | 13.99 | 0.1805 | 0.9165 | 63.34 | 0.4363 |
| Meliaceaesp1 | 0.5944* | 120.3 | 13.99 | 0.1805 | 0.9165 | 63.34 | 0.4363 |
| simaroubaceae | 0.5340 | 122.5 | 13.69 | 0.1809 | 0.9104 | 66.21 | 0.4160 |
| brucea | 0.4532 | 125.3 | 13.28 | 0.1815 | 0.9023 | 70.05 | 0.3888 |
| Bruceamollis | 0.4330* | 126.0 | 13.18 | 0.1817 | 0.9003 | 71.01 | 0.3820 |
| rutaceae | 0.5497 | 124.7 | 13.97 | 0.1811 | 0.9074 | 65.97 | 0.3884 |
| murraya | 0.6589 | 113.3 | 15.39 | 0.1850 | 0.9071 | 63.48 | 0.4175 |
| Murrayaalata | 0.7537* | 109.9 | 15.86 | 0.1844 | 0.9166 | 58.97 | 0.4494 |
| glycosmis | 0.4703 | 120.0 | 14.45 | 0.1864 | 0.8882 | 72.44 | 0.3540 |
| Glycosmissp1 | 0.4390* | 121.1 | 14.30 | 0.1866 | 0.8850 | 73.93 | 0.3435 |
| Glycosmiscochinchinensis | 0.4390* | 121.1 | 14.30 | 0.1866 | 0.8850 | 73.93 | 0.3435 |
| Glycosmiscraibii | 0.4390* | 121.1 | 14.30 | 0.1866 | 0.8850 | 73.93 | 0.3435 |
| clausena | 0.5231 | 118.1 | 14.71 | 0.1860 | 0.8935 | 69.93 | 0.3718 |
| Clausenaexcavata | 0.4820* | 119.5 | 14.51 | 0.1863 | 0.8893 | 71.88 | 0.3580 |
| micromelum | 0.6121 | 114.9 | 15.16 | 0.1854 | 0.9024 | 65.70 | 0.4017 |
| Micromelumfalcatum | 0.6600* | 113.2 | 15.39 | 0.1850 | 0.9072 | 63.42 | 0.4179 |
| tetradium | 0.2887 | 164.5 | 12.18 | 0.2539 | 0.6559 | 136.7 | 0.3361 |
| Tetradiumglabrifolium | 0.2320* | 177.9* | 11.34* | 0.2697* | 0.5995* | 152.6* | 0.3298* |
| acronychia | 0.4687 | 93.24 | 20.30 | 0.2085 | 0.9746 | 72.70 | 0.2612 |
| Acronychiapedunculata | 0.4587* | 90.96* | 21.96* | 0.2188* | 1.037* | 69.46* | 0.2327* |
| melicope | 0.4921 | 96.82 | 15.66 | 0.1769 | 0.7783 | 83.96 | 0.3358 |
| Melicopechunii | 0.4953* | 95.85* | 14.35* | 0.1659* | 0.7075* | 88.74* | 0.3534* |
| zanthoxylum | 0.5667 | 97.21 | 16.41 | 0.1782 | 0.9391 | 57.45 | 0.3708 |
| Zanthoxylumavicennae | 0.6177* | 83.67* | 17.27* | 0.1656* | 0.9967* | 41.92* | 0.3801* |
| Maclurodendron | 0.5499 | 131.4 | 13.75 | 0.1791 | 0.9084 | 65.12 | 0.3594 |
| Maclurodendronoligophlebium | 0.5500* | 138.0* | 13.52* | 0.1770* | 0.9094* | 64.27* | 0.3304* |
| sapindaceae | 0.5646 | 119.5 | 13.33 | 0.1828 | 0.9008 | 64.72 | 0.4702 |
| mischocarpus | 0.7465 | 121.5 | 11.56 | 0.2034 | 0.7638 | 60.33 | 0.6311 |
| Mischocarpushainanensis | 0.7307* | 122.1 | 11.48 | 0.2035 | 0.7622 | 61.08 | 0.6258 |
| Mischocarpuspentapetalus | 0.7307* | 122.1 | 11.48 | 0.2035 | 0.7622 | 61.08 | 0.6258 |
| Mischocarpussundaicus | 0.7800* | 120.3 | 11.73 | 0.2031 | 0.7672 | 58.74 | 0.6424 |
| nephelium | 0.7724 | 125.3 | 10.20 | 0.2153 | 0.6802 | 61.46 | 0.6952 |
| Nepheliumtopengii | 0.7782* | 126.0* | 9.933* | 0.2177* | 0.6635* | 61.66* | 0.7082* |
| dimocarpus | 0.7344 | 124.8 | 10.61 | 0.2107 | 0.7109 | 62.32 | 0.6603 |
| Dimocarpuslongan | 0.7000* | 126.0 | 10.44 | 0.2110 | 0.7074 | 63.96 | 0.6487 |

| | | | | | | | |
|---|---|---|---|---|---|---|---|
| litchi | 0.8115 | 122.0 | 10.99 | 0.2102 | 0.7186 | 58.66 | 0.6862 |
| Litchichinensis | 0.8541* | 120.5 | 11.20 | 0.2099 | 0.7229 | 56.63 | 0.7006 |
| lepisanthes | 0.6471 | 126.0 | 10.77 | 0.2065 | 0.7366 | 65.53 | 0.6087 |
| Lepisanthesrubiginosa | 0.6300* | 126.6 | 10.68 | 0.2066 | 0.7349 | 66.34 | 0.6030 |
| amesiodendron | 0.8166 | 116.2 | 12.80 | 0.1956 | 0.8226 | 55.58 | 0.6215 |
| Amesiodendronchinense | 0.8345* | 115.6 | 12.89 | 0.1955 | 0.8244 | 54.73 | 0.6275 |
| paranephelium | 0.8037 | 116.6 | 12.74 | 0.1957 | 0.8213 | 56.19 | 0.6171 |
| Paranepheliumhainanense | 0.8267* | 115.8 | 12.85 | 0.1955 | 0.8236 | 55.10 | 0.6249 |
| acer | 0.4879 | 124.1 | 12.36 | 0.1882 | 0.8586 | 69.31 | 0.4666 |
| Acerfabri | 0.5145* | 123.1 | 12.49 | 0.1881 | 0.8613 | 68.05 | 0.4755 |
| Acerlaurinum | 0.4300* | 126.1 | 12.07 | 0.1887 | 0.8528 | 72.07 | 0.4471 |
| anacardiaceae | 0.5240 | 116.3 | 13.87 | 0.1639 | 0.9671 | 62.76 | 0.5223 |
| toxicodendron | 0.5620 | 92.36 | 13.53 | 0.1515 | 0.7992 | 43.99 | 0.6814 |
| Toxicodendronvernicifluum | 0.5662* | 89.70* | 13.49* | 0.1501* | 0.7805* | 41.90* | 0.6991* |
| choerospondias | 0.4932 | 117.4 | 13.72 | 0.1641 | 0.9640 | 64.23 | 0.5120 |
| Choerospondiasaxillaris | 0.4870* | 117.6 | 13.69 | 0.1642 | 0.9634 | 64.52 | 0.5099 |
| burseraceae | 0.5159 | 123.0 | 14.16 | 0.1603 | 1.030 | 66.64 | 0.5065 |
| canarium | 0.4860 | 135.6 | 14.82 | 0.1455 | 1.159 | 72.89 | 0.5059 |
| Canariumalbum | 0.4170* | 145.5* | 14.37* | 0.1437* | 0.6627* | 103.8* | 0.3217* |
| Canariumpimela | 0.5450* | 129.9* | 15.49* | 0.1423* | 1.699* | 44.09* | 0.6899* |
| malvales | 0.5699 | 99.24 | 13.38 | 0.1886 | 0.9307 | 60.91 | 0.4899 |
| thymelaeaceae | 0.5296 | 100.7 | 13.18 | 0.1889 | 0.9266 | 62.82 | 0.4764 |
| aquilaria | 0.4134 | 104.8 | 12.60 | 0.1897 | 0.9150 | 68.35 | 0.4373 |
| Aquilariasinensis | 0.3660* | 106.5 | 12.37 | 0.1901 | 0.9102 | 70.60 | 0.4213 |
| wikstroemia | 0.5262 | 100.8 | 13.16 | 0.1889 | 0.9263 | 62.99 | 0.4752 |
| Wikstroemiahainanensis | 0.5294* | 100.7 | 13.18 | 0.1889 | 0.9266 | 62.83 | 0.4763 |
| Wikstroemiaindica | 0.5294* | 100.7 | 13.18 | 0.1889 | 0.9266 | 62.83 | 0.4763 |
| Wikstroemianutans | 0.5294* | 100.7 | 13.18 | 0.1889 | 0.9266 | 62.83 | 0.4763 |
| Wikstroemiapachyrachis | 0.5294* | 100.7 | 13.18 | 0.1889 | 0.9266 | 62.83 | 0.4763 |
| malvaceae | 0.5740 | 99.09 | 13.40 | 0.1886 | 0.9311 | 60.71 | 0.4913 |
| reevesia | 0.4716 | 102.7 | 12.89 | 0.1893 | 0.9208 | 65.58 | 0.4568 |
| Reevesialancifolia | 0.5075* | 101.4 | 13.07 | 0.1891 | 0.9244 | 63.87 | 0.4689 |
| Reevesiathyrsoidea | 0.4350* | 104.0 | 12.71 | 0.1896 | 0.9171 | 67.32 | 0.4445 |
| sterculia | 0.4567 | 103.2 | 12.82 | 0.1894 | 0.9193 | 66.29 | 0.4518 |
| Sterculiahainanensis | 0.4048* | 105.1 | 12.56 | 0.1898 | 0.9141 | 68.75 | 0.4344 |
| Sterculialanceolata | 0.5000* | 101.7 | 13.03 | 0.1891 | 0.9237 | 64.23 | 0.4664 |
| pterospermum | 0.4925 | 102.0 | 13.00 | 0.1892 | 0.9229 | 64.59 | 0.4639 |
| Pterospermumheterophyllum | 0.4460* | 103.6 | 12.77 | 0.1895 | 0.9182 | 66.80 | 0.4482 |
| Pterospermumlanceifolium | 0.5146* | 101.2 | 13.11 | 0.1890 | 0.9251 | 63.54 | 0.4713 |
| Pterospermumxiaoye | 0.5146* | 101.2 | 13.11 | 0.1890 | 0.9251 | 63.54 | 0.4713 |
| microcos | 0.4888 | 102.1 | 12.98 | 0.1892 | 0.9225 | 64.76 | 0.4626 |
| Micrococschungii | 0.4819* | 102.4 | 12.94 | 0.1892 | 0.9218 | 65.09 | 0.4603 |
| dipterocarpaceae | 0.7336 | 93.43 | 14.19 | 0.1874 | 0.9471 | 53.12 | 0.5450 |
| vatica | 0.7577 | 92.58 | 14.31 | 0.1873 | 0.9495 | 51.98 | 0.5531 |
| Vaticamangachapoi | 0.7500* | 92.85 | 14.28 | 0.1873 | 0.9488 | 52.34 | 0.5505 |
| hopea | 0.8353 | 89.83 | 14.70 | 0.1867 | 0.9573 | 48.29 | 0.5792 |
| Hopeahainanensis | 0.8900* | 87.89 | 14.97 | 0.1863 | 0.9628 | 45.69 | 0.5976 |
| brassicales | 0.5866 | 98.65 | 13.46 | 0.1885 | 0.9324 | 60.11 | 0.4955 |
| capparaceae | 0.6725 | 95.60 | 13.89 | 0.1879 | 0.9410 | 56.03 | 0.5244 |
| Capparaceaesp1 | 0.6832* | 95.22 | 13.94 | 0.1878 | 0.9421 | 55.52 | 0.5281 |
| crossosomatales | 0.5384 | 90.13 | 12.98 | 0.1961 | 0.9210 | 61.00 | 0.4841 |
| staphyleaceae | 0.4807 | 92.18 | 12.70 | 0.1966 | 0.9152 | 63.74 | 0.4647 |
| turpinia | 0.4229 | 94.23 | 12.41 | 0.1970 | 0.9094 | 66.49 | 0.4452 |
| Turpiniamontana | 0.3940* | 95.25 | 12.27 | 0.1972 | 0.9065 | 67.86 | 0.4355 |
| saxifragales | 0.6081 | 73.92 | 13.08 | 0.2102 | 0.9350 | 53.75 | 0.5026 |
| hamamelidaceae | 0.5996 | 74.22 | 13.04 | 0.2103 | 0.9342 | 54.15 | 0.4997 |
| eustigma | 0.6330 | 73.04 | 13.21 | 0.2100 | 0.9375 | 52.56 | 0.5110 |
| Eustigmaoblongifolium | 0.6372* | 72.89 | 13.23 | 0.2100 | 0.9379 | 52.36 | 0.5124 |
| exbucklandia | 0.5715 | 75.22 | 12.90 | 0.2105 | 0.9313 | 55.49 | 0.4902 |
| Exbucklandiatonkinensis | 0.5480* | 76.05 | 12.79 | 0.2106 | 0.9290 | 56.61 | 0.4823 |
| chunia | 0.6161 | 73.64 | 13.12 | 0.2102 | 0.9358 | 53.37 | 0.5053 |
| Chuniabucklandioides | 0.6372* | 72.89 | 13.23 | 0.2100 | 0.9379 | 52.36 | 0.5124 |
| daphniphyllaceae | 0.5520 | 75.91 | 12.81 | 0.2106 | 0.9294 | 56.41 | 0.4837 |
| daphniphyllum | 0.5292 | 76.72 | 12.69 | 0.2108 | 0.9271 | 57.50 | 0.4760 |

| | | | | | | | |
|---|---|---|---|---|---|---|---|
| Daphniphyllumcalycinum | 0.5063* | 77.53 | 12.58 | 0.2109 | 0.9248 | 58.59 | 0.4683 |
| altingiaceae | 0.6460 | 72.58 | 13.27 | 0.2099 | 0.9388 | 51.95 | 0.5153 |
| altingia | 0.6731 | 71.62 | 13.41 | 0.2097 | 0.9415 | 50.66 | 0.5244 |
| Altingiaobovata | 0.7003* | 70.65 | 13.54 | 0.2096 | 0.9443 | 49.37 | 0.5336 |
| iteaceae | 0.5875 | 74.65 | 12.98 | 0.2104 | 0.9330 | 54.73 | 0.4956 |
| itea | 0.5834 | 74.80 | 12.96 | 0.2104 | 0.9325 | 54.92 | 0.4943 |
| Iteamacrophylla | 0.5814* | 74.87 | 12.95 | 0.2104 | 0.9323 | 55.02 | 0.4936 |
| Iteaxiaoye | 0.5814* | 74.87 | 12.95 | 0.2104 | 0.9323 | 55.02 | 0.4936 |
| saxifragaceae | 0.5848 | 74.75 | 12.97 | 0.2104 | 0.9327 | 54.86 | 0.4947 |
| Saxifragaceaesp1 | 0.5814* | 74.87 | 12.95 | 0.2104 | 0.9323 | 55.02 | 0.4936 |
| dilleniales | 0.6104 | 72.08 | 13.09 | 0.2138 | 0.9420 | 52.38 | 0.4985 |
| dilleniaceae | 0.6094 | 72.11 | 13.09 | 0.2138 | 0.9419 | 52.42 | 0.4981 |
| dillenia | 0.6063 | 72.22 | 13.07 | 0.2139 | 0.9416 | 52.57 | 0.4971 |
| Dilleniapentagyna | 0.5953* | 72.61 | 13.02 | 0.2139 | 0.9405 | 53.09 | 0.4934 |
| Dilleniaturbinata | 0.6162* | 71.87 | 13.12 | 0.2138 | 0.9426 | 52.10 | 0.5004 |
| gentianales | 0.6145 | 66.18 | 13.19 | 0.2252 | 0.9114 | 49.82 | 0.5056 |
| rubiaceae | 0.6438 | 65.14 | 13.33 | 0.2250 | 0.9143 | 48.42 | 0.5154 |
| Benkara | 0.6406 | 65.26 | 13.32 | 0.2250 | 0.9140 | 48.58 | 0.5143 |
| Benkarahainanensis | 0.6374* | 65.37 | 13.30 | 0.2250 | 0.9137 | 48.73 | 0.5133 |
| antirhea | 0.6404 | 65.26 | 13.32 | 0.2250 | 0.9140 | 48.58 | 0.5143 |
| Antirheachinensis | 0.6374* | 65.37 | 13.30 | 0.2250 | 0.9137 | 48.73 | 0.5133 |
| pertusadina | 0.6847 | 63.69 | 13.54 | 0.2247 | 0.9185 | 46.48 | 0.5292 |
| Pertusadinametcalfii | 0.6800* | 63.86 | 13.51 | 0.2247 | 0.9180 | 46.70 | 0.5276 |
| adina | 0.7215 | 62.39 | 13.72 | 0.2244 | 0.9221 | 44.73 | 0.5416 |
| Adinarubella | 0.7351* | 61.91 | 13.79 | 0.2243 | 0.9235 | 44.08 | 0.5461 |
| nauclea | 0.6423 | 65.20 | 13.33 | 0.2250 | 0.9142 | 48.50 | 0.5149 |
| naucleaofficinalis | 0.6374* | 65.37 | 13.30 | 0.2250 | 0.9137 | 48.73 | 0.5133 |
| diplospora | 0.7015 | 63.10 | 13.62 | 0.2246 | 0.9201 | 45.68 | 0.5348 |
| Diplosporadubia | 0.7000* | 63.15 | 13.61 | 0.2246 | 0.9200 | 45.75 | 0.5343 |
| catunaregam | 0.6913 | 63.46 | 13.57 | 0.2247 | 0.9191 | 46.16 | 0.5314 |
| Catunaregamspinosa | 0.6880* | 63.58 | 13.55 | 0.2247 | 0.9188 | 46.32 | 0.5303 |
| gardenia | 0.6722 | 64.14 | 13.48 | 0.2248 | 0.9172 | 47.07 | 0.5250 |
| Gardeniahainanensis | 0.6681* | 64.28 | 13.46 | 0.2248 | 0.9168 | 47.27 | 0.5236 |
| Gardeniasootepensis | 0.6681* | 64.28 | 13.46 | 0.2248 | 0.9168 | 47.27 | 0.5236 |
| aidia | 0.7404 | 61.72 | 13.81 | 0.2243 | 0.9240 | 43.83 | 0.5479 |
| Aidiacanthioides | 0.7527* | 61.28 | 13.87 | 0.2242 | 0.9253 | 43.25 | 0.5521 |
| Aidiapycnantha | 0.7527* | 61.28 | 13.87 | 0.2242 | 0.9253 | 43.25 | 0.5521 |
| pavetta | 0.6447 | 65.11 | 13.34 | 0.2250 | 0.9144 | 48.38 | 0.5157 |
| Pavettaarenosa | 0.6374* | 65.37 | 13.30 | 0.2250 | 0.9137 | 48.73 | 0.5133 |
| Pavettahongkongensis | 0.6374* | 65.37 | 13.30 | 0.2250 | 0.9137 | 48.73 | 0.5133 |
| tarenna | 0.6783 | 63.92 | 13.51 | 0.2247 | 0.9178 | 46.78 | 0.5270 |
| Tarennaattenuata | 0.6798* | 63.87 | 13.51 | 0.2247 | 0.9180 | 46.71 | 0.5275 |
| Tarennalancilimba | 0.6798* | 63.87 | 13.51 | 0.2247 | 0.9180 | 46.71 | 0.5275 |
| Tarennatsangii | 0.6798* | 63.87 | 13.51 | 0.2247 | 0.9180 | 46.71 | 0.5275 |
| ixora | 0.7755 | 60.48 | 13.99 | 0.2241 | 0.9276 | 42.16 | 0.5597 |
| Ixoranienkui | 0.7929* | 59.86 | 14.07 | 0.2239 | 0.9293 | 41.33 | 0.5656 |
| canthium | 0.6477 | 65.01 | 13.35 | 0.2250 | 0.9147 | 48.24 | 0.5167 |
| Canthiumhorridum | 0.6367* | 65.40 | 13.30 | 0.2250 | 0.9136 | 48.76 | 0.5130 |
| Canthiumsimile | 0.6367* | 65.40 | 13.30 | 0.2250 | 0.9136 | 48.76 | 0.5130 |
| psydrax | 0.7505 | 61.36 | 13.86 | 0.2242 | 0.9251 | 43.35 | 0.5513 |
| Psydraxdicocca | 0.7627* | 60.93 | 13.93 | 0.2241 | 0.9263 | 42.77 | 0.5555 |
| Tarennoidea | 0.6406 | 65.26 | 13.32 | 0.2250 | 0.9140 | 48.58 | 0.5143 |
| Tarennoideawallichii | 0.6374* | 65.37 | 13.30 | 0.2250 | 0.9137 | 48.73 | 0.5133 |
| hedyotis | 0.6367 | 65.40 | 13.30 | 0.2250 | 0.9136 | 48.76 | 0.5130 |
| Hedyotiscathayana | 0.6374* | 65.37 | 13.30 | 0.2250 | 0.9137 | 48.73 | 0.5133 |
| saprosma | 0.6372 | 65.38 | 13.30 | 0.2250 | 0.9137 | 48.74 | 0.5132 |
| Saprosmacrassipes | 0.6374* | 65.37 | 13.30 | 0.2250 | 0.9137 | 48.73 | 0.5133 |
| Saprosmahainanensis | 0.6374* | 65.37 | 13.30 | 0.2250 | 0.9137 | 48.73 | 0.5133 |
| Saprosmamerrillii | 0.6374* | 65.37 | 13.30 | 0.2250 | 0.9137 | 48.73 | 0.5133 |
| Saprosmayueyan | 0.6374* | 65.37 | 13.30 | 0.2250 | 0.9137 | 48.73 | 0.5133 |
| chassalia | 0.6309 | 65.60 | 13.27 | 0.2251 | 0.9130 | 49.04 | 0.5111 |
| Chassaliacurviflora | 0.6374* | 65.37 | 13.30 | 0.2250 | 0.9137 | 48.73 | 0.5133 |
| psychotria | 0.5730 | 67.65 | 12.98 | 0.2255 | 0.9072 | 51.79 | 0.4916 |
| Psychotriaasiatica | 0.5636* | 67.99 | 12.94 | 0.2256 | 0.9063 | 52.23 | 0.4884 |

| | | | | | | | |
|---|---|---|---|---|---|---|---|
| Psychotriastraminea | 0.5636* | 67.99 | 12.94 | 0.2256 | 0.9063 | 52.23 | 0.4884 |
| prismatomeris | 0.6304 | 65.62 | 13.27 | 0.2251 | 0.9130 | 49.06 | 0.5109 |
| Prismatomeristetrandra | 0.6374* | 65.37 | 13.30 | 0.2250 | 0.9137 | 48.73 | 0.5133 |
| lasianthus | 0.6373 | 65.37 | 13.30 | 0.2250 | 0.9137 | 48.73 | 0.5132 |
| Lasianthuschevalieri | 0.6374* | 65.37 | 13.30 | 0.2250 | 0.9137 | 48.73 | 0.5133 |
| Lasianthuscurtisii | 0.6374* | 65.37 | 13.30 | 0.2250 | 0.9137 | 48.73 | 0.5133 |
| Lasianthushirsutus | 0.6374* | 65.37 | 13.30 | 0.2250 | 0.9137 | 48.73 | 0.5133 |
| Lasianthusjaponicus | 0.6374* | 65.37 | 13.30 | 0.2250 | 0.9137 | 48.73 | 0.5133 |
| Lasianthuslancifolius | 0.6374* | 65.37 | 13.30 | 0.2250 | 0.9137 | 48.73 | 0.5133 |
| Lasianthusrhinocerotis | 0.6374* | 65.37 | 13.30 | 0.2250 | 0.9137 | 48.73 | 0.5133 |
| Lasianthustrichophlebus | 0.6374* | 65.37 | 13.30 | 0.2250 | 0.9137 | 48.73 | 0.5133 |
| Caelospermum | 0.6406 | 65.26 | 13.32 | 0.2250 | 0.9140 | 48.58 | 0.5143 |
| Caelospermumtruncatum | 0.6374* | 65.37 | 13.30 | 0.2250 | 0.9137 | 48.73 | 0.5133 |
| Wendlandia | 0.6764 | 63.99 | 13.50 | 0.2248 | 0.9176 | 46.87 | 0.5264 |
| Wendlandiamerrilliana | 0.7337* | 61.96 | 13.78 | 0.2244 | 0.9234 | 44.15 | 0.5457 |
| Wendlandiauvariifolia | 0.6518* | 64.86 | 13.37 | 0.2249 | 0.9152 | 48.04 | 0.5181 |
| apocynaceae | 0.6109 | 66.31 | 13.17 | 0.2252 | 0.9110 | 49.99 | 0.5043 |
| rauvolfia | 0.5001 | 70.24 | 12.62 | 0.2260 | 0.8999 | 55.25 | 0.4671 |
| Rauvolfiaverticillata | 0.4863* | 70.72 | 12.55 | 0.2261 | 0.8985 | 55.91 | 0.4624 |
| kopsia | 0.5740 | 67.62 | 12.99 | 0.2255 | 0.9073 | 51.74 | 0.4919 |
| Kopsiaarborea | 0.5929* | 66.95 | 13.08 | 0.2254 | 0.9092 | 50.84 | 0.4983 |
| wrightia | 0.3411 | 75.87 | 11.83 | 0.2272 | 0.8840 | 62.81 | 0.4136 |
| Wrightialaevis | 0.3120* | 76.91 | 11.69 | 0.2274 | 0.8810 | 64.20 | 0.4038 |
| tabernaemontana | 0.5587 | 68.16 | 12.91 | 0.2256 | 0.9058 | 52.47 | 0.4868 |
| Tabernaemontanabovina | 0.5656* | 67.92 | 12.95 | 0.2256 | 0.9065 | 52.14 | 0.4891 |
| Tabernaemontanabufalina | 0.5656* | 67.92 | 12.95 | 0.2256 | 0.9065 | 52.14 | 0.4891 |
| alstonia | 0.5095 | 69.90 | 12.67 | 0.2260 | 0.9009 | 54.80 | 0.4702 |
| Alstoniarostrata | 0.4640* | 71.52 | 12.44 | 0.2263 | 0.8963 | 56.97 | 0.4549 |
| Hunteria | 0.6654 | 64.38 | 13.44 | 0.2248 | 0.9165 | 47.39 | 0.5227 |
| Hunteriazeylanica | 0.7200* | 62.44 | 13.71 | 0.2244 | 0.9220 | 44.80 | 0.5411 |
| lamiales | 0.6055 | 66.50 | 13.14 | 0.2253 | 0.9105 | 50.24 | 0.5025 |
| bignoniaceae | 0.5817 | 67.34 | 13.03 | 0.2254 | 0.9081 | 51.37 | 0.4945 |
| radermachera | 0.5246 | 69.37 | 12.74 | 0.2259 | 0.9024 | 54.09 | 0.4753 |
| Radermacherasinica | 0.6255* | 65.79 | 13.24 | 0.2251 | 0.9125 | 49.29 | 0.5093 |
| Radermacherafrondosa | 0.4593* | 71.69 | 12.42 | 0.2263 | 0.8958 | 57.20 | 0.4533 |
| Radermacherahainanensis | 0.4593* | 71.69 | 12.42 | 0.2263 | 0.8958 | 57.20 | 0.4533 |
| markhamia | 0.6451 | 65.10 | 13.34 | 0.2250 | 0.9145 | 48.36 | 0.5159 |
| Markhamiastipulata | 0.6755* | 64.02 | 13.49 | 0.2248 | 0.9175 | 46.91 | 0.5261 |
| lamiaceae | 0.5300 | 69.18 | 12.77 | 0.2258 | 0.9029 | 53.83 | 0.4771 |
| gmelina | 0.5624 | 68.03 | 12.93 | 0.2256 | 0.9062 | 52.29 | 0.4880 |
| Gmelinahainanensis | 0.5947* | 66.88 | 13.09 | 0.2253 | 0.9094 | 50.76 | 0.4989 |
| Tsoongia | 0.5533 | 68.35 | 12.88 | 0.2256 | 0.9053 | 52.72 | 0.4850 |
| Tsoongiaaxillariflora | 0.5766* | 67.53 | 13.00 | 0.2255 | 0.9076 | 51.62 | 0.4928 |
| callicarpa | 0.4100 | 73.43 | 12.17 | 0.2267 | 0.8909 | 59.54 | 0.4367 |
| Callicarpabrevipes | 0.3500* | 75.56 | 11.88 | 0.2271 | 0.8849 | 62.39 | 0.4165 |
| Callicarpasp2 | 0.3500* | 75.56 | 11.88 | 0.2271 | 0.8849 | 62.39 | 0.4165 |
| vitex | 0.6354 | 65.44 | 13.29 | 0.2251 | 0.9135 | 48.82 | 0.5126 |
| Vitexpierreana | 0.8545* | 57.67 | 14.38 | 0.2235 | 0.9355 | 38.40 | 0.5863 |
| Vitexquinata | 0.4513* | 71.97 | 12.38 | 0.2264 | 0.8950 | 57.57 | 0.4506 |
| clerodendrum | 0.5702 | 67.75 | 12.97 | 0.2255 | 0.9070 | 51.92 | 0.4906 |
| Clerodendrumhainanense | 0.5735* | 67.64 | 12.99 | 0.2255 | 0.9073 | 51.76 | 0.4918 |
| Clerodendrumkwangtungense | 0.5735* | 67.64 | 12.99 | 0.2255 | 0.9073 | 51.76 | 0.4918 |
| verbenaceae | 0.6258 | 65.78 | 13.24 | 0.2251 | 0.9125 | 49.28 | 0.5094 |
| Verbenaceaesp | 0.6426* | 65.19 | 13.33 | 0.2250 | 0.9142 | 48.48 | 0.5150 |
| oleaceae | 0.6711 | 64.18 | 13.47 | 0.2248 | 0.9171 | 47.13 | 0.5246 |
| olea | 0.6935 | 63.38 | 13.58 | 0.2246 | 0.9193 | 46.06 | 0.5322 |
| Oleabrachiata | 0.6300* | 65.63 | 13.27 | 0.2251 | 0.9130 | 49.08 | 0.5108 |
| Oleaneriifolia | 0.7431* | 61.62 | 13.83 | 0.2243 | 0.9243 | 43.70 | 0.5488 |
| Oleaparvilimba | 0.7431* | 61.62 | 13.83 | 0.2243 | 0.9243 | 43.70 | 0.5488 |
| Oleatsoongii | 0.6300* | 65.63 | 13.27 | 0.2251 | 0.9130 | 49.08 | 0.5108 |
| osmanthus | 0.8323 | 58.46 | 14.27 | 0.2236 | 0.9333 | 39.46 | 0.5788 |
| Osmanthusdidymopetalus | 0.8415* | 58.13 | 14.32 | 0.2236 | 0.9342 | 39.02 | 0.5820 |
| Osmanthushainanensis | 0.8415* | 58.13 | 14.32 | 0.2236 | 0.9342 | 39.02 | 0.5820 |
| Osmanthusmarginatus | 0.8415* | 58.13 | 14.32 | 0.2236 | 0.9342 | 39.02 | 0.5820 |

| | | | | | | | |
|---|---|---|---|---|---|---|---|
| Osmanthusmatsumuranus | 0.8415* | 58.13 | 14.32 | 0.2236 | 0.9342 | 39.02 | 0.5820 |
| chionanthus | 0.7094 | 62.82 | 13.66 | 0.2245 | 0.9209 | 45.30 | 0.5375 |
| Chionanthusbrachythyrsus | 0.6805* | 63.84 | 13.52 | 0.2247 | 0.9180 | 46.68 | 0.5278 |
| Chionanthusramiflorus | 0.7530* | 61.27 | 13.88 | 0.2242 | 0.9253 | 43.23 | 0.5522 |
| boraginales | 0.5693 | 67.78 | 12.96 | 0.2255 | 0.9069 | 51.96 | 0.4904 |
| boraginaceae | 0.5336 | 69.05 | 12.79 | 0.2258 | 0.9033 | 53.66 | 0.4783 |
| ehretia | 0.5140 | 69.74 | 12.69 | 0.2259 | 0.9013 | 54.59 | 0.4718 |
| Ehretialongiflora | 0.5195* | 69.55 | 12.72 | 0.2259 | 0.9019 | 54.33 | 0.4736 |
| cordia | 0.4745 | 71.14 | 12.49 | 0.2262 | 0.8974 | 56.47 | 0.4584 |
| Cordiadichotoma | 0.4512* | 71.97 | 12.38 | 0.2264 | 0.8950 | 57.58 | 0.4506 |
| garryales | 0.5291 | 69.21 | 12.76 | 0.2258 | 0.9028 | 53.87 | 0.4768 |
| icacinaceae | 0.5004 | 70.23 | 12.62 | 0.2260 | 0.9000 | 55.24 | 0.4672 |
| apodytes | 0.5735 | 67.64 | 12.99 | 0.2255 | 0.9073 | 51.77 | 0.4918 |
| Apodytesdimidiata | 0.6100* | 66.34 | 13.17 | 0.2252 | 0.9110 | 50.03 | 0.5040 |
| Platea | 0.4351 | 72.54 | 12.30 | 0.2265 | 0.8934 | 58.34 | 0.4452 |
| Platealatifolia | 0.3400* | 75.91 | 11.83 | 0.2272 | 0.8839 | 62.87 | 0.4132 |
| Plateaparvifolia | 0.4650* | 71.48 | 12.45 | 0.2263 | 0.8964 | 56.92 | 0.4552 |
| apiales | 0.5389 | 68.86 | 12.81 | 0.2257 | 0.9038 | 53.41 | 0.4801 |
| pittosporaceae | 0.5381 | 68.89 | 12.81 | 0.2258 | 0.9037 | 53.45 | 0.4798 |
| pittosporum | 0.6077 | 66.42 | 13.16 | 0.2253 | 0.9107 | 50.14 | 0.5033 |
| Pittosporumbalansae | 0.6135* | 66.22 | 13.18 | 0.2252 | 0.9113 | 49.86 | 0.5052 |
| Pittosporumcrispulum | 0.6135* | 66.22 | 13.18 | 0.2252 | 0.9113 | 49.86 | 0.5052 |
| Pittosporumperryanum | 0.6135* | 66.22 | 13.18 | 0.2252 | 0.9113 | 49.86 | 0.5052 |
| araliaceae | 0.4737 | 71.17 | 12.49 | 0.2262 | 0.8973 | 56.51 | 0.4582 |
| heteropanax | 0.3638 | 75.07 | 11.94 | 0.2270 | 0.8862 | 61.74 | 0.4212 |
| Heteropanaxfragrans | 0.3440* | 75.77 | 11.85 | 0.2271 | 0.8843 | 62.68 | 0.4145 |
| schefflera | 0.3975 | 73.88 | 12.11 | 0.2268 | 0.8896 | 60.13 | 0.4325 |
| Scheffleraha inanensis | 0.3946* | 73.98 | 12.10 | 0.2268 | 0.8893 | 60.27 | 0.4315 |
| Scheffleraheptaphylla | 0.3946* | 73.98 | 12.10 | 0.2268 | 0.8893 | 60.27 | 0.4315 |
| dendropanax | 0.4178 | 73.16 | 12.21 | 0.2266 | 0.8917 | 59.17 | 0.4394 |
| Dendropanaxhainanensis | 0.4198* | 73.08 | 12.22 | 0.2266 | 0.8919 | 59.07 | 0.4400 |
| dipsacales | 0.5508 | 68.44 | 12.87 | 0.2257 | 0.9050 | 52.85 | 0.4841 |
| adoxaceae | 0.5537 | 68.34 | 12.89 | 0.2256 | 0.9053 | 52.71 | 0.4851 |
| sambucus | 0.4821 | 70.88 | 12.53 | 0.2262 | 0.8981 | 56.11 | 0.4610 |
| Sambucusjavanica | 0.4463* | 72.14 | 12.35 | 0.2264 | 0.8945 | 57.81 | 0.4490 |
| viburnum | 0.5923 | 66.97 | 13.08 | 0.2254 | 0.9092 | 50.87 | 0.4981 |
| Viburnumpunctatum | 0.6310* | 65.60 | 13.27 | 0.2251 | 0.9131 | 49.03 | 0.5111 |
| escalloniales | 0.5514 | 68.42 | 12.88 | 0.2257 | 0.9051 | 52.81 | 0.4843 |
| escalloniaceae | 0.5515 | 68.41 | 12.88 | 0.2257 | 0.9051 | 52.81 | 0.4844 |
| polyosma | 0.5519 | 68.40 | 12.88 | 0.2257 | 0.9051 | 52.79 | 0.4845 |
| Polyosmacambodiana | 0.5520* | 68.40 | 12.88 | 0.2257 | 0.9051 | 52.79 | 0.4845 |
| aquifoliales | 0.5599 | 68.12 | 12.92 | 0.2256 | 0.9059 | 52.41 | 0.4872 |
| stemonuraceae | 0.5250 | 69.36 | 12.74 | 0.2258 | 0.9024 | 54.07 | 0.4754 |
| gomphandra | 0.4905 | 70.58 | 12.57 | 0.2261 | 0.8990 | 55.71 | 0.4638 |
| Gomphandratetrandra | 0.4560* | 71.80 | 12.40 | 0.2263 | 0.8955 | 57.35 | 0.4522 |
| cardiopteridaceae | 0.5935 | 66.93 | 13.08 | 0.2254 | 0.9093 | 50.81 | 0.4985 |
| gonocaryum | 0.6275 | 65.72 | 13.25 | 0.2251 | 0.9127 | 49.20 | 0.5099 |
| Gonocaryumlobbianum | 0.6615* | 64.52 | 13.42 | 0.2249 | 0.9161 | 47.58 | 0.5214 |
| aquifoliaceae | 0.5659 | 67.91 | 12.95 | 0.2256 | 0.9065 | 52.13 | 0.4892 |
| ilex | 0.5689 | 67.80 | 12.96 | 0.2255 | 0.9068 | 51.98 | 0.4902 |
| Ilexsp12 | 0.5627* | 68.02 | 12.93 | 0.2256 | 0.9062 | 52.28 | 0.4881 |
| Ilexsp2 | 0.5627* | 68.02 | 12.93 | 0.2256 | 0.9062 | 52.28 | 0.4881 |
| Ilexsp5 | 0.5627* | 68.02 | 12.93 | 0.2256 | 0.9062 | 52.28 | 0.4881 |
| Ilexsterrophylla | 0.6485* | 64.98 | 13.36 | 0.2250 | 0.9148 | 48.20 | 0.5170 |
| Ilextriflora | 0.5627* | 68.02 | 12.93 | 0.2256 | 0.9062 | 52.28 | 0.4881 |
| Ilexangulata | 0.5627* | 68.02 | 12.93 | 0.2256 | 0.9062 | 52.28 | 0.4881 |
| Ilexcochinchinensis | 0.6030* | 66.59 | 13.13 | 0.2253 | 0.9103 | 50.36 | 0.5017 |
| Ilexelmerrilliana | 0.5627* | 68.02 | 12.93 | 0.2256 | 0.9062 | 52.28 | 0.4881 |
| Ilexficoidea | 0.5627* | 68.02 | 12.93 | 0.2256 | 0.9062 | 52.28 | 0.4881 |
| Ilexgodajam | 0.5627* | 68.02 | 12.93 | 0.2256 | 0.9062 | 52.28 | 0.4881 |
| Ilexgoshiensis | 0.5627* | 68.02 | 12.93 | 0.2256 | 0.9062 | 52.28 | 0.4881 |
| Ilexhainanensis | 0.5627* | 68.02 | 12.93 | 0.2256 | 0.9062 | 52.28 | 0.4881 |
| Ilexkobuskiana | 0.5627* | 68.02 | 12.93 | 0.2256 | 0.9062 | 52.28 | 0.4881 |
| Ilexlancilimba | 0.5627* | 68.02 | 12.93 | 0.2256 | 0.9062 | 52.28 | 0.4881 |

| | | | | | | | |
|---|---|---|---|---|---|---|---|
| Ilexnuculicava | 0.5627* | 68.02 | 12.93 | 0.2256 | 0.9062 | 52.28 | 0.4881 |
| Ilexpubescens | 0.5627* | 68.02 | 12.93 | 0.2256 | 0.9062 | 52.28 | 0.4881 |
| Ilexrotunda | 0.5627* | 68.02 | 12.93 | 0.2256 | 0.9062 | 52.28 | 0.4881 |
| Ilexsp | 0.5627* | 68.02 | 12.93 | 0.2256 | 0.9062 | 52.28 | 0.4881 |
| Ilexsp0502 | 0.5627* | 68.02 | 12.93 | 0.2256 | 0.9062 | 52.28 | 0.4881 |
| Ilexsp1 | 0.5627* | 68.02 | 12.93 | 0.2256 | 0.9062 | 52.28 | 0.4881 |
| ericales | 0.5577 | 67.40 | 12.92 | 0.2272 | 0.8981 | 52.29 | 0.4886 |
| pentaphyllacaceae | 0.5621 | 51.81 | 13.01 | 0.2432 | 0.8415 | 49.97 | 0.5508 |
| pentaphylax | 0.5481 | 52.31 | 12.94 | 0.2433 | 0.8401 | 50.64 | 0.5461 |
| Pentaphylaxeuryoides | 0.5340* | 52.80 | 12.87 | 0.2434 | 0.8387 | 51.31 | 0.5413 |
| ternstroemia | 0.5878 | 45.75 | 13.17 | 0.2488 | 0.8207 | 48.05 | 0.5959 |
| Ternstroemiagymnanthera | 0.5814* | 45.98 | 13.14 | 0.2488 | 0.8201 | 48.36 | 0.5937 |
| Ternstroemiahainanensis | 0.5814* | 45.98 | 13.14 | 0.2488 | 0.8201 | 48.36 | 0.5937 |
| anneslea | 0.6425 | 43.82 | 13.45 | 0.2484 | 0.8262 | 45.45 | 0.6143 |
| Annesleafragrans | 0.6843* | 42.33 | 13.65 | 0.2481 | 0.8304 | 43.47 | 0.6284 |
| eurya | 0.5241 | 25.81 | 14.76 | 0.2074 | 0.7681 | 44.07 | 0.6613 |
| Euryaciliata | 0.5200* | 17.43* | 15.67* | 0.1837* | 0.7563* | 41.10* | 0.6852* |
| Euryacuneata | 0.5200* | 25.96 | 14.74 | 0.2075 | 0.7677 | 44.26 | 0.6600 |
| Euryagroffii | 0.5200* | 25.96 | 14.74 | 0.2075 | 0.7677 | 44.26 | 0.6600 |
| Euryahainanensis | 0.5200* | 25.96 | 14.74 | 0.2075 | 0.7677 | 44.26 | 0.6600 |
| Euryaloquaiana | 0.5200* | 25.96 | 14.74 | 0.2075 | 0.7677 | 44.26 | 0.6600 |
| Euryanitida | 0.5300* | 25.60 | 14.79 | 0.2074 | 0.7687 | 43.78 | 0.6633 |
| adinandra | 0.5331 | 52.12 | 11.61 | 0.2893 | 0.7180 | 54.45 | 0.6598 |
| Adinandraangustifolia | 0.5814* | 50.41 | 11.85 | 0.2890 | 0.7228 | 52.16 | 0.6761 |
| Adinandrahainanensis | 0.4700* | 60.53* | 10.87* | 0.2946* | 0.6497* | 59.48* | 0.6733* |
| cleyera | 0.5578 | 42.28 | 11.68 | 0.3090 | 0.8326 | 51.70 | 0.6100 |
| Cleyeraobscurinervia | 0.5677* | 39.13* | 11.26* | 0.3336* | 0.8837* | 51.68* | 0.5900* |
| ebenaceae | 0.6046 | 58.56 | 13.58 | 0.2071 | 0.8929 | 45.25 | 0.4672 |
| diospyros | 0.5900 | 55.47 | 13.99 | 0.1872 | 0.8638 | 41.45 | 0.4902 |
| Diospyrossusarticulata | 0.5643* | 56.38 | 13.86 | 0.1874 | 0.8612 | 42.67 | 0.4815 |
| Diospyroschunii | 0.5643* | 56.38 | 13.86 | 0.1874 | 0.8612 | 42.67 | 0.4815 |
| Diospyroseriantha | 0.7026* | 51.48 | 14.55 | 0.1864 | 0.8751 | 36.10 | 0.5281 |
| Diospyroshainanensis | 0.5643* | 56.38 | 13.86 | 0.1874 | 0.8612 | 42.67 | 0.4815 |
| Diospyrosinflata | 0.5643* | 56.38 | 13.86 | 0.1874 | 0.8612 | 42.67 | 0.4815 |
| Diospyroslongibracteata | 0.5100* | 58.31 | 13.59 | 0.1878 | 0.8558 | 45.26 | 0.4632 |
| Diospyrosmaclurei | 0.7040* | 51.43 | 14.55 | 0.1864 | 0.8752 | 36.03 | 0.5285 |
| Diospyrosmorrisiana | 0.5643* | 54.58* | 14.10* | 0.1773* | 0.8474* | 40.42* | 0.4955* |
| Diospyrosstrigosa | 0.5643* | 56.38 | 13.86 | 0.1874 | 0.8612 | 42.67 | 0.4815 |
| primulaceae | 0.6407 | 59.09 | 13.51 | 0.2169 | 0.9104 | 45.79 | 0.4654 |
| myrsine | 0.7145 | 56.47 | 13.88 | 0.2164 | 0.9178 | 42.28 | 0.4902 |
| Myrsineseguinii | 0.7410* | 55.53 | 14.01 | 0.2162 | 0.9205 | 41.01 | 0.4992 |
| Myrsinestolonifera | 0.7410* | 55.53 | 14.01 | 0.2162 | 0.9205 | 41.01 | 0.4992 |
| ardisia | 0.6111 | 60.14 | 13.37 | 0.2171 | 0.9074 | 47.19 | 0.4555 |
| Ardisiacrassinervosa | 0.6265* | 59.59 | 13.44 | 0.2170 | 0.9090 | 46.46 | 0.4606 |
| Ardisiadensilepidotula | 0.6265* | 59.59 | 13.44 | 0.2170 | 0.9090 | 46.46 | 0.4606 |
| Ardisiaobtusa | 0.5909* | 60.85 | 13.27 | 0.2173 | 0.9054 | 48.15 | 0.4487 |
| Ardisiaquinquegona | 0.5909* | 60.85 | 13.27 | 0.2173 | 0.9054 | 48.15 | 0.4487 |
| Ardisiavillosa | 0.5909* | 60.85 | 13.27 | 0.2173 | 0.9054 | 48.15 | 0.4487 |
| Ardisiavirens | 0.5909* | 60.85 | 13.27 | 0.2173 | 0.9054 | 48.15 | 0.4487 |
| Embelia | 0.6400 | 59.11 | 13.51 | 0.2169 | 0.9103 | 45.82 | 0.4652 |
| Embeliavestita | 0.6393* | 59.14 | 13.51 | 0.2169 | 0.9102 | 45.85 | 0.4649 |
| maesa | 0.6672 | 58.15 | 13.65 | 0.2167 | 0.9130 | 44.53 | 0.4743 |
| Maesaacuminatissima | 0.6760* | 57.84 | 13.69 | 0.2166 | 0.9139 | 44.11 | 0.4773 |
| Maesaconsanguinea | 0.6760* | 57.84 | 13.69 | 0.2166 | 0.9139 | 44.11 | 0.4773 |
| Maesaperlarius | 0.6760* | 57.84 | 13.69 | 0.2166 | 0.9139 | 44.11 | 0.4773 |
| sapotaceae | 0.5887 | 66.13 | 12.71 | 0.2331 | 0.9609 | 53.16 | 0.3672 |
| planchonella | 0.6952 | 69.39 | 10.90 | 0.2431 | 0.9301 | 53.83 | 0.4062 |
| Planchonellaclemensii | 0.7148* | 68.69 | 10.99 | 0.2430 | 0.9320 | 52.90 | 0.4128 |
| chrysophyllum | 0.5699 | 73.83 | 10.27 | 0.2440 | 0.9175 | 59.79 | 0.3640 |
| Chrysophyllumlanceolatum | 0.5230* | 75.49 | 10.04 | 0.2444 | 0.9128 | 62.02 | 0.3483 |
| madhuca | 0.8592 | 65.42 | 9.549 | 0.2626 | 0.9250 | 45.31 | 0.4842 |
| Madhucahainanensis | 0.9257* | 63.99* | 8.798* | 0.2724* | 0.9209* | 41.78* | 0.5181* |
| pouteria | 0.6678 | 72.78 | 13.82 | 0.2025 | 0.9395 | 62.68 | 0.3430 |

| | | | | | | | |
|---|---|---|---|---|---|---|---|
| Pouteriaannamensis | **0.6878*** | **76.34*** | **14.82*** | **0.1822*** | **0.9321*** | **68.54*** | **0.3186*** |
| sarcosperma | **0.5277** | **66.49** | **12.28** | **0.2492** | **1.017** | **52.27** | **0.2996** |
| Sarcospermalaurinum | **0.4667*** | **66.85*** | **11.85*** | **0.2652*** | **1.073*** | **51.37*** | **0.2320*** |
| ericaceae | **0.5276** | **69.63** | **12.85** | **0.2305** | **0.8676** | **53.35** | **0.5012** |
| rhododendron | **0.4984** | **70.66** | **12.70** | **0.2307** | **0.8647** | **54.74** | **0.4914** |
| Rhododendronmoulmainense | **0.4955*** | **70.76** | **12.69** | **0.2308** | **0.8644** | **54.87** | **0.4904** |
| symplocaceae | **0.5244** | **80.57** | **13.95** | **0.2272** | **0.8393** | **55.34** | **0.5102** |
| symplocos | **0.5317** | **80.32** | **13.99** | **0.2272** | **0.8401** | **55.00** | **0.5126** |
| Symplocoscongesta | **0.5335*** | **80.25** | **14.00** | **0.2272** | **0.8402** | **54.91** | **0.5132** |
| Symplocoscrassilimba | **0.5335*** | **80.25** | **14.00** | **0.2272** | **0.8402** | **54.91** | **0.5132** |
| Symplocoseuryoides | **0.5335*** | **80.25** | **14.00** | **0.2272** | **0.8402** | **54.91** | **0.5132** |
| Symplocosglauca | **0.5335*** | **80.25** | **14.00** | **0.2272** | **0.8402** | **54.91** | **0.5132** |
| Symplocosheishanensis | **0.5335*** | **80.25** | **14.00** | **0.2272** | **0.8402** | **54.91** | **0.5132** |
| Symplocoslancifolia | **0.4635*** | **82.73** | **13.65** | **0.2277** | **0.8332** | **58.24** | **0.4897** |
| Symplocospendula | **0.5335*** | **80.25** | **14.00** | **0.2272** | **0.8402** | **54.91** | **0.5132** |
| Symplocospoilanei | **0.5335*** | **80.25** | **14.00** | **0.2272** | **0.8402** | **54.91** | **0.5132** |
| Symplocospseudobarberina | **0.5335*** | **80.25** | **14.00** | **0.2272** | **0.8402** | **54.91** | **0.5132** |
| Symplocosracemosa | **0.5335*** | **80.25** | **14.00** | **0.2272** | **0.8402** | **54.91** | **0.5132** |
| Symplocossp1 | **0.5335*** | **80.25** | **14.00** | **0.2272** | **0.8402** | **54.91** | **0.5132** |
| Symplocossumuntia | **0.5335*** | **80.25** | **14.00** | **0.2272** | **0.8402** | **54.91** | **0.5132** |
| Symplocosviridissima | **0.5335*** | **80.25** | **14.00** | **0.2272** | **0.8402** | **54.91** | **0.5132** |
| Symplocoswikstroemiifolia | **0.5335*** | **80.25** | **14.00** | **0.2272** | **0.8402** | **54.91** | **0.5132** |
| Symplocosadenophylla | **0.6500*** | **76.12** | **14.58** | **0.2263** | **0.8519** | **49.37** | **0.5524** |
| Symplocosanomala | **0.4815*** | **82.10** | **13.74** | **0.2275** | **0.8350** | **57.38** | **0.4957** |
| Symplocoscochinchinensis | **0.5150*** | **80.91** | **13.91** | **0.2273** | **0.8384** | **55.79** | **0.5070** |
| Symplocaceaesp7 | **0.5335*** | **80.25** | **14.00** | **0.2272** | **0.8402** | **54.91** | **0.5132** |
| styracaceae | **0.4394** | **99.74** | **15.69** | **0.2211** | **0.7995** | **62.94** | **0.4646** |
| alniphyllum | **0.3940** | **133.7** | **19.78** | **0.2079** | **0.7324** | **72.21** | **0.4153** |
| Alniphyllumfortunei | **0.3827*** | **142.1*** | **20.80*** | **0.2046*** | **0.7157*** | **74.53*** | **0.4029*** |
| styrax | **0.4165** | **100.6** | **15.57** | **0.2212** | **0.7972** | **64.03** | **0.4568** |
| Styraxagrestis | **0.4050*** | **101.0** | **15.52** | **0.2213** | **0.7961** | **64.57** | **0.4530** |
| Styraxsuberifolius | **0.4050*** | **101.0** | **15.52** | **0.2213** | **0.7961** | **64.57** | **0.4530** |
| theaceae | **0.5304** | **66.97** | **11.87** | **0.2340** | **0.8589** | **51.57** | **0.5477** |
| camellia | **0.5499** | **51.38** | **9.373** | **0.2441** | **0.9817** | **31.81** | **0.6841** |
| Camelliacaudata | **0.5500*** | **51.38** | **9.374** | **0.2441** | **0.9817** | **31.81** | **0.6841** |
| Camelliaoleifera | **0.5500*** | **51.38** | **9.374** | **0.2441** | **0.9817** | **31.81** | **0.6841** |
| Camelliapaucipunctata | **0.5500*** | **51.38** | **9.374** | **0.2441** | **0.9817** | **31.81** | **0.6841** |
| Camelliasinensis | **0.5500*** | **51.38** | **9.374** | **0.2441** | **0.9817** | **31.81** | **0.6841** |
| Camelliasp1 | **0.5500*** | **51.38** | **9.374** | **0.2441** | **0.9817** | **31.81** | **0.6841** |
| Camelliasp3 | **0.5500*** | **51.38** | **9.374** | **0.2441** | **0.9817** | **31.81** | **0.6841** |
| Camelliaxanthochroma | **0.5500*** | **51.38** | **9.374** | **0.2441** | **0.9817** | **31.81** | **0.6841** |
| polyspora | **0.5613** | **50.11** | **8.820** | **0.2490** | **1.033** | **24.85** | **0.7466** |
| Polysporaaxillaris | **0.5677*** | **49.88** | **8.852** | **0.2490** | **1.034** | **24.54** | **0.7487** |
| Polysporahainanensis | **0.5677*** | **49.01*** | **8.243*** | **0.2540*** | **1.084*** | **18.12*** | **0.8074*** |
| pyrenaria | **0.5225** | **45.40** | **9.505** | **0.2360** | **0.9453** | **35.29** | **0.6017** |
| Pyrenariajonquieriana | **0.5183*** | **37.73*** | **9.142*** | **0.2328*** | **0.9617*** | **31.24*** | **0.5857*** |
| Pyrenariamicrocarpa | **0.5183*** | **45.55** | **9.484** | **0.2360** | **0.9449** | **35.49** | **0.6002** |
| Pyrenariaspectabilis | **0.5183*** | **45.55** | **9.484** | **0.2360** | **0.9449** | **35.49** | **0.6002** |
| schima | **0.5430** | **67.95** | **10.73** | **0.2407** | **0.7357** | **67.19** | **0.5448** |
| Schimaremotiserrata | **0.5526*** | **67.62** | **10.78** | **0.2406** | **0.7366** | **66.73** | **0.5480** |
| Schimasuperba | **0.5375*** | **71.53*** | **10.62*** | **0.2424*** | **0.6712*** | **76.41*** | **0.5259*** |
| cornales | **0.5439** | **69.48** | **12.82** | **0.2242** | **0.9119** | **53.39** | **0.4797** |
| nyssaceae | **0.5066** | **70.80** | **12.64** | **0.2244** | **0.9081** | **55.17** | **0.4671** |
| mastixia | **0.4989** | **71.08** | **12.60** | **0.2245** | **0.9074** | **55.53** | **0.4645** |
| Mastixiapentandra | **0.4950*** | **71.22** | **12.58** | **0.2245** | **0.9070** | **55.72** | **0.4632** |
| cornaceae | **0.4976** | **71.12** | **12.59** | **0.2245** | **0.9072** | **55.59** | **0.4641** |
| alangium | **0.4397** | **73.17** | **12.31** | **0.2249** | **0.9014** | **58.34** | **0.4446** |
| Alangiumchinense | **0.4017*** | **74.52** | **12.12** | **0.2252** | **0.8976** | **60.15** | **0.4318** |
| Alangiumkurzii | **0.4200*** | **73.87** | **12.21** | **0.2251** | **0.8995** | **59.28** | **0.4380** |
| cornus | **0.5425** | **69.53** | **12.82** | **0.2242** | **0.9118** | **53.46** | **0.4792** |
| Cornussp1 | **0.5875*** | **67.94** | **13.04** | **0.2238** | **0.9163** | **51.32** | **0.4944** |
| santalales | **0.6368** | **68.58** | **13.24** | **0.2188** | **0.9439** | **49.64** | **0.5046** |
| olacaceae | **0.7150** | **65.81** | **13.63** | **0.2183** | **0.9518** | **45.92** | **0.5309** |
| olax | **0.7523** | **64.49** | **13.81** | **0.2180** | **0.9555** | **44.14** | **0.5435** |

| | | | | | | | |
|---|---|---|---|---|---|---|---|
| Olaximbricata | 0.7710* | 63.82 | 13.91 | 0.2179 | 0.9574 | 43.26 | 0.5497 |
| santalaceae | 0.6849 | 66.88 | 13.48 | 0.2185 | 0.9488 | 47.35 | 0.5208 |
| scleropyrum | 0.6975 | 66.43 | 13.54 | 0.2184 | 0.9500 | 46.75 | 0.5250 |
| Scleropyrumwallichianum | 0.6993* | 66.37 | 13.55 | 0.2184 | 0.9502 | 46.67 | 0.5256 |
| buxales | 0.6429 | 67.23 | 13.22 | 0.2214 | 0.9809 | 47.48 | 0.4905 |
| buxaceae | 0.6693 | 66.30 | 13.35 | 0.2212 | 0.9835 | 46.23 | 0.4993 |
| buxus | 0.6956 | 65.36 | 13.48 | 0.2210 | 0.9861 | 44.97 | 0.5082 |
| Buxusmyrica | 0.7220* | 64.43 | 13.61 | 0.2209 | 0.9888 | 43.72 | 0.5171 |
| proteales | 0.5483 | 68.66 | 12.72 | 0.2263 | 1.000 | 49.89 | 0.4446 |
| proteaceae | 0.5517 | 68.54 | 12.73 | 0.2263 | 1.000 | 49.73 | 0.4458 |
| helicia | 0.5887 | 67.23 | 12.92 | 0.2260 | 1.004 | 47.97 | 0.4582 |
| Heliciacochinchinensis | 0.5510* | 68.56 | 12.73 | 0.2263 | 1.000 | 49.76 | 0.4455 |
| Heliciaformosana | 0.6051* | 66.64 | 13.00 | 0.2259 | 1.006 | 47.19 | 0.4637 |
| Heliciahainanensis | 0.6051* | 66.64 | 13.00 | 0.2259 | 1.006 | 47.19 | 0.4637 |
| Helicialongipetiolata | 0.6051* | 66.64 | 13.00 | 0.2259 | 1.006 | 47.19 | 0.4637 |
| Heliciaobovatifolia | 0.5777* | 67.62 | 12.86 | 0.2261 | 1.003 | 48.50 | 0.4545 |
| Heliciareticulata | 0.6051* | 66.64 | 13.00 | 0.2259 | 1.006 | 47.19 | 0.4637 |
| heliciopsis | 0.4612 | 71.75 | 12.28 | 0.2269 | 0.9914 | 54.04 | 0.4153 |
| Heliciopsislobata | 0.4300* | 72.85 | 12.13 | 0.2271 | 0.9882 | 55.52 | 0.4048 |
| Heliciopsisterminalis | 0.4767* | 71.20 | 12.36 | 0.2268 | 0.9929 | 53.30 | 0.4205 |
| sabiaceae | 0.5297 | 69.32 | 12.62 | 0.2264 | 0.9982 | 50.78 | 0.4384 |
| meliosma | 0.5110 | 69.98 | 12.53 | 0.2265 | 0.9964 | 51.67 | 0.4321 |
| Meliosmaangustifolia | 0.5483* | 68.66 | 12.72 | 0.2263 | 1.000 | 49.89 | 0.4446 |
| Meliosmadumicola | 0.4789* | 71.12 | 12.37 | 0.2268 | 0.9931 | 53.19 | 0.4213 |
| Meliosmafordii | 0.4789* | 71.12 | 12.37 | 0.2268 | 0.9931 | 53.19 | 0.4213 |
| Meliosmalaui | 0.4789* | 71.12 | 12.37 | 0.2268 | 0.9931 | 53.19 | 0.4213 |
| Meliosmarigida | 0.5960* | 66.97 | 12.95 | 0.2259 | 1.005 | 47.63 | 0.4607 |
| Meliosmasquamulata | 0.4985* | 70.42 | 12.47 | 0.2266 | 0.9951 | 52.26 | 0.4279 |
| Meliosmathorelii | 0.4789* | 71.12 | 12.37 | 0.2268 | 0.9931 | 53.19 | 0.4213 |
| monocots | 0.5388 | 66.10 | 12.62 | 0.2326 | 1.042 | 47.22 | 0.4204 |
| arecales | 0.5312 | 66.37 | 12.58 | 0.2327 | 1.041 | 47.58 | 0.4178 |
| arecaceae | 0.5368 | 66.17 | 12.61 | 0.2326 | 1.042 | 47.31 | 0.4197 |
| pinanga | 0.5518 | 65.64 | 12.68 | 0.2325 | 1.044 | 46.60 | 0.4248 |
| Pinangabaviensis | 0.5518* | 65.64 | 12.68 | 0.2325 | 1.044 | 46.60 | 0.4247 |
| livistona | 0.5737 | 64.86 | 12.79 | 0.2324 | 1.046 | 45.56 | 0.4321 |
| Livistonasaribus | 0.5814* | 64.59 | 12.83 | 0.2323 | 1.047 | 45.19 | 0.4347 |
| licuala | 0.5533 | 65.58 | 12.69 | 0.2325 | 1.044 | 46.52 | 0.4253 |
| Licualafordiana | 0.5518* | 65.64 | 12.68 | 0.2325 | 1.044 | 46.60 | 0.4247 |
| Licualahainanensis | 0.5518* | 65.64 | 12.68 | 0.2325 | 1.044 | 46.60 | 0.4247 |
| caryota | 0.5512 | 65.66 | 12.68 | 0.2325 | 1.044 | 46.63 | 0.4245 |
| Caryotamaxima | 0.5518* | 65.64 | 12.68 | 0.2325 | 1.044 | 46.60 | 0.4247 |
| arenga | 0.5512 | 65.66 | 12.68 | 0.2325 | 1.044 | 46.63 | 0.4245 |
| Arengapinnata | 0.5518* | 65.64 | 12.68 | 0.2325 | 1.044 | 46.60 | 0.4247 |
| asparagales | 0.5104 | 67.11 | 12.48 | 0.2328 | 1.039 | 48.57 | 0.4108 |
| asparagaceae | 0.4349 | 69.78 | 12.10 | 0.2334 | 1.032 | 52.15 | 0.3854 |
| dracaena | 0.3594 | 72.46 | 11.73 | 0.2339 | 1.024 | 55.74 | 0.3600 |
| Dracaenaangustifolia | 0.3500* | 72.79 | 11.68 | 0.2340 | 1.023 | 56.19 | 0.3568 |
| magnoliids | 0.5339 | 64.34 | 12.56 | 0.2368 | 1.071 | 45.36 | 0.4047 |
| magnoniales | 0.5151 | 63.90 | 12.61 | 0.2323 | 1.114 | 44.43 | 0.3692 |
| annonaceae | 0.5547 | 62.06 | 13.65 | 0.1981 | 1.134 | 37.72 | 0.3995 |
| miliusa | 0.6161 | 59.89 | 13.96 | 0.1976 | 1.140 | 34.80 | 0.4201 |
| Miliusahorsfieldii | 0.6100* | 60.11 | 13.93 | 0.1977 | 1.140 | 35.09 | 0.4181 |
| popowia | 0.5643 | 61.73 | 13.70 | 0.1980 | 1.135 | 37.26 | 0.4027 |
| Popowiapisocarpa | 0.5450* | 62.41 | 13.60 | 0.1981 | 1.133 | 38.18 | 0.3962 |
| alphonsea | 0.7078 | 56.64 | 14.41 | 0.1970 | 1.150 | 30.44 | 0.4510 |
| Alphonseahainanensis | 0.7380* | 55.57 | 14.56 | 0.1968 | 1.153 | 29.00 | 0.4612 |
| mitrephora | 0.6614 | 58.29 | 14.18 | 0.1973 | 1.145 | 32.65 | 0.4354 |
| Mitrephoratomentosa | 0.6800* | 57.62 | 14.27 | 0.1972 | 1.147 | 31.76 | 0.4416 |
| orophea | 0.5835 | 61.05 | 13.79 | 0.1979 | 1.137 | 36.35 | 0.4092 |
| Oropheahainanensis | 0.5656* | 61.68 | 13.71 | 0.1980 | 1.135 | 37.20 | 0.4031 |
| polyalthia | 0.6103 | 60.10 | 13.93 | 0.1977 | 1.140 | 35.08 | 0.4182 |
| Polyalthiacerasoides | 0.7550* | 54.97 | 14.65 | 0.1966 | 1.154 | 28.20 | 0.4669 |
| Polyalthialaui | 0.5541* | 62.09 | 13.65 | 0.1981 | 1.134 | 37.75 | 0.3993 |
| Polyalthiaobliqua | 0.5541* | 62.09 | 13.65 | 0.1981 | 1.134 | 37.75 | 0.3993 |

| | | | | | | | |
|---|---|---|---|---|---|---|---|
| Polyalthiarumphii | 0.5800* | 61.17 | 13.78 | 0.1979 | 1.137 | 36.52 | 0.4080 |
| disepalum | 0.5637 | 61.33 | 13.92 | 0.1897 | 1.135 | 35.60 | 0.4132 |
| Disepalumplagioneurum | 0.5656* | 61.27 | 13.93 | 0.1897 | 1.136 | 35.51 | 0.4138 |
| uvaria | 0.5649 | 60.05 | 14.60 | 0.1646 | 1.136 | 30.46 | 0.4456 |
| Uvariaboniana | 0.5656* | 60.03 | 14.60 | 0.1646 | 1.136 | 30.43 | 0.4458 |
| dasymaschalon | 0.5648 | 58.81 | 15.27 | 0.1396 | 1.137 | 25.38 | 0.4776 |
| Dasymaschalonrostratum | 0.5656* | 58.37* | 15.50* | 0.1312* | 1.137* | 23.65* | 0.4885* |
| Chieniodendron | 0.5602 | 61.87 | 13.68 | 0.1980 | 1.135 | 37.46 | 0.4013 |
| Chieniodendronhainanense | 0.5656* | 61.68 | 13.71 | 0.1980 | 1.135 | 37.20 | 0.4031 |
| magnoliaceae | 0.4798 | 67.19 | 12.56 | 0.2231 | 1.142 | 48.31 | 0.3165 |
| Manglietia | 0.4517 | 70.99 | 14.13 | 0.2507 | 1.137 | 46.95 | 0.2890 |
| Manglietiafordiana | 0.4235* | 74.80* | 15.69* | 0.2783* | 1.131* | 45.60* | 0.2615* |
| Michelia | 0.5013 | 75.82 | 11.01 | 0.2073 | 1.257 | 42.92 | 0.3275 |
| Micheliabalansae | 0.5423* | 111.1* | 12.06* | 0.1597* | 1.163* | 52.79* | 0.3705* |
| Micheliagioi | 0.4880* | 76.29 | 10.95 | 0.2074 | 1.255 | 43.55 | 0.3230 |
| Micheliamediocris | 0.4950* | 48.68* | 8.481* | 0.2391* | 1.466* | 27.01* | 0.3000* |
| Lirianthe | 0.4607 | 56.90 | 12.37 | 0.2110 | 1.046 | 58.25 | 0.2984 |
| Lirianthechampionii | 0.4415* | 46.61* | 12.17* | 0.1989* | 0.9502* | 68.18* | 0.2804* |
| laurales | 0.5292 | 60.80 | 12.31 | 0.2520 | 1.097 | 42.20 | 0.3972 |
| lauraceae | 0.5482 | 53.01 | 11.63 | 0.2956 | 1.092 | 36.10 | 0.4360 |
| phoebe | 0.5206 | 52.30 | 16.36 | 0.2025 | 0.9065 | 67.21 | 0.3345 |
| Phoebehungmoensis | 0.4900* | 54.74* | 18.02* | 0.1932* | 0.8283* | 78.57* | 0.3097* |
| Phoebetavoyana | 0.5382* | 51.68 | 16.45 | 0.2024 | 0.9082 | 66.38 | 0.3405 |
| machilus | 0.5626 | 44.70 | 11.63 | 0.2266 | 1.092 | 38.52 | 0.3825 |
| Machiluschinensis | 0.5643* | 30.29* | 9.637* | 0.2368* | 1.039* | 33.58* | 0.4085* |
| Machiluscicatricosa | 0.5643* | 60.10* | 14.94* | 0.2296* | 0.9563* | 53.94* | 0.3410* |
| Machilusfoonchewii | 0.5643* | 44.64 | 11.64 | 0.2266 | 1.092 | 38.44 | 0.3831 |
| Machilusgamblei | 0.5643* | 44.64 | 11.64 | 0.2266 | 1.092 | 38.44 | 0.3831 |
| Machilusmonticola | 0.5643* | 40.13* | 9.022* | 0.2188* | 1.312* | 20.91* | 0.4047* |
| Machiluspomifera | 0.5643* | 44.64 | 11.64 | 0.2266 | 1.092 | 38.44 | 0.3831 |
| Machilusrobusta | 0.5643* | 44.64 | 11.64 | 0.2266 | 1.092 | 38.44 | 0.3831 |
| Machilussp | 0.5643* | 44.64 | 11.64 | 0.2266 | 1.092 | 38.44 | 0.3831 |
| Machilusvelutina | 0.5643* | 44.64 | 11.64 | 0.2266 | 1.092 | 38.44 | 0.3831 |
| neolitsea | 0.5343 | 43.42 | 10.60 | 0.1943 | 1.035 | 34.34 | 0.4330 |
| Neolitseacambodiana | 0.5415* | 51.87* | 9.074* | 0.2055* | 0.9582* | 37.11* | 0.4346* |
| Neolitseachui | 0.5415* | 43.17 | 10.63 | 0.1942 | 1.036 | 34.00 | 0.4354 |
| Neolitseaellipsoidea | 0.4527* | 70.99* | 9.478* | 0.1966* | 1.139* | 28.84* | 0.4301* |
| Neolitseaoblongifolia | 0.5300* | 32.76* | 13.92* | 0.1760* | 0.9601* | 48.02* | 0.3559* |
| Neolitseaovatifolia | 0.5415* | 21.44* | 8.757* | 0.1999* | 0.9748* | 27.93* | 0.5542* |
| Neolitseaphanerophlebia | 0.5900* | 41.45 | 10.87 | 0.1939 | 1.041 | 31.69 | 0.4518 |
| Neolitseapulchella | 0.5415* | 39.89* | 11.12* | 0.1849* | 1.111* | 31.26* | 0.3801* |
| alseodaphne | 0.5678 | 65.31 | 11.01 | 0.2044 | 1.397 | 30.98 | 0.3750 |
| Alseodaphnehainanensis | 0.5970* | 73.13* | 10.35* | 0.1660* | 1.576* | 22.83* | 0.3626* |
| lindera | 0.4822 | 51.74 | 14.23 | 0.3627 | 1.086 | 48.90 | 0.4159 |
| Linderacommunis | 0.4519* | 52.81 | 14.08 | 0.3629 | 1.083 | 50.34 | 0.4057 |
| Linderakwangtungensis | 0.5778* | 55.61* | 22.38* | 0.1394* | 0.7492* | 70.48* | 0.4011* |
| Linderametcalfiana | 0.4519* | 52.81 | 14.08 | 0.3629 | 1.083 | 50.34 | 0.4057 |
| Linderanacusua | 0.4519* | 37.51* | 14.26* | 0.7802* | 1.649* | 15.61* | 0.5077* |
| Linderarobusta | 0.4519* | 57.49* | 8.441* | 0.2364* | 0.8426* | 60.69* | 0.3778* |
| cinnamomum | 0.4926 | 52.82 | 8.723 | 0.2526 | 0.9306 | 57.09 | 0.3689 |
| Cinnamomumbejolghota | 0.4671* | 53.72 | 8.596 | 0.2527 | 0.9280 | 58.30 | 0.3603 |
| Cinnamomumburmannii | 0.4900* | 25.01* | 8.371* | 0.1910* | 0.8821* | 49.60* | 0.4087* |
| Cinnamomumliangii | 0.4671* | 53.72 | 8.596 | 0.2527 | 0.9280 | 58.30 | 0.3603 |
| Cinnamomumparthenoxylon | 0.5800* | 66.42* | 8.945* | 0.2587* | 0.8164* | 79.62* | 0.3030* |
| Cinnamomumrigidissimum | 0.4671* | 64.03* | 7.389* | 0.2829* | 1.003* | 46.07* | 0.3983* |
| Cinnamomumsubavenium | 0.5000* | 52.55 | 8.760 | 0.2525 | 0.9313 | 56.74 | 0.3714 |
| Cinnamomumtsoi | 0.4671* | 53.72 | 8.596 | 0.2527 | 0.9280 | 58.30 | 0.3603 |
| cryptocarya | 0.5675 | 43.86 | 10.80 | 0.2398 | 1.124 | 28.17 | 0.4446 |
| Cryptocaryachinensis | 0.5000* | 43.24* | 11.93* | 0.1607* | 1.285* | 21.53* | 0.4040* |
| Cryptocaryachingii | 0.5400* | 41.18* | 8.466* | 0.2529* | 0.9846* | 33.64* | 0.4472* |
| Cryptocaryadensiflora | 0.5357* | 44.99 | 10.64 | 0.2400 | 1.121 | 29.69 | 0.4339 |
| Cryptocaryaimpressinervia | 0.5613* | 44.09 | 10.77 | 0.2398 | 1.123 | 28.47 | 0.4425 |
| Cryptocaryamaclurei | 0.5613* | 44.09 | 10.77 | 0.2398 | 1.123 | 28.47 | 0.4425 |
| Cryptocaryametcalfiana | 0.7600* | 37.04 | 11.76 | 0.2384 | 1.143 | 19.02 | 0.5094 |

| | | | | | | | |
|---|---|---|---|---|---|---|---|
| Cryptocaryasp9 | 0.5613* | 44.09 | 10.77 | 0.2398 | 1.123 | 28.47 | 0.4425 |
| litsea | 0.4348 | 53.19 | 12.45 | 0.3891 | 0.9797 | 45.27 | 0.4203 |
| Litseabaviensis | 0.4353* | 61.06* | 13.13* | 0.2440* | 1.226* | 34.50* | 0.3329* |
| Litseacubeba | 0.3100* | 57.61 | 11.83 | 0.3900 | 0.9672 | 51.20 | 0.3783 |
| Litseaelongata | 0.4255* | 53.52 | 12.40 | 0.3892 | 0.9787 | 45.71 | 0.4172 |
| Litsealancilimba | 0.5317* | 49.75 | 12.93 | 0.3884 | 0.9894 | 40.66 | 0.4529 |
| Litseamonopetala | 0.4228* | 53.61 | 12.39 | 0.3892 | 0.9785 | 45.84 | 0.4163 |
| Litseapseudoelongata | 0.4255* | 53.52 | 12.40 | 0.3892 | 0.9787 | 45.71 | 0.4172 |
| Litseavariabilischinensis | 0.4255* | 53.52 | 12.40 | 0.3892 | 0.9787 | 45.71 | 0.4172 |
| Litseavariabilis | 0.4255* | 47.30* | 13.51* | 0.2430* | 0.8833* | 57.04* | 0.3923* |
| Litseaverticillata | 0.4255* | 49.78* | 12.18* | 0.7621* | 0.7288* | 50.21* | 0.5441* |
| beilschmiedia | 0.5632 | 78.06 | 8.070 | 0.3201 | 1.410 | 18.55 | 0.4883 |
| Beilschmiediaappendiculata | 0.5631* | 78.06 | 8.069 | 0.3201 | 1.410 | 18.55 | 0.4883 |
| Beilschmiediaglauca | 0.5631* | 78.06 | 8.069 | 0.3201 | 1.410 | 18.55 | 0.4883 |
| Beilschmiediaintermedia | 0.5148* | 79.78 | 7.829 | 0.3204 | 1.405 | 20.85 | 0.4720 |
| Beilschmiedialaevis | 0.5631* | 84.90* | 7.209* | 0.3234* | 1.489* | 14.66* | 0.4980* |
| Beilschmiediaobconica | 0.5631* | 78.06 | 8.069 | 0.3201 | 1.410 | 18.55 | 0.4883 |
| Beilschmiediapercoriacea | 0.5631* | 78.06 | 8.069 | 0.3201 | 1.410 | 18.55 | 0.4883 |
| Beilschmiediapergamentacea | 0.5631* | 78.06 | 8.069 | 0.3201 | 1.410 | 18.55 | 0.4883 |
| Beilschmiediaroxburghiana | 0.5631* | 78.06 | 8.069 | 0.3201 | 1.410 | 18.55 | 0.4883 |
| Beilschmiediasp | 0.6063* | 76.53 | 8.284 | 0.3197 | 1.414 | 16.50 | 0.5028 |
| Beilschmiediasp0502 | 0.5631* | 78.06 | 8.069 | 0.3201 | 1.410 | 18.55 | 0.4883 |
| Beilschmiediasp4 | 0.5631* | 78.06 | 8.069 | 0.3201 | 1.410 | 18.55 | 0.4883 |
| Beilschmiediasp6 | 0.5631* | 78.06 | 8.069 | 0.3201 | 1.410 | 18.55 | 0.4883 |
| Beilschmiediatsangii | 0.5631* | 78.06 | 8.069 | 0.3201 | 1.410 | 18.55 | 0.4883 |
| Beilschmiediatungfangensis | 0.5631* | 78.06 | 8.069 | 0.3201 | 1.410 | 18.55 | 0.4883 |
| Beilschmiediawangii | 0.5631* | 78.06 | 8.069 | 0.3201 | 1.410 | 18.55 | 0.4883 |
| endiandra | 0.5923 | 70.19 | 9.075 | 0.3165 | 1.333 | 21.05 | 0.4883 |
| Endiandrahainanensis | 0.6150* | 69.38 | 9.188 | 0.3163 | 1.335 | 19.97 | 0.4959 |
| Lauraceaesp5 | 0.5643* | 52.44 | 11.71 | 0.2955 | 1.093 | 35.33 | 0.4415 |
| Lauraceaesp8 | 0.5643* | 52.44 | 11.71 | 0.2955 | 1.093 | 35.33 | 0.4415 |
| austrobaileyales | 0.5515 | 64.69 | 12.66 | 0.2346 | 1.058 | 45.57 | 0.4176 |
| schisandraceae | 0.5698 | 64.03 | 12.75 | 0.2345 | 1.060 | 44.70 | 0.4238 |
| illicium | 0.5759 | 63.82 | 12.78 | 0.2344 | 1.060 | 44.41 | 0.4259 |
| Illiciumsp1 | 0.5790* | 63.71 | 12.80 | 0.2344 | 1.061 | 44.26 | 0.4269 |
| Illiciumternstroemioides | 0.5790* | 63.71 | 12.80 | 0.2344 | 1.061 | 44.26 | 0.4269 |
| gymnosperms | 0.5044 | 66.35 | 12.43 | 0.2349 | 1.053 | 47.81 | 0.4018 |
| pinales | 0.4858 | 67.01 | 12.34 | 0.2351 | 1.051 | 48.69 | 0.3955 |
| pinaceae | 0.5028 | 66.41 | 12.42 | 0.2350 | 1.053 | 47.89 | 0.4012 |
| pinus | 0.5537 | 64.61 | 12.67 | 0.2346 | 1.058 | 45.47 | 0.4184 |
| Pinuscaribaea | 0.5707* | 64.01 | 12.76 | 0.2345 | 1.060 | 44.66 | 0.4241 |
| podocarpaceae | 0.4644 | 67.77 | 12.23 | 0.2352 | 1.049 | 49.71 | 0.3884 |
| podocarpus | 0.4769 | 67.33 | 12.29 | 0.2351 | 1.050 | 49.12 | 0.3925 |
| Podocarpusneriifolius | 0.4769* | 67.33 | 12.29 | 0.2351 | 1.050 | 49.12 | 0.3925 |
| dacrycarpus | 0.4463 | 68.41 | 12.14 | 0.2354 | 1.047 | 50.57 | 0.3822 |
| Dacrycarpusimbricatus | 0.4133* | 69.58 | 11.98 | 0.2356 | 1.044 | 52.14 | 0.3712 |
| dacrydium | 0.5502 | 64.73 | 12.66 | 0.2346 | 1.058 | 45.63 | 0.4172 |
| Dacrydiumpectinatum | 0.5856* | 63.48 | 12.83 | 0.2344 | 1.061 | 43.95 | 0.4291 |
| cupressaceae | 0.3732 | 71.00 | 11.78 | 0.2359 | 1.040 | 54.05 | 0.3576 |
| cunninghamia | 0.3445 | 72.02 | 11.64 | 0.2361 | 1.037 | 55.41 | 0.3480 |
| Cunninghamialanceolata | 0.3157* | 73.04 | 11.49 | 0.2363 | 1.034 | 56.78 | 0.3383 |
| monilophyte | 0.5347 | 65.28 | 12.58 | 0.2347 | 1.056 | 46.37 | 0.4120 |
| cyatheales | 0.5619 | 64.32 | 12.72 | 0.2345 | 1.059 | 45.07 | 0.4212 |
| cyatheaceae | 0.5736 | 63.90 | 12.77 | 0.2344 | 1.060 | 44.52 | 0.4251 |
| Alsophila | 0.5775 | 63.76 | 12.79 | 0.2344 | 1.061 | 44.33 | 0.4264 |
| Alsophilapodophylla | 0.5814* | 63.63 | 12.81 | 0.2344 | 1.061 | 44.15 | 0.4277 |

#### Cross-validation details

In addition to estimating missing values, PhyloPars performs cross-validation: it temporarily excludes each observation from the available information, in order to calculate its best estimate given al observations and the optimal phylogenetic covariances. This provides a detailed estimate of the error and bias one may expect for the estimated missing feature values. The distributions of estima calculated through cross-validation are shown below; these are compared with two null models (red and green curves) in order to assess the improvement resulting from the PhyloPars evolutionar

**Meanwd**

**Cross-validation error**

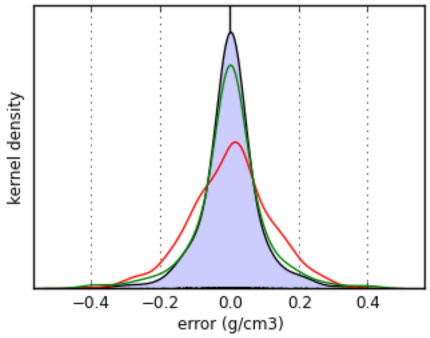


| | evolutionary model | mean model | nearest neighbor model |
|---|---|---|---|
| mean bias | **-0.000581 g/cm3** | 3.3e-16 g/cm3 | -0.000865 g/cm3 |
| mean error | **0.0496 g/cm3** | 0.0884 g/cm3 | 0.0591 g/cm3 |

**Meanleafarea**

**Cross-validation error**

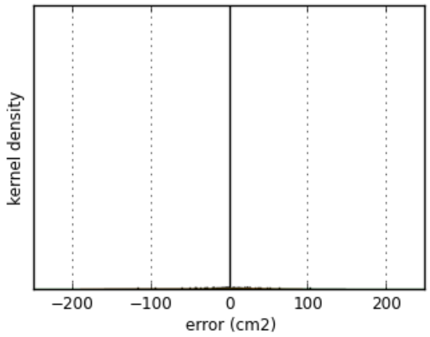


| | evolutionary model | mean model | nearest neighbor model |
|---|---|---|---|
| mean bias | **-0.579 cm2** | -1.23e-14 cm2 | -0.678 cm2 |
| mean error | **23.1 cm2** | 30.2 cm2 | 34.2 cm2 |

**Meansla**

**Cross-validation error**

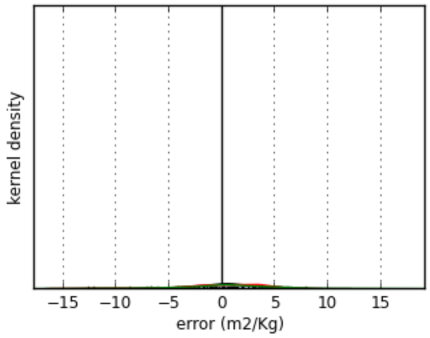


| | evolutionary model | mean model | nearest neighbor model |
|---|---|---|---|
| mean bias | **-0.0725 m2/Kg** | 2e-15 m2/Kg | 0.373 m2/Kg |
| mean error | **2.89 m2/Kg** | 2.88 m2/Kg | 3.7 m2/Kg |

**Meanleafthickness**

**Cross-validation error**

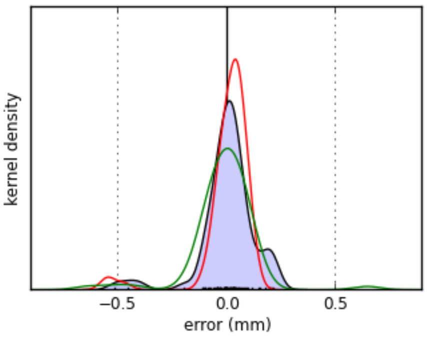


| | evolutionary model | mean model | nearest neighbor model |
|---|---|---|---|
| mean bias | **0.000495 mm** | 1.03e-16 mm | -0.0161 mm |
| mean error | **0.0801 mm** | 0.0678 mm | 0.0876 mm |

#### Meanrootavgdiam

**Cross-validation error**

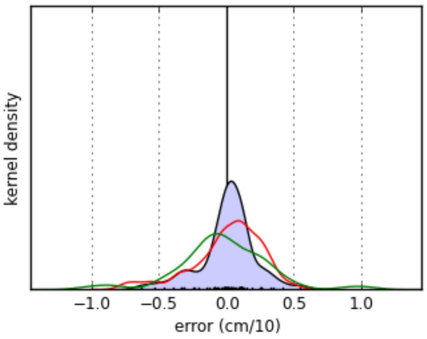


| | evolutionary model | mean model | nearest neighbor model |
|---|---|---|---|
| mean bias | **0.000635 cm/10** | -1.86e-16 cm/10 | -0.0301 cm/10 |
| mean error | **0.143 cm/10** | 0.2 cm/10 | 0.244 cm/10 |

#### Meanspecificrootlength

**Cross-validation error**

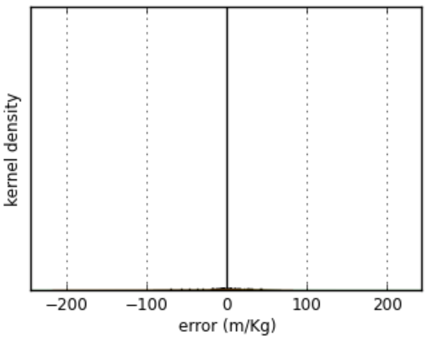


| | evolutionary model | mean model | nearest neighbor model |
|---|---|---|---|
| mean bias | **0.0936 m/Kg** | 4.7e-15 m/Kg | 1.44 m/Kg |
| mean error | **13.7 m/Kg** | 20.6 m/Kg | 27.2 m/Kg |

#### Meanroottd

**Cross-validation error**

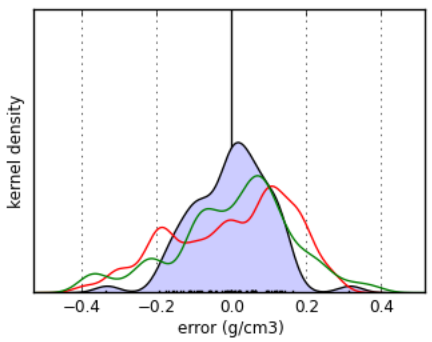


| | evolutionary model | mean model | nearest neighbor model |
|---|---|---|---|
| mean bias | **-0.00257 g/cm3** | 7.04e-18 g/cm3 | 0.00292 g/cm3 |
| mean error | **0.0809 g/cm3** | 0.135 g/cm3 | 0.125 g/cm3 |

**Meanbranchiness**

**Cross-validation error**

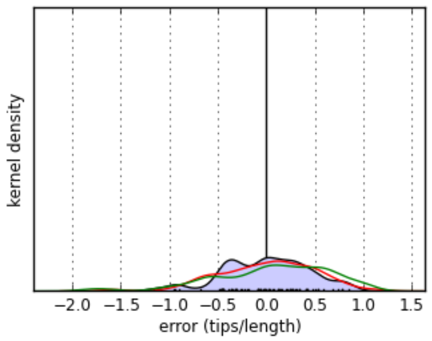


| | evolutionary model | mean model | nearest neighbor model |
|---|---|---|---|
| mean bias | **-0.00825 tips/length** | 3.97e-16 tips/length | 0.0463 tips/length |
| mean error | **0.349 tips/length** | 0.396 tips/length | 0.48 tips/length |

The PhyloPars tool is (c) Jorn Bruggeman 2019. If you are experiencing problems with this tool, please contact jbr [at] pml.ac.uk or bwbrandt [at] few.vu.nl.

### PhyloPars results Model uses traits for individuals only found in the primary forest portion of the transect (i.e. second half of the JFL transect)

#### Contents

Phylogenetic variability
Estimates for missing parameters
Cross-validation details

**Please cite:** Bruggeman J, Heringa J and Brandt BW. (2009) PhyloPars: estimation of missing parameter values using phylogeny. Nucleic Acids Research 37: W179-W184.

#### Phylogenetic variability

PhyloPars first estimates the parameters of the evolutionary model, i.e., the phylogenetic covariances. These are subsequently used to estimate missing feature values and to perform cross-valida

| feature | phylogenetic s.d. | | cross-validation results | | | |
|---|---|---|---|---|---|---|
| | | | error | | bias | |
| meanWD | 0.049 | g/cm3 | 0.05 | g/cm3 | -0.000541 | g/cm3 |
| meanLeafArea | 34.12 | cm2 | 32.9 | cm2 | -4.28 | cm2 |
| meanSLA | 1.935 | m2/Kg | 1.79 | m2/Kg | 0.0902 | m2/Kg |
| meanLeafThickness | 0.03004 | mm | 0.0291 | mm | 0.0041 | mm |
| meanRootAvgDiam | 0.1449 | cm/10 | 0.105 | cm/10 | -0.0123 | cm/10 |
| meanSpecificRootLength | 16.1 | m/Kg | 10.9 | m/Kg | -0.699 | m/Kg |
| meanRootTD | 0.07705 | g/cm3 | 0.0692 | g/cm3 | -0.000553 | g/cm3 |
| meanBranchiness | 0.3966 | tips/length | 0.455 | tips/length | 0.00984 | tips/length |

##### Phylogenetic correlations

| | meanWD | meanLeafArea | meanSLA | meanLeafThickness | meanRootAvgDiam | meanSpecificRootLength | meanRootTD | meanBr |
|---|---|---|---|---|---|---|---|---|
| **meanWD** | 1 | | | | | | | |
| **meanLeafArea** | 0.026 | 1 | | | | | | |
| **meanSLA** | -0.007 | -0.370 | 1 | | | | | |
| **meanLeafThickness** | 0.022 | 0.219 | -0.641 | 1 | | | | |
| **meanRootAvgDiam** | 0.016 | -0.021 | -0.198 | 0.212 | 1 | | | |
| **meanSpecificRootLength** | -0.246 | -0.324 | 0.390 | -0.121 | -0.599 | 1 | | |
| **meanRootTD** | 0.267 | 0.355 | -0.285 | -0.016 | -0.141 | -0.484 | 1 | |
| **meanBranchiness** | 0.106 | 0.028 | 0.027 | 0.067 | -0.225 | 0.223 | -0.113 | 1 |

##### Phylogenetic regression coefficients

Rows represent independent variables and columns dependent variables. A regression of feature *i* on *j* thus can be found at row *j*, column *i*.

| | meanWD | meanLeafArea | meanSLA | meanLeafThickness | meanRootAvgDiam | meanSpecificRootLength | meanRootTD | meanBr |
|---|---|---|---|---|---|---|---|---|
| **meanWD** | 1 | 17.802 | -0.278 | 0.014 | 0.047 | -80.694 | 0.421 | 0.857 |
| **meanLeafArea** | 0.000 | 1 | -0.021 | 0.000 | -0.000 | -0.153 | 0.001 | 0.000 |
| **meanSLA** | -0.000 | -6.522 | 1 | -0.010 | -0.015 | 3.244 | -0.011 | 0.006 |
| **meanLeafThickness** | 0.036 | 248.558 | -41.303 | 1 | 1.025 | -64.862 | -0.041 | 0.889 |
| **meanRootAvgDiam** | 0.005 | -4.834 | -2.650 | 0.044 | 1 | -66.516 | -0.075 | -0.616 |
| **meanSpecificRootLength** | -0.001 | -0.686 | 0.047 | -0.000 | -0.005 | 1 | -0.002 | 0.005 |
| **meanRootTD** | 0.170 | 157.030 | -7.164 | -0.006 | -0.265 | -101.192 | 1 | -0.580 |
| **meanBranchiness** | 0.013 | 2.421 | 0.131 | 0.005 | -0.082 | 9.047 | -0.022 | 1 |

#### Estimates for missing parameters

Using the optimal phylogenetic covariances listed above, PhyloPars estimates the values that were originally missing in the feature matrix. The table below lists all estimated values. If one or more have been provided for a value, this is denoted by a trailing asterisk symbol (*). You can click on an entry to view the contribution of individual observations to the estimate, and to retrieve details su standard deviation of the estimate. You can also download the complete table as a single text file.

| | meanWD (g/cm3) | meanLeafArea (cm2) | meanSLA (m2/Kg) | meanLeafThickness (mm) | meanRootAvgDiam (cm/10) | meanSpecificRootLength (m/Kg) | meanRootTD (g/cm3) | mean (tip |
|---|---|---|---|---|---|---|---|---|

| | | | | | | | |
|---|---|---|---|---|---|---|---|
| root | 0.5269 | 72.25 | 10.91 | 0.2253 | 1.213 | 34.29 | 0.4236 |
| fagales | 0.6168 | 85.95 | 10.24 | 0.1936 | 0.8817 | 45.04 | 0.5587 |
| fagaceae | 0.6190 | 71.00 | 9.663 | 0.1969 | 0.8495 | 45.62 | 0.5746 |
| lithocarpus | 0.6897 | 53.05 | 9.690 | 0.1793 | 0.9902 | 36.62 | 0.5879 |
| Lithocarpusamygdalifolius | 0.7580* | 51.03* | 9.911* | 0.1803* | 0.7312* | 49.81* | 0.5674* |
| Lithocarpusbacgiangensis | 0.6773* | 52.83 | 9.693 | 0.1791 | 0.9896 | 37.63 | 0.5826 |
| Lithocarpusbrachystachyus | 0.6773* | 52.83 | 9.693 | 0.1791 | 0.9896 | 37.63 | 0.5826 |
| Lithocarpuschiungchungensis | 0.6773* | 52.83 | 9.693 | 0.1791 | 0.9896 | 37.63 | 0.5826 |
| Lithocarpuscorneus | 0.8180* | 55.34 | 9.654 | 0.1811 | 0.9962 | 26.27 | 0.6418 |
| Lithocarpuselaeagnifolius | 0.6773* | 52.83 | 9.693 | 0.1791 | 0.9896 | 37.63 | 0.5826 |
| Lithocarpusfenestratus | 0.6773* | 61.76* | 7.953* | 0.2161* | 0.9090* | 40.94* | 0.5314* |
| Lithocarpusfenzelianus | 0.6773* | 38.31* | 8.436* | 0.1933* | 1.367* | 14.23* | 0.6691* |
| Lithocarpushancei | 0.6773* | 52.83 | 9.693 | 0.1791 | 0.9896 | 37.63 | 0.5826 |
| Lithocarpushandelianus | 0.7610* | 54.32 | 9.670 | 0.1803 | 0.9936 | 30.87 | 0.6178 |
| Lithocarpushowii | 0.6773* | 52.83 | 9.693 | 0.1791 | 0.9896 | 37.63 | 0.5826 |
| Lithocarpuslitseifolius | 0.6773* | 52.83 | 9.693 | 0.1791 | 0.9896 | 37.63 | 0.5826 |
| Lithocarpuslongipedicellatus | 0.5995* | 74.02* | 10.81* | 0.1640* | 0.7592* | 50.09* | 0.5846* |
| Lithocarpuspseudovestitus | 0.6773* | 29.85* | 11.21* | 0.1422* | 1.264* | 30.76* | 0.5564* |
| Lithocarpussp0502 | 0.6773* | 52.83 | 9.693 | 0.1791 | 0.9896 | 37.63 | 0.5826 |
| castanopsis | 0.5502 | 73.27 | 10.25 | 0.1676 | 0.7678 | 53.51 | 0.5399 |
| Castanopsiscarlesii | 0.4408* | 71.32 | 10.28 | 0.1661 | 0.7627 | 62.34 | 0.4939 |
| Castanopsisfissa | 0.4446* | 105.2* | 8.717* | 0.1563* | 0.7377* | 58.70* | 0.4834* |
| Castanopsishainanensis | 0.6473* | 75.00 | 10.22 | 0.1689 | 0.7724 | 45.67 | 0.5808 |
| Castanopsishystrix | 0.5650* | 73.53 | 10.24 | 0.1678 | 0.7685 | 52.31 | 0.5462 |
| Castanopsisjianfenglingensis | 0.5680* | 73.58 | 10.24 | 0.1678 | 0.7687 | 52.07 | 0.5474 |
| Castanopsisjucunda | 0.5680* | 73.58 | 10.24 | 0.1678 | 0.7687 | 52.07 | 0.5474 |
| Castanopsistonkinensis | 0.5680* | 46.47* | 11.99* | 0.1728* | 0.7266* | 57.17* | 0.5622* |
| quercus | 0.7003 | 67.46 | 9.734 | 0.1886 | 0.8814 | 38.45 | 0.6032 |
| Quercusacutissima | 0.7334* | 68.05 | 9.725 | 0.1891 | 0.8829 | 35.78 | 0.6172 |
| Cyclobalanopsis | 0.6076 | 58.13 | 9.610 | 0.2043 | 0.8187 | 48.11 | 0.5742 |
| Cyclobalanopsisbella | 0.6062* | 58.10 | 9.610 | 0.2043 | 0.8187 | 48.22 | 0.5736 |
| Cyclobalanopsisblakei | 0.6062* | 20.91* | 13.65* | 0.1627* | 0.7628* | 66.01* | 0.4938* |
| Cyclobalanopsisedithiae | 0.6062* | 129.8* | 5.783* | 0.2971* | 0.8802* | 36.22* | 0.6553* |
| Cyclobalanopsisfleuryi | 0.6062* | 101.5* | 7.120* | 0.2462* | 0.8163* | 43.38* | 0.6119* |
| Cyclobalanopsishui | 0.6062* | 24.17* | 10.46* | 0.1787* | 0.7358* | 56.91* | 0.4811* |
| Cyclobalanopsisneglecta | 0.6062* | 10.32* | 12.35* | 0.1557* | 0.8031* | 45.14* | 0.5617* |
| Cyclobalanopsispatelliformis | 0.6062* | 49.21* | 8.241* | 0.1928* | 0.8836* | 43.24* | 0.6422* |
| Cyclobalanopsisphanera | 0.6062* | 58.10 | 9.610 | 0.2043 | 0.8187 | 48.22 | 0.5736 |
| betulaceae | 0.5671 | 100.6 | 9.128 | 0.1908 | 0.8636 | 46.74 | 0.5628 |
| betula | 0.5567 | 100.4 | 9.130 | 0.1906 | 0.8631 | 47.58 | 0.5584 |
| Betulaalnoides | 0.5515* | 100.3 | 9.132 | 0.1906 | 0.8629 | 48.00 | 0.5562 |
| juglandaceae | 0.5280 | 145.9 | 7.462 | 0.1827 | 0.8625 | 44.52 | 0.5813 |
| engelhardia | 0.4916 | 175.9 | 6.354 | 0.1772 | 0.8613 | 43.88 | 0.5893 |
| Engelhardiahainanensis | 0.4879* | 175.9 | 6.355 | 0.1772 | 0.8612 | 44.17 | 0.5877 |
| Engelhardiaroxburghiana | 0.4879* | 191.2* | 5.796* | 0.1747* | 0.8614* | 42.38* | 0.5994* |
| Engelhardiaspicata | 0.4879* | 175.9 | 6.355 | 0.1772 | 0.8612 | 44.17 | 0.5877 |
| EngelhardiaspicatavarAceriflora | 0.4879* | 175.9 | 6.355 | 0.1772 | 0.8612 | 44.17 | 0.5877 |
| Engelhardiaunijuga | 0.4879* | 175.9 | 6.355 | 0.1772 | 0.8612 | 44.17 | 0.5877 |
| rosales | 0.6484 | 83.78 | 11.95 | 0.1923 | 0.9213 | 43.75 | 0.5420 |
| moraceae | 0.5627 | 70.28 | 14.86 | 0.1863 | 0.9479 | 51.78 | 0.4891 |
| broussonetia | 0.3365 | 90.51 | 14.95 | 0.1957 | 0.7856 | 82.06 | 0.4976 |
| Broussonetiapapyrifera | 0.2900* | 89.68 | 14.96 | 0.1950 | 0.7835 | 85.82 | 0.4780 |
| ficus | 0.3949 | 115.8 | 14.96 | 0.2089 | 0.6367 | 89.38 | 0.6257 |
| Ficusaltissima | 0.4730* | 117.2 | 14.93 | 0.2100 | 0.6404 | 83.08 | 0.6585 |
| Ficusauriculata | 0.4680* | 117.1 | 14.94 | 0.2099 | 0.6401 | 83.49 | 0.6564 |
| Ficusfistulosa | 0.3800* | 115.5 | 14.96 | 0.2087 | 0.6360 | 90.59 | 0.6194 |
| Ficusformosana | 0.4055* | 47.74* | 13.87* | 0.2120* | 0.6709* | 52.32* | 0.7383* |
| Ficusglaberrima | 0.4055* | 116.0 | 14.95 | 0.2091 | 0.6372 | 88.53 | 0.6302 |
| Ficusheteropleura | 0.4055* | 116.0 | 14.95 | 0.2091 | 0.6372 | 88.53 | 0.6302 |
| Ficushirta | 0.4055* | 287.4* | 14.04* | 0.1977* | 0.6322* | 60.22* | 0.7843* |
| Ficuslangkokensis | 0.4055* | 116.0 | 14.95 | 0.2091 | 0.6372 | 88.53 | 0.6302 |
| Ficusnervosa | 0.2800* | 113.8 | 14.99 | 0.2074 | 0.6314 | 98.66 | 0.5774 |
| Ficuspandurata | 0.4055* | 116.0 | 14.95 | 0.2091 | 0.6372 | 88.53 | 0.6302 |

| | | | | | | | |
|---|---|---|---|---|---|---|---|
| Ficuspubigera | 0.4055* | 116.0 | 14.95 | 0.2091 | 0.6372 | 88.53 | 0.6302 |
| Ficussimplicissima | 0.4055* | 116.0 | 14.95 | 0.2091 | 0.6372 | 88.53 | 0.6302 |
| Ficussubpisocarpa | 0.4055* | 116.0 | 14.95 | 0.2091 | 0.6372 | 88.53 | 0.6302 |
| Ficustinctoria | 0.4055* | 116.0 | 14.95 | 0.2091 | 0.6372 | 88.53 | 0.6302 |
| Ficustuphapensis | 0.4055* | 116.0 | 14.95 | 0.2091 | 0.6372 | 88.53 | 0.6302 |
| Ficusvariegata | 0.3270* | 114.6 | 14.97 | 0.2080 | 0.6336 | 94.86 | 0.5971 |
| Ficusvariolosa | 0.4055* | 116.0 | 14.95 | 0.2091 | 0.6372 | 88.53 | 0.6302 |
| Ficusvasculosa | 0.3000* | 23.04* | 16.99* | 0.2223* | 0.5279* | 167.6* | 0.3753* |
| antiaris | 0.3884 | 103.6 | 14.94 | 0.2026 | 0.7122 | 83.90 | 0.5712 |
| Antiaristoxicaria | 0.3831* | 103.5 | 14.95 | 0.2025 | 0.7120 | 84.32 | 0.5689 |
| artocarpus | 0.4959 | 40.05 | 16.01 | 0.1710 | 1.109 | 45.58 | 0.3507 |
| Artocarpusnitidus | 0.4800* | 39.77 | 16.01 | 0.1708 | 1.108 | 46.86 | 0.3440 |
| Artocarpusstyracifolius | 0.5077* | 25.74* | 16.57* | 0.1640* | 1.191* | 38.83* | 0.3005* |
| Artocarpustonkinensis | 0.4667* | 39.53 | 16.01 | 0.1706 | 1.107 | 47.94 | 0.3384 |
| streblus | 0.6898 | 72.54 | 14.82 | 0.1880 | 0.9538 | 41.52 | 0.5426 |
| Streblusilicifolius | 0.7321* | 73.30 | 14.81 | 0.1886 | 0.9558 | 38.11 | 0.5604 |
| Streblusindicus | 0.7321* | 73.30 | 14.81 | 0.1886 | 0.9558 | 38.11 | 0.5604 |
| Streblustaxoides | 0.7321* | 73.30 | 14.81 | 0.1886 | 0.9558 | 38.11 | 0.5604 |
| urticaceae | 0.4506 | 70.68 | 14.31 | 0.1857 | 0.9365 | 60.60 | 0.4453 |
| Oreocnide | 0.3900 | 69.60 | 14.33 | 0.1849 | 0.9337 | 65.49 | 0.4199 |
| Oreocnidetonkinensis | 0.3294* | 68.52 | 14.34 | 0.1841 | 0.9309 | 70.38 | 0.3944 |
| debregeasia | 0.3536 | 68.95 | 14.34 | 0.1844 | 0.9320 | 68.43 | 0.4046 |
| Debregeasiasquamata | 0.3294* | 68.52 | 14.34 | 0.1841 | 0.9309 | 70.38 | 0.3944 |
| cannabaceae | 0.6187 | 76.07 | 13.69 | 0.1890 | 0.9383 | 46.82 | 0.5194 |
| aphananthe | 0.6543 | 76.70 | 13.68 | 0.1895 | 0.9399 | 43.94 | 0.5344 |
| Aphananthecuspidata | 0.6900* | 77.34 | 13.67 | 0.1900 | 0.9416 | 41.06 | 0.5494 |
| celtis | 0.6608 | 76.82 | 13.68 | 0.1896 | 0.9402 | 43.42 | 0.5371 |
| Celtisphilippensis | 0.7030* | 77.57 | 13.66 | 0.1902 | 0.9422 | 40.01 | 0.5549 |
| gironniera | 0.5141 | 74.20 | 13.72 | 0.1876 | 0.9334 | 55.26 | 0.4754 |
| Gironnierasubaequalis | 0.4617* | 73.27 | 13.73 | 0.1869 | 0.9309 | 59.48 | 0.4534 |
| rhamnaceae | 0.7808 | 83.74 | 12.49 | 0.1931 | 0.9336 | 33.29 | 0.5944 |
| rhamnella | 0.7594 | 83.36 | 12.49 | 0.1929 | 0.9326 | 35.02 | 0.5853 |
| Rhamnellarubrinervis | 0.7421* | 83.05 | 12.50 | 0.1926 | 0.9318 | 36.42 | 0.5781 |
| ventilago | 0.9216 | 86.25 | 12.45 | 0.1951 | 0.9402 | 21.93 | 0.6535 |
| Ventilagoleiocarpa | 0.9800* | 87.29 | 12.43 | 0.1959 | 0.9429 | 17.22 | 0.6781 |
| rosaceae | 0.6498 | 83.80 | 11.95 | 0.1923 | 0.9214 | 43.64 | 0.5426 |
| photinia | 0.7217 | 85.08 | 11.93 | 0.1933 | 0.9247 | 37.84 | 0.5729 |
| Photiniabenthamiana | 0.7235* | 85.12 | 11.93 | 0.1933 | 0.9248 | 37.69 | 0.5736 |
| Photiniaprunifolia | 0.7235* | 85.12 | 11.93 | 0.1933 | 0.9248 | 37.69 | 0.5736 |
| eriobotrya | 0.7683 | 85.91 | 11.91 | 0.1939 | 0.9269 | 34.08 | 0.5925 |
| Eriobotryadeflexa | 0.7755* | 86.04 | 11.91 | 0.1940 | 0.9273 | 33.50 | 0.5955 |
| laurocerasus | 0.6522 | 83.85 | 11.95 | 0.1924 | 0.9215 | 43.45 | 0.5436 |
| Laurocerasusphaeosticta | 0.6481* | 83.77 | 11.95 | 0.1923 | 0.9213 | 43.78 | 0.5419 |
| pygeum | 0.6522 | 83.85 | 11.95 | 0.1924 | 0.9215 | 43.45 | 0.5436 |
| Pygeumtopengii | 0.6481* | 83.77 | 11.95 | 0.1923 | 0.9213 | 43.78 | 0.5419 |
| Rhaphiolepis | 0.6487 | 83.78 | 11.95 | 0.1923 | 0.9213 | 43.73 | 0.5421 |
| Rhaphiolepisferruginea | 0.6481* | 83.77 | 11.95 | 0.1923 | 0.9213 | 43.78 | 0.5419 |
| Rhaphiolepisindica | 0.6481* | 83.77 | 11.95 | 0.1923 | 0.9213 | 43.78 | 0.5419 |
| fabales | 0.6463 | 88.36 | 11.36 | 0.1938 | 0.9250 | 44.00 | 0.5346 |
| fabaceae | 0.6541 | 88.50 | 11.36 | 0.1939 | 0.9254 | 43.37 | 0.5379 |
| archidendron | 0.4954 | 85.67 | 11.40 | 0.1918 | 0.9180 | 56.18 | 0.4711 |
| Archidendronclypearia | 0.3233* | 82.61 | 11.45 | 0.1894 | 0.9099 | 70.06 | 0.3987 |
| Archidendronlucidum | 0.5814* | 87.20 | 11.38 | 0.1929 | 0.9220 | 49.24 | 0.5073 |
| Archidendronutile | 0.5814* | 87.20 | 11.38 | 0.1929 | 0.9220 | 49.24 | 0.5073 |
| albizia | 0.4780 | 85.36 | 11.41 | 0.1915 | 0.9172 | 57.58 | 0.4638 |
| Albiziachinensis | 0.3000* | 82.19 | 11.46 | 0.1891 | 0.9089 | 71.95 | 0.3889 |
| Albiziaodoratissima | 0.6385* | 88.22 | 11.36 | 0.1937 | 0.9247 | 44.63 | 0.5313 |
| peltophorum | 0.8257 | 91.55 | 11.31 | 0.1963 | 0.9334 | 29.53 | 0.6100 |
| Peltophorumtonkinense | 0.8486* | 91.96 | 11.30 | 0.1966 | 0.9345 | 27.68 | 0.6197 |
| sindora | 0.8410 | 91.83 | 11.31 | 0.1965 | 0.9341 | 28.29 | 0.6165 |
| Sindoraglabra | 0.8486* | 91.96 | 11.30 | 0.1966 | 0.9345 | 27.68 | 0.6197 |
| ormosia | 0.6027 | 87.58 | 11.37 | 0.1932 | 0.9230 | 47.52 | 0.5162 |
| Ormosiabalansae | 0.4505* | 84.87 | 11.41 | 0.1911 | 0.9159 | 59.80 | 0.4522 |
| Ormosiafordiana | 0.8486* | 91.96 | 11.30 | 0.1966 | 0.9345 | 27.68 | 0.6197 |

| | | | | | | | |
|---|---|---|---|---|---|---|---|
| Ormosiapinnata | 0.5683* | 86.97 | 11.38 | 0.1928 | 0.9214 | 50.29 | 0.5018 |
| Ormosiasemicastrata | 0.6213* | 87.91 | 11.37 | 0.1935 | 0.9239 | 46.02 | 0.5241 |
| Ormosiaxylocarpa | 0.5055* | 85.85 | 11.40 | 0.1919 | 0.9184 | 55.36 | 0.4754 |
| dalbergia | 0.6828 | 89.01 | 11.35 | 0.1943 | 0.9267 | 41.06 | 0.5499 |
| Dalbergiahainanensis | 0.6887* | 89.11 | 11.35 | 0.1944 | 0.9270 | 40.58 | 0.5524 |
| polygalaceae | 0.6705 | 88.79 | 11.35 | 0.1942 | 0.9262 | 42.05 | 0.5448 |
| xanthophyllum | 0.6786 | 88.93 | 11.35 | 0.1943 | 0.9265 | 41.40 | 0.5482 |
| Xanthophyllumhainanense | 0.6867* | 89.08 | 11.35 | 0.1944 | 0.9269 | 40.74 | 0.5516 |
| malpighiales | 0.6420 | 90.51 | 11.35 | 0.1943 | 0.9347 | 44.65 | 0.5228 |
| euphorbiaceae | 0.5468 | 88.81 | 11.38 | 0.1930 | 0.9303 | 52.33 | 0.4828 |
| mallotus | 0.4875 | 87.76 | 11.39 | 0.1922 | 0.9275 | 57.11 | 0.4579 |
| Mallotusanomalus | 0.5038* | 88.05 | 11.39 | 0.1925 | 0.9283 | 55.80 | 0.4647 |
| Mallotuspaniculatus | 0.3450* | 85.22 | 11.43 | 0.1903 | 0.9209 | 68.61 | 0.3979 |
| Mallotusphilippensis | 0.6033* | 89.82 | 11.36 | 0.1938 | 0.9329 | 47.77 | 0.5066 |
| Mallotusyunnanensis | 0.5038* | 88.05 | 11.39 | 0.1925 | 0.9283 | 55.80 | 0.4647 |
| macaranga | 0.4576 | 87.22 | 11.40 | 0.1918 | 0.9261 | 59.53 | 0.4453 |
| Macarangadenticulata | 0.4335* | 86.79 | 11.41 | 0.1915 | 0.9250 | 61.47 | 0.4352 |
| hancea | 0.5215 | 88.36 | 11.39 | 0.1927 | 0.9291 | 54.37 | 0.4721 |
| Hanceahookeriana | 0.5431* | 88.74 | 11.38 | 0.1930 | 0.9301 | 52.63 | 0.4813 |
| cleidion | 0.5048 | 88.06 | 11.39 | 0.1925 | 0.9283 | 55.72 | 0.4651 |
| Cleidionbrevipetiolatum | 0.5167* | 88.27 | 11.39 | 0.1926 | 0.9289 | 54.76 | 0.4701 |
| koilodepas | 0.5339 | 88.58 | 11.38 | 0.1929 | 0.9297 | 53.37 | 0.4774 |
| Koilodepashainanense | 0.5431* | 88.74 | 11.38 | 0.1930 | 0.9301 | 52.63 | 0.4813 |
| claoxylon | 0.3813 | 85.86 | 11.42 | 0.1908 | 0.9226 | 65.68 | 0.4132 |
| Claoxylonindicum | 0.3550* | 85.40 | 11.43 | 0.1904 | 0.9213 | 67.81 | 0.4021 |
| alchornea | 0.4163 | 86.49 | 11.41 | 0.1913 | 0.9242 | 62.86 | 0.4279 |
| Alchornearugosa | 0.4084* | 86.35 | 11.42 | 0.1912 | 0.9238 | 63.49 | 0.4246 |
| croton | 0.5191 | 88.32 | 11.39 | 0.1927 | 0.9290 | 54.56 | 0.4712 |
| Crotoncascarilloides | 0.5104* | 88.16 | 11.39 | 0.1926 | 0.9286 | 55.27 | 0.4675 |
| Crotonlaevigatus | 0.5300* | 88.51 | 11.38 | 0.1928 | 0.9295 | 53.68 | 0.4757 |
| ostodes | 0.3628 | 85.54 | 11.43 | 0.1905 | 0.9217 | 67.17 | 0.4054 |
| Ostodespaniculata | 0.3390* | 85.11 | 11.44 | 0.1902 | 0.9206 | 69.10 | 0.3954 |
| suregada | 0.5980 | 89.72 | 11.36 | 0.1937 | 0.9327 | 48.20 | 0.5043 |
| Suregadamultiflora | 0.6470* | 90.59 | 11.35 | 0.1944 | 0.9350 | 44.24 | 0.5250 |
| triadica | 0.5409 | 88.71 | 11.38 | 0.1930 | 0.9300 | 52.80 | 0.4803 |
| Triadicacochinchinensis | 0.5431* | 88.74 | 11.38 | 0.1930 | 0.9301 | 52.63 | 0.4813 |
| endospermum | 0.3836 | 85.91 | 11.42 | 0.1908 | 0.9227 | 65.50 | 0.4142 |
| Endospermumchinense | 0.3475* | 85.26 | 11.43 | 0.1903 | 0.9210 | 68.41 | 0.3990 |
| Trevia | 0.5450 | 88.78 | 11.38 | 0.1930 | 0.9302 | 52.48 | 0.4820 |
| Trevianudiflora | 0.5431* | 88.74 | 11.38 | 0.1930 | 0.9301 | 52.63 | 0.4813 |
| Euphorbiaceaesp9 | 0.5431* | 88.74 | 11.38 | 0.1930 | 0.9301 | 52.63 | 0.4813 |
| Lasiococca | 0.5450 | 88.78 | 11.38 | 0.1930 | 0.9302 | 52.48 | 0.4820 |
| Lasiococcacomberi | 0.5431* | 88.74 | 11.38 | 0.1930 | 0.9301 | 52.63 | 0.4813 |
| Epiprinus | 0.5450 | 88.78 | 11.38 | 0.1930 | 0.9302 | 52.48 | 0.4820 |
| Epiprinussiletianus | 0.5431* | 88.74 | 11.38 | 0.1930 | 0.9301 | 52.63 | 0.4813 |
| Euphorbiaceaesp11 | 0.5431* | 88.74 | 11.38 | 0.1930 | 0.9301 | 52.63 | 0.4813 |
| Euphorbiaceaesp3 | 0.5431* | 88.74 | 11.38 | 0.1930 | 0.9301 | 52.63 | 0.4813 |
| Euphorbiaceaesp4 | 0.5431* | 88.74 | 11.38 | 0.1930 | 0.9301 | 52.63 | 0.4813 |
| Euphorbiaceaesp5 | 0.5431* | 88.74 | 11.38 | 0.1930 | 0.9301 | 52.63 | 0.4813 |
| phyllanthaceae | 0.6010 | 89.78 | 11.36 | 0.1938 | 0.9328 | 47.95 | 0.5056 |
| antidesma | 0.6284 | 90.26 | 11.36 | 0.1942 | 0.9341 | 45.74 | 0.5171 |
| Antidesmamaclurei | 0.5900* | 89.58 | 11.37 | 0.1936 | 0.9323 | 48.84 | 0.5010 |
| Antidesmamontanum | 0.5900* | 89.58 | 11.37 | 0.1936 | 0.9323 | 48.84 | 0.5010 |
| Antidesmasp1 | 0.6575* | 90.78 | 11.35 | 0.1946 | 0.9355 | 43.39 | 0.5294 |
| Antidesmasp2 | 0.6575* | 90.78 | 11.35 | 0.1946 | 0.9355 | 43.39 | 0.5294 |
| Antidesmahainanense | 0.6575* | 90.78 | 11.35 | 0.1946 | 0.9355 | 43.39 | 0.5294 |
| aporosa | 0.4232 | 86.61 | 11.41 | 0.1914 | 0.9245 | 62.30 | 0.4308 |
| Aporosadioica | 0.3700* | 85.66 | 11.43 | 0.1906 | 0.9220 | 66.60 | 0.4084 |
| baccaurea | 0.5097 | 88.15 | 11.39 | 0.1925 | 0.9286 | 55.32 | 0.4672 |
| Baccaurearamiflora | 0.5431* | 88.74 | 11.38 | 0.1930 | 0.9301 | 52.63 | 0.4813 |
| bischofia | 0.5772 | 89.35 | 11.37 | 0.1935 | 0.9317 | 49.88 | 0.4956 |
| Bischofiapolycarpa | 0.5698* | 89.22 | 11.37 | 0.1934 | 0.9314 | 50.47 | 0.4925 |
| breynia | 0.6215 | 90.14 | 11.36 | 0.1941 | 0.9338 | 46.30 | 0.5142 |
| Breyniafruticosa | 0.6315* | 90.32 | 11.36 | 0.1942 | 0.9342 | 45.49 | 0.5184 |

| | | | | | | | |
|---|---|---|---|---|---|---|---|
| Breyniarostrata | 0.6315* | 90.32 | 11.36 | 0.1942 | 0.9342 | 45.49 | 0.5184 |
| glochidion | 0.5486 | 88.84 | 11.38 | 0.1931 | 0.9304 | 52.18 | 0.4836 |
| Glochidioncoccineum | 0.5576* | 89.00 | 11.38 | 0.1932 | 0.9308 | 51.46 | 0.4873 |
| Glochidionhirsutum | 0.5576* | 89.00 | 11.38 | 0.1932 | 0.9308 | 51.46 | 0.4873 |
| Glochidionsphaerogynum | 0.5576* | 89.00 | 11.38 | 0.1932 | 0.9308 | 51.46 | 0.4873 |
| Glochidiontriandrum | 0.5576* | 89.00 | 11.38 | 0.1932 | 0.9308 | 51.46 | 0.4873 |
| Glochidionzeylanicum | 0.4800* | 87.62 | 11.40 | 0.1921 | 0.9272 | 57.72 | 0.4547 |
| phyllanthus | 0.6033 | 89.82 | 11.36 | 0.1938 | 0.9329 | 47.77 | 0.5066 |
| Phyllanthuspachyphyllus | 0.6126* | 89.98 | 11.36 | 0.1939 | 0.9334 | 47.02 | 0.5105 |
| bridelia | 0.6132 | 89.99 | 11.36 | 0.1940 | 0.9334 | 46.97 | 0.5107 |
| Brideliabalansae | 0.6052* | 89.85 | 11.36 | 0.1938 | 0.9330 | 47.61 | 0.5074 |
| cleistanthus | 0.6404 | 90.48 | 11.35 | 0.1943 | 0.9347 | 44.78 | 0.5222 |
| Cleistanthusconcinnus | 0.6518* | 90.68 | 11.35 | 0.1945 | 0.9352 | 43.86 | 0.5270 |
| leptopus | 0.6282 | 90.26 | 11.36 | 0.1942 | 0.9341 | 45.76 | 0.5170 |
| Leptopushainanensis | 0.6315* | 90.32 | 11.36 | 0.1942 | 0.9342 | 45.49 | 0.5184 |
| actephila | 0.6163 | 90.05 | 11.36 | 0.1940 | 0.9335 | 46.72 | 0.5120 |
| Actephilamerrilliana | 0.6315* | 90.32 | 11.36 | 0.1942 | 0.9342 | 45.49 | 0.5184 |
| ixonanthaceae | 0.6116 | 89.96 | 11.36 | 0.1939 | 0.9333 | 47.10 | 0.5101 |
| ixonanthes | 0.6191 | 90.10 | 11.36 | 0.1940 | 0.9337 | 46.50 | 0.5132 |
| Ixonanthesreticulata | 0.6265* | 90.23 | 11.36 | 0.1941 | 0.9340 | 45.90 | 0.5163 |
| salicaceae | 0.6564 | 90.76 | 11.35 | 0.1945 | 0.9354 | 43.48 | 0.5289 |
| casearia | 0.6394 | 90.46 | 11.35 | 0.1943 | 0.9346 | 44.85 | 0.5218 |
| Caseariamembranacea | 0.6500* | 90.65 | 11.35 | 0.1945 | 0.9351 | 44.00 | 0.5262 |
| Caseariavelutina | 0.6246* | 90.20 | 11.36 | 0.1941 | 0.9339 | 46.05 | 0.5155 |
| homalium | 0.6849 | 91.27 | 11.34 | 0.1949 | 0.9367 | 41.19 | 0.5409 |
| Homaliummollissimum | 0.6957* | 91.46 | 11.34 | 0.1951 | 0.9372 | 40.31 | 0.5455 |
| Homaliumpaniculiflorum | 0.6957* | 91.46 | 11.34 | 0.1951 | 0.9372 | 40.31 | 0.5455 |
| Homaliumphanerophlebium | 0.6400* | 90.47 | 11.35 | 0.1943 | 0.9346 | 44.81 | 0.5220 |
| Homaliumstenophyllum | 0.6957* | 91.46 | 11.34 | 0.1951 | 0.9372 | 40.31 | 0.5455 |
| scolopia | 0.8052 | 93.41 | 11.31 | 0.1966 | 0.9424 | 31.48 | 0.5915 |
| Scolopiasaeva | 0.8300* | 93.85 | 11.30 | 0.1969 | 0.9435 | 29.48 | 0.6019 |
| flacourtia | 0.7518 | 92.46 | 11.32 | 0.1958 | 0.9399 | 35.78 | 0.5691 |
| Flacourtiarukam | 0.7500* | 92.43 | 11.32 | 0.1958 | 0.9398 | 35.93 | 0.5683 |
| achariaceae | 0.6310 | 90.31 | 11.36 | 0.1942 | 0.9342 | 45.53 | 0.5182 |
| hydnocarpus | 0.6311 | 90.31 | 11.36 | 0.1942 | 0.9342 | 45.52 | 0.5183 |
| Hydnocarpushainanensis | 0.6313* | 90.31 | 11.36 | 0.1942 | 0.9342 | 45.51 | 0.5183 |
| hypericaceae | 0.6854 | 91.28 | 11.34 | 0.1949 | 0.9368 | 41.15 | 0.5411 |
| cratoxylum | 0.6915 | 91.39 | 11.34 | 0.1950 | 0.9370 | 40.65 | 0.5437 |
| Cratoxylumcochinchinense | 0.6700* | 91.00 | 11.34 | 0.1947 | 0.9360 | 42.39 | 0.5346 |
| Cratoxylumformosum | 0.7150* | 91.80 | 11.33 | 0.1953 | 0.9381 | 38.76 | 0.5536 |
| calophyllaceae | 0.6797 | 91.18 | 11.34 | 0.1949 | 0.9365 | 41.61 | 0.5387 |
| calophyllum | 0.6700 | 91.00 | 11.34 | 0.1947 | 0.9360 | 42.39 | 0.5346 |
| Calophyllummembranaceum | 0.6692* | 90.99 | 11.34 | 0.1947 | 0.9360 | 42.45 | 0.5343 |
| Calophyllumsp1 | 0.6692* | 90.99 | 11.34 | 0.1947 | 0.9360 | 42.45 | 0.5343 |
| clusiaceae | 0.6745 | 91.08 | 11.34 | 0.1948 | 0.9362 | 42.02 | 0.5365 |
| garcinia | 0.6831 | 91.24 | 11.34 | 0.1949 | 0.9367 | 41.33 | 0.5402 |
| Garciniamultiflora | 0.7416* | 92.28 | 11.32 | 0.1957 | 0.9394 | 36.61 | 0.5648 |
| Garciniaoblongifolia | 0.6268* | 90.23 | 11.36 | 0.1941 | 0.9340 | 45.87 | 0.5165 |
| Clusiaceaesp1 | 0.6692* | 90.99 | 11.34 | 0.1947 | 0.9360 | 42.45 | 0.5343 |
| ochnaceae | 0.7051 | 91.63 | 11.33 | 0.1952 | 0.9377 | 39.56 | 0.5494 |
| ochna | 0.7375 | 92.21 | 11.33 | 0.1957 | 0.9392 | 36.94 | 0.5630 |
| Ochnaintegerrima | 0.7440* | 92.32 | 11.32 | 0.1957 | 0.9395 | 36.42 | 0.5658 |
| Campylospermum | 0.7201 | 91.90 | 11.33 | 0.1954 | 0.9384 | 38.35 | 0.5557 |
| Campylospermumserratum | 0.7351* | 92.16 | 11.33 | 0.1956 | 0.9391 | 37.13 | 0.5620 |
| rhizophoraceae | 0.6974 | 91.49 | 11.34 | 0.1951 | 0.9373 | 40.18 | 0.5461 |
| carallia | 0.6737 | 91.07 | 11.34 | 0.1948 | 0.9362 | 42.09 | 0.5362 |
| Caralliabrachiata | 0.6659* | 90.93 | 11.35 | 0.1947 | 0.9358 | 42.72 | 0.5329 |
| erythroxylaceae | 0.7334 | 92.13 | 11.33 | 0.1956 | 0.9390 | 37.27 | 0.5613 |
| erythroxylum | 0.7615 | 92.63 | 11.32 | 0.1960 | 0.9403 | 35.00 | 0.5731 |
| Erythroxylumsinense | 0.7896* | 93.13 | 11.31 | 0.1964 | 0.9416 | 32.74 | 0.5849 |
| pandaceae | 0.6324 | 90.33 | 11.35 | 0.1942 | 0.9343 | 45.42 | 0.5188 |
| microdesmis | 0.6162 | 90.05 | 11.36 | 0.1940 | 0.9335 | 46.73 | 0.5120 |
| Microdesmiscaseariifolia | 0.6000* | 89.76 | 11.36 | 0.1938 | 0.9328 | 48.04 | 0.5052 |
| putranjivaceae | 0.6839 | 91.25 | 11.34 | 0.1949 | 0.9367 | 41.27 | 0.5405 |

| | | | | | | | |
|---|---|---|---|---|---|---|---|
| drypetes | 0.6978 | 91.50 | 11.34 | 0.1951 | 0.9373 | 40.14 | 0.5463 |
| Drypetescumingii | 0.6945* | 91.44 | 11.34 | 0.1951 | 0.9372 | 40.41 | 0.5449 |
| Drypeteshainanensis | 0.7183* | 91.86 | 11.33 | 0.1954 | 0.9383 | 38.49 | 0.5550 |
| Drypetesperreticulata | 0.6945* | 91.44 | 11.34 | 0.1951 | 0.9372 | 40.41 | 0.5449 |
| oxalidales | 0.6122 | 89.97 | 11.36 | 0.1939 | 0.9333 | 47.05 | 0.5103 |
| elaeocarpaceae | 0.5533 | 88.93 | 11.38 | 0.1931 | 0.9306 | 51.81 | 0.4855 |
| elaeocarpus | 0.4924 | 87.84 | 11.39 | 0.1923 | 0.9277 | 56.72 | 0.4599 |
| Elaeocarpusangustifolius | 0.4029* | 86.25 | 11.42 | 0.1911 | 0.9236 | 63.94 | 0.4223 |
| Elaeocarpusdubius | 0.5240* | 88.40 | 11.38 | 0.1927 | 0.9292 | 54.17 | 0.4732 |
| Elaeocarpusglabripetalus | 0.5032* | 88.04 | 11.39 | 0.1925 | 0.9282 | 55.84 | 0.4645 |
| Elaeocarpushowii | 0.5032* | 88.04 | 11.39 | 0.1925 | 0.9282 | 55.84 | 0.4645 |
| Elaeocarpusjaponicus | 0.5032* | 88.04 | 11.39 | 0.1925 | 0.9282 | 55.84 | 0.4645 |
| Elaeocarpuslimitaneus | 0.5032* | 88.04 | 11.39 | 0.1925 | 0.9282 | 55.84 | 0.4645 |
| Elaeocarpusnitentifolius | 0.5032* | 88.04 | 11.39 | 0.1925 | 0.9282 | 55.84 | 0.4645 |
| Elaeocarpuspoilanei | 0.5032* | 88.04 | 11.39 | 0.1925 | 0.9282 | 55.84 | 0.4645 |
| Elaeocarpussylvestris | 0.4750* | 87.53 | 11.40 | 0.1921 | 0.9269 | 58.12 | 0.4526 |
| sloanea | 0.5458 | 88.79 | 11.38 | 0.1930 | 0.9302 | 52.41 | 0.4824 |
| Sloaneaintegrifolia | 0.6095* | 89.93 | 11.36 | 0.1939 | 0.9332 | 47.27 | 0.5092 |
| Sloaneasinensis | 0.4785* | 87.59 | 11.40 | 0.1921 | 0.9271 | 57.84 | 0.4541 |
| connaraceae | 0.5881 | 89.55 | 11.37 | 0.1936 | 0.9322 | 48.99 | 0.5002 |
| ellipanthus | 0.5847 | 89.49 | 11.37 | 0.1936 | 0.9321 | 49.27 | 0.4988 |
| Ellipanthusglabrifolius | 0.5814* | 89.43 | 11.37 | 0.1935 | 0.9319 | 49.54 | 0.4973 |
| celastrales | 0.6422 | 90.51 | 11.35 | 0.1944 | 0.9347 | 44.63 | 0.5229 |
| celastraceae | 0.6503 | 90.65 | 11.35 | 0.1945 | 0.9351 | 43.98 | 0.5263 |
| Celastraceaesp2 | 0.6797* | 91.18 | 11.34 | 0.1949 | 0.9365 | 41.61 | 0.5387 |
| Celastraceaesp3 | 0.6797* | 91.18 | 11.34 | 0.1949 | 0.9365 | 41.61 | 0.5387 |
| salacia | 0.7478 | 92.39 | 11.32 | 0.1958 | 0.9397 | 36.11 | 0.5674 |
| Salaciachinensis | 0.7600* | 92.61 | 11.32 | 0.1960 | 0.9402 | 35.12 | 0.5725 |
| euonymus | 0.5874 | 89.53 | 11.37 | 0.1936 | 0.9322 | 49.05 | 0.4999 |
| Euonymusgibber | 0.5665* | 89.16 | 11.37 | 0.1933 | 0.9312 | 50.74 | 0.4911 |
| Euonymuslaxiflorus | 0.5665* | 89.16 | 11.37 | 0.1933 | 0.9312 | 50.74 | 0.4911 |
| Euonymusnitidus | 0.5665* | 89.16 | 11.37 | 0.1933 | 0.9312 | 50.74 | 0.4911 |
| myrtales | 0.6326 | 100.4 | 11.31 | 0.1917 | 0.9464 | 48.04 | 0.4889 |
| melastomataceae | 0.6854 | 101.4 | 11.30 | 0.1924 | 0.9488 | 43.77 | 0.5111 |
| blastus | 0.6081 | 99.97 | 11.32 | 0.1914 | 0.9452 | 50.02 | 0.4786 |
| Blastuscochinchinensis | 0.6094* | 100.0 | 11.32 | 0.1914 | 0.9453 | 49.91 | 0.4792 |
| melastoma | 0.4452 | 97.07 | 11.37 | 0.1891 | 0.9376 | 63.16 | 0.4101 |
| Melastomamalabathricum | 0.4400* | 96.98 | 11.37 | 0.1891 | 0.9374 | 63.58 | 0.4079 |
| Melastomapenicillatum | 0.4400* | 96.98 | 11.37 | 0.1891 | 0.9374 | 63.58 | 0.4079 |
| Melastomasanguineum | 0.4400* | 96.98 | 11.37 | 0.1891 | 0.9374 | 63.58 | 0.4079 |
| medinilla | 0.6142 | 100.1 | 11.32 | 0.1915 | 0.9455 | 49.52 | 0.4812 |
| Medinillaassamica | 0.6094* | 100.0 | 11.32 | 0.1914 | 0.9453 | 49.91 | 0.4792 |
| Allomorphia | 0.6474 | 100.7 | 11.31 | 0.1919 | 0.9471 | 46.84 | 0.4951 |
| Allomorphiabalansae | 0.6094* | 100.0 | 11.32 | 0.1914 | 0.9453 | 49.91 | 0.4792 |
| memecylon | 0.7510 | 102.5 | 11.28 | 0.1933 | 0.9519 | 38.49 | 0.5387 |
| Memecylonhainanense | 0.7728* | 102.9 | 11.28 | 0.1936 | 0.9529 | 36.72 | 0.5479 |
| Memecylonligustrifolium | 0.7728* | 102.9 | 11.28 | 0.1936 | 0.9529 | 36.72 | 0.5479 |
| Memecylonnigrescens | 0.7728* | 102.9 | 11.28 | 0.1936 | 0.9529 | 36.72 | 0.5479 |
| myrtaceae | 0.6869 | 101.4 | 11.30 | 0.1924 | 0.9489 | 43.65 | 0.5118 |
| rhodamnia | 0.8244 | 103.8 | 11.26 | 0.1943 | 0.9553 | 32.56 | 0.5696 |
| Rhodamniadumetorum | 0.8787* | 104.8 | 11.25 | 0.1951 | 0.9579 | 28.18 | 0.5924 |
| rhodomyrtus | 0.7069 | 101.7 | 11.29 | 0.1927 | 0.9498 | 42.04 | 0.5202 |
| Rhodomyrtustomentosa | 0.6797* | 101.2 | 11.30 | 0.1923 | 0.9486 | 44.24 | 0.5087 |
| decaspermum | 0.7256 | 102.1 | 11.29 | 0.1930 | 0.9507 | 40.53 | 0.5280 |
| Decaspermummontanum | 0.7170* | 101.9 | 11.29 | 0.1929 | 0.9503 | 41.23 | 0.5244 |
| syzygium | 0.6693 | 101.1 | 11.30 | 0.1922 | 0.9481 | 45.08 | 0.5043 |
| Syzygiumacuminatissimum | 0.6644* | 101.0 | 11.31 | 0.1921 | 0.9479 | 45.47 | 0.5023 |
| Syzygiumaraiocladum | 0.7360* | 102.3 | 11.29 | 0.1931 | 0.9512 | 39.69 | 0.5324 |
| Syzygiumbullockii | 0.6644* | 101.0 | 11.31 | 0.1921 | 0.9479 | 45.47 | 0.5023 |
| Syzygiumbuxifolium | 0.6644* | 101.0 | 11.31 | 0.1921 | 0.9479 | 45.47 | 0.5023 |
| Syzygiumchampionii | 0.6644* | 101.0 | 11.31 | 0.1921 | 0.9479 | 45.47 | 0.5023 |
| Syzygiumchunianum | 0.6644* | 101.0 | 11.31 | 0.1921 | 0.9479 | 45.47 | 0.5023 |
| Syzygiumclaviflorum | 0.6235* | 100.2 | 11.32 | 0.1916 | 0.9459 | 48.77 | 0.4851 |
| Syzygiumcumini | 0.6727* | 101.1 | 11.30 | 0.1923 | 0.9482 | 44.80 | 0.5058 |

| | | | | | | | |
|---|---|---|---|---|---|---|---|
| Syzygiumglobiflorum | 0.6644* | 101.0 | 11.31 | 0.1921 | 0.9479 | 45.47 | 0.5023 |
| Syzygiumhancei | 0.6644* | 101.0 | 11.31 | 0.1921 | 0.9479 | 45.47 | 0.5023 |
| Syzygiumjambos | 0.7000* | 101.6 | 11.30 | 0.1926 | 0.9495 | 42.60 | 0.5173 |
| Syzygiumjienfunicum | 0.6644* | 101.0 | 11.31 | 0.1921 | 0.9479 | 45.47 | 0.5023 |
| Syzygiumlevinei | 0.6644* | 101.0 | 11.31 | 0.1921 | 0.9479 | 45.47 | 0.5023 |
| Syzygiumodoratum | 0.6644* | 101.0 | 11.31 | 0.1921 | 0.9479 | 45.47 | 0.5023 |
| Syzygiumrehderianum | 0.6644* | 101.0 | 11.31 | 0.1921 | 0.9479 | 45.47 | 0.5023 |
| Syzygiumrysopodum | 0.6644* | 101.0 | 11.31 | 0.1921 | 0.9479 | 45.47 | 0.5023 |
| Syzygiumsp | 0.6792* | 101.2 | 11.30 | 0.1923 | 0.9485 | 44.28 | 0.5085 |
| Syzygiumsterrophyllum | 0.6644* | 101.0 | 11.31 | 0.1921 | 0.9479 | 45.47 | 0.5023 |
| Syzygiumtephrodes | 0.6644* | 101.0 | 11.31 | 0.1921 | 0.9479 | 45.47 | 0.5023 |
| Syzygiumtsoongii | 0.6644* | 101.0 | 11.31 | 0.1921 | 0.9479 | 45.47 | 0.5023 |
| sapindales | 0.5606 | 121.6 | 11.24 | 0.1760 | 0.9121 | 61.99 | 0.4181 |
| meliaceae | 0.5769 | 124.9 | 12.42 | 0.1786 | 0.8910 | 67.91 | 0.3687 |
| aphanamixis | 0.5849 | 125.0 | 12.42 | 0.1788 | 0.8913 | 67.26 | 0.3721 |
| Aphanamixispolystachya | 0.5765* | 124.9 | 12.42 | 0.1786 | 0.8909 | 67.94 | 0.3685 |
| aglaia | 0.5994 | 125.3 | 12.41 | 0.1790 | 0.8920 | 66.09 | 0.3782 |
| Aglaiaelaeagnoidea | 0.6300* | 125.8 | 12.41 | 0.1794 | 0.8934 | 63.62 | 0.3910 |
| Aglaiaspectabilis | 0.5750* | 124.9 | 12.42 | 0.1786 | 0.8909 | 68.06 | 0.3679 |
| dysoxylum | 0.5722 | 124.8 | 12.42 | 0.1786 | 0.8907 | 68.29 | 0.3667 |
| Dysoxylumgotadhora | 0.5974* | 125.3 | 12.41 | 0.1789 | 0.8919 | 66.25 | 0.3773 |
| Dysoxylummollissimum | 0.5189* | 123.9 | 12.44 | 0.1779 | 0.8882 | 72.59 | 0.3443 |
| walsura | 0.8326 | 129.5 | 12.35 | 0.1821 | 0.9029 | 47.27 | 0.4763 |
| Walsurapinnata | 0.8680* | 130.1 | 12.34 | 0.1826 | 0.9046 | 44.42 | 0.4911 |
| Walsurarobusta | 0.8680* | 130.1 | 12.34 | 0.1826 | 0.9046 | 44.42 | 0.4911 |
| melia | 0.4818 | 123.2 | 12.45 | 0.1773 | 0.8865 | 75.58 | 0.3287 |
| Meliaazedarach | 0.4378* | 122.4 | 12.46 | 0.1767 | 0.8845 | 79.13 | 0.3102 |
| Heynea | 0.5856 | 125.1 | 12.42 | 0.1788 | 0.8914 | 67.21 | 0.3724 |
| Heyneatrijuga | 0.5944* | 125.2 | 12.42 | 0.1789 | 0.8918 | 66.50 | 0.3760 |
| Meliaceaesp1 | 0.5944* | 125.2 | 12.42 | 0.1789 | 0.8918 | 66.50 | 0.3760 |
| simaroubaceae | 0.5340 | 124.1 | 12.43 | 0.1781 | 0.8890 | 71.37 | 0.3507 |
| brucea | 0.4532 | 122.7 | 12.45 | 0.1770 | 0.8852 | 77.89 | 0.3167 |
| Bruceamollis | 0.4330* | 122.3 | 12.46 | 0.1767 | 0.8842 | 79.52 | 0.3082 |
| rutaceae | 0.5497 | 117.5 | 13.17 | 0.1832 | 0.8847 | 72.15 | 0.3325 |
| murraya | 0.6589 | 102.4 | 13.04 | 0.1761 | 0.9022 | 57.79 | 0.3954 |
| Murrayaalata | 0.7537* | 104.1 | 13.01 | 0.1774 | 0.9066 | 50.15 | 0.4353 |
| glycosmis | 0.4703 | 99.06 | 13.09 | 0.1735 | 0.8934 | 73.01 | 0.3161 |
| Glycosmissp1 | 0.4390* | 98.50 | 13.10 | 0.1731 | 0.8919 | 75.54 | 0.3029 |
| Glycosmiscochinchinensis | 0.4390* | 98.50 | 13.10 | 0.1731 | 0.8919 | 75.54 | 0.3029 |
| Glycosmiscraibii | 0.4390* | 98.50 | 13.10 | 0.1731 | 0.8919 | 75.54 | 0.3029 |
| clausena | 0.5231 | 100.0 | 13.08 | 0.1742 | 0.8958 | 68.75 | 0.3383 |
| Clausenaexcavata | 0.4820* | 99.26 | 13.09 | 0.1737 | 0.8939 | 72.07 | 0.3210 |
| micromelum | 0.6121 | 101.6 | 13.05 | 0.1755 | 0.9000 | 61.57 | 0.3757 |
| Micromelumfalcatum | 0.6600* | 102.4 | 13.04 | 0.1761 | 0.9022 | 57.70 | 0.3959 |
| tetradium | 0.2887 | 78.81 | 13.05 | 0.1624 | 0.8973 | 82.12 | 0.2568 |
| Tetradiumglabrifolium | 0.2320* | 77.80 | 13.07 | 0.1616 | 0.8946 | 86.70 | 0.2329 |
| acronychia | 0.4687 | 71.57 | 12.56 | 0.2070 | 0.9067 | 65.16 | 0.3476 |
| Acronychiapedunculata | 0.4587* | 74.57* | 11.26* | 0.2209* | 0.9497* | 57.00* | 0.3440* |
| melicope | 0.4921 | 58.24 | 16.99 | 0.1688 | 0.7558 | 94.06 | 0.3591 |
| Melicopechunii | 0.4953* | 50.92* | 18.83* | 0.1583* | 0.6910* | 106.6* | 0.3634* |
| zanthoxylum | 0.5667 | 79.45 | 12.39 | 0.1584 | 0.9379 | 53.04 | 0.3788 |
| Zanthoxylumavicennae | 0.6177* | 76.05* | 11.79* | 0.1512* | 0.9680* | 42.27* | 0.4053* |
| Maclurodendron | 0.5499 | 119.1 | 13.95 | 0.1925 | 0.8736 | 76.95 | 0.2992 |
| Maclurodendronoligophlebium | 0.5500* | 120.6* | 14.73* | 0.2018* | 0.8625* | 81.76* | 0.2659* |
| sapindaceae | 0.5646 | 137.2 | 11.44 | 0.1734 | 0.8989 | 65.93 | 0.4020 |
| mischocarpus | 0.7465 | 190.6 | 9.133 | 0.1747 | 0.9391 | 42.87 | 0.6013 |
| Mischocarpushainanensis | 0.7307* | 190.3 | 9.137 | 0.1745 | 0.9383 | 44.15 | 0.5946 |
| Mischocarpuspentapetalus | 0.7307* | 190.3 | 9.137 | 0.1745 | 0.9383 | 44.15 | 0.5946 |
| Mischocarpussundaicus | 0.7800* | 191.2 | 9.123 | 0.1751 | 0.9406 | 40.17 | 0.6153 |
| nephelium | 0.7724 | 219.0 | 7.874 | 0.1744 | 0.9579 | 36.14 | 0.6803 |
| Nepheliumtopengii | 0.7782* | 224.6* | 7.622* | 0.1743* | 0.9617* | 34.74* | 0.6964* |
| dimocarpus | 0.7344 | 207.1 | 8.385 | 0.1741 | 0.9491 | 41.06 | 0.6371 |
| Dimocarpuslongan | 0.7000* | 206.5 | 8.395 | 0.1737 | 0.9475 | 43.84 | 0.6226 |

| | | | | | | | |
|---|---|---|---|---|---|---|---|
| litchi | 0.8115 | 208.5 | 8.364 | 0.1752 | 0.9527 | 34.84 | 0.6695 |
| Litchichinensis | 0.8541* | 209.3 | 8.352 | 0.1758 | 0.9547 | 31.40 | 0.6874 |
| lepisanthes | 0.6471 | 194.4 | 8.910 | 0.1732 | 0.9380 | 49.96 | 0.5731 |
| Lepisanthesrubiginosa | 0.6300* | 194.1 | 8.915 | 0.1730 | 0.9372 | 51.35 | 0.5659 |
| amesiodendron | 0.8166 | 175.1 | 9.864 | 0.1760 | 0.9318 | 40.01 | 0.5898 |
| Amesiodendronchinense | 0.8345* | 175.4 | 9.859 | 0.1763 | 0.9326 | 38.56 | 0.5973 |
| paranephelium | 0.8037 | 174.9 | 9.868 | 0.1759 | 0.9312 | 41.05 | 0.5844 |
| Paranepheliumhainanense | 0.8267* | 175.3 | 9.861 | 0.1762 | 0.9323 | 39.20 | 0.5941 |
| acer | 0.4879 | 147.0 | 10.96 | 0.1721 | 0.9023 | 70.26 | 0.3970 |
| Acerfabri | 0.5145* | 147.4 | 10.95 | 0.1724 | 0.9036 | 68.10 | 0.4082 |
| Acerlaurinum | 0.4300* | 145.9 | 10.97 | 0.1713 | 0.8996 | 74.93 | 0.3727 |
| anacardiaceae | 0.5240 | 147.5 | 10.10 | 0.1552 | 0.8597 | 74.76 | 0.3887 |
| toxicodendron | 0.5620 | 160.0 | 9.316 | 0.1928 | 0.6579 | 106.8 | 0.4426 |
| Toxicodendronvernicifluum | 0.5662* | 161.3* | 9.229* | 0.1970* | 0.6355* | 110.4* | 0.4486* |
| choerospondias | 0.4932 | 147.0 | 10.11 | 0.1548 | 0.8582 | 77.25 | 0.3758 |
| Choerospondiasaxillaris | 0.4870* | 146.8 | 10.12 | 0.1547 | 0.8579 | 77.74 | 0.3732 |
| burseraceae | 0.5159 | 151.7 | 9.770 | 0.1383 | 0.8982 | 68.52 | 0.3780 |
| canarium | 0.4860 | 168.2 | 8.510 | 0.09993 | 0.9458 | 61.99 | 0.3559 |
| Canariumalbum | 0.4170* | 172.7* | 8.106* | 0.08633* | 0.9589* | 64.57* | 0.3237* |
| Canariumpimela | 0.5450* | 169.3 | 8.493 | 0.1007 | 0.9485 | 57.22 | 0.3807 |
| malvales | 0.5699 | 116.2 | 11.26 | 0.1798 | 0.9202 | 59.20 | 0.4322 |
| thymelaeaceae | 0.5296 | 115.5 | 11.28 | 0.1792 | 0.9184 | 62.45 | 0.4152 |
| aquilaria | 0.4134 | 113.4 | 11.31 | 0.1777 | 0.9129 | 71.83 | 0.3663 |
| Aquilariasinensis | 0.3660* | 112.5 | 11.32 | 0.1770 | 0.9107 | 75.66 | 0.3464 |
| wikstroemia | 0.5262 | 115.4 | 11.28 | 0.1792 | 0.9182 | 62.73 | 0.4138 |
| Wikstroemiahainanensis | 0.5294* | 115.5 | 11.28 | 0.1792 | 0.9183 | 62.47 | 0.4151 |
| Wikstroemiaindica | 0.5294* | 115.5 | 11.28 | 0.1792 | 0.9183 | 62.47 | 0.4151 |
| Wikstroemianutans | 0.5294* | 115.5 | 11.28 | 0.1792 | 0.9183 | 62.47 | 0.4151 |
| Wikstroemiapachyrachis | 0.5294* | 115.5 | 11.28 | 0.1792 | 0.9183 | 62.47 | 0.4151 |
| malvaceae | 0.5740 | 116.2 | 11.26 | 0.1798 | 0.9204 | 58.87 | 0.4339 |
| reevesia | 0.4716 | 114.4 | 11.29 | 0.1785 | 0.9156 | 67.14 | 0.3908 |
| Reevesialancifolia | 0.5075* | 115.1 | 11.28 | 0.1789 | 0.9173 | 64.24 | 0.4059 |
| Reevesiathyrsoidea | 0.4350* | 113.8 | 11.30 | 0.1780 | 0.9139 | 70.09 | 0.3754 |
| sterculia | 0.4567 | 114.2 | 11.30 | 0.1782 | 0.9150 | 68.34 | 0.3846 |
| Sterculiahainanensis | 0.4048* | 113.2 | 11.31 | 0.1775 | 0.9125 | 72.52 | 0.3627 |
| Sterculialanceolata | 0.5000* | 114.9 | 11.28 | 0.1788 | 0.9170 | 64.84 | 0.4027 |
| pterospermum | 0.4925 | 114.8 | 11.29 | 0.1787 | 0.9166 | 65.45 | 0.3996 |
| Pterospermumheterophyllum | 0.4460* | 114.0 | 11.30 | 0.1781 | 0.9144 | 69.20 | 0.3800 |
| Pterospermumlanceifolium | 0.5146* | 115.2 | 11.28 | 0.1790 | 0.9177 | 63.67 | 0.4089 |
| Pterospermumxiaoye | 0.5146* | 115.2 | 11.28 | 0.1790 | 0.9177 | 63.67 | 0.4089 |
| microcos | 0.4888 | 114.7 | 11.29 | 0.1787 | 0.9164 | 65.75 | 0.3980 |
| Microcoschungii | 0.4819* | 114.6 | 11.29 | 0.1786 | 0.9161 | 66.31 | 0.3951 |
| dipterocarpaceae | 0.7336 | 119.1 | 11.22 | 0.1820 | 0.9279 | 45.99 | 0.5010 |
| vatica | 0.7577 | 119.5 | 11.21 | 0.1824 | 0.9290 | 44.05 | 0.5111 |
| Vaticamangachapoi | 0.7500* | 119.4 | 11.21 | 0.1823 | 0.9286 | 44.67 | 0.5079 |
| hopea | 0.8353 | 120.9 | 11.19 | 0.1834 | 0.9326 | 37.79 | 0.5438 |
| Hopeahainanensis | 0.8900* | 121.9 | 11.18 | 0.1842 | 0.9352 | 33.37 | 0.5668 |
| brassicales | 0.5866 | 116.5 | 11.26 | 0.1800 | 0.9210 | 57.86 | 0.4392 |
| capparaceae | 0.6725 | 118.0 | 11.24 | 0.1812 | 0.9250 | 50.92 | 0.4753 |
| Capparaceaesp1 | 0.6832* | 118.2 | 11.23 | 0.1813 | 0.9255 | 50.06 | 0.4798 |
| crossosomatales | 0.5384 | 104.4 | 11.32 | 0.1867 | 0.9342 | 57.67 | 0.4392 |
| staphyleaceae | 0.4807 | 103.3 | 11.33 | 0.1859 | 0.9315 | 62.33 | 0.4149 |
| turpinia | 0.4229 | 102.3 | 11.35 | 0.1852 | 0.9288 | 67.00 | 0.3906 |
| Turpiniamontana | 0.3940* | 101.8 | 11.36 | 0.1848 | 0.9275 | 69.33 | 0.3784 |
| saxifragales | 0.6081 | 87.54 | 11.37 | 0.2036 | 0.9882 | 44.51 | 0.4891 |
| hamamelidaceae | 0.5996 | 87.39 | 11.37 | 0.2035 | 0.9878 | 45.19 | 0.4856 |
| eustigma | 0.6330 | 87.98 | 11.36 | 0.2039 | 0.9894 | 42.49 | 0.4996 |
| Eustigmaoblongifolium | 0.6372* | 88.06 | 11.36 | 0.2040 | 0.9896 | 42.16 | 0.5014 |
| exbucklandia | 0.5715 | 86.89 | 11.38 | 0.2031 | 0.9865 | 47.46 | 0.4737 |
| Exbucklandiatonkinensis | 0.5480* | 86.47 | 11.39 | 0.2028 | 0.9854 | 49.36 | 0.4639 |
| chunia | 0.6161 | 87.68 | 11.37 | 0.2037 | 0.9886 | 43.86 | 0.4925 |
| Chuniabucklandioides | 0.6372* | 88.06 | 11.36 | 0.2040 | 0.9896 | 42.16 | 0.5014 |
| daphniphyllaceae | 0.5520 | 86.54 | 11.38 | 0.2028 | 0.9856 | 49.03 | 0.4656 |
| daphniphyllum | 0.5292 | 86.13 | 11.39 | 0.2025 | 0.9846 | 50.87 | 0.4560 |

| | | | | | | | |
|---|---|---|---|---|---|---|---|
| Daphniphyllumcalycinum | 0.5063* | 85.73 | 11.40 | 0.2022 | 0.9835 | 52.72 | 0.4463 |
| altingiaceae | 0.6460 | 88.21 | 11.36 | 0.2041 | 0.9900 | 41.45 | 0.5051 |
| altingia | 0.6731 | 88.70 | 11.35 | 0.2045 | 0.9913 | 39.26 | 0.5165 |
| Altingiaobovata | 0.7002* | 89.18 | 11.34 | 0.2049 | 0.9925 | 37.07 | 0.5279 |
| iteaceae | 0.5875 | 87.17 | 11.37 | 0.2033 | 0.9873 | 46.17 | 0.4805 |
| itea | 0.5834 | 87.10 | 11.38 | 0.2033 | 0.9871 | 46.50 | 0.4788 |
| Iteamacrophylla | 0.5814* | 87.06 | 11.38 | 0.2032 | 0.9870 | 46.66 | 0.4779 |
| Iteaxiaoye | 0.5814* | 87.06 | 11.38 | 0.2032 | 0.9870 | 46.66 | 0.4779 |
| saxifragaceae | 0.5848 | 87.12 | 11.38 | 0.2033 | 0.9872 | 46.39 | 0.4793 |
| Saxifragaceaesp1 | 0.5814* | 87.06 | 11.38 | 0.2032 | 0.9870 | 46.66 | 0.4779 |
| dilleniales | 0.6104 | 84.18 | 11.38 | 0.2079 | 1.006 | 42.58 | 0.4903 |
| dilleniaceae | 0.6094 | 84.16 | 11.38 | 0.2079 | 1.006 | 42.67 | 0.4899 |
| dillenia | 0.6063 | 84.10 | 11.38 | 0.2078 | 1.006 | 42.91 | 0.4886 |
| Dilleniapentagyna | 0.5953* | 83.91 | 11.39 | 0.2077 | 1.005 | 43.80 | 0.4840 |
| Dilleniaturbinata | 0.6162* | 84.28 | 11.38 | 0.2080 | 1.006 | 42.11 | 0.4928 |
| gentianales | 0.6145 | 68.22 | 11.77 | 0.2247 | 0.9914 | 40.14 | 0.5131 |
| rubiaceae | 0.6438 | 68.74 | 11.77 | 0.2251 | 0.9928 | 37.77 | 0.5255 |
| Benkara | 0.6406 | 68.69 | 11.77 | 0.2250 | 0.9927 | 38.03 | 0.5241 |
| Benkarahainanensis | 0.6374* | 68.63 | 11.77 | 0.2250 | 0.9925 | 38.29 | 0.5228 |
| antirhea | 0.6404 | 68.68 | 11.77 | 0.2250 | 0.9927 | 38.05 | 0.5240 |
| Antirheachinensis | 0.6374* | 68.63 | 11.77 | 0.2250 | 0.9925 | 38.29 | 0.5228 |
| pertusadina | 0.6847 | 69.47 | 11.75 | 0.2256 | 0.9947 | 34.47 | 0.5427 |
| Pertusadinametcalfii | 0.6800* | 69.39 | 11.76 | 0.2256 | 0.9945 | 34.85 | 0.5407 |
| adina | 0.7215 | 70.13 | 11.74 | 0.2261 | 0.9964 | 31.51 | 0.5581 |
| Adinarubella | 0.7351* | 70.37 | 11.74 | 0.2263 | 0.9971 | 30.41 | 0.5639 |
| nauclea | 0.6423 | 68.72 | 11.77 | 0.2251 | 0.9927 | 37.90 | 0.5248 |
| naucleaofficinalis | 0.6374* | 68.63 | 11.77 | 0.2250 | 0.9925 | 38.29 | 0.5228 |
| diplospora | 0.7015 | 69.77 | 11.75 | 0.2259 | 0.9955 | 33.12 | 0.5497 |
| Diplosporadubia | 0.7000* | 69.75 | 11.75 | 0.2259 | 0.9954 | 33.24 | 0.5491 |
| catunaregam | 0.6913 | 69.59 | 11.75 | 0.2257 | 0.9950 | 33.94 | 0.5454 |
| Catunaregamspinosa | 0.6880* | 69.53 | 11.75 | 0.2257 | 0.9949 | 34.21 | 0.5441 |
| gardenia | 0.6722 | 69.25 | 11.76 | 0.2255 | 0.9941 | 35.48 | 0.5374 |
| Gardeniahainanensis | 0.6681* | 69.18 | 11.76 | 0.2254 | 0.9939 | 35.81 | 0.5357 |
| Gardeniasootepensis | 0.6681* | 69.18 | 11.76 | 0.2254 | 0.9939 | 35.81 | 0.5357 |
| aidia | 0.7404 | 70.46 | 11.74 | 0.2264 | 0.9973 | 29.98 | 0.5661 |
| Aidiacanthioides | 0.7527* | 70.68 | 11.74 | 0.2266 | 0.9979 | 28.99 | 0.5713 |
| Aidiapycnantha | 0.7527* | 70.68 | 11.74 | 0.2266 | 0.9979 | 28.99 | 0.5713 |
| pavetta | 0.6447 | 68.76 | 11.77 | 0.2251 | 0.9929 | 37.70 | 0.5259 |
| Pavettaarenosa | 0.6374* | 68.63 | 11.77 | 0.2250 | 0.9925 | 38.29 | 0.5228 |
| Pavettahongkongensis | 0.6374* | 68.63 | 11.77 | 0.2250 | 0.9925 | 38.29 | 0.5228 |
| tarenna | 0.6783 | 69.36 | 11.76 | 0.2256 | 0.9944 | 34.98 | 0.5400 |
| Tarennaattenuata | 0.6797* | 69.38 | 11.76 | 0.2256 | 0.9945 | 34.87 | 0.5406 |
| Tarennalancilimba | 0.6797* | 69.38 | 11.76 | 0.2256 | 0.9945 | 34.87 | 0.5406 |
| Tarennatsangii | 0.6797* | 69.38 | 11.76 | 0.2256 | 0.9945 | 34.87 | 0.5406 |
| ixora | 0.7755 | 71.09 | 11.73 | 0.2269 | 0.9990 | 27.15 | 0.5809 |
| Ixoranienkui | 0.7929* | 71.40 | 11.72 | 0.2271 | 0.9998 | 25.74 | 0.5882 |
| canthium | 0.6477 | 68.81 | 11.77 | 0.2251 | 0.9930 | 37.46 | 0.5271 |
| Canthiumhorridum | 0.6367* | 68.62 | 11.77 | 0.2250 | 0.9925 | 38.35 | 0.5225 |
| Canthiumsimile | 0.6367* | 68.62 | 11.77 | 0.2250 | 0.9925 | 38.35 | 0.5225 |
| psydrax | 0.7505 | 70.64 | 11.74 | 0.2265 | 0.9978 | 29.16 | 0.5703 |
| Psydraxdicocca | 0.7627* | 70.86 | 11.73 | 0.2267 | 0.9984 | 28.17 | 0.5755 |
| Tarennoidea | 0.6406 | 68.69 | 11.77 | 0.2250 | 0.9927 | 38.03 | 0.5241 |
| Tarennoideawallichii | 0.6374* | 68.63 | 11.77 | 0.2250 | 0.9925 | 38.29 | 0.5228 |
| hedyotis | 0.6367 | 68.62 | 11.77 | 0.2250 | 0.9925 | 38.35 | 0.5225 |
| Hedyotiscathayana | 0.6374* | 68.63 | 11.77 | 0.2250 | 0.9925 | 38.29 | 0.5228 |
| saprosma | 0.6372 | 68.63 | 11.77 | 0.2250 | 0.9925 | 38.30 | 0.5227 |
| Saprosmacrassipes | 0.6374* | 68.63 | 11.77 | 0.2250 | 0.9925 | 38.29 | 0.5228 |
| Saprosmahainanensis | 0.6374* | 68.63 | 11.77 | 0.2250 | 0.9925 | 38.29 | 0.5228 |
| Saprosmamerrillii | 0.6374* | 68.63 | 11.77 | 0.2250 | 0.9925 | 38.29 | 0.5228 |
| Saprosmayueyan | 0.6374* | 68.63 | 11.77 | 0.2250 | 0.9925 | 38.29 | 0.5228 |
| chassalia | 0.6309 | 68.51 | 11.77 | 0.2249 | 0.9922 | 38.82 | 0.5200 |
| Chassaliacurviflora | 0.6374* | 68.63 | 11.77 | 0.2250 | 0.9925 | 38.29 | 0.5228 |
| psychotria | 0.5730 | 67.48 | 11.79 | 0.2241 | 0.9895 | 43.48 | 0.4957 |
| Psychotriaasiatica | 0.5636* | 67.32 | 11.79 | 0.2240 | 0.9891 | 44.24 | 0.4917 |

| | | | | | | | |
|---|---|---|---|---|---|---|---|
| Psychotriastraminea | 0.5636* | 67.32 | 11.79 | 0.2240 | 0.9891 | 44.24 | 0.4917 |
| prismatomeris | 0.6304 | 68.51 | 11.77 | 0.2249 | 0.9922 | 38.85 | 0.5198 |
| Prismatomeristetrandra | 0.6374* | 68.63 | 11.77 | 0.2250 | 0.9925 | 38.29 | 0.5228 |
| lasianthus | 0.6373 | 68.63 | 11.77 | 0.2250 | 0.9925 | 38.30 | 0.5227 |
| Lasianthuschevalieri | 0.6374* | 68.63 | 11.77 | 0.2250 | 0.9925 | 38.29 | 0.5228 |
| Lasianthuscurtisii | 0.6374* | 68.63 | 11.77 | 0.2250 | 0.9925 | 38.29 | 0.5228 |
| Lasianthushirsutus | 0.6374* | 68.63 | 11.77 | 0.2250 | 0.9925 | 38.29 | 0.5228 |
| Lasianthusjaponicus | 0.6374* | 68.63 | 11.77 | 0.2250 | 0.9925 | 38.29 | 0.5228 |
| Lasianthuslancifolius | 0.6374* | 68.63 | 11.77 | 0.2250 | 0.9925 | 38.29 | 0.5228 |
| Lasianthusrhinocerotis | 0.6374* | 68.63 | 11.77 | 0.2250 | 0.9925 | 38.29 | 0.5228 |
| Lasianthustrichophlebus | 0.6374* | 68.63 | 11.77 | 0.2250 | 0.9925 | 38.29 | 0.5228 |
| Caelospermum | 0.6406 | 68.69 | 11.77 | 0.2250 | 0.9927 | 38.03 | 0.5241 |
| Caelospermumtruncatum | 0.6374* | 68.63 | 11.77 | 0.2250 | 0.9925 | 38.29 | 0.5228 |
| Wendlandia | 0.6764 | 69.33 | 11.76 | 0.2255 | 0.9943 | 35.14 | 0.5392 |
| Wendlandiamerrilliana | 0.7337* | 70.34 | 11.74 | 0.2263 | 0.9970 | 30.52 | 0.5633 |
| Wendlandiauvariifolia | 0.6518* | 68.89 | 11.76 | 0.2252 | 0.9932 | 37.12 | 0.5288 |
| apocynaceae | 0.6109 | 68.16 | 11.78 | 0.2246 | 0.9913 | 40.43 | 0.5116 |
| rauvolfia | 0.5001 | 66.19 | 11.81 | 0.2231 | 0.9861 | 49.37 | 0.4650 |
| Rauvolfiaverticillata | 0.4863* | 65.94 | 11.81 | 0.2229 | 0.9855 | 50.48 | 0.4592 |
| kopsia | 0.5740 | 67.50 | 11.79 | 0.2241 | 0.9896 | 43.41 | 0.4961 |
| Kopsiaarborea | 0.5929* | 67.84 | 11.78 | 0.2244 | 0.9904 | 41.88 | 0.5040 |
| wrightia | 0.3411 | 63.36 | 11.85 | 0.2210 | 0.9787 | 62.20 | 0.3982 |
| Wrightialaevis | 0.3120* | 62.84 | 11.86 | 0.2206 | 0.9773 | 64.55 | 0.3859 |
| tabernaemontana | 0.5587 | 67.23 | 11.79 | 0.2239 | 0.9888 | 44.64 | 0.4897 |
| Tabernaemontanabovina | 0.5656* | 67.35 | 11.79 | 0.2240 | 0.9892 | 44.08 | 0.4926 |
| Tabernaemontanabufalina | 0.5656* | 67.35 | 11.79 | 0.2240 | 0.9892 | 44.08 | 0.4926 |
| alstonia | 0.5095 | 66.35 | 11.80 | 0.2233 | 0.9865 | 48.61 | 0.4690 |
| Alstoniarostrata | 0.4640* | 65.54 | 11.82 | 0.2226 | 0.9844 | 52.28 | 0.4498 |
| Hunteria | 0.6654 | 69.13 | 11.76 | 0.2254 | 0.9938 | 36.03 | 0.5346 |
| Hunteriazeylanica | 0.7200* | 70.10 | 11.75 | 0.2261 | 0.9964 | 31.62 | 0.5575 |
| lamiales | 0.6055 | 68.06 | 11.78 | 0.2246 | 0.9910 | 40.86 | 0.5094 |
| bignoniaceae | 0.5817 | 67.64 | 11.78 | 0.2242 | 0.9899 | 42.78 | 0.4994 |
| radermachera | 0.5246 | 66.62 | 11.80 | 0.2235 | 0.9872 | 47.39 | 0.4753 |
| Radermacherasinica | 0.6255* | 68.42 | 11.77 | 0.2248 | 0.9920 | 39.25 | 0.5178 |
| Radermacherafrondosa | 0.4592* | 65.46 | 11.82 | 0.2226 | 0.9842 | 52.66 | 0.4478 |
| Radermacherahainanensis | 0.4592* | 65.46 | 11.82 | 0.2226 | 0.9842 | 52.66 | 0.4478 |
| markhamia | 0.6451 | 68.77 | 11.77 | 0.2251 | 0.9929 | 37.67 | 0.5260 |
| Markhamiastipulata | 0.6755* | 69.31 | 11.76 | 0.2255 | 0.9943 | 35.21 | 0.5388 |
| lamiaceae | 0.5300 | 66.72 | 11.80 | 0.2235 | 0.9875 | 46.95 | 0.4776 |
| gmelina | 0.5624 | 67.30 | 11.79 | 0.2240 | 0.9890 | 44.34 | 0.4912 |
| Gmelinahainanensis | 0.5947* | 67.87 | 11.78 | 0.2244 | 0.9905 | 41.74 | 0.5048 |
| Tsoongia | 0.5533 | 67.13 | 11.79 | 0.2239 | 0.9886 | 45.07 | 0.4874 |
| Tsoongiaaxillariflora | 0.5766* | 67.55 | 11.78 | 0.2242 | 0.9897 | 43.20 | 0.4972 |
| callicarpa | 0.4100 | 64.58 | 11.83 | 0.2219 | 0.9819 | 56.64 | 0.4271 |
| Callicarpabrevipes | 0.3500* | 63.51 | 11.85 | 0.2211 | 0.9791 | 61.48 | 0.4019 |
| Callicarpasp2 | 0.3500* | 63.51 | 11.85 | 0.2211 | 0.9791 | 61.48 | 0.4019 |
| vitex | 0.6354 | 68.59 | 11.77 | 0.2250 | 0.9924 | 38.45 | 0.5219 |
| Vitexpierreana | 0.8545* | 72.50 | 11.71 | 0.2280 | 1.003 | 20.77 | 0.6141 |
| Vitexquinata | 0.4513* | 65.32 | 11.82 | 0.2225 | 0.9838 | 53.30 | 0.4445 |
| clerodendrum | 0.5702 | 67.43 | 11.79 | 0.2241 | 0.9894 | 43.71 | 0.4945 |
| Clerodendrumhainanense | 0.5735* | 67.49 | 11.79 | 0.2241 | 0.9895 | 43.44 | 0.4959 |
| Clerodendrumkwangtungense | 0.5735* | 67.49 | 11.79 | 0.2241 | 0.9895 | 43.44 | 0.4959 |
| verbenaceae | 0.6258 | 68.42 | 11.77 | 0.2248 | 0.9920 | 39.23 | 0.5179 |
| Verbenaceaesp | 0.6426* | 68.72 | 11.77 | 0.2251 | 0.9928 | 37.87 | 0.5250 |
| oleaceae | 0.6711 | 69.23 | 11.76 | 0.2255 | 0.9941 | 35.57 | 0.5369 |
| olea | 0.6935 | 69.63 | 11.75 | 0.2258 | 0.9951 | 33.76 | 0.5464 |
| Oleabrachiata | 0.6300* | 68.50 | 11.77 | 0.2249 | 0.9922 | 38.89 | 0.5197 |
| Oleaneriifolia | 0.7431* | 70.51 | 11.74 | 0.2264 | 0.9974 | 29.76 | 0.5672 |
| Oleaparvilimba | 0.7431* | 70.51 | 11.74 | 0.2264 | 0.9974 | 29.76 | 0.5672 |
| Oleatsoongii | 0.6300* | 68.50 | 11.77 | 0.2249 | 0.9922 | 38.89 | 0.5197 |
| osmanthus | 0.8323 | 72.10 | 11.71 | 0.2277 | 1.002 | 22.56 | 0.6047 |
| Osmanthusdidymopetalus | 0.8415* | 72.26 | 11.71 | 0.2278 | 1.002 | 21.82 | 0.6086 |
| Osmanthushainanensis | 0.8415* | 72.26 | 11.71 | 0.2278 | 1.002 | 21.82 | 0.6086 |
| Osmanthusmarginatus | 0.8415* | 72.26 | 11.71 | 0.2278 | 1.002 | 21.82 | 0.6086 |

| | | | | | | | |
|---|---|---|---|---|---|---|---|
| Osmanthusmatsumuranus | 0.8415* | 72.26 | 11.71 | 0.2278 | 1.002 | 21.82 | 0.6086 |
| chionanthus | 0.7094 | 69.91 | 11.75 | 0.2260 | 0.9959 | 32.48 | 0.5531 |
| Chionanthusbrachythyrsus | 0.6805* | 69.40 | 11.76 | 0.2256 | 0.9945 | 34.81 | 0.5409 |
| Chionanthusramiflorus | 0.7530* | 70.69 | 11.74 | 0.2266 | 0.9979 | 28.96 | 0.5714 |
| boraginales | 0.5693 | 67.42 | 11.79 | 0.2241 | 0.9893 | 43.78 | 0.4941 |
| boraginaceae | 0.5336 | 66.78 | 11.80 | 0.2236 | 0.9877 | 46.67 | 0.4791 |
| ehretia | 0.5140 | 66.43 | 11.80 | 0.2233 | 0.9868 | 48.24 | 0.4709 |
| Ehretialongiflora | 0.5195* | 66.53 | 11.80 | 0.2234 | 0.9870 | 47.81 | 0.4732 |
| cordia | 0.4745 | 65.73 | 11.81 | 0.2228 | 0.9849 | 51.43 | 0.4543 |
| Cordiadichotoma | 0.4512* | 65.32 | 11.82 | 0.2225 | 0.9838 | 53.31 | 0.4445 |
| garryales | 0.5291 | 66.70 | 11.80 | 0.2235 | 0.9875 | 47.03 | 0.4772 |
| icacinaceae | 0.5004 | 66.19 | 11.81 | 0.2231 | 0.9861 | 49.34 | 0.4651 |
| apodytes | 0.5735 | 67.49 | 11.79 | 0.2241 | 0.9895 | 43.45 | 0.4959 |
| Apodytesdimidiata | 0.6100* | 68.14 | 11.78 | 0.2246 | 0.9912 | 40.50 | 0.5113 |
| Platea | 0.4351 | 65.03 | 11.82 | 0.2222 | 0.9831 | 54.61 | 0.4377 |
| Platealatifolia | 0.3400* | 63.34 | 11.85 | 0.2209 | 0.9786 | 62.29 | 0.3977 |
| Plateaparvifolia | 0.4650* | 65.56 | 11.82 | 0.2226 | 0.9845 | 52.20 | 0.4503 |
| apiales | 0.5389 | 66.88 | 11.80 | 0.2237 | 0.9879 | 46.24 | 0.4813 |
| pittosporaceae | 0.5381 | 66.86 | 11.80 | 0.2236 | 0.9879 | 46.30 | 0.4810 |
| pittosporum | 0.6077 | 68.10 | 11.78 | 0.2246 | 0.9911 | 40.68 | 0.5103 |
| Pittosporumbalansae | 0.6135* | 68.21 | 11.77 | 0.2247 | 0.9914 | 40.22 | 0.5127 |
| Pittosporumcrispulum | 0.6135* | 68.21 | 11.77 | 0.2247 | 0.9914 | 40.22 | 0.5127 |
| Pittosporumperryanum | 0.6135* | 68.21 | 11.77 | 0.2247 | 0.9914 | 40.22 | 0.5127 |
| araliaceae | 0.4737 | 65.72 | 11.81 | 0.2228 | 0.9849 | 51.50 | 0.4539 |
| heteropanax | 0.3638 | 63.76 | 11.84 | 0.2213 | 0.9797 | 60.37 | 0.4077 |
| Heteropanaxfragrans | 0.3440* | 63.41 | 11.85 | 0.2210 | 0.9788 | 61.96 | 0.3994 |
| schefflera | 0.3975 | 64.36 | 11.83 | 0.2217 | 0.9813 | 57.65 | 0.4219 |
| Scheffleraһainanensis | 0.3946* | 64.31 | 11.84 | 0.2217 | 0.9812 | 57.88 | 0.4206 |
| Scheffleraheptaphylla | 0.3946* | 64.31 | 11.84 | 0.2217 | 0.9812 | 57.88 | 0.4206 |
| dendropanax | 0.4178 | 64.72 | 11.83 | 0.2220 | 0.9823 | 56.01 | 0.4304 |
| Dendropanaxhainanensis | 0.4198* | 64.76 | 11.83 | 0.2220 | 0.9824 | 55.84 | 0.4313 |
| dipsacales | 0.5508 | 67.09 | 11.79 | 0.2238 | 0.9885 | 45.28 | 0.4863 |
| adoxaceae | 0.5537 | 67.14 | 11.79 | 0.2239 | 0.9886 | 45.05 | 0.4876 |
| sambucus | 0.4821 | 65.87 | 11.81 | 0.2229 | 0.9853 | 50.82 | 0.4575 |
| Sambucusjavanica | 0.4463* | 65.23 | 11.82 | 0.2224 | 0.9836 | 53.71 | 0.4424 |
| viburnum | 0.5923 | 67.83 | 11.78 | 0.2244 | 0.9904 | 41.93 | 0.5038 |
| Viburnumpunctatum | 0.6310* | 68.52 | 11.77 | 0.2249 | 0.9922 | 38.80 | 0.5201 |
| escalloniales | 0.5514 | 67.10 | 11.79 | 0.2238 | 0.9885 | 45.22 | 0.4866 |
| escalloniaceae | 0.5515 | 67.10 | 11.79 | 0.2238 | 0.9885 | 45.22 | 0.4867 |
| polyosma | 0.5519 | 67.11 | 11.79 | 0.2238 | 0.9885 | 45.19 | 0.4868 |
| Polyosmacambodiana | 0.5520* | 67.11 | 11.79 | 0.2238 | 0.9885 | 45.18 | 0.4869 |
| aquifoliales | 0.5599 | 67.25 | 11.79 | 0.2239 | 0.9889 | 44.54 | 0.4902 |
| stemonuraceae | 0.5250 | 66.63 | 11.80 | 0.2235 | 0.9873 | 47.36 | 0.4755 |
| gomphandra | 0.4905 | 66.02 | 11.81 | 0.2230 | 0.9857 | 50.14 | 0.4610 |
| Gomphandratetrandra | 0.4560* | 65.40 | 11.82 | 0.2225 | 0.9840 | 52.93 | 0.4465 |
| cardiopteridaceae | 0.5935 | 67.85 | 11.78 | 0.2244 | 0.9905 | 41.83 | 0.5043 |
| gonocaryum | 0.6275 | 68.45 | 11.77 | 0.2249 | 0.9920 | 39.09 | 0.5186 |
| Gonocaryumlobbianum | 0.6615* | 69.06 | 11.76 | 0.2253 | 0.9936 | 36.34 | 0.5329 |
| aquifoliaceae | 0.5659 | 67.36 | 11.79 | 0.2240 | 0.9892 | 44.06 | 0.4927 |
| ilex | 0.5689 | 67.41 | 11.79 | 0.2241 | 0.9893 | 43.82 | 0.4939 |
| Ilexsp12 | 0.5627* | 67.30 | 11.79 | 0.2240 | 0.9890 | 44.32 | 0.4914 |
| Ilexsp2 | 0.5627* | 67.30 | 11.79 | 0.2240 | 0.9890 | 44.32 | 0.4914 |
| Ilexsp5 | 0.5627* | 67.30 | 11.79 | 0.2240 | 0.9890 | 44.32 | 0.4914 |
| Ilexsterrophylla | 0.6485* | 68.83 | 11.76 | 0.2252 | 0.9930 | 37.39 | 0.5274 |
| Ilextriflora | 0.5627* | 67.30 | 11.79 | 0.2240 | 0.9890 | 44.32 | 0.4914 |
| Ilexangulata | 0.5627* | 67.30 | 11.79 | 0.2240 | 0.9890 | 44.32 | 0.4914 |
| Ilexcochinchinensis | 0.6030* | 68.02 | 11.78 | 0.2245 | 0.9909 | 41.06 | 0.5083 |
| Ilexelmerrilliana | 0.5627* | 67.30 | 11.79 | 0.2240 | 0.9890 | 44.32 | 0.4914 |
| Ilexficoidea | 0.5627* | 67.30 | 11.79 | 0.2240 | 0.9890 | 44.32 | 0.4914 |
| Ilexgodajam | 0.5627* | 67.30 | 11.79 | 0.2240 | 0.9890 | 44.32 | 0.4914 |
| Ilexgoshiensis | 0.5627* | 67.30 | 11.79 | 0.2240 | 0.9890 | 44.32 | 0.4914 |
| Ilexhainanensis | 0.5627* | 67.30 | 11.79 | 0.2240 | 0.9890 | 44.32 | 0.4914 |
| Ilexkobuskiana | 0.5627* | 67.30 | 11.79 | 0.2240 | 0.9890 | 44.32 | 0.4914 |
| Ilexlancilimba | 0.5627* | 67.30 | 11.79 | 0.2240 | 0.9890 | 44.32 | 0.4914 |

| | | | | | | | |
|---|---|---|---|---|---|---|---|
| Ilexnuculicava | 0.5627* | 67.30 | 11.79 | 0.2240 | 0.9890 | 44.32 | 0.4914 |
| Ilexpubescens | 0.5627* | 67.30 | 11.79 | 0.2240 | 0.9890 | 44.32 | 0.4914 |
| Ilexrotunda | 0.5627* | 67.30 | 11.79 | 0.2240 | 0.9890 | 44.32 | 0.4914 |
| Ilexsp | 0.5627* | 67.30 | 11.79 | 0.2240 | 0.9890 | 44.32 | 0.4914 |
| Ilexsp0502 | 0.5627* | 67.30 | 11.79 | 0.2240 | 0.9890 | 44.32 | 0.4914 |
| Ilexsp1 | 0.5627* | 67.30 | 11.79 | 0.2240 | 0.9890 | 44.32 | 0.4914 |
| ericales | 0.5577 | 64.69 | 11.87 | 0.2264 | 0.9823 | 44.64 | 0.4934 |
| pentaphyllacaceae | 0.5621 | 52.23 | 11.05 | 0.2682 | 0.9147 | 44.37 | 0.5888 |
| pentaphylax | 0.5481 | 51.98 | 11.05 | 0.2680 | 0.9140 | 45.51 | 0.5829 |
| Pentaphylaxeuryoides | 0.5340* | 51.73 | 11.06 | 0.2678 | 0.9133 | 46.64 | 0.5770 |
| ternstroemia | 0.5878 | 51.22 | 10.76 | 0.2812 | 0.8849 | 42.53 | 0.6419 |
| Ternstroemiagymnanthera | 0.5814* | 51.11 | 10.76 | 0.2811 | 0.8846 | 43.05 | 0.6392 |
| Ternstroemiahainanensis | 0.5814* | 51.11 | 10.76 | 0.2811 | 0.8846 | 43.05 | 0.6392 |
| anneslea | 0.6425 | 52.20 | 10.74 | 0.2820 | 0.8875 | 38.12 | 0.6649 |
| Annesleafragrans | 0.6843* | 52.94 | 10.73 | 0.2825 | 0.8894 | 34.74 | 0.6825 |
| eurya | 0.5241 | 26.94 | 12.24 | 0.2321 | 0.8492 | 40.74 | 0.7434 |
| Euryaciliata | 0.5200* | 16.03* | 13.11* | 0.2017* | 0.8480* | 37.47* | 0.7847* |
| Euryacuneata | 0.5200* | 26.87 | 12.24 | 0.2321 | 0.8490 | 41.06 | 0.7417 |
| Euryagroffii | 0.5200* | 26.87 | 12.24 | 0.2321 | 0.8490 | 41.06 | 0.7417 |
| Euryahainanensis | 0.5200* | 26.87 | 12.24 | 0.2321 | 0.8490 | 41.06 | 0.7417 |
| Euryaloquaiana | 0.5200* | 26.87 | 12.24 | 0.2321 | 0.8490 | 41.06 | 0.7417 |
| Euryanitida | 0.5300* | 27.05 | 12.23 | 0.2322 | 0.8494 | 40.26 | 0.7459 |
| adinandra | 0.5331 | 64.20 | 8.873 | 0.3449 | 0.7874 | 59.31 | 0.6340 |
| Adinandraangustifolia | 0.5814* | 65.06 | 8.860 | 0.3455 | 0.7897 | 55.41 | 0.6543 |
| Adinandrahainanensis | 0.4700* | 69.12* | 8.440* | 0.3527* | 0.7504* | 72.69* | 0.5811* |
| cleyera | 0.5578 | 61.94 | 8.609 | 0.3709 | 0.8268 | 44.54 | 0.6966 |
| Cleyeraobscurinervia | 0.5677* | 65.45* | 7.898* | 0.4054* | 0.8313* | 39.26* | 0.7265* |
| ebenaceae | 0.6046 | 52.81 | 11.50 | 0.2423 | 1.016 | 40.01 | 0.4803 |
| diospyros | 0.5900 | 52.55 | 11.50 | 0.2421 | 1.016 | 41.19 | 0.4742 |
| Diospyrossusarticulata | 0.5643* | 52.09 | 11.51 | 0.2418 | 1.014 | 43.26 | 0.4634 |
| Diospyroschunii | 0.5643* | 52.09 | 11.51 | 0.2418 | 1.014 | 43.26 | 0.4634 |
| Diospyroseriantha | 0.7026* | 54.55 | 11.47 | 0.2436 | 1.021 | 32.10 | 0.5215 |
| Diospyroshainanensis | 0.5643* | 52.09 | 11.51 | 0.2418 | 1.014 | 43.26 | 0.4634 |
| Diospyrosinflata | 0.5643* | 52.09 | 11.51 | 0.2418 | 1.014 | 43.26 | 0.4634 |
| Diospyroslongibracteata | 0.5100* | 51.12 | 11.52 | 0.2410 | 1.012 | 47.64 | 0.4405 |
| Diospyrosmaclurei | 0.7040* | 54.58 | 11.47 | 0.2437 | 1.021 | 31.99 | 0.5221 |
| Diospyrosmorrisiana | 0.5643* | 52.09 | 11.51 | 0.2418 | 1.014 | 43.26 | 0.4634 |
| Diospyrosstrigosa | 0.5643* | 52.09 | 11.51 | 0.2418 | 1.014 | 43.26 | 0.4634 |
| primulaceae | 0.6407 | 53.45 | 11.49 | 0.2428 | 1.018 | 37.10 | 0.4955 |
| myrsine | 0.7145 | 54.76 | 11.47 | 0.2438 | 1.021 | 31.14 | 0.5265 |
| Myrsineseguinii | 0.7410* | 55.24 | 11.46 | 0.2442 | 1.023 | 29.00 | 0.5377 |
| Myrsinestolonifera | 0.7410* | 55.24 | 11.46 | 0.2442 | 1.023 | 29.00 | 0.5377 |
| ardisia | 0.6111 | 52.92 | 11.49 | 0.2424 | 1.017 | 39.48 | 0.4831 |
| Ardisiacrassinervosa | 0.6265* | 53.20 | 11.49 | 0.2426 | 1.017 | 38.24 | 0.4895 |
| Ardisiadensilepidotula | 0.6265* | 53.20 | 11.49 | 0.2426 | 1.017 | 38.24 | 0.4895 |
| Ardisiaobtusa | 0.5909* | 52.56 | 11.50 | 0.2421 | 1.016 | 41.11 | 0.4745 |
| Ardisiaquinquegona | 0.5909* | 52.56 | 11.50 | 0.2421 | 1.016 | 41.11 | 0.4745 |
| Ardisiavillosa | 0.5909* | 52.56 | 11.50 | 0.2421 | 1.016 | 41.11 | 0.4745 |
| Ardisiavirens | 0.5909* | 52.56 | 11.50 | 0.2421 | 1.016 | 41.11 | 0.4745 |
| Embelia | 0.6400 | 53.44 | 11.49 | 0.2428 | 1.018 | 37.16 | 0.4952 |
| Embeliavestita | 0.6393* | 53.42 | 11.49 | 0.2428 | 1.018 | 37.21 | 0.4949 |
| maesa | 0.6672 | 53.92 | 11.48 | 0.2432 | 1.019 | 34.96 | 0.5066 |
| Maesaacuminatissima | 0.6760* | 54.08 | 11.48 | 0.2433 | 1.020 | 34.25 | 0.5103 |
| Maesaconsanguinea | 0.6760* | 54.08 | 11.48 | 0.2433 | 1.020 | 34.25 | 0.5103 |
| Maesaperlarius | 0.6760* | 54.08 | 11.48 | 0.2433 | 1.020 | 34.25 | 0.5103 |
| sapotaceae | 0.5887 | 49.42 | 11.39 | 0.2409 | 1.053 | 40.83 | 0.4318 |
| planchonella | 0.6952 | 49.72 | 10.04 | 0.2555 | 1.060 | 37.89 | 0.5016 |
| Planchonellaclemensii | 0.7148* | 50.07 | 10.03 | 0.2558 | 1.061 | 36.31 | 0.5098 |
| chrysophyllum | 0.5699 | 47.49 | 10.07 | 0.2538 | 1.055 | 48.00 | 0.4489 |
| Chrysophyllumlanceolatum | 0.5230* | 46.66 | 10.09 | 0.2532 | 1.052 | 51.78 | 0.4292 |
| madhuca | 0.8592 | 51.58 | 9.115 | 0.2665 | 1.069 | 28.43 | 0.5872 |
| Madhucahainanensis | 0.9257* | 52.23* | 8.656* | 0.2718* | 1.073* | 24.95* | 0.6235* |
| pouteria | 0.6678 | 50.30 | 10.93 | 0.2464 | 1.058 | 36.33 | 0.4735 |

| | | | | | | | |
|---|---|---|---|---|---|---|---|
| Pouteriaannamensis | 0.6878* | 50.66 | 10.92 | 0.2467 | 1.059 | 34.71 | 0.4819 |
| sarcosperma | 0.5277 | 45.77 | 11.73 | 0.2345 | 1.088 | 43.40 | 0.3561 |
| Sarcospermalaurinum | 0.4667* | 42.11* | 12.08* | 0.2281* | 1.122* | 45.97* | 0.2804* |
| ericaceae | 0.5276 | 61.14 | 12.50 | 0.2220 | 0.9546 | 47.08 | 0.4928 |
| rhododendron | 0.4984 | 60.63 | 12.51 | 0.2216 | 0.9532 | 49.43 | 0.4805 |
| Rhododendronmoulmainense | 0.4955* | 60.57 | 12.51 | 0.2216 | 0.9531 | 49.67 | 0.4793 |
| symplocaceae | 0.5244 | 66.10 | 14.23 | 0.2023 | 0.9731 | 42.54 | 0.5003 |
| symplocos | 0.5317 | 66.23 | 14.23 | 0.2024 | 0.9734 | 41.95 | 0.5034 |
| Symplocoscongesta | 0.5335* | 66.26 | 14.23 | 0.2024 | 0.9735 | 41.80 | 0.5041 |
| Symplocoscrassilimba | 0.5335* | 66.26 | 14.23 | 0.2024 | 0.9735 | 41.80 | 0.5041 |
| Symplocoseuryoides | 0.5335* | 66.26 | 14.23 | 0.2024 | 0.9735 | 41.80 | 0.5041 |
| Symplocosglauca | 0.5335* | 66.26 | 14.23 | 0.2024 | 0.9735 | 41.80 | 0.5041 |
| Symplocosheishanensis | 0.5335* | 66.26 | 14.23 | 0.2024 | 0.9735 | 41.80 | 0.5041 |
| Symplocoslancifolia | 0.4635* | 65.02 | 14.25 | 0.2015 | 0.9702 | 47.45 | 0.4747 |
| Symplocospendula | 0.5335* | 66.26 | 14.23 | 0.2024 | 0.9735 | 41.80 | 0.5041 |
| Symplocospoilanei | 0.5335* | 66.26 | 14.23 | 0.2024 | 0.9735 | 41.80 | 0.5041 |
| Symplocospseudobarberina | 0.5335* | 66.26 | 14.23 | 0.2024 | 0.9735 | 41.80 | 0.5041 |
| Symplocosracemosa | 0.5335* | 66.26 | 14.23 | 0.2024 | 0.9735 | 41.80 | 0.5041 |
| Symplocossp1 | 0.5335* | 66.26 | 14.23 | 0.2024 | 0.9735 | 41.80 | 0.5041 |
| Symplocossumuntia | 0.5335* | 66.26 | 14.23 | 0.2024 | 0.9735 | 41.80 | 0.5041 |
| Symplocosviridissima | 0.5335* | 66.26 | 14.23 | 0.2024 | 0.9735 | 41.80 | 0.5041 |
| Symplocoswikstroemiifolia | 0.5335* | 66.26 | 14.23 | 0.2024 | 0.9735 | 41.80 | 0.5041 |
| Symplocosadenophylla | 0.6500* | 68.34 | 14.20 | 0.2040 | 0.9789 | 32.41 | 0.5531 |
| Symplocosanomala | 0.4815* | 65.34 | 14.24 | 0.2017 | 0.9711 | 46.00 | 0.4822 |
| Symplocoscochinchinensis | 0.5150* | 65.93 | 14.23 | 0.2022 | 0.9726 | 43.30 | 0.4963 |
| Symplocaceaesp7 | 0.5335* | 66.26 | 14.23 | 0.2024 | 0.9735 | 41.80 | 0.5041 |
| styracaceae | 0.4394 | 70.54 | 16.77 | 0.1798 | 1.033 | 39.50 | 0.4750 |
| alniphyllum | 0.3940 | 81.62 | 21.81 | 0.1365 | 1.159 | 23.36 | 0.4768 |
| Alniphyllumfortunei | 0.3827* | 84.39* | 23.07* | 0.1257* | 1.191* | 19.33* | 0.4773* |
| styrax | 0.4165 | 70.13 | 16.77 | 0.1795 | 1.032 | 41.35 | 0.4654 |
| Styraxagrestis | 0.4050* | 69.92 | 16.78 | 0.1793 | 1.032 | 42.27 | 0.4605 |
| Styraxsuberifolius | 0.4050* | 69.92 | 16.78 | 0.1793 | 1.032 | 42.27 | 0.4605 |
| theaceae | 0.5304 | 62.30 | 12.19 | 0.2148 | 0.8959 | 52.11 | 0.4960 |
| camellia | 0.5499 | 58.46 | 10.09 | 0.2425 | 0.9350 | 47.58 | 0.5064 |
| Camelliacaudata | 0.5500* | 58.46 | 10.09 | 0.2425 | 0.9350 | 47.57 | 0.5064 |
| Camelliaoleifera | 0.5500* | 58.46 | 10.09 | 0.2425 | 0.9350 | 47.57 | 0.5064 |
| Camelliapaucipunctata | 0.5500* | 58.46 | 10.09 | 0.2425 | 0.9350 | 47.57 | 0.5064 |
| Camelliasinensis | 0.5500* | 58.46 | 10.09 | 0.2425 | 0.9350 | 47.57 | 0.5064 |
| Camelliasp1 | 0.5500* | 58.46 | 10.09 | 0.2425 | 0.9350 | 47.57 | 0.5064 |
| Camelliasp3 | 0.5500* | 58.46 | 10.09 | 0.2425 | 0.9350 | 47.57 | 0.5064 |
| Camelliaxanthochroma | 0.5500* | 58.46 | 10.09 | 0.2425 | 0.9350 | 47.57 | 0.5064 |
| polyspora | 0.5613 | 58.81 | 9.267 | 0.2557 | 1.016 | 37.45 | 0.5075 |
| Polysporaaxillaris | 0.5677* | 58.93 | 9.265 | 0.2558 | 1.016 | 36.94 | 0.5102 |
| Polysporahainanensis | 0.5677* | 59.08* | 8.447* | 0.2689* | 1.097* | 27.73* | 0.5065* |
| pyrenaria | 0.5225 | 54.25 | 11.24 | 0.2286 | 0.7757 | 69.35 | 0.5098 |
| Pyrenariajonquieriana | 0.5182* | 50.60* | 11.56* | 0.2281* | 0.6981* | 80.05* | 0.5192* |
| Pyrenariamicrocarpa | 0.5182* | 54.17 | 11.24 | 0.2285 | 0.7755 | 69.70 | 0.5080 |
| Pyrenariaspectabilis | 0.5182* | 54.17 | 11.24 | 0.2285 | 0.7755 | 69.70 | 0.5080 |
| schima | 0.5430 | 66.56 | 10.83 | 0.1947 | 0.7539 | 64.12 | 0.4815 |
| Schimaremotiserrata | 0.5526* | 66.73 | 10.83 | 0.1948 | 0.7544 | 63.35 | 0.4855 |
| Schimasuperba | 0.5375* | 68.94* | 10.54* | 0.1837* | 0.7052* | 68.53* | 0.4701* |
| cornales | 0.5439 | 69.49 | 11.72 | 0.2212 | 0.9946 | 45.91 | 0.4793 |
| nyssaceae | 0.5066 | 68.83 | 11.73 | 0.2207 | 0.9929 | 48.92 | 0.4636 |
| mastixia | 0.4989 | 68.69 | 11.73 | 0.2206 | 0.9925 | 49.54 | 0.4603 |
| Mastixiapentandra | 0.4950* | 68.62 | 11.73 | 0.2206 | 0.9923 | 49.85 | 0.4587 |
| cornaceae | 0.4976 | 68.67 | 11.73 | 0.2206 | 0.9925 | 49.65 | 0.4598 |
| alangium | 0.4397 | 67.64 | 11.75 | 0.2198 | 0.9898 | 54.31 | 0.4355 |
| Alangiumchinense | 0.4017* | 66.96 | 11.76 | 0.2193 | 0.9880 | 57.39 | 0.4194 |
| Alangiumkurzii | 0.4200* | 67.29 | 11.75 | 0.2195 | 0.9888 | 55.91 | 0.4272 |
| cornus | 0.5425 | 69.47 | 11.72 | 0.2212 | 0.9946 | 46.02 | 0.4787 |
| Cornussp1 | 0.5875* | 70.27 | 11.71 | 0.2218 | 0.9967 | 42.39 | 0.4976 |
| santalales | 0.6368 | 78.72 | 11.46 | 0.2150 | 1.018 | 38.64 | 0.5058 |
| olacaceae | 0.7150 | 80.11 | 11.44 | 0.2161 | 1.022 | 32.33 | 0.5387 |
| olax | 0.7523 | 80.77 | 11.43 | 0.2166 | 1.024 | 29.32 | 0.5544 |

| | | | | | | | |
|---|---|---|---|---|---|---|---|
| Olaximbricata | 0.7710* | 81.11 | 11.43 | 0.2169 | 1.025 | 27.81 | 0.5623 |
| santalaceae | 0.6849 | 79.57 | 11.45 | 0.2157 | 1.021 | 34.76 | 0.5260 |
| scleropyrum | 0.6975 | 79.80 | 11.45 | 0.2159 | 1.021 | 33.74 | 0.5313 |
| Scleropyrumwallichianum | 0.6993* | 79.83 | 11.45 | 0.2159 | 1.021 | 33.60 | 0.5321 |
| buxales | 0.6429 | 79.59 | 11.26 | 0.2162 | 1.073 | 34.90 | 0.4962 |
| buxaceae | 0.6693 | 80.06 | 11.25 | 0.2165 | 1.075 | 32.77 | 0.5073 |
| buxus | 0.6956 | 80.53 | 11.24 | 0.2169 | 1.076 | 30.65 | 0.5184 |
| Buxusmyrica | 0.7220* | 81.00 | 11.24 | 0.2173 | 1.077 | 28.52 | 0.5295 |
| proteales | 0.5483 | 76.15 | 11.16 | 0.2184 | 1.117 | 39.21 | 0.4485 |
| proteaceae | 0.5517 | 76.21 | 11.16 | 0.2185 | 1.117 | 38.94 | 0.4500 |
| helicia | 0.5887 | 76.87 | 11.15 | 0.2190 | 1.119 | 35.95 | 0.4655 |
| Heliciacochinchinensis | 0.5510* | 76.19 | 11.16 | 0.2185 | 1.117 | 38.99 | 0.4496 |
| Heliciaformosana | 0.6051* | 77.16 | 11.14 | 0.2192 | 1.120 | 34.63 | 0.4724 |
| Heliciahainanensis | 0.6051* | 77.16 | 11.14 | 0.2192 | 1.120 | 34.63 | 0.4724 |
| Helicialongipetiolata | 0.6051* | 77.16 | 11.14 | 0.2192 | 1.120 | 34.63 | 0.4724 |
| Heliciaobovatifolia | 0.5777* | 76.67 | 11.15 | 0.2188 | 1.119 | 36.84 | 0.4609 |
| Heliciareticulata | 0.6051* | 77.16 | 11.14 | 0.2192 | 1.120 | 34.63 | 0.4724 |
| heliciopsis | 0.4612 | 74.60 | 11.18 | 0.2173 | 1.113 | 46.24 | 0.4119 |
| Heliciopsislobata | 0.4300* | 74.04 | 11.19 | 0.2168 | 1.112 | 48.76 | 0.3988 |
| Heliciopsisterminalis | 0.4767* | 74.87 | 11.18 | 0.2175 | 1.114 | 44.99 | 0.4184 |
| sabiaceae | 0.5297 | 75.81 | 11.16 | 0.2182 | 1.116 | 40.71 | 0.4407 |
| meliosma | 0.5110 | 75.48 | 11.17 | 0.2179 | 1.115 | 42.22 | 0.4328 |
| Meliosmaangustifolia | 0.5483* | 76.15 | 11.16 | 0.2184 | 1.117 | 39.21 | 0.4485 |
| Meliosmadumicola | 0.4789* | 74.91 | 11.18 | 0.2175 | 1.114 | 44.81 | 0.4193 |
| Meliosmafordii | 0.4789* | 74.91 | 11.18 | 0.2175 | 1.114 | 44.81 | 0.4193 |
| Meliosmalaui | 0.4789* | 74.91 | 11.18 | 0.2175 | 1.114 | 44.81 | 0.4193 |
| Meliosmarigida | 0.5960* | 77.00 | 11.15 | 0.2191 | 1.119 | 35.36 | 0.4686 |
| Meliosmasquamulata | 0.4985* | 75.26 | 11.17 | 0.2178 | 1.115 | 43.23 | 0.4276 |
| Meliosmathorelii | 0.4789* | 74.91 | 11.18 | 0.2175 | 1.114 | 44.81 | 0.4193 |
| monocots | 0.5388 | 73.34 | 10.97 | 0.2237 | 1.189 | 35.00 | 0.4326 |
| arecales | 0.5312 | 73.20 | 10.98 | 0.2236 | 1.189 | 35.61 | 0.4294 |
| arecaceae | 0.5368 | 73.30 | 10.97 | 0.2236 | 1.189 | 35.16 | 0.4318 |
| pinanga | 0.5518 | 73.57 | 10.97 | 0.2238 | 1.190 | 33.94 | 0.4381 |
| Pinangabaviensis | 0.5518* | 73.57 | 10.97 | 0.2238 | 1.190 | 33.95 | 0.4380 |
| livistona | 0.5737 | 73.96 | 10.96 | 0.2241 | 1.191 | 32.18 | 0.4473 |
| Livistonasaribus | 0.5814* | 74.09 | 10.96 | 0.2242 | 1.191 | 31.56 | 0.4505 |
| licuala | 0.5533 | 73.60 | 10.97 | 0.2239 | 1.190 | 33.82 | 0.4387 |
| Licualafordiana | 0.5518* | 73.57 | 10.97 | 0.2238 | 1.190 | 33.95 | 0.4380 |
| Licualahainanensis | 0.5518* | 73.57 | 10.97 | 0.2238 | 1.190 | 33.95 | 0.4380 |
| caryota | 0.5512 | 73.56 | 10.97 | 0.2238 | 1.190 | 34.00 | 0.4378 |
| Caryotamaxima | 0.5518* | 73.57 | 10.97 | 0.2238 | 1.190 | 33.95 | 0.4380 |
| arenga | 0.5512 | 73.56 | 10.97 | 0.2238 | 1.190 | 34.00 | 0.4378 |
| Arengapinnata | 0.5518* | 73.57 | 10.97 | 0.2238 | 1.190 | 33.95 | 0.4380 |
| asparagales | 0.5104 | 72.83 | 10.98 | 0.2233 | 1.188 | 37.29 | 0.4207 |
| asparagaceae | 0.4349 | 71.49 | 11.00 | 0.2222 | 1.184 | 43.38 | 0.3889 |
| dracaena | 0.3594 | 70.14 | 11.02 | 0.2212 | 1.181 | 49.47 | 0.3571 |
| Dracaenaangustifolia | 0.3500* | 69.98 | 11.03 | 0.2211 | 1.180 | 50.23 | 0.3532 |
| magnoliids | 0.5339 | 71.49 | 10.85 | 0.2272 | 1.237 | 32.07 | 0.4226 |
| magnoniales | 0.5151 | 69.38 | 10.75 | 0.2310 | 1.329 | 28.42 | 0.3961 |
| annonaceae | 0.5547 | 71.38 | 11.56 | 0.2068 | 1.317 | 26.16 | 0.4302 |
| miliusa | 0.6161 | 72.47 | 11.54 | 0.2076 | 1.320 | 21.21 | 0.4560 |
| Miliusahorsfieldii | 0.6100* | 72.37 | 11.54 | 0.2075 | 1.320 | 21.70 | 0.4534 |
| popowia | 0.5643 | 71.55 | 11.55 | 0.2069 | 1.318 | 25.39 | 0.4342 |
| Popowiapisocarpa | 0.5450* | 71.21 | 11.56 | 0.2066 | 1.317 | 26.95 | 0.4261 |
| alphonsea | 0.7078 | 74.11 | 11.51 | 0.2088 | 1.325 | 13.81 | 0.4946 |
| Alphonseahainanensis | 0.7380* | 74.64 | 11.51 | 0.2093 | 1.326 | 11.38 | 0.5073 |
| mitrephora | 0.6614 | 73.28 | 11.53 | 0.2082 | 1.322 | 17.56 | 0.4750 |
| Mitrephoratomentosa | 0.6800* | 73.61 | 11.52 | 0.2085 | 1.323 | 16.06 | 0.4829 |
| orophea | 0.5835 | 71.89 | 11.55 | 0.2071 | 1.319 | 23.85 | 0.4423 |
| Oropheahainanensis | 0.5656* | 71.58 | 11.55 | 0.2069 | 1.318 | 25.29 | 0.4347 |
| polyalthia | 0.6103 | 72.37 | 11.54 | 0.2075 | 1.320 | 21.68 | 0.4535 |
| Polyalthiacerasoides | 0.7550* | 74.95 | 11.50 | 0.2095 | 1.327 | 10.00 | 0.5144 |
| Polyalthialaui | 0.5541* | 71.37 | 11.56 | 0.2067 | 1.317 | 26.22 | 0.4299 |
| Polyalthiaobliqua | 0.5541* | 71.37 | 11.56 | 0.2067 | 1.317 | 26.22 | 0.4299 |

| | | | | | | | |
|---|---|---|---|---|---|---|---|
| Polyalthiarumphii | 0.5800* | 71.83 | 11.55 | 0.2071 | 1.319 | 24.13 | 0.4408 |
| disepalum | 0.5637 | 71.98 | 11.82 | 0.1984 | 1.299 | 26.37 | 0.4433 |
| Disepalumplagioneurum | 0.5656* | 72.01 | 11.82 | 0.1984 | 1.299 | 26.22 | 0.4441 |
| uvaria | 0.5649 | 73.32 | 12.63 | 0.1731 | 1.241 | 29.06 | 0.4717 |
| Uvariaboniana | 0.5656* | 73.33 | 12.63 | 0.1731 | 1.241 | 29.01 | 0.4720 |
| dasymaschalon | 0.5648 | 74.64 | 13.43 | 0.1477 | 1.183 | 31.86 | 0.4996 |
| Dasymaschalonrostratum | 0.5656* | 75.09* | 13.70* | 0.1392* | 1.164* | 32.72* | 0.5093* |
| Chieniodendron | 0.5602 | 71.48 | 11.55 | 0.2068 | 1.318 | 25.73 | 0.4325 |
| Chieniodendronhainanense | 0.5656* | 71.58 | 11.55 | 0.2069 | 1.318 | 25.29 | 0.4347 |
| magnoliaceae | 0.4798 | 68.27 | 10.52 | 0.2401 | 1.435 | 26.65 | 0.3508 |
| Manglietia | 0.4517 | 73.27 | 11.92 | 0.2557 | 1.526 | 26.53 | 0.3156 |
| Manglietiafordiana | 0.4235* | 78.27* | 13.31* | 0.2712* | 1.617* | 26.41* | 0.2805* |
| Michelia | 0.5013 | 70.39 | 9.119 | 0.2470 | 1.391 | 28.22 | 0.3345 |
| Micheliabalansae | 0.5423* | 94.04* | 6.963* | 0.2560* | 1.358* | 28.91* | 0.3295* |
| Micheliagioi | 0.4880* | 70.15 | 9.122 | 0.2469 | 1.390 | 29.29 | 0.3289 |
| Micheliamediocris | 0.4950* | 49.08* | 9.866* | 0.2452* | 1.380* | 28.02* | 0.3287* |
| Lirianthe | 0.4607 | 60.24 | 10.29 | 0.2263 | 1.451 | 24.52 | 0.3716 |
| Lirianthechampionii | 0.4415* | 52.20* | 10.06* | 0.2126* | 1.468* | 22.38* | 0.3923* |
| laurales | 0.5292 | 68.79 | 10.64 | 0.2319 | 1.265 | 29.30 | 0.4193 |
| lauraceae | 0.5482 | 65.68 | 10.31 | 0.2370 | 1.185 | 28.50 | 0.4537 |
| phoebe | 0.5206 | 43.52 | 10.10 | 0.2475 | 1.208 | 35.00 | 0.3674 |
| Phoebehungmoensis | 0.4900* | 42.98 | 10.11 | 0.2471 | 1.206 | 37.46 | 0.3545 |
| Phoebetavoyana | 0.5382* | 43.84 | 10.10 | 0.2477 | 1.209 | 33.57 | 0.3748 |
| machilus | 0.5626 | 41.50 | 9.812 | 0.2637 | 1.228 | 31.41 | 0.3809 |
| Machiluschinensis | 0.5643* | 34.26* | 11.85* | 0.2313* | 1.063* | 45.63* | 0.3910* |
| Machiluscicatricosa | 0.5643* | 44.16* | 10.01* | 0.2314* | 1.165* | 28.95* | 0.3989* |
| Machilusfoonchewii | 0.5643* | 41.53 | 9.811 | 0.2637 | 1.228 | 31.27 | 0.3816 |
| Machilusgamblei | 0.5643* | 41.53 | 9.811 | 0.2637 | 1.228 | 31.27 | 0.3816 |
| Machilusmonticola | 0.5643* | 43.39* | 7.304* | 0.3440* | 1.472* | 19.02* | 0.3508* |
| Machiluspomifera | 0.5643* | 41.53 | 9.811 | 0.2637 | 1.228 | 31.27 | 0.3816 |
| Machilusrobusta | 0.5643* | 41.53 | 9.811 | 0.2637 | 1.228 | 31.27 | 0.3816 |
| Machilussp | 0.5643* | 41.53 | 9.811 | 0.2637 | 1.228 | 31.27 | 0.3816 |
| Machilusvelutina | 0.5643* | 41.53 | 9.811 | 0.2637 | 1.228 | 31.27 | 0.3816 |
| neolitsea | 0.5343 | 41.28 | 10.13 | 0.1842 | 1.022 | 36.13 | 0.4074 |
| Neolitseacambodiana | 0.5415* | 47.30* | 9.352* | 0.1939* | 0.8764* | 48.42* | 0.4372* |
| Neolitseachui | 0.5415* | 41.40 | 10.13 | 0.1843 | 1.022 | 35.55 | 0.4104 |
| Neolitseaellipsoidea | 0.4527* | 59.81* | 7.362* | 0.1997* | 1.137* | 25.32* | 0.4107* |
| Neolitseaoblongifolia | 0.5300* | 26.00* | 13.35* | 0.1541* | 1.002* | 37.71* | 0.4355* |
| Neolitseaovatifolia | 0.5415* | 22.22* | 8.611* | 0.2014* | 0.9336* | 42.62* | 0.3703* |
| Neolitseaphanerophlebia | 0.5900* | 42.27 | 10.12 | 0.1850 | 1.024 | 31.63 | 0.4308 |
| Neolitseapulchella | 0.5415* | 47.90* | 11.87* | 0.1631* | 1.119* | 32.21* | 0.3625* |
| alseodaphne | 0.5678 | 58.17 | 10.79 | 0.2059 | 1.350 | 25.23 | 0.3721 |
| Alseodaphnehainanensis | 0.5970* | 62.18* | 10.53* | 0.2029* | 1.508* | 16.78* | 0.3631* |
| lindera | 0.4822 | 51.29 | 13.77 | 0.1880 | 0.9414 | 50.83 | 0.3727 |
| Linderacommunis | 0.4520* | 50.75 | 13.77 | 0.1876 | 0.9400 | 53.27 | 0.3599 |
| Linderakwangtungensis | 0.5778* | 48.84* | 18.56* | 0.1422* | 0.7819* | 60.27* | 0.4201* |
| Linderametcalfiana | 0.4520* | 50.75 | 13.77 | 0.1876 | 0.9400 | 53.27 | 0.3599 |
| Linderanacusua | 0.4520* | 50.75 | 13.77 | 0.1876 | 0.9400 | 53.27 | 0.3599 |
| Linderarobusta | 0.4520* | 51.71* | 10.98* | 0.2188* | 0.9915* | 42.87* | 0.3491* |
| cinnamomum | 0.4926 | 62.43 | 9.055 | 0.2220 | 0.9744 | 43.65 | 0.4174 |
| Cinnamomumbejolghota | 0.4671* | 61.97 | 9.062 | 0.2216 | 0.9732 | 45.70 | 0.4066 |
| Cinnamomumburmannii | 0.4900* | 47.50* | 8.635* | 0.1894* | 0.8786* | 39.91* | 0.5723* |
| Cinnamomumliangii | 0.4671* | 61.97 | 9.062 | 0.2216 | 0.9732 | 45.70 | 0.4066 |
| Cinnamomumparthenoxylon | 0.5800* | 81.23* | 9.790* | 0.2307* | 1.007* | 51.63* | 0.3172* |
| Cinnamomumrigidissimum | 0.4671* | 62.91* | 7.397* | 0.2529* | 0.9844* | 35.97* | 0.4017* |
| Cinnamomumsubavenium | 0.5000* | 62.56 | 9.053 | 0.2221 | 0.9748 | 43.05 | 0.4205 |
| Cinnamomumtsoi | 0.4671* | 61.97 | 9.062 | 0.2216 | 0.9732 | 45.70 | 0.4066 |
| cryptocarya | 0.5675 | 57.62 | 9.652 | 0.2359 | 1.151 | 32.02 | 0.4432 |
| Cryptocaryachinensis | 0.5000* | 51.64* | 9.121* | 0.2335* | 1.489* | 19.64* | 0.3365* |
| Cryptocaryachingii | 0.5400* | 54.36* | 9.633* | 0.2346* | 0.7937* | 56.95* | 0.4846* |
| Cryptocaryadensiflora | 0.5357* | 57.05 | 9.660 | 0.2355 | 1.149 | 34.58 | 0.4298 |
| Cryptocaryaimpressinervia | 0.5613* | 57.51 | 9.653 | 0.2359 | 1.151 | 32.52 | 0.4406 |
| Cryptocaryamaclurei | 0.5613* | 57.51 | 9.653 | 0.2359 | 1.151 | 32.52 | 0.4406 |
| Cryptocaryametcalfiana | 0.7600* | 61.04 | 9.598 | 0.2386 | 1.160 | 16.49 | 0.5241 |

| | | | | | | | |
|---|---|---|---|---|---|---|---|
| Cryptocaryasp9 | 0.5613* | 57.51 | 9.653 | 0.2359 | 1.151 | 32.52 | 0.4406 |
| litsea | 0.4348 | 66.76 | 11.24 | 0.2376 | 1.107 | 33.21 | 0.4681 |
| Litseabaviensis | 0.4353* | 61.68* | 13.89* | 0.2453* | 1.202* | 36.18* | 0.2757* |
| Litseacubeba | 0.3100* | 64.53 | 11.27 | 0.2359 | 1.102 | 43.28 | 0.4156 |
| Litseaelongata | 0.4255* | 66.59 | 11.24 | 0.2374 | 1.107 | 33.96 | 0.4642 |
| Litsealancilimba | 0.5317* | 68.48 | 11.21 | 0.2389 | 1.112 | 25.39 | 0.5089 |
| Litseamonopetala | 0.4228* | 66.54 | 11.24 | 0.2374 | 1.107 | 34.17 | 0.4631 |
| Litseapseudoelongata | 0.4255* | 66.59 | 11.24 | 0.2374 | 1.107 | 33.96 | 0.4642 |
| Litseavariabilischinensis | 0.4255* | 66.59 | 11.24 | 0.2374 | 1.107 | 33.96 | 0.4642 |
| Litseavariabilis | 0.4255* | 51.57* | 10.15* | 0.2447* | 1.223* | 21.33* | 0.6039* |
| Litseaverticillata | 0.4255* | 90.65* | 10.65* | 0.2235* | 0.8450* | 38.96* | 0.5726* |
| beilschmiedia | 0.5632 | 80.70 | 8.177 | 0.2712 | 1.482 | 18.01 | 0.4056 |
| Beilschmiediaappendiculata | 0.5631* | 80.70 | 8.178 | 0.2712 | 1.482 | 18.02 | 0.4056 |
| Beilschmiediaglauca | 0.5631* | 80.70 | 8.178 | 0.2712 | 1.482 | 18.02 | 0.4056 |
| Beilschmiediaintermedia | 0.5148* | 79.84 | 8.191 | 0.2706 | 1.480 | 21.92 | 0.3853 |
| Beilschmiedialaevis | 0.5631* | 84.61* | 7.665* | 0.2794* | 1.561* | 15.65* | 0.3904* |
| Beilschmiediaobconica | 0.5631* | 80.70 | 8.178 | 0.2712 | 1.482 | 18.02 | 0.4056 |
| Beilschmiediapercoriacea | 0.5631* | 80.70 | 8.178 | 0.2712 | 1.482 | 18.02 | 0.4056 |
| Beilschmiediapergamentacea | 0.5631* | 80.70 | 8.178 | 0.2712 | 1.482 | 18.02 | 0.4056 |
| Beilschmiediaroxburghiana | 0.5631* | 80.70 | 8.178 | 0.2712 | 1.482 | 18.02 | 0.4056 |
| Beilschmiediasp | 0.6063* | 81.47 | 8.166 | 0.2718 | 1.484 | 14.54 | 0.4238 |
| Beilschmiediasp0502 | 0.5631* | 80.70 | 8.178 | 0.2712 | 1.482 | 18.02 | 0.4056 |
| Beilschmiediasp4 | 0.5631* | 80.70 | 8.178 | 0.2712 | 1.482 | 18.02 | 0.4056 |
| Beilschmiediasp6 | 0.5631* | 80.70 | 8.178 | 0.2712 | 1.482 | 18.02 | 0.4056 |
| Beilschmiediatsangii | 0.5631* | 80.70 | 8.178 | 0.2712 | 1.482 | 18.02 | 0.4056 |
| Beilschmiediatungfangensis | 0.5631* | 80.70 | 8.178 | 0.2712 | 1.482 | 18.02 | 0.4056 |
| Beilschmiediawangii | 0.5631* | 80.70 | 8.178 | 0.2712 | 1.482 | 18.02 | 0.4056 |
| endiandra | 0.5923 | 77.32 | 8.682 | 0.2634 | 1.404 | 18.03 | 0.4331 |
| Endiandrahainanensis | 0.6150* | 77.72 | 8.676 | 0.2637 | 1.405 | 16.20 | 0.4427 |
| Lauraceaesp5 | 0.5643* | 65.97 | 10.31 | 0.2372 | 1.186 | 27.20 | 0.4605 |
| Lauraceaesp8 | 0.5643* | 65.97 | 10.31 | 0.2372 | 1.186 | 27.20 | 0.4605 |
| austrobaileyales | 0.5515 | 72.68 | 10.91 | 0.2256 | 1.214 | 32.31 | 0.4340 |
| schisandraceae | 0.5698 | 73.01 | 10.90 | 0.2259 | 1.215 | 30.83 | 0.4417 |
| illicium | 0.5759 | 73.12 | 10.90 | 0.2259 | 1.215 | 30.34 | 0.4442 |
| Illiciumsp1 | 0.5790* | 73.17 | 10.90 | 0.2260 | 1.215 | 30.09 | 0.4455 |
| Illiciumternstroemioides | 0.5790* | 73.17 | 10.90 | 0.2260 | 1.215 | 30.09 | 0.4455 |
| gymnosperms | 0.5044 | 71.84 | 10.92 | 0.2250 | 1.212 | 36.11 | 0.4141 |
| pinales | 0.4858 | 71.51 | 10.92 | 0.2247 | 1.211 | 37.61 | 0.4063 |
| pinaceae | 0.5028 | 71.82 | 10.92 | 0.2249 | 1.212 | 36.24 | 0.4135 |
| pinus | 0.5537 | 72.72 | 10.91 | 0.2256 | 1.214 | 32.13 | 0.4349 |
| Pinuscaribaea | 0.5707* | 73.02 | 10.90 | 0.2259 | 1.215 | 30.76 | 0.4420 |
| podocarpaceae | 0.4644 | 71.13 | 10.93 | 0.2244 | 1.210 | 39.33 | 0.3973 |
| podocarpus | 0.4769 | 71.35 | 10.93 | 0.2246 | 1.210 | 38.33 | 0.4026 |
| Podocarpusneriifolius | 0.4769* | 71.35 | 10.93 | 0.2246 | 1.210 | 38.33 | 0.4026 |
| dacrycarpus | 0.4463 | 70.81 | 10.94 | 0.2242 | 1.209 | 40.80 | 0.3897 |
| Dacrycarpusimbricatus | 0.4133* | 70.22 | 10.94 | 0.2237 | 1.207 | 43.46 | 0.3758 |
| dacrydium | 0.5502 | 72.66 | 10.91 | 0.2256 | 1.214 | 32.42 | 0.4334 |
| Dacrydiumpectinatum | 0.5856* | 73.29 | 10.90 | 0.2261 | 1.215 | 29.56 | 0.4483 |
| cupressaceae | 0.3732 | 69.51 | 10.96 | 0.2232 | 1.206 | 46.70 | 0.3590 |
| cunninghamia | 0.3445 | 69.00 | 10.96 | 0.2228 | 1.204 | 49.02 | 0.3469 |
| Cunninghamialanceolata | 0.3157* | 68.49 | 10.97 | 0.2224 | 1.203 | 51.34 | 0.3348 |
| monilophyte | 0.5347 | 72.38 | 10.91 | 0.2254 | 1.213 | 33.67 | 0.4269 |
| cyatheales | 0.5619 | 72.87 | 10.90 | 0.2258 | 1.214 | 31.47 | 0.4383 |
| cyatheaceae | 0.5736 | 73.08 | 10.90 | 0.2259 | 1.215 | 30.53 | 0.4432 |
| Alsophila | 0.5775 | 73.15 | 10.90 | 0.2260 | 1.215 | 30.21 | 0.4449 |
| Alsophilapodophylla | 0.5814* | 73.22 | 10.90 | 0.2260 | 1.215 | 29.90 | 0.4465 |

#### Cross-validation details

In addition to estimating missing values, PhyloPars performs cross-validation: it temporarily excludes each observation from the available information, in order to calculate its best estimate given al observations and the optimal phylogenetic covariances. This provides a detailed estimate of the error and bias one may expect for the estimated missing feature values. The distributions of estima calculated through cross-validation are shown below; these are compared with two null models (red and green curves) in order to assess the improvement resulting from the PhyloPars evolutionar

##### Meanwd

**Cross-validation error**

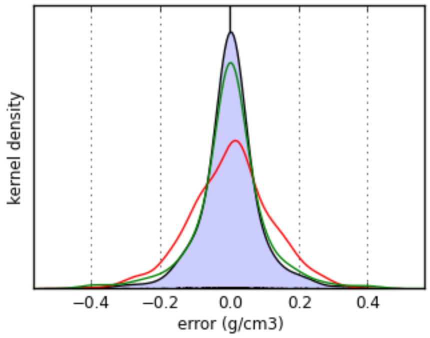


| | evolutionary model | mean model | nearest neighbor model |
|---|---|---|---|
| mean bias | **-0.000541 g/cm3** | 3.3e-16 g/cm3 | -0.000865 g/cm3 |
| mean error | **0.05 g/cm3** | 0.0884 g/cm3 | 0.0591 g/cm3 |

**Meanleafarea**

**Cross-validation error**

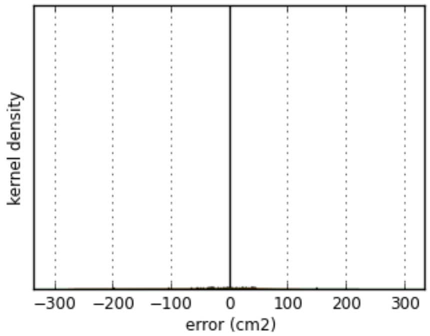


| | evolutionary model | mean model | nearest neighbor model |
|---|---|---|---|
| mean bias | **-4.28 cm2** | -1.73e-14 cm2 | -7.4 cm2 |
| mean error | **32.9 cm2** | 33.7 cm2 | 35.5 cm2 |

**Meansla**

**Cross-validation error**

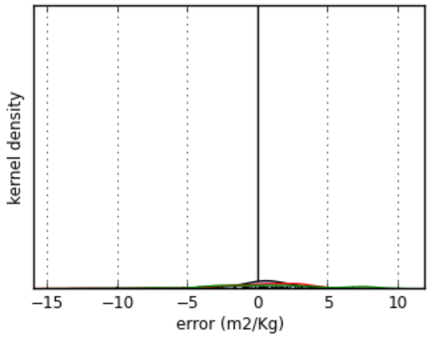


| | evolutionary model | mean model | nearest neighbor model |
|---|---|---|---|
| mean bias | **0.0902 m2/Kg** | -1.43e-15 m2/Kg | 0.484 m2/Kg |
| mean error | **1.79 m2/Kg** | 2.56 m2/Kg | 3.51 m2/Kg |

**Meanleafthickness**

**Cross-validation error**

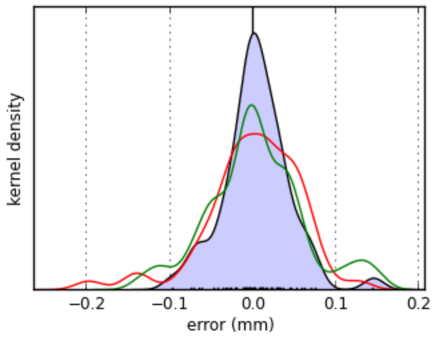


| | evolutionary model | mean model | nearest neighbor model |
|---|---|---|---|
| mean bias | **0.0041 mm** | -6.26e-17 mm | 0.00353 mm |
| mean error | **0.0291 mm** | 0.0423 mm | 0.0432 mm |

#### Meanrootavgdiam

**Cross-validation error**

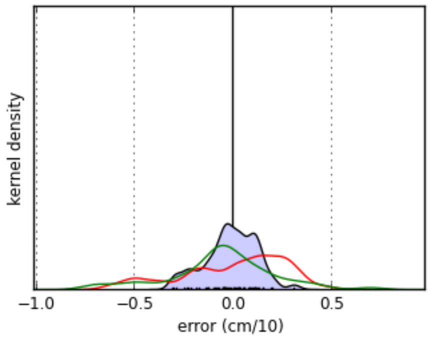


| | evolutionary model | mean model | nearest neighbor model |
|---|---|---|---|
| mean bias | **-0.0123 cm/10** | 6.4e-17 cm/10 | -0.06 cm/10 |
| mean error | **0.105 cm/10** | 0.22 cm/10 | 0.2 cm/10 |

#### Meanspecificrootlength

**Cross-validation error**

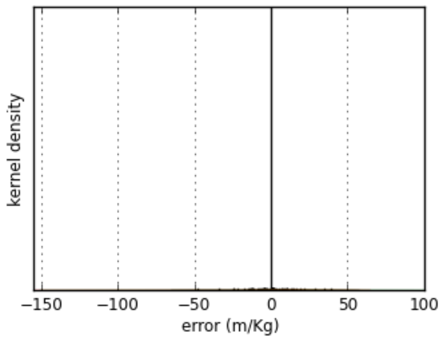


| | evolutionary model | mean model | nearest neighbor model |
|---|---|---|---|
| mean bias | **-0.699 m/Kg** | -2.89e-15 m/Kg | 5.26 m/Kg |
| mean error | **10.9 m/Kg** | 17.6 m/Kg | 23.4 m/Kg |

#### Meanroottd

**Cross-validation error**

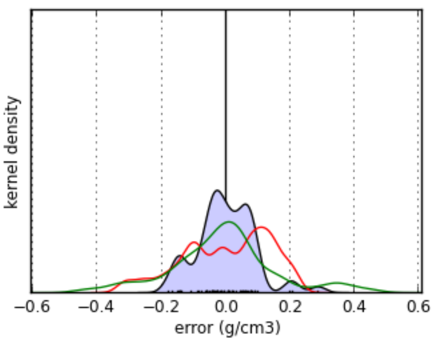


| | evolutionary model | mean model | nearest neighbor model |
|---|---|---|---|
| mean bias | **-0.000553 g/cm3** | 1.01e-16 g/cm3 | -0.0101 g/cm3 |
| mean error | **0.0692 g/cm3** | 0.116 g/cm3 | 0.118 g/cm3 |

#### Meanbranchiness

**Cross-validation error**

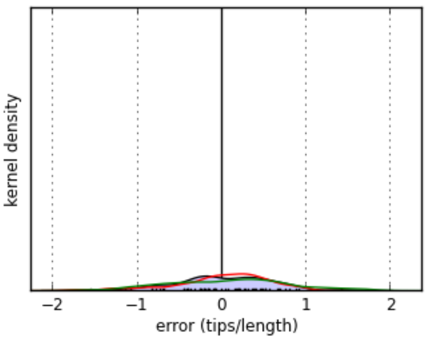


| | evolutionary model | mean model | nearest neighbor model |
|---|---|---|---|
| mean bias | **0.00984 tips/length** | -1.69e-16 tips/length | 0.0764 tips/length |
| mean error | **0.455 tips/length** | 0.432 tips/length | 0.593 tips/length |

The PhyloPars tool is (c) Jorn Bruggeman 2019. If you are experiencing problems with this tool, please contact jbr [at] pml.ac.uk or bwbrandt [at] few.vu.nl.
